# Single cell resolution landscape of equine peripheral blood mononuclear cells reveals diverse immune cell subtypes including T-bet^+^ B cells

**DOI:** 10.1101/2020.05.05.077362

**Authors:** Roosheel S. Patel, Joy E. Tomlinson, Thomas J. Divers, Gerlinde R. Van de Walle, Brad R. Rosenberg

## Abstract

Traditional laboratory model organisms represent a small fraction of the diversity of multicellular life, and findings in any given experimental model often do not translate to other species. Immunology research in non-traditional model organisms can be advantageous or even necessary (e.g. for host-pathogen interaction studies), but presents multiple challenges, many stemming from an incomplete understanding of potentially species-specific immune cell types, frequencies and phenotypes. Identifying and characterizing immune cells in such organisms is frequently limited by the availability of species-reactive immunophenotyping reagents for flow cytometry, and insufficient prior knowledge of cell type-defining markers. Here, we demonstrate the utility of single cell RNA sequencing (scRNA-Seq) to characterize immune cells for which traditional experimental tools are limited. Specifically, we used scRNA-Seq to comprehensively define the cellular diversity of equine peripheral blood mononuclear cells (PBMCs) from healthy horses across different breeds, ages, and sexes. We identified 30 cell type clusters partitioned into five major populations: Monocytes/Dendritic Cells, B cells, CD3^+^PRF1^+^ lymphocytes, CD3^+^PRF1^-^ lymphocytes, and Basophils. Comparative analyses revealed many cell populations analogous to human PBMC, including transcriptionally heterogeneous monocytes and distinct dendritic cell subsets (cDC1, cDC2, plasmacytoid DC). Unexpectedly, we found that a majority of the equine peripheral B cell compartment is comprised of T-bet^+^ B cells; an immune cell subpopulation typically associated with chronic infection and inflammation in human and mouse. Taken together, our results demonstrate the potential of scRNA-Seq for cellular analyses in non-traditional model organisms, and form the basis for an immune cell atlas of horse peripheral blood.

## INTRODUCTION

The utility of studying non-traditional model organisms for biology, and particularly immunology, research is increasingly recognized. Experimental biology has been dominated by work in a relatively short list of well-studied model organisms such as the fruit fly and the mouse. Work in these species has many advantages including short generation times, tractability for genetic manipulation, high quality well-annotated genomes, and a large and diverse toolkit of well-validated reagents and protocols for experimentation (1). Despite the utility of these organisms in uncovering fundamental biological principles, they are not without their limitations. For example, they represent a small fraction of the diversity of multicellular life, and findings in any given experimental model do not always translate between species, as has been particularly well described for mouse-human translational studies (2). Many biological phenomena relevant to human health and society cross species boundaries, such as the circulation of established and/or emerging zoonotic pathogens in animal reservoirs (3) as well as animal health and its contribution to agriculture and the global food supply. As such, a holistic One Health approach to biology and immunology is essential and has been increasingly recognized by public health associations including the World Health Organization (4–6).

While the importance of studying diverse species is recognized, doing so can prove challenging due to a dearth of experimental tools which are available for more commonly investigated laboratory organisms. In immunology, flow cytometry is the traditional “gold standard” technique for defining cell subpopulations (7, 8). As flow cytometry and related mass cytometry technologies advance, with increasing capacity of parameters measured in a single panel, these methods can define immune cell subtypes at ever higher resolution (9). However, irrespective of resolution, flow cytometry ultimately relies on the availability of highly specific, well-defined antibodies against relevant surface markers (7). Such reagents are readily available from commercial sources for widely studied species such as mouse and human, but availability for other species can be limited. Raising, validating and conjugating custom antibodies to every target of interest is laborious, expensive, and not practical for most research laboratories. Furthermore, it requires *a priori* knowledge of cell type-defining surface markers.

Single cell RNA-Sequencing (scRNA-Seq) offers an alternative to flow cytometry as it defines different cell types (and their functional states) by gene expression patterns rather than surface marker expression. Recent advances in scRNA-Seq technology have enabled increased throughput and decreased cost per cell, allowing researchers to process thousands to tens-of-thousands of cells in a single experiment using droplet based microfluidics (10–12). scRNA-Seq offers many potential advantages for characterizing cell types in non-traditional model organisms including i) it can be applied to cells of diverse species without specialized reagents, ii) it does not rely on *a priori* marker selection or reagent availability, and iii) it can be used to identify novel markers for focused studies (13).

In this study, we demonstrate the potential of scRNA-Seq for discerning and discovering cell types in a non-traditional model organism, the horse. Equids are agriculturally and economically important globally, and host multiple zoonotic diseases including eastern equine encephalitis virus, Hendra virus, methicillin resistant *Staphylococcus aureus* (MRSA), *Salmonella spp., Leptospira spp.,* and *Streptococcus equi zooepidemicus* (14). Study of these natural infections in horses could be important from a One Health perspective in multiple aspects: i) to develop tools to prevent infection of horses with zoonotic diseases, ii) to break the chain of horse-to-human transmission, iii) to understand immunologic determinants of protection, clearance, and disease that could translate to improved understanding of human correlates, and iv) to improve the health of an economically important species.

Recent work from our laboratory and others established the current state-of-the-art flow cytometry protocols for immunophenotyping equine PBMC (15). However, compared to mouse and human, many immune cell subtypes remain to be defined at high resolution in the horse. Here, we used scRNA-Seq to characterize equine PBMC at unprecedented cellular resolution and present an immune cell atlas for horse peripheral blood. We identified 30 cell type clusters comprising major CD3^+^ lymphocyte, B cell, Monocyte/Dendritic Cell (DC) and Basophil cell populations. Clusters were annotated based on gene expression signatures, revealing several immune cell subtypes not previously described in horses. Interspecies comparisons with human PBMC scRNA-Seq datasets uncovered conserved blood DC subpopulations, and identified a spectrum of monocyte cell states similar to humans. Remarkably, we found that a large portion of the horse peripheral B cell compartment is comprised of T-bet^+^ B cells. Cellular analogs of this population in human and mouse are associated with chronic infections (16, 17).

## RESULTS

### Single cell RNA-Seq of equine PBMC resolves a diversity of immune cell types

We performed scRNA-Seq on fresh PBMC collected from 7 healthy adult horses. To generate broadly representative datasets, included horses spanned different breeds (Warmblood, Thoroughbred, Quarter Horse), ages (6-10 years), and both sexes (Table 1). In preliminary quality assessments of scRNA-Seq data processed with standard software workflows (10X Genomics Cell Ranger pipeline, EquCab3.0 reference genome with Ensembl v95 transcript annotations), we observed unexpectedly low numbers of genes detected per cell (Fig. S1A). Upon inspection of read mapping patterns for select genes known to be highly expressed in equine PBMC but found to be absent from our datasets, we frequently observed reads mapped immediately downstream of annotated transcript regions (Fig. S1B). This pattern is consistent with effective sequencing of transcript 3’ regions (as expected with 10X Chromium 3’ scRNA-Seq) but omission from quantification due to transcriptome annotations. In particular, incomplete annotation of transcript 3’ untranslated regions (UTRs, the most frequent transcript region captured by 10X Chromium 3’ scRNA-Seq) (18), is common in non-traditional model organisms relative to mouse or human reference transcriptomes (19). We therefore implemented an optimized data processing workflow that included the End Sequence Analysis Toolkit (ESAT, for “rescuing” reads mapped to unannotated 3’ transcript regions) (20), along with additional modifications (manually annotated immunoglobulin genes, quantification strategy for genes with multiple annotations, details in *Materials and Methods)* for this scRNA-Seq dataset. This approach significantly increased the number of genes detected per cell (Fig. S1A), and resulting output was used for all downstream analyses.

**TABLE 1:** Characteristics of horse study subjects

| **Horse** | **Sex** | **Age** | **Breed** | **Cells analyzed** |
| --- | --- | --- | --- | --- |
| Subject 1 | M | 6 | Warmblood | 4618 |
| Subject 2 | F | 8 | Thoroughbred | 5632 |
| Subject 3 | M | 7 | Warmblood | 5878 |
| Subject 4 | M | 8 | Thoroughbred | 4747 |
| Subject 5 | M | 8 | Thoroughbred | 4381 |
| Subject 6 | F | 9 | Quarter horse | 5200 |
| Subject 7 | F | 10 | Warmblood | 4222 |

Unsupervised graph-based clustering of 36,477 cells (post-quality filtering) resolved 31 clusters (Fig. 1A). Based on PCA hierarchical clustering and marker gene expression patterns (Fig. 1B, 1C), we grouped all clusters into 5 “major cell groups”: CD3^+^PRF1^-^ lymphocytes, CD3^+^PRF1^+^ lymphocytes, B cells, Monocytes/Dendritic cells (DCs), and Basophils. Complete marker gene lists for major cell groups are presented in Dataset S1. All major cell groups were represented at similar proportions across all 7 horses (Fig. 1D). To characterize equine PBMC at high resolution and establish a corresponding peripheral blood immune cell atlas, we independently analyzed scRNA-Seq data for the constituent clusters of each major cell group, except Basophils due to the low number of cells.

**Figure 1.**
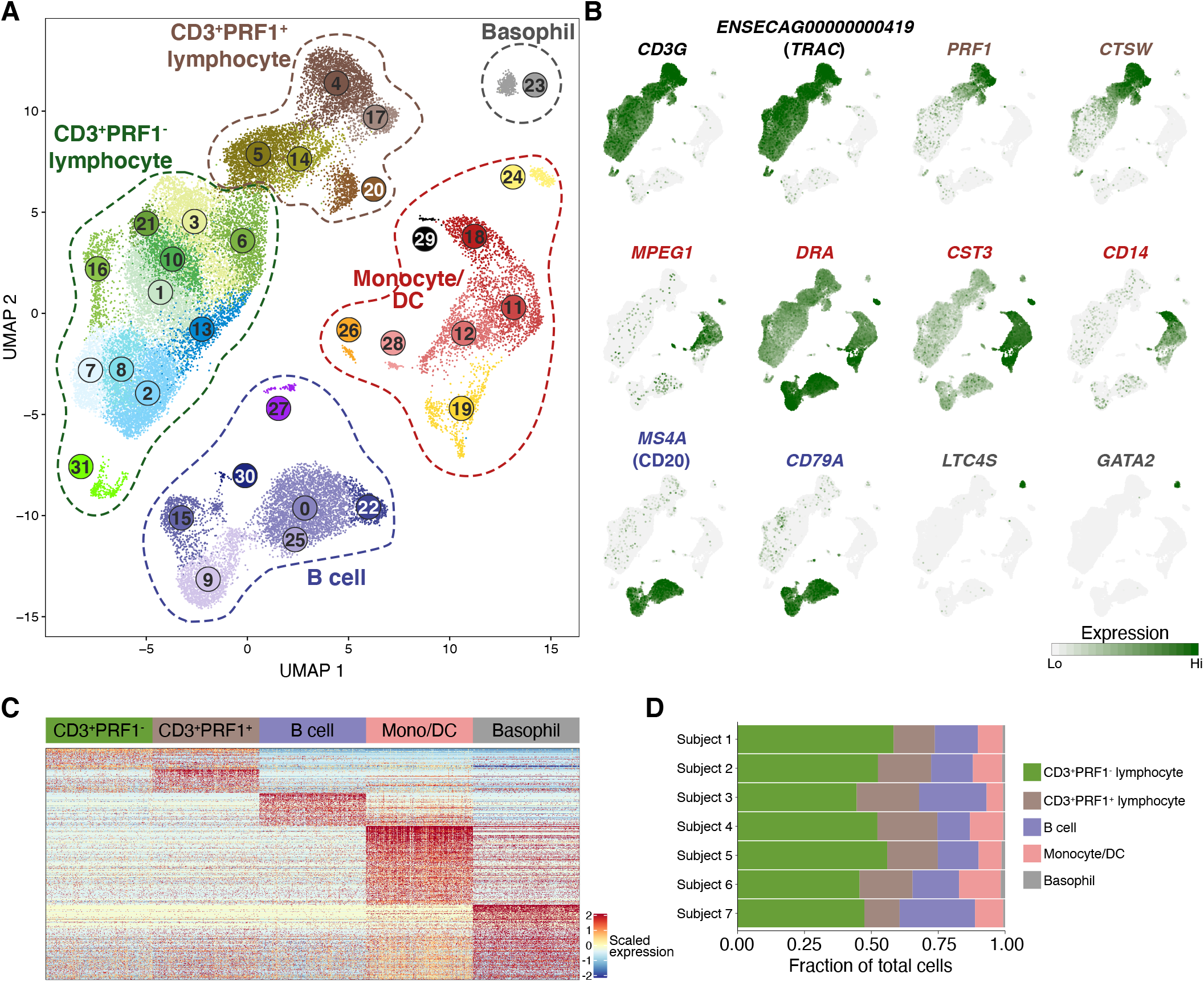
scRNA-Seq analysis of equine PBMC identifies five major cell groups. PBMC from N = 7 horses were processed for scRNA-Seq and analyzed with Seurat v3. Hierarchical clustering identified 31 cell clusters in 5 major cell groups, which were annotated based on marker gene expression. (A) UMAP visualization of equine PBMC (n = 34,677 total cells passing filter). Points (cells) are colored by cluster membership. Dashed outlines indicate major cell groups. (B) Gene expression patterns informing major cell group assignments. Expression values are scaled independently for each plot, ranging from 2.5 to 97.5 percentile of gene expression across all cells. Gene ID ENSECAG00000000419 is annotated as T Cell Receptor Alpha Chain C Region based on protein sequence homology (Ensembl) (C) Heatmap of genes differentially expressed (adjusted p-value < 0.05, log2 fold-change > 1 for each major cell group versus all other major cell groups) by each major cell group. For each major cell group, 30 cells (columns) were randomly selected from each horse for plotting purposes. (D) Frequency of each major cell group in total PBMC per horse.

### Peripheral equine myeloid cells include heterogeneous monocytes and distinct dendritic cell subsets with analogous counterparts in humans

We began with a detailed characterization of the Monocyte/Dendritic cell clusters (Fig. 2A; clusters 11, 12, 18, 19, 24, 26, and 28). Constituent monocyte/dendritic cell clusters were present in similar frequencies across all horses (Fig. 2B). Hierarchical clustering on integrated PCA data suggested two distinct subpopulations (Fig. 2C). Informed by this clustering pattern (Fig. 2C) and differential gene expression analysis (Dataset S2, Fig. 2D), we designated clusters 18, 11, 12, and 28 as monocytes, based on expression of the canonical marker gene *CD14* (21) (Fig. 2D, 2E). Similarly, we initially designated clusters 24, 19, and 26 as presumptive DCs based on high expression of MHC II antigen presentation genes *(DRA, DQA,* with notably elevated relative expression in clusters 19 and 24), and significantly lower *CD14* expression (Fig. 2D, 2F).

**Figure 2.**
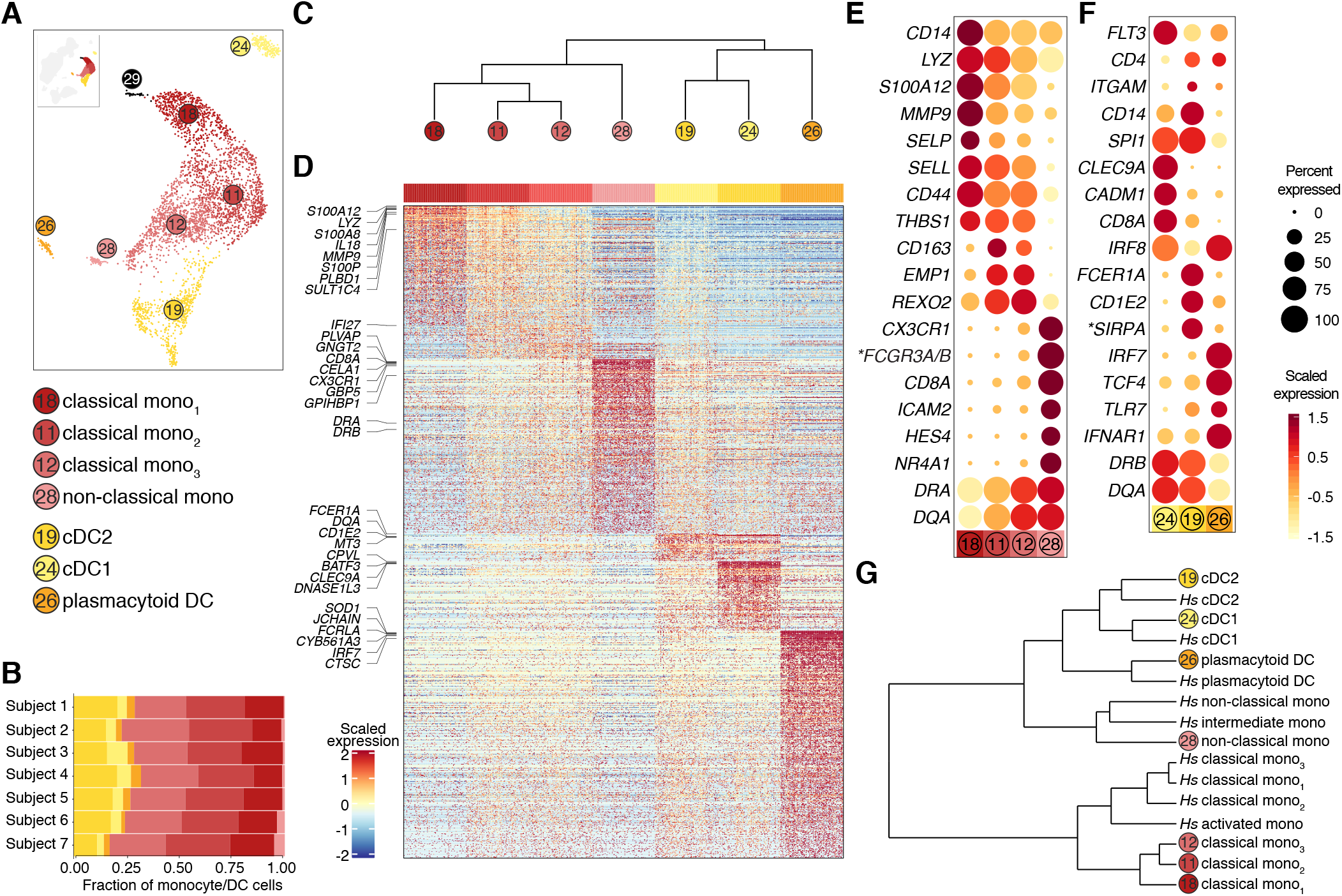
Equine monocyte/DC major cell group is comprised of diverse cell types including a range of monocyte transcriptional states and distinct dendritic cell subtypes. Clusters within the monocyte/DC major cell group were further analyzed and annotated by differential gene expression. Cluster 29 (annotated as neutrophils) was excluded from analysis due to low transcript (UMI) counts. (A) UMAP subset of monocyte/DC clusters with putative cluster annotations. (B) Frequency of each cell cluster within the monocyte/DC group per horse. (C) Hierarchical clustering (integrated PCA dimensions) of monocyte/DC major cell group. Dendrogram indicates two subpopulations, putatively annotated as mono-cytes (clusters 18, 11, 12, 28) and DCs (19, 24, 26). (D) Heatmap of genes differentially expressed (adjusted p-value < 0.05, log2 fold-change > 1 for each cluster versus all other clusters) by each cluster, with select genes labeled at left. (E) Dot plot of select genes differentially expressed across monocyte clusters. Dot size is proportional to number of cells with detectable expression of indicated gene. Dot color intensity indicates gene expression values scaled across plotted clusters. *Gene ID ENSECAG00000006663 is labeled *FCGR3A/B* based on Ensembl/NCBI annotations. (F) Dot plot of select genes differentially expressed across DC clusters. Additional details as in (E). (G) Hierarchical clustering of equine PBMC scRNA-Seq data (monocyte/DC clusters) and human PBMC scRNA-Seq data (monocyte/DC clusters). Median-normalized average expression values for highly variable human/horse one-to-one orthologues were calculated for each cluster, and clustering was performed on Pierson distances by Ward’s method.

Monocytes were composed of 3 abundant clusters (>94% of total monocytes, clusters 18, 11, and 12) and 1 relatively rare cluster (<6% of total monocytes, cluster 28). To assess the distinguishing features of these clusters, we performed differential gene expression analysis for each cluster within the monocyte group (clusters 18, 11, 12, 28). We identified genes with significantly elevated expression in each cluster (Dataset S3), including the monocyte/macrophage marker *CD163* (cluster 11), and numerous genes implicated in cell adhesion and migration such as *MMP9* (cluster 18), thrombospondin 1 *(THBS1* in clusters 11, 18), L-selectin *(SELL* in cluster 11, 18), P-selectin *(SELP* in cluster 18), and *ICAM2* (cluster 28) (Dataset S3). Of note, the top ranked (adjusted p value and log2 fold-change) differentially expressed gene in cluster 28 is ENSECAG00000006663, annotated as *FCGR3A/B* or *CD16,* a canonical marker for non-classical monocytes (CD14loCD16^+^ by flow cytometry) in human PBMC (22). Indeed, hierarchical clustering (Fig. 2C), and heat map (Fig. 2D) visualizations suggest that clusters 18, 11, and 12 exhibit somewhat similar and/or overlapping gene expression patterns, while cluster 28 is notably transcriptionally distinct. Clusters 18, 11, and 12 demonstrate varying expression of genes associated with classical monocytes (CD14^hi^CD16^-^ in humans, Ly6C^hi^CD44^+^ in mice) and/or intermediate monocytes (CD14^++^CD16^+^ in humans), including *CD14, CD44, SELL,* and the MHCII components *DRA* and *DQA* (Fig. 2E). Additional genes with significantly elevated expression levels in cluster 28 include *NR4A1* (transcription factor necessary for differentiation of non-classical monocytes in mice) (23), *CX3CR1* (chemokine receptor characteristic of non-classical monocytes in humans and mice) (24, 25), and *HES4* (target of NOTCH signaling implicated in non-classical monocyte generation) (26) (Fig. 2E). Taken together, these results suggest that equine monocyte populations are analogous to those described in humans and mice, with clusters 18, 11, and 12 most similar to classical monocytes, and cluster 28 to non-classical monocytes.

Presumptive DC clusters (24, 19, 26) were also analyzed by differential gene expression analysis (Dataset S4). Differentially expressed genes in cluster 24 included *CLEC9A, CADM1,* and *BTLA* (Fig. 2F, Dataset S4), all of which are immunophenotyping markers for cDC1 in humans and mice (27) (in mice, *CLEC9A* is also expressed on plasmacytoid DC (28)). Genes with significantly enriched expression in cluster 19 included *FCER1A* and *SIRPA* (Fig. 2F, Dataset S4), which are flow cytometric markers of cDC2 in humans and mice (Reviewed in (27)). DC subsets are also defined by the transcription factors that regulate their development and function, particularly by relative levels of IRF4 and IRF8 (27). Although *IRF4* transcripts were sparsely detected across all DC clusters (likely due to the incomplete sampling depth characteristic of droplet scRNA-Seq), *IRF8* was expressed at high levels in cluster 24 (cDC1) and significantly lower levels in cluster 19 (cDC2). Cluster 24 also exhibited high expression of *BATF3*, another characteristic transcription factor of cDC1 (29). In addition, top ranked differentially expressed genes in cluster 26 included *IRF7* and *TCF4* (E2-2) (Fig. 2F, Dataset S4), both of which are fundamental to plasmacytoid DC (pDC) development and function (30, 31).

To further support our cell type annotations and assess potential differences in monocyte/DC subsets between horses and humans, we performed cross-species hierarchical clustering with a human PBMC public reference scRNA-Seq data set (Fig. S2A-B, Fig. 2G). Equine clusters annotated as classical monocytes clustered first with each other, and next with human classical monocytes (defined by scRNA-Seq gene expression and confirmed with corresponding CD14/CD16 immunophenotyping feature barcoding data). This pattern likely reflects the heterogeneous transcriptional states of classical monocytes defined by *CD14* expression, and suggests broad similarities of this cell type across species. Equine non-classical monocytes clustered with human intermediate and non-classical monocytes. Remarkably, each DC subgroup clustered by cell type rather than species, indicating strong similarities of gene expression patterns between horse and human. These results further support three distinct DC subpopulations in horse peripheral blood that correspond with cDC1 (cluster 24), cDC2 (cluster 19), and pDC (cluster 26) in humans.

### The equine peripheral B cell compartment includes a large proportion of T-bet^+^ B cells

We next performed an in-depth analysis of B cell clusters, as defined by their expression of *MS4A1* (CD20), *CD79A,* MHC-II components (i.e. *DRA),* and/or immunoglobulin transcripts (Fig. 1A, 3A; Clusters 9, 15, 0, 22, 25, 27, and 30). Cluster 25 was excluded from analysis due to low transcript (UMI) counts (Fig. S3A). Remarkably, we observed six B cell clusters with distinct transcriptional profiles represented across all seven horses (Fig. 3B); this heterogeneity was somewhat surprising given our observation of only three B cell clusters (annotated as naïve, memory, and antibody secreting) in human PBMC scRNA-Seq data (Fig. S2A-B and additional datasets, *data not shown).* Hierarchical clustering on integrated PCA data suggested that clusters 27 and 30 were notably dissimilar from other B cell clusters, and based on recognizable gene expression patterns, we annotated cluster 27 as antibody secreting cells (ASCs, expressing *PRDM1*/BLIMP1, *XBP1, IRF4,* high levels of immunoglobulin transcripts) and cluster 30 as proliferating B cells (numerous G2/M associated genes including *PCNA, TOP2A,* and *UBE2C*) (Fig. 3C, D, E, Dataset S5). Of note, ASCs, which consistently exhibited high expression of a single immunoglobulin isotype per cell, demonstrated different isotype frequencies in different horses, perhaps indicative of distinct subclinical immune challenges (Fig. S3B).

**Figure 3.**
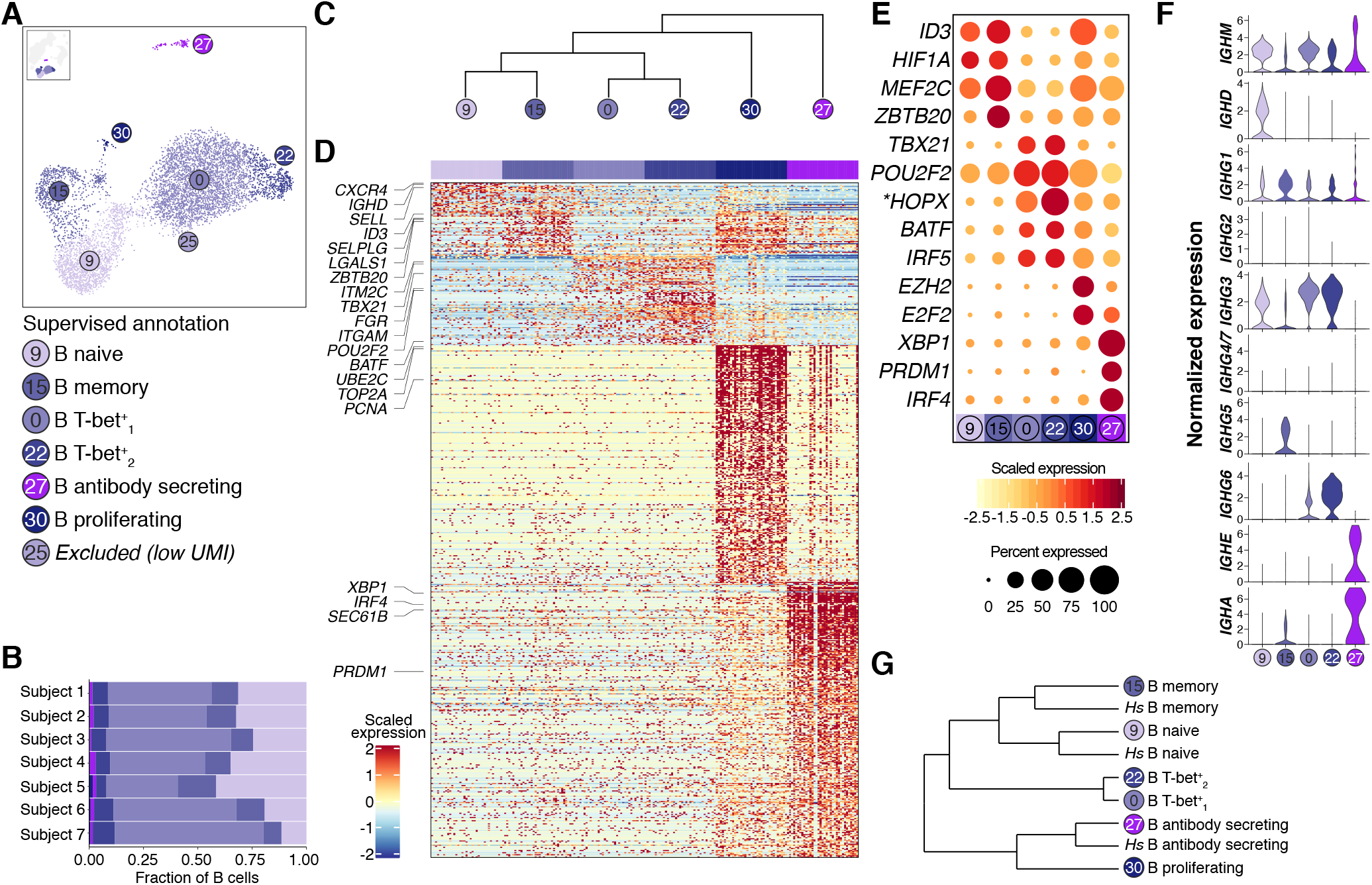
Equine peripheral B cell clusters include several distinct transcriptional states marked by expression of different transcription factors, including T-bet+ B cells. Clusters within the B cell major cell group were further analyzed and annotated by differential gene expression. Cluster 25 was excluded from analysis due to low transcript counts. (A) UMAP subset of B cell clusters with putative cluster annotations. (B) Frequency of each cell cluster within the B cell group per horse. (C) Hierarchical clustering (integrated PCA dimensions) of B cell major cell group. (D) Heatmap of genes differentially expressed (adjusted p-value < 0.05, log2 fold-change > 1 for each cluster versus all other clusters) by each cluster, with select genes labeled at left. (E) Dot plot of select transcription factor genes differentially expressed across B cell clusters. Dot size is proportional to number of cells with detectable expression of indicated gene. Dot color intensity indicates gene expression values scaled across plotted clusters. *Gene ID ENSECAG00000029287 is labeled *HOPX* based on Ensembl/NCBI annotation. (F) Violin plot of immunoglobulin heavy chain isotype transcript expression for indicated B cell clusters. Expression values are log-normalized per cell. (G) Hierarchical clustering of equine PBMC scRNA-Seq data (B cell clusters) and human PBMC scRNA-Seq data (B cell clusters). Median-normalized average expression values for highly variable human/horse one-to-one orthologues were calculated for each cluster, and clustering was performed on Pierson distances by Ward’s method.

**Figure 4.**
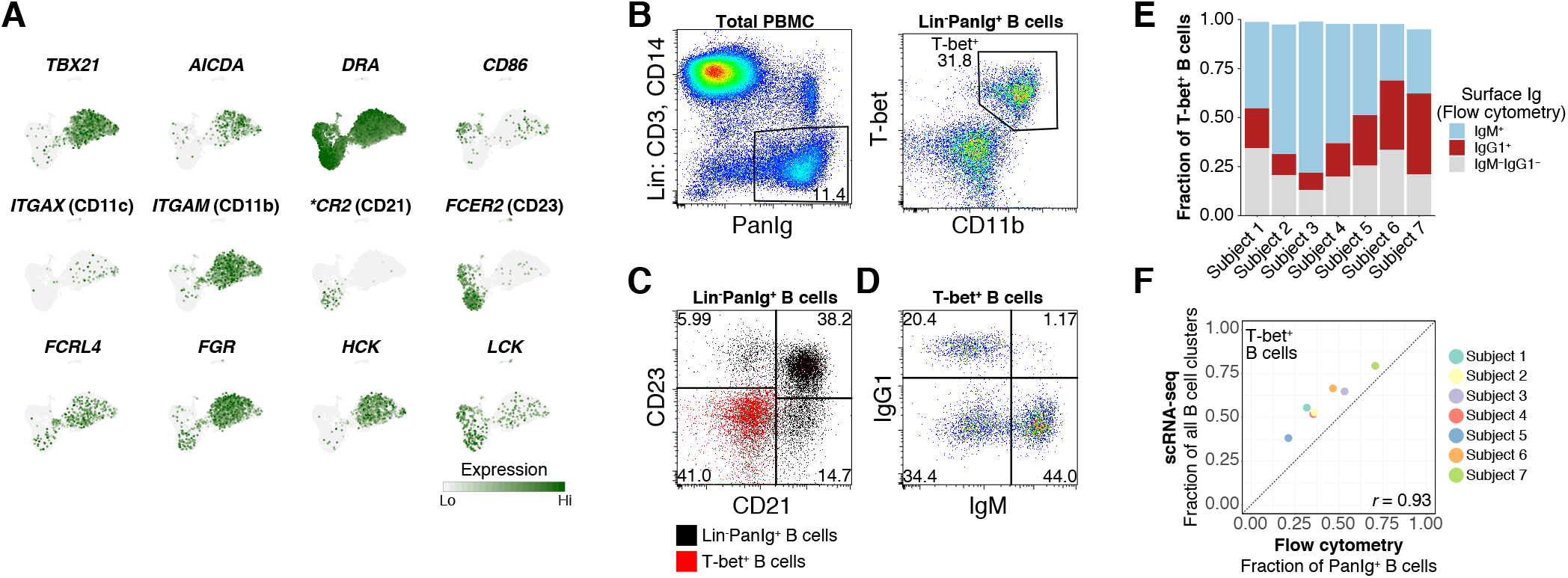
Equine T-bet^+^ B cells exhibit gene expression and cell surface protein marker expression characteristic of human T-bet^+^ B cells, and express diverse surface immunoglobulin isotypes. (A) Expression patterns for select genes associated with T-bet^+^ atypical B cells in humans and/or mice. Expression values are scaled independently for each plot, ranging from 2.5 to 97.5 percentile of gene expression across all B cell clusters. *Gene ID ENSECAG00000031055 labeled as *CR2* (CD21) based on Ensembl/NCBI annotation. (B) Flow cytometry gating strategy and expression of surface markers (PanIg and CD11b) and T-bet. (C) Expression of surface markers CD21 and CD23 in T-bet^4^-B cells. (D) Expression of surface immunoglobulin isotypes (IgM, IgG1) in T-bet^+^ B cells. (E) Quantification of surface immunoglobulin isotype expression by flow cytometry. (F) Correlation of T-bet^+^ B cells as determined by scRNA-Seq compared to flow cytometry. Pearson correlation coefficient, r = 0.93.

In reviewing the differentially expressed gene lists for the remaining B cell clusters, we noted the cluster-restricted expression of several transcription factors important for a variety of immune functions (Fig. 3E). Consistent with hierarchical clustering (Fig. 3C), these results further suggest that B cells in clusters 9 and 15 (expressing transcription factors *ID3*, *HIF1A* and *MEF2C*) employ a different gene regulatory program than B cells in clusters 0 and 22 (defined by specific expression of *TBX21*/T-bet, as well as elevated expression of *POU2F2/*Oct-2) (Fig. 3E). Given the fundamental relationship of immunoglobulin isotypes to B cell biology, we also explored the expression of immunoglobulin transcripts by different clusters. Using an updated immunoglobulin reference annotation (Dataset S6, adapted from (32), details in *Materials and Methods),* we observed different isotype transcript expression patterns in different B cell clusters (Fig. 3F). Importantly, and although we supplemented the transcriptome reference with these updated immunoglobulin transcript definitions during read mapping and quantification, all immunoglobulin genes were excluded from clustering to ensure clusters were not defined by isotype usage. Based on specific expression of *IGHD* transcripts and expression of *IGHM* transcripts, we annotated cluster 9 as naïve B cells. Relatedly, we annotated cluster 15 as likely memory B cells based on an overall gene expression pattern relatively similar to naïve B cells (cluster 9), but with the expression of class-switched isotype transcripts *(IGHG1, IGHG3, IGHG5, IGHA),* and the absence of *IGHD* transcripts. These cells are also defined by expression of *ZBTB20,* a transcription factor associated with antigen-experienced B cells (isotype-switched memory, germinal center, plasma cells) in mice (33), but they do not express plasma cell transcription factors such as *PRDM1*/Blimp-1 and *XBP-1* at appreciable levels (Fig. 3E). The *TBX21*/T-bet^+^ B cells in clusters 0 and 22 exhibited diverse isotype transcript expression patterns, which included both *IGHM* and class-switched isotypes *(IGHG1, IGHG3,* and *IGHG6,* most pronounced in cluster 22). With sequence data restricted to 3’ transcript regions (i.e. without coverage of variable region/constant region exon-exon junction), it was not possible to infer how these RNA expression patterns relate to functional/productive immunoglobulin protein expression.

We next performed cross-species hierarchical clustering on scRNA-Seq data for equine and human B cells (Fig. S2A-B). Consistent with our supervised annotations, equine cells annotated as naïve B cells clustered with human naïve B cells, equine cells annotated as memory B cells clustered with human memory B cells, and the equine cells annotated as antibody secreting clustered with the human ASCs (Fig. 3G). The equine T-bet^+^ B cells (clusters 0 and 22) appeared on a distinct branch of the clustering dendrogram. These results support our annotation of equine naïve and memory B cell populations, and suggest that the T-bet^+^ B cell clusters, which include the most abundant B cell cluster in horse peripheral blood, do not have a corresponding B cell population in PBMC from healthy humans (N = 2).

### Equine T-bet^+^ B cells share gene expression features with human T-bet^+^ B cells and can be identified in equine PBMC by flow cytometry

In humans, T-bet^+^ B cells have been described as “atypical memory B cells,” appearing in the peripheral blood during chronic infection and/or inflammation (17, 34). Although specific markers and/or gene expression patterns vary in different datasets, these B cells are often found to express *ITGAM* (CD11b), *ITGAX* (CD11c), as well as genes that modulate BCR signaling (including *FCRL4, FGR,* and *HCK)* (35–39). Moreover, a recent study of T-bet^+^ B cell populations in the context of chronic HIV infection demonstrated expression of genes associated with germinal center B cells (16, 40). We assessed expression of several of these characteristic genes in B cell clusters, and observed patterns consistent with multiple reports in humans (Fig. 4A). Among B cells, *ITGAM* (CD11b) expression was restricted to clusters 0 and 22, while *FCER2* (CD23) was virtually absent from these clusters. Although sampling for *ITGAX* (CD11c) and *ENSECAG00000031055* (annotated as *CR2*/CD21) was insufficient for differential expression testing, we detected *ITGAX* (CD11 c) positive cells in T-bet^+^ clusters 0 and 22 (Fig. 4A). Moreover, we detected significantly elevated expression of *FCLR4, FGR,* and *HCK* in these clusters (Fig. 4A, Dataset S5).

Based on these scRNA-Seq expression patterns, we developed a flow cytometry panel to identify equine T-bet^+^ B cells by protein expression. Since T-bet exhibits 92% amino acid sequence identity between horses and humans (GenPept accessions XP_023508425.1 and NP_037483.1, respectively), we selected an anti-human T-bet antibody for intracellular labeling. We then adapted a previously validated equine PBMC immunophenotyping panel (15) to identify CD3^-^CD14^-^PanIg^+^ B cells (Fig. S4). As predicted from our scRNA-Seq data, we detected an abundant CD11b^+^ B cell population with high expression of T-bet (Fig. 4B) that did not express surface CD21 or CD23 (Fig. 4C). T-bet expression was not detected in other B cell gates. We also assessed surface isotype usage of T-bet^+^ B cells by flow cytometry; 51 ± 18% T-bet^+^ B cells were IgM^hi^, 23 ± 12% were IgG1^+^, and 24 ± 8% expressed neither IgG1 nor IgM (Fig. 4D, E). It is unclear whether IgM^hi^ T-bet^+^ B cells reflect an antigen-inexperienced naïve subset, a recently activated subset, or a memory cell subset that did not undergo class switch recombination. By flow cytometry, T-bet^+^ B cells comprised 44 ± 17% of total B cells, a percentage that was correlated, but consistently lower, to the percentage observed in scRNA-Seq data (Fig. 4F), perhaps reflecting incomplete sensitivity of PanIg antibody labeling for all B cells.

Taken together, our flow cytometry results validate the existence of a novel population of T-bet^+^ B cells initially identified by scRNA-Seq analysis, which also demonstrated similarities with human T-bet^+^ B cells associated with chronic infection and inflammation.

### CD3^+^PRF1^+^ clusters include lymphocytes with diverse cytotoxic gene expression patterns

The *CD3^+^PRF1^+^* major cell group is composed of five clusters (Fig. 5A; Clusters 4, 5, 14, 17, and 20), which were represented at similar frequencies across all horses examined (Fig. 5B). All clusters were characterized by expression of the cytotoxic effector *PRF1* and *CTSW,* a cathepsin whose expression is associated with cytotoxic capacity (41) (Fig. 1B). Based on hierarchical clustering of integrated PCA data, we partitioned our annotations into three distinct transcriptional programs (Fig. 5C). Although all clusters expressed high levels of CD3 transcripts (*CD3D*, *CD3E*, *CD3G*, Fig. S5A), based on differential gene expression results (Dataset S7), this major cell group likely includes both cytotoxic T cells and NK cells.

**Figure 5.**
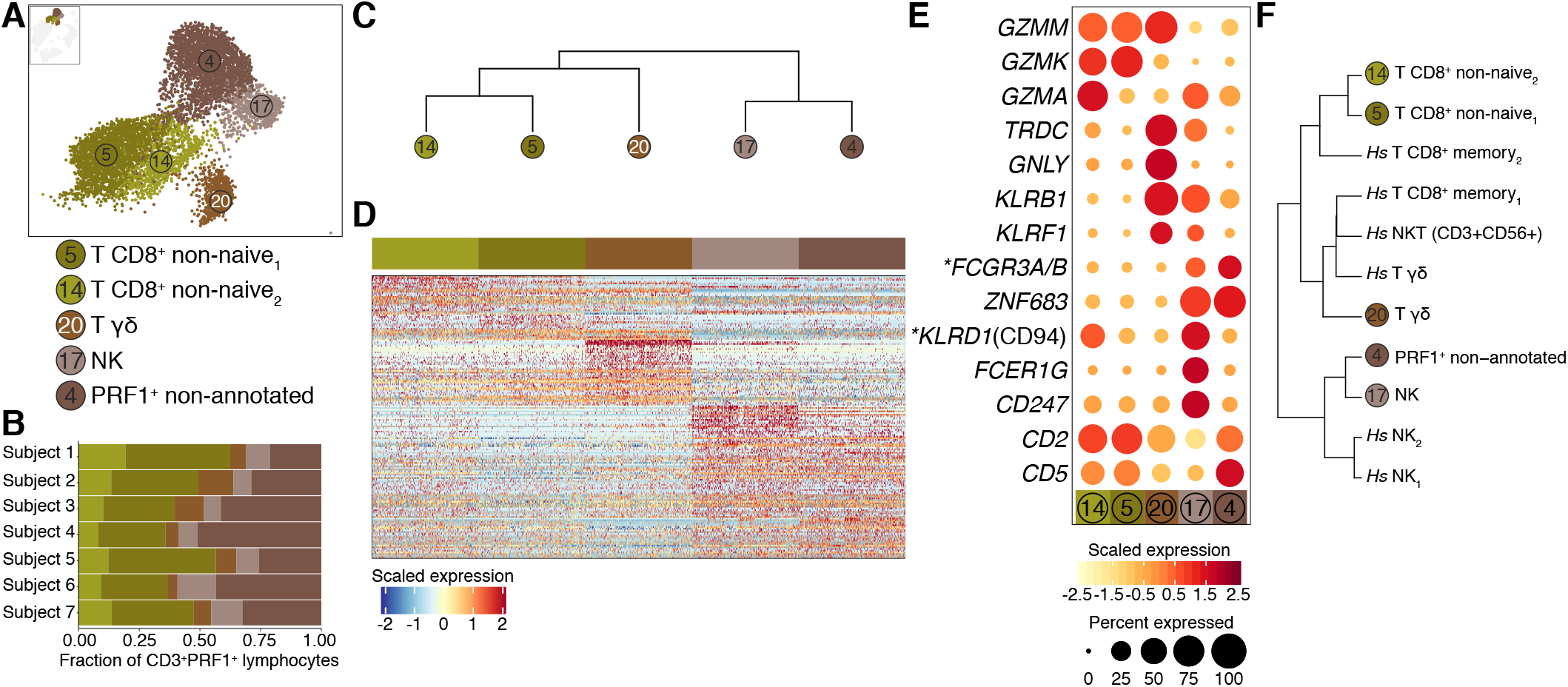
The CD3+PRF1+ clusters contain various cell types with different gene expression patterns characteristic of cytotoxic lymphocyte function. Clusters within the CD3^+^PRF1^+^ lymphocyte major cell group was further analyzed and annotated by differential gene expression. (A) UMAP subset of CD3^+^PRF1^+^ lymphocyte clusters with putative cluster annotations. Selected axis ranges excluded <5 cells in CD3^+^PRF1^+^ group from plot. (B) Frequency of each cell cluster within the CD3^+^PRF1^+^ lymphocyte group per horse. (C) Hierarchical clustering (integrated PCA dimensions) of CD3^+^PRF1 ^+^ lymphocyte clusters major cell group. (D) Heatmap of genes differentially expressed (adjusted p-value < 0.05, log2 fold-change > 0.58 for each cluster versus all other clusters) by each cluster, with select genes labeled at left. (E) Dot plot of select genes associated with cytotoxic lymphocyte populations differentially expressed across CD3^+^PRF1^+^ lymphocyte clusters. Dot size is proportional to number of cells with detectable expression of indicated gene. Dot color intensity indicates gene expression values scaled across plotted clusters. *Gene ID ENSECAG00000006663 is labeled *FCGR3A/B* and Gene ID ENSECAG00000031528 is labeled *KLRD1* (CD94) based on Ensembl/NCBI annotations. (F) Hierarchical clustering of equine PBMC scRNA-Seq data (CD3^+^PRF1^+^ lymphocyte clusters) and human PBMC scRNA-Seq data (non-naive CD8^+^ T cells, NK cells, NKT cells). Median-normalized average expression values for highly variable human/horse one-to-one orthologues were calculated for each cluster, and clustering was performed on Pierson distances by Ward’s method.

Transcriptional profiling studies of human and mouse cells often describe challenges in distinguishing cytotoxic lymphocyte subpopulations, with memory αβ CD8^+^ T cells, NK cells, NKT cells, and γδ T cells exhibiting considerable overlap in gene expression patterns (42–46). Our data suggest similar overlap exists among equine cytotoxic lymphocyte subpopulations. We annotated clusters 5 and 14 as CD8^*+*^ “antigen experienced” or “non-naïve” T cells. While overall quite similar, cluster 5 exhibited features more consistent with CD8^+^ T central memory cells in humans (GZMK/GZMM protein expression, absence of GZMA protein), while cluster 14 exhibited features more consistent with CD8^+^ T effector memory cells (GZMK/GZMM/GZMA protein expression) (47). We emphasize that these expression patterns are not fully distinct and are unlikely to correspond perfectly to subpopulations defined by traditional flow cytometric markers. Furthermore, although these cells appear to share common cytotoxic gene expression programs, we observed notable within-cluster heterogeneity. Indeed, cluster 5 contained mutually exclusive subgroups of cells which expressed either *CD8A* or *CD4* (Fig. S5B).

Although expressing many of the same cytotoxic effector genes, cluster 20 appeared distinct from other cytotoxic lymphocyte clusters (Fig. 5A, 5C). Differential gene expression analysis revealed highly significant elevated expression of *TRDC.* Cells in cluster 20 also expressed lower levels of *TRAC, TRBC1,* and *TRBC2* relative to other cytotoxic lymphocyte clusters (Fig. S5C). Based on these expression patterns, we annotated cluster 20 as cytotoxic γδ T cells. Interestingly, this cluster demonstrated high and specific (or nearly specific) expression of several genes associated with cytotoxicity, including *GNLY, KLRB1,* and *KLRF1.* These results support the existence of equine γδ T cells, which have not been definitively characterized. Moreover, they suggest that these cells employ unique cytolytic mechanisms compared to other equine cytotoxic lymphocytes.

The remaining *CD3^+^PRF1^+^* clusters (17 and 4) exhibited gene expression patterns consistent with both cytotoxic T cells and NK cells. Both clusters demonstrated high expression of TCR complex components, including *CD3D*, *CD3E*, *CD3G*, TCR alpha chain (*TRAC*, ENSECAG00000000419), and TCR beta chain *(TRBC1,* ENSECAG00000033316; *TRBC2,* ENSECAG00000030258) (Fig. S5C). However, both clusters also displayed expression of genes associated with NK cell function, including *FCGR3A/B* (CD16, employed by NK cells for antibody-dependent cellular cytotoxicity), and *ZNF683* (HOBIT, a transcription factor highly expressed by human NK cells (48) and used as a surrogate for equine NK cells by RT-PCR (49) (Fig. 5E); *ZNF683* has also been described in human cytotoxic T cell subsets (50). We annotated cluster 17 as NK cells based on specific expression of ENSECAG00000031528 (annotated as *KLRD1*/*CD94*), which encodes the cell surface lectin central to NKG2 functions (Fig. 5E). This cluster also exhibited specific expression (within the cytotoxic lymphocyte major cell group) of *FCER1G* and *CD247,* both of which are important for NK cell activation signal transduction (51). In addition, and in contrast to cluster 4, this putative NK cell cluster exhibited diminished or absent expression of *CD2* and *CD5*, genes frequently used as T cell markers in humans (52) (Fig. 5E). Of note, multiple descriptions of equine NK cells by flow cytometry or immunohistochemistry have purposefully excluded CD3^+^ cells (53–55). However, consistent with scRNA-Seq, our flow cytometric analysis identified a well-defined CD3^+^CD16^+^ lymphocyte population (Fig. S6A). Given their expression of TCR transcripts, it remains unclear whether these cells have the capacity to respond to specific antigen presented by traditional MHC-I or MHC-II.

Although cluster 4 has gene expression patterns consistent with both cytotoxic T cells and NK cells, the absence of definitive marker genes and/or genes associated with NK cell-restricted functions made it challenging to annotate this cluster. Based on the overlapping gene expression programs described in cytotoxic lymphocytes in better characterized species, we suspect this cluster could include an additional type of CD8^+^ cytotoxic T cells, semi-invariant TCR cytotoxic T cells (e.g. Mucosal-associated invariant T cells, NKT cells), and/or an additional type of NK cell. The latter possibilities are further supported by cross-species comparison to human cytotoxic lymphocytes (Fig. 5F). Alternatively, cluster 4 may represent a novel type of cytotoxic lymphocyte unique to horses.

### CD3^+^ PRFT^-^ clusters include naïve T cells and heterogeneous CD4^+^ T cell populations

The CD3^+^PRF1^-^ T cell major cell group is composed of 11 clusters, including proliferating T cells (cluster 31, Fig. 6A), which were represented at similar frequencies across all horses examined (Fig. 6B). Although the most abundant in both cell and cluster numbers, these subpopulations were also the most challenging to effectively annotate, due in large part to the relatively subtle transcriptional differences detected between most clusters. In our experience, resting T cell populations can be difficult to distinguish by droplet microfluidics scRNA-Seq data. Despite these limitations, we were able to make several observations regarding the non-cytotoxic T cell constituent clusters. First, we distinguished naïve T cells (clusters 2, 7, 8) based on elevated expression of *CCR7, SELL* (L-selectin), and the *LEF1* transcription factor (Fig. 6C). Naïve T cells could be further partitioned into *CD4^+^* (Clusters 2, 8, not significant by differential gene expression) and *CD8^+^* (Cluster 7) subpopulations (Fig. 6C). Although the remaining “non-naïve” clusters (presumably antigen experienced) exhibited significant gene expression differences, we were not able to confidently assign clusters to T cell subsets traditionally defined by flow cytometry (e.g. memory Th1, memory Th2, memory Th17, etc.).

**Figure 6.**
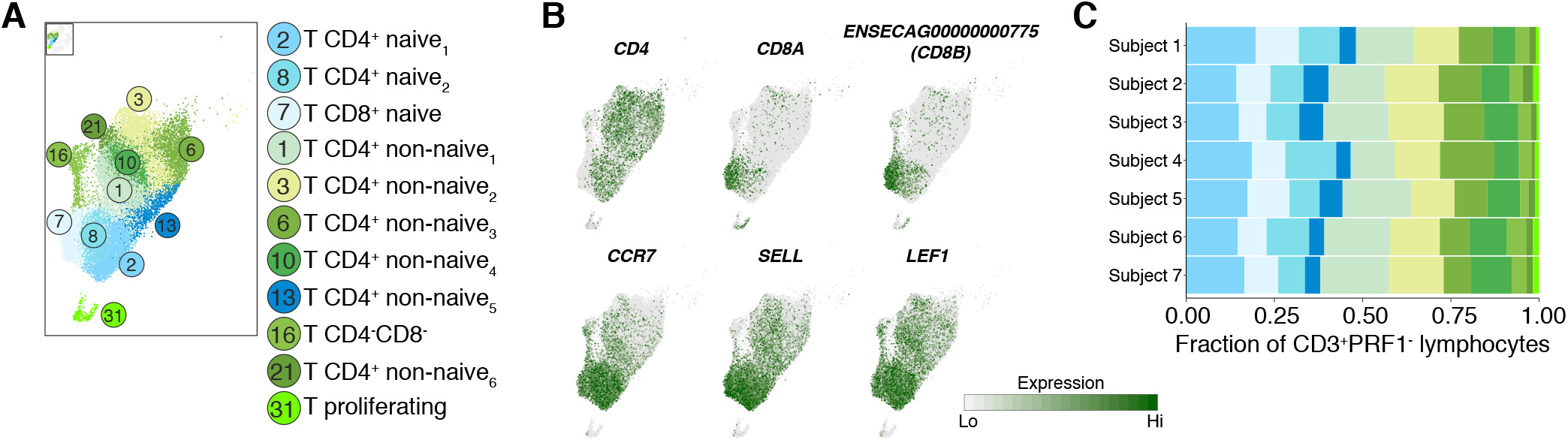
The CD3^+^PRF1^-^ clusters include naïve CD4^+^ and CD8^+^ T cells and additional CD4^+^ T cell populations. Clusters within the CD3^+^PRF1^-^ lymphocyte major cell group were further analyzed and annotated by differential gene expression. (A) UMAP subset of CD3^+^PRF1^-^ lymphocyte clusters with putative cluster annotations. Selected axis ranges excluded <10 cells in CD3^+^PRF1^-^ group from plot. (B) Expression patterns for genes characteristic of CD4/CD8 T cell subsets, and naïve T cell populations *(CCR7, SELL, LEF1).* Expression values are scaled independently for each plot, ranging from 2.5 to 97.5 percentile of gene expression across all CD3^+^PRF1^-^ cells. Gene ID ENSECAG00000000775 labeled as *CD8B* based on Ensembl/NCBI annotation. (C) Frequency of each cell cluster within the CD3^+^PRF1^-^ lymphocyte group per horse.

### High resolution landscape of equine peripheral blood mononuclear cells

Given the improved resolution and novel cell populations identified by scRNA-Seq, we grouped annotated cell clusters into summary populations and provided “reference ranges” for their frequency in healthy horses (Fig. 7A). We also evaluated how the frequencies of major cell groups defined by scRNA-Seq compared to those that can be resolved by current state-of-the-art flow cytometry equine PBMC immunophenotyping (adapted from (15)). Populations defined by flow cytometry gating (Fig. S5) were matched to corresponding scRNA-Seq clusters (grouped as indicated, Fig. 7B). Of note, flow cytometry data was acquired from the same samples processed for scRNA-Seq, and the researcher (J.E.T.) performing gating and analyses was blinded to frequencies determined by scRNA-Seq. Overall, cell frequencies determined by scRNA-Seq were strongly correlated with frequencies determined by flow cytometry (*r* = 0.93 for indicated populations examined). We observed strong concordance in frequencies of CD4^+^ T cells, classical monocytes, and non-classical monocytes (Fig. 7B). We consistently measured higher frequencies of NK cells and B cells by scRNA-Seq, which suggests that current flow cytometry definitions (NK cells: CD3^+^CD16^+^; B cells: Pan-Ig^+^) based on available equine-reactive antibodies might not capture these cell populations entirely.

**Figure 7.**
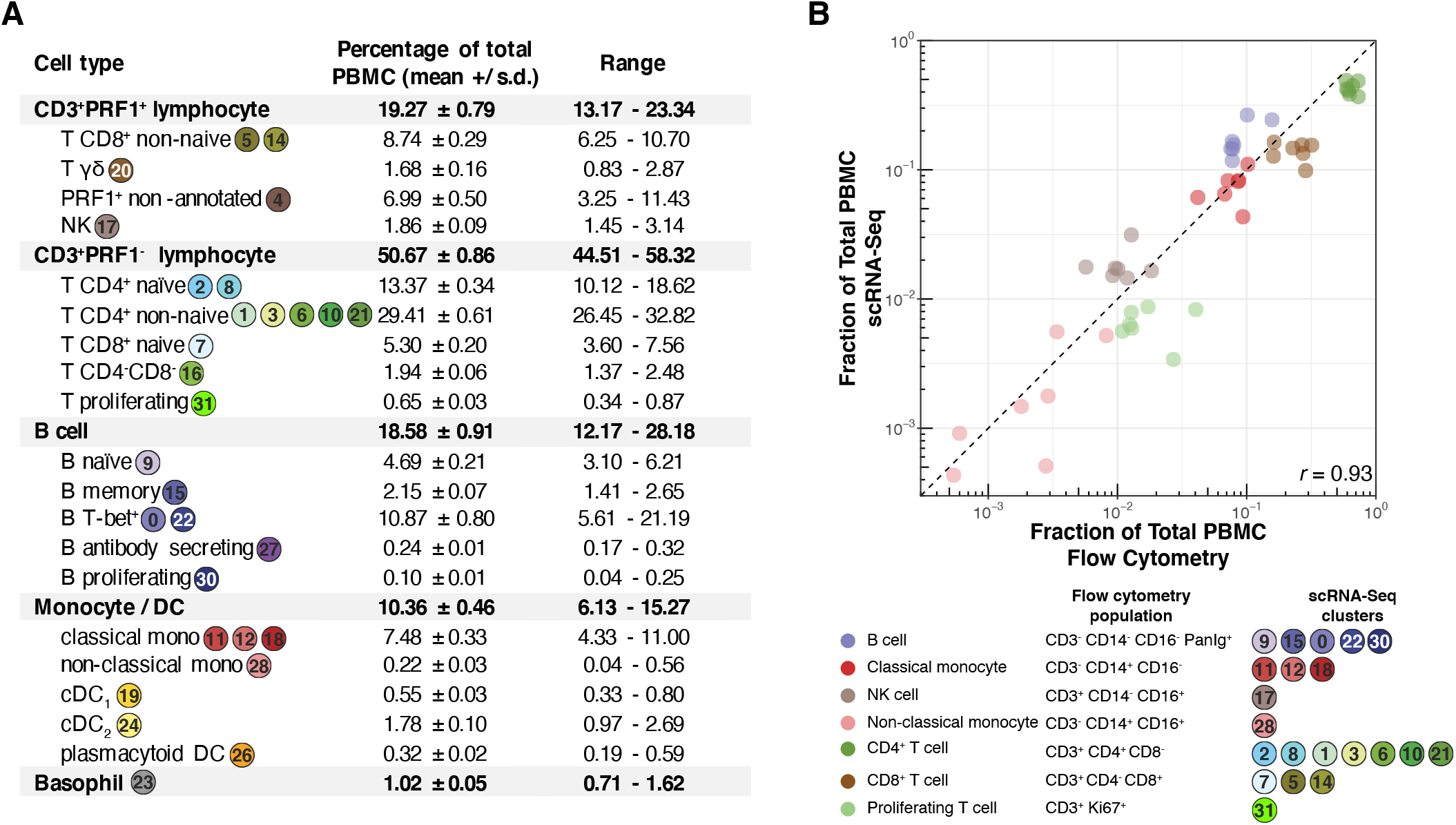
High resolution landscape of equine peripheral blood mononuclear cells. (A) Cell type frequencies determined by scRNA-Seq. Annotated clusters were grouped to summary populations as indicated. (B) Correlation of population frequencies for select cell clusters defined by scRNA-Seq with corresponding populations that can be resolved by current state-of-art flow cytometry. Each point indicates cell population frequency as a fraction of total PBMC from an individual subject. Cell type definitions by flow cytometry markers and scRNA-Seq cluster summaries as indicated. Pearson correlation coefficient, *r* = 0.93.

Taken together, these results demonstrate that our scRNA-Seq cell cluster annotations are consistent with state-of-the-art flow cytometry methods, but can resolve cell populations at much higher resolution and sensitivity.

## DISCUSSION

Here, we used scRNA-Seq to define the cellular landscape of equine peripheral blood immune cells. This study demonstrates proof-of-concept for the utility of high throughput scRNA-Seq as a tool to characterize distinct cell types in species for which traditional flow cytometric tools are limited. Furthermore, interspecies analyses demonstrate the potential of scRNA-Seq for comparative studies exploring the evolution of cellular specialization of the immune system.

Prior to this work, state of the art equine PBMC immunophenotyping methods (15) allowed for effective resolution of CD4^+^ T cells, CD8^+^ T cells, and B cells. Additional panels were developed for this study, which identified proliferating T cells, NK cells, CD14^+^ monocytes, CD16^+^ monocytes, and T-bet+ B cells. Using scRNA-Seq, we resolved 30 cell populations based on variable gene expression patterns. These populations could be partitioned into five major cell groups: Monocyte/DC, B cell, CD3^+^PRF1^+^ lymphocytes, CD3^+^PRF1^-^ lymphocytes, and Basophils. Combining supervised annotation based on prior knowledge and comparative cross-species clustering, we identified multiple distinct subpopulations with immunologically relevant gene expression patterns within these major cell groups. Many of these subpopulations have not been previously described in horse peripheral blood. Cross-species analyses demonstrated that many equine immune cell subpopulations have corresponding populations identifiable in humans. However, we also identified several immune cell populations (e.g. T-bet^+^ B cells, discussed below) absent from healthy human peripheral blood. Overall, our study establishes a cellular atlas of equine PBMC in healthy horses, and demonstrates proof-of-concept for characterizing complex cell populations in non-traditional model organisms.

The mononuclear phagocyte system is comprised of diverse cell types that carry out many different functions in innate and adaptive immune responses. Our analysis of the monocyte/DC major cell group revealed cellular heterogeneity and subpopulations consistent with other species, as well as subpopulation-specific gene expression programs with functional implications for the equine immune system. Monocyte subsets, which are often categorized as classical, intermediate, and non-classical (based on surface expression of CD14 and CD16 in humans (22)), have been described as generally conserved across mammalian species (56), and CD16^+^ monocytes have been previously observed in horses (53). Indeed, our results support the existence of similar monocyte subsets in horse and human peripheral blood. Moreover, the gene expression data supplied by scRNA-Seq offer insight into the putative functions of these cellular subsets. These include distinct trafficking programs (based on adhesion molecules and chemokine receptors) and antigen presentation capacity (expression of MHC-II genes is generally anti-correlated with CD14, with highest expression in the classical mono3 cluster, followed by non-classical mono). These data also include potential novel surface markers (e.g. CD8A for non-classical monocytes) for improved immunophenotyping by flow cytometry, though RNA transcript levels may not necessarily correspond with surface protein expression. We also identified three dendritic cell clusters, with gene expression consistent with the cDC1, cDC2, and pDC subsets described in humans (27) and mice (57). Previous studies of equine DCs have relied on monocyte-derived DCs *in vitro* (58–60). Recently, Ziegler *et al.* identified DCs in equine peripheral blood based on FLT3L binding as detected by flow cytometry (61). Guided by marker expression in other species, they proposed defining three putative DC subsets as cDC1 (Flt3^+^CD4^-^CD13^+^CD14^low^CD172a CADM-1^+^MHCII^high^), cDC2 (Flt3^+^CD4^-^ CD13^-^ CD14^low^CD172a^+^ CADM-1^low^MHCII^high^), and pDC (Flt3^+^CD4^-^CD13^-^CD14^-^CD172a^-^CADM-1^-^MHCII^low^). With the exception of CD13 *(ANPEP,* not detected in our scRNA-Seq data), RNA transcript expression for these surface markers in DC clusters identified in our study was entirely consistent with these definitions. Our transcriptomic data not only provide additional support for these immunophenotyping markers, but also define these subsets by gene expression pattern characteristic of their functions. These include distinguishing transcription factor profiles of cDCs (high *IRF8* in cDC1, low *IRF8* in cDC2, *SPI1/PU.1* expression in both cDC1 and cDC2), elevated expression antigen presentation machinery in cDCs *(CD74, DRA, DRB,* others), and steady-state expression of genes important for virus recognition and interferon response in pDC *(TLR7, IRF7, IFNAR1*). Our data also indicate that, although they share many features with corresponding DC subsets in other species, equine DC subsets exhibit notable differences. For example, we did not detect *ITGAM* (CD11b), a marker for murine cDC1 (57), at appreciable levels in equine cDC1. Informed by the gene expression programs defined here, additional experimental characterizations are necessary to definitively assign functions to these different subsets, each of which is likely to play a critical role in equine immunity.

In addition to identifying and/or validating cell types that we anticipated would be present in equine peripheral blood, the unsupervised clustering approach to scRNA-Seq data also revealed previously undescribed and unexpected cell populations. Within the B cell compartment, we expected to detect clusters consistent with naïve B cells, memory B cells, and antibody secreting cells, based in part on human PBMC scRNA-Seq data and equine PBMC flow cytometry data (15). In addition to these clusters, we were surprised to observe two additional clusters of apparent B cells (as defined by high expression of *MS4A1*/CD20 and immunoglobulin transcripts) with gene expression patterns notably distinct from presumptive naïve and memory clusters. Cells in these clusters, characterized by specific expression (within the B cell major cell group) of the T-bet transcription factor *(TBX21)*, were the most abundant B cells across all seven horses under investigation. Corresponding populations of T-bet^+^ B cells were not observed in human PBMC scRNA-Seq data, and have not been previously described in horses. In mice, T-bet^+^ B cells have been shown to be important for antiviral humoral immunity (34, 62, 63). In humans, T-bet^+^ B cells have been detected in peripheral blood in a variety of chronic inflammatory contexts including systemic lupus erythematosus (64, 65), chronic malaria exposure (66, 67), and chronic viral infection (16, 40, 68). Although a universal definition and designated function for these cells remains elusive, T-bet^+^ B cells are often classified as atypical memory B cells and, at least in some contexts, are thought to arise from repetitive BCR stimulation (17). Indeed, human T-bet^+^ B cells are enriched among virus-specific B cells in HIV infection (16). The equine T-bet^+^ B cell populations identified in the present study share many features with the atypical memory B cell populations described in humans. In addition to *TBX21*/T-bet expression, horse T-bet^+^ B cell populations exhibit similar gene expression patterns, including enriched expression of *FCRL4*, *FGR,* and *HCK,* the protein products of which modulate BCR signaling (38). Furthermore, a recent study of T-bet^+^ B cells in HIV demonstrated that they express genes characteristic of germinal center B cells (16). Interestingly, we detected specific expression of *AICDA* (encoding activation-induced cytidine deaminase) and elevated expression of *APEX1* in equine T-bet^+^ B cells, suggesting that the T-bet^+^ B cells identified here may represent equine equivalents of the T-bet^+^ atypical memory B cells described in humans. If these cells are elicited by chronic antigenic stimulation, it is plausible that horses chronically exposed to numerous pathogens common in standard boarding conditions (e.g. equine alpha and gamma herpesviruses, influenza, rhinitis viruses, hepacivirus, parvovirus-hepatitis, coronavirus, etc.) could expand this population. While viral exposure burdens are likely to be largely similar to burdens experienced by humans, horses in the northeastern U.S. are also frequently exposed to *Borrelia burgdorferi* (agent of Lyme disease) (69) and *Sarcocystis neurona* (agent of Equine protozoal myeloencephalitis) (70), and are continuously infested with or re-exposed to gastrointestinal nematodes (71). The horses in this study did not show signs of active infection or inflammation, as they all had normal complete blood count, serum amyloid A, iron indices, and globulins. Moreover, the surprisingly high frequency of these T-bet^+^ cells within the B cell compartment suggests that they may provide important functions in the sustained immune responses to such pathogens. The impact of pathogen exposure on the genesis of this B cell population might be further explored by experiments in foals and/or in pathogen-free facilities, informed by the scRNA-Seq data and flow cytometric strategies established here. Additionally, horses may represent a useful model organism in which to study further this unique B cell population given their abundant frequency and ready accessibility of large amounts of blood and other tissues, such as lymph nodes.

scRNA-Seq is molecularly compatible with presumably any animal species as most droplet microfluidics scRNA-Seq platforms select mRNAs for barcoding and downstream sequencing based polyA tails, a feature common across metazoans. Additional requirements for scRNA-Seq analysis include a genome (or at minimum, transcriptome) sequence to which reads are mapped, and gene/transcript annotations against which mapped reads can be quantified. Should transcriptome annotations be insufficient for robust scRNA-Seq analysis, as may be the case for less commonly studied organisms, read assignment/quantification strategies can be modified with specialized software tools (e.g. ESAT (20), as implemented here) and/or annotations can be supplemented/replaced with custom annotations derived from bulk RNA-Seq data. Interpretation of scRNA-Seq results can be greatly facilitated by gene/transcript annotations with comprehensive orthologue annotations for multiple species, but this is not a requirement. Without the need for species-specific reagents, and a constantly growing catalog of species with sequenced and annotated genomes, we anticipate that scRNA-Seq will be an increasingly useful research tool for non-traditional model organisms.

Despite the many insights gained from our PBMC analyses, scRNA-Seq, particularly for characterizing cell mixtures from diverse animals, is not without limitations. In the present study, although defining subpopulations with unsupervised clustering methods was reasonably straightforward, assigning putative cell types to each cluster presented challenges. Ideally, automated cell type classification based on external datasets and/or prior knowledge could minimize biases introduced by supervised annotation (72, 73). Recently developed scRNA-Seq data integration and cluster annotation tools have begun to implement this functionality (74–76). We made attempts to apply several of these strategies in comparing equine PBMC to human PBMC, but observed generally poor performance, which we attributed to insufficient interspecies orthologue annotations *(data not shown).* Instead, we adopted a supervised approach based on prior knowledge of human and mouse immune cells to assign likely cell types. We, therefore, emphasize that our presumptive cell type annotations are not definitive, and ultimately require experimental validation by complementary methods, as we pursued with flow cytometry for T-bet^+^ B cells (Fig. 4). Furthermore, for many clusters, most notably in the CD3^+^PRF1^-^ lymphocytes major cell group, we were unable to confidently assign cell types due to limited detection of informative differentially expressed genes. This could be a result of suboptimal clustering (i.e. heterogeneous clusters), relatively low transcript sampling depth, and/or discrepancies in mRNA and corresponding protein expression by which T cell subsets have been previously defined. Many of these issues are likely to be mitigated in the future by perennially improving genome and orthologue annotations, scRNA-Seq methodologies with increased per cell sampling depth, and novel software tools for intra- and inter-species data analyses.

## MATERIALS AND METHODS

### Research subjects and cells

Horses studied here consisted of 3 mares and 4 geldings, 6 to 10 (mean 7.9) years old, 3 Warmbloods, 3 Thoroughbreds, and one Quarter Horse. Horses were determined to be healthy by physical examination, serum biochemistry (including globulins and iron indices), complete blood count, fibrinogen (by heat precipitation method), and serum amyloid A. Samples were processed at the New York State Animal Health Diagnostic Center on automated analyzers ADVIA 2120i (Siemens Healthcare Diagnostics Inc., Tarrytown, NJ, USA) for hematology and Cobas C501 (Roche Diagnostics, Indianapolis, IN, USA) for biochemistry. Subjects 6 and 7 had mildly elevated fibrinogen (400 mg/dL, reference interval < 200 mg/dL) with all other parameters within normal limits, including serum amyloid A < 5 μg/ml (reference interval 0 – 8 μg/ml). Horses were maintained in stalls with partial days spent in pasture (n = 4) or on pasture alone (n=3), and had free access to grass or grass hay. All horses received annual core vaccinations (Eastern and Western Equine Encephalitis, West Nile Virus, Tetanus and Rabies) and at least biannual deworming (products varied). Blood samples were obtained in the morning (8 – 9 am), at least 16 hours after the last grain meal. Subject 1 was sampled in August, Subject 3 in September, and the remaining subjects in November, all 2018.

Approximately 50 mL of blood was collected from each horse by standard jugular venipuncture. Immediately following collection, PBMC were isolated by Ficoll gradient centrifugation, as previously described (15). Residual erythrocytes were removed by ammonium chloride lysis.

All studies were conducted under approval of Cornell University Institutional Animal Care and Use Committee.

### Single cell RNA-Seq

Within one hour of isolation, fresh PBMC were processed for single cell RNA-Seq on the 10X Genomics Chromium platform (10X Genomics). PBMC collection and scRNA-Seq were performed in three independent batches (Batch 1: Subject 1, Batch 2: Subject 3, Batch 3: Subjects 2,4,5,6,7). For each PBMC sample, 9000 cells were loaded to a single lane on the 10X Genomics Chromium instrument. scRNA-Seq libraries were prepared with the 10X Genomics Chromium Single Cell 3’ Reagent Kit (v2), according to manufacturer’s instructions. Libraries were pooled and sequenced on the Illumina NextSeq 500 in paired-end configuration (Read 1, cell barcode: 26 nt; Read 2, transcript: 98 nt) to a target read depth of approximately 35,000 paired-end reads per cell.

### scRNA-Seq Data Processing

scRNA-Seq data will be made available in the GEO repository, *accession number pending. Reference genome and transcript annotations*

The EquCab3.0 reference genome (77) was used in all analyses. Reference transcript annotations (Ensembl v95) were supplemented by manual annotation of the immunoglobulin heavy chain and light chain loci as described by Wagner, et al (Supplemental Dataset S6, (32)).

#### Read mapping and quantification

Reads were assigned to cell barcodes, mapped and quantified per gene using the CellRanger workflow (v 3.0.1, 10X Genomics) with default parameters (“standard workflow”). In our optimized workflow, BAM files generated by CellRanger were reformatted (appending cellular barcode and UMI sequence to alignment read names) and were input to the End Sequence Analysis Toolkit (ESAT, (20)). Briefly, ESAT evaluates reads mapped immediately downstream of annotated genes for potential quantification with the adjacent gene, an approach particularly relevant to 3’ scRNA-Seq data with reference transcriptomes with incomplete 3’UTR annotations. To eliminate ambiguous read assignments due to “overlapping genes” (i.e. exons from two different genes on + and – strands sharing the same genomic coordinates), the immunoglobulin-supplemented reference transcriptome (Ensembl v95) was additionally modified to remove overlapping exon intervals on opposite strands. Reformatted CellRanger BAM files were processed through ESAT in two rounds. First, ESAT was run (2500 nt extension window) with the modified transcriptome reference and set to ignore any duplicated genes. Next, to recover quantification of genes duplicated in the Ensembl v95 reference (n = 185 duplicated genes), ESAT was run (2500 nt extension window) a second time with a filtered reference containing only duplicated genes; resulting read counts were divided across gene duplications and appended to the initial gene x cell count matrix.

#### Doublet removal

Putative “doublet” cell barcodes were identified and removed from downstream analyses with the DoubletDetection tool (78).

### scRNA-Seq Data Analysis - Equine PBMC

Gene-cell count matrices processed in the above workflow were analyzed in Seurat (v3.0.0, (74, 75)) as follows.

#### Filtering, normalization, and data integration

Data were filtered to exclude genes detected in less than 3 cells (per subject), to exclude cells with less than 750 UMIs, and to exclude cells with greater than 5% UMIs assigned to mitochondrial genes (e.g. dead or dying cells). Gene-cell count matrices were independently normalized with SCTransform (79), and the top 5000 most variable genes (variance-stabilizing transformation) were selected for each subject. To minimize subject- and/or batch specific effects, datasets from all subjects were integrated on the first 40 canonical correlation components identified on the union of highly variable genes identified per subject. Immunoglobulin heavy chain and light chain genes were excluded from integration and clustering analysis.

#### Unsupervised graph based clustering

Dimensionality reduction of the integrated dataset was performed by principal component analysis (PCA). Unsupervised graph-based clustering (smart local moving algorithm (80), resolution 1.2) was performed on the first 25 principal components (selected by Scree plot visualization). Data annotated with corresponding clusters were visualized by Uniform manifold approximation and projection (UMAP; n.dims: 25, n.neighbors: 75, cosine metric, min.dist: 0.6)(81).

#### Differential gene expression analysis

Differential gene expression analyses were conducted using *edgeR* v3.26.8 (82, 83), with additional modifications for scRNA-Seq data (84). Gene expression linear models included factors for cellular gene detection rate, subject, and cluster (as identified in Seurat analysis above). Specific contrasts are detailed in relevant Results sections and/or figures. For analyses other than comparisons among CD3^+^PRF1^+^ and CD3^+^PRF1^-^ cell clusters, differential gene expression was defined as adjusted p-value < 0.05 (Benjamini-Hochberg correction) and moderated log2 fold-change > 1 (as determined in edgeR model). Differential gene expression for CD3^+^PRF1^+^ and CD3^+^PRF1^-^ cell comparisons used a less stringent fold-change cutoff (moderated log2 foldchange > 0.58) to account for reduced dynamic range of gene expression observed in these clusters. For all analyses, genes expressed (i.e. greater than or equal to 1 UMI) in less than 25% of cells for at least one group within a contrast were excluded from differential expression results. Resulting differential gene expression lists were further annotated for putative surface protein expression by intersecting one-to-one gene orthologs with the Human Surface Protein Atlas (85).

### scRNA-Seq Data Analysis - Human PBMC

Human PBMC scRNA-Seq datasets (pbmc_10k_v3; pbmc_10k_protein_v3) were obtained from 10X Genomics (https://support.10xgenomics.com/single-cell-gene-expression/datasets). Sample pbmc_10k_v3 included gene expression data from 7,255 human PBMC processed by 10X Chromium 3’ scRNA-Seq v3 chemistry. Sample pbmc_10k_protein_v3, 10,000 cells, also processed by 10X Chromium 3’ scRNA-Seq v3 chemistry, included gene expression data and immunophenotyping feature barcoding data for the following cell surface markers: CD3, CD4, CD8a, CD14, CD15, CD16, CD56, CD19, CD25, CD45RA, CD45RO, PD-1, TIGIT, CD127, IgG2a isotype control, IgG1isotype control, IgG2b isotype control.Gene-cell count matrices (and corresponding antibody count-cell matrix for sample pbmc_10k_protein_v3 were analyzed in Seurat v3.0.0 (74, 75).

Human PBMC scRNA-Seq data were filtered using the same workflow and parameters as above for equine PBMC. Data were normalized by SCTransform (79). The two human PBMC datasets were integrated on the first 30 components identified by CCA. Clustering (smart local moving algorithm (80), resolution 1.2) was performed on the first 35 principal components (selected by Scree plot visualization), and results were visualized by UMAP. Resulting clusters were annotated based on surface marker antibody labeling from sample pbmc_10k_protein_v3, as described in the text and associated figure legends.

### Horse-human PBMC scRNA-Seq cross-species correlation analysis

Cross-species scRNA-Seq correlation analyses were conducted using an approach based on Zilionis et al. (86). Human and horse gene-cell count matrices were filtered to keep only those genes with high confidence 1-to-1 orthologues across species (as defined by Ensembl v95). For each species and each major cell group (monocyte/dendritic cells, B cells, CD3^+^PRF1 ^+^ lymphocytes, CD3^+^PRF1^-^ lymphocytes), following normalization with SCTransform (79) genes were ranked by pearson residual, and genes above the 1.5*inflection point were selected as highly variable genes. Lists of highly variable genes in human and horse datasets were intersected, and the resulting list of orthologs present in both species was used for clustering analysis. Clustering was performed on natural log normalized gene x cell count matrices and clustered on Pearson correlation distance by Ward’s method (87). Results were visualized by dendrogram with the *dend* function in *R*.

### Immunophenotyping of equine PBMC by flow cytometry

The flow cytometric phenotyping protocol was adapted from (15). A list of primary antibodies is included in Table S1. Unconjugated primary antibodies CD23 and IgM were conjugated with Mix-n-Stain fluorescent protein tandem dyes antibody labeling kit for APC-CF750T (Biotium, Fremont, CA, USA) and Mix-n-Stain cf dye antibody labeling kit for CF405M (Biotium, Fremont, CA, USA), respectively, according to manufacturer instructions. All wash steps were 2ml PBS and all labeling was performed at 4°C for live cells and room temperature for fixed cells. Panel M included antibodies against CD3-AF647, CD14-biotin, CD16-unconjugated, PanIg-PE. Cells were blocked with 2% fetal bovine serum for 15 min and incubated with anti-CD16 for 30 min. Cells were washed, blocked with 10% goat serum for 15 minutes, and incubated with secondary antibody for 30 min. Cells were washed, incubated with the remaining monoclonal antibodies to surface antigens for 30 min and washed. Streptavidin-pacific orange was applied for 30 min to label CD14-biotin. Cells were washed and resuspended in PBS with 7AAD viability stain.

Panel T included antibodies against CD3-AF647, CD14-biotin, CD21-BV421, CD4-FITC, CD8-RPE, Ki67-PECy7. Cells were labeled with a fixable viability marker Live/Dead near IR for 30 min, washed, and the surface cocktail followed by streptavidin was applied as for Panel M. Cells were then fixed (eBioscience™ Intracellular fixation and permeabilization buffer set, Thermo Fisher Scientific, Waltham, MA, USA) at room temperature for 30 min, washed in permeabilization buffer, incubated with antibody for the intracellular marker Ki67 for 30min, washed and resuspended in PBS.

Panel B1 included antibodies against PanIg-PE, CD3-AF647, CD14-AF647, Tbet-PECy7, CD21-BV421, CD23-APC-CF750, and CD11b-PerCP-Vio700. Panel B2 included antibodies against PanIg-PE, CD3-AF647, CD14-AF647, Tbet-PECy7, IgM-CF405M, and IgG1-AF488. Cells were labeled with fixable viability marker Live/Dead aqua for 30 min, washed, and the surface cocktail was applied. Cells were then fixed (TrueNuclear™ TF fixation and permeabilization buffer set, BioLegend, San Diego, CA, USA) at room temperature for 60 min, washed in permeabilization buffer, incubated with the intranuclear marker Tbet for 30min, washed and resuspended in PBS.

Fluorescence was measured on a Gallios flow cytometer (Beckman Coulter, Indianapolis, IN, USA) with a minimum 100,000 events collected per sample. Analysis was performed with FlowJo version 10.6.1 (FlowJo LLC, Ashland, OR, USA). Single color controls were used to set the compensation matrix. Gating strategies are shown in Fig. S5. The researcher performing gating analyses (J.E.T) was blinded to scRNA-Seq results. All flow cytometry data is available on Flow Repository, accession number FR-FCM-Z2JN.

## ACKNOWLEDGMENTS

This study was supported by grants from the Agriculture and Food Research initiative Competitive Grant no. 2016-67015-24765 from the USDA National institute of Food and Agriculture and the Jack Lowe Equine Health Funds/Mollie Wilmot Equine Research Fund J.E.T. was supported by the National Institute of Allergy and Infectious Diseases of the National Institutes of Health under Award number K08AI141767. Additional support for B.R.R. and R.S.P from the Department of Microbiology, Icahn School of Medicine at Mount Sinai. We thank Peter Schweitzer and the Cornell Biotechnology Resource Center for scRNA-Seq support. We also thank Charles M. Rice, Troels K.H. Scheel, and Amit Kapoor for helpful discussion and advice. The content is solely the responsibility of the authors and does not necessarily represent the official views of the funders, including the National Institutes of Health.

## REFERENCES

1. J. J. Russell, et al., Non-model model organisms. BMC Biol 15, 55, s12915-017-0391-5 (2017).

2. D. Masopust, C. P. Sivula, S. C. Jameson, Of Mice, Dirty Mice, and Men: Using Mice To Understand Human Immunology. J.I. 199, 383–388 (2017).

3. J. R. Swearengen, Choosing the right animal model for infectious disease research. Animal Model Exp Med 1, 100–108 (2018).

4. S. Ryu, B. I. Kim, J.-S. Lim, C. S. Tan, B. C. Chun, One Health Perspectives on Emerging Public Health Threats. J Prev Med Public Health 50, 411–414 (2017).

5. OneHealth: OIE -World Organisation for Animal Health (March 27, 2020).

6. One Health | CDC (2020) (March 27, 2020).

7. A. Adan, G. Alizada, Y. Kiraz, Y. Baran, A. Nalbant, Flow cytometry: basic principles and applications. Critical Reviews in Biotechnology 37, 163–176 (2017).

8. H. T. Maecker, J. P. McCoy, R. Nussenblatt, Standardizing immunophenotyping for the Human Immunology Project. Nature reviews. Immunology 12, 191–200 (2012).

9. M. H. Spitzer, G. P. Nolan, Mass Cytometry: Single Cells, Many Features. Cell 165, 780–791 (2016).

10. E. Z. Macosko, et al., Highly Parallel Genome-wide Expression Profiling of Individual Cells Using Nanoliter Droplets. Cell 161, 1202–1214 (2015).

11. A. M. Klein, et al., Droplet Barcoding for Single-Cell Transcriptomics Applied to Embryonic Stem Cells. Cell 161, 1187–1201 (2015).

12. G. X. Y. Zheng, et al., Massively parallel digital transcriptional profiling of single cells. Nature communications 8, 14049 (2017).

13. M. J. T. Stubbington, O. Rozenblatt-Rosen, A. Regev, S. A. Teichmann, Single-cell transcriptomics to explore the immune system in health and disease. Science (New York, N.Y.) 358, 58–63 (2017).

14. S. K. Khurana, Zoonotic Pathogens Transmitted from Equines: Diagnosis and Control. Adv. Anim. Vet. Sci. 3, 32–53 (2015).

15. J. E. Tomlinson, B. Wagner, M. J. B. Felippe, G. R. Van de Walle, Multispectral fluorescence-activated cell sorting of B and T cell subpopulations from equine peripheral blood. Veterinary Immunology and Immunopathology 199, 22–31 (2018).

16. J. W. Austin, et al., Overexpression of T-bet in HIV infection is associated with accumulation of B cells outside germinal centers and poor affinity maturation. Science Translational Medicine 11 (2019).

17. J. J. Knox, A. Myles, M. P. Cancro, T-bet+ memory B cells: Generation, function, and fate. Immunol. Rev. 288, 149–160 (2019).

18. X. Zhang, et al., Comparative Analysis of Droplet-Based Ultra-High-Throughput Single-Cell RNA-Seq Systems. Molecular Cell 73, 130–142.e5 (2019).

19. G. Grillo, et al., UTRdb and UTRsite (RELEASE 2010): a collection of sequences and regulatory motifs of the untranslated regions of eukaryotic mRNAs. Nucleic Acids Research 38, D75–D80 (2010).

20. A. Derr, et al., End Sequence Analysis Toolkit (ESAT) expands the extractable information from single-cell RNA-seq data. Genome Res. 26, 1397–1410 (2016).

21. E. Kabithe, J. Hillegas, T. Stokol, J. Moore, B. Wagner, Monoclonal antibodies to equine CD14. Veterinary Immunology and Immunopathology 138, 149–153 (2010).

22. C. V. Jakubzick, G. J. Randolph, P. M. Henson, Monocyte differentiation and antigen-presenting functions. Nat Rev Immunol 17, 349–362 (2017).

23. R. N. Hanna, et al., The transcription factor NR4A1 (Nur77) controls bone marrow differentiation and the survival of Ly6C-monocytes. Nat Immunol 12, 778–785 (2011).

24. F. Geissmann, S. Jung, D. R. Littman, Blood Monocytes Consist of Two Principal Subsets with Distinct Migratory Properties. Immunity 19, 71–82 (2003).

25. J. Cros, et al., Human CD14dim monocytes patrol and sense nucleic acids and viruses via TLR7 and TLR8 receptors. Immunity 33, 375–386 (2010).

26. J. Gamrekelashvili, et al., Regulation of monocyte cell fate by blood vessels mediated by Notch signalling. Nature Communications 7, 12597 (2016).

27. M. Collin, V. Bigley, Human dendritic cell subsets: an update. Immunology 154, 3–20 (2018).

28. I. Caminschi, et al., The dendritic cell subtype-restricted C-type lectin Clec9A is a target for vaccine enhancement. Blood 112, 3264–3273 (2008).

29. K. Hildner, et al., Batf3 Deficiency Reveals a Critical Role for CD8 + Dendritic Cells in Cytotoxic T Cell Immunity. Science 322, 1097–1100 (2008).

30. A. Izaguirre, et al., Comparative analysis of IRF and IFN-alpha expression in human plasmacytoid and monocyte-derived dendritic cells. Journal of Leukocyte Biology 74, 1125–1138 (2003).

31. B. Cisse, et al., Transcription Factor E2-2 Is an Essential and Specific Regulator of Plasmacytoid Dendritic Cell Development. Cell 135, 37–48 (2008).

32. B. Wagner, Immunoglobulins and immunoglobulin genes of the horse. Developmental & Comparative Immunology 30, 155–164 (2006).

33. Y. Wang, D. Bhattacharya, Adjuvant-specific regulation of long-term antibody responses by ZBTB20. The Journal of Experimental Medicine 211,841–856 (2014).

34. K. Rubtsova, A. V. Rubtsov, L. F. van Dyk, J. W. Kappler, P. Marrack, T-box transcription factor T-bet, a key player in a unique type of B-cell activation essential for effective viral clearance. Proc. Natl. Acad. Sci. U.S.A. 110, E3216–3224 (2013).

35. G. R. A. Ehrhardt, et al., Expression of the immunoregulatory molecule FcRH4 defines a distinctive tissue-based population of memory B cells. Journal of Experimental Medicine 202, 783–791 (2005).

36. A. V. Rubtsov, et al., Toll-like receptor 7 (TLR7)-driven accumulation of a novel CD11c+ B-cell population is important for the development of autoimmunity. Blood 118, 1305–1315 (2011).

37. K. Rubtsova, A. V. Rubtsov, M. P. Cancro, P. Marrack, Age-Associated B Cells: A T-bet-Dependent Effector with Roles in Protective and Pathogenic Immunity. J. Immunol. 195, 1933–1937 (2015).

38. Y. Liu, et al., Involvement of the HCK and FGR src-Family Kinases in FCRL4-Mediated Immune Regulation. J.I. 194, 5851–5860 (2015).

39. J. L. Karnell, et al., Role of CD11c+ T-bet+ B cells in human health and disease. Cell. Immunol. 321, 40–45 (2017).

40. J. J. Knox, et al., T-bet+ B cells are induced by human viral infections and dominate the HIV gp140 response. JCI Insight 2 (2017).

41. C. Stoeckle, et al., Cathepsin W expressed exclusively in CD8+ T cells and NK cells, is secreted during target cell killing but is not essential for cytotoxicity in human CTLs. Experimental Hematology 37, 266–275 (2009).

42. J. F. Hedges, J. C. Graff, M. A. Jutila, Transcriptional Profiling of γδ T Cells. J Immunol 171, 4959–4964 (2003).

43. The Immunological Genome Project Consortium, et al., Molecular definition of the identity and activation of natural killer cells. Nature Immunology 13, 1000–1009 (2012).

44. B. J. Schmiedel, et al., Impact of Genetic Polymorphisms on Human Immune Cell Gene Expression. Cell 175, 1701–1715.e16 (2018).

45. Y. Zhao, et al., Single-cell transcriptomic landscape of nucleated cells in umbilical cord blood. GigaScience 8 (2019).

46. O. Franzén, L.-M. Gan, J. L. M. Björkegren, PanglaoDB: a web server for exploration of mouse and human single-cell RNA sequencing data. Database 2019 (2019).

47. M. C. van Aalderen, et al., Label-free Analysis of CD8+ T Cell Subset Proteomes Supports a Progressive Differentiation Model of Human-Virus-Specific T Cells. Cell Reports 19, 1068–1079 (2017).

48. M. Post, et al., The Transcription Factor ZNF683/HOBIT Regulates Human NK-Cell Development. Frontiers in Immunology 8 (2017).

49. R. L. Tallmadge, M. Wang, Q. Sun, M. J. B. Felippe, Transcriptome analysis of immune genes in peripheral blood mononuclear cells of young foals and adult horses. PLoS ONE 13, e0202646 (2018).

50. F. A. Vieira Braga, et al., Blimp-1 homolog Hobit identifies effector-type lymphocytes in humans. European Journal of Immunology 45, 2945–2958 (2015).

51. L. L. Lanier, Up on the tightrope: natural killer cell activation and inhibition. Nature Immunology 9, 495–502 (2008).

52. F. Naeim, Atlas of hematopathology: morphology, immunophenotype, cytogenetics, and molecular approaches, 2nd edition (Elsevier, 2018).

53. L. E. Noronha, R. M. Harman, B. Wagner, D. F. Antczak, Generation and characterization of monoclonal antibodies to equine CD16. Veterinary Immunology and Immunopathology 146, 135–142 (2012).

54. L. E. Noronha, R. M. Harman, B. Wagner, D. F. Antczak, Generation and characterization of monoclonal antibodies to equine NKp46. Veterinary Immunology and Immunopathology 147, 60–68 (2012).

55. D. P. Lunn, J. T. McClure, C. S. Schobert, M. A. Holmes, Abnormal patterns of equine leucocyte differentiation antigen expression in severe combined immunodeficiency foals suggests the phenotype of normal equine natural killer cells. Immunology 84, 495–499 (1995).

56. L. Ziegler-Heitbrock, Monocyte subsets in man and other species. Cellular Immunology 289, 135–139 (2014).

57. A. Mildner, S. Jung, Development and Function of Dendritic Cell Subsets. Immunity 40, 642–656 (2014).

58. D. J. Cavatorta, H. N. Erb, M. J. B. F. Flaminio, Ex vivo generation of mature equine monocyte-derived dendritic cells. Veterinary Immunology and Immunopathology 131,259–267 (2009).

59. Y. Lee, M. Kiupel, G. S. Hussey, Characterization of respiratory dendritic cells from equine lung tissues. BMC Vet Res 13, 1–11 (2017).

60. S. Mauel, F. Steinbach, H. Ludwig, Monocyte-derived dendritic cells from horses differ from dendritic cells of humans and mice. Immunology 117, 463–473 (2006).

61. A. Ziegler, E. Marti, A. Summerfield, A. Baumann, Identification and characterization of equine blood plasmacytoid dendritic cells. Developmental & Comparative Immunology 65, 352–357 (2016).

62. B. E. Barnett, et al., Cutting Edge: B Cell-Intrinsic T-bet Expression Is Required To Control Chronic Viral Infection. J. Immunol. 197, 1017–1022 (2016).

63. D. Piovesan, et al., c-Myb Regulates the T-Bet-Dependent Differentiation Program in B Cells to Coordinate Antibody Responses. Cell Rep 19, 461–470 (2017).

64. S. A. Jenks, et al., Distinct Effector B Cells Induced by Unregulated Toll-like Receptor 7 Contribute to Pathogenic Responses in Systemic Lupus Erythematosus. Immunity 49, 725–739.e6 (2018).

65. S. Wang, et al., IL-21 drives expansion and plasma cell differentiation of autoreactive CD11chiT-bet+ B cells in SLE. Nat Commun 9, 1758 (2018).

66. N. Obeng-Adjei, et al., Malaria-induced interferon-γ drives the expansion of Tbethi atypical memory B cells. PLoS Pathog 13, e1006576 (2017).

67. S. Portugal, et al., Malaria-associated atypical memory B cells exhibit markedly reduced B cell receptor signaling and effector function. eLife 4, e07218 (2015).

68. A. R. Burton, et al., Circulating and intrahepatic antiviral B cells are defective in hepatitis B. J. Clin. Invest. 128, 4588–4603 (2018).

69. T. J. Divers, et al., *Borrelia burgdorferi* Infection and Lyme Disease in North American Horses: A Consensus Statement: Lyme Disease in Horses. J Vet Intern Med 32, 617–632 (2018).

70. S. M. Reed, et al., Equine Protozoal Myeloencephalitis: An Updated Consensus Statement with a Focus on Parasite Biology, Diagnosis, Treatment, and Prevention. J Vet Intern Med 30, 491–502 (2016).

71. A. Raza, A. G. Qamar, K. Hayat, S. Ashraf, A. R. Williams, Anthelmintic resistance and novel control options in equine gastrointestinal nematodes. Parasitology 146, 425–437 (2019).

72. D. Aran, et al., Reference-based analysis of lung single-cell sequencing reveals a transitional profibrotic macrophage. Nature Immunology 20, 163 (2019).

73. Y. Tan, P. Cahan, SingleCellNet: A Computational Tool to Classify Single Cell RNA-Seq Data Across Platforms and Across Species. Cell Systems 9, 207–213.e2 (2019).

74. A. Butler, P. Hoffman, P. Smibert, E. Papalexi, R. Satija, Integrating single-cell transcriptomic data across different conditions, technologies, and species. Nature Biotechnology 36, 411–420 (2018).

75. T. Stuart, et al., Comprehensive Integration of Single-Cell Data. Cell 177, 1888–1902.e21 (2019).

76. J. D. Welch, et al., Single-Cell Multi-omic Integration Compares and Contrasts Features of Brain Cell Identity. Cell 177, 1873–1887.e17 (2019).

77. T. S. Kalbfleisch, et al., Improved reference genome for the domestic horse increases assembly contiguity and composition. Commun Biol 1, 197 (2018).

78. Adam Gayoso, Jonathan Shor, GitHub: DoubletDetection (Zenodo, 2019) https:/doi.org/10.5281/zenodo.2678042 (March 27, 2020).

79. C. Hafemeister, R. Satija, Normalization and variance stabilization of single-cell RNA-seq data using regularized negative binomial regression. Genome Biology 20 (2019).

80. L. Waltman, N. J. van Eck, A smart local moving algorithm for large-scale modularity-based community detection. Eur. Phys. J. B 86, 471 (2013).

81. L. McInnes, J. Healy, N. Saul, L. Großberger, UMAP: Uniform Manifold Approximation and Projection. JOSS 3, 861 (2018).

82. D. J. McCarthy, Y. Chen, G. K. Smyth, Differential expression analysis of multifactor RNA-Seq experiments with respect to biological variation. Nucleic Acids Res. 40, 4288–4297 (2012).

83. M. D. Robinson, D. J. McCarthy, G. K. Smyth, edgeR: a Bioconductor package for differential expression analysis of digital gene expression data. Bioinformatics (Oxford, England) 26, 139–140 (2010).

84. C. Soneson, M. D. Robinson, Bias, robustness and scalability in single-cell differential expression analysis. Nature Methods 15, 255–261 (2018).

85. D. Bausch-Fluck, et al., A Mass Spectrometric-Derived Cell Surface Protein Atlas. PLoS ONE 10, e0121314 (2015).

86. R. Zilionis, et al., Single-Cell Transcriptomics of Human and Mouse Lung Cancers Reveals Conserved Myeloid Populations across Individuals and Species. Immunity (2019) https:/doi.org/10.1016/j.immuni.2019.03.009 (April 24, 2019).

87. J. H. Ward, Hierarchical Grouping to Optimize an Objective Function. Journal of the American Statistical Association 58, 236–244 (1963).

