## Supplemental Tables and Figures for "Single cell resolution landscape of equine peripheral blood mononuclear cells reveals diverse immune cell subtypes including T-bet^+^ B cells"

Figures S1 to S6, Tables S1

Other supplementary materials for this manuscript include the following:

Differential gene expression Datasets S1-5, 7-8

- Dataset S1. Differentially expressed gene marker lists for major cell groups
- Dataset S2. Differentially expressed gene marker lists for monocyte and dendritic cell clusters
- Dataset S3. Differentially expressed gene marker lists for monocyte cell clusters
- Dataset S4. Differentially expressed gene marker lists for dendritic cell clusters
- Dataset S5. Differentially expressed gene marker lists for B cell clusters
- Dataset S7. Differentially expressed gene marker lists for $CD3^+$ $PRF1^+$ lymphocyte cell clusters
- Dataset S8. Differentially expressed gene marker lists for $CD3^+$ $PRF1^-$ lymphocyte cell clusters

Dataset S6. Genome annotation file specifying custom immunoglobulin gene entries; adapted from Wagner et al. [34]

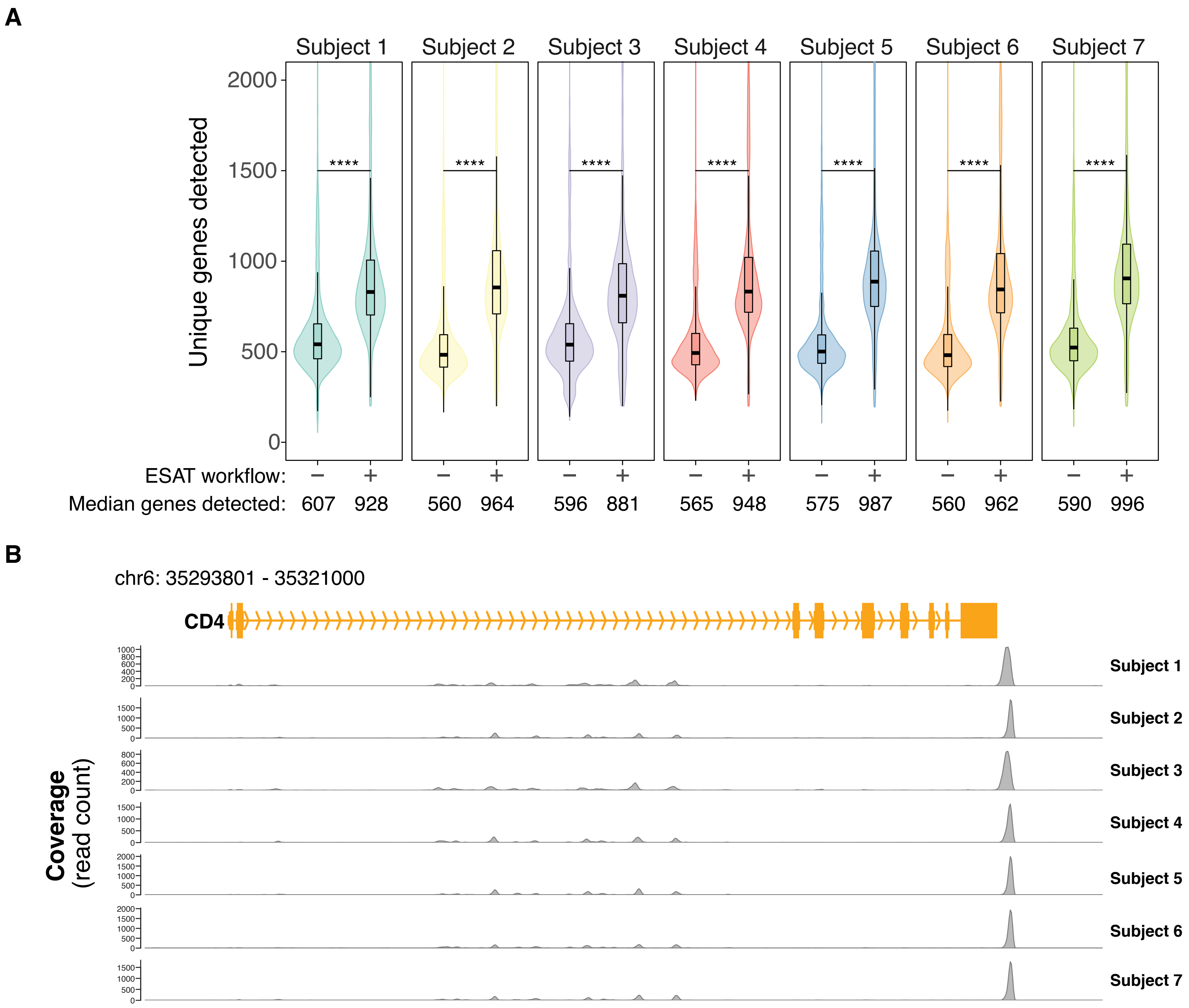


**Figure S1. Optimized scRNA-seq data processing workflow improves per cell gene detection.** (A) Violin plot of number of unique genes detected per cell across each study subject (N = 7) using standard CellRanger (10X Genomics) workflow versus modified workflow incorporating ESAT. Box plot indicate median, 25th percentile and 75th percentile. Unpaired t-tests were conducted on a per subject basis, **** $p < 1 \times 10^{-15}$. (B) scRNA-Seq read mapping pattern at the CD4 locus, known to be abundantly expressed in equine PBMC, demonstrating majority of reads map directly downstream of reference transcript annotation.

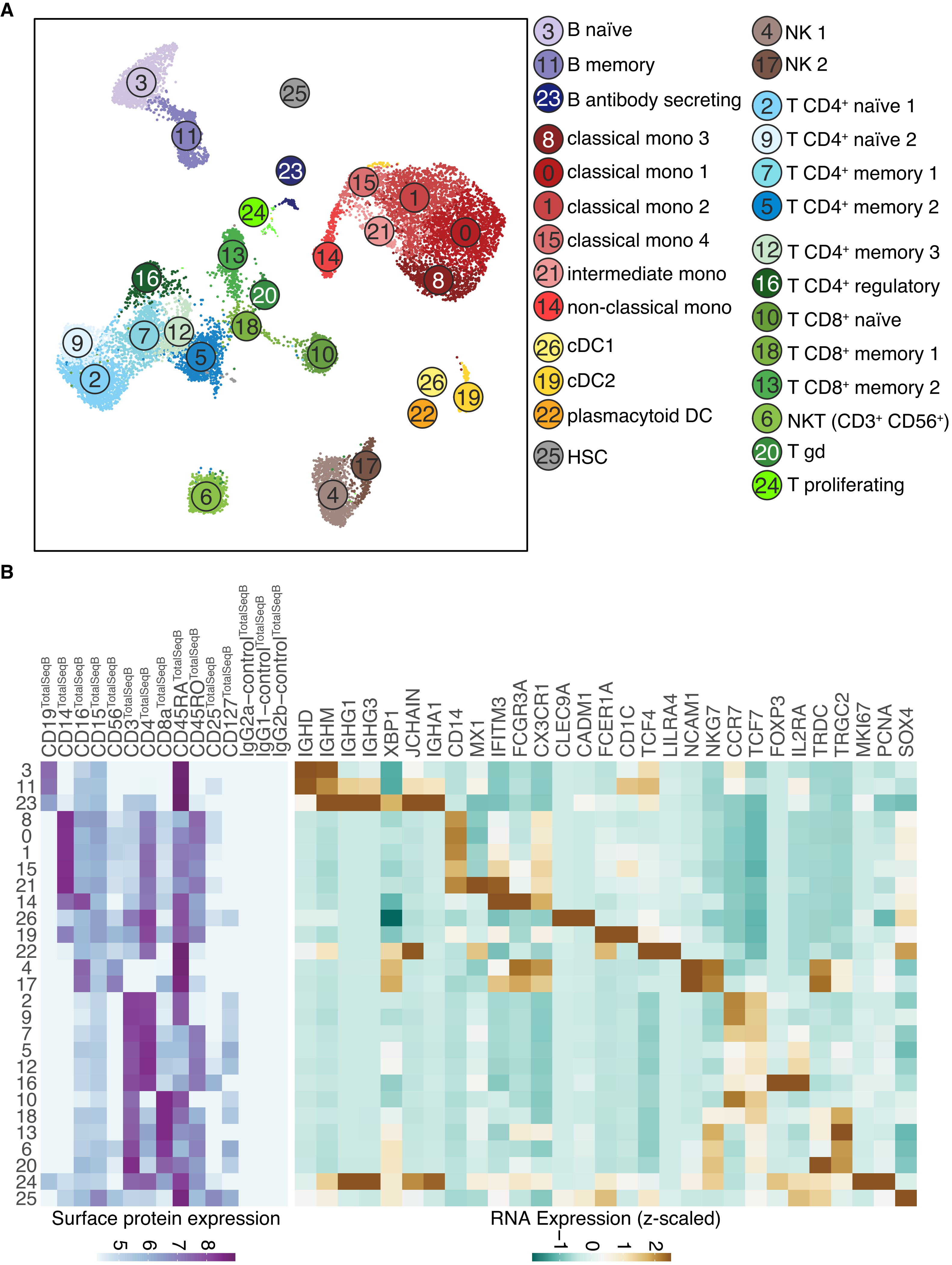


**Figure S2. Human reference scRNA-seq clustering results and annotation.** Human PBMC scRNA-seq data (surface antibody feature barcoding + gene expression) obtained from the 10X Genomics public dataset collection were integrated and analyzed by Seurat v3. Unsupervised clustering analysis of the integrated dataset identified 26 clusters. (A) UMAP representation of human PBMCs (n = 17,255 total cells from two independent datasets passing QC filters). Points are colored by cluster membership. Clusters were annotated primarily by surface marker labeling when available, supplemented by RNA expression of marker genes. (B) Heatmaps of surface antibody feature barcode data (left) and RNA expression for select marker genes (right) utilized to inform cluster annotation.

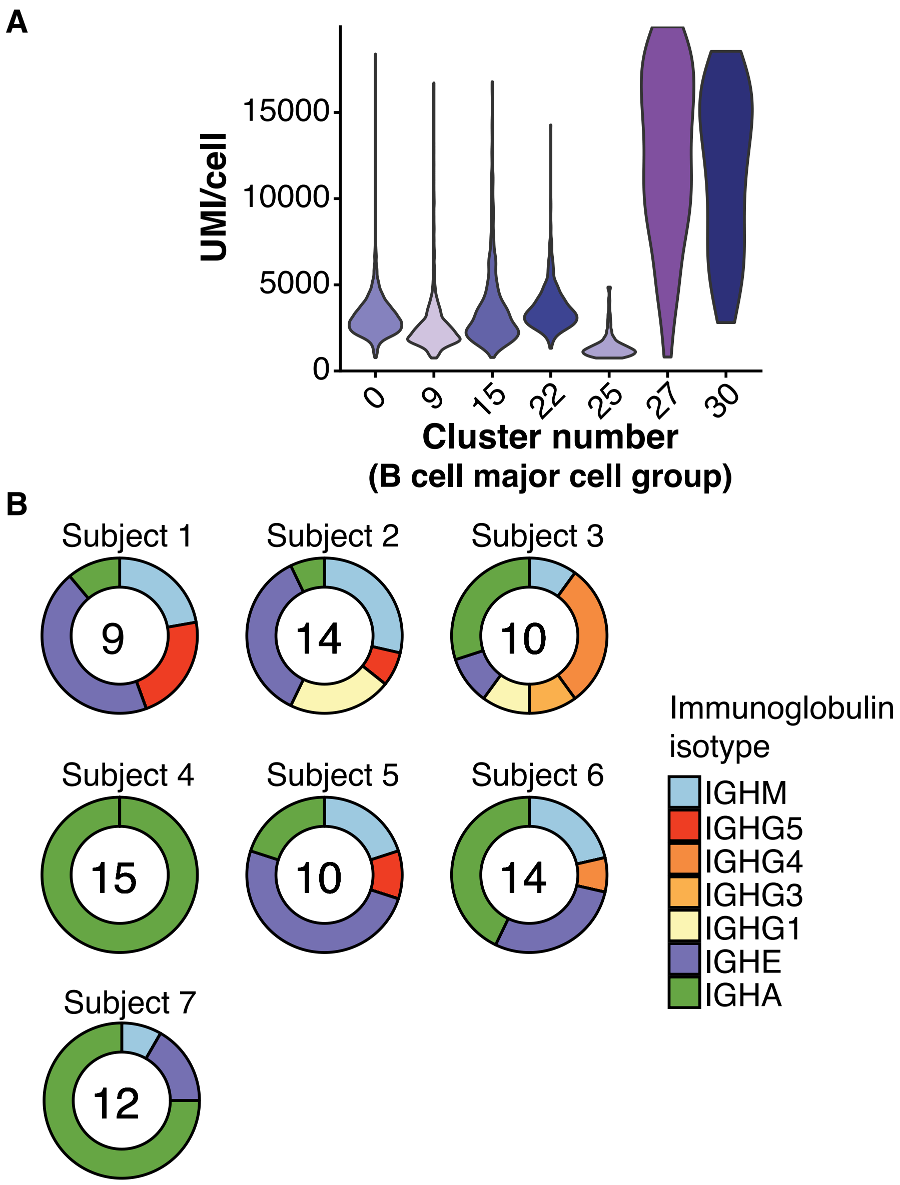


**Figure S3. B cell quality control metrics and antibody secreting cell immunoglobulin isotype usage.** (A) Violin plot of UMI (transcript) counts per cell across B cell clusters identified by unsupervised clustering. Cluster 25 was excluded from downstream analysis due to insufficient UMI counts. (B) Immunoglobulin isotype usage in antibody secreting cells (cluster 27) as determined by scRNA-Seq gene expression data. Center value indicates number of antibody secreting cells (cluster 27) detected by individual subject.

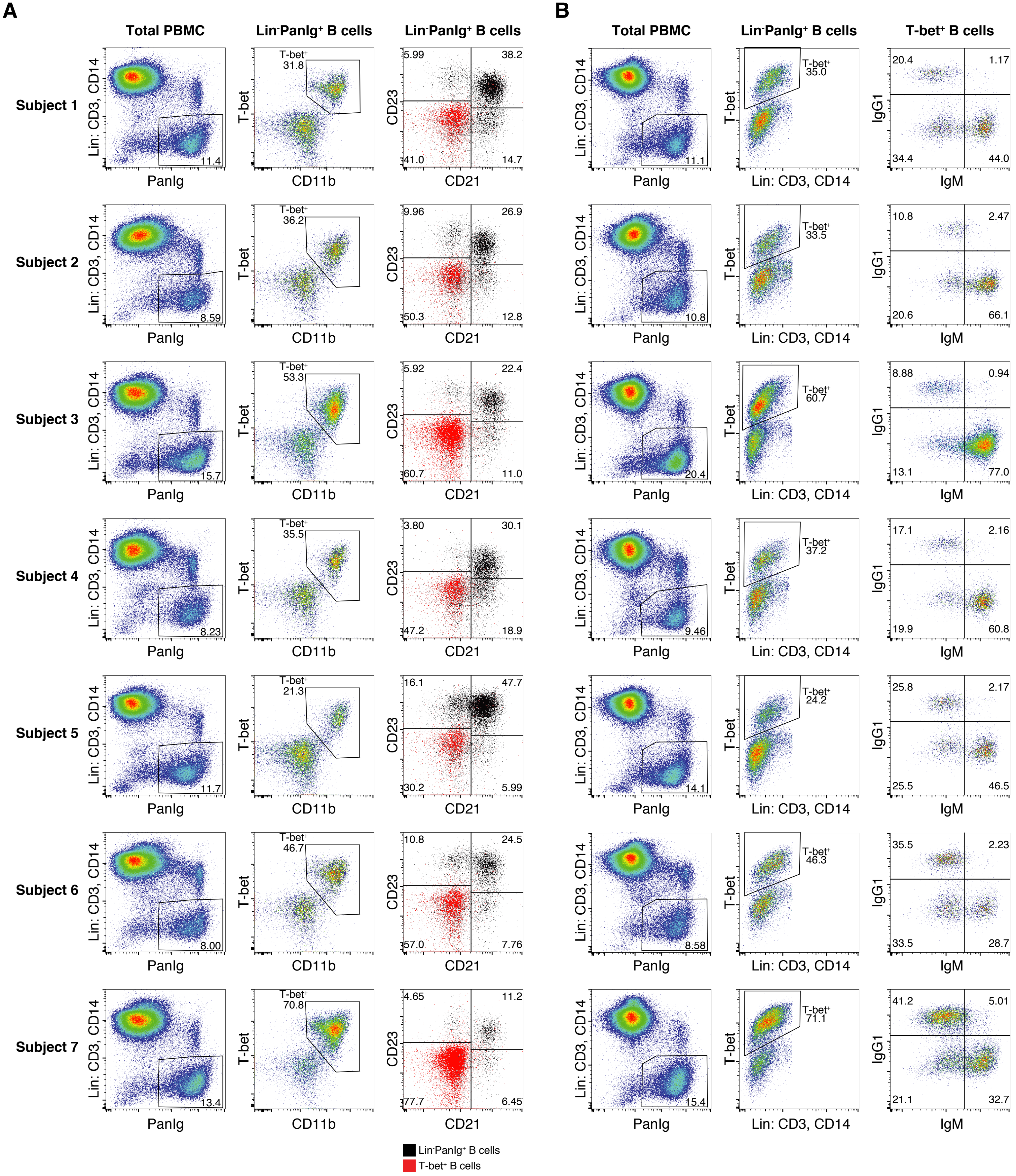


**Figure S4. T-bet$^{+}$ B cells identified by scRNA-Seq are detectable by flow cytometry in all subjects examined.** Flow cytometry gating schemes for T-bet$^{+}$ B cell characterization in equine PBMC across each study subject (N = 7). (A) T-bet$^{+}$ B cell gating for flow cytometry panel including CD11b, CD21 and CD23 labeling. Labels above plots indicate visualized gate. (B) T-bet$^{+}$ B cell gating for flow cytometry panel including IgM and IgG1 surface immunoglobulin labeling. Labels above plots indicate visualized gate.

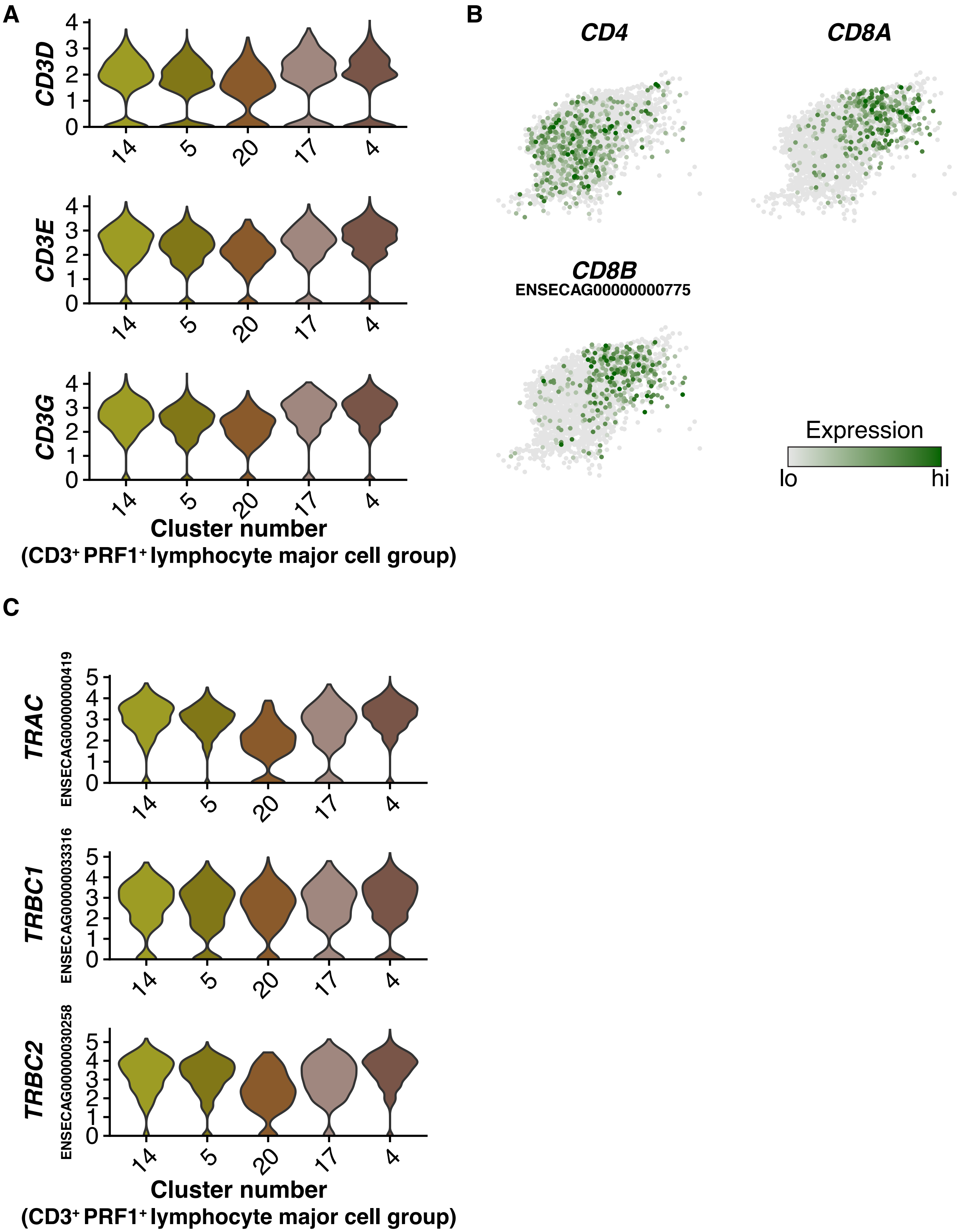


**Figure S5. Select gene expression patterns in CD3⁺ PRF1⁺ lymphocyte major cell group.** (A) Expression of select CD3 transcripts by indicated clusters in CD3⁺PRF1⁺ lymphocyte major cell group. Values plotted as log normalized counts per cell. (B) Expression patterns of *CD4*, *CD8A* and ENSECAG00000000775 (*CD8B*) in cluster 5. Expression values are scaled independently for each plot, ranging from 2.5 to 97.5 percentile of gene expression across all plotted cells. (C) Expression of T cell receptor genes by indicated clusters in CD3⁺PRF1⁺ lymphocyte major cell group. Values plotted as log normalized counts per cell.

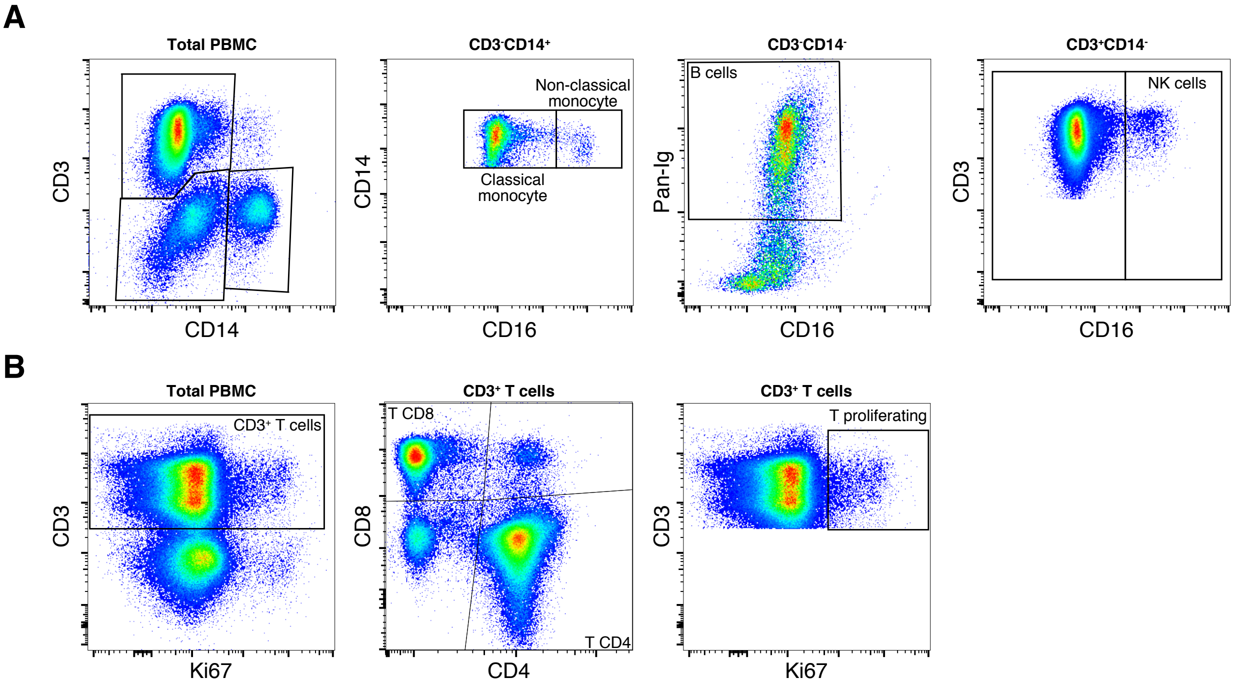


**Figure S6. Representative flow cytometry gating schemes for immunophenotyping of equine PBMC.** PBMC single cell suspensions were labeled with fluorescent-conjugated antibodies and analyzed by flow cytometry as described in *Materials and Methods*. (A) Gating scheme for immunophenotyping panel to resolve monocyte populations, B cells, and NK cells. Labels above plots indicate visualized gate. (B) Gating scheme for immunophenotyping panel to resolve $CD4^+$, $CD8^+$, and proliferating T cell populations. Labels above plots indicate visualized gate.

| Target species | Antibody | Clone | Source | Reference |
|---|---|---|---|---|
| Eq | CD3-AF647 | UC-F6G | UC Davis, Dr. Stott | Tomlinson et al., 2018, Blanchard-Channell et al. 1994; Lunn et al., 1998 |
| Eq | CD4-FITC | CVS4 | Bio Rad Antibodies | Tomlinson et al., 2018; Lunn et al., 1991 |
| Eq | CD8-RPE | CVS8 | Bio Rad Antibodies | Tomlinson et al., 2018; Lunn et al., 1991 |
| Hu | Ki67-PECy7 | B56 | BD Biosciences | This study* |
| Eq | Pan B cells-RPE | CVS36 | Bio Rad Antibodies | Tomlinson et al., 2018; Lunn et al., 1998 |
| Eq | CD16 | 1A2.D11 | Cornell University, Dr. Antczak | Noronha et al., 2012 |
| Hu | CD21-BV421 | B-ly4 | BD Biosciences | Tomlinson et al., 2018; Ibrahim and Steinbach 2012 and 2007 |
| Eq | CD14-Sav | 105 | Cornell University, Dr. Wagner | Kabithe et al., 2010 |
| Eq | CD14-AF647 | 105 | Cornell University, Dr. Wagner | Kabithe et al., 2010 |
| Eq | CD23-APC-CF750 | 51-3 | Cornell University, Dr. Wagner | Wagner et al., 2012 |
| Hu | Tbet-PECy7 | 4B10 | BioLegend | This study** |
| Hu | CD11b-PerCP-Vio700 | M1/70.15.11.5 | Miltenyi Biotec | Ibrahim and Steinbach 2012 |
| Eq | IgM-CF405M | I-22 | Cornell University, Dr. Wagner | Tomlinson et al., 2018; Wagner et al., 2008 |
| Eq | IgG1-AF488 | CVS45 | Cornell University, Dr. Wagner | Tomlinson et al., 2018; Lunn et al., 1998 |
| | Live/Dead | Fixable aqua | ThermoFisher Scientific | |
| | Live/Dead | Fixable near IR | ThermoFisher Scientific | |
| | 7AAD | | BioLegend | |

**Supplemental Table 1. Antibodies used for flow cytometry in this study for equine PBMC immuno-phenotyping.** Eq., horse, Hu., human. *Anti-human Ki67 clone B56. This clone was validated by test on non-replicating primary equine lymphocytes, which were negative, and pokeweed mitogen stimulated replicating primary equine lymphocytes, which were positive (data not shown). **Anti-human T-bet clone 4B10 was validated by co-expression patterns with CD11b, CD21, and CD23 as shown in the main manuscript.

### Table references


Blanchard-Channell, M., Moore, P.F., and Stott, J.L. (1994). Characterization of monoclonal antibodies specific for equine homologues of CD3 and CD5. Immunology *82*, 548–554.

Ibrahim, S., and Steinbach, F. (2007). Non-HLDA8 animal homologue section anti-leukocyte mAbs tested for reactivity with equine leukocytes. Veterinary Immunology and Immunopathology *119*, 81–91.

Ibrahim, S., and Steinbach, F. (2012). Immunoprecipitation of equine CD molecules using anti-human MABs previously analyzed by flow cytometry and immunohistochemistry. Veterinary Immunology and Immunopathology *145*, 7–13.

Lunn, D.P., Holmes, M.A., and Duffus, W.P. (1991). Three monoclonal antibodies identifying antigens on all equine T lymphocytes, and two mutually exclusive T-lymphocyte subsets. Immunology *74*, 251–257.

Lunn, D.P., Holmes, M.A., Antczak, D.F., Agerwal, N., Baker, J., Bendali-Ahcene, S., Blanchard-Channell, M., Byrne, K.M., Cannizzo, K., Davis, W., et al. (1998). Report of the Second Equine Leucocyte Antigen Workshop, Squaw Valley, California, July 1995. Veterinary Immunology and Immunopathology *62*, 101–143.

Wagner, B., Hillegas, J.M., and Babasyan, S. (2012). Monoclonal antibodies to equine CD23 identify the low-affinity receptor for IgE on subpopulations of IgM+ and IgG1+ B-cells in horses. Veterinary Immunology and Immunopathology *146*, 125–134.
