## Supplementary material for "Single cell resolution landscape of equine peripheral blood mononuclear cells reveals diverse immune cell subtypes including T-bet^+^ B cells": Dataset S1

| genes | logFC | logCPM | F | PValue | FDR | percent.exp | Surfacome.Label |
|---|---|---|---|---|---|---|---|
| ENSECAG00000031094 | 5.6047 | 10.892 | 20366 | 0 | 0 | 0.6222257 | n.a. |
| ENSECAG00000031522 | 4.8883 | 10.189 | 10540 | 0 | 0 | 0.2927208 | n.a. |
| CD79A | 4.5604 | 9.7872 | 29197 | 0 | 0 | 0.934813 | surface |
| ENSECAG00000039599 | 4.4224 | 10.091 | 9111 | 0 | 0 | 0.2564023 | n.a. |
| MS4A1 | 4.4071 | 9.7256 | 25078 | 0 | 0 | 0.8907341 | surface |
| ENSECAG00000027826 | 4.3862 | 9.7443 | 20064 | 0 | 0 | 0.8957008 | n.a. |
| Eqca-DOB1 | 3.0937 | 9.4745 | 5418 | 0 | 0 | 0.6406953 | n.a. |
| ENSECAG00000014585 | 3.0711 | 10.271 | 2687 | 0 | 0 | 0.7051063 | n.a. |
| SH3BP5 | 2.7624 | 9.4429 | 4995 | 0 | 0 | 0.5997206 | n.a. |
| TNFRSF13C | 2.7326 | 9.349 | 10061 | 0 | 0 | 0.4564644 | nonsurface |
| BANK1 | 2.622 | 9.413 | 5571 | 0 | 0 | 0.5132702 | n.a. |
| POLD1 | 2.6215 | 9.3555 | 9434 | 0 | 0 | 0.39702 | n.a. |
| ENSECAG00000039021 | 2.5291 | 9.3181 | 13725 | 0 | 0 | 0.384293 | n.a. |
| POU2F2 | 2.5014 | 9.5681 | 3525 | 0 | 0 | 0.6571473 | n.a. |
| IFI30 | 2.4672 | 9.9429 | 4331 | 0 | 0 | 0.823219 | n.a. |
| IGHM | 2.4662 | 9.6407 | 2391 | 0 | 0 | 0.5741114 | n.a. |
| TCF4 | 2.3437 | 9.3454 | 7278 | 0 | 0 | 0.3943815 | n.a. |
| PYGM | 2.3364 | 9.3286 | 6592 | 0 | 0 | 0.3681515 | n.a. |
| DRA | 2.2672 | 11.742 | 6480 | 0 | 0 | 0.994723 | n.a. |
| CYB561A3 | 2.2669 | 9.3685 | 4884 | 0 | 0 | 0.4154897 | nonsurface |
| DQA | 1.9781 | 10.712 | 2812 | 0 | 0 | 0.9658544 | n.a. |
| CD74 | 1.9232 | 12.628 | 7900 | 0 | 0 | 0.9993792 | surface |
| ENSECAG00000022442 | 1.759 | 9.7128 | 1649 | 0 | 0 | 0.6978116 | n.a. |
| DRB | 1.5615 | 11.205 | 2110 | 0 | 0 | 0.987273 | n.a. |
| PTPRCAP | 1.3585 | 10.617 | 1587 | 0 | 0 | 0.9279839 | nonsurface |
| MEF2C | 2.242 | 9.4205 | 5440 | 0 | 0 | 0.410523 | n.a. |
| DQB | 1.7581 | 10.011 | 1440 | 0 | 2E-306 | 0.7504268 | n.a. |
| ALDH2 | 1.9579 | 9.5616 | 3898 | 7E-304 | 1E-301 | 0.5024057 | n.a. |
| IGHG3 | 3.5704 | 9.6047 | 5024 | 4E-303 | 9E-301 | 0.6793419 | n.a. |
| AIM2 | 1.9611 | 9.3585 | 5567 | 3E-284 | 6E-282 | 0.3434735 | n.a. |
| FCRLA | 2.1937 | 9.3304 | 7079 | 3E-279 | 7E-277 | 0.340059 | n.a. |
| CYBA | 1.0532 | 10.509 | 1277 | 1E-274 | 2E-272 | 0.901909 | nonsurface |
| ANKRD2 | 2.0258 | 9.3125 | 8870 | 2E-272 | 3E-270 | 0.2879094 | n.a. |
| ENSECAG00000031569 | 1.1598 | 11.194 | 1246 | 4E-268 | 7E-266 | 0.9382275 | n.a. |
| HVCN1 | 2.0592 | 9.3352 | 5048 | 2E-262 | 4E-260 | 0.3534068 | nonsurface |
| Eqca-DQB1 | 1.3488 | 10.925 | 1190 | 2E-256 | 3E-254 | 0.9714419 | n.a. |
| BHLHE41 | 2.0766 | 9.3067 | 8545 | 4E-252 | 7E-250 | 0.3104144 | n.a. |
| HSF2BP | 2.6688 | 9.3751 | 5312 | 2E-251 | 3E-249 | 0.4698122 | n.a. |
| ENSECAG00000021796 | 1.8738 | 9.6911 | 1600 | 2E-248 | 3E-246 | 0.6754617 | n.a. |
| SPI1 | 1.1699 | 9.6618 | 3311 | 3E-246 | 6E-244 | 0.4240261 | n.a. |
| EMB | 1.5862 | 9.3814 | 2308 | 9E-241 | 1E-238 | 0.3065342 | surface |
| STAP1 | 2.139 | 9.3165 | 7544 | 1E-236 | 2E-234 | 0.285426 | n.a. |
| IGHG1 | 2.6955 | 9.3682 | 4497 | 4E-235 | 6E-233 | 0.2643179 | n.a. |
| FABP3 | 2.071 | 9.3412 | 4056 | 1E-226 | 2E-224 | 0.2784417 | n.a. |

| | | | | | | | |
|---|---|---|---|---|---|---|---|
| CD82 | 1.8261 | 9.4652 | 1411 | 2E-218 | 3E-216 | 0.4912308 | surface |
| CD19 | 1.9609 | 9.2936 | 7938 | 2E-212 | 3E-210 | 0.2956697 | surface |
| BAG2 | 2.1361 | 9.2915 | 7905 | 2E-211 | 2E-209 | 0.2716126 | n.a. |
| APEX1 | 1.7586 | 9.4275 | 1497 | 5E-211 | 6E-209 | 0.4280615 | n.a. |
| LAT2 | 1.4616 | 9.4007 | 2751 | 2E-208 | 3E-206 | 0.3746702 | nonsurface |
| TEX30 | 1.8064 | 9.4443 | 1402 | 3E-207 | 4E-205 | 0.4246469 | n.a. |
| FCRL1 | 2.0333 | 9.2975 | 6232 | 6E-190 | 7E-188 | 0.2840292 | surface |
| UBE2E3 | 1.5572 | 9.3915 | 1456 | 5E-173 | 5E-171 | 0.3211237 | n.a. |
| CYBB | 1.3783 | 9.4653 | 2524 | 7E-171 | 7E-169 | 0.3192612 | surface |
| SWAP70 | 1.852 | 9.3077 | 3624 | 3E-168 | 3E-166 | 0.2857365 | n.a. |
| ENSECAG00000034569 | 1.2484 | 9.7713 | 761.8 | 7E-166 | 6E-164 | 0.3347819 | n.a. |
| POU2AF1 | 1.8837 | 9.2923 | 5458 | 5E-163 | 5E-161 | 0.2562471 | n.a. |
| TFEB | 1.4844 | 9.3544 | 3047 | 8E-160 | 8E-158 | 0.3167779 | n.a. |
| MRPS6 | 1.6352 | 9.4746 | 911.7 | 6E-157 | 5E-155 | 0.4215428 | n.a. |
| HLA-DMA | 1.3938 | 9.6051 | 955.1 | 8E-149 | 7E-147 | 0.5502095 | surface |
| CYSTM1 | 1.1878 | 9.3828 | 1955 | 1E-148 | 9E-147 | 0.2716126 | n.a. |
| RALGPS2 | 1.9727 | 9.3274 | 2687 | 7E-144 | 6E-142 | 0.3496818 | n.a. |
| ENSECAG00000011715 | 1.2557 | 9.8536 | 629.2 | 1E-137 | 1E-135 | 0.6905168 | n.a. |
| HLA-DOA | 1.5466 | 9.3227 | 2531 | 2E-137 | 2E-135 | 0.2616793 | surface |
| HHEX | 1.2511 | 9.4401 | 908.4 | 4E-136 | 3E-134 | 0.4114543 | n.a. |
| SYNGR2 | 1.3057 | 9.3319 | 1869 | 7E-134 | 6E-132 | 0.280925 | n.a. |
| ENSECAG00000014125 | 1.6123 | 9.3437 | 1354 | 1E-133 | 1E-131 | 0.2953593 | n.a. |
| ENSECAG00000030839 | 1.6643 | 9.3107 | 3016 | 3E-124 | 2E-122 | 0.2517461 | n.a. |
| CNPY3 | 1.1659 | 9.4881 | 685.6 | 1E-118 | 8E-117 | 0.413472 | n.a. |
| RPA3 | 1.2912 | 9.6434 | 536.3 | 1E-117 | 7E-116 | 0.5084588 | n.a. |
| NAPSA | 1.0258 | 9.5255 | 1550 | 4E-110 | 3E-108 | 0.3504579 | n.a. |
| ITM2C | 1.4383 | 9.5994 | 606.3 | 5E-110 | 3E-108 | 0.5966165 | surface |
| DNASE2 | 1.1847 | 9.347 | 1393 | 1E-109 | 8E-108 | 0.2739407 | n.a. |
| PYCARD | 1.2709 | 9.489 | 847 | 8E-109 | 5E-107 | 0.4055564 | n.a. |
| ENSECAG00000001010 | 1.7858 | 9.629 | 975.3 | 1E-105 | 9E-104 | 0.4111439 | n.a. |
| BLOC1S2 | 1.2159 | 9.353 | 917.6 | 4E-99 | 2E-97 | 0.2903927 | n.a. |
| FCMR | 1.6644 | 9.3335 | 1459 | 7E-99 | 4E-97 | 0.297377 | n.a. |
| PTPN6 | 1.0635 | 9.58 | 447.2 | 9.4E-95 | 5E-93 | 0.5402763 | n.a. |
| PPP3CA | 1.1244 | 9.3641 | 776.9 | 1E-93 | 5E-92 | 0.297377 | n.a. |
| ENSECAG00000021652 | 1.4853 | 9.3311 | 962.2 | 1.8E-83 | 8E-82 | 0.2678876 | n.a. |
| BIN1 | 1.0475 | 9.382 | 691.2 | 5.5E-82 | 3E-80 | 0.2851156 | n.a. |
| SNRNP200 | 1.3959 | 9.406 | 610.1 | 2.1E-79 | 1E-77 | 0.3811889 | n.a. |
| SNU13 | 1.0404 | 9.6007 | 349.2 | 1.5E-77 | 7E-76 | 0.5407419 | n.a. |
| RTN4 | 1.0579 | 9.4846 | 372.4 | 1.2E-66 | 5E-65 | 0.414248 | n.a. |
| UMAD1 | 1.0507 | 9.3877 | 439.7 | 1.4E-66 | 6E-65 | 0.2742511 | n.a. |
| ENSECAG00000031182 | 1.6561 | 9.2942 | 1727 | 5.6E-54 | 2E-52 | 0.263697 | n.a. |
| STX11 | 1.2148 | 9.3844 | 433.9 | 1.1E-45 | 3E-44 | 0.3377309 | n.a. |

| genes | logFC | logCPM | F | PValue | FDR | percent.exp | Surfacome.Label |
|---|---|---|---|---|---|---|---|
| ENSECAG00000000419 | 3.344 | 10.783 | 12283 | 0 | 0 | 0.9155427 | n.a. |
| ENSECAG00000033316 | 3.1227 | 10.74 | 6382 | 0 | 0 | 0.7868078 | n.a. |
| CD3E | 2.9964 | 10.225 | 10469 | 0 | 0 | 0.8334756 | n.a. |
| ENSECAG00000030258 | 2.945 | 11.389 | 9204 | 0 | 0 | 0.9570315 | n.a. |
| CD3G | 2.8574 | 10.358 | 11036 | 0 | 0 | 0.872005 | surface |
| LTB | 2.5424 | 10.681 | 6953 | 0 | 0 | 0.9003472 | n.a. |
| STMN1 | 2.4702 | 9.7683 | 5479 | 0 | 0 | 0.5056058 | n.a. |
| CD3D | 2.3398 | 9.8018 | 5637 | 0 | 0 | 0.6275112 | surface |
| GIMAP7 | 2.3037 | 10.513 | 6733 | 0 | 0 | 0.8732571 | n.a. |
| SKAP1 | 2.2761 | 9.6803 | 5156 | 0 | 0 | 0.56303 | n.a. |
| LY6E | 2.2078 | 10.241 | 4653 | 0 | 0 | 0.8372318 | surface |
| CD2 | 2.2023 | 9.8338 | 4626 | 0 | 0 | 0.6044619 | surface |
| BCL11B | 2.19 | 9.6169 | 4790 | 0 | 0 | 0.4815321 | n.a. |
| LEF1 | 2.1884 | 9.4498 | 5628 | 0 | 0 | 0.3505777 | n.a. |
| CD5 | 2.0943 | 9.6499 | 4181 | 0 | 0 | 0.4707757 | surface |
| RRP1B | 2.0916 | 9.9368 | 3898 | 0 | 0 | 0.6986512 | n.a. |
| LCK | 2.0488 | 9.7581 | 4164 | 0 | 0 | 0.5567128 | n.a. |
| TRAT1 | 1.8696 | 9.7859 | 3574 | 0 | 0 | 0.4898412 | surface |
| CD27 | 1.8286 | 9.4777 | 3882 | 0 | 0 | 0.3027147 | surface |
| ENSECAG00000034569 | 1.7654 | 9.7713 | 3470 | 0 | 0 | 0.4519379 | n.a. |
| CCR7 | 1.7425 | 9.4631 | 3046 | 0 | 0 | 0.3146662 | surface |
| LAT | 1.7364 | 9.5366 | 3143 | 0 | 0 | 0.407262 | nonsurface |
| SERPINB6 | 1.6892 | 9.5154 | 2767 | 0 | 0 | 0.3683342 | n.a. |
| ID3 | 1.6733 | 9.6697 | 2396 | 0 | 0 | 0.4392465 | n.a. |
| ENSECAG00000014696 | 1.6271 | 9.6 | 2375 | 0 | 0 | 0.4519948 | n.a. |
| ENSECAG00000019029 | 1.5389 | 9.9278 | 2277 | 0 | 0 | 0.6682033 | n.a. |
| EVL | 1.5038 | 9.7253 | 2010 | 0 | 0 | 0.5492573 | n.a. |
| ENSECAG00000000910 | 1.4962 | 9.7673 | 2217 | 0 | 0 | 0.5816971 | n.a. |
| TMEM204 | 1.4872 | 9.4088 | 2367 | 0 | 0 | 0.283137 | surface |
| RCAN3 | 1.4822 | 9.4465 | 2041 | 0 | 0 | 0.3227477 | n.a. |
| ZAP70 | 1.4274 | 9.4322 | 2365 | 0 | 0 | 0.2761937 | n.a. |
| CD69 | 1.4192 | 9.4382 | 2054 | 0 | 0 | 0.2917876 | surface |
| ICAM2 | 1.3218 | 9.4874 | 2186 | 0 | 0 | 0.3173411 | surface |
| NCR3 | 1.3041 | 9.482 | 2151 | 0 | 0 | 0.2876899 | surface |
| FYB1 | 1.2979 | 9.6072 | 1936 | 0 | 0 | 0.4302544 | n.a. |
| ENSECAG00000028632 | 1.2767 | 9.5007 | 2070 | 0 | 0 | 0.2678846 | n.a. |
| ENSECAG00000007663 | 1.2527 | 9.734 | 2019 | 0 | 0 | 0.469296 | n.a. |
| ENSECAG00000029716 | 1.2229 | 9.8771 | 1884 | 0 | 0 | 0.5977463 | n.a. |
| IDO1 | 1.1784 | 9.5783 | 1756 | 0 | 0 | 0.3242274 | n.a. |
| ENSECAG00000040180 | 1.1061 | 9.7881 | 1543 | 0 | 0 | 0.5062888 | n.a. |
| RF00091 | 1.0899 | 12.032 | 5300 | 0 | 0 | 0.9971544 | n.a. |
| RPSA | 1.0861 | 12.095 | 5406 | 0 | 0 | 0.9971544 | n.a. |
| ENSECAG00000020152 | 1.083 | 11.273 | 3566 | 0 | 0 | 0.9392749 | n.a. |
| ENSECAG00000007681 | 1.0581 | 9.6603 | 1378 | 1E-295 | 6E-294 | 0.4120995 | n.a. |

| | | | | | | | |
|---|---|---|---|---|---|---|---|
| INPP4B | 1.1621 | 9.4661 | 1716 | 1E-264 | 7E-263 | 0.2818849 | n.a. |
| PVRIG | 1.0279 | 9.5646 | 1145 | 5E-247 | 3E-245 | 0.3909851 | n.a. |
| GPR183 | 1.1532 | 9.6621 | 1142 | 3E-246 | 2E-244 | 0.3850094 | surface |
| GBP5 | 1.074 | 9.7163 | 1067 | 2E-230 | 8E-229 | 0.4905811 | nonsurface |
| WARS | 1.0457 | 9.8298 | 1017 | 5E-220 | 2E-218 | 0.5852826 | n.a. |
| PSIP1 | 1.0812 | 9.7368 | 1010 | 1E-218 | 6E-217 | 0.538444 | n.a. |
| TRAF3IP3 | 1.0547 | 9.801 | 1002 | 7E-217 | 3E-215 | 0.5730465 | nonsurface |
| GPX4 | 1.0414 | 9.7124 | 898.7 | 6E-195 | 3E-193 | 0.5015082 | n.a. |
| IL7R | 1.2018 | 9.3731 | 2726 | 6E-164 | 2E-162 | 0.4498321 | surface |

| genes | logFC | logCPM | F | PValue | FDR | percent.exp | Surfacome.Label |
|---|---|---|---|---|---|---|---|
| CCL5 | 6.3163 | 11.908 | 67807 | 0 | 0 | 0.9923664 | n.a. |
| CTSW | 4.9141 | 9.899 | 31272 | 0 | 0 | 0.8772639 | n.a. |
| ENSECAG00000031322 | 4.0628 | 10.499 | 16301 | 0 | 0 | 0.9063015 | n.a. |
| KLRK1 | 3.8767 | 9.569 | 27554 | 0 | 0 | 0.7033378 | surface |
| ENSECAG00000029287 | 3.6301 | 10.043 | 14594 | 0 | 0 | 0.8120042 | n.a. |
| PRF1 | 3.513 | 9.4454 | 18581 | 0 | 0 | 0.5265679 | n.a. |
| ENSECAG00000022992 | 3.3938 | 9.4307 | 20110 | 0 | 0 | 0.4481365 | n.a. |
| GZMA | 3.315 | 9.4519 | 11942 | 0 | 0 | 0.290675 | n.a. |
| ENSECAG00000032959 | 3.3148 | 9.6944 | 9606 | 0 | 0 | 0.6936087 | n.a. |
| CD7 | 3.2956 | 9.4726 | 13868 | 0 | 0 | 0.4446939 | surface |
| TRAT1 | 2.9903 | 9.7859 | 7535 | 0 | 0 | 0.6719054 | surface |
| ENSECAG00000008322 | 2.9221 | 9.4726 | 9519 | 0 | 0 | 0.5437809 | n.a. |
| CD3E | 2.9032 | 10.225 | 11057 | 0 | 0 | 0.8693309 | n.a. |
| GZMM | 2.8919 | 9.4961 | 4858 | 0 | 0 | 0.4698398 | n.a. |
| CD3G | 2.8885 | 10.358 | 12563 | 0 | 0 | 0.9166292 | surface |
| NKG7 | 2.8653 | 9.3783 | 12184 | 0 | 0 | 0.4343661 | nonsurface |
| ENSECAG00000000419 | 2.8261 | 10.783 | 9842 | 0 | 0 | 0.9107918 | n.a. |
| ENSECAG00000033316 | 2.813 | 10.74 | 6110 | 0 | 0 | 0.7849124 | n.a. |
| NPY | 2.5398 | 9.5043 | 2948 | 0 | 0 | 0.359527 | n.a. |
| LCK | 2.5031 | 9.7581 | 5847 | 0 | 0 | 0.666517 | n.a. |
| CD8A | 2.489 | 9.4281 | 6325 | 0 | 0 | 0.3189642 | surface |
| GIMAP7 | 2.4653 | 10.513 | 8329 | 0 | 0 | 0.8857955 | n.a. |
| ID2 | 2.451 | 9.6451 | 5635 | 0 | 0 | 0.5542583 | n.a. |
| ENSECAG00000030258 | 2.4385 | 11.389 | 6927 | 0 | 0 | 0.9378836 | n.a. |
| ENSECAG00000039088 | 2.3301 | 9.7274 | 5360 | 0 | 0 | 0.7159108 | n.a. |
| LGALS1 | 2.3298 | 11.109 | 7390 | 0 | 0 | 0.9110912 | nonsurface |
| CD3D | 2.3104 | 9.8018 | 5402 | 0 | 0 | 0.636731 | surface |
| NCR3 | 2.1252 | 9.482 | 4100 | 0 | 0 | 0.3729981 | surface |
| SH2D1A | 2.1193 | 9.4515 | 3870 | 0 | 0 | 0.3203113 | n.a. |
| CD2 | 2.1124 | 9.8338 | 3464 | 0 | 0 | 0.5964676 | surface |
| ENSECAG00000001010 | 2.081 | 9.629 | 4754 | 0 | 0 | 0.3521928 | n.a. |
| ENSECAG00000007681 | 2.0478 | 9.6603 | 3898 | 0 | 0 | 0.5618919 | n.a. |
| S100A4 | 1.9786 | 11.292 | 3839 | 0 | 0 | 0.7124682 | n.a. |
| HCST | 1.9024 | 9.5985 | 3412 | 0 | 0 | 0.5160904 | n.a. |
| ENSECAG00000028632 | 1.7959 | 9.5007 | 3021 | 0 | 0 | 0.3429127 | n.a. |
| IDO1 | 1.7799 | 9.5783 | 2026 | 0 | 0 | 0.4234396 | n.a. |
| SKAP1 | 1.7568 | 9.6803 | 3072 | 0 | 0 | 0.4672953 | n.a. |
| SCML4 | 1.7507 | 9.3954 | 2667 | 0 | 0 | 0.3013022 | n.a. |
| ENSECAG00000029716 | 1.7364 | 9.8771 | 3355 | 0 | 0 | 0.644664 | n.a. |
| ENSECAG00000007668 | 1.7271 | 9.3879 | 2853 | 0 | 0 | 0.2682233 | n.a. |
| RRP1B | 1.6954 | 9.9368 | 2500 | 0 | 0 | 0.5771591 | n.a. |
| ENSECAG00000040180 | 1.6392 | 9.7881 | 2682 | 0 | 0 | 0.5440802 | n.a. |
| ENSECAG00000007663 | 1.6268 | 9.734 | 2930 | 0 | 0 | 0.5081575 | n.a. |
| MARCKSL1 | 1.5744 | 9.5078 | 1781 | 0 | 0 | 0.3700045 | n.a. |

| | | | | | | | |
|---|---|---|---|---|---|---|---|
| YWHAQ | 1.5699 | 9.7075 | 1939 | 0 | 0 | 0.5927256 | n.a. |
| LTB | 1.5613 | 10.681 | 2180 | 0 | 0 | 0.6668163 | n.a. |
| BCL11B | 1.5411 | 9.6169 | 2386 | 0 | 0 | 0.367909 | n.a. |
| FYN | 1.5259 | 9.4235 | 2168 | 0 | 0 | 0.2779524 | n.a. |
| EVL | 1.5195 | 9.7253 | 1824 | 0 | 0 | 0.5261188 | n.a. |
| ZAP70 | 1.4937 | 9.4322 | 2208 | 0 | 0 | 0.2769047 | n.a. |
| ICAM2 | 1.4899 | 9.4874 | 2183 | 0 | 0 | 0.2930699 | surface |
| GBP5 | 1.4838 | 9.7163 | 1629 | 0 | 0 | 0.5092052 | nonsurface |
| PTPRCAP | 1.359 | 10.617 | 3164 | 0 | 0 | 0.8850471 | nonsurface |
| S100A11 | 1.2827 | 9.847 | 1761 | 0 | 0 | 0.5265679 | n.a. |
| ANXA2 | 1.2662 | 9.8875 | 2167 | 0 | 0 | 0.5690765 | n.a. |
| PTPRC | 1.2073 | 10.647 | 2751 | 0 | 0 | 0.9074989 | surface |
| RAC2 | 1.1733 | 10.594 | 2667 | 0 | 0 | 0.9262087 | n.a. |
| ADGRE5 | 1.3257 | 9.5791 | 1480 | 0 | 0 | 0.4854064 | n.a. |
| ZNF683 | 2.3085 | 9.478 | 3232 | 0 | 0 | 0.4171531 | n.a. |
| INPP4B | 1.376 | 9.4661 | 1976 | 3E-304 | 3E-302 | 0.2713666 | n.a. |
| MPC2 | 1.3475 | 9.6359 | 1400 | 2E-300 | 2E-298 | 0.4958838 | nonsurface |
| SERPINB6 | 1.2932 | 9.5154 | 1500 | 2E-290 | 2E-288 | 0.298159 | n.a. |
| TIFA | 1.2464 | 10.029 | 1346 | 4E-289 | 4E-287 | 0.6810358 | n.a. |
| CD5 | 1.4649 | 9.6499 | 1563 | 2E-288 | 2E-286 | 0.4643018 | surface |
| LGALS3 | 1.2731 | 9.5239 | 1927 | 2E-286 | 1E-284 | 0.3620715 | n.a. |
| EMP3 | 1.1715 | 10.126 | 1329 | 2E-285 | 1E-283 | 0.7642568 | surface |
| ENSECAG00000019698 | 1.7296 | 9.4686 | 1833 | 5E-285 | 4E-283 | 0.3869181 | n.a. |
| PVRIG | 1.2447 | 9.5646 | 1302 | 8E-280 | 6E-278 | 0.3854213 | n.a. |
| CTSC | 1.3553 | 9.6133 | 1293 | 6E-278 | 5E-276 | 0.4214938 | n.a. |
| ENSECAG00000014696 | 1.2588 | 9.6 | 1261 | 3E-271 | 2E-269 | 0.3592277 | n.a. |
| ITGAL | 1.3217 | 9.4047 | 1639 | 6E-267 | 5E-265 | 0.3150726 | surface |
| UBD | 1.248 | 9.9806 | 1091 | 1E-235 | 8E-234 | 0.5918276 | n.a. |
| RBL2 | 1.3276 | 9.388 | 1375 | 7E-233 | 5E-231 | 0.2626852 | n.a. |
| SLC9A3R1 | 1.1527 | 9.5551 | 1054 | 9E-228 | 6E-226 | 0.4048795 | n.a. |
| ENSECAG00000038252 | 1.2524 | 9.397 | 1397 | 6E-219 | 4E-217 | 0.2505613 | n.a. |
| ENSECAG00000016543 | 1.1265 | 9.8125 | 999.7 | 2E-216 | 2E-214 | 0.5921269 | n.a. |
| GBP6 | 1.2115 | 9.5114 | 985.8 | 2E-207 | 2E-205 | 0.3008532 | nonsurface |
| ENSECAG00000006071 | 1.1302 | 9.6095 | 930.5 | 1E-201 | 7E-200 | 0.4113157 | n.a. |
| ITGB1 | 1.0549 | 9.5309 | 927.6 | 5E-201 | 3E-199 | 0.3117797 | surface |
| LAT | 1.159 | 9.5366 | 1180 | 3E-199 | 2E-197 | 0.3268972 | nonsurface |
| CD53 | 1.0118 | 9.709 | 903 | 7E-196 | 4E-194 | 0.5529112 | surface |
| GIMAP6 | 1.0621 | 9.7575 | 875.5 | 5E-190 | 3E-188 | 0.5175872 | n.a. |
| FABP5 | 1.2389 | 9.5264 | 1017 | 2E-188 | 1E-186 | 0.299057 | n.a. |
| NPDC1 | 1.146 | 9.5117 | 886.4 | 4E-184 | 2E-182 | 0.3357282 | n.a. |
| SMCO4 | 1.0252 | 9.5303 | 765.6 | 1E-166 | 6E-165 | 0.3524921 | nonsurface |
| NMNAT3 | 1.4571 | 9.394 | 1295 | 4E-164 | 2E-162 | 0.2996557 | n.a. |
| ENSECAG00000003563 | 1.0878 | 9.4315 | 879.7 | 9E-157 | 5E-155 | 0.2797485 | n.a. |
| ENSECAG00000015137 | 2.236 | 9.3925 | 1495 | 1.9E-68 | 5E-67 | 0.3559347 | n.a. |
| GZMK | 2.1413 | 9.5782 | 175 | 1.1E-16 | 9E-16 | 0.3530909 | n.a. |

| genes | logFC | logCPM | F | PValue | FDR | percent.exp | Surfacome.Label |
|---|---|---|---|---|---|---|---|
| FCER1G | 5.5875 | 10.204 | 24540 | 0 | 0 | 0.9688448 | nonsurface |
| SYNE4 | 5.349 | 10.39 | 35150 | 0 | 0 | 0.9988972 | nonsurface |
| CST3 | 4.6466 | 11.921 | 3563 | 0 | 0 | 0.9991729 | n.a. |
| SRGN | 3.1966 | 10.485 | 4045 | 0 | 0 | 0.9762889 | n.a. |
| FTL | 3.0523 | 11.187 | 12177 | 0 | 0 | 0.99807 | n.a. |
| SAT1 | 2.6258 | 9.7271 | 3852 | 0 | 0 | 0.9434795 | n.a. |
| VIM | 2.2376 | 11.286 | 2772 | 0 | 0 | 0.9906259 | nonsurface |
| CYBA | 1.7984 | 10.509 | 1936 | 0 | 0 | 0.9931073 | nonsurface |
| ALDOA | 1.5777 | 9.99 | 1577 | 0 | 0 | 0.970499 | n.a. |
| COTL1 | 1.5122 | 10.529 | 1951 | 0 | 0 | 0.9922801 | n.a. |
| FTH1 | 1.3669 | 10.516 | 1764 | 0 | 0 | 0.9473394 | nonsurface |
| UCP2 | 1.791 | 9.9052 | 1416 | 1E-303 | 1E-301 | 0.9597463 | nonsurface |
| SPI1 | 3.397 | 9.6618 | 4614 | 3E-301 | 4E-299 | 0.9473394 | n.a. |
| ZFP36 | 1.9946 | 9.4955 | 1946 | 6E-283 | 7E-281 | 0.7714364 | n.a. |
| ATP6V1G1 | 1.663 | 9.9335 | 1175 | 3E-253 | 3E-251 | 0.9655363 | n.a. |
| AMDHD2 | 1.5594 | 9.7855 | 1131 | 4E-244 | 4E-242 | 0.9305211 | n.a. |
| SNX3 | 1.4571 | 9.6999 | 1071 | 2E-231 | 2E-229 | 0.8657293 | n.a. |
| PGD | 1.9954 | 9.3424 | 3070 | 3E-222 | 2E-220 | 0.5971878 | n.a. |
| GABARAP | 1.0203 | 10.489 | 928.8 | 2E-201 | 2E-199 | 0.9878688 | n.a. |
| ATP6V0B | 1.8047 | 9.5375 | 1152 | 5E-192 | 4E-190 | 0.8433967 | nonsurface |
| GAPDH | 1.2419 | 10.705 | 876.6 | 3E-190 | 2E-188 | 0.9909016 | n.a. |
| ENSECAG00000030271 | 2.0578 | 9.3599 | 2553 | 2E-189 | 2E-187 | 0.6523297 | n.a. |
| TALDO1 | 1.2385 | 9.9099 | 854.6 | 1E-185 | 1E-183 | 0.9365867 | n.a. |
| TKT | 1.9425 | 9.6069 | 1197 | 7E-176 | 5E-174 | 0.9076372 | n.a. |
| RNF130 | 2.3121 | 9.3597 | 2522 | 3E-164 | 2E-162 | 0.6506755 | surface |
| ITM2B | 1.3469 | 10.174 | 737.9 | 8E-161 | 6E-159 | 0.9346567 | surface |
| PSAP | 2.4695 | 10.167 | 715.2 | 6E-156 | 4E-154 | 0.9671905 | n.a. |
| GRN | 2.4178 | 9.4381 | 1975 | 3E-153 | 2E-151 | 0.8053488 | n.a. |
| C5H1orf162 | 1.9838 | 9.2673 | 8988 | 1E-145 | 6E-144 | 0.2674387 | n.a. |
| NFKBIA | 1.2533 | 9.5431 | 593.9 | 4E-130 | 3E-128 | 0.6275159 | n.a. |
| FGR | 2.8722 | 9.4883 | 1986 | 3E-128 | 2E-126 | 0.8880618 | n.a. |
| ATG3 | 1.2608 | 9.6124 | 598.9 | 1E-126 | 7E-125 | 0.8632479 | n.a. |
| ENSECAG00000003925 | 1.0738 | 9.5059 | 619.8 | 3E-118 | 1E-116 | 0.5789909 | n.a. |
| PLBD1 | 3.3221 | 9.972 | 1339 | 7E-117 | 4E-115 | 0.9054315 | n.a. |
| PNPLA2 | 1.4326 | 9.3533 | 1253 | 2E-111 | 1E-109 | 0.5745795 | nonsurface |
| LYZ | 4.2612 | 11.749 | 490.8 | 5E-108 | 3E-106 | 0.8671078 | n.a. |
| JUNB | 1.231 | 9.5395 | 512.1 | 7E-108 | 3E-106 | 0.7245658 | n.a. |
| CTSS | 3.117 | 9.8161 | 837.1 | 3E-106 | 1E-104 | 0.9517508 | n.a. |
| ENSECAG00000007123 | 1.2067 | 9.3975 | 640.6 | 7E-104 | 3E-102 | 0.5301902 | n.a. |
| TMEM59 | 1.2096 | 9.5994 | 460.7 | 4E-100 | 2E-98 | 0.7714364 | nonsurface |
| PLEK | 1.372 | 9.3075 | 1820 | 8E-100 | 4E-98 | 0.3325062 | n.a. |
| HSBP1 | 1.5801 | 9.5012 | 667.8 | 1E-99 | 7E-98 | 0.8059002 | n.a. |
| EFHD2 | 1.6094 | 9.3994 | 923.5 | 3E-99 | 1E-97 | 0.6148332 | n.a. |
| HSPB1 | 2.5288 | 9.6778 | 604.6 | 2.8E-97 | 1E-95 | 0.9266612 | n.a. |

| | | | | | | | |
|---|---|---|---|---|---|---|---|
| ARPC3 | 1.018 | 10.061 | 435.7 | 3.6E-96 | 2E-94 | 0.9500965 | n.a. |
| ANXA2 | 1.2075 | 9.8875 | 474.7 | 7.6E-96 | 3E-94 | 0.9175627 | n.a. |
| ZYX | 1.6402 | 9.4366 | 707.4 | 2.9E-92 | 1E-90 | 0.6967191 | n.a. |
| ATP5F1D | 1.3251 | 9.6297 | 498.5 | 3.4E-91 | 1E-89 | 0.8855804 | n.a. |
| TSTD3 | 1.737 | 9.4008 | 1025 | 3.3E-88 | 1E-86 | 0.6804522 | n.a. |
| EVI2B | 1.0151 | 9.5277 | 387.4 | 9E-86 | 4E-84 | 0.6087676 | surface |
| VSIR | 1.3748 | 9.4326 | 644 | 1.1E-85 | 4E-84 | 0.700579 | n.a. |
| LAMP1 | 1.564 | 9.6692 | 497.5 | 3.1E-84 | 1E-82 | 0.7956989 | surface |
| ANXA5 | 1.7925 | 9.5163 | 644.9 | 1.1E-83 | 4E-82 | 0.8265784 | n.a. |
| CYSTM1 | 2.1705 | 9.3828 | 1258 | 4.6E-82 | 2E-80 | 0.3733113 | n.a. |
| ENSECAG00000019932 | 1.622 | 9.3315 | 1099 | 1.7E-80 | 7E-79 | 0.4403088 | n.a. |
| RHOB | 1.7651 | 9.2893 | 1889 | 9.4E-79 | 4E-77 | 0.2575131 | n.a. |
| RGS19 | 1.0391 | 9.5283 | 387.5 | 4.9E-78 | 2E-76 | 0.7190516 | n.a. |
| PGK1 | 1.0628 | 9.4881 | 431 | 1.1E-77 | 4E-76 | 0.7198787 | n.a. |
| FGL2 | 2.4336 | 9.488 | 873.6 | 6.4E-76 | 2E-74 | 0.8491867 | n.a. |
| YPEL5 | 1.0852 | 9.4914 | 364.9 | 4.1E-74 | 2E-72 | 0.6040805 | n.a. |
| PGLS | 1.2979 | 9.6893 | 342.7 | 7.2E-72 | 3E-70 | 0.9076372 | n.a. |
| IFNGR1 | 1.3632 | 9.4289 | 524.6 | 8.8E-69 | 3E-67 | 0.643507 | surface |
| LY96 | 1.2122 | 9.3807 | 557 | 1E-68 | 4E-67 | 0.4827681 | n.a. |
| MSRA | 1.1342 | 9.2852 | 1550 | 1.7E-68 | 6E-67 | 0.3540116 | n.a. |
| MXD1 | 1.1103 | 9.3408 | 534.8 | 1.1E-65 | 4E-64 | 0.3410532 | n.a. |
| ATG16L2 | 1.4192 | 9.3409 | 1027 | 1.2E-65 | 4E-64 | 0.4910394 | n.a. |
| RARA | 1.4023 | 9.3162 | 865.9 | 1.1E-64 | 4E-63 | 0.4108078 | n.a. |
| ANXA7 | 1.0268 | 9.5774 | 306 | 2E-63 | 6E-62 | 0.8045216 | n.a. |
| LAMTOR3 | 1.063 | 9.3658 | 479.5 | 1E-60 | 3E-59 | 0.4811139 | n.a. |
| TCIRG1 | 1.5742 | 9.3151 | 1174 | 1.3E-59 | 4E-58 | 0.5166805 | surface |
| RNASE4 | 4.1739 | 10.101 | 514.6 | 3.6E-57 | 1E-55 | 0.9520265 | n.a. |
| ANXA1 | 1.1622 | 9.8295 | 252.3 | 1.3E-56 | 4E-55 | 0.8924731 | n.a. |
| NPC2 | 2.0785 | 9.8367 | 270.5 | 2.1E-55 | 6E-54 | 0.9178384 | n.a. |
| TFEB | 1.2367 | 9.3544 | 1115 | 3.4E-55 | 9E-54 | 0.4877309 | n.a. |
| HK1 | 1.1361 | 9.3001 | 782.4 | 1.2E-54 | 3E-53 | 0.3121037 | n.a. |
| AGPAT2 | 1.1201 | 9.4052 | 391.1 | 1.7E-52 | 5E-51 | 0.554177 | nonsurface |
| LAPTM4A | 1.28 | 9.5901 | 261.7 | 6.5E-52 | 2E-50 | 0.7457954 | nonsurface |
| RNF166 | 1.0827 | 9.3001 | 677.8 | 1.3E-50 | 3E-49 | 0.3209264 | n.a. |
| GSN | 2.3346 | 9.3636 | 1393 | 2.1E-50 | 5E-49 | 0.6619796 | nonsurface |
| DRAXIN | 1.6052 | 9.3226 | 839.6 | 2.5E-50 | 7E-49 | 0.4717397 | n.a. |
| TAX1BP3 | 1.3939 | 9.3543 | 538.3 | 4.2E-50 | 1E-48 | 0.5034464 | n.a. |
| LRMP | 1.0128 | 9.3198 | 631.6 | 2.7E-49 | 7E-48 | 0.3145851 | nonsurface |
| ENSECAG00000035431 | 2.0884 | 9.323 | 2860 | 2.8E-49 | 7E-48 | 0.6258616 | n.a. |
| LDHA | 1.2057 | 9.6788 | 247.9 | 4.4E-49 | 1E-47 | 0.8756548 | n.a. |
| LYN | 1.7661 | 9.3216 | 1073 | 1.3E-47 | 3E-46 | 0.4460987 | surface |
| ENSECAG00000035305 | 1.1693 | 10.092 | 209 | 3.1E-47 | 7E-46 | 0.9161842 | n.a. |
| CYBB | 1.8372 | 9.4653 | 703.8 | 2.6E-46 | 6E-45 | 0.7146402 | surface |
| PCMT1 | 1.1559 | 9.2898 | 817.9 | 6.5E-46 | 2E-44 | 0.3076923 | n.a. |
| MYD88 | 1.0692 | 9.3632 | 371.7 | 1.9E-44 | 4E-43 | 0.4381031 | n.a. |

| | | | | | | | |
|---|---|---|---|---|---|---|---|
| ENSECAG00000017935 | 1.0401 | 9.3545 | 407.8 | 6.4E-44 | 1E-42 | 0.4604356 | n.a. |
| SERPINB1 | 1.7524 | 9.4507 | 376.3 | 1.4E-43 | 3E-42 | 0.72622 | nonsurface |
| DPYD | 1.564 | 9.2972 | 1446 | 1.2E-41 | 3E-40 | 0.4596085 | n.a. |
| TSPO | 2.9953 | 9.7323 | 315.7 | 2.2E-41 | 5E-40 | 0.8596636 | nonsurface |
| RNASE6 | 4.0598 | 10.205 | 329.4 | 2.9E-41 | 6E-40 | 0.9564378 | n.a. |
| CREG1 | 1.7737 | 9.3229 | 2235 | 3.6E-41 | 8E-40 | 0.5596912 | n.a. |
| PAK1 | 2.3112 | 9.331 | 1962 | 1.4E-39 | 3E-38 | 0.6333058 | n.a. |
| SELENOS | 1.4226 | 9.3395 | 481.7 | 3.6E-39 | 7E-38 | 0.4932451 | n.a. |
| CPNE3 | 1.0795 | 9.3269 | 412 | 4.3E-38 | 9E-37 | 0.3655914 | n.a. |
| LAMTOR1 | 1.3468 | 9.5184 | 232.6 | 5.1E-38 | 1E-36 | 0.8012131 | n.a. |
| NAPSA | 2.3976 | 9.5255 | 545.7 | 2E-37 | 4E-36 | 0.8425696 | n.a. |
| ALDH2 | 1.6404 | 9.5616 | 479.4 | 8.2E-37 | 2E-35 | 0.781362 | n.a. |
| GNAI2 | 1.3447 | 9.598 | 204.7 | 2.8E-36 | 5E-35 | 0.870692 | n.a. |
| CD63 | 2.0052 | 9.4159 | 378.8 | 2.9E-36 | 6E-35 | 0.6757651 | surface |
| ETHE1 | 1.0351 | 9.3855 | 300.6 | 6.7E-36 | 1E-34 | 0.5720982 | n.a. |
| REEP3 | 1.0988 | 9.3317 | 397.7 | 1.1E-35 | 2E-34 | 0.4196305 | nonsurface |
| NUDT4 | 1.1655 | 9.2991 | 542.5 | 2.9E-35 | 5E-34 | 0.3192721 | n.a. |
| TMEM70 | 1.014 | 9.2927 | 633.7 | 3.4E-35 | 6E-34 | 0.3272677 | nonsurface |
| H2AFJ | 1.625 | 9.3551 | 500.4 | 1.4E-34 | 2E-33 | 0.6140061 | n.a. |
| IRF2BP2 | 1.4368 | 9.4142 | 279.2 | 1.6E-34 | 3E-33 | 0.6495726 | n.a. |
| FGGY | 1.1054 | 9.2697 | 1754 | 2E-33 | 4E-32 | 0.3085194 | n.a. |
| DUSP1 | 2.6881 | 9.3756 | 440.4 | 3.4E-33 | 6E-32 | 0.5406672 | nonsurface |
| GRINA | 1.2395 | 9.3626 | 306.4 | 2.3E-32 | 4E-31 | 0.4858009 | nonsurface |
| TRABD | 1.0375 | 9.361 | 302.3 | 9.8E-32 | 2E-30 | 0.5312931 | nonsurface |
| GNG10 | 1.7183 | 9.6838 | 165.4 | 2.6E-31 | 4E-30 | 0.8888889 | n.a. |
| PPT1 | 1.6846 | 9.4371 | 260.4 | 1.1E-30 | 2E-29 | 0.7234629 | n.a. |
| ARPC1B | 1.079 | 9.4598 | 247.6 | 2.5E-30 | 4E-29 | 0.4808382 | n.a. |
| IFIT2 | 2.0458 | 9.3061 | 702 | 9.3E-30 | 2E-28 | 0.4066722 | n.a. |
| RHBDD2 | 1.0501 | 9.3113 | 393.6 | 1.5E-29 | 2E-28 | 0.3774469 | nonsurface |
| NFE2L2 | 1.3323 | 9.2896 | 692.5 | 1.9E-29 | 3E-28 | 0.3614557 | n.a. |
| B4GALT1 | 1.143 | 9.2937 | 482.7 | 3E-29 | 5E-28 | 0.311828 | nonsurface |
| RP2 | 1.1109 | 9.3003 | 435.6 | 4.6E-29 | 7E-28 | 0.3338848 | n.a. |
| RNF149 | 1.0867 | 9.3444 | 318.8 | 1.3E-28 | 2E-27 | 0.4891095 | surface |
| NAGK | 2.0399 | 9.431 | 310.8 | 1.8E-28 | 3E-27 | 0.7799835 | n.a. |
| C1H15orf48 | 2.5631 | 9.6929 | 256.8 | 8.9E-28 | 1E-26 | 0.7659222 | n.a. |
| ENSECAG00000016578 | 2.9676 | 9.4164 | 1166 | 1.2E-27 | 2E-26 | 0.8458781 | n.a. |
| RAB24 | 1.0991 | 9.4289 | 174.9 | 1.3E-27 | 2E-26 | 0.4323132 | n.a. |
| STK10 | 1.0122 | 9.392 | 208.2 | 1.4E-27 | 2E-26 | 0.4935208 | n.a. |
| ID2 | 1.0941 | 9.6451 | 145 | 1.5E-27 | 2E-26 | 0.6479184 | n.a. |
| C22H20orf27 | 1.0868 | 9.3454 | 318.3 | 3E-26 | 4E-25 | 0.5613455 | n.a. |
| RAB8B | 1.03 | 9.3036 | 371.6 | 7.4E-26 | 1E-24 | 0.3391232 | n.a. |
| CAMKK2 | 1.0304 | 9.2678 | 906.6 | 1E-25 | 1E-24 | 0.2828784 | n.a. |
| BAZ1A | 1.095 | 9.3687 | 224.8 | 2.6E-25 | 4E-24 | 0.4946237 | n.a. |
| PRNP | 1.7512 | 9.3261 | 525.6 | 8.3E-25 | 1E-23 | 0.5169562 | nonsurface |
| GLIPR2 | 1.1372 | 9.47 | 201.6 | 8.9E-25 | 1E-23 | 0.6859664 | n.a. |

| | | | | | | | |
|---|---|---|---|---|---|---|---|
| ENSECAG00000020136 | 1.3747 | 9.7014 | 106.8 | 1.8E-23 | 2E-22 | 0.7639923 | n.a. |
| ECHDC2 | 1.0312 | 9.3064 | 339 | 1.8E-23 | 2E-22 | 0.3586986 | n.a. |
| REXO2 | 1.0275 | 9.4557 | 178.4 | 2.7E-23 | 4E-22 | 0.6992004 | n.a. |
| ATP6V1B2 | 1.015 | 9.3784 | 205.7 | 2.9E-23 | 4E-22 | 0.573201 | n.a. |
| GM2A | 4.1692 | 9.9979 | 270.7 | 1.9E-22 | 2E-21 | 0.9261097 | n.a. |
| ENSECAG00000017042 | 2.1649 | 9.3458 | 992.3 | 2E-22 | 2E-21 | 0.6379928 | n.a. |
| NFAM1 | 2.4883 | 9.3582 | 620 | 6.2E-22 | 8E-21 | 0.7116074 | surface |
| ENSECAG00000032188 | 1.1675 | 9.5863 | 129.7 | 2.5E-21 | 3E-20 | 0.8444996 | n.a. |
| CLEC12A | 2.9294 | 9.4082 | 814.8 | 7.6E-21 | 9E-20 | 0.787979 | surface |
| SELL | 1.2591 | 9.6904 | 94.1 | 1.2E-20 | 1E-19 | 0.7579267 | surface |
| ITGAX | 1.0297 | 9.2642 | 1005 | 1.5E-20 | 2E-19 | 0.2668872 | surface |
| ENSECAG00000014585 | 1.5921 | 10.271 | 124.8 | 2.8E-20 | 3E-19 | 0.6330301 | n.a. |
| C4BPA | 2.6906 | 9.4013 | 565.3 | 3.4E-20 | 4E-19 | 0.7138131 | n.a. |
| ENSECAG00000012199 | 1.2479 | 9.3154 | 434.3 | 4.3E-20 | 5E-19 | 0.3578715 | n.a. |
| PICALM | 1.2809 | 9.3149 | 283.6 | 5.3E-20 | 6E-19 | 0.4405845 | n.a. |
| IFNGR2 | 1.6817 | 9.3299 | 370 | 9.1E-20 | 1E-18 | 0.5616212 | surface |
| TNFRSF1B | 1.0733 | 9.3575 | 223.2 | 2.6E-19 | 3E-18 | 0.5260546 | surface |
| COL4A3BP | 1.3257 | 9.327 | 250.8 | 4.1E-19 | 4E-18 | 0.4529915 | n.a. |
| NADK | 1.4687 | 9.2968 | 398.6 | 5.3E-19 | 6E-18 | 0.3868211 | n.a. |
| STX7 | 1.0193 | 9.3568 | 213.6 | 7.9E-19 | 9E-18 | 0.4687069 | nonsurface |
| ENSECAG00000015010 | 1.3843 | 9.2872 | 898.2 | 3.8E-17 | 4E-16 | 0.4127378 | n.a. |
| KIAA0513 | 1.2382 | 9.2674 | 578.1 | 1E-16 | 1E-15 | 0.2710229 | n.a. |
| TIMP2 | 1.6794 | 9.2993 | 700.7 | 1.5E-16 | 1E-15 | 0.5125448 | n.a. |
| ALDH1L1 | 1.3227 | 9.2841 | 627.2 | 5.6E-16 | 5E-15 | 0.4154949 | n.a. |
| BLVRB | 1.2419 | 9.4497 | 123 | 2.7E-15 | 2E-14 | 0.7598566 | n.a. |
| ENSECAG00000024719 | 1.0364 | 9.2957 | 221.9 | 2.9E-15 | 3E-14 | 0.2558588 | n.a. |
| PRDX5 | 1.1048 | 9.5264 | 82.62 | 6.5E-15 | 6E-14 | 0.7295285 | n.a. |
| MBOAT7 | 1.2622 | 9.2726 | 422.5 | 9.2E-15 | 8E-14 | 0.2726771 | nonsurface |
| IRF5 | 1.2905 | 9.322 | 311.4 | 1.2E-14 | 1E-13 | 0.4394817 | n.a. |
| NCF1 | 1.7884 | 9.2933 | 470.2 | 1.2E-14 | 1E-13 | 0.4105321 | n.a. |
| LAT2 | 1.417 | 9.4007 | 192 | 2.2E-14 | 2E-13 | 0.6203474 | nonsurface |
| CTSZ | 2.6049 | 9.8631 | 118.1 | 2.7E-14 | 2E-13 | 0.9357596 | n.a. |
| FAM49A | 1.3423 | 9.2748 | 474.8 | 3.5E-14 | 3E-13 | 0.3159636 | n.a. |
| S100A12 | 3.4107 | 10.309 | 143.3 | 3.5E-14 | 3E-13 | 0.7336642 | n.a. |
| RGS18 | 1.8604 | 9.3114 | 269.4 | 4.6E-14 | 4E-13 | 0.3325062 | n.a. |
| EIF4E1B | 1.2576 | 9.2561 | 1050 | 4.8E-14 | 4E-13 | 0.2500689 | n.a. |
| TGFB1 | 1.1235 | 9.3709 | 114.5 | 2.7E-13 | 2E-12 | 0.4830438 | n.a. |
| ENSECAG00000000436 | 2.7119 | 9.9635 | 265.9 | 3E-13 | 2E-12 | 0.7303557 | n.a. |
| C20H6orf62 | 1.1748 | 9.3936 | 97.45 | 4.6E-13 | 4E-12 | 0.475324 | n.a. |
| ATP1B3 | 1.2415 | 9.5391 | 72.63 | 7.8E-13 | 6E-12 | 0.7637166 | surface |
| TREM1 | 2.4112 | 9.4852 | 428.7 | 1.3E-12 | 1E-11 | 0.7063689 | surface |
| RF01956 | 1.5209 | 9.2926 | 298.6 | 1.5E-12 | 1E-11 | 0.5577612 | n.a. |
| HK3 | 1.3064 | 9.284 | 588.4 | 2.6E-12 | 2E-11 | 0.4381031 | n.a. |
| TNFRSF1A | 1.1048 | 9.3254 | 160.3 | 2.8E-12 | 2E-11 | 0.4689826 | surface |
| ATP6AP1 | 1.3607 | 9.3521 | 126.8 | 3.3E-12 | 2E-11 | 0.5704439 | nonsurface |

| | | | | | | | |
|---|---|---|---|---|---|---|---|
| ENSECAG00000034000 | 1.0956 | 9.5622 | 60.19 | 6E-11 | 4E-10 | 0.8136201 | n.a. |
| IVNS1ABP | 1.0165 | 9.3077 | 136.1 | 1.3E-10 | 8E-10 | 0.3253377 | n.a. |
| PLD4 | 1.448 | 9.3359 | 184.4 | 4.9E-10 | 3E-09 | 0.4739454 | n.a. |
| ENSECAG00000033936 | 1.319 | 9.27 | 485.4 | 9.5E-09 | 5E-08 | 0.3418803 | n.a. |
| ENSECAG00000033857 | 2.6649 | 9.3316 | 371.4 | 3E-08 | 2E-07 | 0.6071133 | n.a. |
| CSRP1 | 1.3919 | 9.2812 | 180.1 | 6.9E-08 | 4E-07 | 0.2682658 | n.a. |
| ENSECAG00000019318 | 1.6047 | 9.3455 | 121.9 | 8.1E-08 | 4E-07 | 0.346016 | n.a. |
| LGALS3 | 1.0475 | 9.5239 | 48.94 | 8.3E-08 | 4E-07 | 0.6735594 | n.a. |
| FES | 1.2189 | 9.2936 | 168.6 | 9.2E-08 | 5E-07 | 0.3581472 | n.a. |
| TMEM50B | 1.2428 | 9.2661 | 298.4 | 9.6E-08 | 5E-07 | 0.2828784 | nonsurface |
| QPCT | 1.4083 | 9.3093 | 216.3 | 9.9E-08 | 5E-07 | 0.4772539 | nonsurface |
| ENSECAG00000030777 | 2.1721 | 9.3773 | 196.6 | 1E-07 | 5E-07 | 0.4074993 | n.a. |
| CFP | 2.8474 | 9.4343 | 272.4 | 1.3E-07 | 6E-07 | 0.8207885 | n.a. |
| CHPT1 | 1.304 | 9.2751 | 196.1 | 1.3E-07 | 7E-07 | 0.3041081 | surface |
| DHRS3 | 1.1667 | 9.3111 | 120.7 | 1.8E-07 | 9E-07 | 0.4480287 | nonsurface |
| LST1 | 1.7883 | 9.329 | 99.68 | 2E-07 | 1E-06 | 0.4775296 | nonsurface |
| S100P | 3.6319 | 9.8205 | 121.3 | 2E-07 | 1E-06 | 0.7752964 | n.a. |
| IGSF6 | 2.7636 | 9.3651 | 247.5 | 2E-07 | 1E-06 | 0.719603 | surface |
| C5AR1 | 2.5735 | 9.3446 | 287.1 | 2.8E-07 | 1E-06 | 0.5759581 | surface |
| MMP9 | 2.3491 | 9.4158 | 135.7 | 3.3E-07 | 2E-06 | 0.490488 | n.a. |
| AIF1 | 3.2092 | 9.4438 | 211.4 | 3.5E-07 | 2E-06 | 0.8486352 | n.a. |
| IL18 | 1.3748 | 9.3131 | 206.7 | 4.4E-07 | 2E-06 | 0.2795699 | n.a. |
| PYGL | 1.2475 | 9.2869 | 136.9 | 6.5E-07 | 3E-06 | 0.3363661 | n.a. |
| CEBPB | 2.4968 | 9.407 | 100.9 | 8.8E-07 | 4E-06 | 0.6765922 | n.a. |
| ENSECAG00000013303 | 1.5188 | 9.4002 | 55.1 | 1.1E-06 | 5E-06 | 0.6222774 | n.a. |
| FCGRT | 1.0611 | 9.3372 | 73.56 | 3.4E-06 | 1E-05 | 0.4579542 | surface |
| HEXB | 1.1954 | 9.3642 | 52.02 | 3.5E-06 | 1E-05 | 0.5720982 | n.a. |
| SLC7A7 | 1.0229 | 9.4046 | 26.91 | 0.00015 | 0.0005 | 0.6029777 | n.a. |
| IFI30 | 1.0364 | 9.9429 | 14.75 | 0.00026 | 0.0009 | 0.8461538 | n.a. |
| CD14 | 1.7096 | 9.3178 | 226.2 | 0.0003 | 0.001 | 0.826027 | surface |

| genes | logFC | logCPM | F | PValue | FDR | percent.exp | Surfacome.Label |
|---|---|---|---|---|---|---|---|
| FCER1A | 7.8218 | 9.4154 | 88086 | 0 | 0 | 1 | surface |
| LTC4S | 6.8902 | 9.3026 | 1E+05 | 0 | 0 | 0.9802817 | nonsurface |
| FCER1G | 5.6966 | 10.204 | 46538 | 0 | 0 | 1 | nonsurface |
| ENSECAG00000019875 | 5.4267 | 9.2924 | 30722 | 0 | 0 | 0.9070423 | n.a. |
| RHEX | 5.3002 | 9.2599 | 41773 | 0 | 0 | 0.8422535 | n.a. |
| GCSAML | 5.0467 | 9.253 | 80111 | 0 | 0 | 0.7323944 | n.a. |
| MS4A7 | 4.9216 | 9.3128 | 32071 | 0 | 0 | 0.828169 | n.a. |
| GATA2 | 4.806 | 9.2498 | 74374 | 0 | 0 | 0.7070423 | n.a. |
| GM2A | 4.801 | 9.9979 | 19119 | 0 | 0 | 0.8535211 | n.a. |
| C1H15orf48 | 4.7993 | 9.6929 | 14479 | 0 | 0 | 0.9492958 | n.a. |
| HRH4 | 4.5989 | 9.2474 | 78123 | 0 | 0 | 0.6591549 | surface |
| PRSS57 | 4.4855 | 9.2486 | 52360 | 0 | 0 | 0.5408451 | n.a. |
| MS4A2 | 4.4573 | 9.2465 | 68178 | 0 | 0 | 0.6478873 | n.a. |
| HS3ST1 | 4.4307 | 9.2857 | 19650 | 0 | 0 | 0.6929577 | n.a. |
| CMA1 | 4.3439 | 9.2596 | 26062 | 0 | 0 | 0.5098592 | n.a. |
| PIM1 | 4.2854 | 9.3916 | 4806 | 0 | 0 | 0.7746479 | n.a. |
| TGM3 | 4.2188 | 9.2449 | 52993 | 0 | 0 | 0.5859155 | n.a. |
| TSPO | 4.1611 | 9.7323 | 6970 | 0 | 0 | 0.9380282 | nonsurface |
| ENSECAG00000015137 | 4.0609 | 9.3925 | 12588 | 0 | 0 | 0.3239437 | n.a. |
| ENSECAG00000039088 | 4.0334 | 9.7274 | 4868 | 0 | 0 | 0.9633803 | n.a. |
| ENSECAG00000036100 | 3.9928 | 9.3021 | 25868 | 0 | 0 | 0.5887324 | n.a. |
| ALOX5AP | 3.892 | 9.2637 | 15089 | 0 | 0 | 0.5549296 | nonsurface |
| ITM2B | 3.8626 | 10.174 | 6013 | 0 | 0 | 1 | surface |
| LAP3 | 3.8102 | 9.4403 | 3211 | 0 | 0 | 0.771831 | n.a. |
| ENSECAG00000035094 | 3.7162 | 9.2863 | 9942 | 0 | 0 | 0.6676056 | n.a. |
| ENSECAG00000035834 | 3.6854 | 9.2418 | 34930 | 0 | 0 | 0.4732394 | n.a. |
| ENSECAG00000017717 | 3.6818 | 9.2422 | 34150 | 0 | 0 | 0.4760563 | n.a. |
| CD63 | 3.6179 | 9.4159 | 3988 | 0 | 0 | 0.7464789 | surface |
| SELL | 3.5674 | 9.6904 | 2772 | 0 | 0 | 0.8647887 | surface |
| TP53I11 | 3.5553 | 9.2424 | 34355 | 0 | 0 | 0.4169014 | nonsurface |
| SERPINB10 | 3.5485 | 9.5345 | 9564 | 0 | 0 | 0.5042254 | n.a. |
| ENSECAG00000033857 | 3.5014 | 9.3316 | 21529 | 0 | 0 | 0.5859155 | n.a. |
| eca-mir-223 | 3.4832 | 9.2778 | 16551 | 0 | 0 | 0.4957746 | n.a. |
| SRGN | 3.2705 | 10.485 | 4152 | 0 | 0 | 0.9802817 | n.a. |
| XBP1 | 3.1968 | 9.4868 | 1988 | 0 | 0 | 0.7183099 | n.a. |
| BSG | 3.0046 | 9.6757 | 1818 | 0 | 0 | 0.8901408 | surface |
| PTP4A2 | 2.9994 | 9.7459 | 1943 | 0 | 0 | 0.9126761 | n.a. |
| ENSECAG00000020136 | 2.9917 | 9.7014 | 1899 | 0 | 0 | 0.7746479 | n.a. |
| SYNE4 | 2.9418 | 10.39 | 4101 | 0 | 0 | 0.9014085 | nonsurface |
| PFN1 | 1.6652 | 11.941 | 1855 | 0 | 0 | 0.9971831 | n.a. |
| ENSECAG00000030745 | 1.5973 | 11.502 | 1795 | 0 | 0 | 0.9971831 | n.a. |
| ENSECAG00000040065 | 3.3213 | 9.2435 | 26468 | 0 | 0 | 0.3830986 | n.a. |
| SAT1 | 2.5463 | 9.7271 | 1491 | 0 | 2E-308 | 0.7464789 | n.a. |
| RGS18 | 3.0625 | 9.3114 | 4498 | 0 | 3E-308 | 0.4816901 | n.a. |

| | | | | | | | |
|---|---|---|---|---|---|---|---|
| CYBA | 2.0438 | 10.509 | 1435 | 1E-307 | 2E-305 | 0.9746479 | nonsurface |
| ENSECAG00000032253 | 3.2086 | 9.2484 | 15221 | 7E-302 | 9E-300 | 0.3859155 | n.a. |
| C4orf48 | 2.8243 | 9.5904 | 1384 | 4E-297 | 6E-295 | 0.7521127 | n.a. |
| SPINT2 | 3.2458 | 9.2841 | 4731 | 1E-294 | 2E-292 | 0.4788732 | surface |
| ENSECAG00000012768 | 3.3106 | 9.2463 | 12806 | 2E-292 | 2E-290 | 0.2985915 | n.a. |
| HACD4 | 2.924 | 9.3015 | 13947 | 3E-291 | 4E-289 | 0.3802817 | n.a. |
| ENSECAG00000032032 | 3.2597 | 9.2656 | 5980 | 3E-291 | 4E-289 | 0.484507 | n.a. |
| GNPTAB | 3.3792 | 9.2672 | 5052 | 2E-283 | 3E-281 | 0.4929577 | nonsurface |
| RUNX1 | 3.3901 | 9.2791 | 3814 | 3E-273 | 4E-271 | 0.5042254 | n.a. |
| GALM | 3.2926 | 9.2572 | 6936 | 7E-267 | 7E-265 | 0.3802817 | n.a. |
| ARHGDIB | 1.444 | 11.255 | 1214 | 1E-261 | 2E-259 | 0.9887324 | n.a. |
| COTL1 | 2.0644 | 10.529 | 1166 | 3E-251 | 3E-249 | 0.9352113 | n.a. |
| GCNT2 | 3.1837 | 9.2388 | 42394 | 3E-247 | 2E-245 | 0.3352113 | nonsurface |
| JAK2 | 3.0946 | 9.2772 | 4067 | 9E-244 | 8E-242 | 0.4591549 | n.a. |
| VASP | 2.746 | 9.4309 | 1272 | 2E-235 | 2E-233 | 0.5971831 | n.a. |
| DRAXIN | 2.9746 | 9.3226 | 3225 | 4E-235 | 3E-233 | 0.4422535 | n.a. |
| NFE2 | 2.6318 | 9.2974 | 11631 | 5E-234 | 5E-232 | 0.3380282 | n.a. |
| ITPRID2 | 3.0819 | 9.2614 | 5468 | 3E-233 | 3E-231 | 0.3971831 | n.a. |
| LY96 | 2.8335 | 9.3807 | 1568 | 3E-230 | 3E-228 | 0.4816901 | n.a. |
| ENSECAG00000016948 | 1.3659 | 11.316 | 1059 | 9E-229 | 8E-227 | 0.9971831 | n.a. |
| P2RY10 | 2.731 | 9.4226 | 1186 | 5E-219 | 4E-217 | 0.571831 | surface |
| ENSECAG00000000968 | 2.9485 | 9.2588 | 5524 | 7E-214 | 5E-212 | 0.3295775 | n.a. |
| GATA1 | 2.9444 | 9.2382 | 27884 | 2E-212 | 2E-210 | 0.3042254 | n.a. |
| BTK | 2.6462 | 9.3096 | 4295 | 8E-211 | 6E-209 | 0.3605634 | n.a. |
| ALPL | 2.6426 | 9.2457 | 8385 | 4E-210 | 3E-208 | 0.2985915 | surface |
| ENSECAG00000010232 | 2.9936 | 9.2384 | 24090 | 2E-208 | 1E-206 | 0.284507 | n.a. |
| P2RY14 | 2.5642 | 9.2475 | 11176 | 9E-208 | 6E-206 | 0.2591549 | surface |
| STAP1 | 2.6121 | 9.3165 | 5738 | 1E-207 | 7E-206 | 0.2957746 | n.a. |
| ATP6V1G1 | 2.0331 | 9.9335 | 940.4 | 9E-204 | 6E-202 | 0.8957746 | n.a. |
| DBI | 2.4029 | 9.673 | 931.1 | 8E-202 | 6E-200 | 0.5605634 | n.a. |
| ENSECAG00000013303 | 2.5941 | 9.4002 | 1546 | 2E-194 | 1E-192 | 0.5492958 | n.a. |
| LPCAT2 | 2.442 | 9.2572 | 9584 | 4E-194 | 3E-192 | 0.2591549 | nonsurface |
| NFAM1 | 2.5491 | 9.3582 | 4923 | 6E-194 | 4E-192 | 0.3408451 | surface |
| POC1A | 2.9035 | 9.2936 | 2489 | 4E-186 | 3E-184 | 0.3521127 | n.a. |
| TGFB1 | 2.7039 | 9.3709 | 1333 | 4E-183 | 3E-181 | 0.4788732 | n.a. |
| ALOX5 | 2.6676 | 9.2381 | 25105 | 3E-181 | 2E-179 | 0.2676056 | nonsurface |
| RAB37 | 2.6819 | 9.2869 | 2859 | 9E-179 | 5E-177 | 0.3408451 | n.a. |
| TSTD3 | 2.5685 | 9.4008 | 1646 | 7E-172 | 4E-170 | 0.4591549 | n.a. |
| SAMSN1 | 2.5286 | 9.3692 | 1040 | 2E-165 | 1E-163 | 0.4788732 | n.a. |
| TALDO1 | 1.9889 | 9.9099 | 744.9 | 3E-162 | 2E-160 | 0.7943662 | n.a. |
| ENSECAG00000015753 | 2.6448 | 9.2621 | 3785 | 2E-157 | 1E-155 | 0.2760563 | n.a. |
| ENSECAG00000031913 | 2.4086 | 9.2438 | 8879 | 3E-156 | 2E-154 | 0.2732394 | n.a. |
| MYL6 | 1.4878 | 10.731 | 709.1 | 1E-154 | 6E-153 | 0.9521127 | n.a. |
| SH3BGRL3 | 1.4523 | 10.736 | 704.5 | 1E-153 | 6E-152 | 0.9633803 | n.a. |
| ENSECAG00000037706 | 1.6399 | 10.368 | 697.6 | 3E-152 | 2E-150 | 0.9239437 | n.a. |

| | | | | | | | |
|---|---|---|---|---|---|---|---|
| CFL1 | 1.273 | 11.079 | 688.5 | 3E-150 | 1E-148 | 0.9746479 | n.a. |
| HSBP1 | 2.2536 | 9.5012 | 862.6 | 4E-146 | 2E-144 | 0.5464789 | n.a. |
| C5AR1 | 1.9955 | 9.3446 | 5792 | 2E-143 | 9E-142 | 0.2676056 | surface |
| TNFSF14 | 2.453 | 9.28 | 2816 | 4E-142 | 2E-140 | 0.2676056 | n.a. |
| SPI1 | 1.8346 | 9.6618 | 1837 | 1E-138 | 5E-137 | 0.4253521 | n.a. |
| EID1 | 2.4595 | 9.4085 | 789.5 | 1E-137 | 7E-136 | 0.484507 | n.a. |
| YPEL5 | 2.1593 | 9.4914 | 634.4 | 5E-136 | 2E-134 | 0.5323944 | n.a. |
| ENSECAG00000039232 | 1.0339 | 11.162 | 612.3 | 5E-134 | 2E-132 | 0.9887324 | n.a. |
| ENSECAG00000000910 | 2.16 | 9.7673 | 598.7 | 4E-131 | 2E-129 | 0.5915493 | n.a. |
| CARHSP1 | 2.2412 | 9.4061 | 756.2 | 6E-131 | 3E-129 | 0.4591549 | n.a. |
| SH3KBP1 | 2.2899 | 9.4466 | 669.2 | 9E-128 | 4E-126 | 0.4732394 | n.a. |
| LAT | 2.1872 | 9.5366 | 737.8 | 5E-127 | 2E-125 | 0.428169 | nonsurface |
| ENSECAG00000032131 | 2.3722 | 9.3494 | 951.2 | 5E-127 | 2E-125 | 0.428169 | n.a. |
| MMP9 | 1.9873 | 9.4158 | 2456 | 2E-126 | 8E-125 | 0.2507042 | n.a. |
| LIMD2 | 1.7629 | 10.143 | 565 | 7E-124 | 3E-122 | 0.8450704 | n.a. |
| VAT1 | 2.5075 | 9.2569 | 3197 | 2E-122 | 1E-120 | 0.2704225 | n.a. |
| NT5C | 2.4219 | 9.3389 | 914.4 | 2E-118 | 8E-117 | 0.3774648 | n.a. |
| PRDX1 | 1.9504 | 9.8857 | 531.5 | 1E-116 | 4E-115 | 0.6450704 | n.a. |
| LSP1 | 1.4832 | 10.243 | 526 | 2E-115 | 6E-114 | 0.9042254 | n.a. |
| NPDC1 | 2.1304 | 9.5117 | 551.7 | 1E-113 | 6E-112 | 0.4338028 | n.a. |
| CYSLTR1 | 2.2149 | 9.3142 | 1474 | 3E-112 | 1E-110 | 0.2732394 | surface |
| TMEM14C | 1.8868 | 9.6674 | 508.1 | 1E-111 | 4E-110 | 0.6450704 | nonsurface |
| LDHA | 1.9213 | 9.6788 | 513.3 | 6E-111 | 2E-109 | 0.5577465 | n.a. |
| IGFLR1 | 2.1888 | 9.4383 | 713.3 | 3E-107 | 1E-105 | 0.3774648 | surface |
| RNF130 | 2.1151 | 9.3597 | 1253 | 6E-104 | 2E-102 | 0.3577465 | surface |
| JPT1 | 1.786 | 9.7243 | 469.9 | 2E-103 | 6E-102 | 0.628169 | n.a. |
| RNF11 | 2.2915 | 9.3246 | 984.4 | 2E-103 | 8E-102 | 0.3380282 | n.a. |
| CACYBP | 1.9638 | 9.6062 | 459.7 | 3E-101 | 9E-100 | 0.5605634 | n.a. |
| REXO2 | 2.14 | 9.4557 | 764.7 | 5E-101 | 2E-99 | 0.4112676 | n.a. |
| NFKBIA | 1.8908 | 9.5431 | 457.9 | 6E-101 | 2E-99 | 0.4760563 | n.a. |
| GRB2 | 1.7259 | 9.7762 | 453.3 | 6E-100 | 2E-98 | 0.684507 | n.a. |
| MKNK1 | 2.2792 | 9.2658 | 1988 | 1E-96 | 4E-95 | 0.2535211 | n.a. |
| RALGAPA2 | 2.273 | 9.3573 | 840.8 | 1.7E-96 | 6E-95 | 0.3014085 | nonsurface |
| FTH1 | 1.2327 | 10.516 | 418.8 | 1.6E-92 | 5E-91 | 0.8084507 | nonsurface |
| CD82 | 1.9321 | 9.4652 | 533.8 | 2E-92 | 7E-91 | 0.3746479 | surface |
| CSRP1 | 2.096 | 9.2812 | 1899 | 2.7E-92 | 9E-91 | 0.2647887 | n.a. |
| ARRB2 | 2.0063 | 9.4662 | 517.7 | 7.6E-92 | 3E-90 | 0.4253521 | n.a. |
| NAGK | 1.9904 | 9.431 | 824 | 1.3E-90 | 4E-89 | 0.3605634 | n.a. |
| CREM | 2.2105 | 9.3352 | 739.7 | 1.6E-90 | 5E-89 | 0.3549296 | n.a. |
| TFEB | 1.822 | 9.3544 | 1558 | 4.9E-88 | 2E-86 | 0.2619718 | n.a. |
| ORAI3 | 2.1412 | 9.3598 | 617.7 | 3.1E-86 | 1E-84 | 0.3633803 | n.a. |
| ARPC2 | 1.1538 | 10.591 | 381.3 | 1.8E-84 | 6E-83 | 0.9464789 | n.a. |
| SYNE1 | 2.1082 | 9.3384 | 675.2 | 3.8E-84 | 1E-82 | 0.3352113 | nonsurface |
| SERPINB6 | 1.9001 | 9.5154 | 468.3 | 4.6E-82 | 1E-80 | 0.315493 | n.a. |
| MYL12A | 1.1959 | 10.506 | 359.9 | 7.4E-80 | 2E-78 | 0.9042254 | n.a. |

| | | | | | | | |
|---|---|---|---|---|---|---|---|
| S100A11 | 1.5822 | 9.847 | 359.1 | 1.1E-79 | 3E-78 | 0.4957746 | n.a. |
| CDK2AP2 | 1.6348 | 9.6423 | 356.2 | 4.7E-79 | 1E-77 | 0.5295775 | n.a. |
| TMEM59 | 1.642 | 9.5994 | 352.2 | 3.4E-78 | 1E-76 | 0.5633803 | nonsurface |
| HERPUD2 | 2.0026 | 9.3424 | 578.1 | 4.5E-78 | 1E-76 | 0.3267606 | nonsurface |
| ENSECAG00000016543 | 1.6981 | 9.8125 | 350.5 | 8E-78 | 2E-76 | 0.6197183 | n.a. |
| PPT1 | 1.8976 | 9.4371 | 566.9 | 7.8E-77 | 2E-75 | 0.3774648 | n.a. |
| RASSF3 | 1.959 | 9.3241 | 710.5 | 1.8E-76 | 5E-75 | 0.2873239 | n.a. |
| NDUFA1 | 1.7988 | 9.4956 | 419.7 | 4.2E-75 | 1E-73 | 0.4732394 | nonsurface |
| HCST | 1.5639 | 9.5985 | 352.1 | 1.2E-74 | 3E-73 | 0.4478873 | n.a. |
| GAPDH | 1.1437 | 10.705 | 332.5 | 6.2E-74 | 2E-72 | 0.8816901 | n.a. |
| ATP2A3 | 1.7892 | 9.4984 | 354.1 | 1.9E-73 | 5E-72 | 0.3971831 | n.a. |
| PCBP1 | 1.5337 | 9.785 | 324.4 | 3.4E-72 | 1E-70 | 0.6169014 | n.a. |
| MYH9 | 1.7076 | 9.5442 | 325.6 | 4.8E-72 | 1E-70 | 0.4619718 | n.a. |
| CLIC1 | 1.3547 | 10.07 | 321.2 | 1.7E-71 | 5E-70 | 0.7492958 | n.a. |
| RF01956 | 1.901 | 9.2926 | 1505 | 1.6E-69 | 4E-68 | 0.4535211 | n.a. |
| ADGRE5 | 1.6134 | 9.5791 | 311.5 | 2E-69 | 6E-68 | 0.4788732 | n.a. |
| ARPC3 | 1.3549 | 10.061 | 310.7 | 3E-69 | 8E-68 | 0.7323944 | n.a. |
| MIF | 1.348 | 10.103 | 310.5 | 3.4E-69 | 9E-68 | 0.7323944 | n.a. |
| HK1 | 1.9504 | 9.3001 | 842.1 | 1.4E-68 | 4E-67 | 0.2535211 | n.a. |
| SURF4 | 1.957 | 9.3205 | 656.1 | 2.6E-68 | 7E-67 | 0.2676056 | nonsurface |
| PTPRCAP | 1.1866 | 10.617 | 299.7 | 7.2E-67 | 2E-65 | 0.771831 | nonsurface |
| AGPAT2 | 1.7791 | 9.4052 | 446.5 | 8.6E-66 | 2E-64 | 0.3633803 | nonsurface |
| CYB5R4 | 1.9101 | 9.3322 | 573.2 | 3.4E-65 | 9E-64 | 0.2957746 | n.a. |
| UCP2 | 1.3348 | 9.9052 | 291.5 | 6.7E-65 | 2E-63 | 0.6619718 | nonsurface |
| IDO1 | 1.6255 | 9.5783 | 352.4 | 8.1E-65 | 2E-63 | 0.2985915 | n.a. |
| EVI2B | 1.5363 | 9.5277 | 273.3 | 3.8E-61 | 1E-59 | 0.4507042 | surface |
| GUCD1 | 1.9528 | 9.3108 | 602.7 | 1E-60 | 2E-59 | 0.256338 | n.a. |
| SSR3 | 1.3228 | 9.9724 | 269.6 | 2.3E-60 | 6E-59 | 0.7070423 | nonsurface |
| ENSECAG00000019392 | 1.4294 | 9.7659 | 268.7 | 3.7E-60 | 9E-59 | 0.6394366 | n.a. |
| P4HB | 1.7927 | 9.4063 | 450.1 | 5.6E-60 | 1E-58 | 0.3323944 | n.a. |
| ARL6IP6 | 1.8605 | 9.3181 | 543.6 | 6.6E-60 | 2E-58 | 0.2507042 | nonsurface |
| GABARAP | 1.0112 | 10.489 | 261.9 | 1.1E-58 | 3E-57 | 0.9042254 | n.a. |
| VIM | 1.0763 | 11.286 | 261.4 | 1.4E-58 | 3E-57 | 0.8169014 | nonsurface |
| EHD1 | 1.6633 | 9.4259 | 319.3 | 2E-57 | 5E-56 | 0.3408451 | n.a. |
| HMGB2 | 1.2679 | 10.129 | 253.4 | 7.7E-57 | 2E-55 | 0.7633803 | n.a. |
| ALDOA | 1.251 | 9.99 | 249.8 | 4.4E-56 | 1E-54 | 0.6225352 | n.a. |
| CAPG | 1.5298 | 9.6557 | 248.4 | 3.2E-55 | 7E-54 | 0.4169014 | n.a. |
| CISD2 | 1.5079 | 9.5388 | 242.5 | 1.7E-54 | 4E-53 | 0.4478873 | nonsurface |
| PRELID1 | 1.3736 | 9.7488 | 241.9 | 2.3E-54 | 5E-53 | 0.5464789 | n.a. |
| MICU2 | 1.6587 | 9.4141 | 311 | 1.9E-53 | 4E-52 | 0.3352113 | n.a. |
| RTN3 | 1.7097 | 9.3937 | 366.9 | 3.4E-53 | 8E-52 | 0.315493 | n.a. |
| TMCO1 | 1.6163 | 9.4392 | 288.4 | 8.1E-53 | 2E-51 | 0.3577465 | nonsurface |
| PTPN6 | 1.4374 | 9.58 | 234.6 | 7.6E-51 | 2E-49 | 0.4 | n.a. |
| AP3S1 | 1.4615 | 9.534 | 228.9 | 2.6E-50 | 6E-49 | 0.4338028 | n.a. |
| ARPC5 | 1.2961 | 9.8016 | 214.8 | 1.7E-48 | 4E-47 | 0.5802817 | n.a. |

| | | | | | | | |
|---|---|---|---|---|---|---|---|
| MIR142 | 1.5734 | 9.4165 | 267.5 | 2.9E-48 | 6E-47 | 0.3661972 | n.a. |
| OS9 | 1.567 | 9.3847 | 279.3 | 5.4E-48 | 1E-46 | 0.2732394 | n.a. |
| SF3B6 | 1.4188 | 9.6154 | 209.8 | 2.1E-47 | 4E-46 | 0.4676056 | n.a. |
| UBE2E3 | 1.5402 | 9.3915 | 372.6 | 4.4E-47 | 9E-46 | 0.2929577 | n.a. |
| RGS19 | 1.4373 | 9.5283 | 234.3 | 4.7E-47 | 1E-45 | 0.3887324 | n.a. |
| ARRDC1 | 1.6857 | 9.3734 | 349 | 5.2E-47 | 1E-45 | 0.2957746 | n.a. |
| LAPTM4A | 1.3475 | 9.5901 | 207 | 8.3E-47 | 2E-45 | 0.4647887 | nonsurface |
| GUK1 | 1.4112 | 9.5821 | 204.7 | 2.7E-46 | 6E-45 | 0.4169014 | n.a. |
| PLP2 | 1.1739 | 9.9939 | 203.8 | 4.1E-46 | 8E-45 | 0.6732394 | n.a. |
| RAB24 | 1.4489 | 9.4289 | 268.4 | 4.8E-45 | 1E-43 | 0.315493 | n.a. |
| YIF1B | 1.6998 | 9.3422 | 414.7 | 3.2E-44 | 6E-43 | 0.256338 | nonsurface |
| TPI1 | 1.415 | 9.5703 | 207.2 | 7.6E-44 | 2E-42 | 0.3887324 | n.a. |
| MXD1 | 1.5743 | 9.3408 | 356.7 | 9.9E-44 | 2E-42 | 0.2704225 | n.a. |
| RABAC1 | 1.1856 | 9.8034 | 191.1 | 2.4E-43 | 5E-42 | 0.6084507 | nonsurface |
| SUB1 | 1.0316 | 10.333 | 190.3 | 3.5E-43 | 7E-42 | 0.7605634 | n.a. |
| GPR183 | 1.2954 | 9.6621 | 190.2 | 5.8E-43 | 1E-41 | 0.315493 | surface |
| MISP3 | 1.4105 | 9.507 | 194.1 | 2.2E-42 | 4E-41 | 0.3690141 | n.a. |
| AIP | 1.4535 | 9.5081 | 200 | 6.1E-42 | 1E-40 | 0.371831 | n.a. |
| SPCS3 | 1.264 | 9.6267 | 183 | 2.2E-41 | 4E-40 | 0.4591549 | nonsurface |
| METTL9 | 1.3989 | 9.507 | 184.3 | 9.1E-41 | 2E-39 | 0.3549296 | n.a. |
| GABARAPL2 | 1.2531 | 9.7938 | 178.9 | 1.1E-40 | 2E-39 | 0.5492958 | n.a. |
| CERS2 | 1.5084 | 9.3895 | 255.4 | 1.2E-40 | 2E-39 | 0.2732394 | nonsurface |
| ZFAND5 | 1.5298 | 9.3766 | 268.7 | 1.7E-40 | 3E-39 | 0.3126761 | n.a. |
| RAP1B | 1.2901 | 9.5901 | 177.2 | 4E-40 | 8E-39 | 0.4338028 | n.a. |
| STK10 | 1.5061 | 9.392 | 274.8 | 9.9E-40 | 2E-38 | 0.2732394 | n.a. |
| NDUFB7 | 1.2122 | 9.7474 | 174.4 | 1E-39 | 2E-38 | 0.5802817 | n.a. |
| APOPT1 | 1.5954 | 9.3647 | 305.1 | 1.6E-39 | 3E-38 | 0.2788732 | n.a. |
| LEPROTL1 | 1.3466 | 9.565 | 172.4 | 6.7E-39 | 1E-37 | 0.3887324 | nonsurface |
| TMEM60 | 1.4807 | 9.4303 | 207.8 | 1.8E-38 | 3E-37 | 0.3014085 | n.a. |
| ENSECAG00000020462 | 1.2039 | 9.6751 | 174.2 | 5.9E-38 | 1E-36 | 0.4929577 | n.a. |
| IRF2BP2 | 1.442 | 9.4142 | 287.7 | 1.1E-37 | 2E-36 | 0.2591549 | n.a. |
| NAA38 | 1.5767 | 9.3687 | 293.9 | 1.3E-37 | 2E-36 | 0.2816901 | n.a. |
| ARHGAP15 | 1.2869 | 9.6767 | 161.7 | 5.8E-37 | 1E-35 | 0.4450704 | n.a. |
| TOP1 | 1.4579 | 9.4172 | 220.8 | 1.1E-36 | 2E-35 | 0.2816901 | n.a. |
| ERLEC1 | 1.4577 | 9.3911 | 227.1 | 2E-36 | 4E-35 | 0.2732394 | n.a. |
| TBC1D10C | 1.3998 | 9.4778 | 185.9 | 2.2E-36 | 4E-35 | 0.3070423 | n.a. |
| OSTF1 | 1.0362 | 10.056 | 159 | 2.3E-36 | 4E-35 | 0.656338 | n.a. |
| SEPT7 | 1.193 | 9.8038 | 156.8 | 6.8E-36 | 1E-34 | 0.5183099 | n.a. |
| PRDX5 | 1.3381 | 9.5264 | 203.7 | 7.3E-34 | 1E-32 | 0.3690141 | n.a. |
| ENSECAG00000016499 | 1.3002 | 9.4863 | 177.6 | 1.3E-33 | 2E-32 | 0.3521127 | n.a. |
| CYTIP | 1.1744 | 9.6111 | 145.3 | 2.1E-33 | 4E-32 | 0.3943662 | n.a. |
| CSK | 1.4118 | 9.389 | 215 | 6.7E-33 | 1E-31 | 0.2732394 | n.a. |
| RTF2 | 1.3582 | 9.4173 | 196.6 | 8.1E-33 | 1E-31 | 0.2788732 | n.a. |
| ARL6IP1 | 1.1859 | 9.5522 | 138.8 | 5.7E-32 | 9E-31 | 0.4 | nonsurface |
| NPC2 | 1.0418 | 9.8367 | 143.3 | 1.5E-31 | 2E-30 | 0.4732394 | n.a. |

| | | | | | | | |
|---|---|---|---|---|---|---|---|
| TLN1 | 1.2235 | 9.5187 | 150.7 | 2.2E-31 | 4E-30 | 0.3464789 | n.a. |
| SARAF | 1.0868 | 9.8115 | 135.2 | 3.5E-31 | 6E-30 | 0.5408451 | nonsurface |
| TMEM71 | 1.3634 | 9.4245 | 178 | 7.2E-31 | 1E-29 | 0.2507042 | nonsurface |
| MED28 | 1.3799 | 9.3945 | 195.6 | 9.6E-31 | 2E-29 | 0.256338 | n.a. |
| CDC42SE1 | 1.1401 | 9.6142 | 131.8 | 1.9E-30 | 3E-29 | 0.4676056 | n.a. |
| FAM107B | 1.2278 | 9.5005 | 143.5 | 8.5E-30 | 1E-28 | 0.3014085 | n.a. |
| NDUFS2 | 1.2317 | 9.5195 | 148.2 | 9.2E-30 | 1E-28 | 0.3521127 | n.a. |
| GSTO1 | 1.3155 | 9.4524 | 210.1 | 2.1E-29 | 3E-28 | 0.2647887 | n.a. |
| TMBIM6 | 1.0455 | 9.7742 | 126.1 | 3.2E-29 | 5E-28 | 0.4929577 | nonsurface |
| ANXA6 | 1.232 | 9.5613 | 137 | 1.9E-28 | 3E-27 | 0.3267606 | n.a. |
| PGAM1 | 1.1916 | 9.5435 | 135.3 | 3.3E-28 | 5E-27 | 0.3183099 | n.a. |
| LASP1 | 1.2588 | 9.4194 | 172 | 6E-28 | 9E-27 | 0.2732394 | n.a. |
| SDHA | 1.3259 | 9.4236 | 166.3 | 1.1E-27 | 2E-26 | 0.2591549 | n.a. |
| SRSF5 | 1.0294 | 9.8174 | 118.8 | 1.3E-27 | 2E-26 | 0.5464789 | n.a. |
| ICAM3 | 1.122 | 9.5641 | 123.6 | 1.5E-27 | 2E-26 | 0.3323944 | surface |
| UBE2N | 1.2166 | 9.5201 | 131.5 | 4.8E-27 | 7E-26 | 0.3295775 | n.a. |
| WDR1 | 1.0437 | 9.7464 | 120.1 | 8.2E-27 | 1E-25 | 0.4676056 | nonsurface |
| LCP1 | 1.0247 | 9.7068 | 112.3 | 3.4E-26 | 5E-25 | 0.4647887 | n.a. |
| ACAP1 | 1.1604 | 9.5179 | 124.4 | 3.7E-26 | 5E-25 | 0.2619718 | n.a. |
| TMED2 | 1.0219 | 9.6321 | 107.9 | 3.1E-25 | 4E-24 | 0.4084507 | nonsurface |
| SFT2D1 | 1.1364 | 9.5206 | 114.7 | 9E-25 | 1E-23 | 0.3183099 | nonsurface |
| TMEM50A | 1.0337 | 9.7377 | 105 | 1.3E-24 | 2E-23 | 0.4422535 | nonsurface |
| ACTR2 | 1.0787 | 9.4911 | 115.4 | 1.9E-23 | 3E-22 | 0.3126761 | n.a. |
| WARS | 1.0283 | 9.8298 | 95.98 | 1.2E-22 | 2E-21 | 0.4197183 | n.a. |
| PPP1R12A | 1.119 | 9.4355 | 115.7 | 5.1E-22 | 7E-21 | 0.2760563 | n.a. |
| CHMP2A | 1.0512 | 9.5304 | 103.9 | 6.5E-22 | 8E-21 | 0.3239437 | n.a. |
| CHCHD7 | 1.1702 | 9.4135 | 133.1 | 1E-21 | 1E-20 | 0.2704225 | n.a. |
| PDLIM2 | 1.0725 | 9.5617 | 98.1 | 1.7E-21 | 2E-20 | 0.2929577 | n.a. |
| HHEX | 1.0582 | 9.4401 | 141.1 | 4.2E-21 | 5E-20 | 0.2816901 | n.a. |
| PGK1 | 1.1259 | 9.4881 | 127.8 | 5.1E-21 | 6E-20 | 0.2732394 | n.a. |
| HDAC1 | 1.1229 | 9.4204 | 118 | 5.8E-21 | 7E-20 | 0.2507042 | n.a. |
| RAB1A | 1.0878 | 9.4321 | 114.7 | 6E-20 | 7E-19 | 0.2647887 | n.a. |
| TSC22D4 | 1.1037 | 9.4273 | 108.6 | 1.1E-19 | 1E-18 | 0.2507042 | n.a. |
| PRKAR1A | 1.0581 | 9.4477 | 98.33 | 1.9E-19 | 2E-18 | 0.2507042 | n.a. |
| TMEM179B | 1.0119 | 9.5307 | 91.58 | 4.8E-19 | 6E-18 | 0.3267606 | surface |
| PAFAH1B3 | 1.0071 | 9.5563 | 85.47 | 1.1E-18 | 1E-17 | 0.2788732 | n.a. |
| HCLS1 | 1.0431 | 9.4497 | 103.8 | 5.2E-18 | 6E-17 | 0.2760563 | n.a. |
| KRCC1 | 1.003 | 9.4723 | 96.96 | 1.9E-17 | 2E-16 | 0.2619718 | n.a. |
| ENSECAG00000002541 | 1 | 9.4558 | 87.73 | 3.3E-17 | 4E-16 | 0.2647887 | n.a. |
