## Supplementary material for "Single cell resolution landscape of equine peripheral blood mononuclear cells reveals diverse immune cell subtypes including T-bet^+^ B cells": Dataset S2

| genes | logFC | logCPM | F | PValue | FDR | percent.exp | Surfacome.Label |
|---|---|---|---|---|---|---|---|
| ENSECAG00000000436 | 3.979 | 9.9635 | 13791.7 | 0 | 0 | 0.998997 | n.a. |
| S100A12 | 3.971 | 10.309 | 9234.12 | 0 | 0 | 1 | n.a. |
| LYZ | 3.868 | 11.749 | 9837.96 | 0 | 0 | 1 | n.a. |
| SERPINB10 | 3.273 | 9.5345 | 5966.67 | 0 | 0 | 0.9809428 | n.a. |
| VCAN | 2.889 | 9.3382 | 4815.53 | 0 | 0 | 0.8615848 | n.a. |
| S100A4 | 2.882 | 11.292 | 1912.49 | 0 | 0 | 1 | n.a. |
| THBS1 | 2.789 | 9.3512 | 4465.29 | 0 | 0 | 0.7703109 | n.a. |
| ENSECAG00000024882 | 2.761 | 9.349 | 4224.44 | 0 | 0 | 0.781344 | n.a. |
| PLBD1 | 2.665 | 9.972 | 4282.97 | 0 | 0 | 1 | n.a. |
| S100P | 2.653 | 9.8205 | 7235.55 | 0 | 0 | 1 | n.a. |
| SLPI | 2.519 | 9.3697 | 4241.63 | 0 | 0 | 0.8796389 | n.a. |
| TREM1 | 2.342 | 9.4852 | 6255.79 | 0 | 0 | 0.9578736 | surface |
| SULT1C4 | 2.339 | 9.421 | 4873.76 | 0 | 0 | 0.9538616 | n.a. |
| ENSECAG00000010117 | 2.304 | 9.501 | 5506.1 | 0 | 0 | 0.5416249 | n.a. |
| CTSZ | 2.263 | 9.8631 | 2067.8 | 0 | 0 | 0.998997 | n.a. |
| C1H15orf48 | 2.211 | 9.6929 | 1914.56 | 0 | 0 | 0.9889669 | n.a. |
| MMP9 | 2.187 | 9.4158 | 2829.22 | 0 | 0 | 0.664995 | n.a. |
| GM2A | 2.186 | 9.9979 | 2943.84 | 0 | 0 | 0.998997 | n.a. |
| ECATH-3 | 2.158 | 9.3365 | 5943.08 | 2E-283 | 1E-280 | 0.7873621 | n.a. |
| ENSECAG00000030387 | 2.523 | 9.433 | 1285.24 | 2E-276 | 8E-274 | 0.8655968 | n.a. |
| S100A6 | 2.559 | 10.437 | 1165.19 | 3E-251 | 1E-248 | 0.998997 | n.a. |
| TSPO | 1.747 | 9.7323 | 1146.45 | 3E-247 | 1E-244 | 0.997994 | n.a. |
| ENSECAG00000024886 | 2.545 | 9.319 | 4768.23 | 2E-245 | 7E-243 | 0.8345035 | n.a. |
| RNASE6 | 1.315 | 10.205 | 1127.33 | 3E-243 | 1E-240 | 1 | n.a. |
| S100A8 | 1.563 | 9.3765 | 4340.02 | 9E-239 | 3E-236 | 0.3861585 | n.a. |
| RNASE4 | 1.248 | 10.101 | 1074.97 | 3E-232 | 1E-229 | 1 | n.a. |
| MGST1 | 1.908 | 9.3352 | 2318.43 | 2E-207 | 4E-205 | 0.8214644 | n.a. |
| IFI27 | 2.233 | 9.7314 | 869.67 | 8E-189 | 2E-186 | 0.891675 | n.a. |
| LAMP1 | 1.542 | 9.6692 | 805.729 | 3E-175 | 8E-173 | 0.9829488 | surface |
| S100A5 | 2.018 | 9.5834 | 706.04 | 5E-154 | 1E-151 | 0.8134403 | n.a. |
| VIM | 1.042 | 11.286 | 692.343 | 4E-151 | 1E-148 | 0.998997 | n.a. |
| CYBB | 1.36 | 9.4653 | 796.111 | 2E-149 | 3E-147 | 0.9197593 | surface |
| ALDH2 | 1.36 | 9.5616 | 784.952 | 4E-149 | 8E-147 | 0.9588766 | n.a. |
| NFE2 | 1.697 | 9.2974 | 2962.28 | 6E-149 | 1E-146 | 0.7021063 | n.a. |
| LGALS3 | 1.499 | 9.5239 | 677.68 | 6E-148 | 1E-145 | 0.8986961 | n.a. |
| MECR | 1.949 | 9.3421 | 974.308 | 1E-142 | 2E-140 | 0.8054162 | n.a. |
| ENSECAG00000022247 | 1.837 | 9.2794 | 3576.96 | 8E-139 | 2E-136 | 0.6228686 | n.a. |
| EMP1 | 1.981 | 9.2873 | 2545.14 | 1E-126 | 2E-124 | 0.5827482 | surface |
| C5AR1 | 1.01 | 9.3446 | 2096.72 | 2E-123 | 4E-121 | 0.7552658 | surface |
| QPCT | 1.585 | 9.3093 | 1304.55 | 7E-121 | 1E-118 | 0.7352056 | n.a. |
| CD163 | 1.88 | 9.2595 | 2306.08 | 5E-106 | 9E-104 | 0.446339 | surface |
| ENSECAG00000003345 | 1.292 | 10.17 | 478.333 | 3E-105 | 4E-103 | 0.997994 | n.a. |
| CREG1 | 1.378 | 9.3229 | 1746.89 | 2E-101 | 4E-99 | 0.7673019 | n.a. |
| S100A10 | 1.317 | 10.866 | 454.581 | 3E-100 | 5.2E-98 | 0.998997 | n.a. |
| MHCB3 | 1.618 | 10.265 | 454.22 | 4E-100 | 6E-98 | 0.9187563 | n.a. |
| ENSECAG00000037539 | 1.185 | 9.2665 | 3018.94 | 2E-98 | 3E-96 | 0.4202608 | n.a. |
| PSAP | 1.128 | 10.167 | 444.922 | 4E-98 | 5.4E-96 | 1 | n.a. |
| GSN | 1.131 | 9.3636 | 793.171 | 2E-97 | 3.3E-95 | 0.8686058 | n.a. |
| FGL2 | 1.17 | 9.488 | 440.764 | 3E-97 | 4E-95 | 0.9498495 | n.a. |
| ENSECAG00000019309 | 1.495 | 9.3991 | 488.811 | 1E-96 | 1.6E-94 | 0.7542628 | n.a. |
| BLVRB | 1.391 | 9.4497 | 417.998 | 2E-92 | 2.9E-90 | 0.9358074 | n.a. |
| ENSECAG00000031156 | 1.425 | 9.3403 | 822.269 | 1E-87 | 1.4E-85 | 0.7331996 | n.a. |
| ENSECAG00000020136 | 1.281 | 9.7014 | 390.21 | 2E-86 | 2.6E-84 | 0.9558676 | n.a. |
| DPYD | 1.331 | 9.2972 | 1124.58 | 8E-84 | 8.9E-82 | 0.6539619 | n.a. |
| HNMT | 1.649 | 9.2852 | 2466.41 | 2E-83 | 2.6E-81 | 0.6048144 | n.a. |

| | | | | | | | |
|---|---|---|---|---|---|---|---|
| CXCL14 | 1.484 | 9.2572 | 2342.5 | 2E-80 | 1.9E-78 | 0.4182548 | n.a. |
| ENSECAG00000040180 | 1.406 | 9.7881 | 349.5 | 1E-77 | 1.3E-75 | 0.8806419 | n.a. |
| PLD3 | 1.464 | 9.3271 | 543.623 | 4E-76 | 3.5E-74 | 0.5416249 | n.a. |
| LYPLA1 | 1.503 | 9.3462 | 397.19 | 7E-75 | 6.5E-73 | 0.7251755 | n.a. |
| ENSECAG00000029581 | 1.427 | 9.2817 | 1849.77 | 4E-74 | 4.1E-72 | 0.552658 | n.a. |
| AHNAK | 1.432 | 9.5205 | 313.055 | 1E-69 | 8.5E-68 | 0.8585757 | n.a. |
| HK3 | 1.268 | 9.284 | 1371.53 | 7E-69 | 6.4E-67 | 0.668004 | n.a. |
| ALDH1A1 | 1.348 | 9.2588 | 2430.25 | 3E-67 | 2.6E-65 | 0.3259779 | n.a. |
| NAPSA | 1.011 | 9.5255 | 387.132 | 7E-65 | 5.3E-63 | 0.9689067 | n.a. |
| GIMAP7 | 1.29 | 10.513 | 290.063 | 9E-65 | 7E-63 | 0.9087262 | n.a. |
| TIMP2 | 1.287 | 9.2993 | 904.708 | 1E-62 | 8.2E-61 | 0.6910732 | n.a. |
| ENSECAG00000008826 | 1.256 | 9.376 | 290.704 | 2E-62 | 1.6E-60 | 0.7342026 | n.a. |
| CLEC5A | 1.281 | 9.2656 | 1669.77 | 1E-57 | 9E-56 | 0.4734203 | surface |
| SMS | 1.295 | 9.4498 | 256.478 | 2E-57 | 1.1E-55 | 0.889669 | n.a. |
| HACD4 | 1.175 | 9.3015 | 1447.57 | 7E-56 | 4.7E-54 | 0.663992 | n.a. |
| CD36 | 1.016 | 9.2648 | 1095.26 | 7E-55 | 4.6E-53 | 0.3450351 | surface |
| S100A3 | 1.079 | 9.3159 | 506.052 | 1E-54 | 7.8E-53 | 0.3660983 | n.a. |
| ALDH3A1 | 1.016 | 9.2538 | 2319.14 | 2E-54 | 9.6E-53 | 0.2617854 | n.a. |
| CD14 | 1.107 | 9.3178 | 1680.11 | 9E-52 | 5E-50 | 0.9398195 | surface |
| MNDA | 1.145 | 10.258 | 229.738 | 1E-51 | 5.7E-50 | 0.9829488 | n.a. |
| TMEM150B | 1.209 | 9.2642 | 1340.87 | 9E-51 | 5.1E-49 | 0.3851555 | surface |
| HEBP1 | 1.026 | 9.4026 | 215.159 | 1E-48 | 7.7E-47 | 0.8164493 | n.a. |
| ARL5A | 1.229 | 9.344 | 281.556 | 8E-47 | 3.9E-45 | 0.6389168 | n.a. |
| ENSECAG00000029716 | 1.098 | 9.8771 | 203.983 | 4E-46 | 1.9E-44 | 0.8485456 | n.a. |
| LDHB | 1.065 | 10.019 | 197.381 | 1E-44 | 5E-43 | 0.9849549 | n.a. |
| ENSECAG00000007681 | 1.022 | 9.6603 | 191.835 | 2E-43 | 7.9E-42 | 0.7342026 | n.a. |
| TSPAN4 | 1.072 | 9.2911 | 635.305 | 2E-41 | 1E-39 | 0.4363089 | surface |
| ALDH1L1 | 1.054 | 9.2841 | 556.794 | 4E-39 | 1.7E-37 | 0.5667001 | n.a. |
| CMBL | 1.051 | 9.249 | 1442.89 | 7E-39 | 3.1E-37 | 0.2778335 | n.a. |
| ENSECAG00000036115 | 1.089 | 9.2628 | 1110.65 | 8E-39 | 3.6E-37 | 0.4573721 | n.a. |
| GLUL | 1.104 | 9.3271 | 331.062 | 2E-38 | 7.8E-37 | 0.5827482 | n.a. |
| S100A11 | 1.005 | 9.847 | 164.351 | 2E-37 | 6.4E-36 | 0.9307924 | n.a. |
| SPX | 1.014 | 9.2562 | 1257.71 | 2E-37 | 7.3E-36 | 0.3440321 | n.a. |
| GBP1 | 1.049 | 9.279 | 453.166 | 2E-37 | 8.3E-36 | 0.3851555 | n.a. |
| LGMN | 1.108 | 9.3223 | 269.706 | 1E-35 | 3.8E-34 | 0.4794383 | n.a. |
| ME1 | 1.055 | 9.2478 | 1350.6 | 1E-35 | 5.5E-34 | 0.2828485 | n.a. |
| TTYH3 | 1.097 | 9.2991 | 319.068 | 1E-32 | 4.1E-31 | 0.4383149 | surface |
| TNFSF13 | 1.012 | 9.266 | 916.134 | 4E-32 | 1.4E-30 | 0.4694082 | surface |
| TMEM205 | 1.062 | 9.2912 | 276.487 | 9E-31 | 2.9E-29 | 0.4122367 | n.a. |
| APLP2 | 1.079 | 9.284 | 370.479 | 2E-30 | 6.7E-29 | 0.5145436 | surface |
| CCR2 | 1.079 | 9.2851 | 396.746 | 6E-30 | 1.7E-28 | 0.441324 | surface |

| genes | logFC | logCPM | F | PValue | FDR | percent.exp | Surfacome.Label |
|---|---|---|---|---|---|---|---|
| ENSECAG00000000436 | 3.095 | 9.9635 | 9426.4 | 0 | 0 | 0.962129 | n.a. |
| LYZ | 2.712 | 11.749 | 5529.1 | 0 | 0 | 0.9989765 | n.a. |
| THBS1 | 2.631 | 9.3512 | 4148.13 | 0 | 0 | 0.6990788 | n.a. |
| ENSECAG00000030777 | 2.413 | 9.3773 | 2905.08 | 0 | 0 | 0.7799386 | n.a. |
| GM2A | 1.974 | 9.9979 | 2330.1 | 0 | 0 | 0.9989765 | n.a. |
| RNASE6 | 1.58 | 10.205 | 1572.29 | 0 | 0 | 1 | n.a. |
| RNASE4 | 1.527 | 10.101 | 1552.27 | 0 | 0 | 0.9989765 | n.a. |
| S100P | 1.389 | 9.8205 | 2593.93 | 0 | 0 | 0.9549642 | n.a. |
| SULT1C4 | 2.239 | 9.421 | 4557.65 | 6E-304 | 5E-301 | 0.9365404 | n.a. |
| PLBD1 | 1.459 | 9.972 | 1414.62 | 2E-303 | 1E-300 | 0.9764585 | n.a. |
| TREM1 | 1.864 | 9.4852 | 4501.62 | 2E-300 | 1E-297 | 0.8679632 | surface |
| S100A4 | 2.501 | 11.292 | 1395.31 | 2E-299 | 1E-296 | 0.9979529 | n.a. |
| CTSZ | 1.811 | 9.8631 | 1296.26 | 1E-278 | 6E-276 | 0.9979529 | n.a. |
| IFI27 | 2.513 | 9.7314 | 1107.13 | 5E-239 | 3E-236 | 0.9324463 | n.a. |
| LAMP1 | 1.754 | 9.6692 | 995.894 | 2E-215 | 8E-213 | 0.9856704 | surface |
| ENSECAG00000024882 | 2.204 | 9.349 | 3121.29 | 2E-207 | 1E-204 | 0.593654 | n.a. |
| ENSECAG00000019309 | 2.173 | 9.3991 | 936.217 | 7E-203 | 3E-200 | 0.8495394 | n.a. |
| CYBB | 1.541 | 9.4653 | 1000.75 | 1E-184 | 5E-182 | 0.9293756 | surface |
| S100A10 | 1.688 | 10.866 | 819.648 | 3E-178 | 1E-175 | 0.9989765 | n.a. |
| S100A12 | 1.139 | 10.309 | 1085.15 | 3E-160 | 1E-157 | 0.9344933 | n.a. |
| ENSECAG00000024886 | 2.099 | 9.319 | 3751.79 | 5E-152 | 2E-149 | 0.7144319 | n.a. |
| SERPINB10 | 1.423 | 9.5345 | 1667.21 | 9E-152 | 3E-149 | 0.7635619 | n.a. |
| VIM | 1.011 | 11.286 | 643.029 | 1E-140 | 4E-138 | 0.9989765 | n.a. |
| C4BPA | 1.158 | 9.4013 | 1176.18 | 8E-138 | 2E-135 | 0.8822927 | n.a. |
| CSF1R | 1.737 | 9.3336 | 2106.13 | 2E-135 | 7E-133 | 0.8669396 | surface |
| ALDH2 | 1.301 | 9.5616 | 711.465 | 1E-134 | 3E-132 | 0.9539406 | n.a. |
| EMP1 | 2.007 | 9.2873 | 2576.04 | 1E-128 | 3E-126 | 0.5609007 | surface |
| AHNAK | 1.871 | 9.5205 | 554.681 | 1E-121 | 3E-119 | 0.9037871 | n.a. |
| LGALS3 | 1.332 | 9.5239 | 549.315 | 2E-120 | 4E-118 | 0.8157625 | n.a. |
| MHCB3 | 1.762 | 10.265 | 546.6 | 6E-120 | 1E-117 | 0.9447288 | n.a. |
| ENSECAG00000040634 | 1.925 | 9.9556 | 536.532 | 8E-118 | 2E-115 | 0.9293756 | n.a. |
| TSPAN4 | 1.804 | 9.2911 | 1247.2 | 2E-113 | 3E-111 | 0.5701126 | surface |
| ENSECAG00000029581 | 1.72 | 9.2817 | 2288.45 | 3E-112 | 6E-110 | 0.6305015 | n.a. |
| SLPI | 1.483 | 9.3697 | 2008.28 | 3E-107 | 6E-105 | 0.6550665 | n.a. |
| MPEG1 | 1.332 | 9.3534 | 1017.34 | 4E-104 | 7E-102 | 0.8485159 | surface |
| MNDA | 1.582 | 10.258 | 469.65 | 2E-103 | 3E-101 | 0.9897646 | n.a. |
| FGL2 | 1.199 | 9.488 | 457.7 | 7E-101 | 1.1E-98 | 0.9416581 | n.a. |
| CREG1 | 1.362 | 9.3229 | 1674.7 | 9E-101 | 1.5E-98 | 0.7635619 | n.a. |
| ALDH1A1 | 1.777 | 9.2588 | 3314.76 | 7E-100 | 1.1E-97 | 0.4145343 | n.a. |
| HNMT | 1.845 | 9.2852 | 2864.59 | 7E-99 | 1E-96 | 0.6407369 | n.a. |
| TSPO | 1.044 | 9.7323 | 444.469 | 5E-98 | 7.2E-96 | 0.9570113 | n.a. |
| ENSECAG00000022247 | 1.49 | 9.2794 | 2861.24 | 9E-98 | 1.3E-95 | 0.5097236 | n.a. |
| ABCA6 | 1.885 | 9.28 | 2563.26 | 2E-96 | 3E-94 | 0.5056295 | surface |
| MMP9 | 1.022 | 9.4158 | 1065.41 | 1E-92 | 1.9E-90 | 0.457523 | n.a. |
| ENSECAG00000030387 | 1.525 | 9.433 | 525.3 | 5E-83 | 6.3E-81 | 0.6693961 | n.a. |
| S100A6 | 1.546 | 10.437 | 374.17 | 6E-83 | 8.3E-81 | 0.9570113 | n.a. |
| ENSECAG00000017042 | 1.272 | 9.3458 | 1064.6 | 2E-82 | 2.2E-80 | 0.8607984 | n.a. |
| VCAN | 1.271 | 9.3382 | 1846.07 | 3E-81 | 3.2E-79 | 0.4165814 | n.a. |
| ENSECAG00000021110 | 1.52 | 9.2654 | 1964.38 | 6E-81 | 6.6E-79 | 0.5332651 | n.a. |
| MAFB | 1.581 | 9.2647 | 1592.9 | 4E-78 | 4.7E-76 | 0.4483112 | n.a. |
| PLD3 | 1.464 | 9.3271 | 538.168 | 5E-78 | 5.9E-76 | 0.5199591 | n.a. |
| GLUL | 1.502 | 9.3271 | 554.871 | 1E-76 | 1.5E-74 | 0.6530194 | n.a. |
| PSAP | 1.003 | 10.167 | 344.286 | 2E-76 | 1.8E-74 | 1 | n.a. |
| GIMAP7 | 1.357 | 10.513 | 323.06 | 7E-72 | 6.5E-70 | 0.8781986 | n.a. |
| TIMP1 | 1.348 | 9.3333 | 619.059 | 5E-70 | 4.9E-68 | 0.6059365 | n.a. |

| | | | | | | | |
|---|---|---|---|---|---|---|---|
| ENSECAG00000040180 | 1.341 | 9.7881 | 312.181 | 1E-69 | 1.3E-67 | 0.8505629 | n.a. |
| MECR | 1.42 | 9.3421 | 560.842 | 2E-69 | 2E-67 | 0.6970317 | n.a. |
| ENSECAG00000029196 | 1.381 | 9.2683 | 880.048 | 3E-68 | 3E-66 | 0.3541453 | n.a. |
| TIMP2 | 1.33 | 9.2993 | 941.022 | 1E-67 | 9.6E-66 | 0.6816786 | n.a. |
| GBP1 | 1.358 | 9.279 | 649.275 | 3E-63 | 2E-61 | 0.4176049 | n.a. |
| ENSECAG00000015010 | 1.24 | 9.2872 | 1337.47 | 6E-62 | 4.5E-60 | 0.6110542 | n.a. |
| REXO2 | 1.187 | 9.4557 | 276.567 | 7E-62 | 5.3E-60 | 0.8710338 | n.a. |
| HACD4 | 1.187 | 9.3015 | 1435.33 | 4E-60 | 2.9E-58 | 0.6305015 | n.a. |
| ENSECAG00000029716 | 1.238 | 9.8771 | 265.59 | 2E-59 | 1.2E-57 | 0.8812692 | n.a. |
| PRNP | 1.035 | 9.3261 | 521.439 | 7E-59 | 4.4E-57 | 0.6878199 | surface |
| CD68 | 1.159 | 9.3465 | 576.137 | 3E-56 | 2.2E-54 | 0.8167861 | surface |
| DPYD | 1.011 | 9.2972 | 798.705 | 1E-52 | 8.5E-51 | 0.5578301 | n.a. |
| ARL5A | 1.234 | 9.344 | 280.574 | 6E-48 | 3E-46 | 0.6274309 | n.a. |
| S100A11 | 1.13 | 9.847 | 207.486 | 7E-47 | 3.5E-45 | 0.9426817 | n.a. |
| LDHB | 1.082 | 10.019 | 202.652 | 7E-46 | 3.8E-44 | 0.9764585 | n.a. |
| LRP1 | 1.266 | 9.2501 | 2061.23 | 8E-46 | 4.2E-44 | 0.321392 | surface |
| SPX | 1.129 | 9.2562 | 1398.73 | 1E-45 | 4.9E-44 | 0.3357216 | n.a. |
| CTSB | 1.029 | 9.4471 | 201.474 | 1E-45 | 6.7E-44 | 0.9334698 | n.a. |
| ENSECAG00000039193 | 1.231 | 9.2602 | 1116.03 | 3E-44 | 1.6E-42 | 0.4012282 | n.a. |
| KCNE3 | 1.168 | 9.2985 | 938.379 | 8E-44 | 3.9E-42 | 0.6345957 | n.a. |
| S100A5 | 1.04 | 9.5834 | 190.494 | 3E-43 | 1.5E-41 | 0.5772774 | n.a. |
| CD163 | 1.16 | 9.2595 | 1437.29 | 1E-42 | 6.5E-41 | 0.2845445 | surface |
| N4BP2L1 | 1.098 | 9.4151 | 184.202 | 8E-42 | 3.4E-40 | 0.7922211 | n.a. |
| ENSECAG00000035085 | 1.305 | 9.2904 | 1091.11 | 5E-41 | 2E-39 | 0.3694985 | n.a. |
| RAMP1 | 1.045 | 9.2514 | 1196.73 | 3E-40 | 1.4E-38 | 0.3039918 | n.a. |
| ENSECAG00000027666 | 1.077 | 9.8957 | 176.193 | 4E-40 | 1.8E-38 | 0.9631525 | n.a. |
| CASP4 | 1.05 | 9.4053 | 189.866 | 5E-39 | 2E-37 | 0.567042 | n.a. |
| DOK2 | 1.07 | 9.3423 | 282.334 | 1E-38 | 4.1E-37 | 0.5844422 | n.a. |
| ENSECAG00000003563 | 1.128 | 9.4315 | 176.893 | 1E-38 | 4.5E-37 | 0.593654 | n.a. |
| TBXAS1 | 1.075 | 9.2862 | 941.009 | 9E-38 | 3.5E-36 | 0.593654 | n.a. |
| LGMN | 1.158 | 9.3223 | 287.32 | 2E-36 | 7.4E-35 | 0.4605937 | n.a. |
| RGS10 | 1.088 | 9.5903 | 158.447 | 3E-36 | 1.1E-34 | 0.8096213 | n.a. |
| CMBL | 1.087 | 9.249 | 1537.55 | 8E-36 | 3E-34 | 0.2548618 | n.a. |
| SMS | 1.034 | 9.4498 | 156.412 | 8E-36 | 3.1E-34 | 0.8096213 | n.a. |
| GNS | 1.061 | 9.3016 | 291.447 | 5E-34 | 1.8E-32 | 0.383828 | n.a. |
| ENSECAG00000028267 | 1.18 | 9.2588 | 751.226 | 2E-33 | 6.7E-32 | 0.3715455 | n.a. |
| TTYH3 | 1.069 | 9.2991 | 309.795 | 6E-32 | 2.1E-30 | 0.4073695 | surface |
| APLP2 | 1.016 | 9.284 | 332.381 | 1E-27 | 3.7E-26 | 0.4984647 | surface |

| genes | logFC | logCPM | F | PValue | FDR | percent.exp | Surfacome.Label |
|---|---|---|---|---|---|---|---|
| S100A12 | 5.3 | 10.309 | 19855.1 | 0 | 0 | 0.9967742 | n.a. |
| ENSECAG00000010117 | 5.008 | 9.501 | 13637.9 | 0 | 0 | 0.9274194 | n.a. |
| LYZ | 4.332 | 11.749 | 10784.4 | 0 | 0 | 1 | n.a. |
| ENSECAG00000000436 | 4.219 | 9.9635 | 14142.9 | 0 | 0 | 0.9903226 | n.a. |
| S100A8 | 4.211 | 9.3765 | 12053 | 0 | 0 | 0.7935484 | n.a. |
| IL18 | 4.078 | 9.3131 | 4287.52 | 0 | 0 | 0.8483871 | n.a. |
| MMP9 | 4.054 | 9.4158 | 4883.67 | 0 | 0 | 0.9274194 | n.a. |
| ECATH-3 | 3.921 | 9.3365 | 9313.89 | 0 | 0 | 0.9032258 | n.a. |
| S100A6 | 3.752 | 10.437 | 2140.73 | 0 | 0 | 0.9854839 | n.a. |
| TREM1 | 3.67 | 9.4852 | 9714.4 | 0 | 0 | 0.9580645 | surface |
| C1H15orf48 | 3.534 | 9.6929 | 3625.76 | 0 | 0 | 0.9774194 | n.a. |
| S100P | 3.485 | 9.8205 | 9705.07 | 0 | 0 | 0.9967742 | n.a. |
| VCAN | 3.471 | 9.3382 | 5122.61 | 0 | 0 | 0.7935484 | n.a. |
| ENSECAG00000036100 | 3.424 | 9.3021 | 6512.23 | 0 | 0 | 0.7693548 | n.a. |
| PLBD1 | 3.324 | 9.972 | 5456.91 | 0 | 0 | 0.9903226 | n.a. |
| SERPINB10 | 3.22 | 9.5345 | 5608.6 | 0 | 0 | 0.7193548 | n.a. |
| S100A4 | 3.208 | 11.292 | 2074.17 | 0 | 0 | 0.9758065 | n.a. |
| ENSECAG00000020136 | 3.15 | 9.7014 | 1739.27 | 0 | 0 | 0.9677419 | n.a. |
| SLPI | 3.118 | 9.3697 | 5556.92 | 0 | 0 | 0.6919355 | n.a. |
| TSPO | 2.876 | 9.7323 | 2303.67 | 0 | 0 | 0.9854839 | n.a. |
| GM2A | 2.548 | 9.9979 | 3442.87 | 0 | 0 | 0.9790323 | n.a. |
| CTSZ | 2.227 | 9.8631 | 1699.03 | 0 | 0 | 0.9709677 | n.a. |
| SRGN | 2.078 | 10.485 | 1544.5 | 0 | 0 | 0.9983871 | n.a. |
| RNASE6 | 1.982 | 10.205 | 2040.62 | 0 | 0 | 0.9983871 | n.a. |
| RNASE4 | 1.895 | 10.101 | 1960.81 | 0 | 0 | 0.9919355 | n.a. |
| C5AR1 | 2.395 | 9.3446 | 4015.62 | 0 | 0 | 0.8532258 | surface |
| S100A5 | 3.059 | 9.5834 | 1420.48 | 1E-304 | 3E-302 | 0.7064516 | n.a. |
| ENSECAG00000030387 | 2.789 | 9.433 | 1388.81 | 1E-284 | 4E-282 | 0.6629032 | n.a. |
| ENSECAG00000030745 | 1.209 | 11.502 | 1285.13 | 2E-276 | 6E-274 | 0.9983871 | n.a. |
| PRDX5 | 2.945 | 9.5264 | 1261.78 | 2E-271 | 5E-269 | 0.9354839 | n.a. |
| MGST1 | 2.492 | 9.3352 | 3004.07 | 2E-243 | 4E-241 | 0.6677419 | n.a. |
| LGALS3 | 2.251 | 9.5239 | 1091.45 | 1E-235 | 3E-233 | 0.7725806 | n.a. |
| FTL | 1.277 | 11.187 | 1044.63 | 8E-226 | 2E-223 | 1 | n.a. |
| VIM | 1.428 | 11.286 | 1029.27 | 1E-222 | 3E-220 | 0.9887097 | n.a. |
| ILT11B | 2.809 | 9.2698 | 4345.14 | 2E-213 | 4E-211 | 0.7225806 | n.a. |
| CD14 | 2.576 | 9.3178 | 4881.8 | 2E-210 | 4E-208 | 0.95 | surface |
| ENSECAG00000024882 | 2.569 | 9.349 | 3571.44 | 2E-206 | 5E-204 | 0.5064516 | n.a. |
| CEBPB | 1.784 | 9.407 | 1391.36 | 7E-204 | 1E-201 | 0.7758065 | n.a. |
| SYNE4 | 1.116 | 10.39 | 935.989 | 7E-203 | 1E-200 | 0.9983871 | n.a. |
| ENSECAG00000036967 | 2.565 | 9.2599 | 4749.4 | 5E-201 | 9E-199 | 0.5322581 | n.a. |
| C4BPA | 1.821 | 9.4013 | 1836.35 | 4E-198 | 7E-196 | 0.8435484 | n.a. |
| SULT1C4 | 2.184 | 9.421 | 3967.17 | 9E-195 | 2E-192 | 0.7096774 | n.a. |
| IGSF6 | 1.779 | 9.3651 | 2009.28 | 2E-191 | 3E-189 | 0.8193548 | surface |
| ENSECAG00000037539 | 2.563 | 9.2665 | 4055.85 | 4E-188 | 7E-186 | 0.5677419 | n.a. |
| TKT | 1.74 | 9.6069 | 861.133 | 5E-187 | 9E-185 | 0.9016129 | n.a. |
| THBS1 | 2.538 | 9.3512 | 3190.4 | 2E-176 | 3E-174 | 0.5129032 | n.a. |
| ITM2B | 1.698 | 10.174 | 806.435 | 2E-175 | 4E-173 | 0.9612903 | surface |
| PSAP | 1.655 | 10.167 | 790.174 | 6E-172 | 1E-169 | 1 | n.a. |
| ENSECAG00000006595 | 2.469 | 9.2598 | 2502.3 | 2E-168 | 3E-166 | 0.5451613 | n.a. |
| NFE2 | 2.306 | 9.2974 | 3513.24 | 1E-160 | 2E-158 | 0.5919355 | n.a. |
| VNN2 | 2.28 | 9.2761 | 2225.76 | 1E-157 | 2E-155 | 0.5870968 | surface |
| CD44 | 1.812 | 10.108 | 716.553 | 3E-156 | 4E-154 | 0.9548387 | surface |
| QPCT | 2.245 | 9.3093 | 1750.59 | 3E-152 | 5E-150 | 0.6516129 | n.a. |
| LAMP1 | 1.614 | 9.6692 | 691.515 | 6E-151 | 9E-149 | 0.8564516 | surface |
| CTSS | 1.345 | 9.8161 | 690.406 | 1E-150 | 1E-148 | 0.9532258 | n.a. |

| | | | | | | | |
|---|---|---|---|---|---|---|---|
| DUSP1 | 1.669 | 9.3756 | 689.685 | 2E-150 | 2E-148 | 0.6951613 | n.a. |
| ENSECAG00000003345 | 1.734 | 10.17 | 685.695 | 1E-149 | 1E-147 | 0.9225806 | n.a. |
| GSN | 1.697 | 9.3636 | 1401.19 | 7E-138 | 9E-136 | 0.7177419 | n.a. |
| MAPK13 | 2.424 | 9.2495 | 3514.46 | 9E-138 | 1E-135 | 0.4225806 | n.a. |
| ENSECAG00000010615 | 2.433 | 9.2458 | 1789.73 | 2E-135 | 3E-133 | 0.2870968 | n.a. |
| PADI4 | 2.041 | 9.2553 | 3625.85 | 2E-132 | 3E-130 | 0.4241935 | n.a. |
| GLIPR2 | 1.639 | 9.47 | 609.879 | 4E-129 | 5E-127 | 0.7080645 | n.a. |
| MHCB3 | 2.023 | 10.265 | 578.498 | 9E-127 | 1E-124 | 0.7645161 | n.a. |
| CAPG | 1.923 | 9.6557 | 571.219 | 3E-125 | 4E-123 | 0.7854839 | n.a. |
| ATP1B3 | 1.812 | 9.5391 | 555.174 | 9E-122 | 1E-119 | 0.7903226 | surface |
| RAB24 | 1.688 | 9.4289 | 554.008 | 2E-121 | 2E-119 | 0.7032258 | n.a. |
| ALDOA | 1.341 | 9.99 | 530.76 | 1E-116 | 2E-114 | 0.95 | n.a. |
| ALDH3A1 | 2.44 | 9.2538 | 3534.7 | 4E-116 | 5E-114 | 0.4 | n.a. |
| GAPDH | 1.085 | 10.705 | 521.696 | 1E-114 | 1E-112 | 0.983871 | n.a. |
| ENSECAG00000030271 | 1.51 | 9.3599 | 621 | 6E-111 | 7E-109 | 0.7274194 | n.a. |
| MECR | 2.008 | 9.3421 | 998.621 | 1E-107 | 1E-105 | 0.4612903 | n.a. |
| NAPSA | 1.38 | 9.5255 | 641.755 | 2E-105 | 2E-103 | 0.7774194 | n.a. |
| HMGB2 | 1.578 | 10.129 | 465.464 | 1E-102 | 1E-100 | 0.8306452 | n.a. |
| TUT7 | 1.661 | 9.6301 | 461.57 | 1E-101 | 1E-99 | 0.816129 | n.a. |
| PLP2 | 1.524 | 9.9939 | 457.994 | 6E-101 | 6E-99 | 0.8806452 | n.a. |
| PSTPIP1 | 1.673 | 9.5292 | 456.847 | 1E-100 | 1E-98 | 0.733871 | n.a. |
| ARG2 | 1.767 | 9.2454 | 3243.66 | 5E-100 | 5E-98 | 0.2806452 | n.a. |
| SLC25A5 | 1.069 | 11.478 | 447.05 | 1E-98 | 1.3E-96 | 0.9951613 | n.a. |
| DPYD | 1.792 | 9.2972 | 1270.51 | 5E-97 | 4.5E-95 | 0.5419355 | n.a. |
| FGL2 | 1.399 | 9.488 | 507.329 | 2E-96 | 1.8E-94 | 0.8258065 | n.a. |
| TALDO1 | 1.309 | 9.9099 | 433.463 | 1E-95 | 1E-93 | 0.9467742 | n.a. |
| ENSECAG00000033857 | 1.345 | 9.3316 | 2108.94 | 1E-92 | 1E-90 | 0.6274194 | n.a. |
| SELP | 1.97 | 9.2569 | 2645.03 | 1E-91 | 1.1E-89 | 0.4258065 | surface |
| PGD | 1.411 | 9.3424 | 601.697 | 9E-91 | 7.7E-89 | 0.6274194 | n.a. |
| ENSECAG00000024563 | 1.575 | 9.2493 | 1899.98 | 1E-90 | 1.1E-88 | 0.283871 | n.a. |
| HSPB1 | 1.38 | 9.6778 | 403.94 | 2E-89 | 2.1E-87 | 0.9274194 | n.a. |
| PLD3 | 1.876 | 9.3271 | 686.233 | 3E-89 | 2.9E-87 | 0.383871 | n.a. |
| ENSECAG00000024886 | 1.711 | 9.319 | 2850.86 | 1E-86 | 9.5E-85 | 0.3709677 | n.a. |
| ENSECAG00000040180 | 1.681 | 9.7881 | 391.038 | 1E-86 | 1.2E-84 | 0.7048387 | n.a. |
| DRAXIN | 1.612 | 9.3226 | 743.546 | 1E-84 | 8.8E-83 | 0.5048387 | n.a. |
| NCF4 | 1.562 | 9.2924 | 1230.51 | 1E-80 | 9.9E-79 | 0.3919355 | n.a. |
| MNDA | 1.568 | 10.258 | 361.352 | 4E-80 | 2.9E-78 | 0.9 | n.a. |
| ALDH2 | 1.334 | 9.5616 | 641.946 | 5E-80 | 3.8E-78 | 0.7048387 | n.a. |
| HK3 | 1.521 | 9.284 | 1853.58 | 3E-78 | 2.5E-76 | 0.4290323 | n.a. |
| NFAM1 | 1.342 | 9.3582 | 974.431 | 1E-77 | 9.1E-76 | 0.6451613 | surface |
| LST1 | 1.252 | 9.329 | 576.344 | 2E-76 | 1.2E-74 | 0.5806452 | n.a. |
| BLVRB | 1.398 | 9.4497 | 382.879 | 2E-74 | 1.5E-72 | 0.6451613 | n.a. |
| ENSECAG00000035431 | 1.289 | 9.323 | 1324.05 | 2E-73 | 1.2E-71 | 0.6016129 | n.a. |
| eca-mir-223 | 1.555 | 9.2778 | 1484.93 | 2E-72 | 1.5E-70 | 0.4096774 | n.a. |
| ALOX5AP | 1.72 | 9.2637 | 1537.8 | 2E-72 | 1.6E-70 | 0.2935484 | n.a. |
| RABAC1 | 1.276 | 9.8034 | 325.14 | 2E-72 | 1.7E-70 | 0.8967742 | n.a. |
| NUDT4 | 1.59 | 9.2991 | 548.029 | 1E-71 | 7.4E-70 | 0.4645161 | n.a. |
| ENSECAG00000022247 | 1.509 | 9.2794 | 2583.69 | 1E-71 | 9.3E-70 | 0.3064516 | n.a. |
| CREG1 | 1.419 | 9.3229 | 1824.95 | 1E-71 | 9.4E-70 | 0.4951613 | n.a. |
| AMDHD2 | 1.128 | 9.7855 | 316.447 | 2E-70 | 1.2E-68 | 0.9241935 | n.a. |
| XBP1 | 1.614 | 9.4868 | 315.577 | 3E-70 | 1.9E-68 | 0.5887097 | n.a. |
| ENSECAG00000008826 | 1.616 | 9.376 | 409.692 | 8E-70 | 5.2E-68 | 0.4354839 | n.a. |
| CSF3R | 1.769 | 9.2479 | 3149.11 | 8E-69 | 5.1E-67 | 0.366129 | surface |
| SELL | 1.398 | 9.6904 | 303.741 | 1E-67 | 6.6E-66 | 0.7532258 | surface |
| VSIR | 1.227 | 9.4326 | 302.472 | 2E-67 | 1.2E-65 | 0.6580645 | n.a. |
| BNIP3L | 1.478 | 9.4595 | 300.323 | 5E-67 | 3.5E-65 | 0.6129032 | n.a. |

| | | | | | | | |
|---|---|---|---|---|---|---|---|
| GIMAP7 | 1.525 | 10.513 | 298.175 | 2E-66 | 1E-64 | 0.6225806 | n.a. |
| ICAM3 | 1.408 | 9.5641 | 292.075 | 3E-65 | 2.1E-63 | 0.6048387 | surface |
| GLIPR1 | 1.257 | 9.887 | 280.336 | 1E-62 | 7.1E-61 | 0.7919355 | surface |
| ANXA1 | 1.233 | 9.8295 | 278.464 | 3E-62 | 1.8E-60 | 0.8274194 | n.a. |
| PRELID1 | 1.219 | 9.7488 | 277.37 | 5E-62 | 3E-60 | 0.8435484 | n.a. |
| PGK1 | 1.238 | 9.4881 | 276.431 | 8E-62 | 4.8E-60 | 0.6741935 | n.a. |
| SLC11A1 | 1.294 | 9.3411 | 449.962 | 3E-60 | 1.5E-58 | 0.3225806 | surface |
| ENSECAG00000031962 | 1.617 | 9.2453 | 1494.5 | 5E-60 | 2.7E-58 | 0.2758065 | n.a. |
| ENSECAG00000016578 | 1.014 | 9.4164 | 951.033 | 1E-59 | 7.5E-58 | 0.7596774 | n.a. |
| RGS2 | 1.226 | 9.2835 | 1108.18 | 1E-58 | 5.9E-57 | 0.4516129 | n.a. |
| ITGAM | 1.49 | 9.3026 | 843.095 | 1E-58 | 6.2E-57 | 0.3822581 | surface |
| CERS4 | 1.72 | 9.2984 | 503.255 | 1E-58 | 8.3E-57 | 0.3612903 | n.a. |
| TAGLN2 | 1.108 | 10.164 | 259.935 | 3E-58 | 1.6E-56 | 0.8854839 | n.a. |
| ENSECAG00000007681 | 1.433 | 9.6603 | 270.448 | 7E-58 | 4.2E-56 | 0.5112903 | n.a. |
| PGAM1 | 1.321 | 9.5435 | 256.747 | 1E-57 | 7.9E-56 | 0.6032258 | n.a. |
| CD300LB | 1.271 | 9.263 | 1717.85 | 7E-57 | 3.8E-55 | 0.2870968 | n.a. |
| CD63 | 1.182 | 9.4159 | 271.632 | 3E-56 | 1.7E-54 | 0.5919355 | surface |
| ENSECAG00000024719 | 1.487 | 9.2957 | 483.352 | 1E-55 | 6E-54 | 0.366129 | n.a. |
| TMEM150B | 1.61 | 9.2642 | 1694.19 | 2E-55 | 1E-53 | 0.2790323 | surface |
| ENSECAG00000029003 | 1.179 | 10.041 | 244.482 | 6E-55 | 3.4E-53 | 0.8145161 | n.a. |
| EMP3 | 1.479 | 10.126 | 244.269 | 7E-55 | 3.8E-53 | 0.7790323 | surface |
| CARHSP1 | 1.441 | 9.4061 | 274.796 | 1E-54 | 5.2E-53 | 0.4322581 | n.a. |
| BTG1 | 1.052 | 10.365 | 235.245 | 6E-53 | 3.3E-51 | 0.8241935 | n.a. |
| EQMCE1 | 1.018 | 10.07 | 234.849 | 8E-53 | 4E-51 | 0.866129 | n.a. |
| CLEC4E | 1.46 | 9.2548 | 1842.14 | 1E-52 | 5.1E-51 | 0.2790323 | n.a. |
| LDHA | 1.126 | 9.6788 | 232.396 | 3E-52 | 1.4E-50 | 0.7564516 | n.a. |
| ENSECAG00000032032 | 1.395 | 9.2656 | 868.547 | 6E-52 | 2.8E-50 | 0.2548387 | n.a. |
| SOD2 | 1.298 | 9.3528 | 290.365 | 1E-51 | 5.5E-50 | 0.4516129 | n.a. |
| ENSECAG00000024181 | 1.493 | 9.2586 | 1817.33 | 2E-51 | 8.4E-50 | 0.2919355 | n.a. |
| ENSECAG00000036115 | 1.362 | 9.2628 | 1566.35 | 2E-51 | 1E-49 | 0.2629032 | n.a. |
| CARD19 | 1.324 | 9.4452 | 232.22 | 5E-51 | 2.4E-49 | 0.4548387 | n.a. |
| LYPLA1 | 1.477 | 9.3462 | 348.908 | 8E-51 | 3.8E-49 | 0.383871 | n.a. |
| PKM | 1.19 | 9.8077 | 223.143 | 3E-50 | 1.3E-48 | 0.7032258 | n.a. |
| GPSM3 | 1.016 | 9.9781 | 217.172 | 5E-49 | 2.5E-47 | 0.8403226 | n.a. |
| SMS | 1.308 | 9.4498 | 234.081 | 1E-48 | 5.2E-47 | 0.5225806 | n.a. |
| BSG | 1.112 | 9.6757 | 214.956 | 2E-48 | 7.4E-47 | 0.6806452 | surface |
| PNPLA2 | 1.145 | 9.3533 | 317.954 | 2E-48 | 8.1E-47 | 0.5403226 | n.a. |
| CDA | 1.236 | 9.2512 | 1457.97 | 2E-48 | 9.2E-47 | 0.2919355 | n.a. |
| ENSECAG00000031156 | 1.267 | 9.3403 | 601.286 | 3E-48 | 1.4E-46 | 0.3983871 | n.a. |
| ENSECAG00000015500 | 1.281 | 9.2601 | 1406.61 | 5E-48 | 2.1E-46 | 0.316129 | n.a. |
| CASP4 | 1.472 | 9.4053 | 277.141 | 2E-47 | 7.1E-46 | 0.3419355 | n.a. |
| GTF2A2 | 1.308 | 9.5087 | 208.566 | 4E-47 | 1.7E-45 | 0.5096774 | n.a. |
| SIRPB2 | 1.302 | 9.2571 | 2169.7 | 5E-47 | 2.4E-45 | 0.2596774 | surface |
| GYG1 | 1.216 | 9.5017 | 204.966 | 2E-46 | 1E-44 | 0.583871 | n.a. |
| ENSECAG00000012199 | 1.07 | 9.3154 | 493.983 | 2E-46 | 1E-44 | 0.4048387 | n.a. |
| PGLS | 1.032 | 9.6893 | 203.304 | 5E-46 | 2.3E-44 | 0.8612903 | n.a. |
| TNFSF10 | 1.163 | 9.4268 | 237.717 | 8E-46 | 3.3E-44 | 0.3419355 | n.a. |
| RNF149 | 1.292 | 9.3444 | 297.25 | 1E-45 | 4.8E-44 | 0.5274194 | surface |
| ANXA7 | 1.082 | 9.5774 | 200.711 | 2E-45 | 8.2E-44 | 0.7145161 | n.a. |
| IFIT2 | 1.013 | 9.3061 | 499.734 | 2E-45 | 8.4E-44 | 0.4403226 | n.a. |
| ENSECAG00000033034 | 1.254 | 9.2773 | 1095.29 | 2E-45 | 9.1E-44 | 0.3645161 | n.a. |
| ADIPOR1 | 1.292 | 9.37 | 238.058 | 6E-45 | 2.6E-43 | 0.4354839 | n.a. |
| NPL | 1.167 | 9.2796 | 1072.36 | 9E-45 | 3.8E-43 | 0.3790323 | n.a. |
| EIF4A1 | 1.259 | 9.6961 | 195.253 | 3E-44 | 1.2E-42 | 0.6758065 | n.a. |
| RF00272 | 1.254 | 9.6864 | 194.498 | 4E-44 | 1.8E-42 | 0.6709677 | n.a. |
| AGPAT2 | 1.091 | 9.4052 | 198.736 | 5E-44 | 1.9E-42 | 0.5241935 | n.a. |

| | | | | | | | |
|---|---|---|---|---|---|---|---|
| PPT1 | 1.095 | 9.4371 | 202.436 | 5E-44 | 2E-42 | 0.6177419 | n.a. |
| GALK1 | 1.384 | 9.2866 | 626.84 | 4E-43 | 1.6E-41 | 0.3129032 | n.a. |
| MSN | 1.074 | 9.6123 | 186.305 | 3E-42 | 1E-40 | 0.7016129 | n.a. |
| HEXB | 1.288 | 9.3642 | 246.577 | 9E-42 | 3.6E-40 | 0.4822581 | n.a. |
| SNX10 | 1.075 | 9.274 | 715.126 | 1E-41 | 3.7E-40 | 0.3 | n.a. |
| ENSECAG00000010008 | 1.268 | 9.4684 | 181.718 | 3E-41 | 9.8E-40 | 0.4483871 | n.a. |
| ENSECAG00000019430 | 1.339 | 9.3323 | 293.384 | 7E-41 | 2.5E-39 | 0.2967742 | n.a. |
| YPEL5 | 1.083 | 9.4914 | 178.853 | 1E-40 | 4.1E-39 | 0.5967742 | n.a. |
| VMP1 | 1.421 | 9.3349 | 277.94 | 2E-40 | 7.3E-39 | 0.3548387 | n.a. |
| IVNS1ABP | 1.296 | 9.3077 | 278.851 | 1E-38 | 3.7E-37 | 0.4193548 | n.a. |
| RHBDD2 | 1.214 | 9.3113 | 282.979 | 2E-38 | 6.2E-37 | 0.4177419 | n.a. |
| ENSECAG00000027864 | 1.098 | 9.2631 | 1017.61 | 2E-38 | 7.3E-37 | 0.3258065 | n.a. |
| SAMSN1 | 1.072 | 9.3692 | 228.584 | 4E-38 | 1.6E-36 | 0.2887097 | n.a. |
| ARRB2 | 1.07 | 9.4662 | 181.214 | 5E-38 | 1.6E-36 | 0.4467742 | n.a. |
| ENSECAG00000029716 | 1.156 | 9.8771 | 170.337 | 5E-38 | 1.8E-36 | 0.6290323 | n.a. |
| LTB4R | 1.209 | 9.2469 | 1462.96 | 5E-37 | 1.9E-35 | 0.2548387 | surface |
| CDK2AP2 | 1.008 | 9.6423 | 158.771 | 3E-36 | 8.7E-35 | 0.6967742 | n.a. |
| ENSECAG00000017324 | 1.029 | 9.4228 | 169.448 | 4E-36 | 1.4E-34 | 0.3725806 | n.a. |
| SLC9A3R1 | 1.145 | 9.5551 | 159.734 | 7E-36 | 2.3E-34 | 0.333871 | n.a. |
| RAB5IF | 1.065 | 9.5719 | 148.167 | 5E-34 | 1.7E-32 | 0.6096774 | n.a. |
| IRF2 | 1.053 | 9.4594 | 147.907 | 6E-34 | 1.9E-32 | 0.4370968 | n.a. |
| ENSECAG00000006721 | 1.073 | 9.454 | 153.349 | 1E-33 | 4.5E-32 | 0.55 | n.a. |
| EHD1 | 1.001 | 9.4259 | 168.027 | 1E-33 | 4.6E-32 | 0.3370968 | n.a. |
| TNFRSF1A | 1.08 | 9.3254 | 247.924 | 1E-32 | 3.9E-31 | 0.4435484 | surface |
| SUSD3 | 1.118 | 9.5214 | 141.494 | 1E-32 | 4.5E-31 | 0.3129032 | surface |
| RTN4 | 1.192 | 9.4846 | 146.564 | 3E-32 | 8.9E-31 | 0.433871 | n.a. |
| ENSECAG00000007663 | 1.13 | 9.734 | 157.827 | 6E-32 | 1.9E-30 | 0.4145161 | n.a. |
| GLUL | 1.12 | 9.3271 | 291.524 | 4E-30 | 1.1E-28 | 0.3209677 | n.a. |
| CAMKK2 | 1.204 | 9.2678 | 517.493 | 2E-29 | 6.5E-28 | 0.316129 | n.a. |
| CALCOCO2 | 1.044 | 9.336 | 217.015 | 2E-29 | 6.8E-28 | 0.3096774 | n.a. |
| XG | 1.029 | 9.2905 | 285.925 | 6E-29 | 1.7E-27 | 0.35 | n.a. |
| ENSECAG00000037840 | 1.153 | 9.3288 | 230.736 | 5E-28 | 1.3E-26 | 0.2532258 | n.a. |
| ENSECAG00000036564 | 1.037 | 9.6716 | 118.807 | 1E-27 | 3.4E-26 | 0.5483871 | n.a. |
| CPNE3 | 1.021 | 9.3269 | 178.571 | 3E-27 | 8.4E-26 | 0.4193548 | n.a. |
| ACTN1 | 1.129 | 9.4643 | 127.857 | 3E-27 | 9E-26 | 0.2919355 | n.a. |
| RNF166 | 1.026 | 9.3001 | 227.813 | 2E-26 | 5.4E-25 | 0.3451613 | n.a. |
| TMEM70 | 1.076 | 9.2927 | 308.613 | 5E-26 | 1.2E-24 | 0.3193548 | n.a. |
| HIF1A | 1.042 | 9.3685 | 155.376 | 5E-26 | 1.3E-24 | 0.4048387 | n.a. |
| ALDH1L1 | 1.102 | 9.2841 | 453.939 | 1E-25 | 2.8E-24 | 0.3370968 | n.a. |
| MOSPD3 | 1.051 | 9.3446 | 149.285 | 2E-22 | 4.2E-21 | 0.3 | n.a. |
| CD47 | 1.113 | 9.3081 | 215.104 | 7E-21 | 1.6E-19 | 0.316129 | surface |

| genes | logFC | logCPM | F | PValue | FDR | percent.exp | Surfacome.Label |
|---|---|---|---|---|---|---|---|
| FCER1A | 3.89 | 9.4154 | 7025.63 | 0 | 0 | 0.6699029 | surface |
| ENSECAG00000038313 | 3.031 | 9.3718 | 2089.14 | 0 | 0 | 0.7524272 | n.a. |
| ENSECAG00000014585 | 2.363 | 10.271 | 1994.95 | 0 | 0 | 0.9773463 | n.a. |
| DRA | 2.343 | 11.742 | 2309.06 | 0 | 0 | 0.9983819 | n.a. |
| DRB | 2.31 | 11.205 | 1720.48 | 0 | 0 | 0.9983819 | n.a. |
| CD74 | 1.931 | 12.628 | 2081.05 | 0 | 0 | 1 | surface |
| CST3 | 1.412 | 11.921 | 1641.12 | 0 | 0 | 1 | n.a. |
| DQA | 2.193 | 10.712 | 1317.16 | 4E-283 | 4E-280 | 0.9983819 | n.a. |
| Eqca-DQB1 | 2.034 | 10.925 | 1138.42 | 1E-245 | 1E-242 | 0.9951456 | n.a. |
| ENSECAG00000019318 | 2.156 | 9.3455 | 1174.23 | 7E-229 | 6E-226 | 0.7864078 | n.a. |
| CD1E2 | 2.823 | 9.2674 | 2786.46 | 1E-214 | 7E-212 | 0.6294498 | n.a. |
| MT3 | 2.14 | 9.2863 | 6993.09 | 5E-203 | 3E-200 | 0.4433657 | n.a. |
| DQB | 2.043 | 10.011 | 807.346 | 1E-175 | 7E-173 | 0.9482201 | n.a. |
| FABP3 | 1.97 | 9.3412 | 1058.98 | 4E-160 | 2E-157 | 0.5841424 | n.a. |
| IFI30 | 1.807 | 9.9429 | 724.869 | 5E-158 | 2E-155 | 0.9919094 | n.a. |
| ENSECAG00000029951 | 2.246 | 9.3173 | 976.278 | 2E-154 | 9E-152 | 0.6100324 | n.a. |
| ENSECAG00000038794 | 2.28 | 9.2633 | 3431.58 | 3E-154 | 1E-151 | 0.7216828 | n.a. |
| ID3 | 2.076 | 9.6697 | 592.521 | 9E-130 | 3E-127 | 0.4805825 | n.a. |
| SLAMF9 | 2.046 | 9.2456 | 3848.28 | 7E-129 | 2E-126 | 0.3996764 | surface |
| CKB | 2.033 | 9.3208 | 2320.96 | 2E-127 | 6E-125 | 0.6957929 | n.a. |
| PID1 | 1.862 | 9.2473 | 4409.72 | 2E-120 | 6E-118 | 0.4288026 | n.a. |
| S100A10 | 1.488 | 10.866 | 537.926 | 4E-118 | 1E-115 | 0.9919094 | n.a. |
| CLIC2 | 1.585 | 9.286 | 1978.38 | 9E-117 | 2E-114 | 0.4012945 | n.a. |
| MEF2C | 1.542 | 9.4205 | 658.974 | 1E-111 | 2E-109 | 0.8656958 | n.a. |
| HLA-DOA | 1.8 | 9.3227 | 1004.82 | 1E-111 | 3E-109 | 0.6326861 | surface |
| PKIB | 1.735 | 9.2759 | 1404.86 | 1E-107 | 2E-105 | 0.5032362 | n.a. |
| Eqca-DOB1 | 1.429 | 9.4745 | 1039.14 | 1E-101 | 2E-99 | 0.461165 | n.a. |
| DPP4 | 1.451 | 9.2884 | 909.908 | 4E-97 | 7E-95 | 0.2912621 | surface |
| GPIHBP1 | 1.493 | 9.2751 | 3327.42 | 6E-97 | 9E-95 | 0.4401294 | surface |
| RPSA | 1.038 | 12.095 | 430.685 | 4E-95 | 6.9E-93 | 1 | n.a. |
| RF00091 | 1.06 | 12.032 | 427.193 | 2E-94 | 3.8E-92 | 1 | n.a. |
| ENSECAG00000021796 | 1.542 | 9.6911 | 415.399 | 8E-92 | 1.2E-89 | 0.9724919 | n.a. |
| ASGR2 | 1.964 | 9.2429 | 1488.76 | 9E-92 | 1.2E-89 | 0.3300971 | surface |
| CFP | 1.272 | 9.4343 | 1476.33 | 1E-90 | 2E-88 | 0.9514563 | n.a. |
| CD1B1 | 1.507 | 9.2551 | 1694.09 | 4E-88 | 4.9E-86 | 0.2588997 | n.a. |
| ENSECAG00000031569 | 1.357 | 11.194 | 393.072 | 5E-87 | 6.8E-85 | 0.9967638 | n.a. |
| LGALS1 | 1.356 | 11.109 | 385.054 | 3E-85 | 3.5E-83 | 1 | n.a. |
| ANXA3 | 1.709 | 9.2812 | 792.705 | 3E-85 | 3.6E-83 | 0.5825243 | n.a. |
| FCGRT | 1.474 | 9.3372 | 428.496 | 6E-83 | 6.6E-81 | 0.868932 | surface |
| TIMP1 | 1.631 | 9.3333 | 779.184 | 5E-77 | 5.6E-75 | 0.7022654 | n.a. |
| S100A4 | 1.471 | 11.292 | 345.49 | 1E-76 | 9.8E-75 | 0.97411 | n.a. |
| TSTD1 | 1.35 | 9.2702 | 724.761 | 4E-75 | 4E-73 | 0.3802589 | n.a. |
| ENSECAG00000020152 | 1.082 | 11.273 | 333.22 | 4E-74 | 4.2E-72 | 0.9967638 | n.a. |
| S100A6 | 1.525 | 10.437 | 322.395 | 9E-72 | 9E-70 | 0.9692557 | n.a. |
| CX3CR1 | 1.222 | 9.2887 | 1242.41 | 7E-70 | 6E-68 | 0.4223301 | surface |
| ENSECAG00000040634 | 1.588 | 9.9556 | 310.414 | 4E-69 | 3.2E-67 | 0.9029126 | n.a. |
| ENSECAG00000035085 | 2.046 | 9.2904 | 1985.69 | 7E-69 | 6.2E-67 | 0.3996764 | n.a. |
| ENSECAG00000017092 | 1.408 | 9.2393 | 3601.29 | 5E-66 | 3.8E-64 | 0.276699 | n.a. |
| PECAM1 | 1.097 | 9.2688 | 1219.76 | 4E-65 | 2.8E-63 | 0.3220065 | surface |
| RAB38 | 1.557 | 9.2603 | 1578.74 | 2E-62 | 1.4E-60 | 0.47411 | n.a. |
| YEATS4 | 1.588 | 9.4194 | 277.345 | 5E-62 | 3.8E-60 | 0.7540453 | n.a. |
| HLA-DMA | 1.311 | 9.6051 | 276.976 | 6E-62 | 4.5E-60 | 0.9433657 | surface |
| ENSECAG00000028889 | 1.593 | 9.2871 | 894.953 | 5E-59 | 3.5E-57 | 0.5744337 | n.a. |
| EGLN3 | 1.225 | 9.3249 | 388.387 | 1E-57 | 8.3E-56 | 0.7491909 | n.a. |
| ENSECAG00000016543 | 1.444 | 9.8125 | 253.61 | 7E-57 | 4.6E-55 | 0.8559871 | n.a. |

| | | | | | | | |
|---|---|---|---|---|---|---|---|
| ENSECAG00000009733 | 1.518 | 9.3278 | 327.299 | 3E-56 | 2.1E-54 | 0.5372168 | n.a. |
| LPAR6 | 1.567 | 9.2687 | 1015.41 | 2E-54 | 1.1E-52 | 0.5906149 | surface |
| FGGY | 1.447 | 9.2697 | 809.224 | 3E-53 | 2E-51 | 0.6278317 | n.a. |
| MYCBP2 | 1.346 | 9.5377 | 232.658 | 2E-52 | 1.4E-50 | 0.8915858 | n.a. |
| TSPAN13 | 1.281 | 9.2631 | 474.462 | 7E-48 | 4E-46 | 0.3915858 | surface |
| S100A11 | 1.15 | 9.847 | 191.806 | 2E-43 | 8.9E-42 | 0.8543689 | n.a. |
| ABCA6 | 1.482 | 9.28 | 1395.44 | 2E-41 | 9.2E-40 | 0.4271845 | surface |
| PEA15 | 1.057 | 9.2969 | 306.619 | 3E-39 | 1.6E-37 | 0.4514563 | n.a. |
| S100A5 | 1.094 | 9.5834 | 171.613 | 4E-39 | 2E-37 | 0.6084142 | n.a. |
| TM6SF1 | 1.223 | 9.2848 | 306.22 | 5E-39 | 2.6E-37 | 0.631068 | n.a. |
| NCALD | 1.058 | 9.279 | 400.941 | 6E-35 | 2.7E-33 | 0.3608414 | n.a. |
| CCDC88A | 1.107 | 9.3018 | 450.965 | 2E-33 | 8.2E-32 | 0.7944984 | n.a. |
| ENSECAG00000012400 | 1.068 | 9.3413 | 155.067 | 2E-31 | 8.3E-30 | 0.368932 | n.a. |
| RGS10 | 1.039 | 9.5903 | 127.579 | 2E-29 | 6.2E-28 | 0.8446602 | n.a. |
| ENSECAG00000013393 | 1.244 | 9.2781 | 333.85 | 2E-29 | 6.9E-28 | 0.5776699 | n.a. |
| DAPK1 | 1.114 | 9.2561 | 700.273 | 9E-27 | 3.4E-25 | 0.4676375 | n.a. |
| BTBD1 | 1.009 | 9.2813 | 218.96 | 6E-25 | 2E-23 | 0.420712 | n.a. |
| TCN2 | 1.172 | 9.2547 | 603.549 | 4E-23 | 1.4E-21 | 0.3980583 | n.a. |

| genes | logFC | logCPM | F | PValue | FDR | percent.exp | Surfacome.Label |
|---|---|---|---|---|---|---|---|
| CPVL | 5.186 | 9.3018 | 8463.33 | 0 | 0 | 0.96875 | n.a. |
| MT3 | 5.029 | 9.2863 | 13906.1 | 0 | 0 | 0.9322917 | n.a. |
| ENSECAG00000039383 | 4.511 | 9.2686 | 33228.3 | 0 | 0 | 0.9635417 | n.a. |
| ENSECAG00000014585 | 4.219 | 10.271 | 4803.91 | 0 | 0 | 0.9947917 | n.a. |
| BATF3 | 4.214 | 9.2774 | 3850.1 | 0 | 0 | 0.9739583 | n.a. |
| CLEC9A | 4.194 | 9.2481 | 28129.4 | 0 | 0 | 0.859375 | surface |
| DNASE1L3 | 4.093 | 9.2492 | 15919.1 | 0 | 0 | 0.8385417 | n.a. |
| ENSECAG00000029928 | 4.014 | 9.2723 | 7260.93 | 0 | 0 | 0.9427083 | n.a. |
| ENSECAG00000024996 | 3.86 | 9.2908 | 2604.09 | 0 | 0 | 0.96875 | n.a. |
| GPIHBP1 | 3.711 | 9.2751 | 6161.37 | 0 | 0 | 0.9427083 | surface |
| CST3 | 3.351 | 11.921 | 5993.78 | 0 | 0 | 1 | n.a. |
| ENSECAG00000038313 | 3.209 | 9.3718 | 1888.01 | 0 | 0 | 0.828125 | n.a. |
| DRA | 2.969 | 11.742 | 2759.18 | 0 | 0 | 1 | n.a. |
| DRB | 2.809 | 11.205 | 1860.21 | 0 | 0 | 1 | n.a. |
| CD74 | 2.38 | 12.628 | 2234.39 | 0 | 0 | 1 | surface |
| IRF8 | 3.05 | 9.303 | 1468.81 | 0 | 0 | 0.9791667 | n.a. |
| CKB | 2.989 | 9.3208 | 3070.73 | 2E-302 | 7E-300 | 0.90625 | n.a. |
| DQB | 3.064 | 10.011 | 1407.82 | 5E-302 | 2E-299 | 0.96875 | n.a. |
| ID2 | 3.274 | 9.6451 | 1403.25 | 5E-301 | 2E-298 | 0.984375 | n.a. |
| DQA | 2.624 | 10.712 | 1344.85 | 7E-289 | 3E-286 | 1 | n.a. |
| Eqca-DOB1 | 2.764 | 9.4745 | 2042.29 | 1E-265 | 4E-263 | 0.8020833 | n.a. |
| Eqca-DQB1 | 2.492 | 10.925 | 1223.14 | 2E-263 | 8E-261 | 1 | n.a. |
| LY6E | 3.169 | 10.241 | 1205.29 | 1E-259 | 4E-257 | 0.9791667 | surface |
| CLNK | 3.207 | 9.2461 | 6172.83 | 2E-241 | 5E-239 | 0.7552083 | n.a. |
| LRRK2 | 3.097 | 9.2707 | 1828.44 | 1E-203 | 3E-201 | 0.8802083 | n.a. |
| PKIB | 2.617 | 9.2759 | 2030.64 | 7E-203 | 2E-200 | 0.8854167 | n.a. |
| ENSECAG00000019318 | 2.331 | 9.3455 | 1070.99 | 1E-187 | 4E-185 | 0.9166667 | n.a. |
| CLIC2 | 2.472 | 9.286 | 2411.65 | 1E-186 | 3E-184 | 0.734375 | n.a. |
| ENSECAG00000035094 | 2.204 | 9.2863 | 1222.2 | 1E-176 | 3E-174 | 0.8541667 | n.a. |
| RAB38 | 2.968 | 9.2603 | 2798.24 | 2E-169 | 6E-167 | 0.8802083 | n.a. |
| RPS8 | 1.124 | 12.9 | 727.001 | 2E-158 | 4E-156 | 1 | n.a. |
| RPS27A | 1.223 | 12.702 | 718.616 | 1E-156 | 2E-154 | 1 | n.a. |
| ENSECAG00000035439 | 2.433 | 9.2665 | 4161 | 2E-144 | 5E-142 | 0.6041667 | n.a. |
| RGS1 | 2.559 | 9.3256 | 660.226 | 3E-144 | 6E-142 | 0.7708333 | n.a. |
| RPL35 | 1.076 | 12.908 | 655.225 | 4E-143 | 7E-141 | 1 | n.a. |
| VAC14 | 2.624 | 9.2854 | 734.622 | 1E-142 | 3E-140 | 0.84375 | n.a. |
| RF00324 | 1.047 | 12.723 | 629.622 | 1E-137 | 2E-135 | 1 | n.a. |
| ENSECAG00000005400 | 2.771 | 9.2385 | 21637 | 5E-137 | 1E-134 | 0.6614583 | n.a. |
| NAAA | 2.342 | 9.5818 | 623.68 | 2E-136 | 4E-134 | 0.9479167 | n.a. |
| SPP1 | 2.745 | 9.2475 | 2219.29 | 4E-136 | 7E-134 | 0.5572917 | n.a. |
| CD8A | 2.126 | 9.4281 | 619.266 | 2E-135 | 3E-133 | 0.71875 | surface |
| RPL14 | 1.01 | 12.868 | 596.149 | 1E-130 | 2E-128 | 1 | n.a. |
| RPS18 | 1 | 12.945 | 571.008 | 3E-125 | 6E-123 | 1 | n.a. |
| RUBCNL | 2.16 | 9.2489 | 1979.69 | 2E-120 | 2E-118 | 0.609375 | n.a. |
| HLA-DOA | 2.143 | 9.3227 | 901.66 | 2E-115 | 2E-113 | 0.78125 | surface |
| CSRP1 | 2.153 | 9.2812 | 600.861 | 2E-113 | 3E-111 | 0.8645833 | n.a. |
| ENSECAG00000033359 | 2.253 | 9.258 | 2050.78 | 9E-110 | 1E-107 | 0.5416667 | n.a. |
| BTLA | 1.962 | 9.2816 | 1358.2 | 2E-109 | 2E-107 | 0.546875 | surface |
| NEURL1 | 2.324 | 9.2404 | 4836.85 | 2E-109 | 3E-107 | 0.6458333 | n.a. |
| RAB7B | 2.526 | 9.2391 | 5107.66 | 2E-108 | 2E-106 | 0.6770833 | n.a. |
| PVRIG | 2.026 | 9.5646 | 488.716 | 2E-107 | 2E-105 | 0.8229167 | n.a. |
| ENSECAG00000028889 | 2.238 | 9.2871 | 1177.34 | 7E-107 | 9E-105 | 0.75 | n.a. |
| CST7 | 2.247 | 9.312 | 957.594 | 3E-103 | 3E-101 | 0.640625 | n.a. |
| PID1 | 1.963 | 9.2473 | 3699.05 | 2E-98 | 2.1E-96 | 0.5416667 | n.a. |
| ENSECAG00000031496 | 2.097 | 9.3827 | 445.37 | 3E-98 | 3.5E-96 | 0.8333333 | n.a. |

| | | | | | | | |
|---|---|---|---|---|---|---|---|
| S100A11 | 2.031 | 9.847 | 440.306 | 4E-97 | 4.2E-95 | 1 | n.a. |
| WDFY4 | 2.193 | 9.2866 | 827.619 | 2E-96 | 2.3E-94 | 0.7760417 | n.a. |
| ENSECAG00000034569 | 1.634 | 9.7713 | 419.444 | 1E-92 | 1.2E-90 | 0.5416667 | n.a. |
| DCTPP1 | 2.165 | 9.3868 | 406.268 | 8E-90 | 8E-88 | 0.90625 | n.a. |
| PABPC1 | 1.185 | 10.847 | 397.07 | 7E-88 | 7.4E-86 | 1 | n.a. |
| CADM1 | 2.294 | 9.2459 | 2619.58 | 2E-87 | 1.6E-85 | 0.6770833 | surface |
| AIF1 | 1.272 | 9.4438 | 597.955 | 2E-84 | 1.9E-82 | 0.9739583 | n.a. |
| CFP | 1.33 | 9.4343 | 1077.3 | 1E-82 | 8.9E-81 | 0.9583333 | n.a. |
| S100A10 | 1.491 | 10.866 | 361.282 | 4E-80 | 3.3E-78 | 1 | n.a. |
| FLT3 | 2.174 | 9.2499 | 1362.56 | 1E-78 | 9E-77 | 0.7708333 | surface |
| CRHBP | 1.949 | 9.2452 | 1999.09 | 7E-78 | 6.3E-76 | 0.390625 | n.a. |
| SLAMF7 | 1.694 | 9.2845 | 531.566 | 2E-76 | 1.8E-74 | 0.7447917 | surface |
| TSTD1 | 1.725 | 9.2702 | 700.266 | 4E-72 | 3.3E-70 | 0.5625 | n.a. |
| ENSECAG00000021796 | 1.659 | 9.6911 | 319.336 | 4E-71 | 3.4E-69 | 0.9739583 | n.a. |
| TCTN2 | 1.96 | 9.2411 | 2782.69 | 3E-70 | 2.2E-68 | 0.453125 | surface |
| SNX22 | 1.906 | 9.2475 | 1471.07 | 5E-70 | 3.9E-68 | 0.484375 | n.a. |
| FOS | 1.411 | 9.3156 | 411.344 | 3E-65 | 2.1E-63 | 0.671875 | n.a. |
| ENSECAG00000028100 | 2.043 | 9.2366 | 3901.91 | 5E-65 | 3.3E-63 | 0.4166667 | n.a. |
| ENSECAG00000019029 | 1.692 | 9.9278 | 274.466 | 2E-61 | 1.4E-59 | 0.7291667 | n.a. |
| HLA-DQB1 | 1.434 | 9.2656 | 604.813 | 5E-61 | 3.1E-59 | 0.28125 | surface |
| SORBS3 | 1.514 | 9.2441 | 2142.52 | 1E-60 | 7.8E-59 | 0.3489583 | n.a. |
| ENSECAG00000039164 | 1.762 | 9.2362 | 6307.94 | 1E-58 | 9E-57 | 0.2916667 | n.a. |
| SLAMF8 | 1.923 | 9.2377 | 2215.65 | 7E-58 | 4.4E-56 | 0.4739583 | surface |
| C1QBP | 1.446 | 9.3874 | 246.877 | 2E-55 | 1.1E-53 | 0.78125 | n.a. |
| PLPP1 | 1.72 | 9.2503 | 1065.84 | 4E-54 | 2.1E-52 | 0.4270833 | n.a. |
| ENSECAG00000020136 | 1.352 | 9.7014 | 237.83 | 2E-53 | 1E-51 | 0.984375 | n.a. |
| LPAR6 | 1.773 | 9.2687 | 841.229 | 3E-53 | 1.6E-51 | 0.671875 | surface |
| ATP6V0A2 | 1.834 | 9.2479 | 976.718 | 6E-52 | 3.3E-50 | 0.5520833 | surface |
| PAK1 | 1.294 | 9.331 | 625.31 | 8E-52 | 4.5E-50 | 0.9114583 | n.a. |
| ENSECAG00000006701 | 1.66 | 9.2359 | 2111.25 | 9E-51 | 4.9E-49 | 0.3385417 | n.a. |
| PECAM1 | 1.206 | 9.2688 | 892.66 | 2E-49 | 8.7E-48 | 0.4010417 | surface |
| FBL | 1.577 | 9.7416 | 214.519 | 2E-48 | 1E-46 | 0.96875 | n.a. |
| TUBA1A | 1.46 | 10.091 | 211.677 | 8E-48 | 4.1E-46 | 0.90625 | n.a. |
| ENSECAG00000012400 | 1.479 | 9.3413 | 211.758 | 2E-47 | 9.2E-46 | 0.4375 | n.a. |
| NEXN | 1.607 | 9.2383 | 3957.19 | 7E-47 | 3.6E-45 | 0.3489583 | n.a. |
| LIMCH1 | 1.485 | 9.2424 | 1644.05 | 2E-46 | 7.9E-45 | 0.3645833 | n.a. |
| MPEG1 | 1.28 | 9.3534 | 540.058 | 6E-46 | 2.9E-44 | 0.875 | surface |
| DPP4 | 1.169 | 9.2884 | 465.781 | 2E-45 | 9E-44 | 0.3645833 | surface |
| PDF | 1.671 | 9.2907 | 306.475 | 6E-45 | 2.9E-43 | 0.5989583 | n.a. |
| PTPRCAP | 1.054 | 10.617 | 197.455 | 1E-44 | 4.6E-43 | 0.8802083 | n.a. |
| SLC46A3 | 1.675 | 9.2637 | 386.701 | 1E-44 | 5E-43 | 0.609375 | surface |
| ANXA3 | 1.484 | 9.2812 | 479.953 | 1E-44 | 5E-43 | 0.5364583 | n.a. |
| FAM135A | 1.584 | 9.2372 | 3837.86 | 5E-44 | 2.2E-42 | 0.3489583 | n.a. |
| BCL2A1 | 1.17 | 9.2926 | 499.784 | 2E-43 | 7.2E-42 | 0.34375 | n.a. |
| ENSECAG00000015275 | 1.572 | 9.3043 | 362.339 | 2E-42 | 6.8E-41 | 0.5729167 | n.a. |
| VIPR1 | 1.457 | 9.2359 | 4238.74 | 4E-42 | 1.8E-40 | 0.3177083 | surface |
| ENSECAG00000031154 | 1.292 | 9.2755 | 642.242 | 1E-41 | 5.6E-40 | 0.2760417 | n.a. |
| ARHGEF9 | 1.339 | 9.2412 | 1767.76 | 2E-41 | 7.9E-40 | 0.2916667 | n.a. |
| HLA-DMA | 1.294 | 9.6051 | 182.171 | 2E-41 | 8.6E-40 | 0.9427083 | surface |
| RAB30 | 1.189 | 9.2526 | 1061.19 | 4E-41 | 1.7E-39 | 0.3333333 | n.a. |
| AIG1 | 1.443 | 9.327 | 262.486 | 2E-40 | 8.4E-39 | 0.8385417 | n.a. |
| CCND1 | 1.402 | 9.2529 | 822.615 | 1E-39 | 4.5E-38 | 0.375 | n.a. |
| FGD2 | 1.562 | 9.2999 | 423.213 | 2E-39 | 9E-38 | 0.6770833 | n.a. |
| PPM1J | 1.534 | 9.2407 | 1862.21 | 1E-38 | 4.6E-37 | 0.453125 | n.a. |
| SPCS1 | 1.222 | 9.9456 | 166.849 | 4E-38 | 1.7E-36 | 0.96875 | n.a. |
| PTGER4 | 1.294 | 9.3476 | 178.389 | 5E-38 | 1.8E-36 | 0.5677083 | surface |

| | | | | | | | |
|---|---|---|---|---|---|---|---|
| NDUFB4 | 1.49 | 9.5448 | 166.214 | 6E-38 | 2.3E-36 | 0.9270833 | n.a. |
| PPA1 | 1.509 | 9.4873 | 165.061 | 1E-37 | 4.1E-36 | 0.84375 | n.a. |
| ENSECAG00000032068 | 1.309 | 9.2351 | 6369.38 | 2E-37 | 6.9E-36 | 0.2760417 | n.a. |
| SERPINB9 | 1.175 | 9.3883 | 242.658 | 1E-36 | 3.8E-35 | 0.4375 | n.a. |
| RGS10 | 1.413 | 9.5903 | 159.445 | 2E-36 | 6.5E-35 | 0.921875 | n.a. |
| TRIO | 1.526 | 9.2396 | 1561.6 | 4E-36 | 1.3E-34 | 0.40625 | n.a. |
| CSTA | 1.132 | 9.2431 | 1377.38 | 2E-35 | 5.8E-34 | 0.5104167 | n.a. |
| DHCR24 | 1.081 | 9.3251 | 218.953 | 3E-35 | 1E-33 | 0.5260417 | n.a. |
| CD2 | 1.145 | 9.8338 | 153.571 | 3E-35 | 1.2E-33 | 0.4479167 | surface |
| VPS26B | 1.409 | 9.3122 | 172.778 | 7E-35 | 2.4E-33 | 0.6927083 | n.a. |
| SRP14 | 1.026 | 10.144 | 152.09 | 7E-35 | 2.4E-33 | 1 | n.a. |
| ENSECAG00000038794 | 1.117 | 9.2633 | 1010.37 | 1E-34 | 4.5E-33 | 0.5052083 | n.a. |
| ENSECAG00000023388 | 1.599 | 9.2512 | 1070.67 | 4E-33 | 1.3E-31 | 0.546875 | n.a. |
| TXN | 1.043 | 9.9313 | 143.441 | 5E-33 | 1.8E-31 | 0.984375 | surface |
| SMCO4 | 1.222 | 9.5303 | 141.326 | 2E-32 | 5.1E-31 | 0.6510417 | n.a. |
| GNL1 | 1.421 | 9.2852 | 227.122 | 2E-32 | 5.6E-31 | 0.578125 | n.a. |
| CA12 | 1.356 | 9.2359 | 2699.47 | 2E-32 | 6.2E-31 | 0.2760417 | surface |
| TSPAN2 | 1.183 | 9.2666 | 405.049 | 2E-31 | 7.8E-30 | 0.34375 | surface |
| SGK1 | 1.123 | 9.2522 | 449.73 | 2E-30 | 5E-29 | 0.2760417 | n.a. |
| SMAGP | 1.197 | 9.3065 | 188.148 | 4E-30 | 1.1E-28 | 0.4375 | n.a. |
| TMEM145 | 1.317 | 9.2908 | 277.81 | 4E-30 | 1.2E-28 | 0.3489583 | surface |
| HSP90AB1 | 1.181 | 10.085 | 129.061 | 7E-30 | 2.2E-28 | 0.96875 | n.a. |
| SH3BP1 | 1.193 | 9.4224 | 127.642 | 2E-29 | 4.5E-28 | 0.8177083 | n.a. |
| GCSH | 1.323 | 9.4155 | 122.701 | 2E-28 | 5.2E-27 | 0.7239583 | n.a. |
| MGMT | 1.296 | 9.3322 | 140.171 | 4E-28 | 1E-26 | 0.6145833 | n.a. |
| PPM1M | 1.167 | 9.3461 | 118.221 | 2E-27 | 4.8E-26 | 0.7291667 | n.a. |
| SLC39A10 | 1.204 | 9.3738 | 122.701 | 3E-27 | 8E-26 | 0.59375 | surface |
| C4orf48 | 1.105 | 9.5904 | 117.633 | 4E-27 | 1.2E-25 | 0.546875 | n.a. |
| ENSECAG00000013303 | 1.095 | 9.4002 | 135.664 | 6E-27 | 1.7E-25 | 0.8229167 | n.a. |
| ID3 | 1.192 | 9.6697 | 115.53 | 7E-27 | 1.8E-25 | 0.34375 | n.a. |
| GSTM3 | 1.112 | 9.328 | 167.149 | 4E-26 | 1E-24 | 0.3385417 | n.a. |
| CPNE3 | 1.095 | 9.3269 | 108.059 | 3E-25 | 7.3E-24 | 0.7083333 | n.a. |
| IL21R | 1.068 | 9.3399 | 196.714 | 5E-25 | 1.3E-23 | 0.3020833 | surface |
| TMEM230 | 1.123 | 9.3512 | 105.571 | 1E-24 | 2.5E-23 | 0.6041667 | n.a. |
| ENSECAG00000032128 | 1.03 | 9.2585 | 385.59 | 1E-24 | 2.8E-23 | 0.2864583 | n.a. |
| FMNL2 | 1.054 | 9.2378 | 1784.6 | 2E-24 | 4E-23 | 0.2708333 | n.a. |
| PIK3CB | 1.384 | 9.2443 | 677.883 | 2E-24 | 5.7E-23 | 0.4427083 | n.a. |
| SPINT2 | 1.222 | 9.2841 | 235.676 | 1E-23 | 2.8E-22 | 0.5 | surface |
| ZNF385A | 1.365 | 9.2523 | 662.323 | 5E-23 | 1.1E-21 | 0.5260417 | n.a. |
| TLR10 | 1.039 | 9.286 | 245.421 | 8E-23 | 1.7E-21 | 0.3385417 | surface |
| TBC1D9 | 1.028 | 9.2457 | 575.327 | 3E-22 | 6.4E-21 | 0.359375 | n.a. |
| COPRS | 1.245 | 9.2687 | 239.298 | 5E-22 | 1E-20 | 0.4375 | n.a. |
| ENSECAG00000009733 | 1.116 | 9.3278 | 126.41 | 2E-21 | 3.5E-20 | 0.4375 | n.a. |
| MYOF | 1.073 | 9.247 | 501.614 | 3E-21 | 5.6E-20 | 0.3958333 | surface |
| ENSECAG00000038085 | 1.062 | 9.9701 | 87.4424 | 9E-21 | 2E-19 | 0.9635417 | n.a. |
| C20H6orf62 | 1.025 | 9.3936 | 86.0241 | 2E-20 | 3.9E-19 | 0.8645833 | n.a. |
| PLA2G16 | 1.006 | 9.6144 | 85.4552 | 3E-20 | 5.2E-19 | 0.9739583 | n.a. |
| FCHSD2 | 1.082 | 9.2541 | 341.471 | 4E-20 | 7.4E-19 | 0.2916667 | n.a. |
| ZNF664 | 1.12 | 9.2525 | 374.212 | 2E-19 | 3.4E-18 | 0.2708333 | n.a. |
| NAV1 | 1.082 | 9.2705 | 203.19 | 2E-17 | 4.1E-16 | 0.4895833 | n.a. |
| ST3GAL2 | 1.118 | 9.2628 | 185.645 | 3E-17 | 5.9E-16 | 0.421875 | n.a. |
| GRHPR | 1.001 | 9.3044 | 93.9204 | 8E-17 | 1.4E-15 | 0.5520833 | n.a. |
| NPM3 | 1.015 | 9.6322 | 69.4203 | 8E-17 | 1.5E-15 | 0.8177083 | n.a. |
| ENSECAG00000033970 | 1.001 | 9.6358 | 68.6677 | 1E-16 | 2.1E-15 | 0.8645833 | n.a. |
| RNPEP | 1.186 | 9.2739 | 154.374 | 2E-16 | 2.8E-15 | 0.484375 | n.a. |
| RHOC | 1.03 | 9.2833 | 160.001 | 6E-15 | 9.8E-14 | 0.4375 | n.a. |

| | | | | | | | |
|---|---|---|---|---|---|---|---|
| NISCH | 1.003 | 9.2555 | 197.268 | 9E-15 | 1.5E-13 | 0.2864583 | n.a. |
| CRLF1 | 1.159 | 9.249 | 354.507 | 1E-12 | 1.4E-11 | 0.4375 | n.a. |

| genes | logFC | logCPM | F | PValue | FDR | percent.exp | Surfacome.Label |
|---|---|---|---|---|---|---|---|
| ENSECAG00000003702 | 6.382 | 9.282 | 15784.5 | 0 | 0 | 1 | n.a. |
| ENSECAG00000017313 | 6.24 | 9.2752 | 21049.7 | 0 | 0 | 1 | n.a. |
| ENSECAG00000001923 | 5.307 | 9.6006 | 3032.8 | 0 | 0 | 1 | n.a. |
| SOD1 | 5.253 | 9.8391 | 2629.18 | 0 | 0 | 1 | n.a. |
| JCHAIN | 4.871 | 9.7044 | 5090.05 | 0 | 0 | 0.8108108 | n.a. |
| FCRLA | 4.534 | 9.3304 | 4301.97 | 0 | 0 | 0.9369369 | n.a. |
| ENSECAG00000022644 | 4.416 | 9.2487 | 7826.68 | 0 | 0 | 1 | n.a. |
| CYB561A3 | 4.349 | 9.3685 | 2261.91 | 0 | 0 | 0.981982 | n.a. |
| IRF7 | 4.333 | 9.3097 | 3450.52 | 0 | 0 | 0.9369369 | n.a. |
| TCF4 | 4.018 | 9.3454 | 3099.1 | 0 | 0 | 0.9279279 | n.a. |
| CYSTM1 | 3.29 | 9.3828 | 1627.18 | 0 | 0 | 0.990991 | n.a. |
| ENSECAG00000008542 | 1.609 | 14.412 | 2782.19 | 0 | 0 | 1 | n.a. |
| ENSECAG00000026905 | 1.493 | 13.117 | 1718.81 | 0 | 0 | 1 | n.a. |
| IRF8 | 3.532 | 9.303 | 1755.98 | 1E-291 | 5E-289 | 0.9459459 | n.a. |
| POLD1 | 3.886 | 9.3555 | 2532.76 | 4E-287 | 2E-284 | 0.8108108 | n.a. |
| CTSC | 4.101 | 9.6133 | 1335.51 | 6E-287 | 2E-284 | 0.972973 | n.a. |
| UGCG | 3.846 | 9.3115 | 1235.72 | 5E-266 | 2E-263 | 0.9279279 | n.a. |
| ENSECAG00000019875 | 3.509 | 9.2924 | 2391.09 | 5E-236 | 2E-233 | 0.8738739 | n.a. |
| ENSECAG00000035361 | 3.611 | 9.2396 | 10608.6 | 1E-217 | 4E-215 | 0.7747748 | n.a. |
| EEF1B2 | 1.702 | 12.225 | 999.028 | 3E-216 | 1E-213 | 1 | n.a. |
| ATP13A2 | 3.658 | 9.2517 | 2523.2 | 7E-215 | 2E-212 | 0.8648649 | surface |
| ENSECAG00000015055 | 3.532 | 9.2599 | 2823.09 | 8E-212 | 2E-209 | 0.5945946 | n.a. |
| CCDC50 | 3.314 | 9.3095 | 1624.76 | 3E-203 | 8E-201 | 0.7747748 | n.a. |
| ADORA2B | 3.439 | 9.2559 | 2244.78 | 1E-195 | 3E-193 | 0.8018018 | surface |
| ENSECAG00000039088 | 3.011 | 9.7274 | 861.638 | 4E-187 | 1E-184 | 0.981982 | n.a. |
| ENSECAG00000038338 | 3.318 | 9.2755 | 2842.99 | 6E-186 | 2E-183 | 0.6576577 | n.a. |
| HEBP2 | 3.255 | 9.2913 | 1590.89 | 1E-184 | 3E-182 | 0.7027027 | n.a. |
| ENSECAG00000030777 | 2.606 | 9.3773 | 1886.15 | 1E-180 | 3E-178 | 0.9009009 | n.a. |
| RPS20 | 1.266 | 13.671 | 822.47 | 9E-179 | 2E-176 | 1 | n.a. |
| DUSP5 | 2.886 | 9.267 | 1864.17 | 1E-175 | 3E-173 | 0.6666667 | n.a. |
| ENSECAG00000021212 | 3.418 | 9.2394 | 5349.12 | 1E-174 | 2E-172 | 0.6126126 | n.a. |
| MFAP2 | 3.265 | 9.2367 | 17877.2 | 3E-172 | 5E-170 | 0.4954955 | n.a. |
| RAB24 | 2.872 | 9.4289 | 785.141 | 7E-171 | 1E-168 | 0.963964 | n.a. |
| ENSECAG00000032253 | 3.333 | 9.2484 | 3544.3 | 2E-168 | 3E-166 | 0.5585586 | n.a. |
| SYNGR1 | 3.344 | 9.2464 | 2907.51 | 4E-168 | 8E-166 | 0.7297297 | n.a. |
| BLNK | 3.204 | 9.2737 | 2421.97 | 1E-166 | 3E-164 | 0.7657658 | n.a. |
| ENSECAG00000029287 | 2.838 | 10.043 | 742.077 | 1E-161 | 2E-159 | 0.8018018 | n.a. |
| EPCAM | 2.857 | 9.2602 | 2581.43 | 2E-158 | 3E-156 | 0.5585586 | surface |
| ENSECAG00000010436 | 3.359 | 9.2883 | 1217.93 | 4E-158 | 6E-156 | 0.7207207 | n.a. |
| RPS27 | 1.045 | 13.096 | 680.965 | 1E-148 | 2E-146 | 1 | n.a. |
| ENSECAG00000019029 | 3.199 | 9.9278 | 677.875 | 5E-148 | 8E-146 | 0.8828829 | n.a. |
| CNPY3 | 2.911 | 9.4881 | 657.006 | 1E-143 | 2E-141 | 1 | n.a. |
| MCOLN2 | 3.004 | 9.2516 | 3131.24 | 1E-139 | 2E-137 | 0.6486486 | n.a. |
| ENSECAG00000031985 | 2.936 | 9.238 | 8101.24 | 1E-128 | 2E-126 | 0.5225225 | n.a. |
| NUCB2 | 3 | 9.3402 | 664.537 | 5E-121 | 7E-119 | 0.7387387 | n.a. |
| PTPRCAP | 2.306 | 10.617 | 549.723 | 1E-120 | 2E-118 | 0.972973 | n.a. |
| PROCR | 2.755 | 9.236 | 11803.2 | 2E-120 | 2E-118 | 0.4234234 | surface |
| ST6GALNAC4 | 2.847 | 9.2959 | 672.601 | 4E-120 | 6E-118 | 0.8378378 | n.a. |
| DIPK2A | 2.91 | 9.2586 | 1431.26 | 9E-119 | 1E-116 | 0.7387387 | n.a. |
| ENSECAG00000031496 | 2.746 | 9.3827 | 540.675 | 1E-118 | 2E-116 | 0.8108108 | n.a. |
| FAIM | 2.84 | 9.2661 | 1324.37 | 1E-116 | 2E-114 | 0.6036036 | surface |
| IFNAR1 | 2.641 | 9.352 | 511.028 | 2E-112 | 3E-110 | 0.8648649 | surface |
| FYB1 | 2.654 | 9.6072 | 501.04 | 3E-110 | 4E-108 | 0.9369369 | n.a. |
| ENSECAG00000031522 | 3.133 | 10.189 | 496.409 | 3E-109 | 4E-107 | 0.5765766 | n.a. |
| MPEG1 | 2.234 | 9.3534 | 1450.12 | 2E-108 | 2E-106 | 0.9009009 | surface |

| | | | | | | | |
|---|---|---|---|---|---|---|---|
| CERS6 | 2.845 | 9.2532 | 1585.64 | 1E-107 | 2E-105 | 0.5855856 | n.a. |
| GAPT | 2.561 | 9.2786 | 1212.71 | 6E-107 | 7E-105 | 0.5945946 | n.a. |
| YWHAQ | 2.851 | 9.7075 | 485.791 | 6E-107 | 8E-105 | 0.9279279 | n.a. |
| SLC38A9 | 2.792 | 9.2699 | 878.295 | 1E-106 | 1E-104 | 0.7027027 | surface |
| PPP2R3C | 2.756 | 9.2949 | 623.891 | 3E-104 | 3E-102 | 0.6576577 | n.a. |
| AQP3 | 2.564 | 9.3499 | 756.896 | 2E-101 | 2E-99 | 0.5495495 | n.a. |
| ENSECAG00000002390 | 2.939 | 9.2531 | 1756.38 | 3E-101 | 4E-99 | 0.5315315 | n.a. |
| PPP1R14A | 2.585 | 9.2419 | 5076.63 | 2E-98 | 2.1E-96 | 0.4594595 | n.a. |
| MS4A1 | 2.454 | 9.7256 | 989.359 | 5E-98 | 5.1E-96 | 0.3963964 | surface |
| SRM | 2.791 | 9.2691 | 929.009 | 1E-97 | 1.1E-95 | 0.5495495 | n.a. |
| FAM177A1 | 2.779 | 9.2748 | 849.879 | 5E-96 | 4.8E-94 | 0.5945946 | n.a. |
| LGALS1 | 1.936 | 11.109 | 424.854 | 8E-94 | 8E-92 | 1 | n.a. |
| ENSECAG00000024850 | 2.423 | 9.2353 | 17539.1 | 2E-92 | 1.5E-90 | 0.4504505 | n.a. |
| SNX9 | 2.727 | 9.271 | 796.835 | 2E-92 | 1.8E-90 | 0.7387387 | n.a. |
| FAM129C | 2.68 | 9.2412 | 3897.56 | 7E-91 | 6.7E-89 | 0.5495495 | n.a. |
| SDHB | 2.379 | 9.4571 | 397.341 | 7E-88 | 6.3E-86 | 0.954955 | n.a. |
| ACP5 | 2.531 | 9.7992 | 393.961 | 3E-87 | 3.3E-85 | 0.8918919 | n.a. |
| IL10RB | 2.233 | 9.6008 | 388.751 | 5E-86 | 4.3E-84 | 1 | surface |
| GOT1 | 2.637 | 9.2894 | 635.451 | 2E-84 | 1.4E-82 | 0.6576577 | n.a. |
| C1QBP | 2.277 | 9.3874 | 431.337 | 3E-84 | 2.6E-82 | 0.7837838 | n.a. |
| FCMR | 2.241 | 9.3335 | 737.062 | 3E-84 | 3E-82 | 0.4234234 | n.a. |
| SRP14 | 1.801 | 10.144 | 359.165 | 1E-79 | 9.5E-78 | 0.990991 | n.a. |
| SMIM5 | 2.457 | 9.2402 | 2606.13 | 6E-78 | 4.7E-76 | 0.4324324 | n.a. |
| SLC44A2 | 2.222 | 9.4299 | 348.344 | 2E-77 | 2E-75 | 0.7027027 | surface |
| PLAC8B | 1.647 | 11.113 | 348.074 | 3E-77 | 2.2E-75 | 0.990991 | n.a. |
| L3MBTL3 | 2.493 | 9.2638 | 750.744 | 8E-77 | 6.8E-75 | 0.6486486 | n.a. |
| TFPT | 2.471 | 9.2805 | 537.299 | 3E-76 | 2.7E-74 | 0.7207207 | n.a. |
| ENSECAG00000013303 | 2.109 | 9.4002 | 345.754 | 4E-76 | 3E-74 | 0.9009009 | n.a. |
| SLA2 | 2.386 | 9.2804 | 969.726 | 6E-75 | 4.7E-73 | 0.4324324 | n.a. |
| AKAP9 | 2.439 | 9.4493 | 331.47 | 1E-73 | 8E-72 | 0.8468468 | n.a. |
| PACSIN1 | 2.172 | 9.2564 | 3030.4 | 2E-72 | 1.7E-70 | 0.3873874 | n.a. |
| CILP | 2.055 | 9.2349 | 14055.2 | 1E-71 | 1E-69 | 0.3063063 | n.a. |
| TNFSF13 | 2.46 | 9.266 | 1914.91 | 1E-70 | 8.1E-69 | 0.6936937 | surface |
| MGAT1 | 2.434 | 9.3346 | 385.719 | 3E-70 | 2E-68 | 0.7387387 | n.a. |
| ENSECAG00000034000 | 1.991 | 9.5622 | 311.072 | 3E-69 | 1.9E-67 | 0.972973 | n.a. |
| CD69 | 2.292 | 9.4382 | 438.282 | 2E-68 | 1.6E-66 | 0.4144144 | surface |
| C1QTNF1 | 2.128 | 9.2395 | 3477.83 | 5E-67 | 3.6E-65 | 0.3423423 | n.a. |
| VLDLR | 2.18 | 9.2352 | 2207.53 | 5E-67 | 3.9E-65 | 0.3693694 | surface |
| ATP1B1 | 2.063 | 9.2383 | 4405.4 | 6E-67 | 4.5E-65 | 0.3333333 | surface |
| CALY | 2.155 | 9.2778 | 1027.68 | 1E-65 | 8.4E-64 | 0.3963964 | surface |
| ITM2C | 2.14 | 9.5994 | 292.64 | 2E-65 | 1.7E-63 | 0.8468468 | surface |
| FAM107B | 2.168 | 9.5005 | 292.221 | 3E-65 | 2.1E-63 | 0.6936937 | n.a. |
| RAC2 | 1.456 | 10.594 | 291.21 | 5E-65 | 3.4E-63 | 0.972973 | n.a. |
| CASP7 | 2.148 | 9.2925 | 548.378 | 1E-62 | 8.4E-61 | 0.4774775 | n.a. |
| RF00577 | 1.27 | 10.913 | 277.771 | 4E-62 | 2.7E-60 | 0.9459459 | n.a. |
| PAXX | 2.218 | 9.3487 | 303.159 | 2E-61 | 1.4E-59 | 0.7027027 | n.a. |
| HSP90B1 | 1.983 | 9.5518 | 269.213 | 3E-60 | 1.8E-58 | 0.990991 | n.a. |
| DHCR7 | 2.308 | 9.2548 | 961.978 | 7E-60 | 4.7E-58 | 0.4234234 | n.a. |
| ARRDC5 | 2.136 | 9.2365 | 5976.58 | 1E-59 | 7E-58 | 0.3063063 | n.a. |
| BCL11A | 2.305 | 9.2593 | 1249.4 | 3E-59 | 1.9E-57 | 0.4594595 | n.a. |
| CCS | 2.171 | 9.3726 | 262.592 | 9E-59 | 5.4E-57 | 0.7927928 | n.a. |
| CLEC12A | 1.417 | 9.4082 | 667.346 | 1E-58 | 5.9E-57 | 0.9189189 | surface |
| ENSECAG00000006065 | 1.486 | 10.545 | 261.749 | 1E-58 | 7E-57 | 0.972973 | n.a. |
| RPS21 | 1.093 | 11.698 | 260.423 | 2E-58 | 1.4E-56 | 1 | n.a. |
| TP53I13 | 2.111 | 9.2455 | 1798.9 | 5E-58 | 3E-56 | 0.3693694 | surface |
| IGHM | 1.838 | 9.6407 | 258.185 | 7E-58 | 4.1E-56 | 0.3873874 | n.a. |

| | | | | | | | |
|---|---|---|---|---|---|---|---|
| RPS27L | 1.806 | 9.8828 | 247.514 | 1E-55 | 8.2E-54 | 1 | n.a. |
| ENSECAG00000035141 | 2.225 | 9.2632 | 1597.28 | 3E-55 | 1.4E-53 | 0.3963964 | n.a. |
| CYB5A | 2.3 | 9.3033 | 482.883 | 5E-55 | 2.8E-53 | 0.4504505 | n.a. |
| LAMP3 | 2.067 | 9.2517 | 1089.2 | 9E-55 | 5.3E-53 | 0.4324324 | surface |
| TIGIT | 1.988 | 9.2485 | 1836.89 | 1E-54 | 5.7E-53 | 0.3783784 | surface |
| ENSECAG00000010622 | 1.057 | 12.221 | 242.09 | 2E-54 | 1.2E-52 | 1 | n.a. |
| STX7 | 1.782 | 9.3568 | 244.699 | 3E-53 | 1.5E-51 | 0.7927928 | n.a. |
| ADGRG5 | 2.212 | 9.2595 | 1107.1 | 3E-53 | 1.6E-51 | 0.5855856 | n.a. |
| BEX3 | 2.116 | 9.4183 | 309.483 | 1E-52 | 7.8E-51 | 0.5315315 | n.a. |
| BICDL2 | 2.019 | 9.2439 | 1879.39 | 2E-52 | 1.3E-50 | 0.3513514 | n.a. |
| PDCD4 | 2.13 | 9.6109 | 230.317 | 7E-52 | 4E-50 | 0.9009009 | n.a. |
| ENSECAG00000035548 | 1.932 | 9.2356 | 6234.26 | 1E-51 | 6.4E-50 | 0.2702703 | n.a. |
| CD164 | 2.044 | 9.4651 | 228.525 | 2E-51 | 9.5E-50 | 0.8738739 | surface |
| CD2AP | 2.117 | 9.2797 | 419.257 | 3E-51 | 1.4E-49 | 0.5945946 | n.a. |
| GOLIM4 | 2.199 | 9.26 | 580.588 | 3E-51 | 1.7E-49 | 0.5945946 | n.a. |
| ME3 | 2.005 | 9.2599 | 805.247 | 1E-50 | 6.7E-49 | 0.4054054 | n.a. |
| CTSS | 1.198 | 9.8161 | 224.513 | 1E-50 | 6.7E-49 | 0.972973 | n.a. |
| ENSECAG00000006595 | 1.362 | 9.2598 | 1184.03 | 1E-49 | 7.1E-48 | 0.4414414 | n.a. |
| SMCO4 | 2 | 9.5303 | 219.058 | 2E-49 | 1E-47 | 0.6756757 | n.a. |
| HM13 | 1.831 | 9.3911 | 216.261 | 8E-49 | 4.1E-47 | 0.8648649 | surface |
| RASL11B | 1.756 | 9.2351 | 8075.33 | 1E-48 | 6.9E-47 | 0.2522523 | n.a. |
| UPF3B | 1.957 | 9.3746 | 214.873 | 2E-48 | 8E-47 | 0.7657658 | n.a. |
| ENSECAG00000038714 | 1.975 | 9.3099 | 437.916 | 4E-48 | 1.8E-46 | 0.7207207 | n.a. |
| RNF5 | 1.977 | 9.3876 | 212.907 | 4E-48 | 2.1E-46 | 0.7927928 | n.a. |
| RPL22 | 1.086 | 11.43 | 212.56 | 5E-48 | 2.5E-46 | 1 | n.a. |
| RRBP1 | 2.034 | 9.2913 | 467.154 | 2E-47 | 1E-45 | 0.7027027 | n.a. |
| SLC25A4 | 2.006 | 9.2755 | 641.493 | 2E-47 | 1.1E-45 | 0.4234234 | n.a. |
| CD1B1 | 1.819 | 9.2551 | 1507.08 | 3E-47 | 1.3E-45 | 0.3513514 | n.a. |
| TCHP | 2.089 | 9.2494 | 875.476 | 3E-47 | 1.5E-45 | 0.4054054 | n.a. |
| PRDX2 | 1.877 | 9.2964 | 703.579 | 1E-46 | 4.4E-45 | 0.2612613 | n.a. |
| RNASEH2B | 2.016 | 9.3711 | 248.039 | 3E-46 | 1.3E-44 | 0.7927928 | n.a. |
| PECAM1 | 1.681 | 9.2688 | 1087.21 | 1E-45 | 4.9E-44 | 0.3963964 | surface |
| DMXL2 | 2.059 | 9.2419 | 1950.77 | 2E-45 | 7E-44 | 0.3603604 | n.a. |
| NPHP4 | 1.673 | 9.246 | 2175.18 | 3E-45 | 1.2E-43 | 0.2882883 | n.a. |
| HS3ST1 | 1.91 | 9.2857 | 1287.96 | 7E-45 | 2.9E-43 | 0.3513514 | n.a. |
| CXCR3 | 1.81 | 9.3103 | 643.104 | 2E-44 | 1E-42 | 0.3513514 | surface |
| TEX30 | 1.943 | 9.4443 | 234.028 | 6E-44 | 2.5E-42 | 0.4774775 | n.a. |
| FAM81B | 1.667 | 9.2347 | 2737.77 | 1E-43 | 6.2E-42 | 0.2522523 | n.a. |
| JAML | 1.457 | 9.299 | 352.465 | 2E-42 | 7.5E-41 | 0.7927928 | n.a. |
| CPPED1 | 1.92 | 9.3236 | 287.4 | 2E-40 | 6.6E-39 | 0.5765766 | n.a. |
| SELENOW | 1.733 | 9.7827 | 177.343 | 2E-40 | 9.1E-39 | 0.9459459 | n.a. |
| MESD | 1.88 | 9.4258 | 171.005 | 6E-39 | 2.1E-37 | 0.6936937 | n.a. |
| STMN1 | 1.749 | 9.7683 | 167.969 | 3E-38 | 9.6E-37 | 0.4414414 | n.a. |
| ADA2 | 2.025 | 9.2463 | 1054.31 | 4E-38 | 1.5E-36 | 0.3423423 | n.a. |
| TMED2 | 1.648 | 9.6321 | 166.95 | 4E-38 | 1.6E-36 | 0.9009009 | n.a. |
| PVRIG | 1.532 | 9.5646 | 167.528 | 6E-38 | 2.3E-36 | 0.4954955 | n.a. |
| DNAJB9 | 1.996 | 9.2718 | 387.385 | 6E-38 | 2.4E-36 | 0.5225225 | n.a. |
| TTYH1 | 1.563 | 9.2351 | 2796.69 | 1E-37 | 4.4E-36 | 0.2702703 | surface |
| SPCS3 | 1.569 | 9.6267 | 164.793 | 1E-37 | 4.5E-36 | 0.963964 | n.a. |
| DHCR24 | 1.756 | 9.3251 | 328.244 | 2E-37 | 7E-36 | 0.4324324 | n.a. |
| RAB30 | 1.59 | 9.2526 | 1186.04 | 2E-37 | 7.9E-36 | 0.2882883 | n.a. |
| RGS18 | 1.541 | 9.3114 | 287.537 | 2E-37 | 8.1E-36 | 0.7567568 | n.a. |
| LSS | 1.965 | 9.2458 | 959.003 | 3E-37 | 9.1E-36 | 0.4414414 | n.a. |
| TMEM206 | 1.916 | 9.289 | 435.264 | 6E-37 | 2E-35 | 0.3783784 | n.a. |
| RF00581 | 1.517 | 10.319 | 161.104 | 8E-37 | 2.7E-35 | 0.972973 | n.a. |
| GSTA4 | 1.636 | 9.2357 | 5333.29 | 2E-36 | 5.2E-35 | 0.2612613 | n.a. |

| | | | | | | | |
|---|---|---|---|---|---|---|---|
| B3GNT2 | 1.693 | 9.369 | 162.507 | 6E-36 | 2.1E-34 | 0.6036036 | n.a. |
| PADI2 | 1.825 | 9.2434 | 1702.12 | 7E-36 | 2.5E-34 | 0.4324324 | n.a. |
| SLC4A7 | 1.745 | 9.3287 | 225.009 | 8E-36 | 2.6E-34 | 0.4504505 | surface |
| SEC61G | 1.415 | 9.9637 | 156.452 | 8E-36 | 2.7E-34 | 0.981982 | n.a. |
| HCST | 1.518 | 9.5985 | 155.113 | 2E-35 | 5.2E-34 | 0.6396396 | n.a. |
| IRF2BP2 | 1.502 | 9.4142 | 153.254 | 4E-35 | 1.3E-33 | 0.8648649 | n.a. |
| KAT14 | 1.895 | 9.2878 | 325.275 | 7E-35 | 2.2E-33 | 0.4234234 | n.a. |
| SETX | 1.789 | 9.4027 | 152.032 | 7E-35 | 2.4E-33 | 0.6126126 | n.a. |
| RHOH | 1.638 | 9.4176 | 174.987 | 8E-35 | 2.7E-33 | 0.4594595 | n.a. |
| CD44 | 1.266 | 10.108 | 151.24 | 1E-34 | 3.5E-33 | 0.981982 | surface |
| SEC61B | 1.347 | 9.8749 | 150.008 | 2E-34 | 6.5E-33 | 1 | n.a. |
| SSR3 | 1.419 | 9.9724 | 148.306 | 5E-34 | 1.5E-32 | 1 | n.a. |
| NEURL1 | 1.388 | 9.2404 | 2573.4 | 1E-33 | 3E-32 | 0.2972973 | n.a. |
| ENSECAG00000003681 | 1.121 | 10.937 | 146.652 | 1E-33 | 3.4E-32 | 0.990991 | n.a. |
| WWP1 | 2.019 | 9.2543 | 552.347 | 1E-33 | 3.8E-32 | 0.4864865 | n.a. |
| CYFIP2 | 1.724 | 9.3415 | 264.704 | 2E-33 | 4.7E-32 | 0.4054054 | n.a. |
| SH3BP5 | 1.609 | 9.4429 | 321.375 | 2E-33 | 6.8E-32 | 0.6126126 | n.a. |
| SULF2 | 1.577 | 9.2383 | 2611.05 | 4E-33 | 1.3E-31 | 0.3333333 | n.a. |
| MANF | 1.63 | 9.4674 | 143.307 | 6E-33 | 1.8E-31 | 0.8198198 | n.a. |
| LBH | 1.424 | 9.4636 | 152.166 | 1E-32 | 4.2E-31 | 0.3153153 | n.a. |
| SPCS2 | 1.632 | 9.6657 | 141.517 | 1E-32 | 4.3E-31 | 0.8738739 | n.a. |
| SDF2L1 | 1.65 | 9.3874 | 155.982 | 7E-32 | 2E-30 | 0.7387387 | n.a. |
| GPR183 | 1.717 | 9.6621 | 136.983 | 1E-31 | 4.1E-30 | 0.3963964 | surface |
| AFF3 | 1.523 | 9.2652 | 482.684 | 2E-31 | 6.5E-30 | 0.2612613 | n.a. |
| SEC11C | 1.614 | 9.493 | 135.842 | 2E-31 | 7.1E-30 | 0.8108108 | n.a. |
| PPIB | 1.179 | 10.48 | 135.797 | 3E-31 | 7.2E-30 | 0.990991 | n.a. |
| SEPT11 | 1.711 | 9.2663 | 367.77 | 3E-31 | 8.4E-30 | 0.3603604 | n.a. |
| ENSECAG00000037376 | 1.58 | 9.2452 | 758.585 | 6E-31 | 1.8E-29 | 0.3063063 | n.a. |
| CXHXorf21 | 1.749 | 9.2684 | 462.101 | 8E-31 | 2.2E-29 | 0.4594595 | n.a. |
| H2AFJ | 1.399 | 9.3551 | 176.563 | 1E-30 | 2.8E-29 | 0.8378378 | n.a. |
| BET1 | 1.58 | 9.3715 | 135.628 | 1E-30 | 3.9E-29 | 0.6666667 | n.a. |
| STAMBPL1 | 1.721 | 9.3697 | 211.999 | 1E-30 | 4.1E-29 | 0.3603604 | n.a. |
| SMARCAL1 | 1.662 | 9.2489 | 628.945 | 4E-30 | 1E-28 | 0.3423423 | n.a. |
| SPINT2 | 1.77 | 9.2841 | 343.968 | 5E-30 | 1.3E-28 | 0.4864865 | surface |
| BTG1 | 1.226 | 10.365 | 129.764 | 5E-30 | 1.4E-28 | 0.972973 | n.a. |
| STX8 | 1.593 | 9.3397 | 147.968 | 6E-30 | 1.5E-28 | 0.6036036 | n.a. |
| TPP1 | 1.704 | 9.2798 | 261.346 | 7E-30 | 1.8E-28 | 0.6306306 | n.a. |
| DGKZ | 1.778 | 9.2954 | 252.687 | 7E-30 | 2E-28 | 0.4594595 | n.a. |
| GRN | 1.101 | 9.4381 | 128.616 | 9E-30 | 2.5E-28 | 0.9279279 | n.a. |
| SRPRB | 1.727 | 9.3465 | 166.21 | 1E-29 | 3.4E-28 | 0.5225225 | n.a. |
| TSC22D1 | 1.814 | 9.2501 | 622.194 | 2E-29 | 4.2E-28 | 0.3963964 | n.a. |
| C12H11orf74 | 1.783 | 9.3167 | 213.454 | 2E-29 | 5.2E-28 | 0.6216216 | n.a. |
| CD3G | 1.443 | 10.358 | 143.384 | 3E-29 | 9E-28 | 0.4144144 | surface |
| LY6E | 1.543 | 10.241 | 125.792 | 4E-29 | 9.8E-28 | 0.5585586 | surface |
| HIGD1A | 1.491 | 9.3918 | 125.5 | 4E-29 | 1.1E-27 | 0.6666667 | n.a. |
| LYST | 1.421 | 9.3883 | 124.67 | 7E-29 | 1.7E-27 | 0.7297297 | n.a. |
| TM6SF1 | 1.618 | 9.2848 | 259.39 | 1E-28 | 2.9E-27 | 0.5945946 | n.a. |
| RNF213 | 1.599 | 9.2995 | 171.442 | 1E-28 | 3.6E-27 | 0.6126126 | n.a. |
| CXXC5 | 1.615 | 9.3065 | 292.541 | 1E-28 | 3.7E-27 | 0.3243243 | n.a. |
| POLB | 1.617 | 9.3404 | 151.475 | 2E-28 | 4E-27 | 0.5765766 | n.a. |
| ENSECAG00000034985 | 1.428 | 9.3342 | 154.894 | 2E-28 | 4E-27 | 0.6126126 | n.a. |
| SLC39A10 | 1.607 | 9.3738 | 142.679 | 2E-28 | 5.6E-27 | 0.5135135 | surface |
| TPR | 1.617 | 9.453 | 122.166 | 2E-28 | 5.8E-27 | 0.7117117 | surface |
| SNX30 | 1.791 | 9.2487 | 602.062 | 2E-28 | 5.9E-27 | 0.3963964 | n.a. |
| RECQL5 | 1.729 | 9.2498 | 675.905 | 5E-28 | 1.2E-26 | 0.2882883 | n.a. |
| CBX5 | 1.708 | 9.3148 | 208.229 | 2E-27 | 4E-26 | 0.4414414 | n.a. |

| | | | | | | | |
|---|---|---|---|---|---|---|---|
| DYNLRB2 | 1.581 | 9.2488 | 654.016 | 9E-27 | 2.1E-25 | 0.2792793 | n.a. |
| C4orf48 | 1.481 | 9.5904 | 120.464 | 9E-27 | 2.2E-25 | 0.5765766 | n.a. |
| TMED9 | 1.681 | 9.3567 | 151.831 | 1E-26 | 2.5E-25 | 0.4774775 | n.a. |
| RF01277 | 1.41 | 9.8037 | 113.627 | 2E-26 | 4E-25 | 0.9009009 | n.a. |
| CLK2 | 1.596 | 9.3097 | 166.515 | 2E-26 | 5E-25 | 0.4414414 | n.a. |
| CHRNB1 | 1.568 | 9.2427 | 1051.29 | 3E-26 | 7.2E-25 | 0.2612613 | surface |
| ENSECAG00000031485 | 1.663 | 9.2912 | 235.698 | 7E-26 | 1.6E-24 | 0.4684685 | n.a. |
| ENSECAG00000003617 | 1.535 | 9.398 | 113.788 | 1E-25 | 2.2E-24 | 0.6216216 | n.a. |
| MITD1 | 1.646 | 9.3423 | 168.357 | 1E-25 | 2.8E-24 | 0.3783784 | n.a. |
| IGFLR1 | 1.465 | 9.4383 | 138.306 | 1E-25 | 3.2E-24 | 0.5135135 | surface |
| ENSECAG00000038794 | 1.394 | 9.2633 | 1342.62 | 2E-25 | 3.5E-24 | 0.3693694 | n.a. |
| CPVL | 1.254 | 9.3018 | 787.081 | 2E-25 | 4.5E-24 | 0.3873874 | n.a. |
| ENSECAG00000031921 | 1.475 | 9.4417 | 107.989 | 3E-25 | 6.3E-24 | 0.6846847 | n.a. |
| UCP3 | 1.344 | 9.2375 | 1933.84 | 3E-25 | 7.1E-24 | 0.2522523 | n.a. |
| GBGT1 | 1.639 | 9.2944 | 224.245 | 5E-25 | 1.1E-23 | 0.3873874 | n.a. |
| ITGAE | 1.521 | 9.3387 | 181.59 | 8E-25 | 1.8E-23 | 0.3693694 | surface |
| SNRPN | 1.488 | 9.4022 | 113.86 | 1E-24 | 2.1E-23 | 0.5405405 | n.a. |
| TMEM60 | 1.549 | 9.4303 | 105.174 | 1E-24 | 2.6E-23 | 0.6306306 | n.a. |
| MAD1L1 | 1.614 | 9.2498 | 526.54 | 1E-24 | 2.6E-23 | 0.3513514 | n.a. |
| PPT2-EGFL8 | 1.497 | 9.2729 | 332.236 | 1E-24 | 3E-23 | 0.2972973 | n.a. |
| ASB8 | 1.552 | 9.2901 | 183.984 | 2E-24 | 3.9E-23 | 0.4144144 | n.a. |
| CXCR4 | 1.393 | 9.6177 | 104.078 | 2E-24 | 4.4E-23 | 0.3873874 | surface |
| MRPL34 | 1.579 | 9.4086 | 123.06 | 3E-24 | 7.1E-23 | 0.6306306 | n.a. |
| GINM1 | 1.295 | 9.3701 | 113.907 | 5E-24 | 9.6E-23 | 0.6666667 | surface |
| URI1 | 1.643 | 9.2643 | 289.025 | 7E-24 | 1.4E-22 | 0.3513514 | n.a. |
| ABRAXAS1 | 1.641 | 9.2896 | 239.93 | 2E-23 | 3.1E-22 | 0.3693694 | n.a. |
| FAF1 | 1.649 | 9.2455 | 740.282 | 2E-23 | 3.1E-22 | 0.2972973 | n.a. |
| UVRAG | 1.531 | 9.294 | 178.862 | 3E-23 | 5.4E-22 | 0.4144144 | n.a. |
| GPLD1 | 1.666 | 9.245 | 952.343 | 6E-23 | 1.3E-21 | 0.3423423 | n.a. |
| RF00276 | 1.466 | 9.4503 | 96.7806 | 8E-23 | 1.6E-21 | 0.6486486 | n.a. |
| HMGB1 | 1.037 | 10.086 | 96.2805 | 1E-22 | 2.1E-21 | 0.954955 | n.a. |
| EID1 | 1.499 | 9.4085 | 108.747 | 1E-22 | 2.5E-21 | 0.4504505 | n.a. |
| PARP1 | 1.505 | 9.2948 | 247.729 | 2E-22 | 4E-21 | 0.3153153 | n.a. |
| BRK1 | 1.271 | 9.6928 | 94.4191 | 3E-22 | 5.3E-21 | 0.9189189 | n.a. |
| SLAMF7 | 1.107 | 9.2845 | 209.1 | 3E-22 | 5.3E-21 | 0.3783784 | surface |
| PLD4 | 1.225 | 9.3359 | 190.86 | 3E-22 | 5.4E-21 | 0.7567568 | n.a. |
| RAB2B | 1.537 | 9.269 | 254.071 | 6E-22 | 1.1E-20 | 0.3603604 | n.a. |
| AP3S1 | 1.311 | 9.534 | 92.9408 | 6E-22 | 1.1E-20 | 0.8648649 | n.a. |
| GNG2 | 1.167 | 9.4802 | 92.6707 | 7E-22 | 1.3E-20 | 0.5945946 | n.a. |
| IDI1 | 1.39 | 9.4385 | 92.382 | 8E-22 | 1.4E-20 | 0.6126126 | n.a. |
| TOMM7 | 1.519 | 9.4539 | 107.311 | 1E-21 | 2.2E-20 | 0.5585586 | n.a. |
| CCND1 | 1.241 | 9.2529 | 565.763 | 1E-21 | 2.6E-20 | 0.2612613 | n.a. |
| MTMR14 | 1.39 | 9.3346 | 131.198 | 2E-21 | 3E-20 | 0.4774775 | n.a. |
| MAP4K1 | 1.48 | 9.3262 | 145.271 | 2E-21 | 3.8E-20 | 0.4504505 | n.a. |
| LMAN1 | 1.344 | 9.4114 | 88.8086 | 5E-21 | 8.4E-20 | 0.6576577 | n.a. |
| ENSECAG00000000775 | 1.26 | 9.4621 | 101.802 | 5E-21 | 9.2E-20 | 0.8558559 | n.a. |
| RF00163 | 1.374 | 9.4218 | 106.691 | 7E-21 | 1.2E-19 | 0.4054054 | n.a. |
| TOB1 | 1.331 | 9.3276 | 126.535 | 7E-21 | 1.3E-19 | 0.4594595 | n.a. |
| ENSECAG00000034569 | 1.036 | 9.7713 | 87.7898 | 8E-21 | 1.4E-19 | 0.2702703 | n.a. |
| DKC1 | 1.384 | 9.411 | 94.0709 | 2E-20 | 2.8E-19 | 0.3873874 | n.a. |
| APOBEC3H | 1.368 | 9.3557 | 126.137 | 2E-20 | 3E-19 | 0.4054054 | n.a. |
| AKR1B1 | 1.31 | 9.4302 | 86.178 | 2E-20 | 3.1E-19 | 0.6126126 | n.a. |
| ENSECAG00000038299 | 1.385 | 9.3304 | 110.335 | 2E-20 | 4.1E-19 | 0.5225225 | n.a. |
| ARL1 | 1.438 | 9.4084 | 95.6478 | 3E-20 | 5E-19 | 0.6216216 | n.a. |
| LAT | 1.392 | 9.5366 | 114.947 | 3E-20 | 6.1E-19 | 0.3513514 | n.a. |
| SQLE | 1.45 | 9.2764 | 212.055 | 4E-20 | 6.7E-19 | 0.2972973 | n.a. |

| | | | | | | | |
|---|---|---|---|---|---|---|---|
| PLA2G16 | 1.124 | 9.6144 | 84.0822 | 5E-20 | 8.7E-19 | 0.954955 | n.a. |
| IL7R | 1.24 | 9.3731 | 288.255 | 5E-20 | 8.8E-19 | 0.4054054 | surface |
| CTSB | 1.052 | 9.4471 | 89.8077 | 5E-20 | 9.3E-19 | 0.8378378 | n.a. |
| ZFYVE16 | 1.507 | 9.2508 | 381.444 | 7E-20 | 1.2E-18 | 0.2882883 | n.a. |
| MPPE1 | 1.529 | 9.2717 | 304.072 | 8E-20 | 1.3E-18 | 0.2792793 | n.a. |
| SRPRA | 1.354 | 9.3887 | 87.6805 | 8E-20 | 1.3E-18 | 0.5855856 | n.a. |
| ENSECAG00000032188 | 1.061 | 9.5863 | 82.9736 | 9E-20 | 1.5E-18 | 0.9459459 | n.a. |
| LSM4 | 1.319 | 9.4444 | 82.7367 | 1E-19 | 1.7E-18 | 0.6486486 | n.a. |
| TMEM230 | 1.407 | 9.3512 | 110.599 | 1E-19 | 2E-18 | 0.4864865 | n.a. |
| MTF2 | 1.409 | 9.3258 | 118.233 | 1E-19 | 2.4E-18 | 0.3783784 | n.a. |
| PPA2 | 1.547 | 9.2617 | 269.76 | 2E-19 | 3E-18 | 0.3783784 | n.a. |
| TMED5 | 1.384 | 9.3523 | 104.563 | 2E-19 | 3E-18 | 0.6126126 | n.a. |
| ENSECAG00000040126 | 1.403 | 9.2742 | 200.829 | 2E-19 | 3.3E-18 | 0.3243243 | n.a. |
| RBM34 | 1.364 | 9.3453 | 134.778 | 2E-19 | 3.7E-18 | 0.3603604 | n.a. |
| PRELID1 | 1.033 | 9.7488 | 81.0516 | 2E-19 | 3.8E-18 | 0.981982 | n.a. |
| SLC1A5 | 1.472 | 9.2879 | 180.182 | 3E-19 | 4.6E-18 | 0.4594595 | surface |
| GTF3A | 1.376 | 9.6075 | 80.2685 | 3E-19 | 5.7E-18 | 0.7747748 | n.a. |
| ECHS1 | 1.365 | 9.393 | 92.5777 | 4E-19 | 7.1E-18 | 0.6306306 | n.a. |
| CHMP6 | 1.337 | 9.3694 | 89.938 | 7E-19 | 1.1E-17 | 0.5225225 | n.a. |
| PEA15 | 1.358 | 9.2969 | 196.132 | 1E-18 | 2.4E-17 | 0.4324324 | n.a. |
| HHEX | 1.034 | 9.4401 | 76.7618 | 2E-18 | 3.3E-17 | 0.7117117 | n.a. |
| ENSECAG00000038584 | 1.215 | 10.313 | 76.3191 | 3E-18 | 4.1E-17 | 0.8468468 | n.a. |
| TPST2 | 1.287 | 9.4709 | 75.8777 | 4E-18 | 6.7E-17 | 0.5765766 | n.a. |
| PTBP3 | 1.233 | 9.4001 | 73.9579 | 8E-18 | 1.3E-16 | 0.5585586 | n.a. |
| DNASE2 | 1.419 | 9.347 | 156.153 | 8E-18 | 1.3E-16 | 0.6126126 | n.a. |
| SEC61A1 | 1.387 | 9.3336 | 112.553 | 1E-17 | 2E-16 | 0.5495495 | n.a. |
| CMTM3 | 1.3 | 9.4968 | 81.706 | 2E-17 | 2.6E-16 | 0.6126126 | n.a. |
| NHP2 | 1.292 | 9.4198 | 79.6663 | 2E-17 | 3E-16 | 0.6846847 | n.a. |
| ENSECAG00000016754 | 1.345 | 9.2778 | 156.55 | 2E-17 | 3.1E-16 | 0.4234234 | n.a. |
| CUTA | 1.378 | 9.3533 | 110.659 | 2E-17 | 3.3E-16 | 0.4234234 | n.a. |
| MICAL1 | 1.213 | 9.2809 | 158.215 | 2E-17 | 3.8E-16 | 0.3423423 | n.a. |
| ENSECAG00000020618 | 1.255 | 9.2715 | 251.321 | 4E-17 | 5.6E-16 | 0.2792793 | n.a. |
| TBC1D1 | 1.359 | 9.2859 | 160.401 | 4E-17 | 6.4E-16 | 0.4504505 | n.a. |
| EMC10 | 1.277 | 9.3664 | 79.4183 | 6E-17 | 8.5E-16 | 0.5135135 | n.a. |
| ZNF706 | 1.014 | 9.8395 | 69.9137 | 6E-17 | 9.5E-16 | 0.981982 | n.a. |
| AMPD3 | 1.491 | 9.2518 | 525.358 | 8E-17 | 1.1E-15 | 0.3063063 | n.a. |
| OSTC | 1.203 | 9.5663 | 69.4136 | 8E-17 | 1.2E-15 | 0.8108108 | n.a. |
| EBAG9 | 1.196 | 9.3732 | 72.0868 | 1E-16 | 1.6E-15 | 0.5495495 | n.a. |
| ENSECAG00000023261 | 1.101 | 9.5604 | 68.3696 | 1E-16 | 2E-15 | 0.8558559 | n.a. |
| SPPL2A | 1.395 | 9.2923 | 135.206 | 1E-16 | 2E-15 | 0.4324324 | surface |
| MOCS2 | 1.358 | 9.3218 | 104.275 | 1E-16 | 2.1E-15 | 0.4234234 | n.a. |
| PDIA6 | 1.312 | 9.332 | 98.7787 | 2E-16 | 2.5E-15 | 0.5225225 | n.a. |
| PGAM5 | 1.344 | 9.3326 | 102.664 | 2E-16 | 3E-15 | 0.4144144 | n.a. |
| COPZ1 | 1.293 | 9.391 | 80.0801 | 3E-16 | 4E-15 | 0.5675676 | n.a. |
| ALG5 | 1.26 | 9.3503 | 85.254 | 3E-16 | 4.4E-15 | 0.5495495 | n.a. |
| MED13L | 1.278 | 9.2911 | 112.735 | 5E-16 | 7.2E-15 | 0.4504505 | n.a. |
| ERLEC1 | 1.299 | 9.3911 | 78.512 | 6E-16 | 7.7E-15 | 0.6036036 | n.a. |
| ZDHHC24 | 1.304 | 9.314 | 102.117 | 6E-16 | 7.7E-15 | 0.5225225 | n.a. |
| NEU1 | 1.307 | 9.2661 | 202.402 | 6E-16 | 7.8E-15 | 0.2612613 | n.a. |
| MILR1 | 1.232 | 9.2539 | 401.632 | 6E-16 | 8.4E-15 | 0.3873874 | surface |
| SMIM26 | 1.218 | 9.4437 | 67.6883 | 7E-16 | 9E-15 | 0.7027027 | n.a. |
| CCDC12 | 1.092 | 9.3727 | 65.1878 | 7E-16 | 9.7E-15 | 0.6126126 | n.a. |
| CIRBP | 1.316 | 9.4608 | 69.2034 | 8E-16 | 1.1E-14 | 0.4324324 | n.a. |
| EDEM2 | 1.38 | 9.308 | 120.599 | 1E-15 | 1.7E-14 | 0.4414414 | n.a. |
| PRXL2A | 1.297 | 9.2641 | 229.121 | 2E-15 | 2.8E-14 | 0.2612613 | n.a. |
| PLBD2 | 1.227 | 9.3161 | 110.358 | 2E-15 | 2.9E-14 | 0.6936937 | n.a. |

| | | | | | | | |
|---|---|---|---|---|---|---|---|
| ORMDL2 | 1.306 | 9.3745 | 80.1756 | 2E-15 | 2.9E-14 | 0.4414414 | n.a. |
| CRELD2 | 1.262 | 9.2876 | 137.607 | 2E-15 | 3E-14 | 0.3513514 | n.a. |
| STK17A | 1.044 | 9.6864 | 62.8381 | 2E-15 | 3.1E-14 | 0.7927928 | n.a. |
| DOCK8 | 1.068 | 9.3857 | 62.7789 | 2E-15 | 3.2E-14 | 0.7117117 | n.a. |
| HSP90AA1 | 1.025 | 9.9098 | 61.669 | 4E-15 | 5.5E-14 | 0.8648649 | n.a. |
| RHEB | 1.024 | 9.4952 | 61.5171 | 5E-15 | 6E-14 | 0.8198198 | n.a. |
| CYB5R1 | 1.348 | 9.253 | 289.199 | 6E-15 | 7.9E-14 | 0.2702703 | n.a. |
| NIPA2 | 1.211 | 9.3276 | 84.5799 | 7E-15 | 8.6E-14 | 0.5315315 | n.a. |
| SEC11A | 1.1 | 9.5255 | 59.9524 | 1E-14 | 1.3E-13 | 0.7837838 | n.a. |
| RF00280 | 1.209 | 9.4118 | 78.5525 | 1E-14 | 1.3E-13 | 0.4594595 | n.a. |
| RCL1 | 1.143 | 9.2803 | 159.176 | 1E-14 | 1.5E-13 | 0.3423423 | n.a. |
| SAP30BP | 1.338 | 9.2862 | 141.337 | 1E-14 | 1.7E-13 | 0.2972973 | n.a. |
| ENSECAG00000036779 | 1.166 | 9.5478 | 59.2082 | 1E-14 | 1.9E-13 | 0.6846847 | n.a. |
| TTC14 | 1.205 | 9.3422 | 77.6763 | 2E-14 | 2E-13 | 0.3423423 | n.a. |
| TRAM1 | 1.059 | 9.4843 | 58.6111 | 2E-14 | 2.5E-13 | 0.8198198 | n.a. |
| TSPAN13 | 1.224 | 9.2631 | 249.313 | 3E-14 | 3.2E-13 | 0.3423423 | surface |
| AOAH | 1.036 | 9.3519 | 79.5456 | 3E-14 | 3.7E-13 | 0.5765766 | n.a. |
| ARSB | 1.147 | 9.2518 | 341.587 | 4E-14 | 5E-13 | 0.2702703 | n.a. |
| PRPSAP2 | 1.234 | 9.3039 | 111.11 | 5E-14 | 6.5E-13 | 0.3423423 | n.a. |
| YIPF1 | 1.395 | 9.287 | 152.075 | 1E-13 | 1.2E-12 | 0.3603604 | n.a. |
| SLC2A8 | 1.234 | 9.2576 | 223.48 | 1E-13 | 1.5E-12 | 0.2792793 | surface |
| PRKACB | 1.144 | 9.3195 | 76.3188 | 1E-13 | 1.5E-12 | 0.4144144 | n.a. |
| NEDD1 | 1.192 | 9.2856 | 114.027 | 1E-13 | 1.5E-12 | 0.3423423 | n.a. |
| CSGALNACT2 | 1.228 | 9.2796 | 124.416 | 2E-13 | 2E-12 | 0.3693694 | n.a. |
| MRPS24 | 1.142 | 9.6247 | 54.3575 | 2E-13 | 2E-12 | 0.6756757 | n.a. |
| TMEM41B | 1.255 | 9.2955 | 103.432 | 2E-13 | 2.2E-12 | 0.3783784 | n.a. |
| RPN2 | 1.083 | 9.4695 | 54.1662 | 2E-13 | 2.2E-12 | 0.7027027 | n.a. |
| HAGH | 1.173 | 9.3228 | 80.2457 | 2E-13 | 2.6E-12 | 0.4864865 | n.a. |
| EMD | 1.102 | 9.5059 | 53.8483 | 2E-13 | 2.6E-12 | 0.5945946 | n.a. |
| DPP9 | 1.208 | 9.3491 | 79.0323 | 2E-13 | 2.7E-12 | 0.3693694 | n.a. |
| TMCO1 | 1.01 | 9.4392 | 53.5673 | 3E-13 | 3E-12 | 0.6666667 | n.a. |
| CBFA2T3 | 1.092 | 9.2789 | 160.74 | 3E-13 | 3.6E-12 | 0.3783784 | n.a. |
| SPTBN1 | 1.116 | 9.3474 | 79.2607 | 3E-13 | 3.8E-12 | 0.3333333 | n.a. |
| SSR1 | 1.211 | 9.3182 | 86.5921 | 3E-13 | 3.9E-12 | 0.4234234 | surface |
| GLTP | 1.182 | 9.2919 | 101.446 | 3E-13 | 3.9E-12 | 0.3603604 | n.a. |
| ENSECAG00000026829 | 1.114 | 9.535 | 52.8325 | 4E-13 | 4.3E-12 | 0.6666667 | n.a. |
| STOML2 | 1.163 | 9.3994 | 66.906 | 7E-13 | 7.8E-12 | 0.5225225 | n.a. |
| DECR1 | 1.183 | 9.334 | 78.6259 | 7E-13 | 7.9E-12 | 0.4234234 | n.a. |
| FLT3 | 1.083 | 9.2499 | 440.168 | 8E-13 | 8.9E-12 | 0.3513514 | surface |
| RPAIN | 1.279 | 9.2532 | 233.365 | 8E-13 | 9.3E-12 | 0.2882883 | n.a. |
| CLPP | 1.076 | 9.4622 | 51.2563 | 8E-13 | 9.3E-12 | 0.6126126 | n.a. |
| ST3GAL4 | 1.17 | 9.3052 | 83.9495 | 1E-12 | 1.1E-11 | 0.4054054 | n.a. |
| MRPL36 | 1.115 | 9.4047 | 56.6095 | 1E-12 | 1.2E-11 | 0.5675676 | n.a. |
| FDPS | 1.093 | 9.2939 | 106.807 | 1E-12 | 1.2E-11 | 0.3873874 | n.a. |
| CACYBP | 1.084 | 9.6062 | 50.3072 | 1E-12 | 1.5E-11 | 0.7387387 | n.a. |
| FAM133B | 1.124 | 9.3623 | 62.5617 | 2E-12 | 1.9E-11 | 0.3333333 | n.a. |
| NUDT22 | 1.19 | 9.3226 | 83.0612 | 2E-12 | 2E-11 | 0.5045045 | n.a. |
| ZNF644 | 1.176 | 9.3066 | 89.3394 | 2E-12 | 2E-11 | 0.2882883 | n.a. |
| CDCA4 | 1.173 | 9.2785 | 124.976 | 2E-12 | 2.1E-11 | 0.2882883 | n.a. |
| USO1 | 1.22 | 9.2993 | 94.8846 | 2E-12 | 2.2E-11 | 0.3873874 | n.a. |
| WDFY4 | 1.12 | 9.2866 | 184.772 | 2E-12 | 2.4E-11 | 0.3873874 | n.a. |
| SUMO1 | 1.008 | 9.561 | 49.3064 | 2E-12 | 2.4E-11 | 0.7117117 | n.a. |
| STT3A | 1.194 | 9.2923 | 101.111 | 3E-12 | 2.9E-11 | 0.2972973 | n.a. |
| EIF4E1B | 1.17 | 9.2561 | 319.076 | 3E-12 | 3.2E-11 | 0.4594595 | n.a. |
| PRRC2C | 1.04 | 9.5988 | 48.5755 | 3E-12 | 3.5E-11 | 0.6756757 | n.a. |
| DAPP1 | 1.019 | 9.2902 | 81.6498 | 4E-12 | 3.9E-11 | 0.4504505 | n.a. |

| | | | | | | | |
|---|---|---|---|---|---|---|---|
| SMIM10L1 | 1.086 | 9.5295 | 48.3651 | 4E-12 | 3.9E-11 | 0.5765766 | n.a. |
| IGSF8 | 1.19 | 9.2908 | 100.463 | 4E-12 | 3.9E-11 | 0.3603604 | surface |
| UNC93B1 | 1.178 | 9.2602 | 211.571 | 4E-12 | 4.7E-11 | 0.4414414 | surface |
| CYCS | 1.124 | 9.3968 | 61.165 | 6E-12 | 6.5E-11 | 0.5585586 | n.a. |
| ENSECAG00000032776 | 1.045 | 9.4539 | 47.2077 | 6E-12 | 6.9E-11 | 0.3243243 | n.a. |
| GTF2F1 | 1.189 | 9.3478 | 73.2674 | 7E-12 | 7.1E-11 | 0.4594595 | n.a. |
| TTF1 | 1.129 | 9.2977 | 95.8334 | 7E-12 | 7.3E-11 | 0.3333333 | n.a. |
| ENSECAG00000007978 | 1.072 | 9.3138 | 72.8456 | 7E-12 | 7.3E-11 | 0.3243243 | n.a. |
| AGPAT1 | 1.246 | 9.2783 | 138.27 | 8E-12 | 8.1E-11 | 0.2792793 | n.a. |
| SNRPB2 | 1.064 | 9.5023 | 46.6734 | 9E-12 | 8.9E-11 | 0.6036036 | n.a. |
| INTS12 | 1.132 | 9.3106 | 77.0966 | 9E-12 | 9.6E-11 | 0.3513514 | n.a. |
| KCTD10 | 1.169 | 9.3 | 109.385 | 1E-11 | 1.1E-10 | 0.2522523 | n.a. |
| ZMYND11 | 1.04 | 9.4456 | 46.0385 | 1E-11 | 1.2E-10 | 0.5045045 | n.a. |
| C16orf87 | 1.255 | 9.2689 | 159.919 | 1E-11 | 1.2E-10 | 0.2792793 | n.a. |
| NGLY1 | 1.126 | 9.2901 | 85.5046 | 1E-11 | 1.4E-10 | 0.3603604 | n.a. |
| ERP29 | 1.003 | 9.3518 | 68.734 | 2E-11 | 1.6E-10 | 0.6486486 | n.a. |
| MCM7 | 1.111 | 9.3416 | 74.4716 | 2E-11 | 1.8E-10 | 0.2882883 | n.a. |
| ENSECAG00000015215 | 1.046 | 9.3399 | 59.9466 | 2E-11 | 1.8E-10 | 0.3333333 | n.a. |
| PTPN1 | 1.152 | 9.278 | 135.749 | 2E-11 | 2E-10 | 0.2612613 | n.a. |
| NCSTN | 1.088 | 9.302 | 77.1286 | 2E-11 | 2.4E-10 | 0.4234234 | surface |
| ENSECAG00000036511 | 1.08 | 9.3303 | 75.7884 | 3E-11 | 2.7E-10 | 0.4144144 | n.a. |
| FAM117A | 1.159 | 9.3289 | 74.2189 | 3E-11 | 2.7E-10 | 0.3783784 | n.a. |
| VPS37B | 1.07 | 9.2968 | 104.193 | 3E-11 | 2.9E-10 | 0.2792793 | n.a. |
| ITCH | 1.003 | 9.272 | 87.1479 | 3E-11 | 3.5E-10 | 0.3423423 | n.a. |
| NUDCD2 | 1.024 | 9.3543 | 55.9602 | 4E-11 | 3.6E-10 | 0.4144144 | n.a. |
| COPB1 | 1.197 | 9.3293 | 77.9451 | 4E-11 | 3.8E-10 | 0.4234234 | n.a. |
| ATG12 | 1.118 | 9.2965 | 83.9554 | 4E-11 | 4.2E-10 | 0.4144144 | n.a. |
| ADAM17 | 1.149 | 9.2788 | 115.844 | 4E-11 | 4.4E-10 | 0.2522523 | surface |
| NPDC1 | 1.011 | 9.5117 | 48.4774 | 5E-11 | 4.4E-10 | 0.3513514 | n.a. |
| GLG1 | 1.022 | 9.3396 | 53.8233 | 5E-11 | 4.7E-10 | 0.4414414 | n.a. |
| UTP18 | 1.113 | 9.318 | 77.1198 | 5E-11 | 5.2E-10 | 0.3513514 | n.a. |
| ARMCX3 | 1.005 | 9.3437 | 49.3331 | 6E-11 | 5.8E-10 | 0.4234234 | n.a. |
| FKBP2 | 1.014 | 9.3386 | 62.9921 | 8E-11 | 7.9E-10 | 0.5225225 | n.a. |
| RWDD4 | 1.138 | 9.3262 | 71.7372 | 8E-11 | 8E-10 | 0.4054054 | n.a. |
| PEX16 | 1.085 | 9.3211 | 71.0102 | 1E-10 | 9.3E-10 | 0.2972973 | n.a. |
| UBE2S | 1.048 | 9.3494 | 61.2794 | 1E-10 | 1.2E-09 | 0.3693694 | n.a. |
| BLOC1S2 | 1.066 | 9.353 | 67.9964 | 1E-10 | 1.3E-09 | 0.4864865 | n.a. |
| GLB1 | 1.075 | 9.2938 | 89.2034 | 1E-10 | 1.3E-09 | 0.3333333 | n.a. |
| DZIP3 | 1.094 | 9.2869 | 111.078 | 1E-10 | 1.4E-09 | 0.2972973 | n.a. |
| NUDT3 | 1.072 | 9.2715 | 116.076 | 2E-10 | 1.6E-09 | 0.3063063 | n.a. |
| NSMAF | 1.055 | 9.2696 | 124.548 | 2E-10 | 1.7E-09 | 0.2792793 | n.a. |
| HDAC2 | 1.189 | 9.2975 | 93.7696 | 2E-10 | 2E-09 | 0.3513514 | n.a. |
| GTF2I | 1.051 | 9.3122 | 67.5064 | 2E-10 | 2E-09 | 0.3243243 | n.a. |
| SLC46A3 | 1.155 | 9.2637 | 156.544 | 3E-10 | 2.3E-09 | 0.3333333 | surface |
| RFC1 | 1.046 | 9.3607 | 61.4419 | 3E-10 | 2.5E-09 | 0.3153153 | n.a. |
| MCFD2 | 1.136 | 9.2819 | 101.937 | 3E-10 | 2.7E-09 | 0.3423423 | n.a. |
| BBX | 1.003 | 9.3925 | 45.2696 | 4E-10 | 3.7E-09 | 0.4504505 | n.a. |
| SEC31A | 1.04 | 9.2996 | 66.921 | 4E-10 | 3.9E-09 | 0.3873874 | n.a. |
| ACYP2 | 1.151 | 9.2639 | 142.599 | 5E-10 | 4.1E-09 | 0.2522523 | n.a. |
| EMC9 | 1.19 | 9.2686 | 149.04 | 5E-10 | 4.9E-09 | 0.3063063 | n.a. |
| ACTN4 | 1.106 | 9.3064 | 78.9218 | 6E-10 | 5.5E-09 | 0.4324324 | n.a. |
| TMEM248 | 1.042 | 9.2961 | 78.0809 | 1E-09 | 1.2E-08 | 0.2882883 | n.a. |
| CERS5 | 1.046 | 9.2999 | 73.6107 | 2E-09 | 1.8E-08 | 0.3963964 | n.a. |
| POC1A | 1.039 | 9.2936 | 117.369 | 3E-09 | 2.3E-08 | 0.2702703 | n.a. |
| CETN2 | 1.045 | 9.3883 | 52.4124 | 3E-09 | 2.3E-08 | 0.3693694 | n.a. |
| ATP6V0A1 | 1.078 | 9.2697 | 118.096 | 5E-09 | 3.7E-08 | 0.2612613 | n.a. |

| | | | | | | | |
|---|---|---|---|---|---|---|---|
| PDIA4 | 1.024 | 9.2985 | 78.4562 | 5E-09 | 3.7E-08 | 0.4054054 | n.a. |
| MORF4L2 | 1.011 | 9.2994 | 68.7233 | 6E-09 | 4.9E-08 | 0.3153153 | n.a. |
| PBRM1 | 1.008 | 9.3154 | 59.5166 | 7E-09 | 5.3E-08 | 0.2702703 | n.a. |
| ZNF789 | 1.045 | 9.2679 | 111.926 | 2E-08 | 1.5E-07 | 0.2702703 | n.a. |
| MRPS11 | 1.009 | 9.3077 | 67.4489 | 2E-08 | 1.5E-07 | 0.3063063 | n.a. |
| ENSECAG00000007532 | 1.072 | 9.3313 | 61.2772 | 3E-08 | 2.2E-07 | 0.3783784 | n.a. |
| ACAT1 | 1.017 | 9.2846 | 81.7517 | 4E-08 | 2.9E-07 | 0.3963964 | n.a. |

| genes | logFC | logCPM | F | PValue | FDR | percent.exp | Surfacome.Label |
|---|---|---|---|---|---|---|---|
| ENSECAG00000006663 | 6.089 | 9.3132 | 5712.26 | 0 | 0 | 0.987013 | n.a. |
| ENSECAG00000031322 | 4.877 | 10.499 | 2501.66 | 0 | 0 | 1 | n.a. |
| FCER1G | 2.073 | 10.204 | 1639.98 | 0 | 0 | 1 | n.a. |
| ENSECAG00000032710 | 4.67 | 9.3761 | 1596.6 | 4E-301 | 8E-298 | 0.7662338 | n.a. |
| PLVAP | 4.631 | 9.247 | 5746.3 | 1E-289 | 2E-286 | 0.8441558 | surface |
| RNASE6 | 2.313 | 10.205 | 1124.65 | 1E-242 | 1E-239 | 1 | n.a. |
| GNGT2 | 4.29 | 9.2643 | 3425.66 | 2E-239 | 2E-236 | 0.8181818 | n.a. |
| RNASE4 | 2.259 | 10.101 | 1109.07 | 2E-239 | 2E-236 | 1 | n.a. |
| ALDH2 | 3.167 | 9.5616 | 1421.13 | 5E-219 | 4E-216 | 0.9350649 | n.a. |
| CD8A | 3.924 | 9.4281 | 1197.42 | 2E-204 | 1E-201 | 0.8701299 | surface |
| MNDA | 3.578 | 10.258 | 932.459 | 4E-202 | 3E-199 | 1 | n.a. |
| CELA1 | 4.239 | 9.2376 | 18314.9 | 7E-194 | 4E-191 | 0.6233766 | n.a. |
| ENSECAG00000032959 | 3.835 | 9.6944 | 881.637 | 2E-191 | 1E-188 | 0.7792208 | n.a. |
| CX3CR1 | 3.762 | 9.2887 | 2667.94 | 1E-177 | 8E-175 | 0.7922078 | surface |
| PRNP | 3.253 | 9.3261 | 1345.32 | 8E-169 | 4E-166 | 0.8311688 | surface |
| ATP1B3 | 3.299 | 9.5391 | 773.688 | 2E-168 | 1E-165 | 0.9480519 | surface |
| CEBPB | 2.662 | 9.407 | 1091.83 | 2E-166 | 1E-163 | 0.9480519 | n.a. |
| ENSECAG00000019318 | 3.091 | 9.3455 | 1267.47 | 2E-161 | 9E-159 | 0.7142857 | n.a. |
| FTL | 1.805 | 11.187 | 672.863 | 6E-147 | 3E-144 | 1 | n.a. |
| GBP5 | 3.908 | 9.7163 | 622.859 | 3E-136 | 1E-133 | 0.4415584 | n.a. |
| POU2F2 | 3.354 | 9.5681 | 636.2 | 3E-136 | 1E-133 | 0.8961039 | n.a. |
| C4BPA | 2.456 | 9.4013 | 1393.2 | 2E-129 | 6E-127 | 0.8961039 | n.a. |
| CSF1R | 3.077 | 9.3336 | 2695.69 | 6E-123 | 2E-120 | 0.974026 | surface |
| HMOX1 | 2.81 | 9.3479 | 723.465 | 2E-117 | 6E-115 | 0.8701299 | n.a. |
| NR4A1 | 3.362 | 9.2477 | 3488.13 | 1E-116 | 4E-114 | 0.5714286 | n.a. |
| AIM2 | 3.011 | 9.3585 | 1119.48 | 4E-98 | 1.2E-95 | 0.8181818 | n.a. |
| ABI3 | 3.118 | 9.3324 | 822.043 | 1E-95 | 3.7E-93 | 0.7532468 | n.a. |
| ICAM2 | 3.444 | 9.4874 | 557.245 | 2E-90 | 5E-88 | 0.4935065 | surface |
| TCF7L2 | 3.044 | 9.2572 | 1780.18 | 9E-90 | 2.2E-87 | 0.6493506 | n.a. |
| EQMHCB2 | 1.256 | 12.61 | 401.602 | 8E-89 | 1.9E-86 | 1 | n.a. |
| LST1 | 2.428 | 9.329 | 591.339 | 9E-87 | 2.2E-84 | 0.7792208 | n.a. |
| ENSECAG00000040478 | 3.169 | 9.248 | 3470.52 | 2E-86 | 4.7E-84 | 0.6233766 | n.a. |
| ENSECAG00000033857 | 2.219 | 9.3316 | 1865.74 | 1E-82 | 3.2E-80 | 0.8571429 | n.a. |
| ENSECAG00000015010 | 2.778 | 9.2872 | 1968.93 | 4E-78 | 7.7E-76 | 0.6883117 | n.a. |
| PLA2G16 | 2.438 | 9.6144 | 350.337 | 9E-78 | 1.8E-75 | 1 | n.a. |
| CD74 | 1.433 | 12.628 | 334.824 | 2E-74 | 3.9E-72 | 1 | surface |
| ENSECAG00000031455 | 2.874 | 9.2358 | 5108 | 4E-74 | 7.1E-72 | 0.3766234 | n.a. |
| PLAC8B | 2.006 | 11.113 | 330.999 | 1E-73 | 2.5E-71 | 0.974026 | n.a. |
| H2AFZ | 2.575 | 9.5191 | 330.755 | 1E-73 | 2.7E-71 | 0.7532468 | n.a. |
| ENSECAG00000019100 | 3.013 | 9.2458 | 3997.79 | 2E-72 | 4.2E-70 | 0.5714286 | n.a. |
| ENSECAG00000038143 | 2.96 | 9.2349 | 6033.97 | 4E-72 | 7E-70 | 0.4155844 | n.a. |
| GPIHBP1 | 2.396 | 9.2751 | 2964.51 | 3E-70 | 4.5E-68 | 0.5844156 | surface |
| IFI27 | 2.474 | 9.7314 | 312.091 | 2E-69 | 2.7E-67 | 0.8831169 | n.a. |
| DCSTAMP | 2.744 | 9.2378 | 5354.37 | 4E-69 | 6.7E-67 | 0.3636364 | surface |
| IFI30 | 2.072 | 9.9429 | 301.194 | 3E-67 | 5.7E-65 | 0.961039 | n.a. |
| PTPRCAP | 2.341 | 10.617 | 297.762 | 2E-66 | 3.1E-64 | 0.7402597 | n.a. |
| CD68 | 2.368 | 9.3465 | 947.746 | 2E-66 | 3.5E-64 | 0.8441558 | surface |
| GBP6 | 3.164 | 9.5114 | 341.982 | 2E-65 | 3.2E-63 | 0.3246753 | n.a. |
| CD79A | 2.841 | 9.7872 | 770.523 | 2E-63 | 3.6E-61 | 0.4025974 | surface |
| TNFSF10 | 2.567 | 9.4268 | 320.102 | 8E-63 | 1.2E-60 | 0.7792208 | n.a. |
| ENSECAG00000021110 | 2.682 | 9.2654 | 2385.08 | 1E-62 | 2.1E-60 | 0.5974026 | n.a. |
| ST3GAL6 | 2.807 | 9.5094 | 276.378 | 8E-62 | 1.2E-59 | 0.7532468 | n.a. |
| FLNA | 2.233 | 9.4126 | 274.176 | 2E-61 | 3.4E-59 | 0.7532468 | n.a. |
| CD37 | 2.241 | 9.891 | 270.276 | 2E-60 | 2.3E-58 | 0.961039 | surface |
| P2RY14 | 2.683 | 9.2475 | 1614.57 | 9E-59 | 1.2E-56 | 0.4545455 | surface |

| | | | | | | | |
|---|---|---|---|---|---|---|---|
| SULT1C4 | 2.159 | 9.421 | 1573.7 | 2E-58 | 2.5E-56 | 0.7792208 | n.a. |
| CD300H | 2.631 | 9.2467 | 1838.45 | 3E-58 | 3.6E-56 | 0.5194805 | n.a. |
| CORO1B | 2.192 | 9.4003 | 285.927 | 2E-55 | 2.1E-53 | 0.7922078 | n.a. |
| CD53 | 2.232 | 9.709 | 242.632 | 2E-54 | 2E-52 | 0.9350649 | surface |
| ARHGDIB | 1.274 | 11.255 | 240.54 | 5E-54 | 5.7E-52 | 0.987013 | n.a. |
| HES4 | 2.63 | 9.2428 | 2465.89 | 1E-53 | 1.2E-51 | 0.4415584 | n.a. |
| ENSECAG00000019052 | 2.654 | 9.4644 | 288.825 | 5E-52 | 5.7E-50 | 0.5194805 | n.a. |
| ENSECAG00000036967 | 1.807 | 9.2599 | 2377.69 | 7E-52 | 8E-50 | 0.3506494 | n.a. |
| TPK1 | 2.628 | 9.2622 | 770.036 | 2E-50 | 2.9E-48 | 0.4155844 | n.a. |
| eca-mir-1892 | 2.423 | 9.4799 | 222.122 | 4E-50 | 5.2E-48 | 0.7532468 | n.a. |
| DUSP5 | 2.193 | 9.267 | 1001.15 | 9E-50 | 1E-47 | 0.3896104 | n.a. |
| FABP5 | 2.782 | 9.5264 | 247.972 | 9E-49 | 9.8E-47 | 0.5064935 | n.a. |
| MYO1G | 2.396 | 9.6134 | 215.64 | 1E-48 | 1.3E-46 | 0.8311688 | n.a. |
| MS4A7 | 2.311 | 9.3128 | 1114.51 | 4E-46 | 4.2E-44 | 0.5714286 | n.a. |
| BIN1 | 2.514 | 9.382 | 382.031 | 6E-46 | 6.3E-44 | 0.5064935 | n.a. |
| TXNIP | 1.792 | 10.204 | 202.779 | 7E-46 | 7.2E-44 | 0.974026 | n.a. |
| EVL | 2.578 | 9.7253 | 202.224 | 9E-46 | 9.4E-44 | 0.6103896 | n.a. |
| WARS | 2.356 | 9.8298 | 202.131 | 1E-45 | 9.7E-44 | 0.8051948 | n.a. |
| S100A4 | 1.847 | 11.292 | 201.694 | 1E-45 | 1.2E-43 | 0.9220779 | n.a. |
| TIE1 | 2.541 | 9.2364 | 3774.17 | 1E-45 | 1.2E-43 | 0.3766234 | surface |
| ENSECAG00000040634 | 2.372 | 9.9556 | 195.484 | 3E-44 | 2.6E-42 | 0.7402597 | n.a. |
| SAMD9L | 1.958 | 9.4216 | 191.868 | 2E-42 | 1.5E-40 | 0.6753247 | n.a. |
| Eqca-2 | 1.139 | 11.477 | 186.247 | 3E-42 | 2.5E-40 | 0.961039 | n.a. |
| UBC | 1.307 | 11.249 | 182.173 | 2E-41 | 1.9E-39 | 0.9350649 | n.a. |
| ENSECAG00000004433 | 2.34 | 9.2481 | 1303.39 | 7E-41 | 6.5E-39 | 0.3506494 | n.a. |
| NMI | 2.03 | 9.4149 | 202.969 | 3E-40 | 3E-38 | 0.8181818 | n.a. |
| SAMSN1 | 2.178 | 9.3692 | 232.987 | 4E-40 | 3.3E-38 | 0.5194805 | n.a. |
| ILT11A | 2.031 | 9.2621 | 1377.7 | 6E-40 | 5.3E-38 | 0.5324675 | n.a. |
| RAP1B | 2.029 | 9.5901 | 174.926 | 8E-40 | 6.7E-38 | 0.8831169 | n.a. |
| ISG15 | 2.292 | 9.3193 | 286.7 | 2E-39 | 1.3E-37 | 0.4805195 | n.a. |
| CTNNAL1 | 2.465 | 9.2496 | 1241.98 | 2E-39 | 2.1E-37 | 0.4155844 | n.a. |
| ETS2 | 2.532 | 9.2476 | 1297.21 | 1E-38 | 1.1E-36 | 0.4675325 | n.a. |
| TBC1D8 | 2.396 | 9.2684 | 445.573 | 1E-38 | 1.2E-36 | 0.5454545 | n.a. |
| CYSTM1 | 1.352 | 9.3828 | 168.431 | 2E-38 | 1.6E-36 | 0.8571429 | n.a. |
| MSN | 1.883 | 9.6123 | 168.143 | 2E-38 | 1.9E-36 | 0.8571429 | n.a. |
| ENSECAG00000030777 | 1.751 | 9.3773 | 689.366 | 4E-38 | 3E-36 | 0.6103896 | n.a. |
| SLC7A7 | 1.951 | 9.4046 | 166.479 | 5E-38 | 4.2E-36 | 0.7662338 | n.a. |
| MHCB3 | 2.09 | 10.265 | 164.999 | 1E-37 | 8.8E-36 | 0.8831169 | n.a. |
| GUCY1B1 | 2.333 | 9.2412 | 1389.36 | 3E-37 | 2.6E-35 | 0.3246753 | n.a. |
| SRGN | 1.202 | 10.485 | 159.303 | 2E-36 | 1.5E-34 | 1 | n.a. |
| APP | 1.99 | 9.2487 | 1318.22 | 2E-36 | 1.8E-34 | 0.4155844 | surface |
| GPR31 | 2.127 | 9.2389 | 1269.8 | 5E-36 | 3.7E-34 | 0.2727273 | surface |
| UBD | 2.455 | 9.9806 | 155.484 | 1E-35 | 9.6E-34 | 0.6493506 | n.a. |
| SLC44A2 | 2.157 | 9.4299 | 181.989 | 4E-35 | 2.8E-33 | 0.4935065 | surface |
| ST6GALNAC2 | 2.035 | 9.2783 | 401.333 | 4E-34 | 3E-32 | 0.4805195 | n.a. |
| ENSECAG00000009285 | 2.163 | 9.2377 | 1404.69 | 1E-33 | 6.8E-32 | 0.3636364 | n.a. |
| SPI1 | 1.062 | 9.6618 | 191.927 | 3E-33 | 2.2E-31 | 0.961039 | n.a. |
| CSK | 2.192 | 9.389 | 177.722 | 4E-33 | 2.4E-31 | 0.6883117 | n.a. |
| DUSP12 | 2.352 | 9.3489 | 218.203 | 2E-32 | 1.1E-30 | 0.3896104 | n.a. |
| CYP4F22 | 2.152 | 9.2378 | 2399.38 | 7E-32 | 4.8E-30 | 0.3116883 | n.a. |
| AK3 | 2.07 | 9.4014 | 160.495 | 3E-31 | 1.6E-29 | 0.6883117 | n.a. |
| SDC3 | 2.103 | 9.2391 | 2379.71 | 1E-30 | 6.2E-29 | 0.2597403 | n.a. |
| COX17 | 1.417 | 10.369 | 132.864 | 1E-30 | 7E-29 | 0.974026 | n.a. |
| ENSECAG00000036100 | 1.181 | 9.3021 | 912.542 | 2E-30 | 1.3E-28 | 0.3636364 | n.a. |
| UCP2 | 1.293 | 9.9052 | 131.065 | 3E-30 | 1.7E-28 | 0.961039 | n.a. |
| CYTIP | 1.779 | 9.6111 | 131.015 | 3E-30 | 1.7E-28 | 0.7922078 | n.a. |

| | | | | | | | |
|---|---|---|---|---|---|---|---|
| ADGRE5 | 1.75 | 9.5791 | 130.026 | 5E-30 | 2.8E-28 | 0.7662338 | n.a. |
| LAMP1 | 1.299 | 9.6692 | 129.61 | 6E-30 | 3.4E-28 | 0.8571429 | surface |
| CYBA | 1.064 | 10.509 | 126.915 | 2E-29 | 1.3E-27 | 1 | n.a. |
| ENSECAG00000016543 | 2.041 | 9.8125 | 124.894 | 6E-29 | 3.6E-27 | 0.7272727 | n.a. |
| RF01956 | 1.894 | 9.2926 | 323.978 | 1E-28 | 5.6E-27 | 0.7532468 | n.a. |
| LYN | 1.655 | 9.3216 | 222.871 | 3E-28 | 1.9E-26 | 0.6623377 | surface |
| ENSECAG00000012830 | 2.044 | 9.263 | 605.683 | 4E-28 | 2.1E-26 | 0.2987013 | n.a. |
| HCLS1 | 1.726 | 9.4497 | 120.614 | 5E-28 | 3E-26 | 0.6883117 | n.a. |
| MAP7D3 | 1.897 | 9.41 | 125.363 | 9E-28 | 5.1E-26 | 0.5584416 | n.a. |
| RHOF | 1.967 | 9.4402 | 139.606 | 1E-27 | 7.5E-26 | 0.4155844 | n.a. |
| DHRS3 | 1.856 | 9.3111 | 289.262 | 2E-27 | 1.2E-25 | 0.6103896 | n.a. |
| PLD4 | 1.691 | 9.3359 | 278.059 | 3E-27 | 1.6E-25 | 0.7532468 | n.a. |
| DOK2 | 1.891 | 9.3423 | 221.489 | 5E-27 | 2.7E-25 | 0.4545455 | n.a. |
| RGS10 | 1.765 | 9.5903 | 116.102 | 5E-27 | 2.7E-25 | 0.5584416 | n.a. |
| MRPS6 | 1.978 | 9.4746 | 122.702 | 8E-27 | 4.2E-25 | 0.6493506 | n.a. |
| NAP1L1 | 1.54 | 10.092 | 115.165 | 8E-27 | 4.3E-25 | 0.8701299 | n.a. |
| ENSECAG00000033029 | 1.683 | 9.2619 | 370.742 | 9E-27 | 4.7E-25 | 0.4935065 | n.a. |
| ENSECAG00000031569 | 1.439 | 11.194 | 114.247 | 1E-26 | 6.7E-25 | 0.961039 | n.a. |
| PTPRC | 1.301 | 10.647 | 112.377 | 3E-26 | 1.7E-24 | 0.8961039 | surface |
| RASA3 | 1.993 | 9.5751 | 111.859 | 4E-26 | 2.2E-24 | 0.5064935 | n.a. |
| DUSP1 | 1.113 | 9.3756 | 111.395 | 5E-26 | 2.8E-24 | 0.6883117 | n.a. |
| RF00163 | 2.033 | 9.4218 | 138.944 | 7E-26 | 3.5E-24 | 0.4415584 | n.a. |
| ABCG2 | 1.965 | 9.262 | 481.114 | 9E-26 | 4.7E-24 | 0.3506494 | surface |
| IFIT2 | 1.425 | 9.3061 | 270.364 | 1E-25 | 6.9E-24 | 0.5454545 | n.a. |
| WIPF1 | 1.85 | 9.5089 | 108.21 | 3E-25 | 1.3E-23 | 0.5844156 | n.a. |
| PLEKHO2 | 1.801 | 9.3041 | 189.566 | 5E-25 | 2.5E-23 | 0.5454545 | n.a. |
| UGCG | 1.767 | 9.3115 | 189.833 | 1E-24 | 5.6E-23 | 0.3506494 | n.a. |
| DBI | 1.51 | 9.673 | 102.443 | 5E-24 | 2.4E-22 | 0.8701299 | n.a. |
| GIMAP7 | 1.631 | 10.513 | 102.441 | 5E-24 | 2.4E-22 | 0.6493506 | n.a. |
| ENSECAG00000039428 | 1.046 | 11.047 | 101.944 | 6E-24 | 3E-22 | 1 | n.a. |
| LRRFIP1 | 1.476 | 9.8063 | 100.34 | 1E-23 | 6.7E-22 | 0.9090909 | n.a. |
| ENSECAG00000028581 | 1.942 | 9.2649 | 427.338 | 2E-23 | 1.1E-21 | 0.4285714 | n.a. |
| TBC1D9 | 1.799 | 9.2457 | 936.062 | 4E-23 | 2E-21 | 0.3636364 | n.a. |
| DRA | 1.05 | 11.742 | 97.73 | 5E-23 | 2.4E-21 | 1 | n.a. |
| ENSECAG00000033034 | 1.658 | 9.2773 | 649.451 | 7E-23 | 3.2E-21 | 0.4155844 | n.a. |
| ITM2C | 1.693 | 9.5994 | 96.828 | 8E-23 | 3.8E-21 | 0.4935065 | surface |
| NRN1 | 1.905 | 9.2707 | 460.486 | 1E-22 | 4.7E-21 | 0.3896104 | surface |
| ENSECAG00000031156 | 1.601 | 9.3403 | 303.932 | 1E-22 | 5.5E-21 | 0.4935065 | n.a. |
| ENSECAG00000007681 | 1.583 | 9.6603 | 100.637 | 2E-22 | 7E-21 | 0.5584416 | n.a. |
| ENSECAG00000035431 | 1.346 | 9.323 | 447.092 | 3E-22 | 1.3E-20 | 0.6623377 | n.a. |
| SIVA1 | 1.652 | 9.5677 | 93.7845 | 4E-22 | 1.7E-20 | 0.7532468 | n.a. |
| ENSECAG00000040532 | 1.939 | 9.3029 | 211.721 | 1E-21 | 5.5E-20 | 0.4025974 | n.a. |
| TIFA | 1.958 | 10.029 | 91.1759 | 1E-21 | 6.1E-20 | 0.6363636 | n.a. |
| DRAM2 | 1.874 | 9.402 | 117.493 | 2E-21 | 7.2E-20 | 0.4675325 | n.a. |
| TMC6 | 1.914 | 9.4027 | 118.29 | 6E-21 | 2.5E-19 | 0.3246753 | n.a. |
| ZYX | 1.31 | 9.4366 | 87.6902 | 8E-21 | 3.4E-19 | 0.8311688 | n.a. |
| LAP3 | 1.697 | 9.4403 | 104.644 | 2E-20 | 7.8E-19 | 0.5974026 | n.a. |
| INPP5F | 1.964 | 9.2549 | 478.557 | 3E-20 | 1.2E-18 | 0.2727273 | n.a. |
| DRB | 1.083 | 11.205 | 81.0098 | 2E-19 | 9.5E-18 | 0.974026 | n.a. |
| Eqca-DQB1 | 1.152 | 10.925 | 80.939 | 2E-19 | 9.8E-18 | 0.974026 | n.a. |
| STARD7 | 1.9 | 9.3039 | 190.539 | 2E-19 | 9.9E-18 | 0.4415584 | n.a. |
| ENSECAG00000019430 | 1.797 | 9.3323 | 151.829 | 3E-19 | 1E-17 | 0.4545455 | n.a. |
| SLAMF7 | 1.582 | 9.2845 | 274.582 | 3E-19 | 1.4E-17 | 0.4025974 | surface |
| ENSECAG00000020991 | 1.616 | 9.2681 | 366.104 | 4E-19 | 1.4E-17 | 0.3896104 | n.a. |
| KLF4 | 1.688 | 9.2595 | 695.536 | 4E-19 | 1.4E-17 | 0.3506494 | n.a. |
| ATF3 | 1.793 | 9.2384 | 1181 | 6E-19 | 2.3E-17 | 0.2727273 | n.a. |

| | | | | | | | |
|---|---|---|---|---|---|---|---|
| CYTH4 | 1.685 | 9.2629 | 256.729 | 1E-18 | 4.1E-17 | 0.4805195 | n.a. |
| KIF22 | 1.791 | 9.2545 | 446.729 | 1E-18 | 5.4E-17 | 0.2857143 | n.a. |
| SAT1 | 1.003 | 9.7271 | 76.9138 | 2E-18 | 7.1E-17 | 0.961039 | n.a. |
| PRDX1 | 1.533 | 9.8857 | 76.3286 | 3E-18 | 9.4E-17 | 0.7272727 | n.a. |
| SEPT9 | 1.534 | 9.7694 | 76.203 | 3E-18 | 1E-16 | 0.6753247 | n.a. |
| PAM | 1.825 | 9.2453 | 620.922 | 3E-18 | 1E-16 | 0.2727273 | surface |
| TIMP1 | 1.397 | 9.3333 | 219.108 | 3E-18 | 1.3E-16 | 0.3376623 | n.a. |
| BIN2 | 1.292 | 9.7163 | 75.5979 | 4E-18 | 1.3E-16 | 0.8961039 | n.a. |
| ENSECAG00000036034 | 1.206 | 9.8269 | 75.2205 | 4E-18 | 1.6E-16 | 0.9220779 | n.a. |
| ENSECAG00000027864 | 1.33 | 9.2631 | 534.355 | 9E-18 | 3.1E-16 | 0.3376623 | n.a. |
| ENSECAG00000039324 | 1.108 | 10.369 | 73.129 | 1E-17 | 4.5E-16 | 0.9480519 | n.a. |
| CXCL16 | 1.671 | 9.3053 | 142.732 | 1E-17 | 5.2E-16 | 0.3766234 | surface |
| CORO1A | 1.125 | 10.247 | 70.9559 | 4E-17 | 1.3E-15 | 0.8311688 | n.a. |
| ASCL4 | 1.611 | 9.269 | 519.173 | 6E-17 | 2.1E-15 | 0.4155844 | n.a. |
| ACP5 | 1.611 | 9.7992 | 70.0896 | 6E-17 | 2.1E-15 | 0.5454545 | n.a. |
| ENSECAG00000029716 | 1.322 | 9.8771 | 70.6284 | 7E-17 | 2.6E-15 | 0.6363636 | n.a. |
| ENSECAG00000032131 | 1.56 | 9.3494 | 98.514 | 8E-17 | 2.7E-15 | 0.4935065 | n.a. |
| NEDD9 | 1.756 | 9.2911 | 193.39 | 8E-17 | 2.9E-15 | 0.2987013 | n.a. |
| C20H6orf62 | 1.361 | 9.3936 | 77.5008 | 9E-17 | 3.1E-15 | 0.6363636 | n.a. |
| KLF2 | 1.488 | 9.4082 | 74.1812 | 1E-16 | 5E-15 | 0.4935065 | n.a. |
| ENSECAG00000009162 | 1.023 | 10.473 | 67.8465 | 2E-16 | 6.2E-15 | 0.9350649 | n.a. |
| SQSTM1 | 1.513 | 9.456 | 67.7442 | 2E-16 | 6.5E-15 | 0.5064935 | n.a. |
| JUP | 1.678 | 9.251 | 502.627 | 5E-16 | 1.6E-14 | 0.2597403 | n.a. |
| TUBA4A | 1.346 | 9.5479 | 65.6177 | 6E-16 | 1.9E-14 | 0.4805195 | n.a. |
| P2RY2 | 1.589 | 9.2603 | 497.402 | 7E-16 | 2.3E-14 | 0.3116883 | surface |
| EMP3 | 1.381 | 10.126 | 64.7663 | 9E-16 | 2.8E-14 | 0.7142857 | surface |
| IFIT3 | 1.397 | 9.2748 | 207.697 | 1E-15 | 4.3E-14 | 0.2987013 | n.a. |
| ENSECAG00000034339 | 1.29 | 10.188 | 63.7575 | 1E-15 | 4.6E-14 | 0.7402597 | n.a. |
| EGLN3 | 1.395 | 9.3249 | 195.173 | 2E-15 | 6.8E-14 | 0.6103896 | n.a. |
| NR1H3 | 1.665 | 9.244 | 871.473 | 3E-15 | 1E-13 | 0.2597403 | n.a. |
| TUBA1A | 1.437 | 10.091 | 61.8994 | 4E-15 | 1.2E-13 | 0.5844156 | n.a. |
| NDUFA12 | 1.525 | 9.4672 | 68.83 | 5E-15 | 1.6E-13 | 0.5064935 | n.a. |
| PKIB | 1.102 | 9.2759 | 408.369 | 1E-14 | 4.2E-13 | 0.2857143 | n.a. |
| STK10 | 1.344 | 9.392 | 67.6323 | 2E-14 | 4.8E-13 | 0.5974026 | n.a. |
| XAF1 | 1.558 | 9.2818 | 158.781 | 2E-14 | 5.4E-13 | 0.2597403 | n.a. |
| ENSECAG00000040180 | 1.216 | 9.7881 | 58.8333 | 2E-14 | 5.4E-13 | 0.6363636 | n.a. |
| RF00411 | 1.373 | 9.4519 | 58.4441 | 2E-14 | 6.6E-13 | 0.4545455 | n.a. |
| RPS6KA1 | 1.459 | 9.4019 | 74.9202 | 3E-14 | 8.9E-13 | 0.5714286 | n.a. |
| PLEKHO1 | 1.415 | 9.3156 | 157.445 | 3E-14 | 9.9E-13 | 0.4675325 | n.a. |
| ENSECAG00000009794 | 1.413 | 9.3398 | 81.2009 | 3E-14 | 1.1E-12 | 0.4935065 | n.a. |
| HYPK | 1.473 | 9.4628 | 63.3784 | 4E-14 | 1.1E-12 | 0.5974026 | n.a. |
| CHD9 | 1.359 | 9.4262 | 64.6316 | 4E-14 | 1.3E-12 | 0.5974026 | n.a. |
| BSG | 1.207 | 9.6757 | 56.9782 | 5E-14 | 1.3E-12 | 0.6623377 | surface |
| ENSECAG00000000775 | 1.215 | 9.4621 | 73.226 | 6E-14 | 1.6E-12 | 0.6623377 | n.a. |
| ARL6IP1 | 1.329 | 9.5522 | 56.4419 | 6E-14 | 1.7E-12 | 0.6623377 | n.a. |
| JUNB | 1.134 | 9.5395 | 56.1194 | 7E-14 | 2E-12 | 0.8051948 | n.a. |
| DIPK2A | 1.206 | 9.2586 | 294.557 | 8E-14 | 2.4E-12 | 0.3246753 | n.a. |
| DQB | 1.063 | 10.011 | 54.5329 | 2E-13 | 4.5E-12 | 0.8181818 | n.a. |
| TBXAS1 | 1.219 | 9.2862 | 463.338 | 2E-13 | 4.5E-12 | 0.4415584 | n.a. |
| PAQR4 | 1.426 | 9.2942 | 142.322 | 2E-13 | 5.5E-12 | 0.2597403 | n.a. |
| HPRT1 | 1.198 | 9.5077 | 53.3738 | 3E-13 | 8E-12 | 0.5844156 | n.a. |
| ENSECAG00000011858 | 1.338 | 9.5887 | 53.1453 | 3E-13 | 8.9E-12 | 0.6233766 | n.a. |
| GIMAP6 | 1.273 | 9.7575 | 52.8487 | 4E-13 | 1E-11 | 0.6753247 | n.a. |
| NAAA | 1.206 | 9.5818 | 52.774 | 4E-13 | 1.1E-11 | 0.5974026 | n.a. |
| NDE1 | 1.485 | 9.2796 | 147.766 | 5E-13 | 1.4E-11 | 0.3376623 | n.a. |
| PNRC1 | 1.164 | 9.9575 | 51.8545 | 6E-13 | 1.7E-11 | 0.8181818 | n.a. |

| | | | | | | | |
|---|---|---|---|---|---|---|---|
| MEF2A | 1.452 | 9.302 | 108.276 | 6E-13 | 1.7E-11 | 0.2987013 | n.a. |
| CHMP4B | 1.217 | 9.4518 | 51.4163 | 8E-13 | 2.1E-11 | 0.6623377 | n.a. |
| PTP4A2 | 1.225 | 9.7459 | 50.4964 | 1E-12 | 3.3E-11 | 0.7402597 | n.a. |
| CD48 | 1.01 | 9.9941 | 50.4224 | 1E-12 | 3.4E-11 | 0.8701299 | surface |
| ITGA4 | 1.303 | 9.465 | 54.7847 | 2E-12 | 4.2E-11 | 0.5194805 | surface |
| TMEM14C | 1.123 | 9.6674 | 49.8315 | 2E-12 | 4.5E-11 | 0.7792208 | n.a. |
| CYTH1 | 1.381 | 9.3765 | 63.2125 | 2E-12 | 4.7E-11 | 0.3896104 | n.a. |
| TLR7 | 1.283 | 9.2655 | 297.975 | 2E-12 | 4.8E-11 | 0.3506494 | surface |
| KCNE3 | 1.251 | 9.2985 | 362.922 | 3E-12 | 6.7E-11 | 0.4155844 | n.a. |
| ITGB1 | 1.39 | 9.5309 | 48.665 | 3E-12 | 7.9E-11 | 0.3896104 | surface |
| ARRB2 | 1.163 | 9.4662 | 50.9303 | 4E-12 | 9.8E-11 | 0.5064935 | n.a. |
| LMO2 | 1.339 | 9.3122 | 153.443 | 4E-12 | 1.1E-10 | 0.2987013 | n.a. |
| FCGRT | 1.165 | 9.3372 | 106.431 | 5E-12 | 1.2E-10 | 0.4545455 | surface |
| FYN | 1.461 | 9.4235 | 66.717 | 5E-12 | 1.2E-10 | 0.3246753 | n.a. |
| C9orf116 | 1.313 | 9.3011 | 130.871 | 5E-12 | 1.3E-10 | 0.3116883 | n.a. |
| APRT | 1.224 | 9.6859 | 46.2879 | 1E-11 | 2.5E-10 | 0.6883117 | n.a. |
| PSMB10 | 1.185 | 9.7781 | 45.7203 | 1E-11 | 3.4E-10 | 0.7012987 | n.a. |
| ENSECAG00000038583 | 1.401 | 9.3387 | 81.1365 | 2E-11 | 4.1E-10 | 0.3376623 | n.a. |
| PYCARD | 1.184 | 9.489 | 65.9729 | 2E-11 | 4.3E-10 | 0.6493506 | n.a. |
| LUZP6 | 1.209 | 9.4318 | 49.1396 | 2E-11 | 4.7E-10 | 0.5194805 | n.a. |
| ENSECAG00000034395 | 1.326 | 9.2877 | 160.576 | 2E-11 | 4.8E-10 | 0.2857143 | n.a. |
| MAP3K1 | 1.412 | 9.396 | 57.5088 | 2E-11 | 5E-10 | 0.3246753 | n.a. |
| MAGOH | 1.408 | 9.4803 | 50.0174 | 2E-11 | 5.4E-10 | 0.5194805 | n.a. |
| ZNHIT1 | 1.264 | 9.4252 | 52.3946 | 3E-11 | 6.1E-10 | 0.4805195 | n.a. |
| SH2D3C | 1.313 | 9.2916 | 94.0357 | 3E-11 | 7.3E-10 | 0.2727273 | n.a. |
| MYD88 | 1.179 | 9.3632 | 55.5676 | 4E-11 | 8.7E-10 | 0.5324675 | n.a. |
| MRPL35 | 1.524 | 9.3167 | 87.5982 | 5E-11 | 1.2E-09 | 0.3636364 | n.a. |
| PYCR1 | 1.188 | 9.4844 | 46.3481 | 5E-11 | 1.2E-09 | 0.6623377 | n.a. |
| ITGAL | 1.21 | 9.4047 | 52.9979 | 7E-11 | 1.5E-09 | 0.4545455 | surface |
| ECHDC2 | 1.269 | 9.3064 | 88.6866 | 7E-11 | 1.7E-09 | 0.4155844 | n.a. |
| SNAP23 | 1.433 | 9.3138 | 85.016 | 8E-11 | 1.9E-09 | 0.3376623 | n.a. |
| ENSECAG00000038252 | 1.411 | 9.397 | 64.4756 | 8E-11 | 1.9E-09 | 0.3116883 | n.a. |
| UPF2 | 1.428 | 9.339 | 71.4676 | 1E-10 | 2.4E-09 | 0.3376623 | n.a. |
| PNKD | 1.174 | 9.3004 | 90.4547 | 1E-10 | 2.5E-09 | 0.3116883 | n.a. |
| MYOF | 1.187 | 9.247 | 325.332 | 1E-10 | 3.1E-09 | 0.2597403 | surface |
| ENSECAG00000019309 | 1.049 | 9.3991 | 72.4521 | 2E-10 | 4.3E-09 | 0.3246753 | n.a. |
| CYFIP1 | 1.107 | 9.2506 | 426.149 | 2E-10 | 4.3E-09 | 0.2597403 | n.a. |
| ACAA1 | 1.397 | 9.2784 | 116.627 | 2E-10 | 4.9E-09 | 0.3246753 | n.a. |
| NDUFS5 | 1.149 | 9.6423 | 40.2394 | 2E-10 | 4.9E-09 | 0.7402597 | n.a. |
| RALB | 1.162 | 9.3135 | 66.998 | 2E-10 | 5.2E-09 | 0.4155844 | n.a. |
| ENSECAG00000006071 | 1.309 | 9.6095 | 39.7681 | 3E-10 | 6.2E-09 | 0.4805195 | n.a. |
| PTBP3 | 1.337 | 9.4001 | 55.0067 | 3E-10 | 6.3E-09 | 0.5194805 | n.a. |
| SLC3A2 | 1.397 | 9.3565 | 63.5194 | 3E-10 | 6.9E-09 | 0.3766234 | surface |
| DBNL | 1.041 | 9.4448 | 46.1813 | 4E-10 | 8.3E-09 | 0.7532468 | n.a. |
| ENSECAG00000007663 | 1.166 | 9.734 | 47.4024 | 5E-10 | 1.1E-08 | 0.3766234 | n.a. |
| PDK1 | 1.295 | 9.2628 | 280.685 | 6E-10 | 1.2E-08 | 0.2727273 | n.a. |
| GNA15 | 1.235 | 9.3122 | 68.3887 | 7E-10 | 1.4E-08 | 0.4285714 | n.a. |
| RF01957 | 1.112 | 9.2457 | 361.462 | 7E-10 | 1.5E-08 | 0.2987013 | n.a. |
| TTC9C | 1.239 | 9.3445 | 50.2631 | 8E-10 | 1.6E-08 | 0.3376623 | n.a. |
| PEBP1 | 1.069 | 9.6042 | 37.6136 | 9E-10 | 1.8E-08 | 0.6233766 | n.a. |
| CEPT1 | 1.388 | 9.3104 | 76.4037 | 1E-09 | 1.9E-08 | 0.2857143 | n.a. |
| TK2 | 1.346 | 9.3075 | 86.1482 | 1E-09 | 2.4E-08 | 0.2597403 | n.a. |
| KIAA0513 | 1.008 | 9.2674 | 157.581 | 1E-09 | 2.4E-08 | 0.3116883 | n.a. |
| OSTC | 1.116 | 9.5663 | 36.8149 | 1E-09 | 2.6E-08 | 0.6493506 | n.a. |
| MIF4GD | 1.215 | 9.3713 | 46.0869 | 1E-09 | 2.6E-08 | 0.3506494 | n.a. |
| ENSECAG00000037215 | 1.204 | 9.3741 | 50.8507 | 2E-09 | 3.3E-08 | 0.4545455 | n.a. |

| | | | | | | | |
|---|---|---|---|---|---|---|---|
| YWHAH | 1.063 | 9.5952 | 35.9617 | 2E-09 | 4E-08 | 0.5844156 | n.a. |
| ACSL4 | 1.274 | 9.2833 | 103.088 | 3E-09 | 5.2E-08 | 0.2727273 | n.a. |
| eca-mir-9051 | 1.286 | 9.2854 | 101.884 | 3E-09 | 5.5E-08 | 0.3506494 | n.a. |
| OSBPL8 | 1.106 | 9.334 | 45.8943 | 3E-09 | 5.6E-08 | 0.3896104 | n.a. |
| DNAJC8 | 1.223 | 9.5664 | 35.0834 | 3E-09 | 6.1E-08 | 0.5064935 | n.a. |
| MEF2C | 1.075 | 9.4205 | 102.751 | 4E-09 | 6.8E-08 | 0.4935065 | n.a. |
| NTAN1 | 1.139 | 9.412 | 38.7229 | 4E-09 | 7.8E-08 | 0.4285714 | n.a. |
| BLVRA | 1.222 | 9.2882 | 104.634 | 5E-09 | 9.1E-08 | 0.2857143 | n.a. |
| MPP1 | 1.328 | 9.3612 | 64.5074 | 6E-09 | 1.1E-07 | 0.3636364 | n.a. |
| CMTM3 | 1.105 | 9.4968 | 37.4412 | 7E-09 | 1.2E-07 | 0.3376623 | n.a. |
| REL | 1.215 | 9.3166 | 60.0134 | 7E-09 | 1.3E-07 | 0.2987013 | n.a. |
| AKIRIN2 | 1.135 | 9.4269 | 37.3274 | 8E-09 | 1.4E-07 | 0.4545455 | n.a. |
| ENSA | 1.072 | 9.671 | 33.1585 | 9E-09 | 1.6E-07 | 0.6363636 | n.a. |
| CHCHD7 | 1.159 | 9.4135 | 41.3727 | 1E-08 | 2.1E-07 | 0.3636364 | n.a. |
| RARA | 1.057 | 9.3162 | 52.4451 | 1E-08 | 2.1E-07 | 0.5064935 | n.a. |
| PHF20 | 1.144 | 9.302 | 57.5442 | 1E-08 | 2.5E-07 | 0.2727273 | n.a. |
| NET1 | 1.158 | 9.27 | 121.945 | 1E-08 | 2.6E-07 | 0.3766234 | n.a. |
| EIF4E1B | 1.308 | 9.2561 | 275.574 | 2E-08 | 2.9E-07 | 0.3766234 | n.a. |
| ENSECAG00000037904 | 1.166 | 9.482 | 34.6833 | 2E-08 | 2.9E-07 | 0.3506494 | n.a. |
| PTGER4 | 1.124 | 9.3476 | 51.1479 | 2E-08 | 2.9E-07 | 0.2727273 | surface |
| NDUFS2 | 1.017 | 9.5195 | 33.2995 | 2E-08 | 2.9E-07 | 0.5454545 | n.a. |
| MIER1 | 1.158 | 9.3938 | 37.9466 | 2E-08 | 4.1E-07 | 0.3766234 | n.a. |
| ENSECAG00000019274 | 1.026 | 9.4154 | 32.8137 | 2E-08 | 4.3E-07 | 0.4935065 | n.a. |
| FAM89B | 1.33 | 9.3314 | 70.044 | 3E-08 | 4.4E-07 | 0.3116883 | n.a. |
| ARHGAP15 | 1.12 | 9.6767 | 30.897 | 3E-08 | 4.7E-07 | 0.4935065 | n.a. |
| SDHC | 1.147 | 9.4385 | 37.6579 | 3E-08 | 4.7E-07 | 0.5194805 | n.a. |
| LMAN2 | 1.038 | 9.4812 | 30.6994 | 3E-08 | 5.2E-07 | 0.5064935 | surface |
| UMAD1 | 1.27 | 9.3877 | 46.6215 | 3E-08 | 5.3E-07 | 0.3376623 | n.a. |
| ACSL5 | 1.141 | 9.4151 | 37.9519 | 3E-08 | 5.4E-07 | 0.4155844 | n.a. |
| ARHGEF1 | 1.212 | 9.4652 | 34.6016 | 3E-08 | 5.5E-07 | 0.3506494 | n.a. |
| GARS | 1.221 | 9.3797 | 44.4266 | 3E-08 | 5.7E-07 | 0.3636364 | n.a. |
| LGALS3BP | 1.093 | 9.4204 | 37.6097 | 4E-08 | 6.6E-07 | 0.3896104 | n.a. |
| MRTFA | 1.203 | 9.2972 | 66.9533 | 5E-08 | 9.1E-07 | 0.2727273 | n.a. |
| MCUB | 1.196 | 9.3094 | 74.097 | 6E-08 | 9.3E-07 | 0.3116883 | n.a. |
| POMP | 1.048 | 9.5439 | 31.2134 | 6E-08 | 9.4E-07 | 0.6623377 | n.a. |
| FUT8 | 1.23 | 9.3425 | 53.0264 | 6E-08 | 9.8E-07 | 0.2857143 | n.a. |
| VAMP5 | 1.006 | 9.6046 | 29.2007 | 7E-08 | 1.1E-06 | 0.5454545 | n.a. |
| SNRPG | 1.075 | 9.559 | 32.6952 | 7E-08 | 1.1E-06 | 0.5454545 | n.a. |
| SUN2 | 1.224 | 9.5359 | 29.7655 | 7E-08 | 1.1E-06 | 0.3246753 | n.a. |
| IER5 | 1.134 | 9.2564 | 140.034 | 7E-08 | 1.2E-06 | 0.2597403 | n.a. |
| FUCA1 | 1.04 | 9.3678 | 36.6602 | 8E-08 | 1.2E-06 | 0.2987013 | n.a. |
| MACF1 | 1.095 | 9.3902 | 34.8226 | 8E-08 | 1.2E-06 | 0.2987013 | n.a. |
| RAP2B | 1.213 | 9.3226 | 57.1075 | 1E-07 | 2.3E-06 | 0.2857143 | n.a. |
| ENSECAG00000002652 | 1.208 | 9.2884 | 82.6234 | 1E-07 | 2.3E-06 | 0.2987013 | n.a. |
| SARNP | 1.045 | 9.47 | 27.6057 | 1E-07 | 2.3E-06 | 0.4025974 | n.a. |
| SYNGR2 | 1.149 | 9.3319 | 74.3908 | 2E-07 | 2.7E-06 | 0.4155844 | n.a. |
| UBE2E2 | 1.092 | 9.2944 | 57.759 | 2E-07 | 3.3E-06 | 0.3246753 | n.a. |
| CHMP5 | 1.073 | 9.3711 | 33.1327 | 2E-07 | 3.7E-06 | 0.3246753 | n.a. |
| TRAF3IP3 | 1.151 | 9.801 | 26.2009 | 3E-07 | 4.7E-06 | 0.3636364 | n.a. |
| CCDC12 | 1.022 | 9.3727 | 35.1056 | 4E-07 | 5.5E-06 | 0.4155844 | n.a. |
| VAMP4 | 1.173 | 9.2915 | 72.6805 | 4E-07 | 6.2E-06 | 0.2857143 | n.a. |
| SH3KBP1 | 1.096 | 9.4466 | 30.6863 | 4E-07 | 6.5E-06 | 0.3376623 | n.a. |
| SNU13 | 1.016 | 9.6007 | 25.506 | 4E-07 | 6.6E-06 | 0.4285714 | n.a. |
| SFXN3 | 1.166 | 9.3338 | 52.2899 | 5E-07 | 6.7E-06 | 0.2987013 | n.a. |
| MAP3K8 | 1.009 | 9.3226 | 59.702 | 5E-07 | 7.3E-06 | 0.3376623 | n.a. |
| ZBTB7A | 1.105 | 9.304 | 50.1357 | 7E-07 | 1E-05 | 0.3246753 | n.a. |

| | | | | | | | |
|---|---|---|---|---|---|---|---|
| LRPAP1 | 1.097 | 9.3914 | 34.0488 | 9E-07 | 1.2E-05 | 0.3246753 | n.a. |
| GNAQ | 1.172 | 9.27 | 114.815 | 9E-07 | 1.2E-05 | 0.2727273 | n.a. |
| IL10RA | 1.087 | 9.319 | 54.6258 | 9E-07 | 1.3E-05 | 0.4155844 | surface |
| EQMHCC1 | 1.004 | 9.6459 | 23.7874 | 1E-06 | 1.5E-05 | 0.4285714 | n.a. |
| UQCRC2 | 1.133 | 9.3691 | 39.3232 | 1E-06 | 1.6E-05 | 0.3116883 | n.a. |
| MRPS14 | 1.124 | 9.367 | 38.5716 | 2E-06 | 2.1E-05 | 0.3376623 | n.a. |
| MPHOSPH8 | 1.032 | 9.4682 | 24.5163 | 2E-06 | 2.7E-05 | 0.3636364 | n.a. |
| SERINC1 | 1.058 | 9.3567 | 34.6786 | 2E-06 | 3.3E-05 | 0.3636364 | surface |
| PDHB | 1.109 | 9.3766 | 34.4697 | 3E-06 | 3.5E-05 | 0.2727273 | n.a. |
| RASGRP2 | 1.005 | 9.4643 | 25.2938 | 3E-06 | 4.3E-05 | 0.3636364 | n.a. |
| PSMA6 | 1.025 | 9.4544 | 28.0421 | 4E-06 | 5.5E-05 | 0.4675325 | n.a. |
| TSR2 | 1.042 | 9.3864 | 31.1982 | 6E-06 | 7E-05 | 0.2987013 | n.a. |
| PFDN6 | 1.156 | 9.3579 | 37.6913 | 7E-06 | 8.3E-05 | 0.2597403 | n.a. |
| DNAJC2 | 1.029 | 9.4123 | 28.5446 | 7E-06 | 8.6E-05 | 0.2987013 | n.a. |
| DNASE1L1 | 1.074 | 9.2839 | 58.1919 | 8E-06 | 1E-04 | 0.2987013 | n.a. |
| UBE2N | 1.012 | 9.5201 | 24.3112 | 9E-06 | 0.00011 | 0.3636364 | n.a. |
| SSNA1 | 1.052 | 9.3399 | 37.2934 | 1E-05 | 0.00013 | 0.2597403 | n.a. |
| VTA1 | 1.054 | 9.3204 | 40.0515 | 2E-05 | 0.00019 | 0.2727273 | n.a. |

| genes | logFC | logCPM | F | PValue | FDR | percent.exp | Surfacome.Label |
|---|---|---|---|---|---|---|---|
| S100A8 | 5.766 | 9.3765 | 4777.69 | 0 | 0 | 0.4857143 | n.a. |
| S100A12 | 5.418 | 10.309 | 5011.46 | 0 | 0 | 0.4285714 | n.a. |
| ENSECAG00000010117 | 4.927 | 9.501 | 3327.23 | 0 | 0 | 0.6571429 | n.a. |
| ENSECAG00000030548 | 6.134 | 9.2647 | 5095.44 | 2E-292 | 4E-289 | 0.4 | n.a. |
| C1H15orf48 | 4.51 | 9.6929 | 1553.49 | 1E-277 | 2E-274 | 1 | n.a. |
| ENSECAG00000039739 | 7.579 | 9.2496 | 8791.52 | 1E-252 | 2E-249 | 1 | n.a. |
| ENSECAG00000032321 | 7.882 | 9.2472 | 4082.34 | 7E-217 | 8E-214 | 0.9428571 | n.a. |
| ENSECAG00000039383 | 4.943 | 9.2686 | 8996.34 | 2E-200 | 2E-197 | 0.2571429 | n.a. |
| S100P | 3.499 | 9.8205 | 2334.96 | 3E-199 | 2E-196 | 1 | n.a. |
| RGS2 | 4.347 | 9.2835 | 1978.93 | 3E-129 | 3E-126 | 0.9428571 | n.a. |
| TSPO | 3.378 | 9.7323 | 604.051 | 2E-121 | 1E-118 | 1 | n.a. |
| LYZ | 2.324 | 11.749 | 480.893 | 7E-106 | 4E-103 | 0.8 | n.a. |
| DUSP1 | 3.305 | 9.3756 | 462.181 | 6E-94 | 3.1E-91 | 0.7428571 | n.a. |
| ENSECAG00000036100 | 4.086 | 9.3021 | 2373.04 | 4E-90 | 2.1E-87 | 0.7714286 | n.a. |
| ENSECAG00000016948 | 2.165 | 11.316 | 387.358 | 9E-86 | 4.3E-83 | 1 | n.a. |
| IFIT2 | 3.795 | 9.3061 | 911.994 | 2E-82 | 6.9E-80 | 0.6285714 | n.a. |
| IDO1 | 4.891 | 9.5783 | 450.597 | 2E-81 | 9E-79 | 0.5428571 | n.a. |
| ENSECAG00000019932 | 3.878 | 9.3315 | 658.62 | 1E-80 | 5.5E-78 | 0.8571429 | n.a. |
| G0S2 | 4.107 | 9.2375 | 2716.07 | 1E-72 | 3.3E-70 | 0.4285714 | n.a. |
| ENSECAG00000030745 | 1.931 | 11.502 | 311.762 | 2E-69 | 4.9E-67 | 1 | n.a. |
| PADI4 | 4.053 | 9.2553 | 2705.01 | 2E-68 | 6.4E-66 | 0.7714286 | n.a. |
| ENSECAG00000033029 | 3.785 | 9.2619 | 923.372 | 1E-62 | 3.2E-60 | 0.4571429 | n.a. |
| HCAR1 | 4.88 | 9.239 | 3129.28 | 2E-58 | 3.3E-56 | 0.6571429 | surface |
| ENSECAG00000036967 | 3.88 | 9.2599 | 1754.4 | 2E-58 | 4.8E-56 | 0.8571429 | n.a. |
| ENSECAG00000029928 | 3.23 | 9.2723 | 2721.72 | 1E-53 | 2E-51 | 0.4285714 | n.a. |
| FOS | 3.129 | 9.3156 | 626.226 | 5E-51 | 6.6E-49 | 0.5428571 | n.a. |
| ENSECAG00000030271 | 3.025 | 9.3599 | 382.999 | 3E-49 | 4.1E-47 | 0.8 | n.a. |
| HIST1H1C | 3.781 | 9.2575 | 887.335 | 7E-47 | 8.6E-45 | 0.3428571 | n.a. |
| SERPINB1 | 3.02 | 9.4507 | 285.161 | 1E-46 | 1.4E-44 | 0.5428571 | n.a. |
| SRGN | 2.01 | 10.485 | 205.066 | 2E-46 | 2.4E-44 | 0.9714286 | n.a. |
| ILT11B | 3.621 | 9.2698 | 1398.4 | 2E-45 | 2.4E-43 | 0.6857143 | n.a. |
| LST1 | 2.989 | 9.329 | 428.902 | 2E-43 | 1.6E-41 | 0.7714286 | n.a. |
| ENSECAG00000020136 | 2.824 | 9.7014 | 199.948 | 9E-43 | 9.3E-41 | 0.7142857 | n.a. |
| HMGB2 | 3.082 | 10.129 | 185.796 | 3E-42 | 3.3E-40 | 0.7428571 | n.a. |
| MMP8 | 4.78 | 9.2382 | 1949.27 | 4E-42 | 4.3E-40 | 0.5428571 | n.a. |
| VNN2 | 3.274 | 9.2761 | 808.813 | 8E-42 | 7.1E-40 | 0.7142857 | surface |
| ENSECAG00000035094 | 3.132 | 9.2863 | 627.16 | 4E-40 | 4E-38 | 0.6857143 | n.a. |
| eca-mir-223 | 3.243 | 9.2778 | 961.818 | 4E-39 | 3.7E-37 | 0.6571429 | n.a. |
| EHD1 | 3.468 | 9.4259 | 208.887 | 8E-39 | 7.2E-37 | 0.6 | n.a. |
| C5AR1 | 2.581 | 9.3446 | 855.87 | 1E-38 | 1.1E-36 | 0.8 | surface |
| SNX10 | 3.321 | 9.274 | 856.719 | 3E-38 | 2.4E-36 | 0.5428571 | n.a. |
| PGD | 2.909 | 9.3424 | 371.077 | 2E-37 | 1.3E-35 | 0.6857143 | n.a. |
| DGAT2 | 3.374 | 9.2504 | 1134.46 | 6E-37 | 4.4E-35 | 0.5428571 | n.a. |
| MXD1 | 3.089 | 9.3408 | 227.932 | 6E-37 | 4.7E-35 | 0.6571429 | n.a. |
| TREM1 | 2.433 | 9.4852 | 956.954 | 1E-35 | 9E-34 | 0.7428571 | surface |
| VASP | 3.203 | 9.4309 | 186.208 | 9E-34 | 6.3E-32 | 0.7142857 | n.a. |
| THY1 | 3.55 | 9.2416 | 1725 | 4E-33 | 2.7E-31 | 0.4285714 | surface |
| JAML | 2.863 | 9.299 | 489.131 | 7E-33 | 4.8E-31 | 0.6285714 | n.a. |
| TALDO1 | 2.342 | 9.9099 | 141.069 | 2E-32 | 1.2E-30 | 0.9714286 | n.a. |
| ARHGDIB | 1.781 | 11.255 | 138.743 | 6E-32 | 3.9E-30 | 0.9428571 | n.a. |
| CEBPB | 2.445 | 9.407 | 379.956 | 2E-31 | 1E-29 | 0.6 | n.a. |
| RHOB | 2.929 | 9.2893 | 352.961 | 2E-31 | 1.4E-29 | 0.5428571 | n.a. |
| SOD2 | 2.955 | 9.3528 | 195.302 | 6E-30 | 3.6E-28 | 0.5714286 | n.a. |
| IFIT3 | 3.235 | 9.2748 | 450.706 | 7E-30 | 4.2E-28 | 0.3428571 | n.a. |
| GADD45A | 2.598 | 9.2896 | 443.152 | 2E-29 | 1.4E-27 | 0.6571429 | n.a. |

| | | | | | | | |
|---|---|---|---|---|---|---|---|
| GLIPR1 | 2.601 | 9.887 | 126.874 | 2E-29 | 1.4E-27 | 0.8285714 | surface |
| SLA | 2.831 | 9.3615 | 166.793 | 3E-29 | 1.6E-27 | 0.8 | n.a. |
| ENSECAG00000000910 | 3.779 | 9.7673 | 129.427 | 9E-29 | 5.4E-27 | 0.4571429 | n.a. |
| ACTB | 1.512 | 12.301 | 122.234 | 2E-28 | 1.4E-26 | 1 | n.a. |
| LGALS3 | 2.546 | 9.5239 | 174.214 | 3E-28 | 1.9E-26 | 0.6 | n.a. |
| ENSECAG00000031962 | 3.332 | 9.2453 | 754.647 | 4E-28 | 2.2E-26 | 0.5428571 | n.a. |
| CDA | 2.918 | 9.2512 | 891.589 | 9E-28 | 5.2E-26 | 0.4857143 | n.a. |
| COTL1 | 1.861 | 10.529 | 118.788 | 1E-27 | 7.3E-26 | 1 | n.a. |
| ENSECAG00000006711 | 3.642 | 9.238 | 1432.9 | 1E-27 | 7.7E-26 | 0.5428571 | n.a. |
| LIMD2 | 2.663 | 10.143 | 116.308 | 4E-27 | 2.4E-25 | 0.8571429 | n.a. |
| SERPINB10 | 2.442 | 9.5345 | 577.11 | 7E-27 | 4E-25 | 0.4 | n.a. |
| CD300LB | 2.702 | 9.263 | 1019.48 | 5E-26 | 2.4E-24 | 0.4857143 | n.a. |
| ENSECAG00000031913 | 2.997 | 9.2438 | 1157.33 | 5E-26 | 2.6E-24 | 0.4571429 | n.a. |
| TUBA4A | 2.77 | 9.5479 | 109.563 | 2E-25 | 1.1E-23 | 0.6 | n.a. |
| GNG2 | 2.805 | 9.4802 | 121.676 | 4E-25 | 1.8E-23 | 0.6571429 | n.a. |
| ALOX5AP | 2.963 | 9.2637 | 680.502 | 8E-24 | 3.8E-22 | 0.4857143 | n.a. |
| ARG2 | 2.753 | 9.2454 | 1291.38 | 2E-23 | 1.1E-21 | 0.3428571 | n.a. |
| NPL | 2.617 | 9.2796 | 663.722 | 6E-23 | 2.8E-21 | 0.5142857 | n.a. |
| ENSECAG00000006595 | 2.725 | 9.2598 | 656.163 | 2E-22 | 7.6E-21 | 0.4857143 | n.a. |
| RAC2 | 2.081 | 10.594 | 93.1966 | 5E-22 | 2.4E-20 | 0.8285714 | n.a. |
| ISG20 | 3.296 | 9.3267 | 223.683 | 5E-22 | 2.5E-20 | 0.3428571 | n.a. |
| LITAF | 2.873 | 9.3245 | 253.671 | 1E-21 | 6.2E-20 | 0.5142857 | n.a. |
| ALPL | 2.786 | 9.2457 | 736.692 | 1E-21 | 6.6E-20 | 0.4 | surface |
| ALDOA | 1.806 | 9.99 | 91.105 | 1E-21 | 6.6E-20 | 0.9428571 | n.a. |
| RGS18 | 2.394 | 9.3114 | 241.421 | 2E-21 | 7.6E-20 | 0.5142857 | n.a. |
| GLIPR2 | 2.351 | 9.47 | 139.164 | 6E-21 | 2.6E-19 | 0.6285714 | n.a. |
| NUDT4 | 2.689 | 9.2991 | 216.249 | 1E-20 | 5.5E-19 | 0.5714286 | n.a. |
| UPP1 | 2.974 | 9.2386 | 1302.47 | 2E-20 | 6.5E-19 | 0.4 | n.a. |
| ENSECAG00000015500 | 2.59 | 9.2601 | 772.268 | 3E-20 | 1.3E-18 | 0.4 | n.a. |
| EMB | 3.249 | 9.3814 | 198.074 | 4E-20 | 1.5E-18 | 0.3428571 | surface |
| LTB4R | 2.837 | 9.2469 | 867.554 | 4E-20 | 1.5E-18 | 0.4571429 | surface |
| ENSECAG00000012199 | 2.365 | 9.3154 | 258.432 | 4E-20 | 1.8E-18 | 0.5714286 | n.a. |
| SLC16A3 | 2.648 | 9.3579 | 117.196 | 2E-19 | 9.7E-18 | 0.4 | n.a. |
| ENSECAG00000037706 | 1.813 | 10.368 | 80.0762 | 4E-19 | 1.5E-17 | 0.8571429 | n.a. |
| ENSECAG00000033857 | 1.932 | 9.3316 | 613.146 | 5E-19 | 2E-17 | 0.6571429 | n.a. |
| SAT1 | 1.654 | 9.7271 | 82.9553 | 2E-18 | 6.8E-17 | 0.8857143 | n.a. |
| TMEM229B | 2.741 | 9.2541 | 411.599 | 3E-18 | 1.1E-16 | 0.3428571 | n.a. |
| ARPC2 | 1.442 | 10.591 | 69.9628 | 6E-17 | 2.4E-15 | 0.9428571 | n.a. |
| FTL | 1.013 | 11.187 | 69.3687 | 8E-17 | 3.2E-15 | 0.9714286 | n.a. |
| CASP3 | 2.553 | 9.2592 | 367.322 | 1E-16 | 4.2E-15 | 0.2857143 | n.a. |
| ENSECAG00000032032 | 2.691 | 9.2656 | 372.676 | 1E-16 | 4.3E-15 | 0.3428571 | n.a. |
| ABTB1 | 2.48 | 9.3201 | 129.461 | 1E-16 | 5.1E-15 | 0.5714286 | n.a. |
| PGK1 | 2.24 | 9.4881 | 85.6123 | 2E-16 | 8.5E-15 | 0.6571429 | n.a. |
| NT5C3A | 2.672 | 9.3022 | 154.208 | 3E-16 | 1.1E-14 | 0.3428571 | n.a. |
| CCRL2 | 2.617 | 9.2557 | 486.078 | 4E-16 | 1.6E-14 | 0.2571429 | surface |
| EVI2B | 2.055 | 9.5277 | 65.7791 | 5E-16 | 1.9E-14 | 0.8 | surface |
| HCST | 2.261 | 9.5985 | 71.7341 | 1E-15 | 5E-14 | 0.5428571 | n.a. |
| ALOX5 | 2.519 | 9.2381 | 1676.31 | 4E-15 | 1.5E-13 | 0.2857143 | n.a. |
| ENSECAG00000017324 | 2.334 | 9.4228 | 76.4368 | 6E-15 | 2.1E-13 | 0.4 | n.a. |
| SELL | 2.018 | 9.6904 | 59.934 | 1E-14 | 3.5E-13 | 0.6285714 | surface |
| ENSECAG00000035315 | 3.286 | 9.2353 | 1566.31 | 1E-14 | 3.7E-13 | 0.3142857 | n.a. |
| FCAR | 2.713 | 9.2409 | 987.483 | 1E-14 | 4.3E-13 | 0.3428571 | surface |
| SUSD3 | 2.615 | 9.5214 | 64.7585 | 2E-14 | 5.7E-13 | 0.3714286 | surface |
| CORO1A | 1.874 | 10.247 | 58.9256 | 2E-14 | 5.7E-13 | 0.6857143 | n.a. |
| ARL6IP1 | 2.157 | 9.5522 | 58.6802 | 2E-14 | 6.4E-13 | 0.6285714 | n.a. |
| JUNB | 1.827 | 9.5395 | 58.1598 | 2E-14 | 8.3E-13 | 0.6571429 | n.a. |

| | | | | | | | |
|---|---|---|---|---|---|---|---|
| ENSECAG00000027699 | 1.715 | 9.9492 | 57.8198 | 3E-14 | 9.8E-13 | 0.9428571 | n.a. |
| ANXA11 | 2.163 | 9.4886 | 65.8404 | 5E-14 | 1.7E-12 | 0.6 | n.a. |
| ENSECAG00000028496 | 2.603 | 9.2366 | 1394.56 | 7E-14 | 2.4E-12 | 0.2857143 | n.a. |
| GPSM3 | 1.754 | 9.9781 | 55.2219 | 1E-13 | 3.6E-12 | 0.7142857 | n.a. |
| ENSECAG00000000436 | 1.426 | 9.9635 | 286.321 | 1E-13 | 4.2E-12 | 0.2571429 | n.a. |
| ENSECAG00000018618 | 1.98 | 9.2607 | 550.384 | 2E-13 | 7.9E-12 | 0.4 | n.a. |
| ILT11A | 1.714 | 9.2621 | 515.582 | 3E-13 | 1.1E-11 | 0.3714286 | n.a. |
| ENSECAG00000031851 | 2.444 | 9.2434 | 455.224 | 6E-13 | 1.7E-11 | 0.3714286 | n.a. |
| PFN1 | 1.104 | 11.941 | 51.6842 | 7E-13 | 2.1E-11 | 0.9714286 | n.a. |
| IGSF6 | 1.678 | 9.3651 | 235.633 | 1E-12 | 3.1E-11 | 0.6285714 | surface |
| ENSECAG00000016410 | 2.185 | 9.256 | 293.155 | 1E-12 | 4E-11 | 0.5428571 | n.a. |
| SLC40A1 | 2.486 | 9.2386 | 807.398 | 2E-12 | 6.5E-11 | 0.3142857 | surface |
| VSIR | 1.972 | 9.4326 | 78.4213 | 7E-12 | 2.1E-10 | 0.6285714 | n.a. |
| ARHGAP30 | 1.88 | 9.5616 | 46.836 | 1E-11 | 3E-10 | 0.5714286 | n.a. |
| FTH1 | 1.147 | 10.516 | 46.1134 | 1E-11 | 3.4E-10 | 0.9142857 | n.a. |
| BSG | 1.823 | 9.6757 | 44.3419 | 3E-11 | 8.2E-10 | 0.6 | surface |
| GRB2 | 1.675 | 9.7762 | 43.4489 | 4E-11 | 1.3E-09 | 0.6571429 | n.a. |
| ENSECAG00000007123 | 1.807 | 9.3975 | 54.292 | 6E-11 | 1.8E-09 | 0.5714286 | n.a. |
| FGR | 1.358 | 9.4883 | 88.6342 | 7E-11 | 2E-09 | 0.7714286 | n.a. |
| DDIT3 | 2.074 | 9.287 | 107.107 | 7E-11 | 2.1E-09 | 0.3142857 | n.a. |
| IFNGR2 | 1.82 | 9.3299 | 120.4 | 1E-10 | 3.7E-09 | 0.4857143 | surface |
| NFAM1 | 1.6 | 9.3582 | 185.798 | 1E-10 | 4.2E-09 | 0.5428571 | surface |
| SAMD9L | 1.88 | 9.4216 | 54.908 | 2E-10 | 6E-09 | 0.3428571 | n.a. |
| ENSECAG00000003925 | 1.505 | 9.5059 | 39.9986 | 3E-10 | 7.2E-09 | 0.5714286 | n.a. |
| NCF1 | 1.617 | 9.2933 | 188.036 | 3E-10 | 7.5E-09 | 0.4857143 | n.a. |
| LCP1 | 1.633 | 9.7068 | 39.3868 | 4E-10 | 9.7E-09 | 0.6857143 | n.a. |
| EZR | 1.765 | 9.8342 | 39.3033 | 4E-10 | 1E-08 | 0.5428571 | n.a. |
| DGAT1 | 2.149 | 9.29 | 115.019 | 4E-10 | 1.2E-08 | 0.3142857 | n.a. |
| CD82 | 2.355 | 9.4652 | 58.9566 | 4E-10 | 1.2E-08 | 0.3714286 | surface |
| MBOAT7 | 1.986 | 9.2726 | 174.401 | 5E-10 | 1.4E-08 | 0.4285714 | n.a. |
| YPEL3 | 1.789 | 9.6613 | 38.5514 | 5E-10 | 1.5E-08 | 0.6 | n.a. |
| GRINA | 1.772 | 9.3626 | 57.6079 | 6E-10 | 1.5E-08 | 0.5428571 | n.a. |
| TAGAP | 2.317 | 9.3906 | 71.3435 | 6E-10 | 1.5E-08 | 0.2571429 | n.a. |
| SNX3 | 1.367 | 9.6999 | 38.4347 | 6E-10 | 1.5E-08 | 0.8 | n.a. |
| NCF2 | 2.02 | 9.3332 | 105.397 | 8E-10 | 2E-08 | 0.4285714 | n.a. |
| AMDHD2 | 1.327 | 9.7855 | 37.6522 | 9E-10 | 2.2E-08 | 0.8571429 | n.a. |
| NFKBIA | 1.603 | 9.5431 | 37.182 | 1E-09 | 2.8E-08 | 0.6 | n.a. |
| STK17A | 1.855 | 9.6864 | 36.1699 | 2E-09 | 4.7E-08 | 0.5714286 | n.a. |
| TSC22D3 | 1.658 | 9.6043 | 34.2681 | 5E-09 | 1.2E-07 | 0.3714286 | n.a. |
| LAMTOR3 | 1.812 | 9.3658 | 55.5927 | 7E-09 | 1.7E-07 | 0.4 | n.a. |
| TNFAIP8 | 2.126 | 9.7152 | 33.4569 | 7E-09 | 1.8E-07 | 0.4285714 | n.a. |
| ENSECAG00000024719 | 1.924 | 9.2957 | 96.7354 | 7E-09 | 1.8E-07 | 0.3428571 | n.a. |
| ENSECAG00000020462 | 1.497 | 9.6751 | 35.7301 | 8E-09 | 1.8E-07 | 0.6571429 | n.a. |
| SPI1 | 1.119 | 9.6618 | 80.5658 | 1E-08 | 2.4E-07 | 0.8 | n.a. |
| PTPN6 | 1.737 | 9.58 | 36.8318 | 1E-08 | 2.6E-07 | 0.5714286 | n.a. |
| YPEL5 | 1.544 | 9.4914 | 32.5844 | 2E-08 | 5.5E-07 | 0.4285714 | n.a. |
| CSF3R | 1.823 | 9.2479 | 448.828 | 3E-08 | 6.7E-07 | 0.3428571 | surface |
| LY96 | 1.84 | 9.3807 | 54.7325 | 3E-08 | 7E-07 | 0.4285714 | n.a. |
| KCNJ15 | 1.928 | 9.2429 | 369.025 | 4E-08 | 8.2E-07 | 0.2571429 | n.a. |
| ENSECAG00000032598 | 2.051 | 9.2498 | 216.94 | 4E-08 | 8.5E-07 | 0.3714286 | n.a. |
| ZFP36 | 1.275 | 9.4955 | 34.4459 | 5E-08 | 1E-06 | 0.5714286 | n.a. |
| KLF6 | 1.447 | 9.5982 | 29.5874 | 5E-08 | 1.2E-06 | 0.5714286 | n.a. |
| RAB24 | 1.38 | 9.4289 | 38.2911 | 9E-08 | 1.9E-06 | 0.3714286 | n.a. |
| NAAA | 1.726 | 9.5818 | 30.6966 | 9E-08 | 2E-06 | 0.2857143 | n.a. |
| AGPAT2 | 1.648 | 9.4052 | 41.3872 | 1E-07 | 2.2E-06 | 0.4857143 | n.a. |
| PPP1R12A | 1.719 | 9.4355 | 32.927 | 1E-07 | 2.2E-06 | 0.4 | n.a. |

| | | | | | | | |
|---|---|---|---|---|---|---|---|
| MAFG | 1.84 | 9.2619 | 201.408 | 1E-07 | 2.3E-06 | 0.2571429 | n.a. |
| ENSECAG00000029287 | 1.897 | 10.043 | 32.1527 | 1E-07 | 2.3E-06 | 0.2571429 | n.a. |
| RGS19 | 1.559 | 9.5283 | 33.2344 | 1E-07 | 2.7E-06 | 0.5714286 | n.a. |
| CAP1 | 1.478 | 9.9501 | 27.3794 | 2E-07 | 3.5E-06 | 0.6 | n.a. |
| MYL12A | 1.093 | 10.506 | 27.2949 | 2E-07 | 3.7E-06 | 0.8 | n.a. |
| LSP1 | 1.106 | 10.243 | 27.0185 | 2E-07 | 4.2E-06 | 0.8 | n.a. |
| GNB2 | 1.29 | 9.9204 | 26.9858 | 2E-07 | 4.3E-06 | 0.7714286 | n.a. |
| TMEM154 | 1.965 | 9.3052 | 69.8287 | 2E-07 | 4.3E-06 | 0.2857143 | surface |
| ATG3 | 1.318 | 9.6124 | 30.2748 | 2E-07 | 4.6E-06 | 0.7142857 | n.a. |
| LTB | 2.695 | 10.681 | 26.6838 | 2E-07 | 4.9E-06 | 0.2571429 | n.a. |
| ID2 | 1.395 | 9.6451 | 35.7412 | 3E-07 | 5.4E-06 | 0.4 | n.a. |
| CXCR4 | 1.752 | 9.6177 | 29.9492 | 4E-07 | 8.5E-06 | 0.2571429 | surface |
| CAPZA1 | 1.614 | 9.7535 | 25.4038 | 5E-07 | 9.3E-06 | 0.5714286 | n.a. |
| ENSECAG00000034985 | 1.652 | 9.3342 | 46.0466 | 5E-07 | 9.9E-06 | 0.2857143 | n.a. |
| NFE2 | 1.292 | 9.2974 | 291.928 | 6E-07 | 1.1E-05 | 0.2857143 | n.a. |
| ENSECAG00000027864 | 1.368 | 9.2631 | 251.137 | 6E-07 | 1.2E-05 | 0.3142857 | n.a. |
| XG | 1.684 | 9.2905 | 81.5481 | 9E-07 | 1.8E-05 | 0.3142857 | n.a. |
| FAM107B | 1.941 | 9.5005 | 30.0537 | 9E-07 | 1.8E-05 | 0.3428571 | n.a. |
| IER3 | 1.723 | 9.3418 | 52.4078 | 9E-07 | 1.8E-05 | 0.2571429 | n.a. |
| REEP5 | 1.439 | 9.6201 | 25.5863 | 1E-06 | 1.9E-05 | 0.5714286 | n.a. |
| ACTG1 | 1.141 | 10.745 | 23.9713 | 1E-06 | 1.9E-05 | 0.6571429 | n.a. |
| NADK | 1.491 | 9.2968 | 70.0779 | 1E-06 | 2.1E-05 | 0.3428571 | n.a. |
| UBE2B | 1.435 | 9.8238 | 23.1951 | 1E-06 | 2.7E-05 | 0.6 | n.a. |
| PNPLA2 | 1.498 | 9.3533 | 55.5668 | 2E-06 | 4.1E-05 | 0.4571429 | n.a. |
| ENSECAG00000009892 | 1.692 | 9.3586 | 36.2604 | 3E-06 | 5.4E-05 | 0.2857143 | n.a. |
| CYSTM1 | 1.134 | 9.3828 | 48.7175 | 3E-06 | 5.5E-05 | 0.5714286 | n.a. |
| GSN | 1.408 | 9.3636 | 132.466 | 3E-06 | 5.6E-05 | 0.4571429 | n.a. |
| OAZ2 | 1.824 | 9.3485 | 40.4865 | 4E-06 | 7.1E-05 | 0.2857143 | n.a. |
| PLEK | 1.505 | 9.3075 | 69.2085 | 5E-06 | 9.2E-05 | 0.4571429 | n.a. |
| CDC42SE1 | 1.351 | 9.6142 | 20.6582 | 6E-06 | 9.6E-05 | 0.4571429 | n.a. |
| PTPRC | 1.079 | 10.647 | 20.3932 | 6E-06 | 0.00011 | 0.6285714 | surface |
| HCK | 1.383 | 9.2996 | 110.625 | 7E-06 | 0.00012 | 0.4 | n.a. |
| FBXL5 | 1.55 | 9.3155 | 43.8092 | 8E-06 | 0.00014 | 0.3142857 | n.a. |
| ICAM3 | 1.379 | 9.5641 | 20.4651 | 9E-06 | 0.00016 | 0.4285714 | surface |
| LRCH4 | 1.59 | 9.4087 | 26.1391 | 1E-05 | 0.00022 | 0.3142857 | n.a. |
| IRF2 | 1.376 | 9.4594 | 20.1213 | 1E-05 | 0.00023 | 0.3142857 | n.a. |
| TKT | 1.248 | 9.6069 | 41.0687 | 2E-05 | 0.00026 | 0.5428571 | n.a. |
| ITGB2 | 1.104 | 9.8101 | 18.6144 | 2E-05 | 0.00027 | 0.6285714 | surface |
| STK17B | 1.6 | 9.5111 | 20.7355 | 2E-05 | 0.00029 | 0.3428571 | n.a. |
| SAMD9 | 1.501 | 9.4177 | 25.2768 | 2E-05 | 0.00029 | 0.3714286 | n.a. |
| ARPC5 | 1.196 | 9.8016 | 18.4512 | 2E-05 | 0.00029 | 0.7142857 | n.a. |
| LBR | 1.563 | 9.6093 | 19.0753 | 2E-05 | 0.0003 | 0.3428571 | n.a. |
| LASP1 | 1.416 | 9.4194 | 26.2982 | 2E-05 | 0.0003 | 0.4 | n.a. |
| JPT1 | 1.155 | 9.7243 | 18.0793 | 2E-05 | 0.00037 | 0.5714286 | n.a. |
| NKIRAS2 | 1.688 | 9.3086 | 44.3834 | 2E-05 | 0.0004 | 0.2571429 | n.a. |
| MOB3A | 1.572 | 9.3512 | 29.802 | 3E-05 | 0.00042 | 0.3142857 | n.a. |
| DBNL | 1.275 | 9.4448 | 24.0935 | 4E-05 | 0.00059 | 0.4571429 | n.a. |
| PAIP2 | 1.407 | 9.5555 | 17.0862 | 7E-05 | 0.00109 | 0.4 | n.a. |
| IRF9 | 1.404 | 9.356 | 25.6369 | 9E-05 | 0.00126 | 0.2857143 | n.a. |
| ENSECAG00000039788 | 1.538 | 9.4822 | 28.1287 | 9E-05 | 0.00128 | 0.2571429 | n.a. |
| MAT2B | 1.127 | 9.6743 | 15.1103 | 0.0001 | 0.00147 | 0.5714286 | n.a. |
| ARRDC3 | 1.614 | 9.2691 | 53.8159 | 0.0001 | 0.00163 | 0.2571429 | n.a. |
| OSTF1 | 1.138 | 10.056 | 14.611 | 0.0001 | 0.00187 | 0.5142857 | n.a. |
| KIAA0513 | 1.305 | 9.2674 | 87.4276 | 0.0002 | 0.00221 | 0.2857143 | n.a. |
| RTN3 | 1.461 | 9.3937 | 24.907 | 0.0002 | 0.00221 | 0.3142857 | n.a. |
| RAB1A | 1.36 | 9.4321 | 20.1819 | 0.0002 | 0.0027 | 0.3714286 | n.a. |

| | | | | | | | |
|---|---|---|---|---|---|---|---|
| UBE2D2 | 1.177 | 9.6332 | 13.7382 | 0.0002 | 0.00287 | 0.5428571 | n.a. |
| RCSD1 | 1.264 | 9.6846 | 12.8655 | 0.0003 | 0.00435 | 0.4285714 | n.a. |
| CREG1 | 1.159 | 9.3229 | 148.26 | 0.0003 | 0.00451 | 0.2571429 | n.a. |
| SRI | 1.297 | 9.4117 | 18.8704 | 0.0004 | 0.0048 | 0.3714286 | n.a. |
| C5H1orf162 | 1.107 | 9.2673 | 73.9024 | 0.0005 | 0.00634 | 0.3142857 | n.a. |
| SELPLG | 1.043 | 9.4993 | 14.3495 | 0.0007 | 0.00835 | 0.4 | surface |
| ACTR3 | 1.011 | 9.9778 | 11.3859 | 0.0007 | 0.00889 | 0.6 | n.a. |
| ENSECAG00000009549 | 1.006 | 9.7659 | 11.1208 | 0.0009 | 0.0109 | 0.4571429 | n.a. |
| BZW1 | 1.101 | 9.5286 | 11.2182 | 0.001 | 0.0114 | 0.3714286 | n.a. |
| TNFRSF1A | 1.132 | 9.3254 | 27.534 | 0.001 | 0.01181 | 0.2857143 | surface |
| MICU2 | 1.175 | 9.4141 | 14.9225 | 0.0012 | 0.01414 | 0.3142857 | n.a. |
| PPP1R18 | 1.275 | 9.5073 | 12.251 | 0.0013 | 0.01501 | 0.2571429 | n.a. |
| WDR1 | 1.046 | 9.7464 | 12.6066 | 0.0014 | 0.01593 | 0.4571429 | n.a. |
| RIN3 | 1.205 | 9.3011 | 34.0053 | 0.0014 | 0.01602 | 0.2571429 | n.a. |
| IQGAP1 | 1.071 | 9.4976 | 13.0868 | 0.0015 | 0.01682 | 0.4 | n.a. |
| TOMM6 | 1.035 | 9.7098 | 9.67333 | 0.0019 | 0.0199 | 0.3714286 | n.a. |
| ENSECAG00000014975 | 1.235 | 9.3715 | 17.0162 | 0.0021 | 0.02189 | 0.2571429 | n.a. |
| VAPA | 1.064 | 9.5423 | 11.3274 | 0.0022 | 0.02328 | 0.4 | n.a. |
