## Supplementary material for "Single cell resolution landscape of equine peripheral blood mononuclear cells reveals diverse immune cell subtypes including T-bet^+^ B cells": Dataset S3

| genes | logFC | logCPM | F | PValue | FDR | percent.exp | Surfacome.Label |
|---|---|---|---|---|---|---|---|
| S100A12 | 2.93 | 10.309 | 3205.17 | 0 | 0 | 1 | n.a. |
| ENSECAG00000000436 | 2.006 | 9.9635 | 2561.29 | 0 | 0 | 0.998997 | n.a. |
| LYZ | 1.14 | 11.749 | 1320.38 | 9E-284 | 2E-280 | 1 | n.a. |
| SERPINB10 | 2.306 | 9.5345 | 1749.64 | 2E-264 | 5E-261 | 0.9809428 | n.a. |
| PLBD1 | 1.65 | 9.972 | 1172.51 | 1E-252 | 1E-249 | 1 | n.a. |
| ENSECAG00000010117 | 1.98 | 9.501 | 2355.88 | 2E-158 | 2E-155 | 0.5416249 | n.a. |
| C1H15orf48 | 1.758 | 9.6929 | 604.846 | 2E-132 | 1E-129 | 0.9889669 | n.a. |
| S100A8 | 1.537 | 9.3765 | 2287.13 | 2E-103 | 1E-100 | 0.3861585 | n.a. |
| VCAN | 1.583 | 9.3382 | 1117.92 | 5E-77 | 2E-74 | 0.8615848 | n.a. |
| THBS1 | 1.406 | 9.3512 | 908.453 | 7E-65 | 2E-62 | 0.7703109 | n.a. |
| TREM1 | 1.329 | 9.4852 | 1139.5 | 4E-61 | 9E-59 | 0.9578736 | surface |
| ENSECAG00000024882 | 1.436 | 9.349 | 830.187 | 2E-60 | 5E-58 | 0.781344 | n.a. |
| ENSECAG00000030387 | 1.265 | 9.433 | 250.791 | 3E-53 | 4E-51 | 0.8655968 | n.a. |
| ENSECAG00000024886 | 1.566 | 9.319 | 1204.78 | 1E-52 | 2E-50 | 0.8345035 | n.a. |
| S100A6 | 1.07 | 10.437 | 213.594 | 3E-48 | 4E-46 | 0.998997 | n.a. |
| SLPI | 1.097 | 9.3697 | 499.145 | 7E-48 | 8E-46 | 0.8796389 | n.a. |
| ENSECAG00000020136 | 1.308 | 9.7014 | 191.835 | 2E-43 | 2E-41 | 0.9558676 | n.a. |
| MGST1 | 1.155 | 9.3352 | 536.943 | 3E-43 | 3E-41 | 0.8214644 | nonsurface |
| ECATH-3 | 1.126 | 9.3365 | 1084.71 | 5E-39 | 4E-37 | 0.7873621 | n.a. |
| EMP1 | 1.357 | 9.2873 | 799.885 | 7E-38 | 6E-36 | 0.5827482 | surface |
| CD163 | 1.451 | 9.2595 | 834.934 | 4E-35 | 3E-33 | 0.446339 | surface |
| NFE2 | 1.077 | 9.2974 | 668 | 7E-30 | 4E-28 | 0.7021063 | n.a. |
| TNFSF13 | 1.026 | 9.266 | 480.592 | 2E-18 | 5E-17 | 0.4694082 | nonsurface |

| genes | logFC | logCPM | F | PValue | FDR | percent.exp | Surfacome.Label |
|---|---|---|---|---|---|---|---|
| ENSECAG00000030777 | 2.073 | 9.3773 | 1987.93 | 0 | 0 | 0.7799386 | n.a. |
| S100A10 | 1.09 | 10.866 | 418.66 | 2E-92 | 2E-89 | 0.9989765 | nonsurface |
| ENSECAG00000014585 | 1.046 | 10.271 | 366.629 | 3E-81 | 1E-78 | 0.7144319 | n.a. |
| ENSECAG00000038313 | 1.717 | 9.3718 | 650.987 | 5E-79 | 3E-76 | 0.5291709 | n.a. |
| MPEG1 | 1.059 | 9.3534 | 531.538 | 8E-71 | 3E-68 | 0.8485159 | surface |
| AHNAK | 1.461 | 9.5205 | 281.565 | 6E-63 | 2E-60 | 0.9037871 | n.a. |
| ABCA6 | 1.781 | 9.28 | 1658.71 | 5E-57 | 1E-54 | 0.5056295 | surface |
| ENSECAG00000040634 | 1.085 | 9.9556 | 248.507 | 9E-56 | 2E-53 | 0.9293756 | n.a. |
| ENSECAG00000027666 | 1.069 | 9.8957 | 216.671 | 7E-49 | 1E-46 | 0.9631525 | n.a. |
| THBS1 | 1.221 | 9.3512 | 714.764 | 8E-49 | 1E-46 | 0.6990788 | n.a. |
| CKB | 1.05 | 9.3208 | 670.304 | 4E-44 | 5E-42 | 0.3889458 | n.a. |
| EMP1 | 1.387 | 9.2873 | 822.94 | 2E-39 | 2E-37 | 0.5609007 | surface |
| ENSECAG00000028889 | 1.279 | 9.2871 | 562.173 | 5E-37 | 5E-35 | 0.3520983 | n.a. |
| ALDH1A1 | 1.35 | 9.2588 | 984.695 | 1E-28 | 7E-27 | 0.4145343 | n.a. |
| CD4 | 1.005 | 9.36 | 191.942 | 8E-26 | 5E-24 | 0.2896622 | surface |
| ENSECAG00000013303 | 1.047 | 9.4002 | 150.153 | 1E-24 | 6E-23 | 0.6622313 | n.a. |
| LRP1 | 1.093 | 9.2501 | 967.637 | 6E-22 | 3E-20 | 0.321392 | surface |
| ENSECAG00000024886 | 1.045 | 9.319 | 630.119 | 3E-21 | 1E-19 | 0.7144319 | n.a. |
| HNMT | 1.122 | 9.2852 | 525.011 | 2E-19 | 9E-18 | 0.6407369 | n.a. |
| ENSECAG00000035085 | 1.279 | 9.2904 | 389.409 | 5E-15 | 1E-13 | 0.3694985 | n.a. |

| genes | logFC | logCPM | F | PValue | FDR | percent.exp | Surfacome.Label |
|---|---|---|---|---|---|---|---|
| ENSECAG00000010117 | 5.135 | 9.501 | 7761.48 | 0 | 0 | 0.9274194 | n.a. |
| S100A8 | 4.627 | 9.3765 | 6670.28 | 0 | 0 | 0.7935484 | n.a. |
| S100A12 | 4.481 | 10.309 | 7037.45 | 0 | 0 | 0.9967742 | n.a. |
| ENSECAG00000036100 | 3.374 | 9.3021 | 4280.93 | 0 | 0 | 0.7693548 | n.a. |
| C1H15orf48 | 3.301 | 9.6929 | 2174.74 | 0 | 0 | 0.9774194 | n.a. |
| TREM1 | 2.879 | 9.4852 | 3943.42 | 0 | 0 | 0.9580645 | surface |
| MMP9 | 2.782 | 9.4158 | 1825.82 | 0 | 0 | 0.9274194 | n.a. |
| PLBD1 | 2.419 | 9.972 | 2307.94 | 0 | 0 | 0.9903226 | n.a. |
| ENSECAG00000000436 | 2.285 | 9.9635 | 3341.35 | 0 | 0 | 0.9903226 | n.a. |
| S100P | 1.892 | 9.8205 | 3019.66 | 0 | 0 | 0.9967742 | n.a. |
| LYZ | 1.681 | 11.749 | 2371.12 | 0 | 0 | 1 | n.a. |
| ENSECAG00000020136 | 3.488 | 9.7014 | 1417.42 | 5E-304 | 3E-301 | 0.9677419 | n.a. |
| IL18 | 3.329 | 9.3131 | 2103.89 | 4E-301 | 2E-298 | 0.8483871 | n.a. |
| ECATH-3 | 3.184 | 9.3365 | 4081.02 | 1E-274 | 8E-272 | 0.9032258 | n.a. |
| ENSECAG00000030745 | 1.265 | 11.502 | 1151.22 | 3E-248 | 1E-245 | 0.9983871 | n.a. |
| TSPO | 2.301 | 9.7323 | 1104.26 | 2E-238 | 1E-235 | 0.9854839 | nonsurface |
| S100A6 | 2.462 | 10.437 | 1023.08 | 3E-221 | 1E-218 | 0.9854839 | n.a. |
| SERPINB10 | 2.244 | 9.5345 | 1745.78 | 9E-200 | 4E-197 | 0.7193548 | n.a. |
| PRDX5 | 2.465 | 9.5264 | 850.646 | 9E-185 | 4E-182 | 0.9354839 | n.a. |
| ENSECAG00000006595 | 3.249 | 9.2598 | 2521.19 | 2E-160 | 7E-158 | 0.5451613 | n.a. |
| SRGN | 1.339 | 10.485 | 621.216 | 6E-136 | 2E-133 | 0.9983871 | n.a. |
| ENSECAG00000036967 | 2.559 | 9.2599 | 2562.91 | 2E-131 | 6E-129 | 0.5322581 | n.a. |
| ILT11B | 3.043 | 9.2698 | 2754.53 | 8E-125 | 2E-122 | 0.7225806 | n.a. |
| CD44 | 1.86 | 10.108 | 527.893 | 6E-116 | 2E-113 | 0.9548387 | surface |
| S100A5 | 1.985 | 9.5834 | 524.061 | 4E-115 | 1E-112 | 0.7064516 | n.a. |
| VIM | 1.141 | 11.286 | 512.574 | 1E-112 | 3E-110 | 0.9887097 | nonsurface |
| VCAN | 2.262 | 9.3382 | 1485.73 | 1E-110 | 4E-108 | 0.7935484 | n.a. |
| RAB24 | 1.919 | 9.4289 | 483.023 | 3E-106 | 6E-104 | 0.7032258 | n.a. |
| MAPK13 | 2.715 | 9.2495 | 2851.58 | 5E-103 | 1E-100 | 0.4225806 | n.a. |
| SLPI | 1.795 | 9.3697 | 1614.65 | 7E-100 | 1E-97 | 0.6919355 | n.a. |
| ARG2 | 2.238 | 9.2454 | 3328.17 | 3E-98 | 7E-96 | 0.2806452 | n.a. |
| IGSF6 | 1.454 | 9.3651 | 1005.19 | 9E-95 | 2E-92 | 0.8193548 | surface |
| ENSECAG00000010615 | 2.41 | 9.2458 | 1228.02 | 1E-92 | 2E-90 | 0.2870968 | n.a. |
| MGST1 | 1.836 | 9.3352 | 1145.78 | 9E-89 | 2E-86 | 0.6677419 | nonsurface |
| PADI4 | 2.379 | 9.2553 | 2374.56 | 8E-86 | 1E-83 | 0.4241935 | n.a. |
| ENSECAG00000030387 | 1.575 | 9.433 | 387.406 | 9E-86 | 2E-83 | 0.6629032 | n.a. |
| ALDOA | 1.289 | 9.99 | 385.954 | 2E-85 | 3E-83 | 0.95 | n.a. |
| NAPSA | 1.328 | 9.5255 | 489.745 | 6E-85 | 1E-82 | 0.7774194 | n.a. |
| LGALS3 | 1.678 | 9.5239 | 365.572 | 5E-81 | 7E-79 | 0.7725806 | n.a. |
| GLIPR1 | 1.797 | 9.887 | 355.478 | 7E-79 | 1E-76 | 0.7919355 | surface |
| ENSECAG00000030271 | 1.507 | 9.3599 | 438.316 | 3E-76 | 5E-74 | 0.7274194 | n.a. |
| ENSECAG00000037539 | 2.345 | 9.2665 | 1701.5 | 3E-75 | 4E-73 | 0.5677419 | n.a. |
| PLP2 | 1.506 | 9.9939 | 331.595 | 1E-73 | 1E-71 | 0.8806452 | n.a. |
| DUSP1 | 1.315 | 9.3756 | 325.138 | 2E-72 | 3E-70 | 0.6951613 | nonsurface |
| ALOX5AP | 2.362 | 9.2637 | 1572.7 | 1E-70 | 2E-68 | 0.2935484 | nonsurface |
| HMGB2 | 1.482 | 10.129 | 317.181 | 1E-70 | 2E-68 | 0.8306452 | n.a. |
| ITM2B | 1.092 | 10.174 | 313.556 | 7E-70 | 9E-68 | 0.9612903 | surface |

| | | | | | | | |
|---|---|---|---|---|---|---|---|
| CD14 | 1.47 | 9.3178 | 1349.76 | 9E-70 | 1E-67 | 0.95 | surface |
| CAPG | 1.517 | 9.6557 | 273.81 | 3E-61 | 3E-59 | 0.7854839 | n.a. |
| QPCT | 1.766 | 9.3093 | 745.614 | 1E-59 | 1E-57 | 0.6516129 | nonsurface |
| VNN2 | 2.212 | 9.2761 | 913.4 | 1E-59 | 1E-57 | 0.5870968 | surface |
| ENSECAG00000024563 | 1.957 | 9.2493 | 1204.27 | 2E-59 | 2E-57 | 0.283871 | n.a. |
| SELL | 1.733 | 9.6904 | 261.214 | 2E-58 | 2E-56 | 0.7532258 | surface |
| TUT7 | 1.33 | 9.6301 | 256.699 | 1E-57 | 1E-55 | 0.816129 | n.a. |
| ENSECAG00000003345 | 1.101 | 10.17 | 252.155 | 1E-56 | 1E-54 | 0.9225806 | n.a. |
| C5AR1 | 1.313 | 9.3446 | 636.128 | 2E-56 | 2E-54 | 0.8532258 | surface |
| ENSECAG00000016578 | 1.245 | 9.4164 | 998.991 | 2E-56 | 2E-54 | 0.7596774 | n.a. |
| PGD | 1.301 | 9.3424 | 377.255 | 5E-56 | 4E-54 | 0.6274194 | n.a. |
| TALDO1 | 1.129 | 9.9099 | 241.709 | 3E-54 | 2E-52 | 0.9467742 | n.a. |
| GLIPR2 | 1.161 | 9.47 | 241.295 | 3E-54 | 3E-52 | 0.7080645 | n.a. |
| RGS2 | 1.675 | 9.2835 | 913.566 | 9E-54 | 8E-52 | 0.4516129 | n.a. |
| RABAC1 | 1.24 | 9.8034 | 235.173 | 7E-53 | 5E-51 | 0.8967742 | nonsurface |
| ENSECAG00000024719 | 1.76 | 9.2957 | 447.184 | 7E-53 | 6E-51 | 0.366129 | n.a. |
| BNIP3L | 1.494 | 9.4595 | 231.141 | 5E-52 | 4E-50 | 0.6129032 | nonsurface |
| SOD2 | 1.626 | 9.3528 | 287.83 | 3E-51 | 2E-49 | 0.4516129 | n.a. |
| NFE2 | 1.787 | 9.2974 | 1217.48 | 7E-51 | 6E-49 | 0.5919355 | n.a. |
| HCST | 1.499 | 9.5985 | 236.225 | 2E-50 | 1E-48 | 0.35 | n.a. |
| NUDT4 | 1.718 | 9.2991 | 391.602 | 3E-50 | 2E-48 | 0.4645161 | n.a. |
| CDK2AP2 | 1.273 | 9.6423 | 209.56 | 2E-47 | 2E-45 | 0.6967742 | n.a. |
| NFAM1 | 1.235 | 9.3582 | 553.25 | 3E-46 | 2E-44 | 0.6451613 | surface |
| eca-mir-223 | 1.731 | 9.2778 | 954.817 | 3E-46 | 2E-44 | 0.4096774 | n.a. |
| VASP | 1.55 | 9.4309 | 241.445 | 3E-46 | 2E-44 | 0.3596774 | n.a. |
| GPSM3 | 1.089 | 9.9781 | 200.96 | 2E-45 | 1E-43 | 0.8403226 | n.a. |
| ENSECAG00000032032 | 1.729 | 9.2656 | 721.519 | 6E-45 | 5E-43 | 0.2548387 | n.a. |
| ID2 | 1.316 | 9.6451 | 195.765 | 2E-44 | 2E-42 | 0.5645161 | n.a. |
| BTG1 | 1.013 | 10.365 | 188.531 | 9E-43 | 6E-41 | 0.8241935 | n.a. |
| TAGLN2 | 1.012 | 10.164 | 182.132 | 2E-41 | 1E-39 | 0.8854839 | n.a. |
| PGK1 | 1.173 | 9.4881 | 180.531 | 5E-41 | 3E-39 | 0.6741935 | n.a. |
| ALDH3A1 | 1.96 | 9.2538 | 1185.27 | 6E-39 | 4E-37 | 0.4 | n.a. |
| CPNE3 | 1.393 | 9.3269 | 245.733 | 4E-38 | 2E-36 | 0.4193548 | n.a. |
| XBP1 | 1.186 | 9.4868 | 160.42 | 1E-36 | 6E-35 | 0.5887097 | n.a. |
| GTF2A2 | 1.215 | 9.5087 | 159.415 | 2E-36 | 1E-34 | 0.5096774 | n.a. |
| ICAM3 | 1.192 | 9.5641 | 159.179 | 2E-36 | 1E-34 | 0.6048387 | surface |
| CARHSP1 | 1.246 | 9.4061 | 169.576 | 5E-36 | 3E-34 | 0.4322581 | n.a. |
| SUSD3 | 1.516 | 9.5214 | 156.653 | 7E-36 | 4E-34 | 0.3129032 | surface |
| CERS4 | 1.568 | 9.2984 | 307.403 | 1E-35 | 8E-34 | 0.3612903 | nonsurface |
| IVNS1ABP | 1.361 | 9.3077 | 236.138 | 1E-33 | 5E-32 | 0.4193548 | n.a. |
| RHBDD2 | 1.212 | 9.3113 | 235.343 | 2E-33 | 1E-31 | 0.4177419 | nonsurface |
| NCF4 | 1.38 | 9.2924 | 523.511 | 4E-33 | 2E-31 | 0.3919355 | n.a. |
| ENSECAG00000024882 | 1.212 | 9.349 | 630.959 | 7E-33 | 3E-31 | 0.5064516 | n.a. |
| ADIPOR1 | 1.232 | 9.37 | 169.984 | 1E-32 | 5E-31 | 0.4354839 | nonsurface |
| CSF3R | 1.82 | 9.2479 | 1408.12 | 1E-32 | 6E-31 | 0.366129 | surface |
| EHD1 | 1.157 | 9.4259 | 142.693 | 2E-30 | 1E-28 | 0.3370968 | n.a. |
| FOS | 1.346 | 9.3156 | 450.819 | 3E-30 | 1E-28 | 0.283871 | n.a. |
| GYG1 | 1.035 | 9.5017 | 130.28 | 4E-30 | 2E-28 | 0.583871 | n.a. |

| | | | | | | | |
|---|---|---|---|---|---|---|---|
| ENSECAG00000031962 | 1.946 | 9.2453 | 719.327 | 1E-29 | 5E-28 | 0.2758065 | n.a. |
| RGS18 | 1.31 | 9.3114 | 325.342 | 2E-29 | 1E-27 | 0.3193548 | n.a. |
| LITAF | 1.264 | 9.3245 | 292.333 | 8E-29 | 3E-27 | 0.2564516 | n.a. |
| MXD1 | 1.159 | 9.3408 | 163.378 | 1E-28 | 5E-27 | 0.4048387 | n.a. |
| HK3 | 1.071 | 9.284 | 619.107 | 1E-28 | 6E-27 | 0.4290323 | n.a. |
| SYPL1 | 1.161 | 9.5663 | 122.238 | 2E-28 | 9E-27 | 0.4467742 | surface |
| YPEL5 | 1.054 | 9.4914 | 121.057 | 4E-28 | 2E-26 | 0.5967742 | n.a. |
| JAML | 1.418 | 9.299 | 468.625 | 1E-27 | 4E-26 | 0.3919355 | n.a. |
| SELP | 1.738 | 9.2569 | 920.586 | 1E-27 | 5E-26 | 0.4258065 | surface |
| GNG2 | 1.139 | 9.4802 | 115.936 | 5E-27 | 2E-25 | 0.3403226 | n.a. |
| YPEL3 | 1.096 | 9.6613 | 114.337 | 1E-26 | 5E-25 | 0.6032258 | n.a. |
| ENSECAG00000010008 | 1.027 | 9.4684 | 112.947 | 2E-26 | 9E-25 | 0.4483871 | n.a. |
| ACTN1 | 1.086 | 9.4643 | 112.916 | 2E-26 | 9E-25 | 0.2919355 | n.a. |
| CARD19 | 1.078 | 9.4452 | 115.471 | 3E-26 | 1E-24 | 0.4548387 | n.a. |
| LTB4R | 1.484 | 9.2469 | 919.742 | 2E-25 | 6E-24 | 0.2548387 | surface |
| ENSECAG00000024181 | 1.331 | 9.2586 | 864.55 | 3E-25 | 1E-23 | 0.2919355 | n.a. |
| RNF149 | 1.037 | 9.3444 | 157.556 | 1E-24 | 5E-23 | 0.5274194 | surface |
| RNF166 | 1.143 | 9.3001 | 205.514 | 2E-24 | 5E-23 | 0.3451613 | n.a. |
| DPYD | 1.202 | 9.2972 | 320.321 | 3E-24 | 1E-22 | 0.5419355 | n.a. |
| SNX10 | 1.065 | 9.274 | 374.596 | 5E-24 | 2E-22 | 0.3 | n.a. |
| CD300LB | 1.176 | 9.263 | 638.343 | 7E-23 | 2E-21 | 0.2870968 | n.a. |
| HIF1A | 1.025 | 9.3685 | 120.225 | 3E-21 | 1E-19 | 0.4048387 | n.a. |
| SLC16A3 | 1.034 | 9.3579 | 119.795 | 3E-21 | 1E-19 | 0.2693548 | n.a. |
| ENSECAG00000036115 | 1.033 | 9.2628 | 660.684 | 2E-20 | 5E-19 | 0.2629032 | n.a. |
| CLEC4E | 1.218 | 9.2548 | 582.362 | 1E-17 | 3E-16 | 0.2790323 | n.a. |
| THBS1 | 1.112 | 9.3512 | 380.195 | 2E-15 | 5E-14 | 0.5129032 | n.a. |
| ENSECAG00000015500 | 1.145 | 9.2601 | 457.657 | 4E-15 | 8E-14 | 0.316129 | n.a. |
| CDA | 1.106 | 9.2512 | 370.206 | 8E-15 | 2E-13 | 0.2919355 | nonsurface |

| genes | logFC | logCPM | F | PValue | FDR | percent.exp | Surfacome.Label |
|---|---|---|---|---|---|---|---|
| ENSECAG00000006663 | 5.995 | 9.3132 | 10724.3 | 0 | 0 | 0.987013 | n.a. |
| ENSECAG00000031322 | 5.746 | 10.499 | 4101.29 | 0 | 0 | 1 | n.a. |
| PLVAP | 4.711 | 9.247 | 10787.5 | 0 | 0 | 0.8441558 | surface |
| ENSECAG00000032710 | 4.704 | 9.3761 | 1942.73 | 0 | 0 | 0.7662338 | n.a. |
| GNGT2 | 4.381 | 9.2643 | 6091.49 | 1E-278 | 2E-275 | 0.8181818 | n.a. |
| ENSECAG00000032959 | 4.224 | 9.6944 | 1283.18 | 3E-275 | 2E-272 | 0.7792208 | n.a. |
| CD8A | 4.24 | 9.4281 | 1665.55 | 6E-257 | 5E-254 | 0.8701299 | surface |
| CX3CR1 | 4.108 | 9.2887 | 4879.32 | 2E-237 | 1E-234 | 0.7922078 | surface |
| ENSECAG00000019318 | 3.768 | 9.3455 | 2354.01 | 5E-235 | 3E-232 | 0.7142857 | n.a. |
| CELA1 | 4.223 | 9.2376 | 25498.1 | 1E-198 | 6E-196 | 0.6233766 | n.a. |
| POU2F2 | 3.488 | 9.5681 | 839.813 | 2E-168 | 1E-165 | 0.8961039 | n.a. |
| PTPRCAP | 3.981 | 10.617 | 766.92 | 6E-167 | 3E-164 | 0.7402597 | nonsurface |
| GBP5 | 3.807 | 9.7163 | 678.677 | 3E-148 | 1E-145 | 0.4415584 | nonsurface |
| NR4A1 | 3.515 | 9.2477 | 5931.24 | 6E-144 | 2E-141 | 0.5714286 | nonsurface |
| FCER1G | 1.299 | 10.204 | 649.24 | 7E-142 | 2E-139 | 1 | nonsurface |
| ABI3 | 3.61 | 9.3324 | 1411.85 | 8E-139 | 3E-136 | 0.7532468 | n.a. |
| PLAC8B | 2.561 | 11.113 | 603.115 | 5E-132 | 1E-129 | 0.974026 | n.a. |
| AIM2 | 3.193 | 9.3585 | 1578.81 | 2E-124 | 7E-122 | 0.8181818 | n.a. |
| PLA2G16 | 2.59 | 9.6144 | 560.936 | 5E-123 | 1E-120 | 1 | nonsurface |
| GPIHBP1 | 3.297 | 9.2751 | 6390.04 | 2E-117 | 6E-115 | 0.5844156 | surface |
| CYSTM1 | 2.713 | 9.3828 | 782.792 | 7E-114 | 2E-111 | 0.8571429 | n.a. |
| ARHGDIB | 1.808 | 11.255 | 493.763 | 1E-108 | 3E-106 | 0.987013 | n.a. |
| PRNP | 2.279 | 9.3261 | 620.598 | 1E-102 | 3E-100 | 0.8311688 | nonsurface |
| TCF7L2 | 3.055 | 9.2572 | 2480.54 | 3E-102 | 6E-100 | 0.6493506 | n.a. |
| ICAM2 | 3.439 | 9.4874 | 622.598 | 5E-99 | 1E-96 | 0.4935065 | surface |
| IFI30 | 2.129 | 9.9429 | 437.033 | 2E-96 | 4E-94 | 0.961039 | n.a. |
| CD79A | 3.292 | 9.7872 | 1152.39 | 1E-92 | 2E-90 | 0.4025974 | surface |
| H2AFZ | 2.569 | 9.5191 | 375.212 | 4E-83 | 7E-81 | 0.7532468 | n.a. |
| ALDH2 | 1.606 | 9.5616 | 403.524 | 7E-83 | 1E-80 | 0.9350649 | n.a. |
| ENSECAG00000019052 | 3.133 | 9.4644 | 463.918 | 5E-79 | 9E-77 | 0.5194805 | n.a. |
| ENSECAG00000031455 | 2.921 | 9.2358 | 5991.75 | 2E-78 | 3E-76 | 0.3766234 | n.a. |
| ENSECAG00000040478 | 2.857 | 9.248 | 3089.37 | 2E-76 | 4E-74 | 0.6233766 | n.a. |
| ENSECAG00000038143 | 2.958 | 9.2349 | 7009.1 | 6E-75 | 9E-73 | 0.4155844 | n.a. |
| MNDA | 1.878 | 10.258 | 335.45 | 1E-74 | 2E-72 | 1 | n.a. |
| BIN1 | 3.071 | 9.382 | 659.527 | 4E-74 | 7E-72 | 0.5064935 | n.a. |
| DUSP5 | 2.797 | 9.267 | 1785.47 | 1E-73 | 2E-71 | 0.3896104 | n.a. |
| ST3GAL6 | 2.799 | 9.5094 | 331.03 | 1E-73 | 2E-71 | 0.7532468 | nonsurface |
| ATP1B3 | 1.991 | 9.5391 | 329.659 | 2E-73 | 4E-71 | 0.9480519 | surface |
| CD300H | 2.839 | 9.2467 | 2787.87 | 3E-73 | 5E-71 | 0.5194805 | n.a. |
| P2RY14 | 2.743 | 9.2475 | 2227.08 | 3E-70 | 5E-68 | 0.4545455 | surface |
| HMOX1 | 1.943 | 9.3479 | 337.082 | 4E-68 | 6E-66 | 0.8701299 | nonsurface |
| HES4 | 2.761 | 9.2428 | 3542.74 | 4E-67 | 5E-65 | 0.4415584 | n.a. |
| DCSTAMP | 2.6 | 9.2378 | 5119.68 | 5E-64 | 7E-62 | 0.3636364 | surface |
| GBP6 | 2.909 | 9.5114 | 326.75 | 3E-62 | 4E-60 | 0.3246753 | nonsurface |
| CSF1R | 1.893 | 9.3336 | 1045.98 | 3E-62 | 4E-60 | 0.974026 | surface |
| EVL | 2.759 | 9.7253 | 274.458 | 2E-61 | 3E-59 | 0.6103896 | n.a. |
| eca-mir-1892 | 2.455 | 9.4799 | 271.767 | 8E-61 | 1E-58 | 0.7532468 | n.a. |

| | | | | | | | |
|---|---|---|---|---|---|---|---|
| CD53 | 2.252 | 9.709 | 268.671 | 4E-60 | 5E-58 | 0.9350649 | surface |
| ENSECAG00000019100 | 2.649 | 9.2458 | 3174.76 | 7E-59 | 8E-57 | 0.5714286 | n.a. |
| WARS | 2.478 | 9.8298 | 259.178 | 4E-58 | 5E-56 | 0.8051948 | n.a. |
| SLC44A2 | 2.791 | 9.4299 | 326.215 | 7E-57 | 8E-55 | 0.4935065 | surface |
| CD37 | 2.038 | 9.891 | 250.305 | 4E-56 | 4E-54 | 0.961039 | surface |
| TIE1 | 2.6 | 9.2364 | 4911.63 | 3E-54 | 4E-52 | 0.3766234 | surface |
| TPK1 | 2.566 | 9.2622 | 861.446 | 8E-53 | 8E-51 | 0.4155844 | n.a. |
| FABP5 | 2.747 | 9.5264 | 276.255 | 9E-53 | 1E-50 | 0.5064935 | n.a. |
| RHOF | 2.591 | 9.4402 | 277.706 | 8E-49 | 8E-47 | 0.4155844 | n.a. |
| CORO1B | 1.755 | 9.4003 | 219.443 | 6E-48 | 6E-46 | 0.7922078 | n.a. |
| GPR31 | 2.255 | 9.2389 | 1880.54 | 6E-47 | 6E-45 | 0.2727273 | surface |
| PLD4 | 2.184 | 9.3359 | 558.373 | 7E-47 | 7E-45 | 0.7532468 | n.a. |
| MYO1G | 2.2 | 9.6134 | 204.858 | 2E-46 | 2E-44 | 0.8311688 | n.a. |
| LST1 | 1.769 | 9.329 | 248.701 | 7E-46 | 6E-44 | 0.7792208 | nonsurface |
| CEBPB | 1.382 | 9.407 | 200.836 | 2E-45 | 2E-43 | 0.9480519 | n.a. |
| CD68 | 1.646 | 9.3465 | 543.562 | 3E-44 | 2E-42 | 0.8441558 | surface |
| ENSECAG00000015010 | 1.924 | 9.2872 | 826.307 | 1E-43 | 1E-41 | 0.6883117 | n.a. |
| GUCY1B1 | 2.377 | 9.2412 | 1795.12 | 2E-43 | 1E-41 | 0.3246753 | n.a. |
| TXNIP | 1.639 | 10.204 | 188.197 | 1E-42 | 8E-41 | 0.974026 | n.a. |
| ISG15 | 2.433 | 9.3193 | 316.42 | 1E-41 | 1E-39 | 0.4805195 | n.a. |
| ENSECAG00000009285 | 2.319 | 9.2377 | 1830.36 | 1E-40 | 1E-38 | 0.3636364 | n.a. |
| FTH1 | 1.395 | 10.516 | 177.269 | 2E-40 | 2E-38 | 0.9480519 | nonsurface |
| SLAMF7 | 2.395 | 9.2845 | 687.183 | 5E-40 | 3E-38 | 0.4025974 | surface |
| AK3 | 2.189 | 9.4014 | 211.122 | 8E-40 | 6E-38 | 0.6883117 | n.a. |
| UBC | 1.208 | 11.249 | 173.789 | 1E-39 | 1E-37 | 0.9350649 | n.a. |
| TBC1D8 | 2.223 | 9.2684 | 443.013 | 5E-39 | 3E-37 | 0.5454545 | n.a. |
| ENSECAG00000016543 | 2.202 | 9.8125 | 168.521 | 2E-38 | 1E-36 | 0.7272727 | n.a. |
| NAAA | 2.299 | 9.5818 | 166.052 | 7E-38 | 5E-36 | 0.5974026 | n.a. |
| DUSP12 | 2.398 | 9.3489 | 261.845 | 7E-38 | 5E-36 | 0.3896104 | n.a. |
| UGCG | 2.307 | 9.3115 | 331.022 | 9E-38 | 6E-36 | 0.3506494 | nonsurface |
| RF00163 | 2.37 | 9.4218 | 217.323 | 1E-37 | 7E-36 | 0.4415584 | n.a. |
| SDC3 | 2.155 | 9.2391 | 3238.18 | 1E-37 | 8E-36 | 0.2597403 | nonsurface |
| HCLS1 | 1.87 | 9.4497 | 154.19 | 3E-35 | 2E-33 | 0.6883117 | n.a. |
| MAP7D3 | 2.039 | 9.41 | 165.482 | 1E-34 | 6E-33 | 0.5584416 | n.a. |
| ENSECAG00000036967 | 1.674 | 9.2599 | 1158.1 | 8E-34 | 5E-32 | 0.3506494 | n.a. |
| ENSECAG00000004433 | 2.09 | 9.2481 | 1024.71 | 1E-33 | 7E-32 | 0.3506494 | n.a. |
| CYTIP | 1.798 | 9.6111 | 146.291 | 1E-33 | 8E-32 | 0.7922078 | n.a. |
| SAMSN1 | 1.984 | 9.3692 | 181.314 | 8E-33 | 5E-31 | 0.5194805 | n.a. |
| CSK | 2.013 | 9.389 | 171.167 | 2E-32 | 1E-30 | 0.6883117 | n.a. |
| LYN | 1.684 | 9.3216 | 257.572 | 5E-32 | 3E-30 | 0.6623377 | surface |
| C4BPA | 1.01 | 9.4013 | 202.426 | 2E-31 | 9E-30 | 0.8961039 | n.a. |
| NAP1L1 | 1.552 | 10.092 | 134.617 | 5E-31 | 3E-29 | 0.8701299 | n.a. |
| CYP4F22 | 2.078 | 9.2378 | 2293.61 | 2E-30 | 1E-28 | 0.3116883 | nonsurface |
| TUBA1A | 2.056 | 10.091 | 130.877 | 3E-30 | 2E-28 | 0.5844156 | n.a. |
| TBC1D9 | 2.08 | 9.2457 | 1313.02 | 2E-29 | 1E-27 | 0.3636364 | n.a. |
| PTPRC | 1.341 | 10.647 | 126.409 | 3E-29 | 1E-27 | 0.8961039 | surface |
| C20H6orf62 | 1.802 | 9.3936 | 152.503 | 3E-29 | 2E-27 | 0.6363636 | n.a. |
| ENSECAG00000033857 | 1.302 | 9.3316 | 423.637 | 3E-29 | 2E-27 | 0.8571429 | n.a. |

| | | | | | | | |
|---|---|---|---|---|---|---|---|
| ZYX | 1.483 | 9.4366 | 122.906 | 2E-28 | 8E-27 | 0.8311688 | n.a. |
| PLEKHO2 | 1.939 | 9.3041 | 229.115 | 2E-28 | 8E-27 | 0.5454545 | n.a. |
| ENSECAG00000021110 | 1.75 | 9.2654 | 765.175 | 3E-28 | 1E-26 | 0.5974026 | n.a. |
| SIVA1 | 1.746 | 9.5677 | 120.732 | 5E-28 | 2E-26 | 0.7532468 | n.a. |
| ENSECAG00000040634 | 1.606 | 9.9556 | 119.28 | 1E-27 | 5E-26 | 0.7402597 | n.a. |
| ACP5 | 2.026 | 9.7992 | 118.769 | 1E-27 | 6E-26 | 0.5454545 | n.a. |
| ENSECAG00000036034 | 1.389 | 9.8269 | 112.503 | 3E-26 | 1E-24 | 0.9220779 | n.a. |
| PKIB | 1.829 | 9.2759 | 915.762 | 5E-26 | 2E-24 | 0.2857143 | n.a. |
| UBD | 1.9 | 9.9806 | 109.036 | 2E-25 | 8E-24 | 0.6493506 | n.a. |
| TMC6 | 2.044 | 9.4027 | 148.567 | 2E-25 | 1E-23 | 0.3246753 | nonsurface |
| CYTH4 | 1.893 | 9.2629 | 390.221 | 3E-25 | 1E-23 | 0.4805195 | n.a. |
| ETS2 | 2.098 | 9.2476 | 741.925 | 3E-25 | 2E-23 | 0.4675325 | n.a. |
| ENSECAG00000030777 | 1.302 | 9.3773 | 457.444 | 4E-25 | 2E-23 | 0.6103896 | n.a. |
| ENSECAG00000033029 | 1.889 | 9.2619 | 341.695 | 5E-25 | 2E-23 | 0.4935065 | n.a. |
| CORO1A | 1.345 | 10.247 | 105.648 | 1E-24 | 4E-23 | 0.8311688 | n.a. |
| MRPS6 | 1.717 | 9.4746 | 108.121 | 1E-24 | 6E-23 | 0.6493506 | n.a. |
| LRRFIP1 | 1.439 | 9.8063 | 104.831 | 1E-24 | 6E-23 | 0.9090909 | n.a. |
| FLNA | 1.397 | 9.4126 | 103.801 | 2E-24 | 1E-22 | 0.7532468 | n.a. |
| ENSECAG00000031569 | 1.166 | 11.194 | 103.075 | 3E-24 | 2E-22 | 0.961039 | n.a. |
| DIPK2A | 1.728 | 9.2586 | 638.127 | 7E-24 | 3E-22 | 0.3246753 | n.a. |
| MS4A7 | 1.578 | 9.3128 | 473.866 | 9E-24 | 4E-22 | 0.5714286 | n.a. |
| SELPLG | 1.569 | 9.4993 | 96.7245 | 9E-23 | 3E-21 | 0.6363636 | surface |
| ATF3 | 1.878 | 9.2384 | 1508.3 | 9E-23 | 4E-21 | 0.2727273 | n.a. |
| WIPF1 | 1.601 | 9.5089 | 95.171 | 2E-22 | 7E-21 | 0.5844156 | n.a. |
| CTNNAL1 | 2.018 | 9.2496 | 587.63 | 3E-22 | 1E-20 | 0.4155844 | nonsurface |
| ENSECAG00000032131 | 1.747 | 9.3494 | 136.936 | 3E-22 | 1E-20 | 0.4935065 | n.a. |
| DQB | 1.234 | 10.011 | 93.3265 | 5E-22 | 2E-20 | 0.8181818 | n.a. |
| DBI | 1.262 | 9.673 | 92.8525 | 6E-22 | 2E-20 | 0.8701299 | n.a. |
| RAP1B | 1.363 | 9.5901 | 91.5811 | 1E-21 | 4E-20 | 0.8831169 | n.a. |
| ABCG2 | 1.63 | 9.262 | 382.718 | 2E-21 | 7E-20 | 0.3506494 | surface |
| RGS10 | 1.464 | 9.5903 | 90.4905 | 2E-21 | 7E-20 | 0.5584416 | n.a. |
| UCP2 | 1.039 | 9.9052 | 90.2298 | 2E-21 | 8E-20 | 0.961039 | nonsurface |
| RAC2 | 1.196 | 10.594 | 89.6999 | 3E-21 | 1E-19 | 0.9350649 | n.a. |
| Eqca-DQB1 | 1.012 | 10.925 | 89.3312 | 4E-21 | 1E-19 | 0.974026 | n.a. |
| BIN2 | 1.358 | 9.7163 | 89.0706 | 4E-21 | 1E-19 | 0.8961039 | n.a. |
| ITM2C | 1.566 | 9.5994 | 88.452 | 6E-21 | 2E-19 | 0.4935065 | surface |
| TNFSF10 | 1.455 | 9.4268 | 88.1618 | 6E-21 | 2E-19 | 0.7792208 | n.a. |
| ENSECAG00000012830 | 1.744 | 9.263 | 389.461 | 3E-20 | 9E-19 | 0.2987013 | n.a. |
| MYL12A | 1.042 | 10.506 | 85.0498 | 3E-20 | 1E-18 | 0.9480519 | n.a. |
| SAMD9L | 1.49 | 9.4216 | 84.8498 | 3E-20 | 1E-18 | 0.6753247 | n.a. |
| ADGRE5 | 1.354 | 9.5791 | 84 | 5E-20 | 2E-18 | 0.7662338 | n.a. |
| KIF22 | 1.765 | 9.2545 | 490.187 | 1E-19 | 3E-18 | 0.2857143 | n.a. |
| SEPT9 | 1.511 | 9.7694 | 81.9511 | 1E-19 | 5E-18 | 0.6753247 | n.a. |
| INPP5F | 1.854 | 9.2549 | 454.957 | 2E-19 | 7E-18 | 0.2727273 | n.a. |
| SLC7A7 | 1.275 | 9.4046 | 80.5196 | 3E-19 | 1E-17 | 0.7662338 | n.a. |
| NMI | 1.3 | 9.4149 | 85.93 | 4E-19 | 1E-17 | 0.8181818 | n.a. |
| ST6GALNAC2 | 1.531 | 9.2783 | 166.392 | 2E-18 | 6E-17 | 0.4805195 | nonsurface |
| DBNL | 1.43 | 9.4448 | 91.4394 | 2E-18 | 6E-17 | 0.7532468 | n.a. |

| | | | | | | | |
|---|---|---|---|---|---|---|---|
| RF00411 | 1.531 | 9.4519 | 78.3625 | 2E-18 | 6E-17 | 0.4545455 | n.a. |
| ENSECAG00000009162 | 1.04 | 10.473 | 76.8112 | 2E-18 | 6E-17 | 0.9350649 | n.a. |
| GNA15 | 1.735 | 9.3122 | 150.38 | 2E-18 | 6E-17 | 0.4285714 | n.a. |
| CYTH1 | 1.67 | 9.3765 | 100.63 | 2E-18 | 7E-17 | 0.3896104 | n.a. |
| TIFA | 1.639 | 10.029 | 75.0848 | 5E-18 | 1E-16 | 0.6363636 | n.a. |
| LIMD2 | 1.444 | 10.143 | 74.321 | 7E-18 | 2E-16 | 0.7402597 | n.a. |
| PAM | 1.76 | 9.2453 | 603.17 | 8E-18 | 2E-16 | 0.2727273 | surface |
| CALM2 | 1.101 | 10.275 | 73.6515 | 1E-17 | 3E-16 | 0.8701299 | n.a. |
| HYPK | 1.562 | 9.4628 | 80.5367 | 2E-17 | 4E-16 | 0.5974026 | n.a. |
| JUP | 1.685 | 9.251 | 555.711 | 4E-17 | 1E-15 | 0.2597403 | n.a. |
| ARL6IP1 | 1.487 | 9.5522 | 70.7523 | 4E-17 | 1E-15 | 0.6623377 | nonsurface |
| HPRT1 | 1.31 | 9.5077 | 70.3713 | 5E-17 | 1E-15 | 0.5844156 | n.a. |
| ITGB1 | 1.573 | 9.5309 | 69.9371 | 6E-17 | 2E-15 | 0.3896104 | surface |
| PRDX1 | 1.32 | 9.8857 | 69.8378 | 7E-17 | 2E-15 | 0.7272727 | n.a. |
| TUBA4A | 1.467 | 9.5479 | 69.8119 | 7E-17 | 2E-15 | 0.4805195 | n.a. |
| NEDD9 | 1.68 | 9.2911 | 192.464 | 1E-16 | 4E-15 | 0.2987013 | n.a. |
| KLF2 | 1.467 | 9.4082 | 72.454 | 1E-16 | 4E-15 | 0.4935065 | n.a. |
| NRN1 | 1.505 | 9.2707 | 290.815 | 2E-16 | 4E-15 | 0.3896104 | surface |
| SQSTM1 | 1.481 | 9.456 | 68.1783 | 2E-16 | 4E-15 | 0.5064935 | n.a. |
| MSN | 1.162 | 9.6123 | 67.8971 | 2E-16 | 5E-15 | 0.8571429 | n.a. |
| ITGA4 | 1.409 | 9.465 | 72.7482 | 2E-16 | 5E-15 | 0.5194805 | surface |
| APP | 1.497 | 9.2487 | 397.785 | 5E-16 | 1E-14 | 0.4155844 | surface |
| MEF2A | 1.568 | 9.302 | 142.959 | 5E-16 | 1E-14 | 0.2987013 | n.a. |
| SH2D3C | 1.571 | 9.2916 | 152.119 | 5E-16 | 1E-14 | 0.2727273 | n.a. |
| RPS6KA1 | 1.481 | 9.4019 | 86.0343 | 6E-16 | 2E-14 | 0.5714286 | n.a. |
| PTP4A2 | 1.304 | 9.7459 | 63.995 | 1E-15 | 3E-14 | 0.7402597 | n.a. |
| RASA3 | 1.513 | 9.5751 | 63.8042 | 1E-15 | 4E-14 | 0.5064935 | n.a. |
| IFIT3 | 1.657 | 9.2748 | 210.74 | 2E-15 | 5E-14 | 0.2987013 | n.a. |
| YWHAH | 1.339 | 9.5952 | 62.87 | 2E-15 | 6E-14 | 0.5844156 | n.a. |
| ACTR3 | 1.196 | 9.9778 | 62.3427 | 3E-15 | 7E-14 | 0.8051948 | n.a. |
| ILT11A | 1.406 | 9.2621 | 364.234 | 4E-15 | 8E-14 | 0.5324675 | n.a. |
| RF01956 | 1.391 | 9.2926 | 128.438 | 4E-15 | 9E-14 | 0.7532468 | n.a. |
| FCGRT | 1.378 | 9.3372 | 145.973 | 4E-15 | 1E-13 | 0.4545455 | surface |
| CXCL16 | 1.469 | 9.3053 | 118.287 | 4E-15 | 1E-13 | 0.3766234 | surface |
| ITGAL | 1.401 | 9.4047 | 78.4709 | 5E-15 | 1E-13 | 0.4545455 | surface |
| PTGER4 | 1.589 | 9.3476 | 108.147 | 5E-15 | 1E-13 | 0.2727273 | surface |
| ENSECAG00000040532 | 1.507 | 9.3029 | 127.879 | 7E-15 | 2E-13 | 0.4025974 | n.a. |
| CAP1 | 1.205 | 9.9501 | 60.4588 | 8E-15 | 2E-13 | 0.8701299 | n.a. |
| ENSECAG00000037332 | 1.164 | 10.054 | 60.0829 | 9E-15 | 2E-13 | 0.8181818 | n.a. |
| DHRS3 | 1.282 | 9.3111 | 122.217 | 1E-14 | 3E-13 | 0.6103896 | nonsurface |
| CHD9 | 1.255 | 9.4262 | 62.6179 | 4E-14 | 8E-13 | 0.5974026 | n.a. |
| KLF4 | 1.432 | 9.2595 | 414.294 | 4E-14 | 1E-12 | 0.3506494 | n.a. |
| PNRC1 | 1.18 | 9.9575 | 56.3668 | 6E-14 | 1E-12 | 0.8181818 | n.a. |
| LUZP6 | 1.268 | 9.4318 | 60.3365 | 1E-13 | 2E-12 | 0.5194805 | n.a. |
| TTC9C | 1.442 | 9.3445 | 76.4089 | 1E-13 | 3E-12 | 0.3376623 | n.a. |
| FYN | 1.455 | 9.4235 | 75.2579 | 2E-13 | 4E-12 | 0.3246753 | n.a. |
| PTBP3 | 1.492 | 9.4001 | 74.9678 | 2E-13 | 5E-12 | 0.5194805 | n.a. |
| TMEM14C | 1.101 | 9.6674 | 53.423 | 3E-13 | 6E-12 | 0.7792208 | nonsurface |

| | | | | | | | |
|---|---|---|---|---|---|---|---|
| CHMP4B | 1.197 | 9.4518 | 53.4445 | 3E-13 | 6E-12 | 0.6623377 | n.a. |
| DDX5 | 1.061 | 10.25 | 52.5219 | 4E-13 | 9E-12 | 0.7922078 | n.a. |
| ENSECAG00000003925 | 1.105 | 9.5059 | 52.1188 | 5E-13 | 1E-11 | 0.6753247 | n.a. |
| ENSECAG00000009794 | 1.297 | 9.3398 | 70.7175 | 9E-13 | 2E-11 | 0.4935065 | n.a. |
| CCDC12 | 1.408 | 9.3727 | 68.9793 | 2E-12 | 4E-11 | 0.4155844 | nonsurface |
| CMTM3 | 1.342 | 9.4968 | 56.0877 | 2E-12 | 4E-11 | 0.3376623 | nonsurface |
| REL | 1.407 | 9.3166 | 90.7477 | 3E-12 | 6E-11 | 0.2987013 | n.a. |
| LAP3 | 1.175 | 9.4403 | 58.0567 | 3E-12 | 7E-11 | 0.5974026 | n.a. |
| ARHGAP15 | 1.381 | 9.6767 | 48.3727 | 4E-12 | 7E-11 | 0.4935065 | n.a. |
| NDUFS5 | 1.186 | 9.6423 | 48.2309 | 4E-12 | 8E-11 | 0.7402597 | n.a. |
| UPF2 | 1.45 | 9.339 | 82.0678 | 4E-12 | 8E-11 | 0.3376623 | n.a. |
| ENSECAG00000034339 | 1.066 | 10.188 | 47.5897 | 5E-12 | 1E-10 | 0.7402597 | n.a. |
| MAGOH | 1.376 | 9.4803 | 53.1727 | 6E-12 | 1E-10 | 0.5194805 | n.a. |
| NR1H3 | 1.403 | 9.244 | 628.465 | 7E-12 | 1E-10 | 0.2597403 | n.a. |
| MRPL35 | 1.512 | 9.3167 | 95.3594 | 7E-12 | 1E-10 | 0.3636364 | n.a. |
| ENSECAG00000028581 | 1.453 | 9.2649 | 171.524 | 7E-12 | 1E-10 | 0.4285714 | n.a. |
| STARD7 | 1.369 | 9.3039 | 101.97 | 8E-12 | 2E-10 | 0.4415584 | n.a. |
| ENSECAG00000000775 | 1.012 | 9.4621 | 58.1766 | 8E-12 | 2E-10 | 0.6623377 | n.a. |
| TK2 | 1.459 | 9.3075 | 111.453 | 9E-12 | 2E-10 | 0.2597403 | n.a. |
| SNAP23 | 1.432 | 9.3138 | 93.8423 | 1E-11 | 2E-10 | 0.3376623 | n.a. |
| PAQR4 | 1.395 | 9.2942 | 116.794 | 1E-11 | 3E-10 | 0.2597403 | nonsurface |
| NET1 | 1.365 | 9.27 | 183.811 | 2E-11 | 3E-10 | 0.3766234 | n.a. |
| MEF2C | 1.158 | 9.4205 | 127.046 | 2E-11 | 5E-10 | 0.4935065 | n.a. |
| DRAM2 | 1.236 | 9.402 | 52.1182 | 2E-11 | 5E-10 | 0.4675325 | nonsurface |
| SYNGR2 | 1.374 | 9.3319 | 120.287 | 3E-11 | 6E-10 | 0.4155844 | n.a. |
| STK10 | 1.101 | 9.392 | 48.3534 | 5E-11 | 9E-10 | 0.5974026 | n.a. |
| NDUFA12 | 1.189 | 9.4672 | 46.7599 | 6E-11 | 1E-09 | 0.5064935 | n.a. |
| ACSL4 | 1.348 | 9.2833 | 123.079 | 8E-11 | 2E-09 | 0.2727273 | nonsurface |
| ENY2 | 1.12 | 9.4726 | 44.8742 | 9E-11 | 2E-09 | 0.6233766 | n.a. |
| ENSECAG00000011858 | 1.104 | 9.5887 | 41.8417 | 1E-10 | 2E-09 | 0.6233766 | n.a. |
| RHOB | 1.394 | 9.2893 | 128.484 | 1E-10 | 2E-09 | 0.3766234 | n.a. |
| ANXA6 | 1.345 | 9.5613 | 41.4659 | 1E-10 | 2E-09 | 0.3636364 | n.a. |
| GARS | 1.35 | 9.3797 | 60.6585 | 2E-10 | 3E-09 | 0.3636364 | n.a. |
| EGLN3 | 1.042 | 9.3249 | 113.167 | 2E-10 | 3E-09 | 0.6103896 | n.a. |
| PEBP1 | 1.055 | 9.6042 | 40.5845 | 2E-10 | 3E-09 | 0.6233766 | n.a. |
| MYD88 | 1.115 | 9.3632 | 51.3798 | 2E-10 | 4E-09 | 0.5324675 | n.a. |
| NDE1 | 1.284 | 9.2796 | 106.698 | 2E-10 | 4E-09 | 0.3376623 | n.a. |
| ENSECAG00000020991 | 1.087 | 9.2681 | 143.766 | 2E-10 | 4E-09 | 0.3896104 | n.a. |
| PHF20 | 1.237 | 9.302 | 73.2419 | 2E-10 | 4E-09 | 0.2727273 | n.a. |
| ENSECAG00000038583 | 1.28 | 9.3387 | 65.8595 | 3E-10 | 6E-09 | 0.3376623 | n.a. |
| OSTC | 1.088 | 9.5663 | 39.066 | 4E-10 | 7E-09 | 0.6493506 | nonsurface |
| RIPOR2 | 1.267 | 9.4745 | 42.799 | 7E-10 | 1E-08 | 0.3246753 | n.a. |
| DNAJC8 | 1.203 | 9.5664 | 37.5757 | 9E-10 | 1E-08 | 0.5064935 | n.a. |
| OSBPL8 | 1.038 | 9.334 | 44.9054 | 2E-09 | 3E-08 | 0.3896104 | nonsurface |
| EIF4E1B | 1.358 | 9.2561 | 319.991 | 2E-09 | 3E-08 | 0.3766234 | n.a. |
| MAP3K8 | 1.24 | 9.3226 | 86.6445 | 2E-09 | 3E-08 | 0.3376623 | n.a. |
| ACAA1 | 1.289 | 9.2784 | 106.394 | 2E-09 | 3E-08 | 0.3246753 | nonsurface |
| CYFIP1 | 1.064 | 9.2506 | 343.375 | 2E-09 | 4E-08 | 0.2597403 | n.a. |

| | | | | | | | |
|---|---|---|---|---|---|---|---|
| ENSA | 1.046 | 9.671 | 35.2783 | 3E-09 | 5E-08 | 0.6363636 | n.a. |
| MACF1 | 1.178 | 9.3902 | 40.2774 | 5E-09 | 7E-08 | 0.2987013 | n.a. |
| MAP3K1 | 1.161 | 9.396 | 41.0316 | 6E-09 | 1E-07 | 0.3246753 | n.a. |
| NUCKS1 | 1.142 | 9.7354 | 33.4487 | 7E-09 | 1E-07 | 0.5064935 | n.a. |
| FUT8 | 1.29 | 9.3425 | 60.4977 | 7E-09 | 1E-07 | 0.2857143 | nonsurface |
| HHEX | 1.061 | 9.4401 | 43.9157 | 8E-09 | 1E-07 | 0.5584416 | n.a. |
| RALB | 1.031 | 9.3135 | 52.4976 | 9E-09 | 1E-07 | 0.4155844 | n.a. |
| RASGRP2 | 1.206 | 9.4643 | 37.8311 | 1E-08 | 2E-07 | 0.3636364 | n.a. |
| UBE2E2 | 1.142 | 9.2944 | 69.8415 | 1E-08 | 2E-07 | 0.3246753 | n.a. |
| PLEKHO1 | 1.013 | 9.3156 | 76.5359 | 2E-08 | 3E-07 | 0.4675325 | n.a. |
| AKIRIN2 | 1.072 | 9.4269 | 35.8204 | 2E-08 | 3E-07 | 0.4545455 | n.a. |
| SDHC | 1.11 | 9.4385 | 39.1342 | 2E-08 | 3E-07 | 0.5194805 | nonsurface |
| POLR2K | 1.226 | 9.3995 | 44.9253 | 2E-08 | 3E-07 | 0.3766234 | n.a. |
| ENSECAG00000037215 | 1.068 | 9.3741 | 43.9355 | 2E-08 | 3E-07 | 0.4545455 | n.a. |
| MIER1 | 1.102 | 9.3938 | 37.59 | 2E-08 | 3E-07 | 0.3766234 | n.a. |
| MRTFA | 1.207 | 9.2972 | 69.9972 | 2E-08 | 3E-07 | 0.2727273 | n.a. |
| SRSF5 | 1.005 | 9.8174 | 31.0132 | 3E-08 | 4E-07 | 0.5844156 | n.a. |
| CEPT1 | 1.208 | 9.3104 | 60.7267 | 3E-08 | 4E-07 | 0.2857143 | n.a. |
| ENSECAG00000037904 | 1.127 | 9.482 | 32.869 | 4E-08 | 5E-07 | 0.3506494 | n.a. |
| RALA | 1.166 | 9.3729 | 42.8256 | 4E-08 | 5E-07 | 0.2727273 | n.a. |
| ENSECAG00000038252 | 1.189 | 9.397 | 44.7921 | 4E-08 | 6E-07 | 0.3116883 | n.a. |
| SH3KBP1 | 1.18 | 9.4466 | 36.8173 | 5E-08 | 7E-07 | 0.3376623 | n.a. |
| XAF1 | 1.096 | 9.2818 | 66.8601 | 6E-08 | 9E-07 | 0.2597403 | n.a. |
| MPHOSPH8 | 1.119 | 9.4682 | 31.3896 | 7E-08 | 9E-07 | 0.3636364 | n.a. |
| SMIM10L1 | 1.04 | 9.5295 | 28.741 | 8E-08 | 1E-06 | 0.4155844 | n.a. |
| RNF7 | 1.007 | 9.4313 | 28.9766 | 1E-07 | 1E-06 | 0.4025974 | n.a. |
| ARL8A | 1.026 | 9.3214 | 44.7181 | 1E-07 | 2E-06 | 0.3116883 | n.a. |
| AKR1B1 | 1.204 | 9.4302 | 38.4149 | 2E-07 | 2E-06 | 0.3116883 | n.a. |
| CHCHD7 | 1.019 | 9.4135 | 33.1779 | 2E-07 | 2E-06 | 0.3636364 | n.a. |
| NUB1 | 1.038 | 9.4301 | 28.9805 | 2E-07 | 2E-06 | 0.3376623 | n.a. |
| ENSECAG00000038714 | 1.116 | 9.3099 | 102.798 | 2E-07 | 2E-06 | 0.3376623 | n.a. |
| TRAF3IP3 | 1.116 | 9.801 | 27.0167 | 2E-07 | 3E-06 | 0.3636364 | nonsurface |
| SLC3A2 | 1.102 | 9.3565 | 40.8118 | 3E-07 | 3E-06 | 0.3766234 | surface |
| ENSECAG00000002652 | 1.149 | 9.2884 | 75.3664 | 3E-07 | 3E-06 | 0.2987013 | n.a. |
| ENSECAG00000038022 | 1.011 | 9.3419 | 40.2851 | 3E-07 | 4E-06 | 0.2987013 | n.a. |
| DNAJC2 | 1.178 | 9.4123 | 36.3564 | 3E-07 | 4E-06 | 0.2987013 | n.a. |
| ENSECAG00000019430 | 1.01 | 9.3323 | 43.4906 | 4E-07 | 5E-06 | 0.4545455 | n.a. |
| TLR7 | 1.012 | 9.2655 | 139.811 | 4E-07 | 5E-06 | 0.3506494 | surface |
| PDHB | 1.149 | 9.3766 | 40.5876 | 5E-07 | 6E-06 | 0.2727273 | n.a. |
| SUN2 | 1.076 | 9.5359 | 25.373 | 5E-07 | 7E-06 | 0.3246753 | nonsurface |
| SNU13 | 1.006 | 9.6007 | 24.9506 | 6E-07 | 7E-06 | 0.4285714 | n.a. |
| PFDN6 | 1.221 | 9.3579 | 46.1366 | 6E-07 | 8E-06 | 0.2597403 | n.a. |
| CCDC88A | 1.049 | 9.3018 | 176.424 | 7E-07 | 8E-06 | 0.5324675 | n.a. |
| DEGS1 | 1.01 | 9.3656 | 34.7864 | 8E-07 | 1E-05 | 0.3636364 | nonsurface |
| PSMD14 | 1.154 | 9.3754 | 42.4399 | 8E-07 | 1E-05 | 0.2727273 | n.a. |
| MRPS14 | 1.125 | 9.367 | 38.8632 | 9E-07 | 1E-05 | 0.3376623 | n.a. |
| RIN3 | 1.118 | 9.3011 | 65.769 | 1E-06 | 1E-05 | 0.2857143 | n.a. |
| UMAD1 | 1.077 | 9.3877 | 35.3432 | 1E-06 | 1E-05 | 0.3376623 | n.a. |

| | | | | | | | |
|---|---|---|---|---|---|---|---|
| eca-mir-9051 | 1.014 | 9.2854 | 60.6798 | 1E-06 | 1E-05 | 0.3506494 | n.a. |
| WAS | 1.075 | 9.441 | 32.0954 | 1E-06 | 2E-05 | 0.4155844 | n.a. |
| PIK3R1 | 1.028 | 9.3963 | 28.8815 | 3E-06 | 4E-05 | 0.2727273 | n.a. |
| UQCRC2 | 1.033 | 9.3691 | 35.8671 | 3E-06 | 4E-05 | 0.3116883 | n.a. |
| ARHGAP4 | 1.048 | 9.3823 | 33.9169 | 4E-06 | 4E-05 | 0.3376623 | n.a. |
| TSR2 | 1.045 | 9.3864 | 31.3269 | 4E-06 | 4E-05 | 0.2987013 | n.a. |
| VTA1 | 1.082 | 9.3204 | 44.5179 | 5E-06 | 5E-05 | 0.2727273 | n.a. |
| AKAP13 | 1.09 | 9.3016 | 52.9442 | 5E-06 | 5E-05 | 0.2857143 | n.a. |
| MPP1 | 1.004 | 9.3612 | 39.677 | 7E-06 | 7E-05 | 0.3636364 | n.a. |
| UBE2N | 1.015 | 9.5201 | 23.7328 | 8E-06 | 8E-05 | 0.3636364 | n.a. |
| ENSECAG00000016410 | 1.013 | 9.256 | 142.781 | 2E-05 | 0.0002 | 0.3636364 | n.a. |
| FAM89B | 1.012 | 9.3314 | 40.1627 | 2E-05 | 0.0002 | 0.3116883 | n.a. |
