## Supplementary material for "Single cell resolution landscape of equine peripheral blood mononuclear cells reveals diverse immune cell subtypes including T-bet^+^ B cells": Dataset S4

| genes | logFC | logCPM | F | PValue | FDR | percent.exp | Surfacome.Label |
|---|---|---|---|---|---|---|---|
| S100A4 | 5.057 | 11.292 | 2185.55 | 0 | 0 | 0.97411 | n.a. |
| FCER1A | 4.093 | 9.4154 | 4933.38 | 0 | 0 | 0.6699029 | surface |
| LYZ | 3.866 | 11.749 | 1768.79 | 0 | 0 | 0.5954693 | n.a. |
| FCER1G | 2.104 | 10.204 | 2409.95 | 0 | 0 | 1 | nonsurface |
| RNASE6 | 2.876 | 10.205 | 1400.19 | 2E-300 | 2E-297 | 0.9514563 | n.a. |
| RNASE4 | 2.705 | 10.101 | 1344.44 | 8E-289 | 8E-286 | 0.9368932 | n.a. |
| S100A6 | 3.512 | 10.437 | 1037.57 | 3E-224 | 2E-221 | 0.9692557 | n.a. |
| DRA | 1.764 | 11.742 | 1009.84 | 2E-218 | 1E-215 | 0.9983819 | n.a. |
| ENSECAG00000035085 | 3.132 | 9.2904 | 3552.9 | 5E-208 | 3E-205 | 0.3996764 | n.a. |
| CD74 | 1.328 | 12.628 | 909.593 | 3E-197 | 2E-194 | 1 | surface |
| DRB | 1.9 | 11.205 | 859.198 | 1E-186 | 7E-184 | 0.9983819 | n.a. |
| SPI1 | 1.847 | 9.6618 | 885.831 | 6E-166 | 3E-163 | 0.9902913 | n.a. |
| S100A5 | 2.962 | 9.5834 | 729.631 | 5E-159 | 2E-156 | 0.6084142 | n.a. |
| TIMP1 | 3.01 | 9.3333 | 1328.42 | 2E-154 | 6E-152 | 0.7022654 | n.a. |
| CD1E2 | 2.629 | 9.2674 | 1789.92 | 6E-151 | 2E-148 | 0.6294498 | n.a. |
| S100A10 | 2.061 | 10.866 | 651.924 | 2E-142 | 7E-140 | 0.9919094 | nonsurface |
| MNDA | 2.642 | 10.258 | 619.582 | 1E-135 | 5E-133 | 0.9320388 | n.a. |
| ENSECAG00000040634 | 2.975 | 9.9556 | 594.627 | 3E-130 | 1E-127 | 0.9029126 | n.a. |
| CST3 | 1.077 | 11.921 | 594.41 | 3E-130 | 1E-127 | 1 | n.a. |
| PLBD1 | 1.491 | 9.972 | 651.37 | 5E-129 | 2E-126 | 0.828479 | n.a. |
| DQA | 1.635 | 10.712 | 579.094 | 7E-127 | 2E-124 | 0.9983819 | n.a. |
| ENSECAG00000038313 | 2.18 | 9.3718 | 770.723 | 4E-125 | 1E-122 | 0.7524272 | n.a. |
| CFP | 2.217 | 9.4343 | 2231.03 | 9E-124 | 2E-121 | 0.9514563 | n.a. |
| SYNE4 | 1.118 | 10.39 | 543.005 | 3E-119 | 9E-117 | 0.9967638 | nonsurface |
| S100P | 2.55 | 9.8205 | 1739.17 | 2E-117 | 5E-115 | 0.276699 | n.a. |
| TSPO | 2.137 | 9.7323 | 489.343 | 3E-107 | 6E-105 | 0.6925566 | nonsurface |
| Eqca-DQB1 | 1.365 | 10.925 | 446.617 | 2E-98 | 3E-96 | 0.9951456 | n.a. |
| VIM | 1.204 | 11.286 | 444.868 | 4E-98 | 8E-96 | 1 | nonsurface |
| ENSECAG00000033857 | 2.4 | 9.3316 | 2745.46 | 8E-96 | 2E-93 | 0.512945 | n.a. |
| ENSECAG00000040180 | 2.324 | 9.7881 | 420.459 | 7E-93 | 1E-90 | 0.7346278 | n.a. |
| HACD4 | 2.353 | 9.3015 | 2436.15 | 1E-92 | 2E-90 | 0.5711974 | n.a. |
| ENSECAG00000019318 | 1.845 | 9.3455 | 479.993 | 7E-92 | 1E-89 | 0.7864078 | n.a. |
| ABCA6 | 2.311 | 9.28 | 2258.01 | 7E-92 | 1E-89 | 0.4271845 | surface |
| CSF1R | 2.137 | 9.3336 | 2014.05 | 5E-90 | 9E-88 | 0.5404531 | surface |
| ENSECAG00000029716 | 1.961 | 9.8771 | 404.711 | 2E-89 | 3E-87 | 0.8462783 | n.a. |
| MS4A7 | 2.187 | 9.3128 | 1704.95 | 6E-88 | 1E-85 | 0.5679612 | n.a. |
| ENSECAG00000029951 | 1.955 | 9.3173 | 431.338 | 3E-87 | 4E-85 | 0.6100324 | n.a. |
| IFI30 | 1.356 | 9.9429 | 390.99 | 2E-86 | 2E-84 | 0.9919094 | n.a. |
| ANXA2 | 1.769 | 9.8875 | 386.03 | 2E-85 | 3E-83 | 0.9320388 | n.a. |
| SERPINB1 | 1.973 | 9.4507 | 379.487 | 5E-84 | 7E-82 | 0.8139159 | nonsurface |
| CTSZ | 1.258 | 9.8631 | 377.018 | 2E-83 | 2E-81 | 0.8980583 | n.a. |
| PRNP | 2.202 | 9.3261 | 941.836 | 1E-80 | 2E-78 | 0.4967638 | nonsurface |
| GIMAP7 | 2.019 | 10.513 | 360.678 | 5E-80 | 7E-78 | 0.7152104 | n.a. |
| ENSECAG00000007681 | 2.213 | 9.6603 | 354.872 | 9E-79 | 1E-76 | 0.6229773 | n.a. |
| ENSECAG00000035431 | 2.107 | 9.323 | 1739.26 | 1E-78 | 2E-76 | 0.5631068 | n.a. |
| FCGRT | 1.866 | 9.3372 | 409.13 | 5E-77 | 6E-75 | 0.868932 | surface |
| IFI27 | 2.001 | 9.7314 | 311.802 | 2E-69 | 2E-67 | 0.381877 | nonsurface |

| | | | | | | | |
|---|---|---|---|---|---|---|---|
| ENSECAG00000031569 | 1.236 | 11.194 | 311.296 | 2E-69 | 3E-67 | 0.9967638 | n.a. |
| EGLN3 | 1.321 | 9.3249 | 367.031 | 1E-68 | 1E-66 | 0.7491909 | n.a. |
| ENSECAG00000007663 | 1.765 | 9.734 | 303.696 | 1E-67 | 1E-65 | 0.5987055 | n.a. |
| HSPB1 | 1.49 | 9.6778 | 303.369 | 1E-67 | 1E-65 | 0.9482201 | n.a. |
| KCNE3 | 1.976 | 9.2985 | 1394.86 | 2E-62 | 2E-60 | 0.5711974 | n.a. |
| MEF2C | 1.163 | 9.4205 | 306.167 | 3E-62 | 3E-60 | 0.8656958 | n.a. |
| ASGR2 | 1.97 | 9.2429 | 1154.06 | 3E-61 | 3E-59 | 0.3300971 | surface |
| ENSECAG00000021796 | 1.358 | 9.6911 | 269.378 | 3E-60 | 3E-58 | 0.9724919 | n.a. |
| MHCB3 | 1.893 | 10.265 | 256.214 | 2E-57 | 2E-55 | 0.6812298 | n.a. |
| FGGY | 1.839 | 9.2697 | 832.895 | 4E-57 | 4E-55 | 0.6278317 | n.a. |
| ACTG1 | 1.269 | 10.745 | 254.43 | 4E-57 | 4E-55 | 1 | n.a. |
| SLAMF9 | 1.794 | 9.2456 | 1454.88 | 2E-56 | 2E-54 | 0.3996764 | surface |
| ENSECAG00000015010 | 1.817 | 9.2872 | 1551.66 | 3E-56 | 3E-54 | 0.3883495 | n.a. |
| HLA-DMA | 1.468 | 9.6051 | 248.844 | 7E-56 | 6E-54 | 0.9433657 | surface |
| CD68 | 1.353 | 9.3465 | 576.502 | 4E-55 | 4E-53 | 0.7281553 | surface |
| FGL2 | 1.387 | 9.488 | 268.735 | 3E-52 | 2E-50 | 0.7961165 | n.a. |
| YEATS4 | 1.636 | 9.4194 | 229.025 | 1E-51 | 1E-49 | 0.7540453 | n.a. |
| NFAM1 | 1.601 | 9.3582 | 673.223 | 3E-51 | 2E-49 | 0.7621359 | surface |
| FGR | 1.024 | 9.4883 | 247.402 | 2E-50 | 1E-48 | 0.9158576 | n.a. |
| PKM | 1.48 | 9.8077 | 213.372 | 4E-48 | 3E-46 | 0.8656958 | n.a. |
| SAMD9L | 1.674 | 9.4216 | 244.802 | 2E-46 | 1E-44 | 0.3834951 | n.a. |
| FABP3 | 1.226 | 9.3412 | 252.068 | 1E-45 | 1E-43 | 0.5841424 | n.a. |
| MSN | 1.609 | 9.6123 | 199.583 | 3E-45 | 2E-43 | 0.8074434 | n.a. |
| ENSECAG00000019309 | 1.583 | 9.3991 | 304.823 | 8E-45 | 6E-43 | 0.3640777 | n.a. |
| REXO2 | 1.322 | 9.4557 | 192.313 | 1E-43 | 9E-42 | 0.7216828 | n.a. |
| BLVRB | 1.338 | 9.4497 | 199.333 | 1E-43 | 9E-42 | 0.7249191 | n.a. |
| ALDH2 | 1.524 | 9.5616 | 385.993 | 2E-43 | 1E-41 | 0.5825243 | n.a. |
| GLIPR2 | 1.535 | 9.47 | 229.932 | 4E-43 | 3E-41 | 0.5938511 | n.a. |
| IL13RA1 | 1.497 | 9.3015 | 823.555 | 1E-42 | 9E-41 | 0.6860841 | surface |
| CD14 | 1.611 | 9.3178 | 1761.75 | 2E-42 | 1E-40 | 0.7200647 | surface |
| CKB | 1.518 | 9.3208 | 1070.03 | 5E-42 | 3E-40 | 0.6957929 | n.a. |
| MYCBP2 | 1.426 | 9.5377 | 182.649 | 2E-41 | 1E-39 | 0.8915858 | n.a. |
| HNMT | 1.441 | 9.2852 | 1767.2 | 3E-41 | 2E-39 | 0.2621359 | n.a. |
| HK3 | 1.606 | 9.284 | 1287.97 | 5E-41 | 3E-39 | 0.3559871 | n.a. |
| SAT1 | 1.129 | 9.7271 | 180.05 | 6E-41 | 4E-39 | 0.9368932 | n.a. |
| LGALS3 | 1.406 | 9.5239 | 226.306 | 7E-40 | 4E-38 | 0.3559871 | n.a. |
| ENSECAG00000016543 | 1.467 | 9.8125 | 172.895 | 2E-39 | 1E-37 | 0.8559871 | n.a. |
| ENSECAG00000024181 | 1.439 | 9.2586 | 1484.29 | 3E-39 | 2E-37 | 0.3009709 | n.a. |
| PAK1 | 1.293 | 9.331 | 702.905 | 7E-39 | 4E-37 | 0.7847896 | n.a. |
| FABP5 | 1.577 | 9.5264 | 167.25 | 4E-38 | 2E-36 | 0.3834951 | n.a. |
| ANXA5 | 1.156 | 9.5163 | 165.551 | 8E-38 | 5E-36 | 0.8430421 | n.a. |
| SRGN | 1.036 | 10.485 | 161.926 | 5E-37 | 3E-35 | 0.9563107 | n.a. |
| IGSF6 | 1.095 | 9.3651 | 538.37 | 1E-36 | 7E-35 | 0.6440129 | surface |
| RGS10 | 1.42 | 9.5903 | 159.279 | 2E-36 | 1E-34 | 0.8446602 | n.a. |
| ENSECAG00000017042 | 1.167 | 9.3458 | 546.32 | 2E-36 | 1E-34 | 0.7071197 | n.a. |
| TKT | 1.106 | 9.6069 | 157.509 | 5E-36 | 3E-34 | 0.8770227 | n.a. |
| CAPN2 | 1.387 | 9.3317 | 210.395 | 9E-35 | 5E-33 | 0.5016181 | n.a. |
| S100A11 | 1.268 | 9.847 | 151.247 | 1E-34 | 6E-33 | 0.8543689 | n.a. |

| | | | | | | | |
|---|---|---|---|---|---|---|---|
| DOK2 | 1.404 | 9.3423 | 277.762 | 1E-34 | 7E-33 | 0.3414239 | n.a. |
| DHRS3 | 1.428 | 9.3111 | 344.822 | 1E-33 | 6E-32 | 0.5533981 | nonsurface |
| NMI | 1.274 | 9.4149 | 146.051 | 1E-33 | 7E-32 | 0.579288 | n.a. |
| TCN2 | 1.292 | 9.2547 | 687.4 | 7E-33 | 4E-31 | 0.3980583 | n.a. |
| PLEKHO1 | 1.304 | 9.3156 | 273.61 | 9E-33 | 4E-31 | 0.6666667 | n.a. |
| JUNB | 1.314 | 9.5395 | 141.551 | 1E-32 | 7E-31 | 0.7524272 | n.a. |
| NCF2 | 1.262 | 9.3332 | 208.773 | 1E-31 | 5E-30 | 0.5 | n.a. |
| KLF4 | 1.418 | 9.2595 | 971.28 | 9E-31 | 4E-29 | 0.4627832 | n.a. |
| C1H15orf48 | 1.22 | 9.6929 | 189.843 | 1E-30 | 4E-29 | 0.4352751 | n.a. |
| CORO1B | 1.084 | 9.4003 | 142.1 | 3E-30 | 1E-28 | 0.8317152 | n.a. |
| ENSECAG00000029003 | 1.008 | 10.041 | 131.074 | 3E-30 | 1E-28 | 0.9417476 | n.a. |
| ANXA1 | 1.106 | 9.8295 | 128.396 | 1E-29 | 4E-28 | 0.8754045 | n.a. |
| EMP3 | 1.054 | 10.126 | 126.919 | 2E-29 | 9E-28 | 0.8802589 | surface |
| MYADM | 1.29 | 9.3852 | 144.281 | 5E-29 | 2E-27 | 0.5307443 | surface |
| ENSECAG00000009733 | 1.312 | 9.3278 | 178.414 | 1E-28 | 5E-27 | 0.5372168 | n.a. |
| LDHA | 1.044 | 9.6788 | 122.436 | 2E-28 | 8E-27 | 0.9061489 | n.a. |
| CX3CR1 | 1.352 | 9.2887 | 477.198 | 6E-28 | 2E-26 | 0.4223301 | surface |
| GLUL | 1.325 | 9.3271 | 276.801 | 2E-27 | 7E-26 | 0.3867314 | n.a. |
| AIM2 | 1.267 | 9.3585 | 299.732 | 2E-27 | 8E-26 | 0.6634304 | n.a. |
| ENSECAG00000019932 | 1.212 | 9.3315 | 181.553 | 3E-27 | 1E-25 | 0.5889968 | n.a. |
| TCF7L2 | 1.286 | 9.2572 | 512.019 | 3E-27 | 1E-25 | 0.328479 | n.a. |
| CASP4 | 1.198 | 9.4053 | 148.345 | 1E-26 | 4E-25 | 0.2993528 | n.a. |
| FLNA | 1.125 | 9.4126 | 137.107 | 1E-26 | 5E-25 | 0.4466019 | n.a. |
| HIF1A | 1.026 | 9.3685 | 113.536 | 8E-26 | 3E-24 | 0.6456311 | n.a. |
| RF01956 | 1.309 | 9.2926 | 374.392 | 5E-25 | 2E-23 | 0.579288 | n.a. |
| LY86 | 1.079 | 9.286 | 392.032 | 8E-25 | 3E-23 | 0.3122977 | n.a. |
| ASCL4 | 1.153 | 9.269 | 720.832 | 1E-24 | 4E-23 | 0.2847896 | n.a. |
| ENSECAG00000017092 | 1.17 | 9.2393 | 1049.61 | 5E-24 | 2E-22 | 0.276699 | n.a. |
| UBE2E3 | 1.049 | 9.3915 | 129.886 | 6E-24 | 2E-22 | 0.5453074 | n.a. |
| LPAR6 | 1.073 | 9.2687 | 410.744 | 1E-23 | 4E-22 | 0.5906149 | surface |
| ID3 | 1.115 | 9.6697 | 100.177 | 2E-23 | 5E-22 | 0.4805825 | n.a. |
| ITGAX | 1.265 | 9.2642 | 499.31 | 2E-23 | 7E-22 | 0.3802589 | surface |
| TFEB | 1.22 | 9.3544 | 259.597 | 2E-23 | 7E-22 | 0.605178 | n.a. |
| NRN1 | 1.211 | 9.2707 | 398.17 | 2E-23 | 7E-22 | 0.3867314 | surface |
| C4BPA | 1.041 | 9.4013 | 376.54 | 8E-23 | 2E-21 | 0.3673139 | n.a. |
| ENSECAG00000028889 | 1.09 | 9.2871 | 374.96 | 1E-22 | 3E-21 | 0.5744337 | n.a. |
| eca-mir-223 | 1.191 | 9.2778 | 675.872 | 4E-22 | 1E-20 | 0.2864078 | n.a. |
| TMEM106A | 1.146 | 9.2932 | 159.251 | 6E-22 | 2E-20 | 0.4466019 | surface |
| CASP1 | 1.185 | 9.2737 | 499.342 | 8E-22 | 2E-20 | 0.4401294 | n.a. |
| CEBPB | 1.087 | 9.407 | 261.133 | 1E-21 | 4E-20 | 0.2669903 | n.a. |
| MECR | 1.041 | 9.3421 | 220.398 | 1E-19 | 3E-18 | 0.276699 | n.a. |
| DPP4 | 1.036 | 9.2884 | 197.578 | 2E-19 | 5E-18 | 0.2912621 | surface |
| LCP2 | 1.04 | 9.4951 | 78.2187 | 1E-18 | 2E-17 | 0.4919094 | n.a. |
| DDIT4 | 1.12 | 9.3328 | 138.672 | 2E-18 | 4E-17 | 0.3317152 | n.a. |
| CREG1 | 1.005 | 9.3229 | 546.458 | 2E-18 | 6E-17 | 0.2718447 | n.a. |
| HMOX1 | 1.034 | 9.3479 | 163.737 | 6E-18 | 1E-16 | 0.4110032 | nonsurface |
| ENSECAG00000034395 | 1.03 | 9.2877 | 209.794 | 6E-18 | 1E-16 | 0.3236246 | n.a. |
| RARG | 1.028 | 9.278 | 214.621 | 7E-18 | 2E-16 | 0.2718447 | n.a. |

| genes | logFC | logCPM | F | PValue | FDR | percent.exp | Surfacome.Label |
|---|---|---|---|---|---|---|---|
| ENSECAG00000039383 | 5.335 | 9.2686 | 24114.1 | 0 | 0 | 0.9635417 | n.a. |
| ID2 | 4.437 | 9.6451 | 1558.59 | 0 | 0 | 0.984375 | n.a. |
| CPVL | 4.324 | 9.3018 | 5716.33 | 0 | 0 | 0.96875 | n.a. |
| ENSECAG00000029928 | 4.287 | 9.2723 | 5436.85 | 0 | 0 | 0.9427083 | n.a. |
| CLEC9A | 4.205 | 9.2481 | 20849.3 | 0 | 0 | 0.859375 | surface |
| ENSECAG00000024996 | 4.161 | 9.2908 | 2249.08 | 0 | 0 | 0.96875 | n.a. |
| BATF3 | 4.159 | 9.2774 | 2995.88 | 0 | 0 | 0.9739583 | n.a. |
| MT3 | 3.725 | 9.2863 | 5094.25 | 0 | 0 | 0.9322917 | n.a. |
| DNASE1L3 | 3.682 | 9.2492 | 9536.46 | 0 | 0 | 0.8385417 | n.a. |
| CST3 | 3.622 | 11.921 | 5142.67 | 0 | 0 | 1 | n.a. |
| ENSECAG00000014585 | 2.901 | 10.271 | 2046.19 | 0 | 0 | 0.9947917 | n.a. |
| DRA | 2.585 | 11.742 | 1844.52 | 0 | 0 | 1 | n.a. |
| CD74 | 1.917 | 12.628 | 1511.98 | 0 | 0 | 1 | surface |
| ENSECAG00000009190 | 1.381 | 14.441 | 1491.73 | 0 | 0 | 1 | n.a. |
| GM2A | 2.481 | 9.9979 | 1477.7 | 7E-295 | 3E-292 | 0.9583333 | n.a. |
| GPIHBP1 | 3.359 | 9.2751 | 3325.63 | 6E-292 | 2E-289 | 0.9427083 | surface |
| DRB | 2.555 | 11.205 | 1330.35 | 7E-286 | 3E-283 | 1 | n.a. |
| CKB | 2.773 | 9.3208 | 2030.39 | 1E-194 | 4E-192 | 0.90625 | n.a. |
| LRRK2 | 3.226 | 9.2707 | 1536.14 | 1E-192 | 3E-190 | 0.8802083 | n.a. |
| DQA | 2.2 | 10.712 | 873.081 | 2E-189 | 5E-187 | 1 | n.a. |
| CLNK | 3.185 | 9.2461 | 3937.41 | 4E-189 | 1E-186 | 0.7552083 | n.a. |
| ENSECAG00000020136 | 3.291 | 9.7014 | 829.983 | 2E-180 | 7E-178 | 0.984375 | n.a. |
| Eqca-DQB1 | 1.966 | 10.925 | 754.906 | 2E-164 | 5E-162 | 1 | n.a. |
| DQB | 2.325 | 10.011 | 708.228 | 2E-154 | 4E-152 | 0.96875 | n.a. |
| ENSECAG00000038313 | 2.414 | 9.3718 | 869.822 | 3E-146 | 8E-144 | 0.828125 | n.a. |
| RNASE6 | 2.172 | 10.205 | 650.456 | 7E-141 | 2E-138 | 0.9739583 | n.a. |
| ENSECAG00000005400 | 2.775 | 9.2385 | 17325.9 | 1E-136 | 3E-134 | 0.6614583 | n.a. |
| CD8A | 2.77 | 9.4281 | 604.888 | 2E-132 | 4E-130 | 0.71875 | surface |
| RNASE4 | 2.087 | 10.101 | 655.613 | 5E-129 | 1E-126 | 0.96875 | n.a. |
| ENSECAG00000035094 | 2.495 | 9.2863 | 877.691 | 1E-124 | 3E-122 | 0.8541667 | n.a. |
| CFP | 2.294 | 9.4343 | 2052.89 | 2E-124 | 5E-122 | 0.9583333 | n.a. |
| RGS2 | 2.767 | 9.2835 | 2027.16 | 3E-124 | 5E-122 | 0.5104167 | n.a. |
| S100A10 | 2.066 | 10.866 | 543.564 | 3E-119 | 5E-117 | 1 | nonsurface |
| SPP1 | 2.648 | 9.2475 | 1684.21 | 3E-116 | 6E-114 | 0.5572917 | n.a. |
| RAB7B | 2.607 | 9.2391 | 4273.32 | 5E-112 | 9E-110 | 0.6770833 | n.a. |
| CST7 | 2.473 | 9.312 | 921.908 | 4E-105 | 6E-103 | 0.640625 | n.a. |
| VAC14 | 2.395 | 9.2854 | 473.912 | 2E-104 | 4E-102 | 0.84375 | n.a. |
| FOS | 2.101 | 9.3156 | 620.492 | 3E-104 | 5E-102 | 0.671875 | n.a. |
| ENSECAG00000035439 | 2.419 | 9.2665 | 2818.38 | 6E-104 | 1E-101 | 0.6041667 | n.a. |
| S100A11 | 2.425 | 9.847 | 458.593 | 4E-101 | 7E-99 | 1 | n.a. |
| RUBCNL | 2.037 | 9.2489 | 1306.68 | 2E-96 | 2E-94 | 0.609375 | n.a. |
| ENSECAG00000019318 | 2.074 | 9.3455 | 595.646 | 2E-93 | 2E-91 | 0.9166667 | n.a. |
| PKIB | 2.125 | 9.2759 | 1021.87 | 1E-92 | 2E-90 | 0.8854167 | n.a. |
| RAB38 | 2.382 | 9.2603 | 1378.7 | 3E-91 | 5E-89 | 0.8802083 | n.a. |
| SPI1 | 1.573 | 9.6618 | 529.68 | 1E-89 | 2E-87 | 0.9791667 | n.a. |
| NAAA | 2.49 | 9.5818 | 393.412 | 5E-87 | 6E-85 | 0.9479167 | n.a. |
| CADM1 | 2.353 | 9.2459 | 1910.7 | 5E-84 | 7E-82 | 0.6770833 | surface |

| | | | | | | | |
|---|---|---|---|---|---|---|---|
| LY6E | 1.84 | 10.241 | 374.218 | 6E-83 | 8E-81 | 0.9791667 | surface |
| PAK1 | 1.933 | 9.331 | 1148.14 | 2E-76 | 2E-74 | 0.9114583 | n.a. |
| IRF8 | 1.406 | 9.303 | 328.089 | 5E-73 | 7E-71 | 0.9791667 | n.a. |
| ENSECAG00000028889 | 1.936 | 9.2871 | 740.905 | 2E-69 | 2E-67 | 0.75 | n.a. |
| PABPC1 | 1.031 | 10.847 | 296.513 | 4E-66 | 4E-64 | 1 | n.a. |
| CSRP1 | 1.653 | 9.2812 | 296.117 | 4E-66 | 5E-64 | 0.8645833 | n.a. |
| PPT1 | 1.807 | 9.4371 | 275.018 | 2E-61 | 2E-59 | 0.84375 | n.a. |
| ENSECAG00000028100 | 2.014 | 9.2366 | 3022.67 | 2E-61 | 2E-59 | 0.4166667 | n.a. |
| TCTN2 | 1.985 | 9.2411 | 1995.95 | 2E-61 | 2E-59 | 0.453125 | surface |
| CLIC2 | 1.919 | 9.286 | 738.094 | 1E-60 | 1E-58 | 0.734375 | n.a. |
| ENSECAG00000021796 | 1.512 | 9.6911 | 265.572 | 2E-59 | 2E-57 | 0.9739583 | n.a. |
| ENSECAG00000039164 | 1.77 | 9.2362 | 5406.94 | 4E-59 | 4E-57 | 0.2916667 | n.a. |
| ENSECAG00000033359 | 1.873 | 9.258 | 750.156 | 9E-56 | 8E-54 | 0.5416667 | n.a. |
| SLAMF8 | 1.931 | 9.2377 | 1654.28 | 3E-55 | 3E-53 | 0.4739583 | surface |
| SNX22 | 1.858 | 9.2475 | 964.969 | 5E-55 | 4E-53 | 0.484375 | n.a. |
| RGS1 | 1.68 | 9.3256 | 239.741 | 7E-54 | 6E-52 | 0.7708333 | n.a. |
| Eqca-2 | 1.034 | 11.477 | 239.009 | 1E-53 | 8E-52 | 0.9791667 | n.a. |
| DCTPP1 | 1.843 | 9.3868 | 237.633 | 2E-53 | 2E-51 | 0.90625 | n.a. |
| FGD2 | 1.944 | 9.2999 | 541.814 | 3E-53 | 2E-51 | 0.6770833 | n.a. |
| WDFY4 | 1.441 | 9.2866 | 280.088 | 5E-53 | 4E-51 | 0.7760417 | nonsurface |
| PLEK | 1.628 | 9.3075 | 272.968 | 6E-53 | 5E-51 | 0.71875 | n.a. |
| PSMB8 | 1.337 | 10.346 | 231.781 | 4E-52 | 3E-50 | 0.9895833 | n.a. |
| Eqca-DOB1 | 1.378 | 9.4745 | 308.351 | 6E-52 | 4E-50 | 0.8020833 | n.a. |
| RGS10 | 1.91 | 9.5903 | 228.259 | 2E-51 | 2E-49 | 0.921875 | n.a. |
| ENSECAG00000034569 | 1.569 | 9.7713 | 223.197 | 3E-50 | 2E-48 | 0.5416667 | n.a. |
| SRGN | 1.362 | 10.485 | 221.306 | 7E-50 | 5E-48 | 0.9791667 | n.a. |
| BTLA | 1.374 | 9.2816 | 506.169 | 8E-50 | 6E-48 | 0.546875 | surface |
| HLA-DMA | 1.446 | 9.6051 | 202.926 | 6E-46 | 4E-44 | 0.9427083 | surface |
| NEURL1 | 1.443 | 9.2404 | 989.733 | 3E-45 | 2E-43 | 0.6458333 | n.a. |
| PPA1 | 1.711 | 9.4873 | 192.454 | 1E-43 | 8E-42 | 0.84375 | n.a. |
| SGK1 | 1.545 | 9.2522 | 650.406 | 2E-43 | 1E-41 | 0.2760417 | n.a. |
| PLEKHO2 | 1.692 | 9.3041 | 250.061 | 3E-43 | 2E-41 | 0.6510417 | n.a. |
| AIG1 | 1.447 | 9.327 | 242.619 | 5E-43 | 3E-41 | 0.8385417 | nonsurface |
| PSTPIP1 | 1.697 | 9.5292 | 184.587 | 6E-42 | 4E-40 | 0.6979167 | n.a. |
| SORBS3 | 1.494 | 9.2441 | 1245.7 | 6E-41 | 3E-39 | 0.3489583 | n.a. |
| ENSECAG00000023388 | 1.724 | 9.2512 | 1172.34 | 1E-40 | 6E-39 | 0.546875 | n.a. |
| ENSECAG00000012400 | 1.282 | 9.3413 | 218.274 | 8E-40 | 4E-38 | 0.4375 | n.a. |
| PDF | 1.621 | 9.2907 | 223.139 | 2E-38 | 9E-37 | 0.5989583 | n.a. |
| GADD45A | 1.654 | 9.2896 | 637.332 | 2E-38 | 1E-36 | 0.5 | n.a. |
| ENSECAG00000006701 | 1.609 | 9.2359 | 1235.75 | 5E-38 | 3E-36 | 0.3385417 | n.a. |
| MYOF | 1.667 | 9.247 | 916.594 | 1E-37 | 6E-36 | 0.3958333 | surface |
| PCMT1 | 1.561 | 9.2898 | 235.111 | 3E-37 | 1E-35 | 0.6510417 | n.a. |
| FAM135A | 1.506 | 9.2372 | 2692.69 | 4E-37 | 2E-35 | 0.3489583 | n.a. |
| ENSECAG00000015275 | 1.565 | 9.3043 | 285.041 | 2E-36 | 8E-35 | 0.5729167 | n.a. |
| CPNE3 | 1.44 | 9.3269 | 157.383 | 5E-36 | 3E-34 | 0.7083333 | n.a. |
| PLPP1 | 1.665 | 9.2503 | 588.844 | 9E-36 | 5E-34 | 0.4270833 | n.a. |
| SNX3 | 1.204 | 9.6999 | 155.555 | 1E-35 | 6E-34 | 0.9947917 | n.a. |
| CA12 | 1.403 | 9.2359 | 2726.41 | 3E-35 | 2E-33 | 0.2760417 | surface |

| | | | | | | | |
|---|---|---|---|---|---|---|---|
| CSTA | 1.691 | 9.2431 | 1472.5 | 4E-34 | 2E-32 | 0.5104167 | n.a. |
| FGL2 | 1.288 | 9.488 | 193.169 | 1E-33 | 5E-32 | 0.796875 | n.a. |
| PPM1M | 1.401 | 9.3461 | 140.995 | 2E-32 | 8E-31 | 0.7291667 | n.a. |
| SLC46A3 | 1.119 | 9.2637 | 160.493 | 8E-32 | 4E-30 | 0.609375 | surface |
| HLA-DOA | 1.346 | 9.3227 | 218.01 | 2E-31 | 1E-29 | 0.78125 | surface |
| FLT3 | 1.173 | 9.2499 | 257.44 | 6E-31 | 2E-29 | 0.7708333 | surface |
| LDHB | 1.161 | 10.019 | 132.718 | 1E-30 | 5E-29 | 0.9791667 | n.a. |
| ENSECAG00000032068 | 1.293 | 9.2351 | 4540.89 | 3E-30 | 1E-28 | 0.2760417 | n.a. |
| HLA-DQB1 | 1.05 | 9.2656 | 236.495 | 3E-30 | 1E-28 | 0.28125 | surface |
| ENSECAG00000013276 | 1.266 | 9.3936 | 130.371 | 4E-30 | 2E-28 | 0.8020833 | n.a. |
| FBL | 1.174 | 9.7416 | 128.792 | 8E-30 | 3E-28 | 0.96875 | n.a. |
| ATP6V0A2 | 1.245 | 9.2479 | 348.101 | 9E-30 | 4E-28 | 0.5520833 | surface |
| LPAR6 | 1.343 | 9.2687 | 436.802 | 1E-29 | 6E-28 | 0.671875 | surface |
| TUBA1A | 1.269 | 10.091 | 126.703 | 2E-29 | 9E-28 | 0.90625 | n.a. |
| TXN | 1.07 | 9.9313 | 126.01 | 3E-29 | 1E-27 | 0.984375 | n.a. |
| PTGER4 | 1.32 | 9.3476 | 126.561 | 1E-28 | 4E-27 | 0.5677083 | surface |
| TSTD1 | 1.408 | 9.2702 | 323.046 | 2E-28 | 6E-27 | 0.5625 | n.a. |
| HCK | 1.426 | 9.2996 | 396.836 | 3E-28 | 1E-26 | 0.59375 | n.a. |
| ARHGEF9 | 1.343 | 9.2412 | 1020.67 | 8E-28 | 3E-26 | 0.2916667 | n.a. |
| VIPR1 | 1.39 | 9.2359 | 2138.1 | 7E-26 | 2E-24 | 0.3177083 | surface |
| MGMT | 1.267 | 9.3322 | 117.276 | 2E-25 | 5E-24 | 0.6145833 | n.a. |
| ENSECAG00000031154 | 1.248 | 9.2755 | 341.027 | 7E-25 | 2E-23 | 0.2760417 | n.a. |
| NEXN | 1.353 | 9.2383 | 1608.08 | 4E-24 | 1E-22 | 0.3489583 | n.a. |
| PIK3CB | 1.286 | 9.2443 | 529.678 | 5E-24 | 1E-22 | 0.4427083 | n.a. |
| ENSECAG00000016410 | 1.26 | 9.256 | 676.815 | 5E-24 | 2E-22 | 0.390625 | n.a. |
| CRHBP | 1.172 | 9.2452 | 304.097 | 1E-23 | 5E-22 | 0.390625 | n.a. |
| RHOB | 1.172 | 9.2893 | 175.755 | 2E-23 | 5E-22 | 0.484375 | n.a. |
| TBXAS1 | 1.286 | 9.2862 | 719.676 | 2E-23 | 8E-22 | 0.46875 | nonsurface |
| WDR41 | 1.187 | 9.3199 | 111.694 | 6E-23 | 2E-21 | 0.578125 | n.a. |
| GNL1 | 1.096 | 9.2852 | 116.729 | 6E-23 | 2E-21 | 0.578125 | n.a. |
| CD40 | 1.308 | 9.2962 | 258.19 | 8E-23 | 3E-21 | 0.4427083 | surface |
| OPTN | 1.227 | 9.4432 | 105.836 | 3E-22 | 1E-20 | 0.390625 | n.a. |
| TBC1D9 | 1.272 | 9.2457 | 644.822 | 5E-22 | 1E-20 | 0.359375 | n.a. |
| MPST | 1.203 | 9.2994 | 154.013 | 9E-22 | 3E-20 | 0.4427083 | n.a. |
| TRIO | 1.247 | 9.2396 | 738.826 | 1E-21 | 4E-20 | 0.40625 | n.a. |
| NDUFB4 | 1.04 | 9.5448 | 90.6432 | 2E-21 | 5E-20 | 0.9270833 | nonsurface |
| DUSP1 | 1.172 | 9.3756 | 155.339 | 2E-21 | 6E-20 | 0.4270833 | nonsurface |
| TMEM145 | 1.241 | 9.2908 | 182.127 | 3E-21 | 7E-20 | 0.3489583 | surface |
| SMAGP | 1.208 | 9.3065 | 132.064 | 9E-21 | 3E-19 | 0.4375 | nonsurface |
| SUSD3 | 1.237 | 9.5214 | 87.2427 | 1E-20 | 3E-19 | 0.46875 | surface |
| NFAM1 | 1.091 | 9.3582 | 325.235 | 1E-20 | 4E-19 | 0.671875 | surface |
| COPRS | 1.199 | 9.2687 | 191.033 | 2E-20 | 4E-19 | 0.4375 | n.a. |
| APRT | 1.121 | 9.6859 | 84.7115 | 4E-20 | 1E-18 | 0.8958333 | n.a. |
| ITGA4 | 1.082 | 9.465 | 84.2236 | 5E-20 | 1E-18 | 0.7708333 | surface |
| LASP1 | 1.123 | 9.4194 | 83.7719 | 6E-20 | 2E-18 | 0.765625 | n.a. |
| BCL2A1 | 1.14 | 9.2926 | 200.683 | 9E-20 | 2E-18 | 0.34375 | n.a. |
| TMEM50B | 1.141 | 9.2661 | 263.277 | 2E-19 | 5E-18 | 0.484375 | nonsurface |
| ENSECAG00000031913 | 1.147 | 9.2438 | 572.873 | 1E-18 | 3E-17 | 0.296875 | n.a. |

| | | | | | | | |
|---|---|---|---|---|---|---|---|
| ENSECAG00000037904 | 1.166 | 9.482 | 77.6946 | 1E-18 | 3E-17 | 0.59375 | n.a. |
| PYGL | 1.074 | 9.2869 | 202.313 | 1E-18 | 3E-17 | 0.4635417 | n.a. |
| LGMN | 1.138 | 9.3223 | 155.423 | 2E-18 | 4E-17 | 0.3020833 | n.a. |
| TAP1 | 1.127 | 9.3381 | 100.671 | 2E-18 | 5E-17 | 0.5052083 | n.a. |
| GCSH | 1.07 | 9.4155 | 74.3467 | 7E-18 | 2E-16 | 0.7239583 | n.a. |
| LY5 | 1.135 | 9.2865 | 164.617 | 1E-17 | 3E-16 | 0.3645833 | n.a. |
| ENSECAG00000005359 | 1.139 | 9.311 | 117.932 | 7E-17 | 2E-15 | 0.4635417 | n.a. |
| OSCAR | 1.046 | 9.2533 | 404.487 | 4E-16 | 8E-15 | 0.4322917 | n.a. |
| IL21R | 1.093 | 9.3399 | 117.44 | 7E-16 | 1E-14 | 0.3020833 | surface |
| ZNF664 | 1.061 | 9.2525 | 259.379 | 7E-16 | 1E-14 | 0.2708333 | n.a. |
| ZNF385A | 1.011 | 9.2523 | 341.396 | 1E-15 | 2E-14 | 0.5260417 | n.a. |
| QPCT | 1.067 | 9.3093 | 323.141 | 2E-15 | 5E-14 | 0.3177083 | nonsurface |
| LIMCH1 | 1.009 | 9.2424 | 436.2 | 4E-14 | 8E-13 | 0.3645833 | n.a. |
| RNPEP | 1.024 | 9.2739 | 99.606 | 7E-14 | 1E-12 | 0.484375 | n.a. |

| genes | logFC | logCPM | F | PValue | FDR | percent.exp | Surfacome.Label |
|---|---|---|---|---|---|---|---|
| ENSECAG00000003702 | 5.549 | 9.282 | 10390.8 | 0 | 0 | 1 | n.a. |
| ENSECAG00000017313 | 5.518 | 9.2752 | 13080 | 0 | 0 | 1 | n.a. |
| IRF7 | 4.838 | 9.3097 | 2531.69 | 0 | 0 | 0.9369369 | n.a. |
| JCHAIN | 4.792 | 9.7044 | 4527.01 | 0 | 0 | 0.8108108 | n.a. |
| SOD1 | 4.644 | 9.8391 | 2555.99 | 0 | 0 | 1 | n.a. |
| FCRLA | 4.557 | 9.3304 | 4149.98 | 0 | 0 | 0.9369369 | n.a. |
| CYB561A3 | 4.494 | 9.3685 | 2538.89 | 0 | 0 | 0.981982 | nonsurface |
| RAB24 | 4.448 | 9.4289 | 1537.65 | 0 | 0 | 0.963964 | n.a. |
| ENSECAG00000022644 | 4.422 | 9.2487 | 7947.44 | 0 | 0 | 1 | n.a. |
| ENSECAG00000039088 | 4.338 | 9.7274 | 1684.31 | 0 | 0 | 0.981982 | n.a. |
| ENSECAG00000001923 | 4.081 | 9.6006 | 1729.78 | 0 | 0 | 1 | n.a. |
| ENSECAG00000008542 | 1.332 | 14.412 | 2022.79 | 0 | 0 | 1 | n.a. |
| CTSC | 4.107 | 9.6133 | 1488.39 | 0 | 0 | 0.972973 | n.a. |
| ENSECAG00000030777 | 3.717 | 9.3773 | 3649.92 | 0 | 0 | 0.9009009 | n.a. |
| UGCG | 4.09 | 9.3115 | 1417.72 | 4E-304 | 2E-301 | 0.9279279 | nonsurface |
| CYSTM1 | 3.11 | 9.3828 | 1340.57 | 5E-288 | 2E-285 | 0.990991 | n.a. |
| TCF4 | 3.386 | 9.3454 | 1922.8 | 4E-255 | 1E-252 | 0.9279279 | n.a. |
| ATP13A2 | 3.848 | 9.2517 | 3425.56 | 1E-244 | 3E-242 | 0.8648649 | surface |
| POLD1 | 3.669 | 9.3555 | 1906.61 | 9E-236 | 3E-233 | 0.8108108 | n.a. |
| ENSECAG00000019875 | 3.123 | 9.2924 | 1943.28 | 1E-198 | 4E-196 | 0.8738739 | n.a. |
| EEF1B2 | 1.462 | 12.225 | 852.621 | 3E-185 | 8E-183 | 1 | n.a. |
| ENSECAG00000029287 | 3.432 | 10.043 | 846.754 | 6E-184 | 1E-181 | 0.8018018 | n.a. |
| ADORA2B | 3.622 | 9.2559 | 2601.48 | 2E-180 | 3E-178 | 0.8018018 | surface |
| CTSS | 2.15 | 9.8161 | 813.661 | 6E-177 | 1E-174 | 0.972973 | n.a. |
| RAC2 | 2.52 | 10.594 | 809.898 | 4E-176 | 9E-174 | 0.972973 | n.a. |
| ENSECAG00000038338 | 3.411 | 9.2755 | 2809.1 | 6E-173 | 1E-170 | 0.6576577 | n.a. |
| DUSP5 | 3.32 | 9.267 | 2007.28 | 2E-170 | 4E-168 | 0.6666667 | n.a. |
| NUCB2 | 3.326 | 9.3402 | 912.459 | 8E-163 | 2E-160 | 0.7387387 | n.a. |
| BLNK | 3.361 | 9.2737 | 2493.39 | 1E-160 | 2E-158 | 0.7657658 | n.a. |
| FYB1 | 3.089 | 9.6072 | 733.789 | 6E-160 | 1E-157 | 0.9369369 | n.a. |
| ENSECAG00000015055 | 3.35 | 9.2599 | 2239.33 | 5E-157 | 9E-155 | 0.5945946 | n.a. |
| SYNGR1 | 3.329 | 9.2464 | 3067.9 | 3E-154 | 5E-152 | 0.7297297 | n.a. |
| IFNAR1 | 3.006 | 9.352 | 695.047 | 1E-151 | 2E-149 | 0.8648649 | surface |
| CCDC50 | 2.941 | 9.3095 | 1001.54 | 3E-151 | 5E-149 | 0.7747748 | n.a. |
| IRF8 | 2.038 | 9.303 | 679.79 | 2E-148 | 3E-146 | 0.9459459 | n.a. |
| MPEG1 | 2.193 | 9.3534 | 1778.55 | 4E-147 | 6E-145 | 0.9009009 | surface |
| HEBP2 | 3.092 | 9.2913 | 1090.3 | 5E-137 | 7E-135 | 0.7027027 | n.a. |
| IL10RB | 2.816 | 9.6008 | 609.061 | 3E-133 | 4E-131 | 1 | surface |
| ENSECAG00000035361 | 3.355 | 9.2396 | 6397.93 | 9E-132 | 1E-129 | 0.7747748 | n.a. |
| MFAP2 | 3.261 | 9.2367 | 13601.5 | 4E-131 | 5E-129 | 0.4954955 | n.a. |
| ITM2C | 2.954 | 9.5994 | 588.471 | 6E-129 | 9E-127 | 0.8468468 | surface |
| SDHB | 2.663 | 9.4571 | 582.769 | 1E-127 | 1E-125 | 0.954955 | n.a. |
| ENSECAG00000010436 | 3.041 | 9.2883 | 886.861 | 2E-126 | 3E-124 | 0.7207207 | n.a. |
| ST6GALNAC4 | 2.801 | 9.2959 | 684.289 | 2E-125 | 3E-123 | 0.8378378 | nonsurface |
| ENSECAG00000021212 | 3.299 | 9.2394 | 4358.86 | 3E-124 | 3E-122 | 0.6126126 | n.a. |
| FCER1G | 1.375 | 10.204 | 654.465 | 4E-123 | 5E-121 | 0.981982 | nonsurface |
| GAPT | 2.897 | 9.2786 | 1469.65 | 9E-118 | 1E-115 | 0.5945946 | nonsurface |

| | | | | | | | |
|---|---|---|---|---|---|---|---|
| PPP2R3C | 2.825 | 9.2949 | 688.145 | 3E-115 | 4E-113 | 0.6576577 | n.a. |
| TNFSF13 | 3 | 9.266 | 3320.17 | 5E-115 | 5E-113 | 0.6936937 | nonsurface |
| CLEC12A | 1.829 | 9.4082 | 1295.34 | 3E-113 | 3E-111 | 0.9189189 | surface |
| ENSECAG00000031522 | 3.395 | 10.189 | 508.857 | 7E-112 | 8E-110 | 0.5765766 | n.a. |
| CNPY3 | 2.333 | 9.4881 | 500.649 | 4E-110 | 5E-108 | 1 | n.a. |
| YWHAQ | 2.645 | 9.7075 | 489.226 | 1E-107 | 1E-105 | 0.9279279 | n.a. |
| ENSECAG00000032253 | 2.836 | 9.2484 | 2099.18 | 7E-107 | 7E-105 | 0.5585586 | n.a. |
| ENSECAG00000031985 | 2.88 | 9.238 | 6403.93 | 1E-104 | 1E-102 | 0.5225225 | n.a. |
| CERS6 | 2.872 | 9.2532 | 1570.49 | 2E-102 | 2E-100 | 0.5855856 | nonsurface |
| DIPK2A | 2.751 | 9.2586 | 1100.03 | 4E-102 | 4E-100 | 0.7387387 | n.a. |
| SNX9 | 2.71 | 9.271 | 820.266 | 1E-100 | 1E-98 | 0.7387387 | n.a. |
| ENSECAG00000002390 | 2.988 | 9.2531 | 1881.99 | 3E-100 | 3E-98 | 0.5315315 | n.a. |
| PROCR | 2.75 | 9.236 | 9426.07 | 2E-98 | 2E-96 | 0.4234234 | surface |
| MCOLN2 | 2.817 | 9.2516 | 2142.08 | 2E-97 | 2E-95 | 0.6486486 | nonsurface |
| ACP5 | 2.574 | 9.7992 | 432.163 | 2E-95 | 2E-93 | 0.8918919 | n.a. |
| FAM129C | 2.741 | 9.2412 | 4361.59 | 8E-93 | 7E-91 | 0.5495495 | n.a. |
| SRM | 2.61 | 9.2691 | 779.997 | 5E-91 | 4E-89 | 0.5495495 | n.a. |
| MS4A1 | 2.52 | 9.7256 | 912.085 | 2E-90 | 2E-88 | 0.3963964 | surface |
| L3MBTL3 | 2.7 | 9.2638 | 935.427 | 3E-90 | 3E-88 | 0.6486486 | n.a. |
| FAM177A1 | 2.648 | 9.2748 | 748.004 | 5E-88 | 5E-86 | 0.5945946 | n.a. |
| GOT1 | 2.587 | 9.2894 | 611.468 | 3E-85 | 2E-83 | 0.6576577 | n.a. |
| CD164 | 2.537 | 9.4651 | 382.179 | 1E-84 | 1E-82 | 0.8738739 | surface |
| HM13 | 2.258 | 9.3911 | 379.219 | 5E-84 | 4E-82 | 0.8648649 | surface |
| FAIM | 2.581 | 9.2661 | 859.799 | 6E-84 | 5E-82 | 0.6036036 | surface |
| SLC44A2 | 2.534 | 9.4299 | 372.862 | 1E-82 | 1E-80 | 0.7027027 | surface |
| FAM107B | 2.611 | 9.5005 | 369.419 | 7E-82 | 5E-80 | 0.6936937 | n.a. |
| HSP90B1 | 2.097 | 9.5518 | 363.942 | 1E-80 | 8E-79 | 0.990991 | n.a. |
| FCMR | 2.372 | 9.3335 | 692.276 | 1E-80 | 9E-79 | 0.4234234 | n.a. |
| CD44 | 1.844 | 10.108 | 360.807 | 5E-80 | 4E-78 | 0.981982 | surface |
| BTG1 | 2.127 | 10.365 | 360.549 | 5E-80 | 4E-78 | 0.972973 | n.a. |
| PAXX | 2.372 | 9.3487 | 369.702 | 1E-76 | 1E-74 | 0.7027027 | n.a. |
| ENSECAG00000034000 | 1.924 | 9.5622 | 342.658 | 4E-76 | 3E-74 | 0.972973 | n.a. |
| PLAC8B | 1.568 | 11.113 | 341.342 | 8E-76 | 6E-74 | 0.990991 | n.a. |
| ENSECAG00000024850 | 2.426 | 9.2353 | 13797.8 | 4E-75 | 3E-73 | 0.4504505 | n.a. |
| AKAP9 | 2.321 | 9.4493 | 327.899 | 6E-73 | 4E-71 | 0.8468468 | n.a. |
| ENSECAG00000006595 | 2.428 | 9.2598 | 2104.16 | 2E-70 | 2E-68 | 0.4414414 | n.a. |
| RGS18 | 2.036 | 9.3114 | 522.52 | 5E-70 | 4E-68 | 0.7567568 | n.a. |
| GNG2 | 2.489 | 9.4802 | 337.914 | 7E-70 | 5E-68 | 0.5945946 | n.a. |
| EPCAM | 1.986 | 9.2602 | 1149.08 | 3E-69 | 2E-67 | 0.5585586 | surface |
| JAML | 2.066 | 9.299 | 596.594 | 4E-69 | 3E-67 | 0.7927928 | n.a. |
| SLC38A9 | 2.035 | 9.2699 | 411.24 | 8E-69 | 5E-67 | 0.7027027 | surface |
| NCF1 | 2.39 | 9.2933 | 1232.31 | 8E-68 | 5E-66 | 0.5945946 | n.a. |
| SLA2 | 2.423 | 9.2804 | 843.686 | 6E-67 | 4E-65 | 0.4324324 | n.a. |
| ATP1B1 | 2.083 | 9.2383 | 4283.16 | 1E-65 | 9E-64 | 0.3333333 | surface |
| PACSIN1 | 2.242 | 9.2564 | 2656.92 | 2E-65 | 1E-63 | 0.3873874 | n.a. |
| PRELID1 | 1.9 | 9.7488 | 293.578 | 2E-65 | 1E-63 | 0.981982 | n.a. |
| GINM1 | 2.225 | 9.3701 | 379.963 | 6E-65 | 3E-63 | 0.6666667 | surface |
| SMIM5 | 2.403 | 9.2402 | 2112.01 | 3E-64 | 2E-62 | 0.4324324 | nonsurface |

| | | | | | | | |
|---|---|---|---|---|---|---|---|
| CASP7 | 2.335 | 9.2925 | 533.72 | 1E-63 | 6E-62 | 0.4774775 | n.a. |
| RNASEH2B | 2.182 | 9.3711 | 323.258 | 8E-63 | 5E-61 | 0.7927928 | n.a. |
| ENSECAG00000035141 | 2.351 | 9.2632 | 1883.84 | 8E-63 | 5E-61 | 0.3963964 | n.a. |
| AQP3 | 2.256 | 9.3499 | 404.247 | 1E-62 | 7E-61 | 0.5495495 | n.a. |
| CD69 | 2.374 | 9.4382 | 392.077 | 2E-61 | 1E-59 | 0.4144144 | surface |
| CILP | 2.057 | 9.2349 | 11041.9 | 3E-61 | 1E-59 | 0.3063063 | n.a. |
| H2AFJ | 1.808 | 9.3551 | 340.119 | 3E-61 | 2E-59 | 0.8378378 | n.a. |
| ARRDC5 | 2.102 | 9.2365 | 5375.54 | 2E-60 | 1E-58 | 0.3063063 | n.a. |
| VLDLR | 2.163 | 9.2352 | 2097.8 | 1E-59 | 8E-58 | 0.3693694 | surface |
| IGHM | 2.107 | 9.6407 | 266.174 | 2E-59 | 1E-57 | 0.3873874 | n.a. |
| PDCD4 | 2.092 | 9.6109 | 265.29 | 2E-59 | 1E-57 | 0.9009009 | n.a. |
| ENSECAG00000038714 | 1.89 | 9.3099 | 484.16 | 4E-59 | 2E-57 | 0.7207207 | n.a. |
| DMXL2 | 2.223 | 9.2419 | 2620.48 | 5E-59 | 3E-57 | 0.3603604 | nonsurface |
| CALY | 2.227 | 9.2778 | 893.489 | 1E-58 | 7E-57 | 0.3963964 | surface |
| GRN | 1.469 | 9.4381 | 261.25 | 2E-58 | 8E-57 | 0.9279279 | n.a. |
| MGAT1 | 2.066 | 9.3346 | 282.886 | 5E-58 | 3E-56 | 0.7387387 | nonsurface |
| ENSECAG00000034985 | 2.274 | 9.3342 | 371.424 | 1E-57 | 5E-56 | 0.6126126 | n.a. |
| TFPT | 2.013 | 9.2805 | 345.221 | 1E-57 | 8E-56 | 0.7207207 | n.a. |
| B3GNT2 | 2.146 | 9.369 | 278.874 | 3E-57 | 2E-55 | 0.6036036 | nonsurface |
| UBC | 1.232 | 11.249 | 253.796 | 6E-57 | 3E-55 | 0.972973 | n.a. |
| LAMP3 | 2.052 | 9.2517 | 1109.27 | 2E-56 | 1E-54 | 0.4324324 | surface |
| GOLIM4 | 2.092 | 9.26 | 563.141 | 4E-56 | 2E-54 | 0.5945946 | nonsurface |
| TMED2 | 1.864 | 9.6321 | 247.272 | 2E-55 | 8E-54 | 0.9009009 | nonsurface |
| SPCS3 | 1.808 | 9.6267 | 246.467 | 2E-55 | 1E-53 | 0.963964 | nonsurface |
| LYST | 1.885 | 9.3883 | 241.176 | 3E-54 | 2E-52 | 0.7297297 | n.a. |
| SH3BP5 | 2.143 | 9.4429 | 542.984 | 7E-53 | 3E-51 | 0.6126126 | n.a. |
| UPF3B | 1.956 | 9.3746 | 232.116 | 3E-52 | 1E-50 | 0.7657658 | n.a. |
| ENSECAG00000006065 | 1.237 | 10.545 | 229.557 | 1E-51 | 5E-50 | 0.972973 | n.a. |
| MANF | 1.914 | 9.4674 | 228.733 | 2E-51 | 8E-50 | 0.8198198 | n.a. |
| ENSECAG00000031921 | 2.014 | 9.4417 | 228.527 | 2E-51 | 9E-50 | 0.6846847 | n.a. |
| PPP1R14A | 2.19 | 9.2419 | 2000.7 | 1E-50 | 5E-49 | 0.4594595 | n.a. |
| SEC61B | 1.487 | 9.8749 | 224.543 | 1E-50 | 6E-49 | 1 | nonsurface |
| CCS | 1.909 | 9.3726 | 224.106 | 2E-50 | 8E-49 | 0.7927928 | nonsurface |
| PRDX2 | 2.017 | 9.2964 | 758.197 | 1E-48 | 5E-47 | 0.2612613 | n.a. |
| RPS27L | 1.513 | 9.8828 | 215.599 | 1E-48 | 5E-47 | 1 | n.a. |
| RRBP1 | 1.812 | 9.2913 | 377.542 | 2E-48 | 8E-47 | 0.7027027 | nonsurface |
| ADGRG5 | 2.082 | 9.2595 | 907.224 | 3E-48 | 1E-46 | 0.5855856 | n.a. |
| DHCR7 | 2.123 | 9.2548 | 694.045 | 4E-48 | 2E-46 | 0.4234234 | nonsurface |
| TIGIT | 1.997 | 9.2485 | 1534.89 | 6E-48 | 2E-46 | 0.3783784 | surface |
| TCHP | 2.007 | 9.2494 | 819.798 | 3E-47 | 1E-45 | 0.4054054 | n.a. |
| MESD | 1.929 | 9.4258 | 204.722 | 3E-46 | 1E-44 | 0.6936937 | n.a. |
| CTSB | 1.465 | 9.4471 | 204.184 | 3E-46 | 1E-44 | 0.8378378 | n.a. |
| DNAJB9 | 2.12 | 9.2718 | 477.499 | 4E-46 | 2E-44 | 0.5225225 | n.a. |
| PTPRCAP | 1.45 | 10.617 | 203.679 | 4E-46 | 2E-44 | 0.972973 | nonsurface |
| SEPT11 | 1.938 | 9.2663 | 575.484 | 5E-46 | 2E-44 | 0.3603604 | n.a. |
| ENSECAG00000019029 | 1.681 | 9.9278 | 201.909 | 1E-45 | 4E-44 | 0.8828829 | n.a. |
| SRP14 | 1.246 | 10.144 | 198.741 | 5E-45 | 2E-43 | 0.990991 | n.a. |
| SPCS2 | 1.807 | 9.6657 | 198.573 | 6E-45 | 2E-43 | 0.8738739 | nonsurface |

| | | | | | | | |
|---|---|---|---|---|---|---|---|
| CPPED1 | 2.012 | 9.3236 | 305.879 | 7E-45 | 3E-43 | 0.5765766 | n.a. |
| NPHP4 | 1.73 | 9.246 | 2190.56 | 3E-44 | 1E-42 | 0.2882883 | n.a. |
| DGKZ | 2.074 | 9.2954 | 398.381 | 9E-44 | 4E-42 | 0.4594595 | n.a. |
| C1QTNF1 | 2.021 | 9.2395 | 1985.77 | 1E-43 | 5E-42 | 0.3423423 | n.a. |
| HCST | 1.798 | 9.5985 | 189.941 | 4E-43 | 2E-41 | 0.6396396 | n.a. |
| TMEM206 | 2.023 | 9.289 | 502.185 | 5E-43 | 2E-41 | 0.3783784 | n.a. |
| STX7 | 1.494 | 9.3568 | 186.387 | 3E-42 | 1E-40 | 0.7927928 | nonsurface |
| RASL11B | 1.774 | 9.2351 | 6803.99 | 3E-42 | 1E-40 | 0.2522523 | n.a. |
| STK17A | 1.858 | 9.6864 | 184.8 | 6E-42 | 2E-40 | 0.7927928 | n.a. |
| TEX30 | 1.995 | 9.4443 | 225.966 | 6E-42 | 2E-40 | 0.4774775 | n.a. |
| HIGD1A | 1.736 | 9.3918 | 182.795 | 2E-41 | 6E-40 | 0.6666667 | nonsurface |
| RNF5 | 1.738 | 9.3876 | 182.088 | 2E-41 | 8E-40 | 0.7927928 | nonsurface |
| BICDL2 | 1.88 | 9.2439 | 1319.25 | 4E-41 | 1E-39 | 0.3513514 | n.a. |
| TMED9 | 1.984 | 9.3567 | 239.164 | 4E-41 | 2E-39 | 0.4774775 | n.a. |
| NPC2 | 1.3 | 9.8367 | 178.408 | 1E-40 | 5E-39 | 0.981982 | n.a. |
| STX8 | 1.784 | 9.3397 | 204.686 | 2E-40 | 6E-39 | 0.6036036 | nonsurface |
| CYFIP2 | 2.092 | 9.3415 | 313.436 | 3E-40 | 1E-38 | 0.4054054 | n.a. |
| SEC61G | 1.343 | 9.9637 | 174.967 | 8E-40 | 3E-38 | 0.981982 | nonsurface |
| SRPRB | 1.9 | 9.3465 | 226.664 | 1E-39 | 4E-38 | 0.5225225 | nonsurface |
| CD2AP | 1.691 | 9.2797 | 247.068 | 2E-39 | 6E-38 | 0.5945946 | n.a. |
| TMCO1 | 1.791 | 9.4392 | 173.202 | 2E-39 | 7E-38 | 0.6666667 | nonsurface |
| BCL11A | 1.887 | 9.2593 | 630.513 | 4E-39 | 1E-37 | 0.4594595 | n.a. |
| LSS | 1.913 | 9.2458 | 992.51 | 5E-39 | 2E-37 | 0.4414414 | n.a. |
| SLC1A5 | 2.005 | 9.2879 | 403.493 | 7E-39 | 2E-37 | 0.4594595 | surface |
| SELENOW | 1.544 | 9.7827 | 170.566 | 7E-39 | 2E-37 | 0.9459459 | n.a. |
| SEC11C | 1.634 | 9.493 | 170.252 | 8E-39 | 3E-37 | 0.8108108 | nonsurface |
| ENSECAG00000035548 | 1.887 | 9.2356 | 4127.42 | 1E-38 | 5E-37 | 0.2702703 | n.a. |
| LGALS1 | 1.019 | 11.109 | 165.946 | 7E-38 | 2E-36 | 1 | nonsurface |
| AOAH | 1.75 | 9.3519 | 245.796 | 1E-36 | 4E-35 | 0.5765766 | n.a. |
| WWP1 | 1.957 | 9.2543 | 583.348 | 2E-36 | 6E-35 | 0.4864865 | n.a. |
| CXCR3 | 1.805 | 9.3103 | 504.655 | 4E-36 | 1E-34 | 0.3513514 | surface |
| RABAC1 | 1.434 | 9.8034 | 157.414 | 5E-36 | 2E-34 | 0.9099099 | nonsurface |
| TP53I13 | 1.815 | 9.2455 | 880.118 | 8E-36 | 3E-34 | 0.3693694 | surface |
| TMED5 | 1.734 | 9.3523 | 190.449 | 8E-36 | 3E-34 | 0.6126126 | nonsurface |
| HS3ST1 | 1.861 | 9.2857 | 946.733 | 2E-35 | 7E-34 | 0.3513514 | n.a. |
| RNF213 | 1.678 | 9.2995 | 205.187 | 2E-34 | 6E-33 | 0.6126126 | nonsurface |
| CYB5A | 1.847 | 9.3033 | 256.682 | 2E-34 | 6E-33 | 0.4504505 | nonsurface |
| SETX | 1.68 | 9.4027 | 149.377 | 3E-34 | 9E-33 | 0.6126126 | n.a. |
| RHEB | 1.501 | 9.4952 | 144.977 | 3E-33 | 8E-32 | 0.8198198 | n.a. |
| FAM81B | 1.629 | 9.2347 | 1834.37 | 7E-33 | 2E-31 | 0.2522523 | n.a. |
| BEX3 | 1.834 | 9.4183 | 174.14 | 8E-33 | 3E-31 | 0.5315315 | n.a. |
| TMEM60 | 1.681 | 9.4303 | 142.538 | 9E-33 | 3E-31 | 0.6306306 | n.a. |
| KAT14 | 1.787 | 9.2878 | 277.106 | 4E-32 | 1E-30 | 0.4234234 | n.a. |
| C1QBP | 1.278 | 9.3874 | 138.508 | 6E-32 | 2E-30 | 0.7837838 | n.a. |
| ME3 | 1.739 | 9.2599 | 421.338 | 7E-32 | 2E-30 | 0.4054054 | n.a. |
| CXCR4 | 1.717 | 9.6177 | 138.074 | 8E-32 | 2E-30 | 0.3873874 | surface |
| LAMP1 | 1.648 | 9.6692 | 177.662 | 3E-31 | 8E-30 | 0.5675676 | surface |
| PADI2 | 1.625 | 9.2434 | 1344.51 | 5E-31 | 1E-29 | 0.4324324 | n.a. |

| | | | | | | | |
|---|---|---|---|---|---|---|---|
| SELL | 1.521 | 9.6904 | 133.697 | 7E-31 | 2E-29 | 0.8108108 | surface |
| ADA2 | 1.857 | 9.2463 | 783.162 | 9E-31 | 2E-29 | 0.3423423 | n.a. |
| SEC11A | 1.549 | 9.5255 | 131.478 | 2E-30 | 6E-29 | 0.7837838 | nonsurface |
| TPST2 | 1.726 | 9.4709 | 144.146 | 3E-30 | 9E-29 | 0.5765766 | nonsurface |
| XBP1 | 1.558 | 9.4868 | 133.437 | 7E-30 | 2E-28 | 0.6846847 | n.a. |
| SSR3 | 1.151 | 9.9724 | 125.66 | 4E-29 | 1E-27 | 1 | nonsurface |
| CLK2 | 1.586 | 9.3097 | 176.103 | 2E-28 | 5E-27 | 0.4414414 | n.a. |
| RECQL5 | 1.698 | 9.2498 | 643.962 | 2E-28 | 6E-27 | 0.2882883 | n.a. |
| ENSECAG00000032188 | 1.201 | 9.5863 | 121.683 | 3E-28 | 8E-27 | 0.9459459 | n.a. |
| GSTA4 | 1.555 | 9.2357 | 3502.14 | 4E-28 | 1E-26 | 0.2612613 | n.a. |
| LBH | 1.582 | 9.4636 | 127.351 | 4E-28 | 1E-26 | 0.3153153 | n.a. |
| BET1 | 1.409 | 9.3715 | 119.486 | 9E-28 | 2E-26 | 0.6666667 | nonsurface |
| ARSB | 1.57 | 9.2518 | 828.679 | 1E-27 | 3E-26 | 0.2702703 | n.a. |
| C12H11orf74 | 1.54 | 9.3167 | 172.868 | 3E-27 | 7E-26 | 0.6216216 | n.a. |
| DYNLRB2 | 1.623 | 9.2488 | 660.861 | 3E-27 | 8E-26 | 0.2792793 | n.a. |
| TPP1 | 1.436 | 9.2798 | 191.969 | 3E-27 | 9E-26 | 0.6306306 | n.a. |
| CXHXorf21 | 1.501 | 9.2684 | 325.782 | 3E-27 | 9E-26 | 0.4594595 | n.a. |
| SNX30 | 1.61 | 9.2487 | 519.675 | 4E-27 | 9E-26 | 0.3963964 | n.a. |
| LMAN1 | 1.427 | 9.4114 | 116.519 | 4E-27 | 1E-25 | 0.6576577 | nonsurface |
| IRF2BP2 | 1.245 | 9.4142 | 116.298 | 5E-27 | 1E-25 | 0.8648649 | n.a. |
| SMARCAL1 | 1.559 | 9.2489 | 499.099 | 5E-27 | 1E-25 | 0.3423423 | n.a. |
| MAD1L1 | 1.644 | 9.2498 | 568.607 | 7E-27 | 2E-25 | 0.3513514 | n.a. |
| TRAM1 | 1.412 | 9.4843 | 115.104 | 8E-27 | 2E-25 | 0.8198198 | nonsurface |
| GPLD1 | 1.668 | 9.245 | 1062.44 | 9E-27 | 2E-25 | 0.3423423 | n.a. |
| SELENOS | 1.449 | 9.3395 | 163.487 | 2E-26 | 4E-25 | 0.5585586 | n.a. |
| CYBB | 1.371 | 9.4653 | 267.835 | 2E-26 | 5E-25 | 0.5405405 | surface |
| ITGAE | 1.622 | 9.3387 | 188.987 | 3E-26 | 7E-25 | 0.3693694 | surface |
| AP3S1 | 1.377 | 9.534 | 111.445 | 5E-26 | 1E-24 | 0.8648649 | n.a. |
| CCDC12 | 1.398 | 9.3727 | 110.089 | 1E-25 | 2E-24 | 0.6126126 | nonsurface |
| CBX5 | 1.615 | 9.3148 | 188.071 | 1E-25 | 3E-24 | 0.4414414 | n.a. |
| SLC25A4 | 1.592 | 9.2755 | 280.015 | 1E-25 | 3E-24 | 0.4234234 | nonsurface |
| ENSECAG00000000775 | 1.143 | 9.4621 | 108.99 | 2E-25 | 4E-24 | 0.8558559 | n.a. |
| ENSECAG00000003617 | 1.421 | 9.398 | 108.943 | 2E-25 | 4E-24 | 0.6216216 | n.a. |
| SNRPN | 1.476 | 9.4022 | 116.202 | 2E-25 | 5E-24 | 0.5405405 | n.a. |
| POLB | 1.403 | 9.3404 | 119.455 | 3E-25 | 6E-24 | 0.5765766 | n.a. |
| HHEX | 1.239 | 9.4401 | 107.824 | 3E-25 | 7E-24 | 0.7117117 | n.a. |
| TOB1 | 1.56 | 9.3276 | 166.503 | 3E-25 | 8E-24 | 0.4594595 | n.a. |
| STMN1 | 1.547 | 9.7683 | 107.643 | 4E-25 | 9E-24 | 0.4414414 | n.a. |
| CD3G | 1.566 | 10.358 | 126.718 | 4E-25 | 1E-23 | 0.4144144 | surface |
| RF00581 | 1.066 | 10.319 | 106.924 | 5E-25 | 1E-23 | 0.972973 | n.a. |
| H2AFZ | 1.379 | 9.5191 | 105.566 | 1E-24 | 2E-23 | 0.6846847 | n.a. |
| MITD1 | 1.563 | 9.3423 | 153.105 | 1E-24 | 3E-23 | 0.3783784 | n.a. |
| MRPL34 | 1.461 | 9.4086 | 114.606 | 1E-24 | 3E-23 | 0.6306306 | n.a. |
| SULF2 | 1.572 | 9.2383 | 1589.43 | 3E-24 | 6E-23 | 0.3333333 | n.a. |
| CD1B1 | 1.261 | 9.2551 | 751.198 | 6E-24 | 1E-22 | 0.3513514 | n.a. |
| HMGB1 | 1.047 | 10.086 | 101.858 | 6E-24 | 1E-22 | 0.954955 | n.a. |
| CXXC5 | 1.506 | 9.3065 | 221.407 | 9E-24 | 2E-22 | 0.3243243 | n.a. |
| APBB1IP | 1.253 | 9.6791 | 100.674 | 1E-23 | 2E-22 | 0.8648649 | n.a. |

| | | | | | | | |
|---|---|---|---|---|---|---|---|
| STAMBPL1 | 1.573 | 9.3697 | 150.463 | 1E-23 | 3E-22 | 0.3603604 | n.a. |
| SEC61A1 | 1.483 | 9.3336 | 145.332 | 2E-23 | 4E-22 | 0.5495495 | nonsurface |
| URI1 | 1.531 | 9.2643 | 254.235 | 2E-23 | 4E-22 | 0.3513514 | n.a. |
| TPR | 1.354 | 9.453 | 99.0158 | 3E-23 | 6E-22 | 0.7117117 | surface |
| YPEL3 | 1.462 | 9.6613 | 98.7153 | 3E-23 | 7E-22 | 0.5945946 | n.a. |
| ENSECAG00000031496 | 1.238 | 9.3827 | 98.0957 | 4E-23 | 9E-22 | 0.8108108 | n.a. |
| SRPRA | 1.352 | 9.3887 | 98.0354 | 4E-23 | 9E-22 | 0.5855856 | n.a. |
| ZFYVE16 | 1.543 | 9.2508 | 438.551 | 5E-23 | 1E-21 | 0.2882883 | n.a. |
| CHRNB1 | 1.501 | 9.2427 | 840.779 | 8E-23 | 2E-21 | 0.2612613 | surface |
| SDF2L1 | 1.22 | 9.3874 | 97.3042 | 1E-22 | 3E-21 | 0.7387387 | n.a. |
| DOCK8 | 1.281 | 9.3857 | 95.6656 | 1E-22 | 3E-21 | 0.7117117 | n.a. |
| MTF2 | 1.417 | 9.3258 | 123.447 | 2E-22 | 4E-21 | 0.3783784 | n.a. |
| NUDT3 | 1.533 | 9.2715 | 294.243 | 3E-22 | 5E-21 | 0.3063063 | n.a. |
| ABRAXAS1 | 1.568 | 9.2896 | 212.585 | 7E-22 | 1E-20 | 0.3693694 | n.a. |
| APOBEC3H | 1.509 | 9.3557 | 133.937 | 7E-22 | 1E-20 | 0.4054054 | n.a. |
| ALOX5AP | 1.517 | 9.2637 | 775.873 | 7E-22 | 1E-20 | 0.2522523 | nonsurface |
| TLR7 | 1.37 | 9.2655 | 631.082 | 1E-21 | 3E-20 | 0.2972973 | surface |
| ENSECAG00000038199 | 1.426 | 9.2818 | 407.728 | 2E-21 | 3E-20 | 0.3153153 | n.a. |
| GPR183 | 1.471 | 9.6621 | 90.1599 | 2E-21 | 4E-20 | 0.3963964 | surface |
| ALG5 | 1.322 | 9.3503 | 104.53 | 3E-21 | 7E-20 | 0.5495495 | nonsurface |
| ASB8 | 1.349 | 9.2901 | 139.654 | 5E-21 | 9E-20 | 0.4144144 | n.a. |
| TSC22D1 | 1.441 | 9.2501 | 357.324 | 1E-20 | 2E-19 | 0.3963964 | n.a. |
| YIPF1 | 1.607 | 9.287 | 229.251 | 2E-20 | 3E-19 | 0.3603604 | n.a. |
| ZDHHC24 | 1.366 | 9.314 | 125.971 | 2E-20 | 3E-19 | 0.5225225 | nonsurface |
| ENSECAG00000037332 | 1.063 | 10.054 | 85.6718 | 2E-20 | 4E-19 | 0.954955 | n.a. |
| GTF3A | 1.301 | 9.6075 | 85.5822 | 2E-20 | 4E-19 | 0.7747748 | n.a. |
| MOCS2 | 1.398 | 9.3218 | 123.467 | 3E-20 | 5E-19 | 0.4234234 | n.a. |
| MED13L | 1.348 | 9.2911 | 142.183 | 3E-20 | 6E-19 | 0.4504505 | n.a. |
| ARMCX3 | 1.337 | 9.3437 | 101.466 | 4E-20 | 7E-19 | 0.4234234 | nonsurface |
| FAF1 | 1.495 | 9.2455 | 574.836 | 4E-20 | 8E-19 | 0.2972973 | n.a. |
| IGFLR1 | 1.249 | 9.4383 | 101.167 | 4E-20 | 8E-19 | 0.5135135 | surface |
| UVRAG | 1.358 | 9.294 | 137.87 | 6E-20 | 1E-18 | 0.4144144 | n.a. |
| RHOH | 1.325 | 9.4176 | 85.4804 | 8E-20 | 1E-18 | 0.4594595 | n.a. |
| SPPL2A | 1.397 | 9.2923 | 152.643 | 8E-20 | 1E-18 | 0.4324324 | surface |
| PTBP3 | 1.251 | 9.4001 | 83.0757 | 8E-20 | 1E-18 | 0.5585586 | n.a. |
| CTSA | 1.376 | 9.2925 | 198.034 | 9E-20 | 2E-18 | 0.2702703 | n.a. |
| EIF4E1B | 1.397 | 9.2561 | 555.414 | 1E-19 | 2E-18 | 0.4594595 | n.a. |
| RASGRP2 | 1.337 | 9.4643 | 82.2902 | 1E-19 | 2E-18 | 0.4864865 | n.a. |
| RNF13 | 1.278 | 9.3579 | 91.5832 | 1E-19 | 2E-18 | 0.4594595 | surface |
| NEU1 | 1.345 | 9.2661 | 240.689 | 1E-19 | 2E-18 | 0.2612613 | n.a. |
| TTYH1 | 1.426 | 9.2351 | 1167.52 | 1E-19 | 2E-18 | 0.2702703 | surface |
| ENSECAG00000040126 | 1.395 | 9.2742 | 192.9 | 2E-19 | 3E-18 | 0.3243243 | n.a. |
| CHMP6 | 1.26 | 9.3694 | 87.2236 | 3E-19 | 4E-18 | 0.5225225 | n.a. |
| SRP19 | 1.244 | 9.5262 | 80.7827 | 3E-19 | 5E-18 | 0.7387387 | n.a. |
| PLBD2 | 1.205 | 9.3161 | 122.437 | 3E-19 | 5E-18 | 0.6936937 | n.a. |
| GBGT1 | 1.347 | 9.2944 | 152.145 | 3E-19 | 5E-18 | 0.3873874 | nonsurface |
| SUMO1 | 1.223 | 9.561 | 80.0345 | 4E-19 | 7E-18 | 0.7117117 | n.a. |
| DNAJC3 | 1.19 | 9.3287 | 96.4539 | 5E-19 | 8E-18 | 0.5495495 | nonsurface |

| | | | | | | | |
|---|---|---|---|---|---|---|---|
| SLC4A7 | 1.199 | 9.3287 | 86.7931 | 5E-19 | 9E-18 | 0.4504505 | surface |
| PARP1 | 1.446 | 9.2948 | 195.01 | 5E-19 | 9E-18 | 0.3153153 | nonsurface |
| LEPROTL1 | 1.303 | 9.565 | 79.3899 | 5E-19 | 9E-18 | 0.6486486 | nonsurface |
| HSP90AA1 | 1.075 | 9.9098 | 78.7599 | 7E-19 | 1E-17 | 0.8648649 | n.a. |
| CD53 | 1.236 | 9.709 | 78.7267 | 7E-19 | 1E-17 | 0.8108108 | surface |
| AGA | 1.157 | 9.3481 | 91.9178 | 8E-19 | 1E-17 | 0.5405405 | n.a. |
| ENSECAG00000013303 | 1.001 | 9.4002 | 78.6547 | 8E-19 | 1E-17 | 0.9009009 | n.a. |
| VPS37B | 1.387 | 9.2968 | 198.874 | 8E-19 | 1E-17 | 0.2792793 | n.a. |
| ENSECAG00000016754 | 1.399 | 9.2778 | 177.988 | 9E-19 | 1E-17 | 0.4234234 | n.a. |
| SEC31A | 1.335 | 9.2996 | 133.348 | 1E-18 | 2E-17 | 0.3873874 | n.a. |
| GRID2IP | 1.235 | 9.4806 | 78.0816 | 1E-18 | 2E-17 | 0.6126126 | n.a. |
| SSR1 | 1.351 | 9.3182 | 121.743 | 1E-18 | 2E-17 | 0.4234234 | surface |
| TMEM41B | 1.398 | 9.2955 | 149.967 | 1E-18 | 2E-17 | 0.3783784 | nonsurface |
| ND3 | 1.019 | 9.8247 | 77.2477 | 2E-18 | 3E-17 | 0.9099099 | n.a. |
| IGSF8 | 1.412 | 9.2908 | 165.519 | 2E-18 | 3E-17 | 0.3603604 | surface |
| OSTC | 1.154 | 9.5663 | 76.7828 | 2E-18 | 3E-17 | 0.8108108 | nonsurface |
| S1PR4 | 1.397 | 9.4734 | 91.3039 | 2E-18 | 3E-17 | 0.3333333 | surface |
| RF00163 | 1.37 | 9.4218 | 88.5156 | 3E-18 | 5E-17 | 0.4054054 | n.a. |
| TBC1D1 | 1.35 | 9.2859 | 161.643 | 3E-18 | 6E-17 | 0.4504505 | nonsurface |
| ST3GAL4 | 1.373 | 9.3052 | 134.51 | 4E-18 | 7E-17 | 0.4054054 | nonsurface |
| EMC10 | 1.192 | 9.3664 | 75.452 | 4E-18 | 7E-17 | 0.5135135 | n.a. |
| TM9SF2 | 1.282 | 9.362 | 90.9209 | 5E-18 | 8E-17 | 0.4414414 | surface |
| ARL1 | 1.241 | 9.4084 | 77.9104 | 5E-18 | 9E-17 | 0.6216216 | nonsurface |
| BRK1 | 1.052 | 9.6928 | 74.6559 | 6E-18 | 9E-17 | 0.9189189 | n.a. |
| USO1 | 1.331 | 9.2993 | 131.22 | 9E-18 | 1E-16 | 0.3873874 | n.a. |
| IL7R | 1.317 | 9.3731 | 255.578 | 1E-17 | 1E-16 | 0.4054054 | surface |
| DKC1 | 1.24 | 9.411 | 75.8976 | 1E-17 | 2E-16 | 0.3873874 | n.a. |
| EBAG9 | 1.16 | 9.3732 | 73.6234 | 1E-17 | 2E-16 | 0.5495495 | nonsurface |
| PDIA6 | 1.186 | 9.332 | 92.0036 | 1E-17 | 2E-16 | 0.5225225 | n.a. |
| CHID1 | 1.32 | 9.3333 | 119.54 | 1E-17 | 2E-16 | 0.4594595 | n.a. |
| ACTN4 | 1.421 | 9.3064 | 146.676 | 1E-17 | 2E-16 | 0.4324324 | n.a. |
| CIRBP | 1.33 | 9.4608 | 74.9285 | 1E-17 | 2E-16 | 0.4324324 | n.a. |
| MTMR14 | 1.259 | 9.3346 | 101.17 | 1E-17 | 2E-16 | 0.4774775 | n.a. |
| RPN2 | 1.151 | 9.4695 | 72.6198 | 2E-17 | 2E-16 | 0.7027027 | nonsurface |
| PPA2 | 1.409 | 9.2617 | 217.009 | 2E-17 | 3E-16 | 0.3783784 | n.a. |
| TTF1 | 1.328 | 9.2977 | 144.694 | 2E-17 | 4E-16 | 0.3333333 | n.a. |
| ECHS1 | 1.169 | 9.393 | 75.4492 | 3E-17 | 4E-16 | 0.6306306 | n.a. |
| SMIM26 | 1.175 | 9.4437 | 71.125 | 3E-17 | 5E-16 | 0.7027027 | n.a. |
| KCTD10 | 1.412 | 9.3 | 173.11 | 3E-17 | 5E-16 | 0.2522523 | n.a. |
| RNF149 | 1.25 | 9.3444 | 113.786 | 4E-17 | 5E-16 | 0.4954955 | surface |
| MICAL1 | 1.286 | 9.2809 | 149.74 | 4E-17 | 7E-16 | 0.3423423 | n.a. |
| ENSECAG00000031485 | 1.334 | 9.2912 | 134.697 | 5E-17 | 7E-16 | 0.4684685 | n.a. |
| RAB2B | 1.291 | 9.269 | 165.865 | 5E-17 | 7E-16 | 0.3603604 | n.a. |
| DPP9 | 1.232 | 9.3491 | 90.9299 | 5E-17 | 8E-16 | 0.3693694 | n.a. |
| ITCH | 1.238 | 9.272 | 148.816 | 5E-17 | 8E-16 | 0.3423423 | n.a. |
| RPAIN | 1.325 | 9.2532 | 300.652 | 6E-17 | 8E-16 | 0.2882883 | n.a. |
| LAT | 1.416 | 9.5366 | 95.803 | 7E-17 | 1E-15 | 0.3513514 | nonsurface |
| PPT2-EGFL8 | 1.167 | 9.2729 | 196.039 | 1E-16 | 2E-15 | 0.2972973 | n.a. |

| | | | | | | | |
|---|---|---|---|---|---|---|---|
| SAP30BP | 1.351 | 9.2862 | 154.424 | 1E-16 | 2E-15 | 0.2972973 | n.a. |
| EDEM2 | 1.288 | 9.308 | 115.636 | 2E-16 | 3E-15 | 0.4414414 | n.a. |
| MILR1 | 1.239 | 9.2539 | 345.806 | 3E-16 | 4E-15 | 0.3873874 | surface |
| PRXL2A | 1.276 | 9.2641 | 218.675 | 4E-16 | 6E-15 | 0.2612613 | n.a. |
| ARL5A | 1.12 | 9.344 | 97.7921 | 6E-16 | 8E-15 | 0.3513514 | n.a. |
| COPZ1 | 1.206 | 9.391 | 76.4536 | 8E-16 | 1E-14 | 0.5675676 | n.a. |
| NHP2 | 1.077 | 9.4198 | 64.4737 | 1E-15 | 1E-14 | 0.6846847 | n.a. |
| SORL1 | 1.196 | 9.2707 | 262.875 | 1E-15 | 2E-14 | 0.2612613 | surface |
| FLII | 1.255 | 9.3044 | 118.647 | 2E-15 | 2E-14 | 0.4144144 | n.a. |
| SDF2 | 1.268 | 9.387 | 77.2181 | 2E-15 | 2E-14 | 0.4504505 | n.a. |
| CACYBP | 1.141 | 9.6062 | 63.0749 | 2E-15 | 3E-14 | 0.7387387 | n.a. |
| FAM117A | 1.284 | 9.3289 | 101.21 | 2E-15 | 3E-14 | 0.3783784 | n.a. |
| TOMM7 | 1.184 | 9.4539 | 68.4713 | 3E-15 | 4E-14 | 0.5585586 | nonsurface |
| GLG1 | 1.154 | 9.3396 | 77.1866 | 3E-15 | 5E-14 | 0.4414414 | n.a. |
| SURF4 | 1.166 | 9.3205 | 91.1825 | 4E-15 | 5E-14 | 0.3873874 | nonsurface |
| LMAN2 | 1.117 | 9.4812 | 61.5731 | 4E-15 | 6E-14 | 0.6306306 | surface |
| RNF10 | 1.155 | 9.3557 | 76.0742 | 5E-15 | 6E-14 | 0.5135135 | nonsurface |
| PTPN1 | 1.276 | 9.278 | 178.258 | 5E-15 | 6E-14 | 0.2612613 | nonsurface |
| CDCA4 | 1.219 | 9.2785 | 148.822 | 5E-15 | 6E-14 | 0.2882883 | n.a. |
| LRMP | 1.054 | 9.3198 | 84.0855 | 6E-15 | 8E-14 | 0.3513514 | nonsurface |
| RF00276 | 1.094 | 9.4503 | 60.7829 | 7E-15 | 8E-14 | 0.6486486 | n.a. |
| TMEM165 | 1.06 | 9.4054 | 64.4894 | 7E-15 | 9E-14 | 0.6216216 | nonsurface |
| CYB5R1 | 1.281 | 9.253 | 271.902 | 7E-15 | 9E-14 | 0.2702703 | n.a. |
| LSM4 | 1.055 | 9.4444 | 60.5802 | 7E-15 | 9E-14 | 0.6486486 | n.a. |
| FBXW2 | 1.155 | 9.3881 | 66.3363 | 7E-15 | 1E-13 | 0.3963964 | n.a. |
| ADAM17 | 1.247 | 9.2788 | 153.874 | 1E-14 | 1E-13 | 0.2522523 | surface |
| SNRPB2 | 1.154 | 9.5023 | 59.56 | 1E-14 | 2E-13 | 0.6036036 | n.a. |
| ZNF644 | 1.228 | 9.3066 | 105.755 | 1E-14 | 2E-13 | 0.2882883 | n.a. |
| FAM133B | 1.15 | 9.3623 | 68.8582 | 1E-14 | 2E-13 | 0.3333333 | n.a. |
| RF00280 | 1.104 | 9.4118 | 68.2008 | 1E-14 | 2E-13 | 0.4594595 | n.a. |
| ATG12 | 1.243 | 9.2965 | 110.249 | 2E-14 | 2E-13 | 0.4144144 | n.a. |
| RWDD4 | 1.258 | 9.3262 | 96.4643 | 2E-14 | 2E-13 | 0.4054054 | n.a. |
| MCM7 | 1.215 | 9.3416 | 92.414 | 2E-14 | 2E-13 | 0.2882883 | n.a. |
| CMTM3 | 1.099 | 9.4968 | 62.4928 | 2E-14 | 2E-13 | 0.6126126 | nonsurface |
| EID1 | 1.139 | 9.4085 | 62.6957 | 2E-14 | 3E-13 | 0.4504505 | n.a. |
| COPB1 | 1.266 | 9.3293 | 97.6414 | 2E-14 | 3E-13 | 0.4234234 | n.a. |
| ENSECAG00000038299 | 1.078 | 9.3304 | 70.8074 | 2E-14 | 3E-13 | 0.5225225 | n.a. |
| SEMA4D | 1.182 | 9.315 | 108.801 | 3E-14 | 3E-13 | 0.2882883 | surface |
| DNASE2 | 1.133 | 9.347 | 114.381 | 3E-14 | 3E-13 | 0.6126126 | n.a. |
| PRPSAP2 | 1.245 | 9.3039 | 110.462 | 3E-14 | 3E-13 | 0.3423423 | n.a. |
| MCFD2 | 1.228 | 9.2819 | 140.444 | 3E-14 | 3E-13 | 0.3423423 | n.a. |
| NCSTN | 1.182 | 9.302 | 100.116 | 3E-14 | 4E-13 | 0.4234234 | surface |
| ENSECAG00000036779 | 1.094 | 9.5478 | 57.4225 | 4E-14 | 4E-13 | 0.6846847 | n.a. |
| UBXN4 | 1.033 | 9.4336 | 56.3845 | 6E-14 | 7E-13 | 0.6216216 | n.a. |
| ENSECAG00000040274 | 1.029 | 9.269 | 115.25 | 7E-14 | 9E-13 | 0.4324324 | n.a. |
| ORMDL2 | 1.15 | 9.3745 | 66.4155 | 8E-14 | 9E-13 | 0.4414414 | nonsurface |
| CYB5R4 | 1.214 | 9.3322 | 91.238 | 8E-14 | 9E-13 | 0.3963964 | n.a. |
| CAT | 1.053 | 9.3771 | 65.3601 | 1E-13 | 1E-12 | 0.5405405 | n.a. |

| | | | | | | | |
|---|---|---|---|---|---|---|---|
| ZNF789 | 1.242 | 9.2679 | 179.982 | 1E-13 | 1E-12 | 0.2702703 | n.a. |
| TTC14 | 1.061 | 9.3422 | 64.5903 | 1E-13 | 1E-12 | 0.3423423 | n.a. |
| ENSECAG00000026829 | 1.058 | 9.535 | 55.1146 | 1E-13 | 1E-12 | 0.6666667 | n.a. |
| CSGALNACT2 | 1.168 | 9.2796 | 116.445 | 1E-13 | 1E-12 | 0.3693694 | nonsurface |
| UCP3 | 1.08 | 9.2375 | 816.156 | 1E-13 | 1E-12 | 0.2522523 | n.a. |
| NIPA2 | 1.004 | 9.3276 | 64.216 | 2E-13 | 2E-12 | 0.5315315 | nonsurface |
| ENSECAG00000019052 | 1.174 | 9.4644 | 62.241 | 2E-13 | 2E-12 | 0.3153153 | n.a. |
| CYBC1 | 1.063 | 9.3514 | 62.5586 | 2E-13 | 2E-12 | 0.4954955 | n.a. |
| FKBP2 | 1.053 | 9.3386 | 75.7736 | 2E-13 | 2E-12 | 0.5225225 | n.a. |
| C16orf87 | 1.235 | 9.2689 | 170.182 | 2E-13 | 3E-12 | 0.2792793 | n.a. |
| LGALS3BP | 1.053 | 9.4204 | 55.5094 | 2E-13 | 3E-12 | 0.5315315 | n.a. |
| GLTP | 1.166 | 9.2919 | 99.9894 | 3E-13 | 3E-12 | 0.3603604 | n.a. |
| SDF4 | 1.218 | 9.343 | 83.0399 | 3E-13 | 3E-12 | 0.3783784 | n.a. |
| GTF2F1 | 1.142 | 9.3478 | 73.6504 | 3E-13 | 3E-12 | 0.4594595 | n.a. |
| ERLEC1 | 1.074 | 9.3911 | 61.3081 | 3E-13 | 3E-12 | 0.6036036 | n.a. |
| GET4 | 1.134 | 9.3366 | 75.5804 | 3E-13 | 4E-12 | 0.2972973 | n.a. |
| PRRC2C | 1.041 | 9.5988 | 52.8652 | 4E-13 | 4E-12 | 0.6756757 | n.a. |
| EMD | 1.055 | 9.5059 | 52.6847 | 4E-13 | 4E-12 | 0.5945946 | nonsurface |
| STT3A | 1.123 | 9.2923 | 94.8998 | 5E-13 | 5E-12 | 0.2972973 | n.a. |
| MRPL36 | 1.03 | 9.4047 | 53.7883 | 5E-13 | 5E-12 | 0.5675676 | n.a. |
| RF00578 | 1.111 | 9.2683 | 166.759 | 6E-13 | 6E-12 | 0.2792793 | n.a. |
| UBE2S | 1.133 | 9.3494 | 74.6968 | 7E-13 | 8E-12 | 0.3693694 | n.a. |
| CRELD2 | 1.077 | 9.2876 | 110.202 | 9E-13 | 1E-11 | 0.3513514 | n.a. |
| NEDD1 | 1.06 | 9.2856 | 89.024 | 9E-13 | 1E-11 | 0.3423423 | n.a. |
| MPPE1 | 1.275 | 9.2717 | 168.754 | 1E-12 | 1E-11 | 0.2792793 | nonsurface |
| AFF3 | 1.013 | 9.2652 | 117.863 | 2E-12 | 2E-11 | 0.2612613 | n.a. |
| ENSECAG00000012199 | 1.014 | 9.3154 | 186.004 | 2E-12 | 2E-11 | 0.2882883 | n.a. |
| ADD1 | 1.086 | 9.3658 | 60.6601 | 2E-12 | 2E-11 | 0.4144144 | n.a. |
| CUTA | 1.112 | 9.3533 | 68.3255 | 2E-12 | 3E-11 | 0.4234234 | n.a. |
| SH3KBP1 | 1.119 | 9.4466 | 54.2995 | 3E-12 | 3E-11 | 0.3693694 | n.a. |
| HDAC2 | 1.191 | 9.2975 | 107.189 | 3E-12 | 4E-11 | 0.3513514 | n.a. |
| FAM45A | 1.104 | 9.3673 | 68.5373 | 4E-12 | 4E-11 | 0.2792793 | n.a. |
| DECR1 | 1.075 | 9.334 | 67.7202 | 4E-12 | 4E-11 | 0.4234234 | n.a. |
| MAP4K1 | 1.014 | 9.3262 | 65.0455 | 7E-12 | 8E-11 | 0.4504505 | n.a. |
| CLEC2B | 1.042 | 9.382 | 82.4257 | 8E-12 | 8E-11 | 0.2792793 | n.a. |
| CLPTM1 | 1.024 | 9.302 | 77.3999 | 9E-12 | 9E-11 | 0.2792793 | nonsurface |
| ENSECAG00000032776 | 1.025 | 9.4539 | 45.9189 | 1E-11 | 1E-10 | 0.3243243 | n.a. |
| PGAM5 | 1.018 | 9.3326 | 61.5412 | 1E-11 | 1E-10 | 0.4144144 | n.a. |
| SIRT7 | 1.023 | 9.3196 | 66.5398 | 2E-11 | 2E-10 | 0.4324324 | n.a. |
| SLC2A8 | 1.084 | 9.2576 | 169.135 | 2E-11 | 2E-10 | 0.2792793 | surface |
| ENSECAG00000030463 | 1.084 | 9.2585 | 175.198 | 2E-11 | 2E-10 | 0.2522523 | n.a. |
| IMPA1 | 1.027 | 9.2927 | 81.1142 | 3E-11 | 3E-10 | 0.2882883 | n.a. |
| TUBA4A | 1.103 | 9.5479 | 48.8647 | 3E-11 | 3E-10 | 0.3333333 | n.a. |
| JARID2 | 1.045 | 9.2792 | 97.1236 | 3E-11 | 3E-10 | 0.3243243 | n.a. |
| SYVN1 | 1.124 | 9.3092 | 118.192 | 3E-11 | 3E-10 | 0.2522523 | nonsurface |
| CARHSP1 | 1.031 | 9.4061 | 53.5885 | 5E-11 | 5E-10 | 0.3333333 | n.a. |
| ENSECAG00000037376 | 1.074 | 9.2452 | 190.306 | 5E-11 | 5E-10 | 0.3063063 | n.a. |
| ENSECAG00000007532 | 1.165 | 9.3313 | 79.882 | 1E-10 | 1E-09 | 0.3783784 | n.a. |

| | | | | | | | |
|---|---|---|---|---|---|---|---|
| CERS5 | 1.052 | 9.2999 | 77.8608 | 1E-10 | 1E-09 | 0.3963964 | nonsurface |
| AMPD3 | 1.111 | 9.2518 | 272.062 | 1E-10 | 1E-09 | 0.3063063 | n.a. |
| INTS12 | 1.016 | 9.3106 | 65.7569 | 1E-10 | 1E-09 | 0.3513514 | n.a. |
| RABL6 | 1.086 | 9.3067 | 80.5453 | 2E-10 | 2E-09 | 0.2972973 | n.a. |
| ATP6V0A1 | 1.081 | 9.2697 | 129.045 | 3E-10 | 2E-09 | 0.2612613 | n.a. |
| USP24 | 1.016 | 9.2996 | 74.5585 | 3E-10 | 2E-09 | 0.2702703 | n.a. |
| INPP4A | 1.039 | 9.3149 | 71.2928 | 3E-10 | 2E-09 | 0.2792793 | n.a. |
| UTP18 | 1.038 | 9.318 | 68.2491 | 4E-10 | 3E-09 | 0.3513514 | n.a. |
| PLIN3 | 1.047 | 9.2703 | 120.58 | 5E-10 | 5E-09 | 0.2882883 | n.a. |
| VDAC3 | 1.005 | 9.3496 | 56.0552 | 7E-10 | 6E-09 | 0.3153153 | n.a. |
| CRBN | 1.012 | 9.3164 | 67.5163 | 9E-10 | 8E-09 | 0.2612613 | n.a. |
| MTIF3 | 1.053 | 9.2963 | 84.4801 | 1E-09 | 1E-08 | 0.2612613 | n.a. |
| EMC9 | 1.084 | 9.2686 | 136.782 | 2E-09 | 2E-08 | 0.3063063 | n.a. |
| POC1A | 1.063 | 9.2936 | 106.421 | 2E-08 | 1E-07 | 0.2702703 | n.a. |
