## Supplementary material for "Single cell resolution landscape of equine peripheral blood mononuclear cells reveals diverse immune cell subtypes including T-bet^+^ B cells": Dataset S5

| genes | logFC | logCPM | F | PValue | FDR | percent.exp | Surfacome.Label |
|---|---|---|---|---|---|---|---|
| ENSECAG00000014585 | 3.61 | 10.271 | 3938.4 | 0 | 0 | 0.9881782 | n.a. |
| PLAC8B | 2.869 | 11.113 | 2235.4 | 0 | 0 | 0.9957563 | n.a. |
| IGHG3 | 2.763 | 9.6047 | 3171 | 0 | 0 | 0.930585 | n.a. |
| ENSECAG00000027826 | 1.357 | 9.7443 | 1164.6 | 4E-251 | 3E-248 | 0.9536223 | n.a. |
| MS4A1 | 1.131 | 9.7256 | 947.15 | 3E-205 | 2E-202 | 0.9439224 | surface |
| ENSECAG00000001010 | 1.288 | 9.629 | 921.86 | 7E-200 | 4E-197 | 0.6095787 | n.a. |
| HSF2BP | 1.598 | 9.3751 | 1059.6 | 4E-198 | 2E-195 | 0.6095787 | n.a. |
| SH3BP5 | 1.562 | 9.4429 | 849.32 | 2E-184 | 9E-182 | 0.7356775 | n.a. |
| ENSECAG00000029287 | 1.652 | 10.043 | 775.74 | 7E-169 | 3E-166 | 0.5159139 | n.a. |
| CMTM3 | 2.069 | 9.4968 | 735.75 | 2E-160 | 1E-157 | 0.6041225 | n.a. |
| BANK1 | 1.457 | 9.413 | 784.2 | 4E-160 | 2E-157 | 0.6535314 | n.a. |
| ANKRD2 | 1.608 | 9.3125 | 1435.3 | 2E-150 | 7E-148 | 0.4319491 | n.a. |
| PTPRCAP | 1.296 | 10.617 | 659.2 | 5E-144 | 2E-141 | 0.9839345 | n.a. |
| AHNAK | 1.716 | 9.5205 | 642.43 | 2E-140 | 6E-138 | 0.4149742 | n.a. |
| TBX21 | 1.658 | 9.3099 | 972.24 | 2E-139 | 6E-137 | 0.3110033 | n.a. |
| IGHG6 | 1.326 | 9.3177 | 1345.4 | 3E-124 | 9E-122 | 0.2664444 | n.a. |
| ABI3 | 1.581 | 9.3324 | 736.62 | 5E-120 | 1E-117 | 0.2891785 | n.a. |
| ITM2C | 1.595 | 9.5994 | 527.54 | 7E-116 | 2E-113 | 0.7796302 | surface |
| BIN1 | 1.465 | 9.382 | 501.35 | 3E-110 | 7E-108 | 0.3946651 | n.a. |
| FGR | 1.476 | 9.4883 | 923.49 | 2E-101 | 3E-99 | 0.2612913 | n.a. |
| ENSECAG00000031156 | 1.585 | 9.3403 | 1061.8 | 5E-101 | 9E-99 | 0.2809942 | n.a. |
| NAPSA | 1.25 | 9.5255 | 677.34 | 5E-96 | 9E-94 | 0.429221 | n.a. |
| SERPINB1 | 1.458 | 9.4507 | 517.48 | 1E-93 | 2E-91 | 0.285238 | n.a. |
| POU2F2 | 1.209 | 9.5681 | 391.68 | 1E-86 | 2E-84 | 0.7959988 | n.a. |
| RAC1 | 1.148 | 9.9054 | 390.82 | 2E-86 | 2E-84 | 0.8169142 | n.a. |
| ENSECAG00000030839 | 1.356 | 9.3107 | 795.42 | 3E-86 | 4E-84 | 0.2843286 | n.a. |
| RALGPS2 | 1.171 | 9.3274 | 443.54 | 2E-84 | 3E-82 | 0.4528645 | n.a. |
| ENSECAG00000039788 | 1.192 | 9.4822 | 370.7 | 4E-82 | 5E-80 | 0.3216126 | n.a. |
| SERPINB9 | 1.281 | 9.3883 | 367.43 | 2E-81 | 2E-79 | 0.4086087 | n.a. |
| FGD2 | 1.203 | 9.2999 | 572.12 | 2E-80 | 3E-78 | 0.2846317 | n.a. |
| TBXA2R | 1.31 | 9.2964 | 552.36 | 4E-79 | 5E-77 | 0.2512883 | surface |
| TFEB | 1.128 | 9.3544 | 544.04 | 1E-78 | 1E-76 | 0.3986056 | n.a. |
| CORO1B | 1.328 | 9.4003 | 444.42 | 4E-69 | 4E-67 | 0.2543195 | n.a. |
| ATP6V1F | 1.16 | 9.6787 | 310.21 | 4E-69 | 4E-67 | 0.6771749 | n.a. |
| ITGAM | 1.169 | 9.3026 | 668.13 | 6E-59 | 5E-57 | 0.2525008 | surface |
| FYTTD1 | 1.033 | 9.337 | 247.72 | 1E-55 | 1E-53 | 0.2652319 | n.a. |
| SYVN1 | 1.073 | 9.3092 | 301.9 | 7E-55 | 5E-53 | 0.3113065 | n.a. |
| ENSECAG00000031182 | 1.136 | 9.2942 | 572.66 | 3E-54 | 2E-52 | 0.3185814 | n.a. |
| ENSECAG00000013060 | 1.065 | 9.2815 | 607.33 | 5E-53 | 3E-51 | 0.2882692 | n.a. |
| CD37 | 1.004 | 9.891 | 213.43 | 3E-48 | 2E-46 | 0.7229463 | surface |
| ARSI | 1.009 | 9.2746 | 672.74 | 7E-48 | 4E-46 | 0.2606851 | n.a. |
| FOXP1 | 1.045 | 9.5063 | 174.03 | 1E-39 | 6E-38 | 0.3949682 | n.a. |

| genes | logFC | logCPM | F | PValue | FDR | percent.exp | Surfacome.Label |
|---|---|---|---|---|---|---|---|
| CXCR4 | 2.93 | 9.6177 | 1035.1 | 8E-224 | 2E-220 | 0.6697417 | surface |
| DRA | 1.095 | 11.742 | 732.43 | 1E-159 | 1E-156 | 0.999385 | n.a. |
| PDE4B | 2.033 | 9.326 | 725.82 | 3E-158 | 3E-155 | 0.3554736 | n.a. |
| IGHD | 2.565 | 9.2851 | 1868.5 | 3E-145 | 2E-142 | 0.4151292 | n.a. |
| BTG1 | 1.647 | 10.365 | 661.84 | 1E-144 | 9E-142 | 0.9157442 | n.a. |
| STAP1 | 1.662 | 9.3165 | 1118.5 | 6E-142 | 4E-139 | 0.5184502 | n.a. |
| MNDA | 1.616 | 10.258 | 617.08 | 5E-135 | 3E-132 | 0.9311193 | n.a. |
| FCMR | 1.635 | 9.3335 | 599.25 | 3E-131 | 2E-128 | 0.4920049 | n.a. |
| DRB | 1.084 | 11.205 | 569.69 | 7E-125 | 3E-122 | 0.9926199 | n.a. |
| Eqca-DOB1 | 1.277 | 9.4745 | 620.29 | 6E-118 | 2E-115 | 0.7183272 | n.a. |
| ENSECAG00000030839 | 1.747 | 9.3107 | 1026.7 | 2E-117 | 7E-115 | 0.3302583 | n.a. |
| SELL | 1.787 | 9.6904 | 522.38 | 9E-115 | 3E-112 | 0.4261993 | surface |
| Eqca-DQB1 | 1.157 | 10.925 | 507.06 | 2E-111 | 6E-109 | 0.9772448 | n.a. |
| HVCN1 | 1.499 | 9.3352 | 687.06 | 7E-109 | 2E-106 | 0.4876999 | n.a. |
| MARCKSL1 | 1.816 | 9.5078 | 494.09 | 1E-108 | 3E-106 | 0.2872079 | n.a. |
| UBD | 1.891 | 9.9806 | 485.03 | 9E-107 | 3E-104 | 0.7761378 | n.a. |
| DQB | 1.213 | 10.011 | 393.91 | 4E-87 | 9E-85 | 0.7699877 | n.a. |
| AIM2 | 1.334 | 9.3585 | 740.22 | 7E-86 | 2E-83 | 0.4280443 | n.a. |
| ENSECAG00000000910 | 1.449 | 9.7673 | 376.98 | 2E-83 | 3E-81 | 0.4926199 | n.a. |
| EIF1B | 1.541 | 9.5637 | 373.35 | 1E-82 | 2E-80 | 0.6340713 | n.a. |
| SESN1 | 1.59 | 9.306 | 439.95 | 1E-77 | 2E-75 | 0.2527675 | n.a. |
| EVL | 1.433 | 9.7253 | 310.2 | 4E-69 | 6E-67 | 0.2669127 | n.a. |
| ENSECAG00000034569 | 1.082 | 9.7713 | 298.93 | 1E-66 | 2E-64 | 0.5086101 | n.a. |
| ITGAE | 1.205 | 9.3387 | 270.58 | 1E-60 | 2E-58 | 0.303813 | surface |
| ENSECAG00000032776 | 1.326 | 9.4539 | 246.83 | 2E-55 | 2E-53 | 0.4065191 | n.a. |
| GNG10 | 1.035 | 9.6838 | 240.86 | 4E-54 | 4E-52 | 0.3665437 | n.a. |
| FCRLA | 1.011 | 9.3304 | 539.03 | 2E-51 | 2E-49 | 0.3690037 | n.a. |
| ZFP36L1 | 1.175 | 9.6197 | 204 | 4E-46 | 3E-44 | 0.4858549 | n.a. |
| CD37 | 1.027 | 9.891 | 188.87 | 7E-43 | 6E-41 | 0.6371464 | surface |
| FOXP1 | 1.141 | 9.5063 | 175.9 | 5E-40 | 3E-38 | 0.3708487 | n.a. |
| P2RY10 | 1.005 | 9.4226 | 151.58 | 9E-35 | 5E-33 | 0.2583026 | surface |

| genes | logFC | logCPM | F | PValue | FDR | percent.exp | Surfacome.Label |
|---|---|---|---|---|---|---|---|
| IGHG5 | 2.313 | 9.2999 | 3063.4 | 0 | 0 | 0.3355705 | n.a. |
| SH3BGRL3 | 1.571 | 10.736 | 978.99 | 6E-212 | 2E-208 | 0.957047 | n.a. |
| SELPLG | 2.13 | 9.4993 | 949.66 | 1E-205 | 2E-202 | 0.3422819 | surface |
| LGALS1 | 1.906 | 11.109 | 801.64 | 2E-174 | 3E-171 | 0.6161074 | n.a. |
| DRA | 1.076 | 11.742 | 649.43 | 6E-142 | 5E-139 | 1 | n.a. |
| ENSECAG00000038584 | 1.629 | 10.313 | 641.72 | 3E-140 | 2E-137 | 0.9422819 | n.a. |
| C11H17orf99 | 1.905 | 9.251 | 1886.1 | 2E-138 | 1E-135 | 0.3073826 | n.a. |
| ID3 | 1.927 | 9.6697 | 619.54 | 1E-135 | 9E-133 | 0.6281879 | n.a. |
| ENSECAG00000038338 | 1.522 | 9.2755 | 869.27 | 1E-133 | 6E-131 | 0.4389262 | n.a. |
| MEF2C | 1.259 | 9.4205 | 623.17 | 2E-129 | 8E-127 | 0.7691275 | n.a. |
| Eqca-DOB1 | 1.312 | 9.4745 | 617.87 | 2E-122 | 8E-120 | 0.7825503 | n.a. |
| LTB | 1.398 | 10.681 | 494.36 | 9E-109 | 4E-106 | 0.7248322 | n.a. |
| ENSECAG00000031569 | 1.029 | 11.194 | 475.85 | 9E-105 | 3E-102 | 0.9919463 | n.a. |
| GNG2 | 1.374 | 9.4802 | 469.7 | 2E-103 | 7E-101 | 0.3275168 | n.a. |
| ZBTB20 | 1.855 | 9.3268 | 468.45 | 3E-103 | 1E-100 | 0.566443 | n.a. |
| STAP1 | 1.395 | 9.3165 | 854.87 | 5E-103 | 2E-100 | 0.4926174 | n.a. |
| DQA | 1.114 | 10.712 | 445.83 | 2E-98 | 8E-96 | 0.990604 | n.a. |
| GLIPR1 | 1.511 | 9.887 | 429 | 1E-94 | 3E-92 | 0.6738255 | surface |
| ENSECAG00000029003 | 1.425 | 10.041 | 416.55 | 5E-92 | 1E-89 | 0.6442953 | n.a. |
| ITGB1 | 1.641 | 9.5309 | 411.92 | 5E-91 | 1E-88 | 0.2724832 | surface |
| Eqca-DQB1 | 1.096 | 10.925 | 408.74 | 2E-90 | 6E-88 | 0.9946309 | n.a. |
| REV1 | 1.525 | 9.2709 | 564.05 | 3E-87 | 8E-85 | 0.2536913 | n.a. |
| DQB | 1.244 | 10.011 | 375.89 | 3E-83 | 7E-81 | 0.8187919 | n.a. |
| SELL | 1.592 | 9.6904 | 372.83 | 1E-82 | 3E-80 | 0.4147651 | surface |
| ENSECAG00000040634 | 1.43 | 9.9556 | 366.24 | 3E-81 | 7E-79 | 0.7744966 | n.a. |
| IFI30 | 1.031 | 9.9429 | 356.46 | 4E-79 | 9E-77 | 0.8939597 | n.a. |
| FCMR | 1.185 | 9.3335 | 354.39 | 4E-75 | 7E-73 | 0.4550336 | n.a. |
| SMAGP | 1.293 | 9.3065 | 348.16 | 1E-74 | 3E-72 | 0.3463087 | n.a. |
| S100A6 | 1.301 | 10.437 | 317.16 | 1E-70 | 2E-68 | 0.6134228 | n.a. |
| FABP3 | 1.116 | 9.3412 | 396.79 | 6E-67 | 9E-65 | 0.5127517 | n.a. |
| VIM | 1.002 | 11.286 | 275.43 | 1E-61 | 2E-59 | 0.7798658 | n.a. |
| BTG1 | 1.112 | 10.365 | 274.28 | 2E-61 | 4E-59 | 0.8684564 | n.a. |
| ACTG1 | 1.028 | 10.745 | 268.41 | 4E-60 | 7E-58 | 0.7194631 | n.a. |
| AIM2 | 1.209 | 9.3585 | 594.11 | 4E-58 | 7E-56 | 0.4724832 | n.a. |
| PLP2 | 1.03 | 9.9939 | 249.95 | 4E-56 | 6E-54 | 0.8389262 | n.a. |
| ENSECAG00000034569 | 1.077 | 9.7713 | 249.87 | 4E-56 | 6E-54 | 0.5812081 | n.a. |
| MDH2 | 1.109 | 9.573 | 244.64 | 6E-55 | 8E-53 | 0.5583893 | n.a. |
| DBNL | 1.111 | 9.4448 | 230.38 | 7E-52 | 1E-49 | 0.5208054 | n.a. |
| S100A11 | 1.264 | 9.847 | 228.72 | 2E-51 | 2E-49 | 0.2979866 | n.a. |
| CXCR4 | 1.181 | 9.6177 | 226.23 | 6E-51 | 8E-49 | 0.3181208 | surface |
| HLA-DMA | 1.019 | 9.6051 | 223.19 | 3E-50 | 3E-48 | 0.6442953 | surface |
| UBD | 1.357 | 9.9806 | 220.04 | 1E-49 | 2E-47 | 0.6711409 | n.a. |
| GNG10 | 1.014 | 9.6838 | 193.23 | 8E-44 | 1E-41 | 0.4818792 | n.a. |
| ACTN1 | 1.045 | 9.4643 | 188.94 | 7E-43 | 8E-41 | 0.2765101 | n.a. |
| PDLIM2 | 1.14 | 9.5617 | 185.15 | 5E-42 | 5E-40 | 0.3597315 | n.a. |
| ENSECAG00000013303 | 1.185 | 9.4002 | 266.49 | 2E-39 | 2E-37 | 0.3127517 | n.a. |
| HIF1A | 1.01 | 9.3685 | 187.83 | 2E-36 | 1E-34 | 0.3543624 | n.a. |

| | | | | | | | |
|---|---|---|---|---|---|---|---|
| S100A4 | 1.085 | 11.292 | 100.64 | 1E-23 | 6E-22 | 0.261745 | n.a. |

| genes | logFC | logCPM | F | PValue | FDR | percent.exp | Surfacome.Label |
|---|---|---|---|---|---|---|---|
| IGHG6 | 4.396 | 9.3177 | 3802.6 | 0 | 0 | 0.7139831 | n.a. |
| ENSECAG00000014585 | 3.88 | 10.271 | 4019.5 | 0 | 0 | 1 | n.a. |
| ENSECAG00000001010 | 3.335 | 9.629 | 2579.8 | 0 | 0 | 0.9724576 | n.a. |
| ENSECAG00000029287 | 3.32 | 10.043 | 1898.9 | 0 | 0 | 0.9004237 | n.a. |
| PLAC8B | 2.979 | 11.113 | 2078.3 | 0 | 0 | 1 | n.a. |
| IGHG3 | 2.395 | 9.6047 | 2207.9 | 0 | 0 | 0.7372881 | n.a. |
| RPS20 | 1.823 | 13.671 | 4107.1 | 0 | 0 | 1 | n.a. |
| LAPTM5 | 1.566 | 11.361 | 1279.6 | 3E-275 | 3E-272 | 1 | n.a. |
| ENSECAG00000027826 | 1.587 | 9.7443 | 1178.8 | 5E-254 | 4E-251 | 0.9851695 | n.a. |
| CMTM3 | 2.979 | 9.4968 | 1032.8 | 3E-223 | 2E-220 | 0.8326271 | n.a. |
| ENSECAG00000035439 | 2.398 | 9.2665 | 2386.8 | 5E-209 | 3E-206 | 0.5635593 | n.a. |
| BANK1 | 1.949 | 9.413 | 976.07 | 1E-194 | 6E-192 | 0.7966102 | n.a. |
| ANKRD2 | 2.275 | 9.3125 | 1814.5 | 2E-189 | 1E-186 | 0.5995763 | n.a. |
| HSF2BP | 1.969 | 9.3751 | 1142.7 | 4E-188 | 2E-185 | 0.7033898 | n.a. |
| ENSECAG00000012199 | 2.157 | 9.3154 | 1039.9 | 5E-169 | 2E-166 | 0.5487288 | n.a. |
| PTPRCAP | 1.622 | 10.617 | 724.74 | 5E-158 | 2E-155 | 0.9978814 | n.a. |
| ENSECAG00000033804 | 2.01 | 9.2877 | 1818.2 | 8E-153 | 3E-150 | 0.4004237 | n.a. |
| ENSECAG00000039788 | 2.151 | 9.4822 | 753.67 | 1E-146 | 5E-144 | 0.529661 | n.a. |
| TSPAN4 | 2.275 | 9.2911 | 1709.8 | 8E-145 | 3E-142 | 0.3305085 | surface |
| SH3BP5 | 1.668 | 9.4429 | 740.84 | 2E-143 | 6E-141 | 0.7987288 | n.a. |
| ITM2C | 2.028 | 9.5994 | 612.72 | 4E-134 | 1E-131 | 0.8919492 | surface |
| GNGT2 | 2.05 | 9.2643 | 1841.4 | 1E-130 | 3E-128 | 0.2648305 | n.a. |
| ADORA2A | 2.037 | 9.2768 | 951.75 | 1E-126 | 3E-124 | 0.4173729 | surface |
| CYSTM1 | 1.764 | 9.3828 | 647.33 | 2E-125 | 6E-123 | 0.6228814 | n.a. |
| CXXC5 | 2.063 | 9.3065 | 695.5 | 3E-125 | 9E-123 | 0.4427966 | n.a. |
| BPIFB4 | 2.168 | 9.249 | 2022.8 | 1E-124 | 3E-122 | 0.375 | n.a. |
| MS4A1 | 1.006 | 9.7256 | 553.42 | 2E-121 | 5E-119 | 0.940678 | surface |
| AHNAK | 2.024 | 9.5205 | 597.27 | 3E-121 | 9E-119 | 0.5317797 | n.a. |
| TBX21 | 2.056 | 9.3099 | 1038.6 | 3E-119 | 8E-117 | 0.4258475 | n.a. |
| SERPINB1 | 2.172 | 9.4507 | 710.74 | 2E-115 | 5E-113 | 0.4724576 | n.a. |
| ABI3 | 2.085 | 9.3324 | 830.36 | 3E-114 | 7E-112 | 0.4258475 | n.a. |
| RF01277 | 1.624 | 9.8037 | 515.46 | 3E-113 | 6E-111 | 0.9216102 | n.a. |
| SERPINB9 | 1.878 | 9.3883 | 501.72 | 2E-110 | 5E-108 | 0.5847458 | n.a. |
| CALM2 | 1.261 | 10.275 | 442.86 | 1E-97 | 2E-95 | 0.9830508 | n.a. |
| ATP6V1F | 1.705 | 9.6787 | 436.73 | 2E-96 | 4E-94 | 0.8411017 | n.a. |
| SNRNP200 | 1.736 | 9.406 | 396.24 | 1E-87 | 2E-85 | 0.6631356 | n.a. |
| FGL2 | 1.702 | 9.488 | 709.98 | 1E-82 | 2E-80 | 0.3601695 | n.a. |
| POU2F2 | 1.401 | 9.5681 | 368.52 | 1E-81 | 2E-79 | 0.8538136 | n.a. |
| BIN1 | 1.566 | 9.382 | 403.62 | 3E-81 | 5E-79 | 0.4491525 | n.a. |
| BHLHE41 | 1.399 | 9.3067 | 740.11 | 1E-80 | 2E-78 | 0.5614407 | n.a. |
| EMP3 | 1.455 | 10.126 | 357.34 | 3E-79 | 4E-77 | 0.7690678 | surface |
| TNNI3 | 1.891 | 9.2525 | 1185.2 | 4E-79 | 7E-77 | 0.3177966 | n.a. |
| CYBB | 1.538 | 9.4653 | 692.63 | 7E-76 | 9E-74 | 0.5550847 | surface |
| TBXA2R | 1.674 | 9.2964 | 592.62 | 3E-74 | 4E-72 | 0.3241525 | surface |
| ARSI | 1.597 | 9.2746 | 946.84 | 2E-71 | 2E-69 | 0.4152542 | n.a. |
| APEX1 | 1.382 | 9.4275 | 304.53 | 7E-68 | 8E-66 | 0.6864407 | n.a. |
| PLEK | 1.625 | 9.3075 | 605.42 | 8E-66 | 1E-63 | 0.279661 | n.a. |

| | | | | | | | |
|---|---|---|---|---|---|---|---|
| POLR2E | 1.519 | 9.4718 | 288.57 | 2E-64 | 2E-62 | 0.6292373 | n.a. |
| POLD1 | 1.055 | 9.3555 | 305.54 | 2E-58 | 2E-56 | 0.5995763 | n.a. |
| CYB561A3 | 1.15 | 9.3685 | 310.83 | 1E-56 | 1E-54 | 0.5932203 | n.a. |
| FUT8 | 1.264 | 9.3425 | 234.53 | 9E-53 | 9E-51 | 0.3961864 | n.a. |
| STX11 | 1.282 | 9.3844 | 237.69 | 8E-50 | 8E-48 | 0.5 | n.a. |
| ITGB2 | 1.228 | 9.8101 | 214.82 | 2E-48 | 2E-46 | 0.625 | surface |
| ENSECAG00000021652 | 1.178 | 9.3311 | 214.13 | 3E-48 | 3E-46 | 0.4724576 | n.a. |
| TK2 | 1.381 | 9.3075 | 350.93 | 4E-48 | 4E-46 | 0.279661 | n.a. |
| CCND3 | 1.254 | 9.7279 | 212.82 | 5E-48 | 4E-46 | 0.7097458 | n.a. |
| PYCARD | 1.085 | 9.489 | 225.2 | 6E-48 | 5E-46 | 0.6165254 | n.a. |
| TFEB | 1.209 | 9.3544 | 420.5 | 5E-47 | 4E-45 | 0.4661017 | n.a. |
| MMD | 1.237 | 9.3359 | 233.42 | 7E-47 | 6E-45 | 0.345339 | n.a. |
| TIFA | 1.114 | 10.029 | 205.28 | 2E-46 | 2E-44 | 0.8580508 | n.a. |
| PDCD5 | 1.245 | 9.4004 | 191.15 | 2E-43 | 2E-41 | 0.4427966 | n.a. |
| PRDX1 | 1.028 | 9.8857 | 189.66 | 5E-43 | 4E-41 | 0.8898305 | n.a. |
| ITGAM | 1.3 | 9.3026 | 597.11 | 7E-43 | 5E-41 | 0.2902542 | surface |
| PYGM | 1.059 | 9.3286 | 367.06 | 1E-42 | 9E-41 | 0.4978814 | n.a. |
| NCOA7 | 1.245 | 9.3234 | 231.83 | 2E-41 | 1E-39 | 0.3898305 | n.a. |
| PKIG | 1.103 | 9.271 | 595.2 | 2E-39 | 1E-37 | 0.3474576 | n.a. |
| PLD4 | 1.184 | 9.3359 | 400.08 | 1E-38 | 7E-37 | 0.3919492 | n.a. |
| BATF | 1.176 | 9.3505 | 230.8 | 2E-37 | 1E-35 | 0.3220339 | n.a. |
| AKT2 | 1.209 | 9.2915 | 273.13 | 1E-36 | 8E-35 | 0.279661 | n.a. |
| RALGPS2 | 1.032 | 9.3274 | 260.53 | 1E-36 | 9E-35 | 0.4639831 | n.a. |
| CHID1 | 1.053 | 9.3333 | 203.78 | 1E-35 | 9E-34 | 0.2987288 | n.a. |
| CRACR2B | 1.08 | 9.2743 | 328.99 | 2E-35 | 1E-33 | 0.2902542 | n.a. |
| DESI1 | 1.113 | 9.3256 | 188.56 | 4E-35 | 2E-33 | 0.3813559 | n.a. |
| IRF5 | 1.063 | 9.322 | 333.19 | 4E-35 | 3E-33 | 0.3771186 | n.a. |
| ENSECAG00000031156 | 1.251 | 9.3403 | 549.94 | 1E-34 | 7E-33 | 0.2605932 | n.a. |
| ACADM | 1.147 | 9.4176 | 151.03 | 1E-34 | 7E-33 | 0.434322 | n.a. |
| TCEA3 | 1.219 | 9.3171 | 279.45 | 1E-34 | 8E-33 | 0.3008475 | n.a. |
| FYTTD1 | 1.148 | 9.337 | 192.85 | 1E-33 | 6E-32 | 0.3199153 | n.a. |
| ARHGAP15 | 1.072 | 9.6767 | 145.74 | 2E-33 | 9E-32 | 0.720339 | n.a. |
| NUDT8 | 1.058 | 9.3058 | 210.61 | 9E-33 | 5E-31 | 0.309322 | n.a. |
| TMEM206 | 1.183 | 9.289 | 284.15 | 2E-32 | 1E-30 | 0.2944915 | n.a. |
| NUAK2 | 1.15 | 9.2961 | 224.84 | 2E-30 | 8E-29 | 0.3283898 | n.a. |
| APBB1IP | 1.009 | 9.6791 | 125.98 | 3E-29 | 2E-27 | 0.5995763 | n.a. |
| PXK | 1.056 | 9.2957 | 200.62 | 2E-28 | 1E-26 | 0.2775424 | n.a. |
| UBE2E1 | 1.087 | 9.3227 | 175.84 | 5E-28 | 2E-26 | 0.2542373 | n.a. |
| FGD2 | 1.029 | 9.2999 | 318.08 | 1E-27 | 5E-26 | 0.3008475 | n.a. |
| BLNK | 1.012 | 9.2737 | 401.74 | 2E-24 | 9E-23 | 0.2817797 | n.a. |
| IFNGR1 | 1.023 | 9.4289 | 131.89 | 1E-23 | 5E-22 | 0.2521186 | surface |

| genes | logFC | logCPM | F | PValue | FDR | percent.exp | Surfacome.Label |
|---|---|---|---|---|---|---|---|
| JCHAIN | 8.319 | 9.7044 | 11336 | 0 | 0 | 0.952381 | n.a. |
| IGHA | 8.201 | 9.6684 | 11984 | 0 | 0 | 0.5833333 | n.a. |
| ENSECAG00000031094 | 4.887 | 10.892 | 4544.7 | 0 | 0 | 0.7619048 | n.a. |
| ENSECAG00000031522 | 4.866 | 10.189 | 2129.9 | 0 | 0 | 0.8095238 | n.a. |
| ENSECAG00000039599 | 4.86 | 10.091 | 2580.3 | 0 | 0 | 0.6785714 | n.a. |
| SLC23A1 | 4.77 | 9.2465 | 6990.5 | 0 | 0 | 0.8333333 | surface |
| ENSECAG00000015109 | 4.012 | 9.6255 | 1583.1 | 0 | 0 | 0.7738095 | n.a. |
| TXNDC5 | 4.381 | 9.249 | 3835.9 | 0 | 0 | 0.8214286 | n.a. |
| IGHM | 3.86 | 9.6407 | 1410.8 | 1E-302 | 7E-300 | 0.2857143 | n.a. |
| HSP90B1 | 3.913 | 9.5518 | 1143.2 | 1E-246 | 7E-244 | 0.9285714 | n.a. |
| XBP1 | 3.705 | 9.4868 | 1080.4 | 2E-233 | 1E-230 | 0.8452381 | n.a. |
| SELENOS | 3.497 | 9.3395 | 1115.9 | 3E-230 | 1E-227 | 0.8452381 | n.a. |
| SEC61B | 2.975 | 9.8749 | 869.76 | 8E-189 | 3E-186 | 0.952381 | n.a. |
| PPIB | 2.472 | 10.48 | 866.27 | 4E-188 | 2E-185 | 0.9761905 | n.a. |
| PRDX2 | 3.098 | 9.2964 | 863.87 | 1E-187 | 5E-185 | 0.7142857 | n.a. |
| PDIA4 | 3.28 | 9.2985 | 1029.6 | 4E-181 | 1E-178 | 0.7738095 | n.a. |
| SEC11C | 3.464 | 9.493 | 826.41 | 1E-179 | 4E-177 | 0.9047619 | n.a. |
| SSR3 | 2.772 | 9.9724 | 752.09 | 8E-164 | 2E-161 | 0.9166667 | n.a. |
| AQP3 | 2.886 | 9.3499 | 942.33 | 3E-162 | 8E-160 | 0.6190476 | n.a. |
| CRELD2 | 3.172 | 9.2876 | 731.39 | 2E-159 | 6E-157 | 0.8809524 | n.a. |
| SDF2L1 | 3.292 | 9.3874 | 727.98 | 1E-158 | 3E-156 | 0.8214286 | n.a. |
| RRBP1 | 3.048 | 9.2913 | 1843.9 | 4E-154 | 1E-151 | 0.547619 | n.a. |
| MANF | 3.245 | 9.4674 | 693.15 | 3E-151 | 7E-149 | 0.8690476 | n.a. |
| EAF2 | 3.521 | 9.2421 | 3642.1 | 2E-150 | 4E-148 | 0.7380952 | n.a. |
| RABAC1 | 2.847 | 9.8034 | 681.25 | 1E-148 | 2E-146 | 0.9404762 | n.a. |
| TENT5C | 3.549 | 9.2576 | 1333.2 | 2E-147 | 5E-145 | 0.7857143 | n.a. |
| PDIA6 | 3.162 | 9.332 | 656.92 | 2E-143 | 3E-141 | 0.75 | n.a. |
| REXO2 | 2.797 | 9.4557 | 644.01 | 9E-141 | 2E-138 | 0.8452381 | n.a. |
| ENSECAG00000036762 | 3.302 | 9.2422 | 3364.7 | 1E-139 | 3E-137 | 0.797619 | n.a. |
| PRDX4 | 3.203 | 9.3564 | 616.14 | 8E-135 | 1E-132 | 0.8095238 | n.a. |
| DUSP5 | 2.784 | 9.267 | 678.14 | 1E-120 | 2E-118 | 0.7619048 | n.a. |
| IRF4 | 3.133 | 9.2503 | 1186.4 | 1E-119 | 2E-117 | 0.6071429 | n.a. |
| FKBP11 | 3.136 | 9.2432 | 2421.9 | 2E-119 | 3E-117 | 0.6428571 | n.a. |
| FKBP2 | 2.688 | 9.3386 | 525.07 | 2E-115 | 4E-113 | 0.8095238 | n.a. |
| ENSECAG00000026891 | 1.887 | 9.2736 | 1774.7 | 8E-115 | 1E-112 | 0.3452381 | n.a. |
| DHRS3 | 2.495 | 9.3111 | 1014.3 | 6E-113 | 1E-110 | 0.6428571 | n.a. |
| LMAN1 | 2.911 | 9.4114 | 502.54 | 2E-110 | 2E-108 | 0.8571429 | n.a. |
| PIM1 | 2.579 | 9.3916 | 501.97 | 2E-110 | 3E-108 | 0.8214286 | n.a. |
| CALR | 2.599 | 9.7942 | 501.92 | 2E-110 | 3E-108 | 0.9166667 | n.a. |
| HM13 | 2.585 | 9.3911 | 463.98 | 3E-102 | 4E-100 | 0.8333333 | surface |
| STMN1 | 2.173 | 9.7683 | 461.48 | 1E-101 | 1E-99 | 0.797619 | n.a. |
| ITGA4 | 2.318 | 9.465 | 461.36 | 1E-101 | 2E-99 | 0.7261905 | surface |
| SPCS2 | 2.659 | 9.6657 | 457.48 | 8E-101 | 1E-98 | 0.9285714 | n.a. |
| BLVRB | 3.137 | 9.4497 | 809.35 | 4E-95 | 5E-93 | 0.3452381 | n.a. |
| CRYBB1 | 2.797 | 9.2422 | 2982.4 | 2E-94 | 2E-92 | 0.547619 | n.a. |
| ENSECAG00000001010 | 1.751 | 9.629 | 426.44 | 4E-94 | 4E-92 | 0.4166667 | n.a. |
| SPCS3 | 2.411 | 9.6267 | 419.01 | 1E-92 | 2E-90 | 0.952381 | n.a. |

| | | | | | | | |
|---|---|---|---|---|---|---|---|
| DNAJC3 | 2.464 | 9.3287 | 451.88 | 8E-91 | 9E-89 | 0.7619048 | n.a. |
| BRI3BP | 2.377 | 9.2708 | 610.39 | 5E-83 | 6E-81 | 0.6666667 | n.a. |
| P4HB | 2.258 | 9.4063 | 369.68 | 6E-82 | 7E-80 | 0.7857143 | n.a. |
| SSR1 | 2.415 | 9.3182 | 428.03 | 1E-81 | 1E-79 | 0.6666667 | surface |
| SPG21 | 2.548 | 9.3532 | 364.41 | 8E-81 | 9E-79 | 0.7380952 | n.a. |
| OSTC | 2.396 | 9.5663 | 359.4 | 1E-79 | 1E-77 | 0.8928571 | n.a. |
| DNAJB9 | 2.592 | 9.2718 | 711.27 | 4E-77 | 4E-75 | 0.5714286 | n.a. |
| PDIA5 | 2.734 | 9.2449 | 1820.1 | 8E-77 | 9E-75 | 0.6547619 | n.a. |
| ALG5 | 2.445 | 9.3503 | 342.74 | 4E-76 | 4E-74 | 0.7619048 | n.a. |
| UBE2J1 | 2.466 | 9.2895 | 446.53 | 5E-76 | 5E-74 | 0.7619048 | n.a. |
| SSR2 | 1.867 | 10.096 | 341.11 | 8E-76 | 8E-74 | 0.9761905 | n.a. |
| COX2 | 1.147 | 11.529 | 330.84 | 1E-73 | 1E-71 | 0.9642857 | n.a. |
| TPST2 | 2.412 | 9.4709 | 325.98 | 2E-72 | 2E-70 | 0.8928571 | n.a. |
| RPN1 | 2.324 | 9.4173 | 311.46 | 2E-69 | 2E-67 | 0.797619 | surface |
| GRID2IP | 2.376 | 9.4806 | 310.44 | 4E-69 | 3E-67 | 0.8809524 | n.a. |
| MAN1A1 | 2.323 | 9.2616 | 840.08 | 8E-69 | 7E-67 | 0.5714286 | n.a. |
| TMED2 | 2.163 | 9.6321 | 302.78 | 2E-67 | 1E-65 | 0.8928571 | n.a. |
| KRTCAP2 | 2.142 | 9.5812 | 288.2 | 2E-64 | 2E-62 | 0.8928571 | n.a. |
| SSR4 | 1.566 | 10.501 | 277.87 | 4E-62 | 3E-60 | 0.9761905 | n.a. |
| SEC22B | 2.155 | 9.3214 | 276.75 | 7E-62 | 6E-60 | 0.7261905 | n.a. |
| TMBIM6 | 2.08 | 9.7742 | 270.78 | 1E-60 | 1E-58 | 0.9285714 | n.a. |
| SEC61G | 1.706 | 9.9637 | 269.64 | 2E-60 | 2E-58 | 0.952381 | n.a. |
| ITM2B | 2.053 | 10.174 | 259.94 | 3E-58 | 2E-56 | 0.8452381 | surface |
| SRPRB | 2.161 | 9.3465 | 258.64 | 6E-58 | 4E-56 | 0.702381 | n.a. |
| S100A10 | 1.541 | 10.866 | 256.66 | 1E-57 | 1E-55 | 0.9047619 | n.a. |
| PQLC3 | 2.265 | 9.3249 | 388.91 | 6E-57 | 4E-55 | 0.5238095 | n.a. |
| ENSECAG00000032916 | 1.55 | 10.218 | 248.03 | 1E-55 | 8E-54 | 0.9642857 | n.a. |
| MLLT3 | 2.146 | 9.2897 | 572.83 | 1E-55 | 9E-54 | 0.5238095 | n.a. |
| SEC61A1 | 1.959 | 9.3336 | 261.79 | 2E-55 | 2E-53 | 0.7142857 | n.a. |
| ENSECAG00000014391 | 2.082 | 9.3144 | 243.02 | 1E-54 | 1E-52 | 0.7142857 | n.a. |
| HYOU1 | 2.284 | 9.257 | 847.01 | 2E-54 | 1E-52 | 0.5238095 | n.a. |
| PDIA3 | 1.973 | 9.7363 | 239.55 | 7E-54 | 5E-52 | 0.9047619 | n.a. |
| ENSECAG00000023261 | 1.945 | 9.5604 | 237.68 | 2E-53 | 1E-51 | 0.8809524 | n.a. |
| TRAM1 | 2.045 | 9.4843 | 237.24 | 2E-53 | 2E-51 | 0.7619048 | n.a. |
| TSPAN13 | 1.834 | 9.2631 | 785.22 | 2E-52 | 1E-50 | 0.3928571 | surface |
| SYS1 | 2.12 | 9.288 | 341.95 | 2E-52 | 1E-50 | 0.6071429 | n.a. |
| EMC9 | 1.872 | 9.2686 | 608.56 | 6E-51 | 4E-49 | 0.4880952 | n.a. |
| BICDL2 | 2.177 | 9.2439 | 1376 | 4E-49 | 3E-47 | 0.5119048 | n.a. |
| CREB3L2 | 1.947 | 9.2412 | 1963.3 | 8E-49 | 5E-47 | 0.4166667 | n.a. |
| TMED9 | 2.053 | 9.3567 | 216.32 | 8E-49 | 5E-47 | 0.6547619 | n.a. |
| SRP14 | 1.563 | 10.144 | 216.24 | 8E-49 | 5E-47 | 0.9404762 | n.a. |
| ERLEC1 | 2.041 | 9.3911 | 208.06 | 5E-47 | 3E-45 | 0.7738095 | n.a. |
| H2AFX | 1.711 | 9.3048 | 292.55 | 2E-45 | 1E-43 | 0.452381 | n.a. |
| KDELR1 | 1.961 | 9.4158 | 200.21 | 3E-45 | 2E-43 | 0.7380952 | n.a. |
| PRDM1 | 1.877 | 9.2548 | 867.76 | 5E-45 | 3E-43 | 0.4285714 | n.a. |
| TUT7 | 1.925 | 9.6301 | 190.63 | 3E-43 | 2E-41 | 0.7857143 | n.a. |
| SPCS1 | 1.581 | 9.9456 | 188.72 | 8E-43 | 5E-41 | 0.9166667 | n.a. |
| ASPH | 1.552 | 9.2561 | 1026.9 | 2E-42 | 1E-40 | 0.2738095 | n.a. |

| | | | | | | | |
|---|---|---|---|---|---|---|---|
| RNF149 | 1.731 | 9.3444 | 217.14 | 4E-41 | 2E-39 | 0.6666667 | surface |
| CDCA3 | 1.424 | 9.2462 | 2868.6 | 8E-41 | 5E-39 | 0.3214286 | n.a. |
| MGAT1 | 1.829 | 9.3346 | 179.5 | 8E-41 | 5E-39 | 0.7380952 | n.a. |
| B4GALT3 | 2.139 | 9.273 | 380.8 | 9E-41 | 5E-39 | 0.547619 | n.a. |
| RPN2 | 1.859 | 9.4695 | 179.14 | 9E-41 | 5E-39 | 0.8095238 | n.a. |
| ENSECAG00000006192 | 1.646 | 9.2352 | 3595.5 | 2E-40 | 1E-38 | 0.2857143 | n.a. |
| ANXA2 | 1.362 | 9.8875 | 169.49 | 1E-38 | 6E-37 | 0.5595238 | n.a. |
| SELPLG | 1.278 | 9.4993 | 170.66 | 1E-38 | 8E-37 | 0.5714286 | surface |
| RRM2 | 1.2 | 9.252 | 756.27 | 3E-37 | 1E-35 | 0.2619048 | n.a. |
| CYTIP | 1.787 | 9.6111 | 162.39 | 4E-37 | 2E-35 | 0.702381 | n.a. |
| ERGIC2 | 1.731 | 9.3256 | 155.87 | 1E-35 | 5E-34 | 0.7857143 | n.a. |
| IFI6 | 1.607 | 9.3792 | 155.7 | 1E-35 | 6E-34 | 0.6428571 | n.a. |
| GMPPB | 1.836 | 9.2571 | 465.35 | 2E-35 | 8E-34 | 0.5 | n.a. |
| MTDH | 1.548 | 9.6915 | 154.8 | 2E-35 | 9E-34 | 0.9166667 | n.a. |
| GOLGB1 | 1.725 | 9.3504 | 146.01 | 5E-33 | 3E-31 | 0.7261905 | n.a. |
| EDEM2 | 1.686 | 9.308 | 169.25 | 7E-33 | 3E-31 | 0.5357143 | n.a. |
| HDLBP | 1.595 | 9.337 | 142.59 | 8E-33 | 4E-31 | 0.6904762 | n.a. |
| ENSECAG00000010480 | 1.611 | 9.3685 | 139.96 | 3E-32 | 1E-30 | 0.5952381 | n.a. |
| H2AFZ | 1.5 | 9.5191 | 136.7 | 2E-31 | 7E-30 | 0.6071429 | n.a. |
| MAN2A1 | 1.821 | 9.2929 | 266.55 | 5E-31 | 2E-29 | 0.5 | n.a. |
| REEP4 | 1.227 | 9.2747 | 186 | 8E-31 | 4E-29 | 0.4404762 | n.a. |
| ENSECAG00000012455 | 1.65 | 9.3151 | 167.78 | 2E-30 | 1E-28 | 0.5714286 | n.a. |
| CREB3 | 1.591 | 9.3037 | 140.03 | 3E-30 | 1E-28 | 0.6190476 | n.a. |
| CD99 | 1.725 | 9.2965 | 255.91 | 2E-29 | 8E-28 | 0.5595238 | n.a. |
| TMED10 | 1.646 | 9.3206 | 187.66 | 2E-28 | 1E-26 | 0.4642857 | n.a. |
| DERL1 | 1.555 | 9.3343 | 120.31 | 6E-28 | 3E-26 | 0.6666667 | n.a. |
| NANS | 1.509 | 9.4296 | 119.24 | 1E-27 | 4E-26 | 0.7857143 | n.a. |
| C1H15orf48 | 1.027 | 9.6929 | 118.97 | 1E-27 | 5E-26 | 0.5 | n.a. |
| ENSECAG00000036511 | 1.526 | 9.3303 | 117.52 | 2E-27 | 1E-25 | 0.7857143 | n.a. |
| ENSECAG00000024654 | 1.434 | 9.3419 | 117.43 | 3E-27 | 1E-25 | 0.4761905 | n.a. |
| PLEKHA8 | 1.567 | 9.2354 | 3767.8 | 5E-27 | 2E-25 | 0.3333333 | n.a. |
| VSIR | 1.365 | 9.4326 | 139.04 | 6E-27 | 2E-25 | 0.5238095 | n.a. |
| SURF4 | 1.542 | 9.3205 | 173.84 | 1E-26 | 6E-25 | 0.5357143 | n.a. |
| CKAP4 | 1.376 | 9.2455 | 904.03 | 2E-26 | 7E-25 | 0.297619 | n.a. |
| MLEC | 1.676 | 9.2587 | 381.07 | 3E-26 | 1E-24 | 0.4404762 | n.a. |
| DDOST | 1.451 | 9.3389 | 118.08 | 7E-26 | 3E-24 | 0.6547619 | n.a. |
| ENSECAG00000032940 | 1.381 | 9.2852 | 244.43 | 7E-26 | 3E-24 | 0.3690476 | n.a. |
| STT3A | 1.484 | 9.2923 | 154.74 | 2E-25 | 9E-24 | 0.4404762 | n.a. |
| TMEM59 | 1.484 | 9.5994 | 107.56 | 4E-25 | 1E-23 | 0.8333333 | n.a. |
| ARL4C | 1.281 | 9.3066 | 243.15 | 7E-25 | 3E-23 | 0.3333333 | n.a. |
| CCR10 | 1.558 | 9.2411 | 1699.8 | 9E-25 | 3E-23 | 0.3452381 | surface |
| ERP29 | 1.333 | 9.3518 | 105 | 1E-24 | 5E-23 | 0.5714286 | n.a. |
| CDK2AP2 | 1.343 | 9.6423 | 104.28 | 2E-24 | 7E-23 | 0.8214286 | n.a. |
| ENSECAG00000027666 | 1.186 | 9.8957 | 100.21 | 1E-23 | 5E-22 | 0.8928571 | n.a. |
| SYPL1 | 1.316 | 9.5663 | 99.84 | 2E-23 | 6E-22 | 0.8809524 | surface |
| ZNF706 | 1.157 | 9.8395 | 99.742 | 2E-23 | 6E-22 | 0.952381 | n.a. |
| SRM | 1.481 | 9.2691 | 148.6 | 2E-23 | 7E-22 | 0.4166667 | n.a. |
| HERPUD1 | 1.472 | 9.4297 | 98.732 | 3E-23 | 1E-21 | 0.702381 | n.a. |

| | | | | | | | |
|---|---|---|---|---|---|---|---|
| CANX | 1.383 | 9.3959 | 99.739 | 5E-23 | 2E-21 | 0.7261905 | n.a. |
| SLC35C1 | 1.35 | 9.2505 | 405.47 | 2E-22 | 6E-21 | 0.3214286 | n.a. |
| SRPRA | 1.427 | 9.3887 | 94.331 | 3E-22 | 9E-21 | 0.6428571 | n.a. |
| PRR5 | 1.121 | 9.2846 | 249.37 | 5E-22 | 2E-20 | 0.3095238 | n.a. |
| LMAN2 | 1.386 | 9.4812 | 93.281 | 5E-22 | 2E-20 | 0.7619048 | surface |
| USO1 | 1.485 | 9.2993 | 146.89 | 6E-22 | 2E-20 | 0.5238095 | n.a. |
| EDEM3 | 1.49 | 9.2417 | 789.87 | 7E-22 | 2E-20 | 0.297619 | n.a. |
| EIF5 | 1.338 | 9.6081 | 91.821 | 1E-21 | 3E-20 | 0.8333333 | n.a. |
| OGFR | 1.344 | 9.3302 | 92.326 | 2E-21 | 5E-20 | 0.5952381 | n.a. |
| MDH1 | 1.366 | 9.4669 | 89.896 | 3E-21 | 8E-20 | 0.8214286 | n.a. |
| ENSECAG00000006721 | 1.302 | 9.454 | 88.83 | 5E-21 | 1E-19 | 0.6904762 | n.a. |
| UFM1 | 1.362 | 9.3419 | 88.517 | 5E-21 | 2E-19 | 0.5833333 | n.a. |
| BET1 | 1.358 | 9.3715 | 86.34 | 2E-20 | 5E-19 | 0.6666667 | n.a. |
| SEL1L | 1.398 | 9.2493 | 419.1 | 3E-20 | 1E-18 | 0.3333333 | n.a. |
| GMDS | 1.359 | 9.2963 | 105.12 | 4E-20 | 1E-18 | 0.6547619 | n.a. |
| MOGS | 1.336 | 9.2692 | 157.99 | 4E-20 | 1E-18 | 0.4404762 | n.a. |
| FAM117A | 1.287 | 9.3289 | 94.633 | 4E-20 | 1E-18 | 0.5119048 | n.a. |
| RNF5 | 1.361 | 9.3876 | 84.534 | 4E-20 | 1E-18 | 0.6666667 | n.a. |
| FAM219A | 1.354 | 9.2405 | 726.56 | 5E-20 | 1E-18 | 0.3095238 | n.a. |
| SND1 | 1.371 | 9.3185 | 118.89 | 5E-20 | 1E-18 | 0.5952381 | n.a. |
| SEC24A | 1.434 | 9.2501 | 344.84 | 7E-20 | 2E-18 | 0.3809524 | n.a. |
| C14H5orf30 | 1.332 | 9.2435 | 643.35 | 1E-19 | 3E-18 | 0.2738095 | n.a. |
| CCL5 | 1.565 | 11.908 | 82.183 | 1E-19 | 4E-18 | 0.297619 | n.a. |
| ENSECAG00000035182 | 1.125 | 9.2638 | 96.981 | 3E-19 | 7E-18 | 0.3571429 | n.a. |
| SAR1A | 1.258 | 9.3164 | 81.724 | 5E-19 | 1E-17 | 0.5714286 | n.a. |
| TMEM41B | 1.081 | 9.2955 | 104.18 | 6E-19 | 2E-17 | 0.3928571 | n.a. |
| MAPK6 | 1.147 | 9.2551 | 302.39 | 6E-19 | 2E-17 | 0.2738095 | n.a. |
| GPATCH11 | 1.329 | 9.3358 | 77.723 | 1E-18 | 3E-17 | 0.6547619 | n.a. |
| GPR15 | 1.007 | 9.265 | 278.25 | 1E-18 | 4E-17 | 0.3452381 | surface |
| ENSECAG00000024071 | 1.231 | 9.3207 | 112.36 | 2E-18 | 5E-17 | 0.4404762 | n.a. |
| GOLT1B | 1.4 | 9.296 | 120.99 | 5E-18 | 1E-16 | 0.4047619 | n.a. |
| MAP4K4 | 1.404 | 9.2623 | 321.17 | 1E-17 | 3E-16 | 0.3214286 | n.a. |
| UBXN4 | 1.243 | 9.4336 | 71.921 | 2E-17 | 6E-16 | 0.8095238 | n.a. |
| DIPK1A | 1.301 | 9.2939 | 86.732 | 3E-17 | 8E-16 | 0.7142857 | n.a. |
| BET1L | 1.274 | 9.3466 | 76.735 | 6E-17 | 1E-15 | 0.6428571 | n.a. |
| DDRGK1 | 1.25 | 9.3153 | 93.243 | 6E-17 | 2E-15 | 0.5595238 | n.a. |
| DPP4 | 1.182 | 9.2884 | 109.49 | 1E-16 | 2E-15 | 0.5119048 | surface |
| ELL2 | 1.264 | 9.2391 | 883.12 | 1E-16 | 3E-15 | 0.2857143 | n.a. |
| ERGIC3 | 1.187 | 9.4683 | 68.481 | 1E-16 | 3E-15 | 0.75 | n.a. |
| AUP1 | 1.191 | 9.3833 | 68.421 | 1E-16 | 3E-15 | 0.6428571 | n.a. |
| ARFGAP3 | 1.204 | 9.2974 | 84.484 | 2E-16 | 4E-15 | 0.4642857 | n.a. |
| TMEM14A | 1.003 | 9.3121 | 132.2 | 3E-16 | 7E-15 | 0.3571429 | n.a. |
| GMPPA | 1.402 | 9.274 | 147.65 | 4E-16 | 1E-14 | 0.5 | n.a. |
| MKRN1 | 1.19 | 9.3263 | 77.434 | 4E-16 | 1E-14 | 0.5357143 | n.a. |
| YIPF2 | 1.252 | 9.3109 | 97.327 | 6E-16 | 1E-14 | 0.4642857 | n.a. |
| VPS37B | 1.269 | 9.2968 | 100.14 | 6E-16 | 1E-14 | 0.5238095 | n.a. |
| NCF1 | 1.182 | 9.2933 | 249.49 | 7E-16 | 2E-14 | 0.4642857 | n.a. |
| TMEM208 | 1.247 | 9.3185 | 83.311 | 9E-16 | 2E-14 | 0.5833333 | n.a. |

| | | | | | | | |
|---|---|---|---|---|---|---|---|
| PSENEN | 1.157 | 9.5496 | 64.526 | 1E-15 | 2E-14 | 0.8333333 | n.a. |
| ENSECAG00000026829 | 1.186 | 9.535 | 64.499 | 1E-15 | 2E-14 | 0.797619 | n.a. |
| ENSECAG00000018072 | 1.094 | 9.2931 | 64.246 | 1E-15 | 3E-14 | 0.4642857 | n.a. |
| PDXDC1 | 1.264 | 9.2633 | 146.85 | 1E-15 | 3E-14 | 0.4880952 | n.a. |
| DNAJC25 | 1.334 | 9.2588 | 215.87 | 1E-15 | 3E-14 | 0.3214286 | n.a. |
| AGA | 1.216 | 9.3481 | 99.449 | 1E-15 | 3E-14 | 0.4880952 | n.a. |
| SERP1 | 1.162 | 9.3042 | 119.35 | 2E-15 | 4E-14 | 0.452381 | n.a. |
| MAGT1 | 1.157 | 9.334 | 70.993 | 2E-15 | 4E-14 | 0.4880952 | n.a. |
| TMED7 | 1.155 | 9.3905 | 63.111 | 2E-15 | 4E-14 | 0.5833333 | surface |
| TBL2 | 1.388 | 9.2545 | 246.61 | 4E-15 | 9E-14 | 0.3452381 | n.a. |
| NUDT4 | 1.256 | 9.2991 | 138.85 | 5E-15 | 1E-13 | 0.297619 | n.a. |
| TMEM165 | 1.13 | 9.4054 | 60.873 | 6E-15 | 1E-13 | 0.6190476 | n.a. |
| GPX7 | 1.254 | 9.2463 | 374.93 | 7E-15 | 1E-13 | 0.3809524 | n.a. |
| MFSD11 | 1.059 | 9.259 | 244.8 | 7E-15 | 2E-13 | 0.297619 | surface |
| ARF4 | 1.139 | 9.4745 | 59.564 | 1E-14 | 2E-13 | 0.7857143 | n.a. |
| PREB | 1.082 | 9.2646 | 131.09 | 1E-14 | 3E-13 | 0.3452381 | n.a. |
| CASP8 | 1.452 | 9.2993 | 118.64 | 2E-14 | 3E-13 | 0.3809524 | n.a. |
| REEP3 | 1.165 | 9.3317 | 84.117 | 2E-14 | 3E-13 | 0.4285714 | n.a. |
| BNIP3L | 1.489 | 9.4595 | 80.28 | 2E-14 | 4E-13 | 0.4761905 | n.a. |
| PAXX | 1.09 | 9.3487 | 58.32 | 2E-14 | 5E-13 | 0.5595238 | n.a. |
| BCAP29 | 1.095 | 9.3956 | 60.536 | 3E-14 | 6E-13 | 0.5357143 | n.a. |
| SRP19 | 1.114 | 9.5262 | 57.592 | 3E-14 | 7E-13 | 0.7857143 | n.a. |
| ENSECAG00000012339 | 1.058 | 9.328 | 59.535 | 4E-14 | 7E-13 | 0.5238095 | n.a. |
| GBP5 | 1.068 | 9.7163 | 57.056 | 4E-14 | 8E-13 | 0.6666667 | n.a. |
| REEP5 | 1.063 | 9.6201 | 56.528 | 6E-14 | 1E-12 | 0.8214286 | n.a. |
| COPE | 1.056 | 9.5903 | 55.983 | 7E-14 | 1E-12 | 0.8452381 | n.a. |
| TCEAL9 | 1.221 | 9.2671 | 113.42 | 9E-14 | 2E-12 | 0.3571429 | n.a. |
| STRADB | 1.431 | 9.2591 | 191.31 | 1E-13 | 2E-12 | 0.3571429 | n.a. |
| SNRPN | 1.136 | 9.4022 | 55.453 | 1E-13 | 2E-12 | 0.6785714 | n.a. |
| NTAN1 | 1.058 | 9.412 | 55.091 | 1E-13 | 2E-12 | 0.702381 | n.a. |
| NGLY1 | 1.274 | 9.2901 | 98.115 | 1E-13 | 3E-12 | 0.4404762 | n.a. |
| GORASP2 | 1.194 | 9.3269 | 71.219 | 1E-13 | 3E-12 | 0.5952381 | n.a. |
| VDAC3 | 1.152 | 9.3496 | 54.689 | 1E-13 | 3E-12 | 0.5833333 | n.a. |
| PRKCSH | 1.144 | 9.2832 | 92.128 | 2E-13 | 3E-12 | 0.4166667 | n.a. |
| SLC30A7 | 1.058 | 9.289 | 106.44 | 2E-13 | 3E-12 | 0.2857143 | n.a. |
| ERP44 | 1.098 | 9.4288 | 54.005 | 2E-13 | 4E-12 | 0.797619 | n.a. |
| GINM1 | 1.003 | 9.3701 | 57.466 | 2E-13 | 4E-12 | 0.5357143 | surface |
| YARS | 1.119 | 9.2689 | 151.67 | 2E-13 | 4E-12 | 0.3571429 | n.a. |
| SEC63 | 1.088 | 9.4017 | 54.122 | 3E-13 | 5E-12 | 0.5357143 | n.a. |
| EI24 | 1.149 | 9.2692 | 145.43 | 3E-13 | 5E-12 | 0.2857143 | n.a. |
| PPP3CC | 1.122 | 9.3183 | 67.128 | 5E-13 | 8E-12 | 0.5357143 | n.a. |
| DERL2 | 1.055 | 9.3504 | 52.275 | 5E-13 | 8E-12 | 0.5714286 | n.a. |
| YIPF3 | 1.101 | 9.4584 | 52.211 | 5E-13 | 9E-12 | 0.75 | n.a. |
| TRABD | 1.013 | 9.361 | 59.471 | 5E-13 | 9E-12 | 0.5714286 | n.a. |
| NME1 | 1.107 | 9.4477 | 51.427 | 8E-13 | 1E-11 | 0.6904762 | n.a. |
| STX5 | 1.013 | 9.3118 | 54.426 | 1E-12 | 2E-11 | 0.6071429 | n.a. |
| TMEM263 | 1.216 | 9.331 | 85.297 | 3E-12 | 5E-11 | 0.4047619 | n.a. |
| TMCO1 | 1.079 | 9.4392 | 48.713 | 3E-12 | 5E-11 | 0.6547619 | n.a. |

| | | | | | | | |
|---|---|---|---|---|---|---|---|
| EHBP1 | 1.146 | 9.326 | 101.01 | 3E-12 | 5E-11 | 0.3214286 | n.a. |
| YIF1A | 1.201 | 9.2778 | 104.01 | 3E-12 | 5E-11 | 0.3452381 | n.a. |
| DAD1 | 1.053 | 9.4477 | 48.289 | 4E-12 | 6E-11 | 0.6785714 | n.a. |
| CDKN2C | 1.018 | 9.2805 | 121.85 | 4E-12 | 6E-11 | 0.3452381 | n.a. |
| FAHD2A | 1.05 | 9.2946 | 70.351 | 5E-12 | 7E-11 | 0.4166667 | n.a. |
| SMCHD1 | 1.085 | 9.3994 | 49.652 | 5E-12 | 7E-11 | 0.6666667 | n.a. |
| CHPF2 | 1.027 | 9.254 | 236.79 | 5E-12 | 8E-11 | 0.2619048 | n.a. |
| LAPTM4A | 1.033 | 9.5901 | 47.702 | 5E-12 | 8E-11 | 0.7619048 | n.a. |
| UBA5 | 1.118 | 9.2783 | 91.426 | 5E-12 | 8E-11 | 0.4761905 | n.a. |
| GALE | 1.036 | 9.2603 | 100.32 | 7E-12 | 1E-10 | 0.3214286 | n.a. |
| MESD | 1.074 | 9.4258 | 46.864 | 8E-12 | 1E-10 | 0.6190476 | n.a. |
| GPATCH2 | 1.262 | 9.2506 | 245.64 | 1E-11 | 2E-10 | 0.3333333 | n.a. |
| EIF2D | 1.356 | 9.275 | 134.58 | 1E-11 | 2E-10 | 0.4166667 | n.a. |
| PLPP5 | 1.223 | 9.2785 | 115.67 | 1E-11 | 2E-10 | 0.4166667 | n.a. |
| PTPA | 1.156 | 9.292 | 75.212 | 2E-11 | 3E-10 | 0.3809524 | n.a. |
| TOP1 | 1.02 | 9.4172 | 44.445 | 3E-11 | 4E-10 | 0.7261905 | n.a. |
| ARFIP2 | 1.089 | 9.2783 | 70.603 | 3E-11 | 4E-10 | 0.452381 | n.a. |
| ENSECAG00000036054 | 1.048 | 9.2783 | 72.997 | 3E-11 | 4E-10 | 0.4404762 | n.a. |
| RF00569 | 1.002 | 9.279 | 45.48 | 5E-11 | 8E-10 | 0.452381 | n.a. |
| ALG12 | 1.029 | 9.268 | 90.652 | 6E-11 | 8E-10 | 0.3690476 | n.a. |
| HIST1H1C | 1.155 | 9.2575 | 146.73 | 1E-10 | 2E-09 | 0.2738095 | n.a. |
| TIMM23 | 1.031 | 9.2994 | 54.275 | 1E-10 | 2E-09 | 0.5357143 | n.a. |
| SDF4 | 1.008 | 9.343 | 50.907 | 1E-10 | 2E-09 | 0.4642857 | n.a. |
| SEC23B | 1.039 | 9.2732 | 88.618 | 3E-10 | 4E-09 | 0.297619 | n.a. |
| HIST1H1D | 1.106 | 9.281 | 105.66 | 3E-10 | 4E-09 | 0.3095238 | n.a. |
| ENSECAG00000032710 | 1.012 | 9.3761 | 52.935 | 4E-10 | 5E-09 | 0.3452381 | n.a. |
| POU2AF1 | 1.205 | 9.2923 | 113.53 | 4E-10 | 5E-09 | 0.7738095 | n.a. |
| SEC31A | 1.029 | 9.2996 | 67.35 | 6E-10 | 8E-09 | 0.3809524 | n.a. |
| PRKD2 | 1.001 | 9.3027 | 52.958 | 1E-09 | 2E-08 | 0.5119048 | n.a. |
| TUBB4A | 1.11 | 9.3253 | 67.437 | 3E-09 | 4E-08 | 0.4404762 | n.a. |
| SETDB2 | 1.076 | 9.2506 | 175.69 | 4E-09 | 5E-08 | 0.2619048 | n.a. |
| ELK3 | 1.036 | 9.3026 | 71.892 | 4E-08 | 4E-07 | 0.4047619 | n.a. |
| TNFRSF13B | 1.086 | 9.2609 | 60.786 | 6E-08 | 6E-07 | 0.6190476 | surface |
| SS18L2 | 1.173 | 9.3178 | 59.331 | 1E-06 | 1E-05 | 0.3214286 | n.a. |

| genes | logFC | logCPM | F | PValue | FDR | percent.exp | Surfacome.Label |
|---|---|---|---|---|---|---|---|
| ENSECAG00000026891 | 3.979 | 9.2736 | 3590.8 | 2E-287 | 2E-283 | 0.8857143 | n.a. |
| STMN1 | 4.289 | 9.7683 | 1313.4 | 3E-282 | 1E-278 | 1 | n.a. |
| UBE2C | 4.288 | 9.2563 | 5409.2 | 1E-266 | 4E-263 | 0.8857143 | n.a. |
| HMGB2 | 3.294 | 10.129 | 702.07 | 4E-153 | 7E-150 | 1 | n.a. |
| H2AFX | 3.321 | 9.3048 | 588.8 | 5E-129 | 9E-126 | 0.8857143 | n.a. |
| NUSAP1 | 3.463 | 9.2433 | 6401.6 | 2E-126 | 3E-123 | 0.6285714 | n.a. |
| TOP2A | 3.551 | 9.245 | 5054 | 4E-123 | 5E-120 | 0.6571429 | n.a. |
| ENSECAG00000036105 | 3.159 | 9.2441 | 6693.7 | 8E-107 | 8E-104 | 0.8 | n.a. |
| CENPF | 3.216 | 9.2409 | 4487.2 | 2E-104 | 2E-101 | 0.6 | n.a. |
| RRM2 | 3.04 | 9.252 | 1823.8 | 1E-99 | 9E-97 | 0.8571429 | n.a. |
| H2AFZ | 3.082 | 9.5191 | 415.31 | 9E-92 | 6E-89 | 1 | n.a. |
| CDCA3 | 2.729 | 9.2462 | 3092.2 | 1E-91 | 8E-89 | 0.6857143 | n.a. |
| CKS2 | 3.109 | 9.2847 | 410.33 | 1E-90 | 6E-88 | 0.8 | n.a. |
| CCNB1 | 3.022 | 9.2437 | 2195.2 | 1E-83 | 7E-81 | 0.6285714 | n.a. |
| TUBA1A | 2.664 | 10.091 | 372.41 | 2E-82 | 8E-80 | 0.9714286 | n.a. |
| TPX2 | 2.876 | 9.2428 | 3700.5 | 1E-81 | 7E-79 | 0.8285714 | n.a. |
| BIRC5 | 2.783 | 9.243 | 4506.9 | 2E-77 | 8E-75 | 0.6857143 | n.a. |
| CDC20 | 2.799 | 9.2401 | 4816.9 | 6E-77 | 3E-74 | 0.6 | n.a. |
| CCNB2 | 2.76 | 9.2382 | 6246 | 2E-74 | 1E-71 | 0.6285714 | n.a. |
| CCNA2 | 2.911 | 9.2425 | 5213.5 | 2E-72 | 6E-70 | 0.6857143 | n.a. |
| CENPA | 2.608 | 9.2413 | 3263.4 | 6E-70 | 2E-67 | 0.5142857 | n.a. |
| eca-mir-8997 | 2.669 | 9.2408 | 5168.9 | 3E-68 | 1E-65 | 0.6571429 | n.a. |
| ASF1B | 2.599 | 9.241 | 4267.6 | 4E-67 | 1E-64 | 0.6 | n.a. |
| CDCA8 | 2.746 | 9.2406 | 3353.3 | 1E-66 | 4E-64 | 0.6285714 | n.a. |
| ENSECAG00000037332 | 2.344 | 10.054 | 292.29 | 3E-65 | 9E-63 | 1 | n.a. |
| HMGB1 | 2.11 | 10.086 | 257.91 | 8E-58 | 2E-55 | 1 | n.a. |
| TYMS | 2.715 | 9.3134 | 253.17 | 8E-57 | 2E-54 | 0.8857143 | n.a. |
| TACC3 | 2.565 | 9.2769 | 345.05 | 7E-56 | 2E-53 | 0.8 | n.a. |
| CDKN3 | 2.249 | 9.2388 | 3943.9 | 2E-53 | 4E-51 | 0.5142857 | n.a. |
| SPC25 | 2.609 | 9.2421 | 1918.1 | 4E-53 | 1E-50 | 0.7142857 | n.a. |
| ENSECAG00000039428 | 1.293 | 11.047 | 203.56 | 5E-46 | 1E-43 | 1 | n.a. |
| CENPE | 2.391 | 9.2392 | 3386.3 | 6E-46 | 2E-43 | 0.5428571 | n.a. |
| TCF19 | 2.313 | 9.2869 | 275.34 | 2E-44 | 5E-42 | 0.7142857 | n.a. |
| HMGB3 | 2.41 | 9.2519 | 821.95 | 4E-43 | 8E-41 | 0.6571429 | n.a. |
| CDK1 | 2.356 | 9.2397 | 2570.9 | 6E-43 | 1E-40 | 0.4857143 | n.a. |
| UBE2S | 2.427 | 9.3494 | 185.02 | 5E-42 | 1E-39 | 0.7714286 | n.a. |
| DEPDC1B | 2.1 | 9.2381 | 3244.3 | 2E-41 | 3E-39 | 0.4857143 | n.a. |
| PCNA | 2.372 | 9.3865 | 180.55 | 5E-41 | 9E-39 | 0.7428571 | n.a. |
| MXD3 | 2.373 | 9.2429 | 1068.9 | 2E-40 | 3E-38 | 0.5714286 | n.a. |
| PBK | 2.04 | 9.2383 | 3318.1 | 2E-40 | 4E-38 | 0.5142857 | n.a. |
| SGO1 | 2.215 | 9.2375 | 4843.3 | 4E-40 | 7E-38 | 0.5142857 | n.a. |
| CENPM | 2.574 | 9.247 | 506.13 | 1E-39 | 2E-37 | 0.8285714 | n.a. |
| KIF11 | 2.206 | 9.2384 | 2522.7 | 2E-39 | 3E-37 | 0.5714286 | n.a. |
| ENSECAG00000033690 | 2.371 | 9.2632 | 272.11 | 4E-38 | 7E-36 | 0.6857143 | n.a. |
| NUF2 | 2.156 | 9.2372 | 3966.7 | 2E-37 | 3E-35 | 0.5428571 | n.a. |
| TUBB4A | 2.368 | 9.3253 | 162.46 | 4E-37 | 6E-35 | 0.7428571 | n.a. |
| AURKB | 1.978 | 9.2364 | 2953.9 | 2E-36 | 2E-34 | 0.3714286 | n.a. |

| | | | | | | | |
|---|---|---|---|---|---|---|---|
| UCP2 | 1.804 | 9.9052 | 156.35 | 8E-36 | 1E-33 | 0.9714286 | n.a. |
| KIF22 | 2.154 | 9.2545 | 450.48 | 5E-35 | 7E-33 | 0.6571429 | n.a. |
| ARHGEF39 | 2.206 | 9.2399 | 1369.5 | 1E-34 | 2E-32 | 0.5714286 | n.a. |
| RHNO1 | 1.917 | 9.2779 | 351.42 | 2E-34 | 2E-32 | 0.6 | n.a. |
| KIFC1 | 2.223 | 9.2409 | 1955.1 | 3E-34 | 4E-32 | 0.6285714 | n.a. |
| REEP4 | 2.114 | 9.2747 | 257 | 3E-34 | 4E-32 | 0.6857143 | n.a. |
| CDC25B | 1.922 | 9.3048 | 160.34 | 5E-34 | 7E-32 | 0.7142857 | n.a. |
| CKS1B | 2.221 | 9.345 | 146.88 | 1E-33 | 1E-31 | 0.7714286 | n.a. |
| ENSECAG00000035897 | 2.078 | 9.3428 | 146.27 | 1E-33 | 2E-31 | 0.8571429 | n.a. |
| LMNB1 | 2.078 | 9.2729 | 222.74 | 2E-33 | 2E-31 | 0.8 | n.a. |
| ID3 | 1.989 | 9.6697 | 138.69 | 6E-32 | 7E-30 | 0.8285714 | n.a. |
| TROAP | 1.979 | 9.2354 | 5113.8 | 9E-32 | 1E-29 | 0.4 | n.a. |
| HMMR | 2.332 | 9.2372 | 2526.2 | 1E-31 | 2E-29 | 0.5142857 | n.a. |
| UBE2T | 1.952 | 9.2403 | 1292.4 | 2E-31 | 2E-29 | 0.4285714 | n.a. |
| PLK1 | 1.974 | 9.239 | 1923 | 5E-31 | 6E-29 | 0.3428571 | n.a. |
| ACTG1 | 1.555 | 10.745 | 128.74 | 9E-30 | 1E-27 | 1 | n.a. |
| SMC4 | 2.139 | 9.3283 | 122.16 | 2E-28 | 3E-26 | 0.9428571 | n.a. |
| CKAP2 | 1.719 | 9.2353 | 4988.4 | 1E-27 | 1E-25 | 0.3428571 | n.a. |
| MYBL2 | 1.803 | 9.2383 | 3741.9 | 1E-27 | 1E-25 | 0.4 | n.a. |
| H2AFV | 1.737 | 9.7995 | 118.54 | 1E-27 | 2E-25 | 1 | n.a. |
| ENSECAG00000038338 | 1.733 | 9.2755 | 171.37 | 2E-26 | 2E-24 | 0.7142857 | n.a. |
| MIS18A | 1.859 | 9.2371 | 2537.9 | 5E-26 | 5E-24 | 0.5142857 | n.a. |
| NRM | 1.809 | 9.2939 | 116.04 | 5E-26 | 5E-24 | 0.7142857 | n.a. |
| CEP55 | 1.706 | 9.2368 | 2007.7 | 6E-26 | 6E-24 | 0.4 | n.a. |
| CENPP | 2.088 | 9.2531 | 309.28 | 1E-25 | 1E-23 | 0.7428571 | n.a. |
| S100A6 | 1.671 | 10.437 | 106.95 | 5E-25 | 5E-23 | 0.8857143 | n.a. |
| NEIL3 | 1.634 | 9.2353 | 157.76 | 1E-24 | 1E-22 | 0.3142857 | n.a. |
| FBXO5 | 1.688 | 9.2403 | 1477.9 | 1E-24 | 1E-22 | 0.4 | n.a. |
| ENSECAG00000012818 | 1.309 | 9.2495 | 503.94 | 3E-24 | 3E-22 | 0.3142857 | n.a. |
| KIF15 | 1.901 | 9.2382 | 1647.9 | 3E-24 | 3E-22 | 0.4571429 | n.a. |
| CENPW | 2.122 | 9.242 | 1036 | 4E-24 | 3E-22 | 0.5142857 | n.a. |
| CDKN2C | 1.785 | 9.2805 | 184.89 | 4E-24 | 4E-22 | 0.6571429 | n.a. |
| PKMYT1 | 1.94 | 9.2511 | 213.71 | 1E-23 | 1E-21 | 0.6571429 | n.a. |
| KIF23 | 1.579 | 9.2359 | 2554.4 | 1E-23 | 1E-21 | 0.4285714 | n.a. |
| CLSPN | 1.823 | 9.2395 | 2046.6 | 3E-23 | 2E-21 | 0.4857143 | n.a. |
| KNSTRN | 2.085 | 9.2554 | 250.5 | 6E-23 | 5E-21 | 0.6857143 | n.a. |
| MIS18BP1 | 1.945 | 9.2452 | 612.33 | 2E-22 | 2E-20 | 0.5142857 | n.a. |
| ZWINT | 1.637 | 9.2392 | 1129 | 5E-22 | 5E-20 | 0.4571429 | n.a. |
| TLX2 | 1.911 | 9.2358 | 2308.1 | 8E-22 | 7E-20 | 0.4571429 | n.a. |
| NCAPG2 | 1.911 | 9.2424 | 600.84 | 1E-21 | 1E-19 | 0.4285714 | n.a. |
| CDC45 | 1.482 | 9.2362 | 2584.3 | 5E-21 | 4E-19 | 0.3428571 | n.a. |
| KIF20B | 1.903 | 9.2938 | 107.35 | 8E-21 | 6E-19 | 0.6857143 | n.a. |
| EZH2 | 1.667 | 9.245 | 512.66 | 1E-20 | 1E-18 | 0.4857143 | n.a. |
| BARD1 | 1.767 | 9.2398 | 1274 | 3E-20 | 2E-18 | 0.5714286 | n.a. |
| NCAPG | 1.859 | 9.243 | 442.68 | 4E-20 | 3E-18 | 0.5142857 | n.a. |
| SGO2 | 1.738 | 9.2385 | 1145.5 | 5E-20 | 3E-18 | 0.4 | n.a. |
| ASPM | 1.362 | 9.2353 | 3404.1 | 5E-20 | 4E-18 | 0.2571429 | n.a. |
| UHRF1 | 1.344 | 9.2365 | 2390.3 | 6E-20 | 4E-18 | 0.2571429 | n.a. |

| | | | | | | | |
|---|---|---|---|---|---|---|---|
| BUB1 | 1.443 | 9.2359 | 2439.3 | 7E-20 | 5E-18 | 0.3714286 | n.a. |
| MTFR2 | 1.657 | 9.2373 | 2009.1 | 7E-20 | 5E-18 | 0.4 | n.a. |
| KIF18B | 1.332 | 9.2346 | 1824 | 8E-20 | 6E-18 | 0.3714286 | n.a. |
| SHCBP1 | 1.223 | 9.2367 | 2689 | 6E-19 | 4E-17 | 0.3714286 | n.a. |
| PRDX2 | 1.326 | 9.2964 | 78.105 | 1E-18 | 7E-17 | 0.7142857 | n.a. |
| MNS1 | 1.669 | 9.2425 | 526.41 | 1E-18 | 9E-17 | 0.4857143 | n.a. |
| KNL1 | 1.952 | 9.2399 | 768.3 | 2E-18 | 1E-16 | 0.4857143 | n.a. |
| PRC1 | 1.745 | 9.24 | 817.65 | 2E-18 | 1E-16 | 0.4285714 | n.a. |
| CDKN2A | 1.525 | 9.249 | 276.95 | 3E-18 | 2E-16 | 0.4 | n.a. |
| MCM7 | 1.728 | 9.3416 | 75.494 | 4E-18 | 2E-16 | 0.6857143 | n.a. |
| AURKA | 1.845 | 9.247 | 298.96 | 9E-18 | 6E-16 | 0.5714286 | n.a. |
| PLK4 | 1.653 | 9.2404 | 563.08 | 9E-18 | 6E-16 | 0.4 | n.a. |
| ENSECAG00000019780 | 1.43 | 9.2353 | 2937.1 | 1E-17 | 8E-16 | 0.3428571 | n.a. |
| NEK2 | 1.651 | 9.2368 | 1326.1 | 2E-17 | 1E-15 | 0.4 | n.a. |
| ITGA4 | 1.069 | 9.465 | 71.77 | 3E-17 | 2E-15 | 0.7428571 | surface |
| CIP2A | 1.774 | 9.25 | 251.4 | 8E-17 | 5E-15 | 0.4571429 | n.a. |
| SSBP2 | 1.26 | 9.2735 | 173.4 | 9E-17 | 5E-15 | 0.3142857 | n.a. |
| TUBA4A | 1.559 | 9.5479 | 67.566 | 2E-16 | 1E-14 | 0.8857143 | n.a. |
| CIT | 1.298 | 9.2349 | 2775.8 | 3E-16 | 2E-14 | 0.3142857 | n.a. |
| FEN1 | 1.767 | 9.2609 | 157.3 | 4E-16 | 2E-14 | 0.6571429 | n.a. |
| C1H15orf48 | 1.054 | 9.6929 | 65.666 | 6E-16 | 3E-14 | 0.4857143 | n.a. |
| ACOT7 | 1.417 | 9.2472 | 294.43 | 6E-16 | 3E-14 | 0.4571429 | n.a. |
| CDCA2 | 1.175 | 9.2358 | 1594.5 | 9E-16 | 5E-14 | 0.2857143 | n.a. |
| SPC24 | 1.477 | 9.2428 | 486.42 | 1E-15 | 7E-14 | 0.4285714 | n.a. |
| SKA3 | 1.278 | 9.2375 | 872.3 | 2E-15 | 1E-13 | 0.3714286 | n.a. |
| CCNE2 | 1.253 | 9.2372 | 1414 | 3E-15 | 1E-13 | 0.3428571 | n.a. |
| ENSECAG00000033471 | 1.493 | 9.244 | 325.51 | 3E-15 | 2E-13 | 0.5428571 | n.a. |
| CDCA7 | 1.576 | 9.2634 | 79.122 | 3E-15 | 2E-13 | 0.4571429 | n.a. |
| INCENP | 1.606 | 9.2578 | 211.16 | 4E-15 | 2E-13 | 0.4571429 | n.a. |
| DEK | 1.486 | 9.6176 | 61.774 | 4E-15 | 2E-13 | 0.8571429 | n.a. |
| E2F2 | 1.261 | 9.2429 | 483.51 | 4E-15 | 2E-13 | 0.4285714 | n.a. |
| WEE1 | 1.276 | 9.2393 | 842.25 | 6E-15 | 3E-13 | 0.4 | n.a. |
| SMC1A | 1.525 | 9.3103 | 66.06 | 6E-15 | 3E-13 | 0.5428571 | n.a. |
| POC1A | 1.414 | 9.2936 | 65.246 | 7E-15 | 4E-13 | 0.7428571 | n.a. |
| TPI1 | 1.43 | 9.5703 | 60.498 | 8E-15 | 4E-13 | 0.9428571 | n.a. |
| CKAP2L | 1.471 | 9.2529 | 209.55 | 1E-14 | 6E-13 | 0.4 | n.a. |
| VIM | 1.012 | 11.286 | 59.437 | 1E-14 | 7E-13 | 1 | n.a. |
| RAD21 | 1.566 | 9.3865 | 58.968 | 2E-14 | 8E-13 | 0.7142857 | n.a. |
| CHAF1A | 1.268 | 9.2395 | 701.32 | 6E-14 | 3E-12 | 0.3142857 | n.a. |
| GPSM2 | 1.361 | 9.2363 | 1616.1 | 6E-14 | 3E-12 | 0.3142857 | n.a. |
| S100A5 | 1.256 | 9.5834 | 55.3 | 1E-13 | 5E-12 | 0.4285714 | n.a. |
| GTSE1 | 1.249 | 9.2367 | 1641.5 | 1E-13 | 7E-12 | 0.2857143 | n.a. |
| NCAPH | 1.125 | 9.2367 | 1267.1 | 2E-13 | 7E-12 | 0.3428571 | n.a. |
| ENSECAG00000020532 | 1.242 | 9.7393 | 53.501 | 3E-13 | 1E-11 | 1 | n.a. |
| NDC80 | 1.1 | 9.2498 | 307.31 | 3E-13 | 1E-11 | 0.3428571 | n.a. |
| RFC3 | 1.517 | 9.2557 | 128.66 | 3E-13 | 1E-11 | 0.5714286 | n.a. |
| EXO1 | 1.141 | 9.2351 | 1891.8 | 3E-13 | 2E-11 | 0.3142857 | n.a. |
| ATAD2 | 1.559 | 9.2578 | 104.36 | 4E-13 | 2E-11 | 0.5714286 | n.a. |

| | | | | | | | |
|---|---|---|---|---|---|---|---|
| ENSECAG00000018918 | 1.505 | 9.237 | 1095.2 | 1E-12 | 4E-11 | 0.3714286 | n.a. |
| CENPL | 1.099 | 9.2357 | 1708.5 | 1E-12 | 5E-11 | 0.2857143 | n.a. |
| CCNF | 1.149 | 9.2356 | 1666.6 | 1E-12 | 5E-11 | 0.2571429 | n.a. |
| KIF4A | 1.345 | 9.2349 | 1982 | 1E-12 | 5E-11 | 0.2857143 | n.a. |
| ENSECAG00000015297 | 1.321 | 9.2596 | 96.163 | 2E-12 | 7E-11 | 0.5428571 | n.a. |
| JPT1 | 1.259 | 9.7243 | 49.688 | 2E-12 | 8E-11 | 0.8 | n.a. |
| CDCA4 | 1.511 | 9.2785 | 111.01 | 2E-12 | 9E-11 | 0.4571429 | n.a. |
| MELK | 1.468 | 9.2366 | 1278.2 | 2E-12 | 9E-11 | 0.2571429 | n.a. |
| GINS1 | 1.377 | 9.2464 | 295.45 | 3E-12 | 1E-10 | 0.3428571 | n.a. |
| HSPA2 | 1.26 | 9.2405 | 516.47 | 5E-12 | 2E-10 | 0.2857143 | n.a. |
| SYNE2 | 1.606 | 9.2424 | 245.15 | 1E-11 | 6E-10 | 0.4 | n.a. |
| CORT | 1.488 | 9.2439 | 211.4 | 2E-11 | 6E-10 | 0.5428571 | n.a. |
| TKT | 1.074 | 9.6069 | 44.989 | 2E-11 | 8E-10 | 0.9142857 | n.a. |
| CALCOCO2 | 1.134 | 9.336 | 44.925 | 2E-11 | 8E-10 | 0.6285714 | n.a. |
| DNMT1 | 1.319 | 9.2975 | 84.549 | 2E-11 | 9E-10 | 0.4571429 | n.a. |
| ORC1 | 1.259 | 9.237 | 923.06 | 3E-11 | 1E-09 | 0.2571429 | n.a. |
| GINS2 | 1.08 | 9.239 | 596.11 | 4E-11 | 2E-09 | 0.3142857 | n.a. |
| H2AFY | 1.217 | 9.3852 | 45.301 | 4E-11 | 2E-09 | 0.6571429 | n.a. |
| DHDDS | 1.241 | 9.3007 | 43.17 | 5E-11 | 2E-09 | 0.5714286 | n.a. |
| NCAPD2 | 1.603 | 9.2437 | 293.8 | 5E-11 | 2E-09 | 0.4857143 | n.a. |
| GLUL | 1.273 | 9.3271 | 115.83 | 6E-11 | 2E-09 | 0.3428571 | n.a. |
| ENSECAG00000038539 | 1.14 | 9.3369 | 64.92 | 7E-11 | 3E-09 | 0.5142857 | n.a. |
| DEGS2 | 1.102 | 9.2366 | 426.1 | 7E-11 | 3E-09 | 0.2571429 | n.a. |
| CENPK | 1.456 | 9.253 | 136.54 | 7E-11 | 3E-09 | 0.4571429 | n.a. |
| CD69 | 1.202 | 9.4382 | 45.682 | 9E-11 | 3E-09 | 0.2857143 | surface |
| ORC6 | 1.445 | 9.2453 | 224.38 | 1E-10 | 5E-09 | 0.4 | n.a. |
| TRIM59 | 1.287 | 9.3315 | 42.69 | 2E-10 | 6E-09 | 0.5714286 | n.a. |
| CD86 | 1.035 | 9.2873 | 152.67 | 2E-10 | 6E-09 | 0.3428571 | surface |
| CENPU | 1.174 | 9.2762 | 83.043 | 3E-10 | 1E-08 | 0.4285714 | n.a. |
| FABP3 | 1.23 | 9.3412 | 39.283 | 4E-10 | 1E-08 | 0.8571429 | n.a. |
| RPA2 | 1.301 | 9.3129 | 40.874 | 9E-10 | 3E-08 | 0.7142857 | n.a. |
| GMNN | 1.412 | 9.2659 | 85.403 | 1E-09 | 3E-08 | 0.5428571 | n.a. |
| UBE2E3 | 1.168 | 9.3915 | 35.426 | 3E-09 | 9E-08 | 1 | n.a. |
| ENSECAG00000009520 | 1.183 | 9.3369 | 34.377 | 5E-09 | 1E-07 | 0.6857143 | n.a. |
| ENSECAG00000036180 | 1.098 | 9.4114 | 34.037 | 5E-09 | 2E-07 | 0.9142857 | n.a. |
| NCAPH2 | 1.148 | 9.2984 | 36.722 | 7E-09 | 2E-07 | 0.6285714 | n.a. |
| H1FX | 1.183 | 9.3523 | 34.339 | 7E-09 | 2E-07 | 0.5714286 | n.a. |
| BRCA2 | 1.077 | 9.2516 | 113.02 | 7E-09 | 2E-07 | 0.2857143 | n.a. |
| DBNL | 1.139 | 9.4448 | 33.413 | 8E-09 | 2E-07 | 0.8857143 | n.a. |
| ARL6IP1 | 1.227 | 9.5522 | 33.298 | 8E-09 | 2E-07 | 0.6285714 | n.a. |
| ENSECAG00000034985 | 1.06 | 9.3342 | 32.527 | 1E-08 | 3E-07 | 0.4 | n.a. |
| SSBP3 | 1.15 | 9.2712 | 65.67 | 2E-08 | 4E-07 | 0.3714286 | n.a. |
| NUCKS1 | 1.061 | 9.7354 | 31.958 | 2E-08 | 4E-07 | 0.9714286 | n.a. |
| MAZ | 1.215 | 9.275 | 76.504 | 2E-08 | 7E-07 | 0.4 | n.a. |
| GSTK1 | 1.34 | 9.2598 | 81.742 | 3E-08 | 7E-07 | 0.4857143 | n.a. |
| ENSECAG00000035662 | 1.136 | 9.4047 | 30.746 | 3E-08 | 8E-07 | 0.8857143 | n.a. |
| SMAGP | 1.056 | 9.3065 | 30.487 | 3E-08 | 9E-07 | 0.6285714 | n.a. |
| DDX39A | 1.192 | 9.3461 | 30.168 | 4E-08 | 1E-06 | 0.7714286 | n.a. |

| | | | | | | | |
|---|---|---|---|---|---|---|---|
| MCM4 | 1.061 | 9.262 | 61.738 | 4E-08 | 1E-06 | 0.4 | n.a. |
| RAD18 | 1.176 | 9.2966 | 48.325 | 4E-08 | 1E-06 | 0.4857143 | n.a. |
| PALM | 1.142 | 9.2423 | 147.61 | 4E-08 | 1E-06 | 0.2857143 | n.a. |
| ODF2 | 1.194 | 9.2583 | 93.728 | 5E-08 | 1E-06 | 0.3714286 | n.a. |
| HIST1H1D | 1.071 | 9.281 | 51.632 | 6E-08 | 1E-06 | 0.4285714 | n.a. |
| MCUB | 1.244 | 9.3094 | 56.587 | 9E-08 | 2E-06 | 0.5428571 | n.a. |
| ANP32E | 1.184 | 9.2631 | 51.953 | 1E-07 | 3E-06 | 0.3428571 | n.a. |
| SSNA1 | 1.032 | 9.3399 | 27.928 | 1E-07 | 3E-06 | 0.5714286 | n.a. |
| ENSECAG00000017935 | 1.014 | 9.3545 | 27.278 | 2E-07 | 4E-06 | 0.5714286 | n.a. |
| FDFT1 | 1.032 | 9.3105 | 26.87 | 2E-07 | 5E-06 | 0.6 | n.a. |
| CCDC34 | 1.444 | 9.2605 | 118.92 | 2E-07 | 6E-06 | 0.4 | n.a. |
| WRAP73 | 1.183 | 9.2804 | 48.245 | 2E-07 | 6E-06 | 0.4285714 | n.a. |
| CBFB | 1.096 | 9.3467 | 31.661 | 3E-07 | 6E-06 | 0.5428571 | n.a. |
| CKAP5 | 1.178 | 9.2595 | 94.139 | 3E-07 | 8E-06 | 0.3142857 | n.a. |
| KIF2A | 1.019 | 9.3544 | 25.975 | 3E-07 | 8E-06 | 0.6857143 | n.a. |
| ENSECAG00000026827 | 1.148 | 9.2436 | 141.05 | 4E-07 | 9E-06 | 0.4 | n.a. |
| ENSECAG00000005186 | 1.055 | 9.2505 | 94.812 | 4E-07 | 9E-06 | 0.3428571 | n.a. |
| SMC3 | 1.084 | 9.4311 | 24.884 | 6E-07 | 1E-05 | 0.8571429 | n.a. |
| IDI1 | 1.02 | 9.4385 | 24.85 | 6E-07 | 1E-05 | 0.4857143 | n.a. |
| FDPS | 1.015 | 9.2939 | 32.758 | 7E-07 | 2E-05 | 0.5142857 | n.a. |
| PROSER3 | 1.039 | 9.2431 | 220.17 | 8E-07 | 2E-05 | 0.2571429 | n.a. |
| MCM6 | 1.149 | 9.313 | 24.398 | 8E-07 | 2E-05 | 0.6 | n.a. |
| KCNS3 | 1.07 | 9.2415 | 174.09 | 1E-06 | 2E-05 | 0.3428571 | n.a. |
| USP1 | 1.017 | 9.3606 | 22.761 | 2E-06 | 4E-05 | 0.7428571 | n.a. |
| NCAPD3 | 1.097 | 9.2575 | 80.088 | 2E-06 | 4E-05 | 0.3428571 | n.a. |
| HP1BP3 | 1.032 | 9.4292 | 22.197 | 2E-06 | 5E-05 | 0.8571429 | n.a. |
| ENSECAG00000020955 | 1.294 | 9.2591 | 38.531 | 3E-06 | 6E-05 | 0.6 | n.a. |
| ENSECAG00000006493 | 1.047 | 9.2594 | 54.76 | 3E-06 | 7E-05 | 0.4571429 | n.a. |
| MLF1 | 1.422 | 9.2374 | 402.31 | 4E-06 | 8E-05 | 0.3714286 | n.a. |
| CCSAP | 1.005 | 9.2445 | 109.53 | 1E-05 | 0.0002 | 0.2857143 | n.a. |
| AKIP1 | 1.07 | 9.2726 | 57.685 | 1E-05 | 0.0003 | 0.4 | n.a. |
| ENSECAG00000017450 | 1.011 | 9.2479 | 66.045 | 1E-05 | 0.0003 | 0.2571429 | n.a. |
| HAUS5 | 1.042 | 9.2685 | 43.73 | 2E-05 | 0.0003 | 0.4285714 | n.a. |
| DCAKD | 1.082 | 9.2565 | 66.781 | 2E-05 | 0.0004 | 0.5142857 | n.a. |
| CENPH | 1.119 | 9.2558 | 74.871 | 2E-05 | 0.0004 | 0.4285714 | n.a. |
| NSD2 | 1.019 | 9.2457 | 170.86 | 2E-05 | 0.0004 | 0.2857143 | n.a. |
