## Supplementary material for "Single cell resolution landscape of equine peripheral blood mononuclear cells reveals diverse immune cell subtypes including T-bet^+^ B cells": Dataset S6

```
24        ensembl   gene      47209642  47215788  .         -         .
gene_id "ENSECAG00000009625"; gene_version "2"; gene_name "IGHA"; gene_source
"ensembl"; gene_biotype "protein_coding";
24        ensembl   transcript 47209642  47215788  .         -         .
gene_id "ENSECAG00000009625"; gene_version "2"; transcript_id "ENSECAT00000009807";
transcript_version "2"; gene_name "IGHA"; gene_source "ensembl"; gene_biotype
"protein_coding"; transcript_source "ensembl"; transcript_biotype "protein_coding";
24        ensembl   exon      47215726  47215788  .         -         .
gene_id "ENSECAG00000009625"; gene_version "2"; transcript_id "ENSECAT00000009807";
transcript_version "2"; exon_number "1"; gene_name "IGHA"; gene_source "ensembl";
gene_biotype "protein_coding"; transcript_source "ensembl"; transcript_biotype
"protein_coding"; exon_id "ENSECAE00000219832"; exon_version "1";
24        ensembl   CDS       47215726  47215788  .         -         0
gene_id "ENSECAG00000009625"; gene_version "2"; transcript_id "ENSECAT00000009807";
transcript_version "2"; exon_number "1"; gene_name "IGHA"; gene_source "ensembl";
gene_biotype "protein_coding"; transcript_source "ensembl"; transcript_biotype
"protein_coding"; protein_id "ENSECAP00000007504"; protein_version "2";
24        ensembl   exon      47210820  47211153  .         -         .
gene_id "ENSECAG00000009625"; gene_version "2"; transcript_id "ENSECAT00000009807";
transcript_version "2"; exon_number "2"; gene_name "IGHA"; gene_source "ensembl";
gene_biotype "protein_coding"; transcript_source "ensembl"; transcript_biotype
"protein_coding"; exon_id "ENSECAE00000051120"; exon_version "2";
24        ensembl   CDS       47210820  47211153  .         -         0
gene_id "ENSECAG00000009625"; gene_version "2"; transcript_id "ENSECAT00000009807";
transcript_version "2"; exon_number "2"; gene_name "IGHA"; gene_source "ensembl";
gene_biotype "protein_coding"; transcript_source "ensembl"; transcript_biotype
"protein_coding"; protein_id "ENSECAP00000007504"; protein_version "2";
24        ensembl   exon      47210248  47210592  .         -         .
gene_id "ENSECAG00000009625"; gene_version "2"; transcript_id "ENSECAT00000009807";
transcript_version "2"; exon_number "3"; gene_name "IGHA"; gene_source "ensembl";
gene_biotype "protein_coding"; transcript_source "ensembl"; transcript_biotype
"protein_coding"; exon_id "ENSECAE00000051147"; exon_version "2";
24        ensembl   CDS       47210248  47210592  .         -         2
gene_id "ENSECAG00000009625"; gene_version "2"; transcript_id "ENSECAT00000009807";
transcript_version "2"; exon_number "3"; gene_name "IGHA"; gene_source "ensembl";
gene_biotype "protein_coding"; transcript_source "ensembl"; transcript_biotype
"protein_coding"; protein_id "ENSECAP00000007504"; protein_version "2";
24        ensembl   exon      47209642  47210036  .         -         .
gene_id "ENSECAG00000009625"; gene_version "2"; transcript_id "ENSECAT00000009807";
transcript_version "2"; exon_number "4"; gene_name "IGHA"; gene_source "ensembl";
gene_biotype "protein_coding"; transcript_source "ensembl"; transcript_biotype
"protein_coding"; exon_id "ENSECAE00000293426"; exon_version "1";
24        ensembl   CDS       47209645  47210036  .         -         2
gene_id "ENSECAG00000009625"; gene_version "2"; transcript_id "ENSECAT00000009807";
transcript_version "2"; exon_number "4"; gene_name "IGHA"; gene_source "ensembl";
gene_biotype "protein_coding"; transcript_source "ensembl"; transcript_biotype
"protein_coding"; protein_id "ENSECAP00000007504"; protein_version "2";
24        ensembl   stop_codon 47209642  47209644  .         -         0
gene_id "ENSECAG00000009625"; gene_version "2"; transcript_id "ENSECAT00000009807";
transcript_version "2"; exon_number "4"; gene_name "IGHA"; gene_source "ensembl";
gene_biotype "protein_coding"; transcript_source "ensembl"; transcript_biotype
"protein_coding";
24        ensembl   gene      47221642  47223613  .         -         .
gene_id "ENSECAG00000009575"; gene_version "2"; gene_name "IGHE"; gene_source
"ensembl"; gene_biotype "protein_coding";
24        ensembl   transcript 47221642  47223613  .         -         .
gene_id "ENSECAG00000009575"; gene_version "2"; transcript_id "ENSECAT00000009796";
transcript_version "2"; gene_name "IGHE"; gene_source "ensembl"; gene_biotype
"protein_coding"; transcript_source "ensembl"; transcript_biotype "protein_coding";
24        ensembl   exon      47223074  47223613  .         -         .
gene_id "ENSECAG00000009575"; gene_version "2"; transcript_id "ENSECAT00000009796";
transcript_version "2"; exon_number "1"; gene_name "IGHE"; gene_source "ensembl";
gene_biotype "protein_coding"; transcript_source "ensembl"; transcript_biotype
"protein_coding"; exon_id "ENSECAE00000246955"; exon_version "1";
24        ensembl   CDS       47223074  47223314  .         -         0
gene_id "ENSECAG00000009575"; gene_version "2"; transcript_id "ENSECAT00000009796";
```

```
transcript_version "2"; exon_number "1"; gene_name "IGHE"; gene_source "ensembl";
gene_biotype "protein_coding"; transcript_source "ensembl"; transcript_biotype
"protein_coding"; protein_id "ENSECAP00000007494"; protein_version "2";
24	ensembl	exon	47222590	47222913	.	-	.
gene_id "ENSECAG00000009575"; gene_version "2"; transcript_id "ENSECAT00000009796";
transcript_version "2"; exon_number "2"; gene_name "IGHE"; gene_source "ensembl";
gene_biotype "protein_coding"; transcript_source "ensembl"; transcript_biotype
"protein_coding"; exon_id "ENSECAE00000050683"; exon_version "1";
24	ensembl	CDS	47222590	47222913	.	-	2
gene_id "ENSECAG00000009575"; gene_version "2"; transcript_id "ENSECAT00000009796";
transcript_version "2"; exon_number "2"; gene_name "IGHE"; gene_source "ensembl";
gene_biotype "protein_coding"; transcript_source "ensembl"; transcript_biotype
"protein_coding"; protein_id "ENSECAP00000007494"; protein_version "2";
24	ensembl	exon	47222172	47222492	.	-	.
gene_id "ENSECAG00000009575"; gene_version "2"; transcript_id "ENSECAT00000009796";
transcript_version "2"; exon_number "3"; gene_name "IGHE"; gene_source "ensembl";
gene_biotype "protein_coding"; transcript_source "ensembl"; transcript_biotype
"protein_coding"; exon_id "ENSECAE00000050851"; exon_version "1";
24	ensembl	CDS	47222172	47222492	.	-	2
gene_id "ENSECAG00000009575"; gene_version "2"; transcript_id "ENSECAT00000009796";
transcript_version "2"; exon_number "3"; gene_name "IGHE"; gene_source "ensembl";
gene_biotype "protein_coding"; transcript_source "ensembl"; transcript_biotype
"protein_coding"; protein_id "ENSECAP00000007494"; protein_version "2";
24	ensembl	exon	47221642	47222099	.	-	.
gene_id "ENSECAG00000009575"; gene_version "2"; transcript_id "ENSECAT00000009796";
transcript_version "2"; exon_number "4"; gene_name "IGHE"; gene_source "ensembl";
gene_biotype "protein_coding"; transcript_source "ensembl"; transcript_biotype
"protein_coding"; exon_id "ENSECAE00000051024"; exon_version "2";
24	ensembl	CDS	47221765	47222099	.	-	2
gene_id "ENSECAG00000009575"; gene_version "2"; transcript_id "ENSECAT00000009796";
transcript_version "2"; exon_number "4"; gene_name "IGHE"; gene_source "ensembl";
gene_biotype "protein_coding"; transcript_source "ensembl"; transcript_biotype
"protein_coding"; protein_id "ENSECAP00000007494"; protein_version "2";
24	ensembl	stop_codon	47221762	47221764	.	-	0
gene_id "ENSECAG00000009575"; gene_version "2"; transcript_id "ENSECAT00000009796";
transcript_version "2"; exon_number "4"; gene_name "IGHE"; gene_source "ensembl";
gene_biotype "protein_coding"; transcript_source "ensembl"; transcript_biotype
"protein_coding";
24	ensembl	five_prime_utr	47223315	47223613	.	-	.
	gene_id "ENSECAG00000009575"; gene_version "2"; transcript_id
"ENSECAT00000009796"; transcript_version "2"; gene_name "IGHE"; gene_source "ensembl";
gene_biotype "protein_coding"; transcript_source "ensembl"; transcript_biotype
"protein_coding";
24	ensembl	three_prime_utr	47221642	47221761	.	-	.
	gene_id "ENSECAG00000009575"; gene_version "2"; transcript_id
"ENSECAT00000009796"; transcript_version "2"; gene_name "IGHE"; gene_source "ensembl";
gene_biotype "protein_coding"; transcript_source "ensembl"; transcript_biotype
"protein_coding";
24	ensembl	gene	47283376	47277433	.	-	.
gene_id "ENSECAG00000009474"; gene_version "2"; gene_name "IGHG6"; gene_source
"ensembl"; gene_biotype "protein_coding";
24	ensembl	transcript	47283376	47277433	.	-	.
gene_id "ENSECAG00000009474"; gene_version "2"; transcript_id "ENSECAT00000009646";
transcript_version "2"; gene_name "IGHG6"; gene_source "ensembl"; gene_biotype
"protein_coding"; transcript_source "ensembl"; transcript_biotype "protein_coding";
24	ensembl	exon	47283376	47283517	.	-	.
gene_id "ENSECAG00000009474"; gene_version "2"; transcript_id "ENSECAT00000009646";
transcript_version "2"; exon_number "1"; gene_name "IGHG6"; gene_source "ensembl";
gene_biotype "protein_coding"; transcript_source "ensembl"; transcript_biotype
"protein_coding"; exon_id "ENSECAE00000259340"; exon_version "1";
24	ensembl	CDS	47283376	47283517	.	-	0
gene_id "ENSECAG00000009474"; gene_version "2"; transcript_id "ENSECAT00000009646";
transcript_version "2"; exon_number "1"; gene_name "IGHG6"; gene_source "ensembl";
gene_biotype "protein_coding"; transcript_source "ensembl"; transcript_biotype
"protein_coding"; protein_id "ENSECAP00000007368"; protein_version "2";
24	ensembl	start_codon	47283515	47283517	.	-	0
```

```
          gene_id "ENSECAG00000009474"; gene_version "2"; transcript_id
"ENSECAT00000009646"; transcript_version "2"; exon_number "1"; gene_name "IGHG6";
gene_source "ensembl"; gene_biotype "protein_coding"; transcript_source "ensembl";
transcript_biotype "protein_coding";
24        ensembl   exon      47280183  47280476  .         -         .
gene_id "ENSECAG00000009474"; gene_version "2"; transcript_id "ENSECAT00000009646";
transcript_version "2"; exon_number "2"; gene_name "IGHG6"; gene_source "ensembl";
gene_biotype "protein_coding"; transcript_source "ensembl"; transcript_biotype
"protein_coding"; exon_id "ENSECAE00000014626"; exon_version "2";
24        ensembl   CDS       47280183  47280476  .         -         2
gene_id "ENSECAG00000009474"; gene_version "2"; transcript_id "ENSECAT00000009646";
transcript_version "2"; exon_number "2"; gene_name "IGHG6"; gene_source "ensembl";
gene_biotype "protein_coding"; transcript_source "ensembl"; transcript_biotype
"protein_coding"; protein_id "ENSECAP00000007368"; protein_version "2";
24        ensembl   exon      47279863  47279898  .         -         .
gene_id "ENSECAG00000009474"; gene_version "2"; transcript_id "ENSECAT00000009646";
transcript_version "2"; exon_number "3"; gene_name "IGHG6"; gene_source "ensembl";
gene_biotype "protein_coding"; transcript_source "ensembl"; transcript_biotype
"protein_coding"; exon_id "ENSECAE00000049480"; exon_version "1";
24        ensembl   CDS       47279863  47279898  .         -         2
gene_id "ENSECAG00000009474"; gene_version "2"; transcript_id "ENSECAT00000009646";
transcript_version "2"; exon_number "3"; gene_name "IGHG6"; gene_source "ensembl";
gene_biotype "protein_coding"; transcript_source "ensembl"; transcript_biotype
"protein_coding"; protein_id "ENSECAP00000007368"; protein_version "2";
24        ensembl   exon      47279409  47279738  .         -         .
gene_id "ENSECAG00000009474"; gene_version "2"; transcript_id "ENSECAT00000009646";
transcript_version "2"; exon_number "4"; gene_name "IGHG6"; gene_source "ensembl";
gene_biotype "protein_coding"; transcript_source "ensembl"; transcript_biotype
"protein_coding"; exon_id "ENSECAE00000049514"; exon_version "2";
24        ensembl   CDS       47279409  47279738  .         -         2
gene_id "ENSECAG00000009474"; gene_version "2"; transcript_id "ENSECAT00000009646";
transcript_version "2"; exon_number "4"; gene_name "IGHG6"; gene_source "ensembl";
gene_biotype "protein_coding"; transcript_source "ensembl"; transcript_biotype
"protein_coding"; protein_id "ENSECAP00000007368"; protein_version "2";
24        ensembl   exon      47279005  47279325  .         -         .
gene_id "ENSECAG00000009474"; gene_version "2"; transcript_id "ENSECAT00000009646";
transcript_version "2"; exon_number "5"; gene_name "IGHG6"; gene_source "ensembl";
gene_biotype "protein_coding"; transcript_source "ensembl"; transcript_biotype
"protein_coding"; exon_id "ENSECAE00000049529"; exon_version "2";
24        ensembl   CDS       47279005  47279325  .         -         2
gene_id "ENSECAG00000009474"; gene_version "2"; transcript_id "ENSECAT00000009646";
transcript_version "2"; exon_number "5"; gene_name "IGHG6"; gene_source "ensembl";
gene_biotype "protein_coding"; transcript_source "ensembl"; transcript_biotype
"protein_coding"; protein_id "ENSECAP00000007368"; protein_version "2";
24        ensembl   exon      47277303  47277433  .         -         .
gene_id "ENSECAG00000009474"; gene_version "2"; transcript_id "ENSECAT00000009646";
transcript_version "2"; exon_number "6"; gene_name "IGHG6"; gene_source "ensembl";
gene_biotype "protein_coding"; transcript_source "ensembl"; transcript_biotype
"protein_coding"; exon_id "ENSECAE00000241476"; exon_version "1";
24        ensembl   CDS       47277303  47277433  .         -         2
gene_id "ENSECAG00000009474"; gene_version "2"; transcript_id "ENSECAT00000009646";
transcript_version "2"; exon_number "6"; gene_name "IGHG6"; gene_source "ensembl";
gene_biotype "protein_coding"; transcript_source "ensembl"; transcript_biotype
"protein_coding"; protein_id "ENSECAP00000007368"; protein_version "2";
24        ensembl   stop_codon 47277431  47277433  .         -         0
gene_id "ENSECAG00000009474"; gene_version "2"; transcript_id "ENSECAT00000009646";
transcript_version "2"; exon_number "7"; gene_name "IGHG6"; gene_source "ensembl";
gene_biotype "protein_coding"; transcript_source "ensembl"; transcript_biotype
"protein_coding";
24        ensembl   gene      47248314  47244995  .         -         .
gene_id "ENSECAG10000009474"; gene_version "2"; gene_name "IGHG5"; gene_source
"ensembl"; gene_biotype "protein_coding";
24        ensembl   transcript 47248314  47244995  .         -         .
gene_id "ENSECAG10000009474"; gene_version "2"; transcript_id "ENSECAT00000003731";
transcript_version "2"; gene_name "IGHG5"; gene_source "ensembl"; gene_biotype
"protein_coding"; transcript_source "ensembl"; transcript_biotype "protein_coding";
```

```
24	ensembl	exon	47248314	47248346	.	-	.
gene_id "ENSECAG10000009474"; gene_version "2"; transcript_id "ENSECAT00000003731";
transcript_version "2"; exon_number "1"; gene_name "IGHG5"; gene_source "ensembl";
gene_biotype "protein_coding"; transcript_source "ensembl"; transcript_biotype
"protein_coding"; exon_id "ENSECAE00000050431"; exon_version "1";
24	ensembl	CDS	47248314	47248346	.	-	2
gene_id "ENSECAG10000009474"; gene_version "2"; transcript_id "ENSECAT00000003731";
transcript_version "2"; exon_number "1"; gene_name "IGHG5"; gene_source "ensembl";
gene_biotype "protein_coding"; transcript_source "ensembl"; transcript_biotype
"protein_coding"; protein_id "ENSECAP00000002609"; protein_version "2";
24	ensembl	start_codon	47248314	47248316	.	-	0
	gene_id "ENSECAG10000009474"; gene_version "2"; transcript_id
"ENSECAT00000003731"; transcript_version "2"; exon_number "1"; gene_name "IGHG5";
gene_source "ensembl"; gene_biotype "protein_coding"; transcript_source "ensembl";
transcript_biotype "protein_coding";
24	ensembl	exon	47247855	47248184	.	-	.
gene_id "ENSECAG10000009474"; gene_version "2"; transcript_id "ENSECAT00000003731";
transcript_version "2"; exon_number "2"; gene_name "IGHG5"; gene_source "ensembl";
gene_biotype "protein_coding"; transcript_source "ensembl"; transcript_biotype
"protein_coding"; exon_id "ENSECAE00000014795"; exon_version "2";
24	ensembl	CDS	47247855	47248184	.	-	2
gene_id "ENSECAG10000009474"; gene_version "2"; transcript_id "ENSECAT00000003731";
transcript_version "2"; exon_number "2"; gene_name "IGHG5"; gene_source "ensembl";
gene_biotype "protein_coding"; transcript_source "ensembl"; transcript_biotype
"protein_coding"; protein_id "ENSECAP00000002609"; protein_version "2";
24	ensembl	exon	47247450	47247770	.	-	.
gene_id "ENSECAG10000009474"; gene_version "2"; transcript_id "ENSECAT00000003731";
transcript_version "2"; exon_number "3"; gene_name "IGHG5"; gene_source "ensembl";
gene_biotype "protein_coding"; transcript_source "ensembl"; transcript_biotype
"protein_coding"; exon_id "ENSECAE00000014882"; exon_version "2";
24	ensembl	CDS	47247450	47247770	.	-	2
gene_id "ENSECAG10000009474"; gene_version "2"; transcript_id "ENSECAT00000003731";
transcript_version "2"; exon_number "3"; gene_name "IGHG5"; gene_source "ensembl";
gene_biotype "protein_coding"; transcript_source "ensembl"; transcript_biotype
"protein_coding"; protein_id "ENSECAP00000002609"; protein_version "2";
24	ensembl	exon	47245708	47245838	.	-	.
gene_id "ENSECAG10000009474"; gene_version "2"; transcript_id "ENSECAT00000003731";
transcript_version "2"; exon_number "4"; gene_name "IGHG5"; gene_source "ensembl";
gene_biotype "protein_coding"; transcript_source "ensembl"; transcript_biotype
"protein_coding"; exon_id "ENSECAE00000270785"; exon_version "1";
24	ensembl	CDS	47245708	47245838	.	-	2
gene_id "ENSECAG10000009474"; gene_version "2"; transcript_id "ENSECAT00000003731";
transcript_version "2"; exon_number "4"; gene_name "IGHG5"; gene_source "ensembl";
gene_biotype "protein_coding"; transcript_source "ensembl"; transcript_biotype
"protein_coding"; protein_id "ENSECAP00000002609"; protein_version "2";
24	ensembl	exon	47244993	47245076	.	-	.
gene_id "ENSECAG10000009474"; gene_version "2"; transcript_id "ENSECAT00000003731";
transcript_version "2"; exon_number "5"; gene_name "IGHG5"; gene_source "ensembl";
gene_biotype "protein_coding"; transcript_source "ensembl"; transcript_biotype
"protein_coding"; exon_id "ENSECAE00000236271"; exon_version "1";
24	ensembl	CDS	47244996	47245076	.	-	0
gene_id "ENSECAG10000009474"; gene_version "2"; transcript_id "ENSECAT00000003731";
transcript_version "2"; exon_number "5"; gene_name "IGHG5"; gene_source "ensembl";
gene_biotype "protein_coding"; transcript_source "ensembl"; transcript_biotype
"protein_coding"; protein_id "ENSECAP00000002609"; protein_version "2";
24	ensembl	stop_codon	47244993	47244995	.	-	0
gene_id "ENSECAG10000009474"; gene_version "2"; transcript_id "ENSECAT00000003731";
transcript_version "2"; exon_number "5"; gene_name "IGHG5"; gene_source "ensembl";
gene_biotype "protein_coding"; transcript_source "ensembl"; transcript_biotype
"protein_coding";
24	ensembl	gene	47303436	47302594	.	-	.
gene_id "ENSECAG10000007258"; gene_version "2"; gene_name "IGHG4"; gene_source
"ensembl"; gene_biotype "protein_coding";
24	ensembl	transcript	47303436	47302594	.	-	.
gene_id "ENSECAG10000007258"; gene_version "2"; transcript_id "ENSECAT10000007376";
transcript_version "2"; gene_name "IGHG4"; gene_source "ensembl"; gene_biotype
```

```
"protein_coding"; transcript_source "ensembl"; transcript_biotype "protein_coding";
24	ensembl	exon	47303436	47303729	.	-	.
gene_id "ENSECAG10000007258"; gene_version "2"; transcript_id "ENSECAT10000007376";
transcript_version "2"; exon_number "1"; gene_name "IGHG4"; gene_source "ensembl";
gene_biotype "protein_coding"; transcript_source "ensembl"; transcript_biotype
"protein_coding"; exon_id "ENSECAE00000050431"; exon_version "1";
24	ensembl	CDS	47303436	47303729	.	-	2
gene_id "ENSECAG10000007258"; gene_version "2"; transcript_id "ENSECAT10000007376";
transcript_version "2"; exon_number "1"; gene_name "IGHG4"; gene_source "ensembl";
gene_biotype "protein_coding"; transcript_source "ensembl"; transcript_biotype
"protein_coding"; protein_id "ENSECAP00000002609"; protein_version "2";
24	ensembl	start_codon	47303436	47303438	.	-	0
	gene_id "ENSECAG10000007258"; gene_version "2"; transcript_id
"ENSECAT10000007376"; transcript_version "2"; exon_number "1"; gene_name "IGHG4";
gene_source "ensembl"; gene_biotype "protein_coding"; transcript_source "ensembl";
transcript_biotype "protein_coding";
24	ensembl	exon	47303120	47303170	.	-	.
gene_id "ENSECAG10000007258"; gene_version "2"; transcript_id "ENSECAT10000007376";
transcript_version "2"; exon_number "2"; gene_name "IGHG4"; gene_source "ensembl";
gene_biotype "protein_coding"; transcript_source "ensembl"; transcript_biotype
"protein_coding"; exon_id "ENSECAE00000014795"; exon_version "2";
24	ensembl	CDS	47303120	47303170	.	-	2
gene_id "ENSECAG10000007258"; gene_version "2"; transcript_id "ENSECAT10000007376";
transcript_version "2"; exon_number "2"; gene_name "IGHG4"; gene_source "ensembl";
gene_biotype "protein_coding"; transcript_source "ensembl"; transcript_biotype
"protein_coding"; protein_id "ENSECAP00000002609"; protein_version "2";
24	ensembl	exon	47302661	47302992	.	-	.
gene_id "ENSECAG10000007258"; gene_version "2"; transcript_id "ENSECAT10000007376";
transcript_version "2"; exon_number "3"; gene_name "IGHG4"; gene_source "ensembl";
gene_biotype "protein_coding"; transcript_source "ensembl"; transcript_biotype
"protein_coding"; exon_id "ENSECAE00000014882"; exon_version "2";
24	ensembl	CDS	47302661	47302992	.	-	2
gene_id "ENSECAG10000007258"; gene_version "2"; transcript_id "ENSECAT10000007376";
transcript_version "2"; exon_number "3"; gene_name "IGHG4"; gene_source "ensembl";
gene_biotype "protein_coding"; transcript_source "ensembl"; transcript_biotype
"protein_coding"; protein_id "ENSECAP00000002609"; protein_version "2";
24	ensembl	exon	47302258	47302594	.	-	.
gene_id "ENSECAG10000007258"; gene_version "2"; transcript_id "ENSECAT10000007376";
transcript_version "2"; exon_number "4"; gene_name "IGHG4"; gene_source "ensembl";
gene_biotype "protein_coding"; transcript_source "ensembl"; transcript_biotype
"protein_coding"; exon_id "ENSECAE00000270785"; exon_version "1";
24	ensembl	CDS	47302258	47302594	.	-	2
gene_id "ENSECAG10000007258"; gene_version "2"; transcript_id "ENSECAT10000007376";
transcript_version "2"; exon_number "4"; gene_name "IGHG4"; gene_source "ensembl";
gene_biotype "protein_coding"; transcript_source "ensembl"; transcript_biotype
"protein_coding"; protein_id "ENSECAP00000002609"; protein_version "2";
24	ensembl	stop_codon	47302592	47302594	.	-	0
gene_id "ENSECAG10000007258"; gene_version "2"; transcript_id "ENSECAT10000007376";
transcript_version "2"; exon_number "5"; gene_name "IGHG4"; gene_source "ensembl";
gene_biotype "protein_coding"; transcript_source "ensembl"; transcript_biotype
"protein_coding";
24	ensembl	gene	47323097	47332110	.	-	.
gene_id "ENSECAG00000006095"; gene_version "2"; gene_name "IGHG3"; gene_source
"ensembl"; gene_biotype "protein_coding";
24	ensembl	transcript	47323097	47332110	.	-	.
gene_id "ENSECAG00000006095"; gene_version "2"; transcript_id "ENSECAT00000003796";
transcript_version "2"; gene_name "IGHG3"; gene_source "ensembl"; gene_biotype
"protein_coding"; transcript_source "ensembl"; transcript_biotype "protein_coding";
24	ensembl	exon	47332095	47332110	.	-	.
gene_id "ENSECAG00000006095"; gene_version "2"; transcript_id "ENSECAT00000003796";
transcript_version "2"; exon_number "1"; gene_name "IGHG3"; gene_source "ensembl";
gene_biotype "protein_coding"; transcript_source "ensembl"; transcript_biotype
"protein_coding"; exon_id "ENSECAE00000244021"; exon_version "1";
24	ensembl	CDS	47332095	47332110	.	-	0
gene_id "ENSECAG00000006095"; gene_version "2"; transcript_id "ENSECAT00000003796";
transcript_version "2"; exon_number "1"; gene_name "IGHG3"; gene_source "ensembl";
```

```
gene_biotype "protein_coding"; transcript_source "ensembl"; transcript_biotype
"protein_coding"; protein_id "ENSECAP00000002646"; protein_version "2";
24        ensembl   start_codon         47332108  47332110  .         -         0
          gene_id "ENSECAG00000006095"; gene_version "2"; transcript_id
"ENSECAT00000003796"; transcript_version "2"; exon_number "1"; gene_name "IGHG3";
gene_source "ensembl"; gene_biotype "protein_coding"; transcript_source "ensembl";
transcript_biotype "protein_coding";
24        ensembl   exon      47327486  47327779  .         -         .
gene_id "ENSECAG00000006095"; gene_version "2"; transcript_id "ENSECAT00000003796";
transcript_version "2"; exon_number "2"; gene_name "IGHG3"; gene_source "ensembl";
gene_biotype "protein_coding"; transcript_source "ensembl"; transcript_biotype
"protein_coding"; exon_id "ENSECAE00000015003"; exon_version "2";
24        ensembl   CDS       47327486  47327779  .         -         2
gene_id "ENSECAG00000006095"; gene_version "2"; transcript_id "ENSECAT00000003796";
transcript_version "2"; exon_number "2"; gene_name "IGHG3"; gene_source "ensembl";
gene_biotype "protein_coding"; transcript_source "ensembl"; transcript_biotype
"protein_coding"; protein_id "ENSECAP00000002646"; protein_version "2";
24        ensembl   exon      47327122  47327232  .         -         .
gene_id "ENSECAG00000006095"; gene_version "2"; transcript_id "ENSECAT00000003796";
transcript_version "2"; exon_number "3"; gene_name "IGHG3"; gene_source "ensembl";
gene_biotype "protein_coding"; transcript_source "ensembl"; transcript_biotype
"protein_coding"; exon_id "ENSECAE00000021936"; exon_version "1";
24        ensembl   CDS       47327122  47327232  .         -         2
gene_id "ENSECAG00000006095"; gene_version "2"; transcript_id "ENSECAT00000003796";
transcript_version "2"; exon_number "3"; gene_name "IGHG3"; gene_source "ensembl";
gene_biotype "protein_coding"; transcript_source "ensembl"; transcript_biotype
"protein_coding"; protein_id "ENSECAP00000002646"; protein_version "2";
24        ensembl   exon      47326660  47326989  .         -         .
gene_id "ENSECAG00000006095"; gene_version "2"; transcript_id "ENSECAT00000003796";
transcript_version "2"; exon_number "4"; gene_name "IGHG3"; gene_source "ensembl";
gene_biotype "protein_coding"; transcript_source "ensembl"; transcript_biotype
"protein_coding"; exon_id "ENSECAE00000015081"; exon_version "2";
24        ensembl   CDS       47326660  47326989  .         -         2
gene_id "ENSECAG00000006095"; gene_version "2"; transcript_id "ENSECAT00000003796";
transcript_version "2"; exon_number "4"; gene_name "IGHG3"; gene_source "ensembl";
gene_biotype "protein_coding"; transcript_source "ensembl"; transcript_biotype
"protein_coding"; protein_id "ENSECAP00000002646"; protein_version "2";
24        ensembl   exon      47326264  47326584  .         -         .
gene_id "ENSECAG00000006095"; gene_version "2"; transcript_id "ENSECAT00000003796";
transcript_version "2"; exon_number "5"; gene_name "IGHG3"; gene_source "ensembl";
gene_biotype "protein_coding"; transcript_source "ensembl"; transcript_biotype
"protein_coding"; exon_id "ENSECAE00000015121"; exon_version "2";
24        ensembl   CDS       47326264  47326584  .         -         2
gene_id "ENSECAG00000006095"; gene_version "2"; transcript_id "ENSECAT00000003796";
transcript_version "2"; exon_number "5"; gene_name "IGHG3"; gene_source "ensembl";
gene_biotype "protein_coding"; transcript_source "ensembl"; transcript_biotype
"protein_coding"; protein_id "ENSECAP00000002646"; protein_version "2";
24        ensembl   exon      47324973  47325103  .         -         .
gene_id "ENSECAG00000006095"; gene_version "2"; transcript_id "ENSECAT00000003796";
transcript_version "2"; exon_number "6"; gene_name "IGHG3"; gene_source "ensembl";
gene_biotype "protein_coding"; transcript_source "ensembl"; transcript_biotype
"protein_coding"; exon_id "ENSECAE00000294111"; exon_version "1";
24        ensembl   CDS       47324973  47325103  .         -         2
gene_id "ENSECAG00000006095"; gene_version "2"; transcript_id "ENSECAT00000003796";
transcript_version "2"; exon_number "6"; gene_name "IGHG3"; gene_source "ensembl";
gene_biotype "protein_coding"; transcript_source "ensembl"; transcript_biotype
"protein_coding"; protein_id "ENSECAP00000002646"; protein_version "2";
24        ensembl   exon      47323097  47324418  .         -         .
gene_id "ENSECAG00000006095"; gene_version "2"; transcript_id "ENSECAT00000003796";
transcript_version "2"; exon_number "7"; gene_name "IGHG3"; gene_source "ensembl";
gene_biotype "protein_coding"; transcript_source "ensembl"; transcript_biotype
"protein_coding"; exon_id "ENSECAE00000241616"; exon_version "1";
24        ensembl   CDS       47324338  47324418  .         -         0
gene_id "ENSECAG00000006095"; gene_version "2"; transcript_id "ENSECAT00000003796";
transcript_version "2"; exon_number "7"; gene_name "IGHG3"; gene_source "ensembl";
gene_biotype "protein_coding"; transcript_source "ensembl"; transcript_biotype
```

"protein_coding"; protein_id "ENSECAP00000002646"; protein_version "2";
24 ensembl stop_codon 47324335 47324337 . - 0
gene_id "ENSECAG00000006095"; gene_version "2"; transcript_id "ENSECAT00000003796"; transcript_version "2"; exon_number "7"; gene_name "IGHG3"; gene_source "ensembl"; gene_biotype "protein_coding"; transcript_source "ensembl"; transcript_biotype "protein_coding";
24 ensembl three_prime_utr 47323097 47324334 . - .
gene_id "ENSECAG00000006095"; gene_version "2"; transcript_id "ENSECAT00000003796"; transcript_version "2"; gene_name "IGHG3"; gene_source "ensembl"; gene_biotype "protein_coding"; transcript_source "ensembl"; transcript_biotype "protein_coding";
24 ensembl gene 47352977 47352095 . - .
gene_id "ENSECAG00000003774"; gene_version "2"; gene_name "IGHG2"; gene_source "ensembl"; gene_biotype "protein_coding";
24 ensembl transcript 47352977 47352095 . - .
gene_id "ENSECAG00000003774"; gene_version "2"; transcript_id "ENSECAT00000042165"; transcript_version "1"; gene_name "IGHG2"; gene_source "ensembl"; gene_biotype "protein_coding"; transcript_source "ensembl"; transcript_biotype "protein_coding";
24 ensembl exon 47352977 47353270 . - .
gene_id "ENSECAG00000003774"; gene_version "2"; transcript_id "ENSECAT00000042165"; transcript_version "1"; exon_number "1"; gene_name "IGHG2"; gene_source "ensembl"; gene_biotype "protein_coding"; transcript_source "ensembl"; transcript_biotype "protein_coding"; exon_id "ENSECAE00000242419"; exon_version "1";
24 ensembl CDS 47352977 47353270 . - 0
gene_id "ENSECAG00000003774"; gene_version "2"; transcript_id "ENSECAT00000042165"; transcript_version "1"; exon_number "1"; gene_name "IGHG2"; gene_source "ensembl"; gene_biotype "protein_coding"; transcript_source "ensembl"; transcript_biotype "protein_coding"; protein_id "ENSECAP00000031549"; protein_version "1";
24 ensembl start_codon 47352977 47352979 . - 0
gene_id "ENSECAG00000003774"; gene_version "2"; transcript_id "ENSECAT00000042165"; transcript_version "1"; exon_number "1"; gene_name "IGHG2"; gene_source "ensembl"; gene_biotype "protein_coding"; transcript_source "ensembl"; transcript_biotype "protein_coding";
24 ensembl exon 47352640 47352714 . - .
gene_id "ENSECAG00000003774"; gene_version "2"; transcript_id "ENSECAT00000042165"; transcript_version "1"; exon_number "2"; gene_name "IGHG2"; gene_source "ensembl"; gene_biotype "protein_coding"; transcript_source "ensembl"; transcript_biotype "protein_coding"; exon_id "ENSECAE00000274052"; exon_version "1";
24 ensembl CDS 47352640 47352714 . - 2
gene_id "ENSECAG00000003774"; gene_version "2"; transcript_id "ENSECAT00000042165"; transcript_version "1"; exon_number "2"; gene_name "IGHG2"; gene_source "ensembl"; gene_biotype "protein_coding"; transcript_source "ensembl"; transcript_biotype "protein_coding"; protein_id "ENSECAP00000031549"; protein_version "1";
24 ensembl exon 47352179 47352512 . - .
gene_id "ENSECAG00000003774"; gene_version "2"; transcript_id "ENSECAT00000042165"; transcript_version "1"; exon_number "3"; gene_name "IGHG2"; gene_source "ensembl"; gene_biotype "protein_coding"; transcript_source "ensembl"; transcript_biotype "protein_coding"; exon_id "ENSECAE00000014724"; exon_version "1";
24 ensembl CDS 47352179 47352512 . - 2
gene_id "ENSECAG00000003774"; gene_version "2"; transcript_id "ENSECAT00000042165"; transcript_version "1"; exon_number "3"; gene_name "IGHG2"; gene_source "ensembl"; gene_biotype "protein_coding"; transcript_source "ensembl"; transcript_biotype "protein_coding"; protein_id "ENSECAP00000031549"; protein_version "1";
24 ensembl exon 47351772 47352095 . - .
gene_id "ENSECAG00000003774"; gene_version "2"; transcript_id "ENSECAT00000042165"; transcript_version "1"; exon_number "4"; gene_name "IGHG2"; gene_source "ensembl"; gene_biotype "protein_coding"; transcript_source "ensembl"; transcript_biotype "protein_coding"; exon_id "ENSECAE00000244690"; exon_version "1";
24 ensembl CDS 47351772 47352095 . - 2
gene_id "ENSECAG00000003774"; gene_version "2"; transcript_id "ENSECAT00000042165"; transcript_version "1"; exon_number "4"; gene_name "IGHG2"; gene_source "ensembl"; gene_biotype "protein_coding"; transcript_source "ensembl"; transcript_biotype "protein_coding"; protein_id "ENSECAP00000031549"; protein_version "1";
24 ensembl stop_codon 47352093 47352095 . - 0
gene_id "ENSECAG00000003774"; gene_version "2"; transcript_id "ENSECAT00000042165"; transcript_version "1"; exon_number "7"; gene_name "IGHG2"; gene_source "ensembl";

```
gene_biotype "protein_coding"; transcript_source "ensembl"; transcript_biotype
"protein_coding";
24	ensembl	gene	47384898	47381485	.	-	.
gene_id "ENSECAG10000003774"; gene_version "2"; gene_name "IGHG1"; gene_source
"ensembl"; gene_biotype "protein_coding";
24	ensembl	transcript	47384898	47381485	.	-	.
gene_id "ENSECAG10000003774"; gene_version "2"; transcript_id "ENSECAT00000052499";
transcript_version "1"; gene_name "IGHG1"; gene_source "ensembl"; gene_biotype
"protein_coding"; transcript_source "ensembl"; transcript_biotype "protein_coding";
24	ensembl	exon	47384898	47385191	.	-	.
gene_id "ENSECAG10000003774"; gene_version "2"; transcript_id "ENSECAT00000052499";
transcript_version "1"; exon_number "1"; gene_name "IGHG1"; gene_source "ensembl";
gene_biotype "protein_coding"; transcript_source "ensembl"; transcript_biotype
"protein_coding"; exon_id "ENSECAE00000274052"; exon_version "1";
24	ensembl	CDS	47384898	47385191	.	-	2
gene_id "ENSECAG10000003774"; gene_version "2"; transcript_id "ENSECAT00000052499";
transcript_version "1"; exon_number "1"; gene_name "IGHG1"; gene_source "ensembl";
gene_biotype "protein_coding"; transcript_source "ensembl"; transcript_biotype
"protein_coding"; protein_id "ENSECAP00000042575"; protein_version "1";
24	ensembl	start_codon	47460643	47460645	.	-	0
	gene_id "ENSECAG10000003774"; gene_version "2"; transcript_id
"ENSECAT00000052499"; transcript_version "1"; exon_number "1"; gene_name "IGHG1";
gene_source "ensembl"; gene_biotype "protein_coding"; transcript_source "ensembl";
transcript_biotype "protein_coding";
24	ensembl	exon	47384554	47384613	.	-	.
gene_id "ENSECAG10000003774"; gene_version "2"; transcript_id "ENSECAT00000052499";
transcript_version "1"; exon_number "2"; gene_name "IGHG1"; gene_source "ensembl";
gene_biotype "protein_coding"; transcript_source "ensembl"; transcript_biotype
"protein_coding"; exon_id "ENSECAE00000014724"; exon_version "1";
24	ensembl	CDS	47384554	47384613	.	-	2
gene_id "ENSECAG10000003774"; gene_version "2"; transcript_id "ENSECAT00000052499";
transcript_version "1"; exon_number "2"; gene_name "IGHG1"; gene_source "ensembl";
gene_biotype "protein_coding"; transcript_source "ensembl"; transcript_biotype
"protein_coding"; protein_id "ENSECAP00000042575"; protein_version "1";
24	ensembl	exon	47384092	47384421	.	-	.
gene_id "ENSECAG10000003774"; gene_version "2"; transcript_id "ENSECAT00000052499";
transcript_version "1"; exon_number "3"; gene_name "IGHG1"; gene_source "ensembl";
gene_biotype "protein_coding"; transcript_source "ensembl"; transcript_biotype
"protein_coding"; exon_id "ENSECAE00000244690"; exon_version "1";
24	ensembl	CDS	47384092	47384421	.	-	2
gene_id "ENSECAG10000003774"; gene_version "2"; transcript_id "ENSECAT00000052499";
transcript_version "1"; exon_number "3"; gene_name "IGHG1"; gene_source "ensembl";
gene_biotype "protein_coding"; transcript_source "ensembl"; transcript_biotype
"protein_coding"; protein_id "ENSECAP00000042575"; protein_version "1";
24	ensembl	exon	47383687	47384007	.	-	.
gene_id "ENSECAG10000003774"; gene_version "2"; transcript_id "ENSECAT00000052499";
transcript_version "1"; exon_number "4"; gene_name "IGHG1"; gene_source "ensembl";
gene_biotype "protein_coding"; transcript_source "ensembl"; transcript_biotype
"protein_coding"; exon_id "ENSECAE00000022102"; exon_version "2";
24	ensembl	CDS	47383687	47384007	.	-	2
gene_id "ENSECAG10000003774"; gene_version "2"; transcript_id "ENSECAT00000052499";
transcript_version "1"; exon_number "4"; gene_name "IGHG1"; gene_source "ensembl";
gene_biotype "protein_coding"; transcript_source "ensembl"; transcript_biotype
"protein_coding"; protein_id "ENSECAP00000042575"; protein_version "1";
24	ensembl	exon	47381970	47382100	.	-	.
gene_id "ENSECAG10000003774"; gene_version "2"; transcript_id "ENSECAT00000052499";
transcript_version "1"; exon_number "5"; gene_name "IGHG1"; gene_source "ensembl";
gene_biotype "protein_coding"; transcript_source "ensembl"; transcript_biotype
"protein_coding"; exon_id "ENSECAE00000252397"; exon_version "1";
24	ensembl	CDS	47381970	47382100	.	-	2
gene_id "ENSECAG10000003774"; gene_version "2"; transcript_id "ENSECAT00000052499";
transcript_version "1"; exon_number "5"; gene_name "IGHG1"; gene_source "ensembl";
gene_biotype "protein_coding"; transcript_source "ensembl"; transcript_biotype
"protein_coding"; protein_id "ENSECAP00000042575"; protein_version "1";
24	ensembl	exon	47381483	47381566	.	-	.
gene_id "ENSECAG10000003774"; gene_version "2"; transcript_id "ENSECAT00000052499";
```

transcript_version "1"; exon_number "6"; gene_name "IGHG1"; gene_source "ensembl";
gene_biotype "protein_coding"; transcript_source "ensembl"; transcript_biotype
"protein_coding"; exon_id "ENSECAE00000281683"; exon_version "1";
24 ensembl CDS 47381486 47381566 . - 0
gene_id "ENSECAG10000003774"; gene_version "2"; transcript_id "ENSECAT00000052499";
transcript_version "1"; exon_number "6"; gene_name "IGHG1"; gene_source "ensembl";
gene_biotype "protein_coding"; transcript_source "ensembl"; transcript_biotype
"protein_coding"; protein_id "ENSECAP00000042575"; protein_version "1";
24 ensembl stop_codon 47381483 47381485 . - 0
gene_id "ENSECAG10000003774"; gene_version "2"; transcript_id "ENSECAT00000052499";
transcript_version "1"; exon_number "6"; gene_name "IGHG1"; gene_source "ensembl";
gene_biotype "protein_coding"; transcript_source "ensembl"; transcript_biotype
"protein_coding";
24 ensembl gene 47430083 47436268 . - .
gene_id "ENSECAG00000040576"; gene_version "1"; gene_name "IGHD"; gene_source
"ensembl"; gene_biotype "protein_coding";
24 ensembl transcript 47430083 47436268 . - .
gene_id "ENSECAG00000040576"; gene_version "1"; transcript_id "ENSECAT00000058461";
transcript_version "1"; gene_name "IGHD"; gene_source "ensembl"; gene_biotype
"protein_coding"; transcript_source "ensembl"; transcript_biotype "protein_coding";
24 ensembl exon 47435605 47436268 . - .
gene_id "ENSECAG00000040576"; gene_version "1"; transcript_id "ENSECAT00000058461";
transcript_version "1"; exon_number "1"; gene_name "IGHD"; gene_source "ensembl";
gene_biotype "protein_coding"; transcript_source "ensembl"; transcript_biotype
"protein_coding"; exon_id "ENSECAE00000303750"; exon_version "1";
24 ensembl CDS 47435605 47436268 . - 0
gene_id "ENSECAG00000040576"; gene_version "1"; transcript_id "ENSECAT00000058461";
transcript_version "1"; exon_number "1"; gene_name "IGHD"; gene_source "ensembl";
gene_biotype "protein_coding"; transcript_source "ensembl"; transcript_biotype
"protein_coding"; protein_id "ENSECAP00000043488"; protein_version "1";
24 ensembl start_codon 47436266 47436268 . - 0
gene_id "ENSECAG00000040576"; gene_version "1"; transcript_id
"ENSECAT00000058461"; transcript_version "1"; exon_number "1"; gene_name "IGHD";
gene_source "ensembl"; gene_biotype "protein_coding"; transcript_source "ensembl";
transcript_biotype "protein_coding";
24 ensembl exon 47430543 47430682 . - .
gene_id "ENSECAG00000040576"; gene_version "1"; transcript_id "ENSECAT00000058461";
transcript_version "1"; exon_number "2"; gene_name "IGHD"; gene_source "ensembl";
gene_biotype "protein_coding"; transcript_source "ensembl"; transcript_biotype
"protein_coding"; exon_id "ENSECAE00000270880"; exon_version "1";
24 ensembl CDS 47430543 47430682 . - 2
gene_id "ENSECAG00000040576"; gene_version "1"; transcript_id "ENSECAT00000058461";
transcript_version "1"; exon_number "2"; gene_name "IGHD"; gene_source "ensembl";
gene_biotype "protein_coding"; transcript_source "ensembl"; transcript_biotype
"protein_coding"; protein_id "ENSECAP00000043488"; protein_version "1";
24 ensembl exon 47430083 47430091 . - .
gene_id "ENSECAG00000040576"; gene_version "1"; transcript_id "ENSECAT00000058461";
transcript_version "1"; exon_number "3"; gene_name "IGHD"; gene_source "ensembl";
gene_biotype "protein_coding"; transcript_source "ensembl"; transcript_biotype
"protein_coding"; exon_id "ENSECAE00000251497"; exon_version "1";
24 ensembl CDS 47430086 47430091 . - 0
gene_id "ENSECAG00000040576"; gene_version "1"; transcript_id "ENSECAT00000058461";
transcript_version "1"; exon_number "3"; gene_name "IGHD"; gene_source "ensembl";
gene_biotype "protein_coding"; transcript_source "ensembl"; transcript_biotype
"protein_coding"; protein_id "ENSECAP00000043488"; protein_version "1";
24 ensembl stop_codon 47430083 47430085 . - 0
gene_id "ENSECAG00000040576"; gene_version "1"; transcript_id "ENSECAT00000058461";
transcript_version "1"; exon_number "3"; gene_name "IGHD"; gene_source "ensembl";
gene_biotype "protein_coding"; transcript_source "ensembl"; transcript_biotype
"protein_coding";
24 ensembl gene 47444165 47449425 . - .
gene_id "ENSECAG00000000548"; gene_version "2"; gene_name "IGHM"; gene_source
"ensembl"; gene_biotype "protein_coding";
24 ensembl transcript 47444165 47449246 . - .
gene_id "ENSECAG00000000548"; gene_version "2"; transcript_id "ENSECAT00000052719";
transcript_version "1"; gene_name "IGHM"; gene_source "ensembl"; gene_biotype

```
"protein_coding"; transcript_source "ensembl"; transcript_biotype "protein_coding";
24	ensembl	exon	47448742	47449246	.	-	.
gene_id "ENSECAG00000000548"; gene_version "2"; transcript_id "ENSECAT00000052719";
transcript_version "1"; exon_number "1"; gene_name "IGHM"; gene_source "ensembl";
gene_biotype "protein_coding"; transcript_source "ensembl"; transcript_biotype
"protein_coding"; exon_id "ENSECAE00000003932"; exon_version "2";
24	ensembl	CDS	47448742	47449246	.	-	0
gene_id "ENSECAG00000000548"; gene_version "2"; transcript_id "ENSECAT00000052719";
transcript_version "1"; exon_number "1"; gene_name "IGHM"; gene_source "ensembl";
gene_biotype "protein_coding"; transcript_source "ensembl"; transcript_biotype
"protein_coding"; protein_id "ENSECAP00000029691"; protein_version "1";
24	ensembl	start_codon		47449244	47449246	.	-	0
	gene_id "ENSECAG00000000548"; gene_version "2"; transcript_id
"ENSECAT00000052719"; transcript_version "1"; exon_number "1"; gene_name "IGHM";
gene_source "ensembl"; gene_biotype "protein_coding"; transcript_source "ensembl";
transcript_biotype "protein_coding";
24	ensembl	exon	47447676	47448656	.	-	.
gene_id "ENSECAG00000000548"; gene_version "2"; transcript_id "ENSECAT00000052719";
transcript_version "1"; exon_number "2"; gene_name "IGHM"; gene_source "ensembl";
gene_biotype "protein_coding"; transcript_source "ensembl"; transcript_biotype
"protein_coding"; exon_id "ENSECAE00000289448"; exon_version "1";
24	ensembl	CDS	47447676	47448656	.	-	2
gene_id "ENSECAG00000000548"; gene_version "2"; transcript_id "ENSECAT00000052719";
transcript_version "1"; exon_number "2"; gene_name "IGHM"; gene_source "ensembl";
gene_biotype "protein_coding"; transcript_source "ensembl"; transcript_biotype
"protein_coding"; protein_id "ENSECAP00000029691"; protein_version "1";
24	ensembl	exon	47447177	47447509	.	-	.
gene_id "ENSECAG00000000548"; gene_version "2"; transcript_id "ENSECAT00000052719";
transcript_version "1"; exon_number "3"; gene_name "IGHM"; gene_source "ensembl";
gene_biotype "protein_coding"; transcript_source "ensembl"; transcript_biotype
"protein_coding"; exon_id "ENSECAE00000220225"; exon_version "1";
24	ensembl	CDS	47447177	47447509	.	-	2
gene_id "ENSECAG00000000548"; gene_version "2"; transcript_id "ENSECAT00000052719";
transcript_version "1"; exon_number "3"; gene_name "IGHM"; gene_source "ensembl";
gene_biotype "protein_coding"; transcript_source "ensembl"; transcript_biotype
"protein_coding"; protein_id "ENSECAP00000029691"; protein_version "1";
24	ensembl	exon	47444165	47444403	.	-	.
gene_id "ENSECAG00000000548"; gene_version "2"; transcript_id "ENSECAT00000052719";
transcript_version "1"; exon_number "4"; gene_name "IGHM"; gene_source "ensembl";
gene_biotype "protein_coding"; transcript_source "ensembl"; transcript_biotype
"protein_coding"; exon_id "ENSECAE00000307728"; exon_version "1";
24	ensembl	CDS	47444168	47444403	.	-	2
gene_id "ENSECAG00000000548"; gene_version "2"; transcript_id "ENSECAT00000052719";
transcript_version "1"; exon_number "4"; gene_name "IGHM"; gene_source "ensembl";
gene_biotype "protein_coding"; transcript_source "ensembl"; transcript_biotype
"protein_coding"; protein_id "ENSECAP00000029691"; protein_version "1";
24	ensembl	stop_codon	47444165	47444167	.	-	0
gene_id "ENSECAG00000000548"; gene_version "2"; transcript_id "ENSECAT00000052719";
transcript_version "1"; exon_number "4"; gene_name "IGHM"; gene_source "ensembl";
gene_biotype "protein_coding"; transcript_source "ensembl"; transcript_biotype
"protein_coding";
24	ensembl	transcript	47446961	47449425	.	-	.
gene_id "ENSECAG00000000548"; gene_version "2"; transcript_id "ENSECAT00000000429";
transcript_version "2"; gene_name "IGHM"; gene_source "ensembl"; gene_biotype
"protein_coding"; transcript_source "ensembl"; transcript_biotype "protein_coding";
24	ensembl	exon	47448742	47449425	.	-	.
gene_id "ENSECAG00000000548"; gene_version "2"; transcript_id "ENSECAT00000000429";
transcript_version "2"; exon_number "1"; gene_name "IGHM"; gene_source "ensembl";
gene_biotype "protein_coding"; transcript_source "ensembl"; transcript_biotype
"protein_coding"; exon_id "ENSECAE00000254333"; exon_version "1";
24	ensembl	CDS	47448742	47449246	.	-	0
gene_id "ENSECAG00000000548"; gene_version "2"; transcript_id "ENSECAT00000000429";
transcript_version "2"; exon_number "1"; gene_name "IGHM"; gene_source "ensembl";
gene_biotype "protein_coding"; transcript_source "ensembl"; transcript_biotype
"protein_coding"; protein_id "ENSECAP00000000334"; protein_version "2";
24	ensembl	start_codon		47449244	47449246	.	-	0
```

```
          gene_id "ENSECAG00000000548"; gene_version "2"; transcript_id
"ENSECAT00000000429"; transcript_version "2"; exon_number "1"; gene_name "IGHM";
gene_source "ensembl"; gene_biotype "protein_coding"; transcript_source "ensembl";
transcript_biotype "protein_coding";
24        ensembl   exon      47448315  47448656  .         -         .
gene_id "ENSECAG00000000548"; gene_version "2"; transcript_id "ENSECAT00000000429";
transcript_version "2"; exon_number "2"; gene_name "IGHM"; gene_source "ensembl";
gene_biotype "protein_coding"; transcript_source "ensembl"; transcript_biotype
"protein_coding"; exon_id "ENSECAE00000003948"; exon_version "1";
24        ensembl   CDS       47448315  47448656  .         -         2
gene_id "ENSECAG00000000548"; gene_version "2"; transcript_id "ENSECAT00000000429";
transcript_version "2"; exon_number "2"; gene_name "IGHM"; gene_source "ensembl";
gene_biotype "protein_coding"; transcript_source "ensembl"; transcript_biotype
"protein_coding"; protein_id "ENSECAP00000000334"; protein_version "2";
24        ensembl   exon      47447676  47447993  .         -         .
gene_id "ENSECAG00000000548"; gene_version "2"; transcript_id "ENSECAT00000000429";
transcript_version "2"; exon_number "3"; gene_name "IGHM"; gene_source "ensembl";
gene_biotype "protein_coding"; transcript_source "ensembl"; transcript_biotype
"protein_coding"; exon_id "ENSECAE00000003973"; exon_version "1";
24        ensembl   CDS       47447676  47447993  .         -         2
gene_id "ENSECAG00000000548"; gene_version "2"; transcript_id "ENSECAT00000000429";
transcript_version "2"; exon_number "3"; gene_name "IGHM"; gene_source "ensembl";
gene_biotype "protein_coding"; transcript_source "ensembl"; transcript_biotype
"protein_coding"; protein_id "ENSECAP00000000334"; protein_version "2";
24        ensembl   exon      47446961  47447509  .         -         .
gene_id "ENSECAG00000000548"; gene_version "2"; transcript_id "ENSECAT00000000429";
transcript_version "2"; exon_number "4"; gene_name "IGHM"; gene_source "ensembl";
gene_biotype "protein_coding"; transcript_source "ensembl"; transcript_biotype
"protein_coding"; exon_id "ENSECAE00000004015"; exon_version "2";
24        ensembl   CDS       47447118  47447509  .         -         2
gene_id "ENSECAG00000000548"; gene_version "2"; transcript_id "ENSECAT00000000429";
transcript_version "2"; exon_number "4"; gene_name "IGHM"; gene_source "ensembl";
gene_biotype "protein_coding"; transcript_source "ensembl"; transcript_biotype
"protein_coding"; protein_id "ENSECAP00000000334"; protein_version "2";
24        ensembl   stop_codon 47447115  47447117  .        -         0
gene_id "ENSECAG00000000548"; gene_version "2"; transcript_id "ENSECAT00000000429";
transcript_version "2"; exon_number "4"; gene_name "IGHM"; gene_source "ensembl";
gene_biotype "protein_coding"; transcript_source "ensembl"; transcript_biotype
"protein_coding";
24        ensembl   five_prime_utr      47449247  47449425  .         -         .
          gene_id "ENSECAG00000000548"; gene_version "2"; transcript_id
"ENSECAT00000000429"; transcript_version "2"; gene_name "IGHM"; gene_source "ensembl";
gene_biotype "protein_coding"; transcript_source "ensembl"; transcript_biotype
"protein_coding";
24        ensembl   three_prime_utr     47446961  47447114  .         -         .
          gene_id "ENSECAG00000000548"; gene_version "2"; transcript_id
"ENSECAT00000000429"; transcript_version "2"; gene_name "IGHM"; gene_source "ensembl";
gene_biotype "protein_coding"; transcript_source "ensembl"; transcript_biotype
"protein_coding";
```
