## Supplementary material for "Single cell resolution landscape of equine peripheral blood mononuclear cells reveals diverse immune cell subtypes including T-bet^+^ B cells": Dataset S7

| genes | logFC | logCPM | F | PValue | FDR | percent.exp | Surfacome.Label |
|---|---|---|---|---|---|---|---|
| GZMK | 3.3492 | 9.57824 | 7818.59 | 0 | 0 | 0.5800757 | n.a. |
| GZMA | 2.5096 | 9.45194 | 5059.96 | 0 | 0 | 0.7288777 | n.a. |
| CGA | 2 | 9.30276 | 2927.74 | 1E-219 | 3E-216 | 0.4312736 | n.a. |
| ENSECAG00000031528 | 1.3213 | 9.35154 | 1356.1 | 2E-181 | 3E-178 | 0.4350567 | n.a. |
| SH2D1A | 1.3441 | 9.45149 | 773.754 | 2E-166 | 2E-163 | 0.593947 | n.a. |
| COTL1 | 1.2774 | 10.5286 | 674.665 | 2E-147 | 3E-144 | 0.8259773 | n.a. |
| CD27 | 1.4637 | 9.47772 | 764.368 | 4E-122 | 4E-119 | 0.3278689 | surface |
| GZMM | 1.0607 | 9.49614 | 681.583 | 1E-114 | 1E-111 | 0.6443884 | n.a. |
| FABP5 | 1.3265 | 9.52638 | 455.385 | 2E-100 | 1E-97 | 0.4527112 | n.a. |
| S100A6 | 1.2954 | 10.4366 | 432.322 | 2E-95 | 1E-92 | 0.5523329 | n.a. |
| ACTG1 | 0.992 | 10.7454 | 388.519 | 5E-86 | 3E-83 | 0.7919294 | n.a. |
| MNDA | 0.8988 | 10.2583 | 304.662 | 6E-68 | 3E-65 | 0.8007566 | n.a. |
| ENSECAG00000000419 | 0.5955 | 10.7827 | 274.137 | 2E-61 | 1E-58 | 0.925599 | n.a. |
| LY6E | 0.8577 | 10.2408 | 271.929 | 7E-61 | 3E-58 | 0.703657 | surface |
| ENSECAG00000028035 | 1.0498 | 9.3107 | 804.79 | 1E-59 | 4E-57 | 0.2736444 | n.a. |
| EOMES | 1.0309 | 9.28343 | 732.794 | 2E-51 | 7E-49 | 0.3165195 | n.a. |
| CD7 | 0.5983 | 9.47261 | 222.936 | 2E-49 | 5E-47 | 0.517024 | surface |
| RF00581 | 0.7252 | 10.3191 | 217.684 | 4E-49 | 1E-46 | 0.7553594 | n.a. |
| CD44 | 0.689 | 10.108 | 155.106 | 2E-35 | 3E-33 | 0.6494325 | surface |
| ENSECAG00000020136 | 0.6637 | 9.70141 | 110.35 | 9E-26 | 1E-23 | 0.4211854 | n.a. |
| RF02216 | 0.6286 | 9.67166 | 98.2219 | 4E-23 | 5E-21 | 0.3883985 | n.a. |
| ITGA4 | 0.6543 | 9.46502 | 93.3192 | 1E-20 | 1E-18 | 0.2875158 | surface |
| ENSECAG00000032776 | 0.6507 | 9.45395 | 85.7494 | 1E-19 | 1E-17 | 0.2900378 | n.a. |

| genes | logFC | logCPM | F | PValue | FDR | percent.exp | Surfacome.Label |
|---|---|---|---|---|---|---|---|
| FCER1G | 4.3429 | 10.2039 | 14731.8 | 0 | 0 | 0.507764 | n.a. |
| NPY | 3.2007 | 9.50433 | 4520.41 | 0 | 0 | 0.7701863 | n.a. |
| ENSECAG00000001010 | 2.6362 | 9.62902 | 4114.82 | 0 | 0 | 0.6614907 | n.a. |
| KLRB1 | 2.6233 | 9.64052 | 4239.76 | 0 | 0 | 0.6071429 | surface |
| SYNE4 | 2.6074 | 10.39 | 2598.59 | 0 | 0 | 0.4021739 | n.a. |
| ZNF683 | 2.5327 | 9.47802 | 3266.21 | 0 | 0 | 0.7468944 | n.a. |
| CCL3 | 2.5122 | 9.29866 | 5578.92 | 0 | 0 | 0.5062112 | n.a. |
| CD247 | 2.4977 | 9.41444 | 1876.19 | 0 | 0 | 0.576087 | n.a. |
| ENSECAG00000031528 | 2.2605 | 9.35154 | 3931.25 | 0 | 0 | 0.6040373 | n.a. |
| CD7 | 2.0776 | 9.47261 | 2494.46 | 0 | 0 | 0.8400621 | surface |
| TRDC | 1.9682 | 9.41245 | 2070.82 | 0 | 0 | 0.3664596 | n.a. |
| PRF1 | 1.9175 | 9.44536 | 2662.49 | 0 | 0 | 0.8757764 | n.a. |
| GZMA | 1.6853 | 9.45194 | 1782.16 | 0 | 0 | 0.4968944 | n.a. |
| ENSECAG00000029287 | 1.4478 | 10.0425 | 1407.34 | 6E-302 | 3E-299 | 0.9860248 | n.a. |
| ENSECAG00000032627 | 2.384 | 9.28916 | 4759.91 | 2E-299 | 1E-296 | 0.363354 | n.a. |
| ENSECAG00000030236 | 2.6599 | 9.25249 | 8426.74 | 3E-281 | 1E-278 | 0.3586957 | n.a. |
| CCL5 | 0.8682 | 11.9076 | 1161.91 | 2E-250 | 7E-248 | 0.9937888 | n.a. |
| ENSECAG00000031322 | 1.2753 | 10.4994 | 1124.57 | 1E-242 | 5E-240 | 0.9875776 | n.a. |
| TIMP1 | 2.5006 | 9.33328 | 3565.74 | 1E-242 | 6E-240 | 0.3012422 | n.a. |
| C1H15orf48 | 2.1792 | 9.69292 | 1864.97 | 8E-240 | 3E-237 | 0.3944099 | n.a. |
| ETNK2 | 2.2784 | 9.31226 | 2470.48 | 2E-233 | 8E-231 | 0.4440994 | n.a. |
| ENSECAG00000032959 | 1.3111 | 9.69441 | 872.351 | 2E-189 | 8E-187 | 0.8757764 | n.a. |
| ENSECAG00000016880 | 1.8853 | 9.26997 | 3740.31 | 2E-165 | 6E-163 | 0.3509317 | n.a. |
| LITAF | 1.7744 | 9.32446 | 1332.59 | 9E-154 | 3E-151 | 0.4906832 | n.a. |
| CX3CR1 | 1.8439 | 9.28867 | 3713.08 | 4E-135 | 1E-132 | 0.2717391 | surface |
| KLRK1 | 0.9646 | 9.56899 | 792.188 | 4E-134 | 9E-132 | 0.8618012 | surface |
| ENSECAG00000038714 | 1.7386 | 9.3099 | 1920.91 | 7E-118 | 2E-115 | 0.3385093 | n.a. |
| ADGRG1 | 1.6097 | 9.28684 | 2131.23 | 3E-103 | 6E-101 | 0.363354 | n.a. |
| IGF2 | 1.4111 | 9.32565 | 789.354 | 4E-95 | 9E-93 | 0.4487578 | surface |
| ENSECAG00000039088 | 1.02 | 9.72742 | 430.598 | 5E-95 | 9E-93 | 0.8757764 | n.a. |
| ENSECAG00000015137 | 1.2265 | 9.39252 | 1288.48 | 3E-93 | 6E-91 | 0.4083851 | n.a. |
| FGR | 1.6144 | 9.48827 | 1139.25 | 4E-88 | 7E-86 | 0.2748447 | n.a. |
| ENSECAG00000006663 | 1.419 | 9.31319 | 1533.32 | 4E-81 | 7E-79 | 0.257764 | n.a. |
| CTSW | 0.6683 | 9.89902 | 351.738 | 4E-78 | 7E-76 | 0.9425466 | n.a. |
| SERPINB9 | 1.3559 | 9.38828 | 503.554 | 1E-75 | 2E-73 | 0.3819876 | n.a. |
| ID2 | 0.9187 | 9.64515 | 315.535 | 3E-70 | 4E-68 | 0.7111801 | n.a. |
| RETREG1 | 1.2537 | 9.29791 | 863.707 | 8E-67 | 1E-64 | 0.378882 | n.a. |
| PTPN22 | 1.2658 | 9.3108 | 472.697 | 4E-57 | 6E-55 | 0.2763975 | n.a. |
| SCML4 | 1.0912 | 9.3954 | 308.631 | 2E-56 | 3E-54 | 0.4549689 | n.a. |
| S100A4 | 0.8003 | 11.2922 | 249.797 | 5E-56 | 6E-54 | 0.9130435 | n.a. |
| ABI3 | 1.2512 | 9.33242 | 566.38 | 1E-53 | 1E-51 | 0.2531056 | n.a. |
| ATP2A3 | 1.0311 | 9.49836 | 221.913 | 5E-50 | 6E-48 | 0.5031056 | n.a. |
| PRDX1 | 0.9518 | 9.88573 | 186.626 | 2E-42 | 2E-40 | 0.5093168 | n.a. |
| CST7 | 0.9687 | 9.31203 | 416.629 | 2E-41 | 2E-39 | 0.363354 | n.a. |
| DDIT4 | 1.1008 | 9.33284 | 348.654 | 3E-41 | 3E-39 | 0.257764 | n.a. |
| ANXA2 | 0.7307 | 9.88751 | 174.132 | 1E-39 | 1E-37 | 0.6754658 | n.a. |
| ICAM2 | 0.9266 | 9.48738 | 195.554 | 2E-39 | 2E-37 | 0.4130435 | surface |

| | | | | | | | |
|---|---|---|---|---|---|---|---|
| TRAT1 | 0.6457 | 9.78592 | 151.495 | 1E-34 | 8E-33 | 0.742236 | surface |
| RAP1B | 0.8243 | 9.59007 | 145.392 | 2E-33 | 2E-31 | 0.5403727 | n.a. |
| BZW2 | 0.9089 | 9.4112 | 154.17 | 4E-30 | 3E-28 | 0.3059006 | n.a. |
| HMGB1 | 0.6611 | 10.0857 | 128.756 | 9E-30 | 6E-28 | 0.7531056 | n.a. |
| LGALS3 | 0.7914 | 9.52391 | 170.033 | 2E-29 | 1E-27 | 0.4285714 | n.a. |
| CD8A | 0.6878 | 9.42808 | 162.417 | 7E-29 | 5E-27 | 0.4301242 | surface |
| LUZP6 | 0.964 | 9.43178 | 168.937 | 1E-28 | 8E-27 | 0.2531056 | n.a. |
| NMNAT3 | 0.7549 | 9.39404 | 140.729 | 6E-25 | 3E-23 | 0.3090062 | n.a. |
| S1PR5 | 0.8173 | 9.29327 | 304.928 | 6E-25 | 3E-23 | 0.2810559 | surface |
| ENSECAG00000038252 | 0.7772 | 9.39703 | 132.416 | 1E-24 | 6E-23 | 0.3276398 | n.a. |
| YWHAQ | 0.6261 | 9.70751 | 104.166 | 2E-24 | 1E-22 | 0.6801242 | n.a. |
| SRGN | 0.5801 | 10.4845 | 103.782 | 2E-24 | 1E-22 | 0.7298137 | n.a. |
| TUBA1A | 0.6699 | 10.0912 | 100.6 | 1E-23 | 6E-22 | 0.6381988 | n.a. |
| ENSECAG00000035305 | 0.6914 | 10.0916 | 100.345 | 1E-23 | 7E-22 | 0.6428571 | n.a. |
| LBH | 0.7784 | 9.46361 | 106.458 | 1E-22 | 6E-21 | 0.3198758 | n.a. |
| EID1 | 0.7925 | 9.40846 | 115.783 | 2E-22 | 8E-21 | 0.2841615 | n.a. |
| IL2RB | 0.732 | 9.31659 | 174.237 | 2E-22 | 1E-20 | 0.3431677 | surface |
| SLC44A2 | 0.7855 | 9.42987 | 107.646 | 5E-21 | 2E-19 | 0.2841615 | surface |
| PTPN6 | 0.7607 | 9.58003 | 89.8096 | 2E-20 | 1E-18 | 0.257764 | n.a. |
| CDK2AP2 | 0.6114 | 9.6423 | 82.5184 | 1E-19 | 4E-18 | 0.5139752 | n.a. |
| RAB1A | 0.7523 | 9.43205 | 100.397 | 2E-18 | 6E-17 | 0.2732919 | n.a. |
| ADD3 | 0.6076 | 9.52569 | 63.5771 | 2E-15 | 5E-14 | 0.3649068 | n.a. |
| CARHSP1 | 0.6208 | 9.40606 | 74.6898 | 4E-14 | 1E-12 | 0.2965839 | n.a. |
| TMEM71 | 0.5951 | 9.42449 | 64.0719 | 1E-13 | 3E-12 | 0.2981366 | n.a. |
| ENSECAG00000022488 | 0.6225 | 9.34948 | 79.8239 | 5E-12 | 1E-10 | 0.2562112 | n.a. |

| genes | logFC | logCPM | F | PValue | FDR | percent.exp |
|---|---|---|---|---|---|---|
| GNLY | 4.0016 | 9.3314 | 10831.2 | 0 | 0 | 0.7869416 |
| TRDC | 3.4874 | 9.41245 | 6634.39 | 0 | 0 | 0.790378 |
| KLRB1 | 3.4601 | 9.64052 | 7420.79 | 0 | 0 | 0.9123711 |
| RPLP0 | 0.7168 | 13.1133 | 1261.81 | 2E-271 | 3E-268 | 1 |
| LTB | 1.6216 | 10.6814 | 1258.54 | 8E-271 | 1E-267 | 0.9415808 |
| GZMM | 1.5989 | 9.49614 | 1203.49 | 1E-242 | 1E-239 | 0.838488 |
| ENSECAG00000010622 | 0.9104 | 12.2214 | 1059.08 | 7E-229 | 6E-226 | 0.9948454 |
| RPS12 | 0.6288 | 13.4813 | 1024.35 | 2E-221 | 1E-218 | 1 |
| RPS8 | 0.6677 | 12.8998 | 1020.28 | 1E-220 | 8E-218 | 1 |
| EMB | 2.142 | 9.38136 | 1584.13 | 8E-203 | 5E-200 | 0.4690722 |
| TPT1 | 0.7392 | 12.0825 | 924.383 | 2E-200 | 1E-197 | 0.9931271 |
| RPS15A | 0.598 | 12.8209 | 889.43 | 5E-193 | 3E-190 | 1 |
| SH2D1A | 1.4271 | 9.45149 | 695.718 | 2E-150 | 9E-148 | 0.7079038 |
| KLRF1 | 1.8218 | 9.26172 | 2332.54 | 8E-141 | 2E-138 | 0.338488 |
| COTL1 | 1.2532 | 10.5286 | 579.931 | 4E-127 | 1E-124 | 0.9072165 |
| ENSECAG00000032694 | 1.8248 | 9.28976 | 1345.02 | 4E-115 | 1E-112 | 0.2972509 |
| RPL36A-HNRNPH2 | 0.671 | 11.8526 | 521.011 | 2E-114 | 4E-112 | 0.9879725 |
| TNFRSF25 | 1.7712 | 9.27421 | 1961.22 | 1E-109 | 2E-107 | 0.2508591 |
| CFAP20 | 1.5882 | 9.36468 | 586.289 | 2E-97 | 3E-95 | 0.3814433 |
| TNFSF13B | 1.5421 | 9.32484 | 691.779 | 1E-88 | 1E-86 | 0.3814433 |
| HSP90AB1 | 1.2423 | 10.0849 | 380.065 | 3E-84 | 4E-82 | 0.6821306 |
| ENSECAG00000003681 | 0.7864 | 10.9372 | 375.64 | 3E-83 | 4E-81 | 0.9175258 |
| PLAC8B | 0.9492 | 11.1131 | 363.767 | 1E-80 | 1E-78 | 0.9381443 |
| CAMK4 | 1.5457 | 9.32258 | 775.742 | 6E-79 | 7E-77 | 0.2955326 |
| IL7R | 1.2782 | 9.37306 | 790.106 | 4E-63 | 4E-61 | 0.5549828 |
| ENSECAG00000007668 | 1.0789 | 9.38792 | 288.867 | 2E-58 | 2E-56 | 0.5652921 |
| LAG3 | 1.2227 | 9.29765 | 468.289 | 3E-54 | 3E-52 | 0.3917526 |
| GBP5 | 0.9058 | 9.71631 | 217.655 | 4E-49 | 3E-47 | 0.7800687 |
| PTPMT1 | 1.0197 | 9.41133 | 219.15 | 2E-46 | 1E-44 | 0.4725086 |
| SATB1 | 1.1446 | 9.36591 | 296.292 | 6E-46 | 4E-44 | 0.3436426 |
| GBP6 | 0.9189 | 9.5114 | 183.649 | 1E-41 | 6E-40 | 0.5309278 |
| AQP3 | 1.0991 | 9.3499 | 318.954 | 5E-40 | 3E-38 | 0.2972509 |
| PCMTD1 | 1.0076 | 9.32831 | 241.589 | 2E-35 | 1E-33 | 0.2611684 |
| ZNF706 | 0.8239 | 9.83949 | 151.108 | 1E-34 | 6E-33 | 0.5687285 |
| SSR4 | 0.6002 | 10.501 | 140.192 | 3E-32 | 1E-30 | 0.8350515 |
| PDCD4 | 0.8191 | 9.61093 | 138.145 | 8E-32 | 4E-30 | 0.5498282 |
| TNFSF14 | 0.9942 | 9.28 | 367.908 | 1E-30 | 7E-29 | 0.2783505 |
| CELF1 | 0.8895 | 9.36908 | 157.569 | 3E-29 | 1E-27 | 0.3264605 |
| ENSECAG00000040180 | 0.5999 | 9.7881 | 116.234 | 5E-27 | 2E-25 | 0.7233677 |
| GLIPR2 | 0.6898 | 9.46997 | 131.586 | 8E-23 | 3E-21 | 0.3487973 |
| TSPO | 0.686 | 9.73227 | 121.944 | 5E-22 | 2E-20 | 0.3075601 |
| PIK3R1 | 0.6497 | 9.39634 | 83.3678 | 3E-19 | 9E-18 | 0.4175258 |
| IDO1 | 0.5865 | 9.57828 | 77.3679 | 1E-18 | 5E-17 | 0.5790378 |
| INPP4B | 0.5812 | 9.46609 | 73.3974 | 1E-17 | 4E-16 | 0.4175258 |
| EPB41 | 0.6572 | 9.4445 | 76.4584 | 8E-17 | 2E-15 | 0.3539519 |
| EOMES | 0.6125 | 9.28343 | 198.572 | 2E-16 | 7E-15 | 0.3075601 |
| XPA | 0.5932 | 9.38679 | 66.8224 | 2E-15 | 6E-14 | 0.3608247 |
| SH2D2A | 0.6394 | 9.30769 | 111.052 | 6E-15 | 2E-13 | 0.3591065 |

| Surfacome.Label |
| --- |
| n.a. |
| n.a. |
| surface |
| n.a. |
| n.a. |
| n.a. |
| n.a. |
| n.a. |
| n.a. |
| surface |
| n.a. |
| n.a. |
| n.a. |
| surface |
| n.a. |
| n.a. |
| n.a. |
| surface |
| n.a. |
| surface |
| n.a. |
| n.a. |
| n.a. |
| n.a. |
| surface |
| n.a. |
| surface |
| n.a. |
| n.a. |
| n.a. |
| n.a. |
| n.a. |
| n.a. |
| n.a. |
| n.a. |
| n.a. |
| n.a. |
| n.a. |
| n.a. |
| n.a. |
| n.a. |
| n.a. |
| n.a. |
| n.a. |
| n.a. |
| n.a. |
| n.a. |
| n.a. |

| genes | logFC | logCPM | F | PValue | FDR | percent.exp | Surfacome.Label |
|---|---|---|---|---|---|---|---|
| ENSECAG00000001010 | 2.9007 | 9.62902 | 9167.17 | 0 | 0 | 0.6712046 | n.a. |
| ZNF683 | 2.8526 | 9.47802 | 7527.31 | 0 | 0 | 0.8089934 | n.a. |
| ENSECAG00000022193 | 2.8506 | 9.33233 | 9794.92 | 0 | 0 | 0.4797855 | n.a. |
| ENSECAG00000015137 | 2.6708 | 9.39252 | 9575.45 | 0 | 0 | 0.644802 | n.a. |
| ENSECAG00000008322 | 2.5057 | 9.47262 | 9149.11 | 0 | 0 | 0.8490099 | n.a. |
| ENSECAG00000006663 | 2.2719 | 9.31319 | 6818.08 | 0 | 0 | 0.3910891 | n.a. |
| NPY | 1.8256 | 9.50433 | 2166.14 | 0 | 0 | 0.5082508 | n.a. |
| ENSECAG00000029287 | 1.2972 | 10.0425 | 2303.22 | 0 | 0 | 0.9760726 | n.a. |
| KLRK1 | 1.0787 | 9.56899 | 1911.76 | 0 | 0 | 0.8737624 | surface |
| ENSECAG00000031322 | 1.0397 | 10.4994 | 1519.24 | 0 | 0 | 0.9645215 | n.a. |
| CCL5 | 0.934 | 11.9076 | 2824 | 0 | 0 | 0.9995875 | n.a. |
| B2M | 0.6437 | 13.8602 | 2879.8 | 0 | 0 | 1 | n.a. |
| ETNK2 | 1.8587 | 9.31226 | 3169.46 | 0 | 7E-307 | 0.3358086 | n.a. |
| CD5 | 1.5775 | 9.64992 | 1350.41 | 5E-290 | 2E-287 | 0.5928218 | surface |
| ENSECAG00000009413 | 0.8594 | 10.8806 | 1305.57 | 1E-280 | 4E-278 | 0.9682343 | n.a. |
| CX3CR1 | 1.8275 | 9.28867 | 6158.08 | 2E-257 | 5E-255 | 0.2693894 | surface |
| ADGRG1 | 1.8072 | 9.28684 | 4779.69 | 3E-247 | 7E-245 | 0.4026403 | n.a. |
| S100A4 | 1.044 | 11.2922 | 1058.69 | 9E-229 | 2E-226 | 0.9150165 | n.a. |
| ENSECAG00000032959 | 0.9624 | 9.69441 | 951.016 | 5E-206 | 9E-204 | 0.7908416 | n.a. |
| CD7 | 0.9645 | 9.47261 | 982.996 | 7E-201 | 1E-198 | 0.5969472 | surface |
| ENSECAG00000000419 | 0.7716 | 10.7827 | 841.157 | 9E-183 | 1E-180 | 0.9467822 | n.a. |
| NMNAT3 | 1.3667 | 9.39404 | 961.232 | 3E-178 | 4E-176 | 0.4315182 | n.a. |
| ENSECAG00000019698 | 1.2413 | 9.46862 | 817.276 | 2E-176 | 3E-174 | 0.4892739 | n.a. |
| ENSECAG00000038714 | 1.4038 | 9.3099 | 2646.31 | 2E-175 | 3E-173 | 0.2557756 | n.a. |
| CD8A | 1.0094 | 9.42808 | 763.202 | 5E-166 | 6E-164 | 0.3997525 | surface |
| ENSECAG00000030258 | 0.7655 | 11.3888 | 743.655 | 5E-162 | 6E-160 | 0.960396 | n.a. |
| PRDX1 | 1.2146 | 9.88573 | 734.366 | 5E-160 | 6E-158 | 0.5511551 | n.a. |
| CTSW | 0.594 | 9.89902 | 628.141 | 2E-137 | 2E-135 | 0.9335809 | n.a. |
| CD3G | 0.6044 | 10.3583 | 580.567 | 3E-127 | 3E-125 | 0.9302805 | surface |
| RETREG1 | 1.1871 | 9.29791 | 1488.97 | 4E-124 | 4E-122 | 0.3605611 | n.a. |
| CCL3 | 1.1102 | 9.29866 | 2068.5 | 4E-124 | 4E-122 | 0.2590759 | n.a. |
| LGALS3 | 1.0372 | 9.52391 | 672.324 | 1E-120 | 1E-118 | 0.4674092 | n.a. |
| SERPINB9 | 1.1443 | 9.38828 | 774.273 | 2E-119 | 2E-117 | 0.3333333 | n.a. |
| ID2 | 0.7751 | 9.64515 | 513.015 | 9E-113 | 8E-111 | 0.6914191 | n.a. |
| IFI27 | 1.0818 | 9.73139 | 506.892 | 2E-111 | 2E-109 | 0.4261551 | n.a. |
| DDIT4 | 1.1729 | 9.33284 | 851.716 | 2E-102 | 2E-100 | 0.2582508 | n.a. |
| ANXA2 | 0.7099 | 9.88751 | 393.909 | 4E-87 | 2E-85 | 0.6431518 | n.a. |
| ENSECAG00000039088 | 0.6409 | 9.72742 | 375.095 | 4E-83 | 2E-81 | 0.7871287 | n.a. |
| S1PR5 | 0.9537 | 9.29327 | 847.973 | 6E-72 | 3E-70 | 0.2924917 | surface |
| ADGRE5 | 0.7366 | 9.57906 | 304.543 | 7E-68 | 4E-66 | 0.5540429 | n.a. |
| ATP2A3 | 0.7776 | 9.49836 | 289.926 | 9E-65 | 5E-63 | 0.4026403 | n.a. |
| YWHAQ | 0.6625 | 9.70751 | 280.952 | 8E-63 | 4E-61 | 0.6588284 | n.a. |
| ALDOA | 0.6659 | 9.98996 | 252.946 | 9E-57 | 5E-55 | 0.5631188 | n.a. |
| RAB1A | 0.8436 | 9.43205 | 300.722 | 1E-56 | 6E-55 | 0.2636139 | n.a. |
| CALR | 0.6981 | 9.79425 | 249.916 | 4E-56 | 2E-54 | 0.5346535 | n.a. |
| ITGAL | 0.7152 | 9.4047 | 278.185 | 3E-53 | 1E-51 | 0.3869637 | surface |
| CDK2AP2 | 0.6517 | 9.6423 | 228.274 | 2E-51 | 1E-49 | 0.4938119 | n.a. |
| ITGB2 | 0.6269 | 9.81007 | 220.456 | 1E-49 | 5E-48 | 0.5759076 | surface |
| ITM2C | 0.6867 | 9.59936 | 222.687 | 4E-49 | 2E-47 | 0.3997525 | surface |
| SOD1 | 0.6712 | 9.83911 | 206.948 | 9E-47 | 4E-45 | 0.5173267 | n.a. |

| | | | | | | | |
|---|---|---|---|---|---|---|---|
| ENSECAG00000035021 | 0.6812 | 9.50832 | 209.039 | 9E-46 | 4E-44 | 0.3729373 | n.a. |
| NPC2 | 0.6034 | 9.83667 | 195.786 | 2E-44 | 1E-42 | 0.4872112 | n.a. |
| BATF | 0.7192 | 9.3505 | 295.962 | 3E-43 | 1E-41 | 0.2673267 | n.a. |
| FLNA | 0.7386 | 9.4126 | 270.563 | 5E-41 | 2E-39 | 0.2801155 | n.a. |
| SCML4 | 0.6116 | 9.3954 | 213.335 | 2E-38 | 9E-37 | 0.355198 | n.a. |
| MSN | 0.5841 | 9.61231 | 166.801 | 5E-38 | 2E-36 | 0.44967 | n.a. |
| ENSECAG00000038252 | 0.6157 | 9.39703 | 197.273 | 4E-37 | 1E-35 | 0.3023927 | n.a. |
| CCND3 | 0.6018 | 9.72794 | 160.663 | 1E-36 | 4E-35 | 0.4657591 | n.a. |
| ARRB2 | 0.6109 | 9.46619 | 172.733 | 1E-33 | 4E-32 | 0.3234323 | n.a. |
| LGALS3BP | 0.6197 | 9.42042 | 160.357 | 2E-29 | 5E-28 | 0.2520627 | n.a. |

| genes | logFC | logCPM | F | PValue | FDR | percent.exp | Surfacome.Label |
|---|---|---|---|---|---|---|---|
| GZMK | 3.6008 | 9.57824 | 13770.8 | 0 | 0 | 0.7859696 | n.a. |
| CD27 | 1.8854 | 9.47772 | 1993.44 | 0 | 0 | 0.4557641 | surface |
| RPS19 | 0.5876 | 13.4903 | 2249.67 | 0 | 0 | 0.9995532 | n.a. |
| DAPL1 | 1.8503 | 9.31506 | 2733.58 | 3E-253 | 9E-251 | 0.2904379 | n.a. |
| GZMM | 1.0873 | 9.49614 | 1238.34 | 4E-213 | 9E-211 | 0.7533512 | n.a. |
| ENSECAG00000028035 | 1.5109 | 9.3107 | 2412.72 | 2E-202 | 4E-200 | 0.4048257 | n.a. |
| GPR183 | 1.5558 | 9.66209 | 898.899 | 5E-195 | 9E-193 | 0.3346738 | surface |
| COTL1 | 1.0077 | 10.5286 | 723.207 | 1E-157 | 2E-155 | 0.8722073 | n.a. |
| LY6E | 0.9538 | 10.2408 | 660.803 | 2E-144 | 3E-142 | 0.7940125 | surface |
| CXCR3 | 1.3036 | 9.31033 | 1610.5 | 6E-134 | 7E-132 | 0.2993744 | surface |
| LTB | 0.7261 | 10.6814 | 468.148 | 4E-103 | 3E-101 | 0.883378 | n.a. |
| CXCR4 | 1.1329 | 9.61766 | 457.085 | 9E-101 | 7E-99 | 0.3476318 | surface |
| PLAC8B | 0.729 | 11.1131 | 443.647 | 7E-98 | 5E-96 | 0.8981233 | n.a. |
| IL7R | 1.0085 | 9.37306 | 1070.63 | 1E-78 | 8E-77 | 0.4553172 | surface |
| CD2 | 0.6311 | 9.83383 | 293.038 | 2E-65 | 1E-63 | 0.74084 | surface |
| ENSECAG00000019029 | 0.7921 | 9.92782 | 270.688 | 1E-60 | 7E-59 | 0.471403 | n.a. |
| FYB1 | 0.7296 | 9.60722 | 252.116 | 1E-56 | 7E-55 | 0.4647006 | n.a. |
| FUCA1 | 0.7847 | 9.36777 | 315.191 | 2E-50 | 7E-49 | 0.3092046 | n.a. |
| NPM3 | 0.7775 | 9.63221 | 216.948 | 6E-49 | 3E-47 | 0.3194817 | n.a. |
| IDO1 | 0.6157 | 9.57828 | 211.821 | 8E-48 | 3E-46 | 0.5630027 | n.a. |
| HSP90AB1 | 0.6099 | 10.0849 | 175.958 | 5E-40 | 2E-38 | 0.5357462 | n.a. |
