## Supplementary material for "Single cell resolution landscape of equine peripheral blood mononuclear cells reveals diverse immune cell subtypes including T-bet^+^ B cells": Dataset S8

| genes | logFC | logCPM | F | PValue | FDR | percent.exp | Surfacome.Label |
|---|---|---|---|---|---|---|---|
| STMN1 | 1.034 | 9.7683 | 1720 | 0 | 0 | 0.7903814 | n.a. |
| LTB | 0.687 | 10.681 | 986.98 | 1E-213 | 3E-210 | 0.960199 | n.a. |
| GPR183 | 0.823 | 9.6621 | 753.67 | 4E-164 | 4E-161 | 0.5276949 | surface |
| ENSECAG00000034569 | 0.689 | 9.7713 | 573.96 | 8E-126 | 6E-123 | 0.5864013 | n.a. |
| C4orf48 | 0.651 | 9.5904 | 460.22 | 2E-101 | 1E-98 | 0.5605307 | n.a. |

| genes | logFC | logCPM | F | PValue | FDR | percent.exp | Surfacome.Label |
|---|---|---|---|---|---|---|---|
| LGALS1 | 1.587 | 11.109 | 2115.5 | 0 | 0 | 0.792926 | n.a. |
| GPR183 | 1.577 | 9.6621 | 1852.7 | 0 | 0 | 0.7241158 | surface |
| S100A11 | 1.299 | 9.847 | 1534.6 | 0 | 0 | 0.6707395 | n.a. |
| ENSECAG00000029575 | 1.575 | 9.2895 | 5714.4 | 0 | 0 | 0.2610932 | n.a. |
| TNFRSF18 | 1.55 | 9.34 | 2358.5 | 0 | 0 | 0.4102894 | surface |
| GATA3 | 1.484 | 9.3436 | 2012.5 | 5E-277 | 4E-274 | 0.3993569 | n.a. |
| S100A4 | 1.229 | 11.292 | 1144.2 | 9E-247 | 6E-244 | 0.7459807 | n.a. |
| TNFRSF4 | 1.343 | 9.2923 | 3097.1 | 1E-220 | 8E-218 | 0.2861736 | surface |
| C4orf48 | 1.122 | 9.5904 | 879.1 | 8E-191 | 4E-188 | 0.6649518 | n.a. |
| INPP4B | 1.095 | 9.4661 | 832.07 | 2E-159 | 1E-156 | 0.4508039 | n.a. |
| SRGN | 0.882 | 10.485 | 650.47 | 4E-142 | 1E-139 | 0.7736334 | n.a. |
| IGFLR1 | 0.932 | 9.4383 | 699.79 | 4E-133 | 2E-130 | 0.4212219 | surface |
| SMCO4 | 0.997 | 9.5303 | 592.87 | 7E-130 | 3E-127 | 0.4707395 | n.a. |
| F2R | 1.115 | 9.3054 | 1324.8 | 5E-117 | 2E-114 | 0.266881 | surface |
| CD82 | 1.008 | 9.4652 | 601.85 | 1E-104 | 3E-102 | 0.266881 | surface |
| EMP3 | 0.762 | 10.126 | 474.37 | 2E-104 | 5E-102 | 0.807717 | surface |
| PDCD4 | 0.879 | 9.6109 | 394.44 | 3E-87 | 7E-85 | 0.4700965 | n.a. |
| PKM | 0.747 | 9.8077 | 375.87 | 3E-83 | 7E-81 | 0.65209 | n.a. |
| PLA2G16 | 0.802 | 9.6144 | 390.59 | 4E-82 | 9E-80 | 0.34791 | n.a. |
| ACP5 | 0.749 | 9.7992 | 357.08 | 3E-79 | 7E-77 | 0.6765273 | n.a. |
| CYBA | 0.614 | 10.509 | 277.96 | 4E-62 | 6E-60 | 0.6392283 | n.a. |
| MISP3 | 0.679 | 9.507 | 238.9 | 1E-53 | 1E-51 | 0.3890675 | n.a. |
| HSPA1A | 0.781 | 9.3501 | 362.29 | 2E-52 | 3E-50 | 0.2636656 | n.a. |
| ENSECAG00000004031 | 0.641 | 9.5374 | 217.05 | 6E-49 | 7E-47 | 0.3935691 | n.a. |
| REEP5 | 0.649 | 9.6201 | 215.84 | 1E-48 | 1E-46 | 0.4527331 | n.a. |
| SEC61B | 0.581 | 9.8749 | 209.69 | 2E-47 | 3E-45 | 0.5286174 | n.a. |
| eca-mir-1892 | 0.617 | 9.4799 | 218.09 | 1E-44 | 1E-42 | 0.2797428 | n.a. |

| genes | logFC | logCPM | F | PValue | FDR | percent.exp | Surfacome.Label |
|---|---|---|---|---|---|---|---|
| RPS12 | 0.629 | 13.481 | 1924.8 | 0 | 0 | 1 | n.a. |
| LGALS1 | 0.803 | 11.109 | 367.47 | 2E-81 | 3E-79 | 0.6288981 | n.a. |
| ITGB1 | 0.773 | 9.5309 | 346.68 | 5E-77 | 7E-75 | 0.3575884 | surface |
| HSP90AB1 | 0.743 | 10.085 | 292.95 | 2E-65 | 2E-63 | 0.7494802 | n.a. |
| CISH | 0.729 | 9.3863 | 296.08 | 7E-48 | 5E-46 | 0.2713098 | n.a. |
| INPP4B | 0.645 | 9.4661 | 204.68 | 3E-43 | 2E-41 | 0.3555094 | n.a. |
| IL7R | 0.689 | 9.3731 | 474.92 | 4E-43 | 3E-41 | 0.5561331 | surface |
| NPM3 | 0.681 | 9.6322 | 169.22 | 1E-38 | 9E-37 | 0.510395 | n.a. |

| genes | logFC | logCPM | F | PValue | FDR | percent.exp | Surfacome.Label |
|---|---|---|---|---|---|---|---|
| CD200 | 2.206 | 9.2494 | 18157 | 0 | 0 | 0.2793462 | surface |
| ID3 | 1.9 | 9.6697 | 1579.1 | 0 | 0 | 0.8306092 | n.a. |
| S100A6 | 1.898 | 10.437 | 1687.6 | 0 | 0 | 0.8514116 | n.a. |
| CD27 | 1.686 | 9.4777 | 1245.8 | 4E-268 | 7E-265 | 0.6448737 | surface |
| S100A5 | 1.647 | 9.5834 | 1194.1 | 3E-257 | 4E-254 | 0.3878158 | n.a. |
| UBAC2 | 2.015 | 9.3221 | 1827.6 | 7E-239 | 7E-236 | 0.2986627 | surface |
| DRA | 1.343 | 11.742 | 1088.9 | 4E-235 | 3E-232 | 0.9673105 | n.a. |
| GPR183 | 1.647 | 9.6621 | 1031.9 | 4E-223 | 3E-220 | 0.7013373 | surface |
| SH2D1A | 1.512 | 9.4515 | 958.37 | 1E-207 | 1E-204 | 0.4769688 | n.a. |
| DRB | 1.248 | 11.205 | 819.13 | 4E-178 | 3E-175 | 0.922734 | n.a. |
| S100A11 | 1.289 | 9.847 | 786.71 | 3E-171 | 2E-168 | 0.5943536 | n.a. |
| CD74 | 0.856 | 12.628 | 704.58 | 1E-153 | 5E-151 | 0.9940565 | surface |
| CPPED1 | 1.526 | 9.3236 | 1025.5 | 2E-147 | 8E-145 | 0.332838 | n.a. |
| STK39 | 1.331 | 9.3069 | 1121.6 | 5E-146 | 2E-143 | 0.3239227 | n.a. |
| PLP2 | 1.263 | 9.9939 | 659.25 | 5E-144 | 2E-141 | 0.8068351 | n.a. |
| DQA | 1.279 | 10.712 | 620.37 | 1E-135 | 3E-133 | 0.7102526 | n.a. |
| WARS | 1.243 | 9.8298 | 600.22 | 2E-131 | 6E-129 | 0.8023774 | n.a. |
| NKG7 | 1.371 | 9.3783 | 1382 | 6E-127 | 2E-124 | 0.2897474 | n.a. |
| Eqca-DQB1 | 1.054 | 10.925 | 532.48 | 6E-117 | 2E-114 | 0.8885587 | n.a. |
| MNDA | 1.214 | 10.258 | 521.73 | 1E-114 | 3E-112 | 0.6849926 | n.a. |
| IER3 | 1.394 | 9.3418 | 716.27 | 2E-105 | 6E-103 | 0.3447251 | n.a. |
| CTSW | 1.174 | 9.899 | 628.04 | 7E-104 | 2E-101 | 0.3016345 | n.a. |
| POU2F2 | 1.38 | 9.5681 | 560.72 | 2E-100 | 4E-98 | 0.3135215 | n.a. |
| ANXA2 | 1.021 | 9.8875 | 451.89 | 1E-99 | 3E-97 | 0.5720654 | n.a. |
| ENSECAG00000034569 | 1.087 | 9.7713 | 433.93 | 9E-96 | 2E-93 | 0.5913819 | n.a. |
| SYTL3 | 1.233 | 9.3361 | 894.58 | 1E-93 | 2E-91 | 0.2927192 | n.a. |
| DQB | 1.158 | 10.011 | 395.99 | 1E-87 | 2E-85 | 0.5572065 | n.a. |
| DCTD | 1.23 | 9.3429 | 602.4 | 3E-87 | 4E-85 | 0.2704309 | n.a. |
| RF00581 | 0.805 | 10.319 | 357.14 | 3E-79 | 4E-77 | 0.872214 | n.a. |
| IDO1 | 1.016 | 9.5783 | 315.13 | 3E-70 | 4E-68 | 0.4769688 | n.a. |
| YPEL3 | 1.065 | 9.6613 | 314.72 | 4E-70 | 5E-68 | 0.5884101 | n.a. |
| ENSECAG00000000419 | 0.596 | 10.783 | 296.08 | 4E-66 | 5E-64 | 0.9554235 | n.a. |
| ACTN1 | 1.012 | 9.4643 | 291.53 | 4E-65 | 5E-63 | 0.48737 | n.a. |
| FYN | 0.917 | 9.4235 | 285.74 | 8E-64 | 8E-62 | 0.3580981 | n.a. |
| HCST | 0.92 | 9.5985 | 251.57 | 2E-56 | 2E-54 | 0.4962853 | n.a. |
| GID8 | 0.972 | 9.4665 | 242.7 | 2E-54 | 1E-52 | 0.3759287 | n.a. |
| FABP5 | 0.68 | 9.5264 | 241.64 | 3E-54 | 2E-52 | 0.2808321 | n.a. |
| TESC | 0.855 | 9.3671 | 272.65 | 2E-48 | 2E-46 | 0.2734027 | n.a. |
| ARPC3 | 0.748 | 10.061 | 210.52 | 1E-47 | 1E-45 | 0.6062407 | n.a. |
| ANXA1 | 0.836 | 9.8295 | 196.98 | 1E-44 | 1E-42 | 0.5750371 | n.a. |
| AQP3 | 0.888 | 9.3499 | 314.9 | 4E-44 | 3E-42 | 0.2882615 | n.a. |
| ACTG1 | 0.636 | 10.745 | 191.8 | 2E-43 | 1E-41 | 0.8320951 | n.a. |
| P2RY10 | 0.869 | 9.4226 | 199.96 | 1E-42 | 9E-41 | 0.3610698 | surface |
| HLA-DMA | 0.924 | 9.6051 | 243.27 | 3E-41 | 2E-39 | 0.3150074 | surface |
| NAAA | 0.856 | 9.5818 | 180.03 | 6E-41 | 4E-39 | 0.4502229 | n.a. |
| SAMHD1 | 0.797 | 9.4265 | 179.47 | 8E-41 | 6E-39 | 0.3521545 | n.a. |
| RF02216 | 0.751 | 9.6717 | 178.08 | 2E-40 | 1E-38 | 0.5156018 | n.a. |

| | | | | | | | |
|---|---|---|---|---|---|---|---|
| ENSECAG00000010008 | 0.818 | 9.4684 | 170.41 | 7E-39 | 5E-37 | 0.3135215 | n.a. |
| MRPS6 | 0.855 | 9.4746 | 175.63 | 4E-37 | 3E-35 | 0.2971768 | n.a. |
| ENSECAG00000020136 | 0.752 | 9.7014 | 162.3 | 4E-37 | 3E-35 | 0.4130758 | n.a. |
| CTSC | 0.745 | 9.6133 | 162.23 | 4E-37 | 3E-35 | 0.4962853 | n.a. |
| ENSECAG00000038584 | 0.604 | 10.313 | 157.97 | 4E-36 | 2E-34 | 0.8558692 | n.a. |
| CD44 | 0.598 | 10.108 | 143.73 | 5E-33 | 3E-31 | 0.7132244 | surface |
| ENSECAG00000032776 | 0.776 | 9.4539 | 138.73 | 6E-32 | 3E-30 | 0.2867756 | n.a. |
| UBE2R2 | 0.762 | 9.356 | 179.08 | 3E-31 | 1E-29 | 0.2897474 | n.a. |
| RAP1B | 0.739 | 9.5901 | 131.29 | 2E-30 | 1E-28 | 0.384844 | n.a. |
| RCSD1 | 0.712 | 9.6846 | 129.48 | 6E-30 | 3E-28 | 0.5453195 | n.a. |
| ENSECAG00000027676 | 0.655 | 9.7058 | 123.75 | 1E-28 | 5E-27 | 0.4680535 | n.a. |
| ENSECAG00000032710 | 0.664 | 9.3761 | 126.6 | 3E-28 | 1E-26 | 0.2867756 | n.a. |
| ITGB2 | 0.631 | 9.8101 | 116.04 | 5E-27 | 2E-25 | 0.5616642 | surface |
| MYCBP2 | 0.676 | 9.5377 | 111.78 | 4E-26 | 2E-24 | 0.3818722 | n.a. |
| FAM107B | 0.663 | 9.5005 | 108.67 | 2E-25 | 9E-24 | 0.4249629 | n.a. |
| ENSECAG00000036564 | 0.614 | 9.6716 | 105.86 | 9E-25 | 4E-23 | 0.5661218 | n.a. |
| eca-mir-9023 | 0.709 | 9.4296 | 118.06 | 4E-24 | 2E-22 | 0.3179792 | n.a. |
| NPC2 | 0.603 | 9.8367 | 99.275 | 2E-23 | 9E-22 | 0.4279346 | n.a. |
| ITGA4 | 0.582 | 9.465 | 94.86 | 2E-22 | 8E-21 | 0.3239227 | surface |
| ANXA11 | 0.617 | 9.4886 | 93.808 | 4E-22 | 1E-20 | 0.282318 | n.a. |
| TCF7 | 0.589 | 9.3798 | 124.66 | 6E-20 | 2E-18 | 0.3120357 | n.a. |
| XPA | 0.595 | 9.3868 | 90 | 1E-18 | 4E-17 | 0.2526003 | n.a. |

| genes | logFC | logCPM | F | PValue | FDR | percent.exp | Surfacome.Label |
|---|---|---|---|---|---|---|---|
| ENSECAG00000030502 | 2.428 | 9.3968 | 8450.9 | 0 | 0 | 0.531293 | n.a. |
| CCR7 | 1.674 | 9.4631 | 3015.4 | 0 | 0 | 0.5876891 | surface |
| SELL | 1.455 | 9.6904 | 1752.8 | 0 | 0 | 0.5818432 | surface |
| ENSECAG00000028304 | 1.155 | 9.3449 | 1698.6 | 9E-215 | 1E-212 | 0.2709766 | n.a. |
| OXNAD1 | 1.269 | 9.3622 | 1623.9 | 7E-214 | 1E-211 | 0.3590096 | n.a. |
| RGS10 | 1.092 | 9.5903 | 961.3 | 4E-208 | 5E-206 | 0.4993122 | n.a. |
| LEF1 | 1.001 | 9.4498 | 1062.2 | 3E-176 | 4E-174 | 0.4951857 | n.a. |
| TXNIP | 0.879 | 10.204 | 789.02 | 1E-171 | 1E-169 | 0.8304677 | n.a. |
| PLAC8B | 0.589 | 11.113 | 437.67 | 1E-96 | 9E-95 | 0.8641678 | n.a. |
| ENSECAG00000034569 | 0.657 | 9.7713 | 433.71 | 1E-95 | 7E-94 | 0.5870014 | n.a. |
| FKBP5 | 0.785 | 9.3784 | 587.34 | 2E-93 | 1E-91 | 0.3184319 | n.a. |
| ENSECAG00000030387 | 0.832 | 9.433 | 784.41 | 1E-92 | 9E-91 | 0.2506878 | n.a. |
| TRAT1 | 0.606 | 9.7859 | 345.03 | 1E-76 | 7E-75 | 0.5952545 | surface |
| ADGB | 0.661 | 9.3514 | 496.37 | 3E-62 | 2E-60 | 0.2647868 | n.a. |
| CD69 | 0.65 | 9.4382 | 333.31 | 8E-62 | 4E-60 | 0.3480055 | surface |
| RF00213 | 0.683 | 9.3791 | 371.15 | 2E-61 | 1E-59 | 0.2568776 | n.a. |
| PLA2G16 | 0.621 | 9.6144 | 315.89 | 4E-50 | 2E-48 | 0.30674 | n.a. |

| genes | logFC | logCPM | F | PValue | FDR | percent.exp | Surfacome.Label |
|---|---|---|---|---|---|---|---|
| LGALS1 | 1.985 | 11.109 | 1479.9 | 0 | 0 | 0.8430769 | n.a. |
| ENSECAG00000009190 | 0.656 | 14.441 | 884.21 | 7E-192 | 3E-188 | 1 | n.a. |
| ACTB | 0.925 | 12.301 | 698.31 | 2E-152 | 6E-149 | 1 | n.a. |
| PFN1 | 0.808 | 11.941 | 645.06 | 5E-141 | 1E-137 | 1 | n.a. |
| ZNF683 | 1.564 | 9.478 | 1050.2 | 6E-131 | 9E-128 | 0.2584615 | n.a. |
| IFI27 | 1.536 | 9.7314 | 565.06 | 7E-124 | 9E-121 | 0.6061538 | n.a. |
| COX1 | 0.725 | 12.246 | 551.36 | 6E-121 | 6E-118 | 1 | n.a. |
| ARPC3 | 1.296 | 10.061 | 504.38 | 7E-111 | 6E-108 | 0.9353846 | n.a. |
| ACTG1 | 1.109 | 10.745 | 447.76 | 9E-99 | 7E-96 | 0.9784615 | n.a. |
| SUB1 | 1.066 | 10.333 | 420.45 | 7E-93 | 5E-90 | 0.9876923 | n.a. |
| ENSECAG00000037706 | 1 | 10.368 | 396.36 | 1E-87 | 6E-85 | 0.9907692 | n.a. |
| CCL5 | 1.42 | 11.908 | 387.09 | 1E-85 | 6E-83 | 0.2830769 | n.a. |
| TNFRSF18 | 1.468 | 9.34 | 625.04 | 1E-84 | 7E-82 | 0.4892308 | surface |
| SH3BGRL3 | 0.857 | 10.736 | 356.72 | 4E-79 | 2E-76 | 1 | n.a. |
| ENSECAG00000029003 | 1.068 | 10.041 | 353.13 | 2E-78 | 1E-75 | 0.9753846 | n.a. |
| COTL1 | 0.976 | 10.529 | 345.81 | 8E-77 | 3E-74 | 0.9692308 | n.a. |
| UCP2 | 1.155 | 9.9052 | 344.02 | 2E-76 | 7E-74 | 0.7907692 | n.a. |
| ENSECAG00000030745 | 0.611 | 11.502 | 338.2 | 4E-75 | 1E-72 | 1 | n.a. |
| SLC25A5 | 0.65 | 11.478 | 335.55 | 1E-74 | 5E-72 | 1 | n.a. |
| MNDA | 1.202 | 10.258 | 334.4 | 2E-74 | 8E-72 | 0.8215385 | n.a. |
| ENSECAG00000029575 | 1.333 | 9.2895 | 1164.7 | 2E-73 | 7E-71 | 0.2553846 | n.a. |
| S100A4 | 1.046 | 11.292 | 322.11 | 1E-71 | 3E-69 | 0.7138462 | n.a. |
| S100A11 | 0.977 | 9.847 | 320.82 | 2E-71 | 6E-69 | 0.7015385 | n.a. |
| IGFLR1 | 1.181 | 9.4383 | 314.26 | 5E-70 | 1E-67 | 0.7138462 | surface |
| ENSECAG00000034569 | 1.136 | 9.7713 | 293.54 | 2E-65 | 4E-63 | 0.6092308 | n.a. |
| FABP5 | 0.94 | 9.5264 | 288.37 | 2E-64 | 5E-62 | 0.5661538 | n.a. |
| IFI30 | 1.105 | 9.9429 | 287.52 | 3E-64 | 8E-62 | 0.5907692 | n.a. |
| MYL6 | 0.771 | 10.731 | 284.72 | 1E-63 | 3E-61 | 1 | n.a. |
| ANXA2 | 0.903 | 9.8875 | 253.48 | 7E-57 | 1E-54 | 0.7846154 | n.a. |
| ARPC2 | 0.756 | 10.591 | 248.02 | 1E-55 | 2E-53 | 0.9876923 | n.a. |
| ENSECAG00000000419 | 0.706 | 10.783 | 239.65 | 7E-54 | 1E-51 | 0.9938462 | n.a. |
| ENSECAG00000031569 | 0.764 | 11.194 | 236.14 | 4E-53 | 8E-51 | 0.9969231 | n.a. |
| ENSECAG00000027676 | 1.038 | 9.7058 | 226.16 | 6E-51 | 1E-48 | 0.7569231 | n.a. |
| RAC2 | 0.708 | 10.594 | 213.62 | 3E-48 | 5E-46 | 0.9938462 | n.a. |
| RF00581 | 0.78 | 10.319 | 206.08 | 1E-46 | 2E-44 | 0.9753846 | n.a. |
| STK39 | 1.063 | 9.3069 | 386.56 | 2E-46 | 2E-44 | 0.3876923 | n.a. |
| EMB | 0.731 | 9.3814 | 318.49 | 4E-46 | 6E-44 | 0.2584615 | surface |
| SAMHD1 | 1.073 | 9.4265 | 192.64 | 1E-43 | 2E-41 | 0.6246154 | n.a. |
| WDR1 | 0.912 | 9.7464 | 189.86 | 4E-43 | 6E-41 | 0.7753846 | n.a. |
| SRGN | 0.852 | 10.485 | 189.76 | 5E-43 | 7E-41 | 0.8615385 | n.a. |
| ITGB1 | 0.819 | 9.5309 | 183.4 | 1E-41 | 2E-39 | 0.3630769 | surface |
| TNFRSF4 | 1.05 | 9.2923 | 549.95 | 1E-40 | 2E-38 | 0.3076923 | surface |
| LSP1 | 0.731 | 10.243 | 177.91 | 2E-40 | 2E-38 | 0.9415385 | n.a. |
| CD44 | 0.777 | 10.108 | 166.05 | 7E-38 | 8E-36 | 0.9107692 | surface |
| ENSECAG00000006071 | 0.901 | 9.6095 | 160.19 | 1E-36 | 1E-34 | 0.7384615 | n.a. |
| GLIPR1 | 0.812 | 9.887 | 147.48 | 7E-34 | 8E-32 | 0.7538462 | surface |
| ENSECAG00000019073 | 0.775 | 9.919 | 147.13 | 9E-34 | 9E-32 | 0.8615385 | n.a. |

| | | | | | | | |
|---|---|---|---|---|---|---|---|
| CIB1 | 0.817 | 9.4458 | 142.34 | 1E-32 | 1E-30 | 0.5261538 | n.a. |
| DYNLRB1 | 0.797 | 9.8334 | 141.37 | 2E-32 | 2E-30 | 0.9015385 | n.a. |
| LY6E | 0.685 | 10.241 | 140.99 | 2E-32 | 2E-30 | 0.96 | surface |
| ENSECAG00000029287 | 0.62 | 10.043 | 129.92 | 5E-30 | 5E-28 | 0.3015385 | n.a. |
| PTPRCAP | 0.602 | 10.617 | 128.87 | 8E-30 | 8E-28 | 0.9630769 | n.a. |
| ENSECAG00000027666 | 0.739 | 9.8957 | 128.17 | 1E-29 | 1E-27 | 0.8184615 | n.a. |
| DBI | 0.753 | 9.673 | 122.44 | 2E-28 | 2E-26 | 0.64 | n.a. |
| ENSECAG00000020136 | 0.779 | 9.7014 | 122.06 | 2E-28 | 2E-26 | 0.6123077 | n.a. |
| CAPZA1 | 0.776 | 9.7535 | 118.16 | 2E-27 | 1E-25 | 0.8123077 | n.a. |
| CORO1A | 0.608 | 10.247 | 115.12 | 8E-27 | 7E-25 | 0.9723077 | n.a. |
| DCTD | 0.792 | 9.3429 | 163.51 | 1E-26 | 9E-25 | 0.3446154 | n.a. |
| JPT1 | 0.73 | 9.7243 | 111.92 | 4E-26 | 3E-24 | 0.7384615 | n.a. |
| HSBP1 | 0.72 | 9.5012 | 108.29 | 3E-25 | 2E-23 | 0.4676923 | n.a. |
| RAB37 | 0.808 | 9.2869 | 306.19 | 4E-25 | 3E-23 | 0.2553846 | n.a. |
| GATA3 | 0.894 | 9.3436 | 190.14 | 5E-25 | 3E-23 | 0.3784615 | n.a. |
| ENSECAG00000019698 | 0.804 | 9.4686 | 99.397 | 2E-23 | 1E-21 | 0.5569231 | n.a. |
| ARPC5 | 0.682 | 9.8016 | 99.363 | 2E-23 | 1E-21 | 0.8215385 | n.a. |
| CAPZB | 0.694 | 9.814 | 98.55 | 3E-23 | 2E-21 | 0.8061538 | n.a. |
| ATP2A3 | 0.743 | 9.4984 | 91.808 | 1E-21 | 6E-20 | 0.5446154 | n.a. |
| CD5 | 0.685 | 9.6499 | 90.27 | 2E-21 | 1E-19 | 0.7784615 | surface |
| ATP5F1C | 0.637 | 9.7509 | 88.812 | 5E-21 | 3E-19 | 0.7784615 | n.a. |
| ENSECAG00000009520 | 0.765 | 9.3369 | 111.72 | 6E-21 | 3E-19 | 0.4307692 | n.a. |
| ENSECAG00000028107 | 0.594 | 9.3832 | 87.086 | 1E-20 | 6E-19 | 0.3784615 | n.a. |
| IDH2 | 0.647 | 9.6001 | 86.785 | 1E-20 | 7E-19 | 0.6830769 | n.a. |
| ITGAL | 0.695 | 9.4047 | 104.12 | 2E-20 | 1E-18 | 0.3415385 | surface |
| NPC2 | 0.646 | 9.8367 | 83.865 | 6E-20 | 3E-18 | 0.6646154 | n.a. |
| EMP3 | 0.581 | 10.126 | 83.667 | 6E-20 | 3E-18 | 0.8738462 | surface |
| PPP1CA | 0.632 | 9.7882 | 83.58 | 6E-20 | 3E-18 | 0.8215385 | n.a. |
| ENSECAG00000028768 | 0.656 | 9.3068 | 163.83 | 1E-19 | 6E-18 | 0.2707692 | n.a. |
| PSMB2 | 0.653 | 9.5898 | 82.138 | 1E-19 | 7E-18 | 0.6646154 | n.a. |
| CAPG | 0.662 | 9.6557 | 82.033 | 1E-19 | 7E-18 | 0.6923077 | n.a. |
| SAT1 | 0.633 | 9.7271 | 81.777 | 2E-19 | 8E-18 | 0.3907692 | n.a. |
| RAC1 | 0.607 | 9.9054 | 80.74 | 3E-19 | 1E-17 | 0.7692308 | n.a. |
| ENSECAG00000010008 | 0.692 | 9.4684 | 80.291 | 3E-19 | 2E-17 | 0.5138462 | n.a. |
| NAA10 | 0.68 | 9.5568 | 79 | 7E-19 | 3E-17 | 0.7261538 | n.a. |
| APOBEC3H | 0.706 | 9.3557 | 83.551 | 2E-18 | 9E-17 | 0.4061538 | n.a. |
| SULT2B1 | 0.87 | 9.3135 | 111.94 | 4E-18 | 2E-16 | 0.4061538 | n.a. |
| TPST2 | 0.658 | 9.4709 | 74.214 | 7E-18 | 3E-16 | 0.5353846 | n.a. |
| PSMB4 | 0.636 | 9.6149 | 73.496 | 1E-17 | 5E-16 | 0.6738462 | n.a. |
| ENSECAG00000020462 | 0.602 | 9.6751 | 73.276 | 1E-17 | 5E-16 | 0.6 | n.a. |
| RF02216 | 0.618 | 9.6717 | 72.297 | 2E-17 | 8E-16 | 0.6830769 | n.a. |
| PSMD9 | 0.648 | 9.3271 | 124.29 | 2E-17 | 9E-16 | 0.2707692 | n.a. |
| ENSECAG00000004031 | 0.667 | 9.5374 | 72.142 | 2E-17 | 9E-16 | 0.5476923 | n.a. |
| SLC11A1 | 0.68 | 9.3411 | 107.44 | 4E-17 | 2E-15 | 0.3169231 | surface |
| SLC9A3R1 | 0.651 | 9.5551 | 69.803 | 7E-17 | 3E-15 | 0.7046154 | n.a. |
| CMTM3 | 0.634 | 9.4968 | 69.437 | 8E-17 | 3E-15 | 0.4553846 | n.a. |
| PGLS | 0.619 | 9.6893 | 67.891 | 2E-16 | 7E-15 | 0.6215385 | n.a. |
| PSTPIP1 | 0.617 | 9.5292 | 67.404 | 2E-16 | 9E-15 | 0.48 | n.a. |

| | | | | | | | |
|---|---|---|---|---|---|---|---|
| MPG | 0.599 | 9.4375 | 66.613 | 3E-16 | 1E-14 | 0.5230769 | n.a. |
| TRAPPC1 | 0.632 | 9.5261 | 66.111 | 4E-16 | 2E-14 | 0.5907692 | n.a. |
| RF00152 | 0.669 | 9.3407 | 78.115 | 5E-16 | 2E-14 | 0.3784615 | n.a. |
| ITGA4 | 0.604 | 9.465 | 65.631 | 6E-16 | 2E-14 | 0.4369231 | surface |
| MYO1F | 0.66 | 9.3593 | 113.62 | 7E-16 | 3E-14 | 0.3292308 | n.a. |
| ARFRP1 | 0.599 | 9.3836 | 63.934 | 1E-15 | 5E-14 | 0.3876923 | n.a. |
| RGS19 | 0.603 | 9.5283 | 63.765 | 1E-15 | 5E-14 | 0.4861538 | n.a. |
| LCP1 | 0.594 | 9.7068 | 63.003 | 2E-15 | 8E-14 | 0.6307692 | n.a. |
| STK10 | 0.628 | 9.392 | 72.11 | 4E-15 | 1E-13 | 0.3538462 | n.a. |
| ATP6V1D | 0.59 | 9.3519 | 83.967 | 5E-15 | 2E-13 | 0.32 | n.a. |
| RF00592 | 0.654 | 9.3558 | 67.908 | 1E-14 | 4E-13 | 0.4276923 | n.a. |
| PDLIM2 | 0.608 | 9.5617 | 56.708 | 5E-14 | 2E-12 | 0.7292308 | n.a. |
| KDELR1 | 0.602 | 9.4158 | 64.142 | 6E-14 | 2E-12 | 0.4030769 | n.a. |
| ICOS | 0.712 | 9.2888 | 142.62 | 6E-14 | 2E-12 | 0.2615385 | surface |
| LEPROTL1 | 0.6 | 9.565 | 56.221 | 7E-14 | 2E-12 | 0.7261538 | n.a. |
| SAMD9L | 0.608 | 9.4216 | 59.522 | 1E-13 | 3E-12 | 0.3476923 | n.a. |
| FYN | 0.602 | 9.4235 | 65.64 | 2E-13 | 7E-12 | 0.4769231 | n.a. |
| SIT1 | 0.626 | 9.3941 | 53.74 | 2E-13 | 7E-12 | 0.5138462 | surface |
| P2RY10 | 0.589 | 9.4226 | 52.749 | 4E-13 | 1E-11 | 0.4184615 | surface |
| ORAI3 | 0.61 | 9.3598 | 67.157 | 2E-11 | 6E-10 | 0.3046154 | n.a. |

| genes | logFC | logCPM | F | PValue | FDR | percent.exp | Surfacome.Label |
|---|---|---|---|---|---|---|---|
| S100A4 | 3.167 | 11.292 | 16596 | 0 | 0 | 0.9886594 | n.a. |
| LGALS1 | 2.1 | 11.109 | 5955.3 | 0 | 0 | 0.8622924 | n.a. |
| ITGB1 | 1.918 | 9.5309 | 3880.6 | 0 | 0 | 0.5556906 | surface |
| S100A11 | 1.69 | 9.847 | 3797 | 0 | 0 | 0.7379506 | n.a. |
| ENSECAG00000029287 | 1.454 | 10.043 | 2276.8 | 0 | 0 | 0.454435 | n.a. |
| ENSECAG00000031569 | 1.164 | 11.194 | 2809 | 0 | 0 | 0.9797489 | n.a. |
| S100A6 | 1.078 | 10.437 | 1319.8 | 1E-283 | 1E-280 | 0.7808829 | n.a. |
| VIM | 0.857 | 11.286 | 1240.5 | 5E-267 | 4E-264 | 0.9056298 | n.a. |
| CD74 | 0.641 | 12.628 | 1180.7 | 2E-254 | 1E-251 | 0.9696233 | surface |
| EMP3 | 0.963 | 10.126 | 1179.6 | 3E-254 | 2E-251 | 0.8525719 | surface |
| AHNAK | 1.233 | 9.5205 | 1588.7 | 1E-243 | 5E-241 | 0.2847307 | n.a. |
| S100A10 | 0.656 | 10.866 | 1012.5 | 5E-219 | 2E-216 | 0.9704334 | n.a. |
| C4orf48 | 0.998 | 9.5904 | 999.13 | 3E-216 | 1E-213 | 0.6573512 | n.a. |
| GLIPR1 | 0.97 | 9.887 | 983.55 | 6E-213 | 3E-210 | 0.690968 | surface |
| S100A5 | 0.914 | 9.5834 | 995.45 | 1E-207 | 5E-205 | 0.3005265 | n.a. |
| ANXA2 | 0.842 | 9.8875 | 927.26 | 5E-201 | 2E-198 | 0.6119887 | n.a. |
| TSPO | 1.128 | 9.7323 | 1251.6 | 9E-179 | 3E-176 | 0.2847307 | n.a. |
| INPP4B | 0.944 | 9.4661 | 892.06 | 2E-171 | 5E-169 | 0.4224382 | n.a. |
| FABP5 | 0.7 | 9.5264 | 733.17 | 9E-160 | 3E-157 | 0.3256379 | n.a. |
| LTB | 0.628 | 10.681 | 731.69 | 2E-159 | 5E-157 | 0.9384366 | n.a. |
| CD2 | 0.735 | 9.8338 | 687.88 | 4E-150 | 1E-147 | 0.7403807 | surface |
| GLIPR2 | 0.835 | 9.47 | 1043.4 | 3E-148 | 7E-146 | 0.3232078 | n.a. |
| SRGN | 0.735 | 10.485 | 658.96 | 6E-144 | 1E-141 | 0.7748076 | n.a. |
| LGALS3 | 0.893 | 9.5239 | 941.27 | 3E-139 | 6E-137 | 0.2766302 | n.a. |
| NCR3 | 0.796 | 9.482 | 606.88 | 8E-133 | 2E-130 | 0.4467396 | surface |
| IFI27 | 0.768 | 9.7314 | 587.81 | 9E-129 | 2E-126 | 0.3507493 | n.a. |
| CD40LG | 0.872 | 9.3506 | 1097.1 | 1E-125 | 2E-123 | 0.2831106 | surface |
| KLF6 | 0.874 | 9.5982 | 557.37 | 3E-122 | 5E-120 | 0.4390441 | n.a. |
| DRA | 0.602 | 11.742 | 553.22 | 2E-121 | 4E-119 | 0.8521669 | n.a. |
| JPT1 | 0.75 | 9.7243 | 551.6 | 5E-121 | 9E-119 | 0.600648 | n.a. |
| DRB | 0.623 | 11.205 | 550.03 | 1E-120 | 2E-118 | 0.8043742 | n.a. |
| SUB1 | 0.585 | 10.333 | 536.94 | 7E-118 | 1E-115 | 0.8776833 | n.a. |
| CYBA | 0.673 | 10.509 | 517.78 | 9E-114 | 1E-111 | 0.6771972 | n.a. |
| IGFLR1 | 0.685 | 9.4383 | 578.5 | 1E-108 | 2E-106 | 0.3924666 | surface |
| Eqca-DQB1 | 0.605 | 10.925 | 489.77 | 9E-108 | 1E-105 | 0.8100446 | n.a. |
| PLP2 | 0.652 | 9.9939 | 468.33 | 4E-103 | 5E-101 | 0.7509113 | n.a. |
| DQA | 0.63 | 10.712 | 393.88 | 4E-87 | 5E-85 | 0.6010531 | n.a. |
| ENSECAG00000040634 | 0.619 | 9.9556 | 380.01 | 4E-84 | 4E-82 | 0.7197246 | n.a. |
| ANXA1 | 0.638 | 9.8295 | 354.77 | 1E-78 | 1E-76 | 0.5605508 | n.a. |
| IL7R | 0.688 | 9.3731 | 912.66 | 8E-77 | 8E-75 | 0.5573107 | surface |
| CAST | 0.649 | 9.5123 | 338.86 | 3E-75 | 3E-73 | 0.345484 | n.a. |
| CISH | 0.671 | 9.3863 | 481.46 | 5E-73 | 5E-71 | 0.2563791 | n.a. |
| CMTM6 | 0.621 | 9.4988 | 283.33 | 6E-63 | 5E-61 | 0.344674 | n.a. |
| TEX30 | 0.619 | 9.4443 | 341.96 | 7E-60 | 6E-58 | 0.254354 | n.a. |
| ADGRE5 | 0.611 | 9.5791 | 241.33 | 3E-54 | 2E-52 | 0.2891859 | n.a. |

| genes | logFC | logCPM | F | PValue | FDR | percent.exp | Surfacome.Label |
|---|---|---|---|---|---|---|---|
| CCL5 | 5.052 | 11.908 | 7226.2 | 0 | 0 | 0.7048458 | n.a. |
| ENSECAG00000026891 | 4.414 | 9.2736 | 11666 | 0 | 0 | 0.8237885 | n.a. |
| GZMA | 4.337 | 9.4519 | 5068.2 | 0 | 0 | 0.4757709 | n.a. |
| UBE2C | 3.736 | 9.2563 | 7573.5 | 0 | 0 | 0.6299559 | n.a. |
| KLRK1 | 3.116 | 9.569 | 2806 | 0 | 0 | 0.5022026 | surface |
| HMGB2 | 3.115 | 10.129 | 2229.5 | 0 | 0 | 0.9911894 | n.a. |
| ENSECAG00000037332 | 2.754 | 10.054 | 1629.8 | 0 | 0 | 1 | n.a. |
| ENSECAG00000029287 | 2.74 | 10.043 | 1467.6 | 0 | 0 | 0.6740088 | n.a. |
| H2AFZ | 3.202 | 9.5191 | 1426.4 | 6E-306 | 3E-303 | 0.9251101 | n.a. |
| CGA | 2.875 | 9.3028 | 3148.9 | 2E-253 | 7E-251 | 0.5330396 | n.a. |
| H2AFX | 3.165 | 9.3048 | 1356 | 2E-237 | 5E-235 | 0.6696035 | n.a. |
| CTSW | 2.272 | 9.899 | 989.73 | 3E-214 | 7E-212 | 0.5374449 | n.a. |
| ACTB | 1.3 | 12.301 | 989.48 | 4E-214 | 8E-212 | 1 | n.a. |
| CDCA3 | 3.167 | 9.2462 | 12299 | 5E-210 | 1E-207 | 0.6211454 | n.a. |
| PCNA | 3.019 | 9.3865 | 965.65 | 4E-209 | 8E-207 | 0.7753304 | n.a. |
| RRM2 | 3.136 | 9.252 | 5890.8 | 7E-203 | 1E-200 | 0.6079295 | n.a. |
| TYMS | 2.868 | 9.3134 | 848.91 | 2E-184 | 3E-182 | 0.722467 | n.a. |
| HMGB1 | 2.015 | 10.086 | 846.55 | 7E-184 | 1E-181 | 0.9823789 | n.a. |
| TUBA1A | 2.217 | 10.091 | 831.46 | 1E-180 | 2E-178 | 0.9427313 | n.a. |
| ENSECAG00000035897 | 2.485 | 9.3428 | 761.99 | 6E-166 | 9E-164 | 0.9118943 | n.a. |
| STMN1 | 2.305 | 9.7683 | 757.82 | 5E-165 | 7E-163 | 0.9779736 | n.a. |
| CKS2 | 2.447 | 9.2847 | 1049.1 | 2E-163 | 2E-161 | 0.6431718 | n.a. |
| ENSECAG00000036105 | 2.899 | 9.2441 | 11885 | 2E-158 | 2E-156 | 0.5506608 | n.a. |
| H2AFV | 2.067 | 9.7995 | 718.21 | 1E-156 | 2E-154 | 0.9911894 | n.a. |
| TCF19 | 2.857 | 9.2869 | 1147 | 7E-153 | 8E-151 | 0.7444934 | n.a. |
| LMNB1 | 2.554 | 9.2729 | 1446.5 | 4E-149 | 4E-147 | 0.7268722 | n.a. |
| FABP5 | 2.091 | 9.5264 | 678.47 | 4E-148 | 4E-146 | 0.9295154 | n.a. |
| ANXA2 | 1.743 | 9.8875 | 673.81 | 4E-147 | 4E-145 | 0.9823789 | n.a. |
| TOP2A | 2.79 | 9.245 | 6727.8 | 6E-146 | 6E-144 | 0.4449339 | n.a. |
| BIRC5 | 2.82 | 9.243 | 11748 | 3E-143 | 3E-141 | 0.5462555 | n.a. |
| NUSAP1 | 2.653 | 9.2433 | 8795.3 | 2E-142 | 2E-140 | 0.3656388 | n.a. |
| S100A11 | 1.706 | 9.847 | 636.48 | 4E-139 | 3E-137 | 0.9251101 | n.a. |
| S100A4 | 1.608 | 11.292 | 620.93 | 7E-136 | 7E-134 | 0.876652 | n.a. |
| PFN1 | 0.951 | 11.941 | 599.08 | 3E-131 | 3E-129 | 1 | n.a. |
| CCNA2 | 2.627 | 9.2425 | 9213.1 | 3E-126 | 3E-124 | 0.4449339 | n.a. |
| TPX2 | 2.57 | 9.2428 | 7293.9 | 1E-119 | 8E-118 | 0.5418502 | n.a. |
| LGALS1 | 1.405 | 11.109 | 540 | 2E-118 | 1E-116 | 0.9207048 | n.a. |
| ASF1B | 2.481 | 9.241 | 9911.6 | 3E-118 | 2E-116 | 0.5066079 | n.a. |
| CENPA | 2.391 | 9.2413 | 7529.3 | 9E-116 | 7E-114 | 0.3964758 | n.a. |
| CENPF | 2.187 | 9.2409 | 6529.6 | 9E-115 | 7E-113 | 0.3480176 | n.a. |
| TUBB4A | 2.125 | 9.3253 | 521.61 | 1E-114 | 1E-112 | 0.5726872 | n.a. |
| ENSECAG00000031322 | 1.872 | 10.499 | 493.03 | 2E-108 | 1E-106 | 0.7577093 | n.a. |
| C1H15orf48 | 1.174 | 9.6929 | 557.26 | 4E-103 | 3E-101 | 0.2643172 | n.a. |
| PRF1 | 1.912 | 9.4454 | 781.38 | 5E-103 | 3E-101 | 0.4801762 | n.a. |
| eca-mir-8997 | 2.387 | 9.2408 | 9632.3 | 2E-99 | 1E-97 | 0.5066079 | n.a. |
| CENPM | 1.757 | 9.247 | 3127.1 | 1E-97 | 6E-96 | 0.4096916 | n.a. |
| CDC20 | 2.224 | 9.2401 | 9988.1 | 8E-96 | 5E-94 | 0.3215859 | n.a. |

| | | | | | | | |
|---|---|---|---|---|---|---|---|
| ENSECAG00000009190 | 0.583 | 14.441 | 425.44 | 6E-94 | 4E-92 | 1 | n.a. |
| CDKN2C | 2.007 | 9.2805 | 761.37 | 9E-93 | 6E-91 | 0.5330396 | n.a. |
| CCNB2 | 1.882 | 9.2382 | 9403.5 | 2E-92 | 2E-90 | 0.3348018 | n.a. |
| SMC4 | 2.045 | 9.3283 | 409.4 | 2E-90 | 1E-88 | 0.7180617 | n.a. |
| UCP2 | 1.424 | 9.9052 | 389.62 | 3E-86 | 2E-84 | 0.9647577 | n.a. |
| ENSECAG00000022442 | 1.611 | 9.7128 | 381.94 | 1E-84 | 8E-83 | 0.9471366 | n.a. |
| SPC25 | 2.142 | 9.2421 | 3897.7 | 2E-83 | 9E-82 | 0.4669604 | n.a. |
| NKG7 | 1.842 | 9.3783 | 545.63 | 2E-81 | 1E-79 | 0.5374449 | n.a. |
| ENSECAG00000033690 | 2.342 | 9.2632 | 843.84 | 5E-81 | 3E-79 | 0.5550661 | n.a. |
| CDKN3 | 1.931 | 9.2388 | 8112.5 | 1E-80 | 7E-79 | 0.3876652 | n.a. |
| ENSECAG00000039428 | 0.889 | 11.047 | 359.8 | 8E-80 | 4E-78 | 0.9911894 | n.a. |
| VIM | 1.085 | 11.286 | 359.21 | 1E-79 | 6E-78 | 0.9823789 | n.a. |
| KIFC1 | 2.163 | 9.2409 | 4600.4 | 5E-76 | 3E-74 | 0.4449339 | n.a. |
| CDKN2A | 1.83 | 9.249 | 1902 | 4E-75 | 2E-73 | 0.3920705 | n.a. |
| UBE2S | 1.808 | 9.3494 | 331.31 | 1E-73 | 6E-72 | 0.6123348 | n.a. |
| CENPE | 1.789 | 9.2392 | 5870.4 | 1E-73 | 6E-72 | 0.3656388 | n.a. |
| DEK | 1.667 | 9.6176 | 331.18 | 1E-73 | 6E-72 | 0.938326 | n.a. |
| DHDDS | 1.645 | 9.3007 | 342.47 | 2E-73 | 8E-72 | 0.5374449 | n.a. |
| CCNB1 | 2.141 | 9.2437 | 2417.6 | 3E-70 | 2E-68 | 0.3964758 | n.a. |
| MCM5 | 2.174 | 9.2776 | 693.36 | 5E-70 | 2E-68 | 0.5903084 | n.a. |
| NRM | 1.711 | 9.2939 | 461.96 | 7E-70 | 3E-68 | 0.5506608 | n.a. |
| CD8A | 1.516 | 9.4281 | 361.77 | 1E-68 | 5E-67 | 0.2819383 | surface |
| MNDA | 1.336 | 10.258 | 307.95 | 1E-68 | 6E-67 | 0.9295154 | n.a. |
| PRDX2 | 1.147 | 9.2964 | 306.72 | 2E-68 | 1E-66 | 0.2599119 | n.a. |
| FBXO5 | 1.937 | 9.2403 | 4372.4 | 6E-67 | 3E-65 | 0.3964758 | n.a. |
| LSP1 | 1.128 | 10.243 | 299.19 | 9E-67 | 4E-65 | 0.9911894 | n.a. |
| DEPDC1B | 1.858 | 9.2381 | 6007.9 | 4E-65 | 2E-63 | 0.3524229 | n.a. |
| MCM7 | 1.743 | 9.3416 | 291.84 | 4E-65 | 2E-63 | 0.6431718 | n.a. |
| CDCA8 | 1.995 | 9.2406 | 3583.7 | 4E-65 | 2E-63 | 0.3524229 | n.a. |
| REEP4 | 1.941 | 9.2747 | 669.55 | 1E-64 | 5E-63 | 0.5418502 | n.a. |
| FEN1 | 2.032 | 9.2609 | 769.51 | 1E-64 | 6E-63 | 0.5374449 | n.a. |
| S100A6 | 1.282 | 10.437 | 288.46 | 2E-64 | 9E-63 | 0.907489 | n.a. |
| ENSECAG00000028889 | 1.155 | 9.2871 | 997.88 | 7E-64 | 3E-62 | 0.2511013 | n.a. |
| MCM6 | 1.861 | 9.313 | 291.9 | 2E-63 | 7E-62 | 0.7577093 | n.a. |
| CDK1 | 1.925 | 9.2397 | 4237 | 1E-62 | 5E-61 | 0.3876652 | n.a. |
| TACC3 | 1.859 | 9.2769 | 537.87 | 2E-62 | 1E-60 | 0.5330396 | n.a. |
| UHRF1 | 1.647 | 9.2365 | 9226.5 | 5E-62 | 2E-60 | 0.3171806 | n.a. |
| MXD3 | 1.714 | 9.2429 | 2193.9 | 6E-62 | 2E-60 | 0.3039648 | n.a. |
| KIF11 | 1.863 | 9.2384 | 6566.1 | 2E-61 | 1E-59 | 0.3832599 | n.a. |
| ARPC2 | 0.935 | 10.591 | 271.18 | 1E-60 | 5E-59 | 1 | n.a. |
| PBK | 1.733 | 9.2383 | 4295.3 | 2E-59 | 7E-58 | 0.3524229 | n.a. |
| GAPDH | 0.912 | 10.705 | 264.87 | 2E-59 | 1E-57 | 0.9955947 | n.a. |
| HNRNPA2B1 | 1.188 | 10.068 | 259.06 | 4E-58 | 2E-56 | 0.9823789 | n.a. |
| KIF15 | 1.548 | 9.2382 | 5036.8 | 4E-57 | 2E-55 | 0.2819383 | n.a. |
| CST7 | 1.294 | 9.312 | 676.16 | 5E-57 | 2E-55 | 0.3744493 | n.a. |
| CENPU | 1.518 | 9.2762 | 568.45 | 3E-56 | 1E-54 | 0.4625551 | n.a. |
| SIVA1 | 1.47 | 9.5677 | 245.2 | 4E-55 | 2E-53 | 0.7929515 | n.a. |
| ZNF683 | 1.204 | 9.478 | 323.3 | 9E-55 | 4E-53 | 0.2951542 | n.a. |

| | | | | | | | |
|---|---|---|---|---|---|---|---|
| CLSPN | 1.87 | 9.2395 | 3847.2 | 1E-54 | 4E-53 | 0.4405286 | n.a. |
| NCAPG | 1.822 | 9.243 | 1983.2 | 5E-54 | 2E-52 | 0.4273128 | n.a. |
| CYTB | 0.784 | 11.109 | 238.66 | 1E-53 | 4E-52 | 0.9955947 | n.a. |
| CDC45 | 1.564 | 9.2362 | 6409.7 | 4E-53 | 2E-51 | 0.3259912 | n.a. |
| USP1 | 1.535 | 9.3606 | 243.57 | 5E-53 | 2E-51 | 0.6828194 | n.a. |
| NCAPG2 | 1.766 | 9.2424 | 2318.9 | 6E-53 | 2E-51 | 0.4096916 | n.a. |
| MCM4 | 1.812 | 9.262 | 597.08 | 2E-52 | 6E-51 | 0.5682819 | n.a. |
| MCM3 | 1.977 | 9.2749 | 478.39 | 2E-52 | 6E-51 | 0.6123348 | n.a. |
| ACTG1 | 0.999 | 10.745 | 229.88 | 9E-52 | 3E-50 | 0.9911894 | n.a. |
| DDX39A | 1.435 | 9.3461 | 229 | 2E-51 | 6E-50 | 0.7356828 | n.a. |
| GLUL | 1.112 | 9.3271 | 556.96 | 3E-51 | 1E-49 | 0.277533 | n.a. |
| NUCKS1 | 1.338 | 9.7354 | 225.84 | 7E-51 | 3E-49 | 0.9515419 | n.a. |
| ATAD2 | 1.689 | 9.2578 | 714.96 | 2E-50 | 6E-49 | 0.4757709 | n.a. |
| SGO1 | 1.621 | 9.2375 | 5615 | 2E-50 | 6E-49 | 0.2731278 | n.a. |
| ENSECAG00000033471 | 1.721 | 9.244 | 1442 | 9E-50 | 3E-48 | 0.5286344 | n.a. |
| KNL1 | 1.373 | 9.2399 | 2454.6 | 1E-49 | 5E-48 | 0.30837 | n.a. |
| CENPW | 1.749 | 9.242 | 2319.3 | 8E-49 | 3E-47 | 0.4008811 | n.a. |
| ENSECAG00000032959 | 1.354 | 9.6944 | 215.96 | 1E-48 | 3E-47 | 0.6167401 | n.a. |
| KIF22 | 1.748 | 9.2545 | 730.44 | 1E-48 | 4E-47 | 0.4052863 | n.a. |
| GALK1 | 1.107 | 9.2866 | 387.53 | 4E-47 | 1E-45 | 0.4845815 | n.a. |
| MYBL2 | 1.794 | 9.2383 | 7851.4 | 2E-46 | 6E-45 | 0.4361233 | n.a. |
| MTFR2 | 1.537 | 9.2373 | 5018.5 | 1E-45 | 4E-44 | 0.3524229 | n.a. |
| ID2 | 1.194 | 9.6451 | 200.25 | 2E-45 | 8E-44 | 0.5374449 | n.a. |
| CFL1 | 0.671 | 11.079 | 198.99 | 5E-45 | 2E-43 | 1 | n.a. |
| S100A5 | 1.061 | 9.5834 | 197.43 | 1E-44 | 3E-43 | 0.5462555 | n.a. |
| ENSECAG00000026827 | 1.51 | 9.2436 | 1585.2 | 1E-44 | 4E-43 | 0.3876652 | n.a. |
| ENSECAG00000037706 | 0.872 | 10.368 | 187.42 | 2E-42 | 5E-41 | 0.9955947 | n.a. |
| MNS1 | 1.312 | 9.2425 | 1902.5 | 2E-42 | 7E-41 | 0.277533 | n.a. |
| GMNN | 1.737 | 9.2659 | 411.72 | 5E-42 | 1E-40 | 0.5594714 | n.a. |
| ENSECAG00000020532 | 1.126 | 9.7393 | 182.83 | 1E-41 | 5E-40 | 0.9779736 | n.a. |
| LIG1 | 1.556 | 9.2648 | 417.53 | 9E-41 | 3E-39 | 0.4845815 | n.a. |
| HMMR | 1.25 | 9.2372 | 4205.5 | 1E-40 | 4E-39 | 0.2511013 | n.a. |
| HIST1H1D | 1.478 | 9.281 | 357.89 | 1E-40 | 4E-39 | 0.4185022 | n.a. |
| ENSECAG00000022254 | 0.879 | 9.3218 | 284.54 | 1E-40 | 4E-39 | 0.4273128 | n.a. |
| NUF2 | 1.467 | 9.2372 | 4415.4 | 3E-40 | 9E-39 | 0.2995595 | n.a. |
| PKMYT1 | 1.479 | 9.2511 | 752.85 | 3E-40 | 1E-38 | 0.3920705 | n.a. |
| MRPS16 | 1.403 | 9.3709 | 176.43 | 4E-40 | 1E-38 | 0.7400881 | n.a. |
| E2F2 | 1.206 | 9.2429 | 1744.1 | 4E-40 | 1E-38 | 0.2951542 | n.a. |
| COX8A | 0.915 | 10.168 | 175.41 | 6E-40 | 2E-38 | 1 | n.a. |
| CCNE2 | 1.372 | 9.2372 | 3903.3 | 7E-40 | 2E-38 | 0.2731278 | n.a. |
| SMC1A | 1.394 | 9.3103 | 228.14 | 7E-40 | 2E-38 | 0.5462555 | n.a. |
| NCAPH | 1.321 | 9.2367 | 4997.8 | 1E-39 | 3E-38 | 0.2731278 | n.a. |
| DPP4 | 0.902 | 9.2884 | 296.02 | 3E-39 | 8E-38 | 0.3656388 | surface |
| CDC6 | 1.355 | 9.2363 | 5948.1 | 4E-39 | 1E-37 | 0.2643172 | n.a. |
| SHCBP1 | 1.451 | 9.2367 | 5589.5 | 5E-39 | 1E-37 | 0.2863436 | n.a. |
| BATF | 1.287 | 9.3505 | 240.13 | 5E-39 | 1E-37 | 0.5374449 | n.a. |
| CDK2AP2 | 1.139 | 9.6423 | 170.83 | 6E-39 | 2E-37 | 0.8414097 | n.a. |
| EFHD2 | 0.945 | 9.3994 | 183.86 | 1E-38 | 3E-37 | 0.4889868 | n.a. |

| | | | | | | | |
|---|---|---|---|---|---|---|---|
| MIS18A | 1.438 | 9.2371 | 4492.3 | 2E-38 | 5E-37 | 0.3215859 | n.a. |
| EZR | 1.116 | 9.8342 | 168 | 2E-38 | 7E-37 | 0.9559471 | n.a. |
| HMGB3 | 1.631 | 9.2519 | 716.73 | 3E-38 | 9E-37 | 0.4140969 | n.a. |
| ENSECAG00000015275 | 1.13 | 9.3043 | 332.07 | 6E-38 | 2E-36 | 0.4140969 | n.a. |
| MSC | 1.139 | 9.2493 | 862.61 | 2E-37 | 5E-36 | 0.2511013 | n.a. |
| BARD1 | 1.516 | 9.2398 | 2492.5 | 2E-37 | 5E-36 | 0.3436123 | n.a. |
| RPA2 | 1.406 | 9.3129 | 198.88 | 3E-37 | 7E-36 | 0.6079295 | n.a. |
| TPI1 | 1.166 | 9.5703 | 159.89 | 1E-36 | 4E-35 | 0.8986784 | n.a. |
| ARPC3 | 0.9 | 10.061 | 159.65 | 2E-36 | 4E-35 | 0.9955947 | n.a. |
| ENSECAG00000015297 | 1.636 | 9.2596 | 481.69 | 4E-36 | 1E-34 | 0.4669604 | n.a. |
| GZMM | 0.934 | 9.4961 | 230.76 | 2E-35 | 5E-34 | 0.3920705 | n.a. |
| TYMP | 0.99 | 9.3627 | 188.69 | 3E-35 | 7E-34 | 0.4801762 | n.a. |
| YWHAQ | 1.16 | 9.7075 | 153.61 | 3E-35 | 9E-34 | 0.938326 | n.a. |
| LSM3 | 1.213 | 9.5393 | 152.67 | 5E-35 | 1E-33 | 0.907489 | n.a. |
| RAD21 | 1.234 | 9.3865 | 152.55 | 6E-35 | 1E-33 | 0.6387665 | n.a. |
| RRM1 | 1.487 | 9.2608 | 467.02 | 8E-35 | 2E-33 | 0.4273128 | n.a. |
| UBE2T | 1.407 | 9.2403 | 1547.1 | 1E-34 | 3E-33 | 0.2995595 | n.a. |
| PEBP1 | 1.084 | 9.6042 | 148.79 | 4E-34 | 9E-33 | 0.8810573 | n.a. |
| MYL6 | 0.682 | 10.731 | 147.92 | 6E-34 | 1E-32 | 1 | n.a. |
| NCAPD2 | 1.294 | 9.2437 | 1150.5 | 6E-34 | 2E-32 | 0.3303965 | n.a. |
| DNMT1 | 1.597 | 9.2975 | 225.74 | 7E-34 | 2E-32 | 0.6079295 | n.a. |
| SNRPG | 1.15 | 9.559 | 146.26 | 1E-33 | 3E-32 | 0.9339207 | n.a. |
| CAP1 | 0.958 | 9.9501 | 145.79 | 2E-33 | 4E-32 | 0.9735683 | n.a. |
| CLIC1 | 0.896 | 10.07 | 145.36 | 2E-33 | 5E-32 | 0.9647577 | n.a. |
| POC1A | 1.326 | 9.2936 | 236.25 | 3E-33 | 6E-32 | 0.5991189 | n.a. |
| ARHGEF39 | 1.392 | 9.2399 | 1532.7 | 3E-33 | 8E-32 | 0.2731278 | n.a. |
| RFC2 | 1.25 | 9.2986 | 210.14 | 5E-33 | 1E-31 | 0.5506608 | n.a. |
| ZWINT | 1.691 | 9.2392 | 1839.7 | 5E-33 | 1E-31 | 0.4361233 | n.a. |
| DBI | 0.973 | 9.673 | 143.06 | 7E-33 | 2E-31 | 0.8986784 | n.a. |
| NANS | 1.125 | 9.4296 | 142.72 | 8E-33 | 2E-31 | 0.7753304 | n.a. |
| CDCA7 | 1.621 | 9.2634 | 154.59 | 2E-32 | 5E-31 | 0.6387665 | n.a. |
| SMC3 | 1.25 | 9.4311 | 139.93 | 3E-32 | 7E-31 | 0.7621145 | n.a. |
| ENSECAG00000012818 | 1.103 | 9.2495 | 492.82 | 3E-32 | 8E-31 | 0.2643172 | n.a. |
| GLIPR2 | 0.788 | 9.47 | 138.94 | 5E-32 | 1E-30 | 0.6651982 | n.a. |
| CSRP1 | 0.946 | 9.2812 | 369.72 | 1E-31 | 3E-30 | 0.2643172 | n.a. |
| CEP55 | 1.245 | 9.2368 | 2300 | 1E-31 | 3E-30 | 0.2687225 | n.a. |
| ENSECAG00000016018 | 0.735 | 9.3138 | 278.76 | 3E-31 | 7E-30 | 0.277533 | n.a. |
| ENSECAG00000031569 | 0.69 | 11.194 | 133.5 | 8E-31 | 2E-29 | 0.9911894 | n.a. |
| GINS1 | 1.345 | 9.2464 | 837.48 | 1E-30 | 3E-29 | 0.2907489 | n.a. |
| RFC3 | 1.609 | 9.2557 | 483.91 | 2E-30 | 4E-29 | 0.4801762 | n.a. |
| SSNA1 | 1.166 | 9.3399 | 133.91 | 3E-30 | 6E-29 | 0.6299559 | n.a. |
| UNG | 1.428 | 9.26 | 322.05 | 4E-30 | 8E-29 | 0.3480176 | n.a. |
| CORT | 1.628 | 9.2439 | 767.94 | 4E-30 | 8E-29 | 0.5242291 | n.a. |
| FLNA | 1.023 | 9.4126 | 129.39 | 6E-30 | 1E-28 | 0.5418502 | n.a. |
| PRC1 | 1.27 | 9.24 | 1519.8 | 8E-30 | 2E-28 | 0.2951542 | n.a. |
| CDC25B | 1.272 | 9.3048 | 131.42 | 1E-29 | 2E-28 | 0.5110132 | n.a. |
| COTL1 | 0.747 | 10.529 | 128.39 | 1E-29 | 2E-28 | 0.9779736 | n.a. |
| GPS2 | 1.095 | 9.4995 | 125.64 | 4E-29 | 9E-28 | 0.7577093 | n.a. |

| | | | | | | | |
|---|---|---|---|---|---|---|---|
| GINS2 | 1.584 | 9.239 | 2258.2 | 6E-29 | 1E-27 | 0.3612335 | n.a. |
| LCP1 | 0.989 | 9.7068 | 124.26 | 8E-29 | 2E-27 | 0.9118943 | n.a. |
| CHAF1A | 1.154 | 9.2395 | 1481 | 1E-28 | 2E-27 | 0.277533 | n.a. |
| SH3BGRL3 | 0.621 | 10.736 | 121.09 | 4E-28 | 9E-27 | 0.9911894 | n.a. |
| CST3 | 0.769 | 11.921 | 120.99 | 4E-28 | 9E-27 | 0.7885463 | n.a. |
| AURKAIP1 | 1.101 | 9.4351 | 120.24 | 6E-28 | 1E-26 | 0.845815 | n.a. |
| CIAO2A | 0.88 | 9.316 | 154.3 | 8E-28 | 2E-26 | 0.5330396 | n.a. |
| ENSECAG00000018242 | 1.167 | 9.2923 | 201.04 | 1E-27 | 2E-26 | 0.5242291 | n.a. |
| EZH2 | 1.256 | 9.245 | 942.65 | 2E-27 | 5E-26 | 0.3524229 | n.a. |
| SDF2L1 | 0.978 | 9.3874 | 116.66 | 4E-27 | 8E-26 | 0.7180617 | n.a. |
| CRELD2 | 0.678 | 9.2876 | 180.58 | 5E-27 | 1E-25 | 0.2907489 | n.a. |
| AURKA | 1.338 | 9.247 | 762.14 | 5E-27 | 1E-25 | 0.2951542 | n.a. |
| CKS1B | 1.276 | 9.345 | 115.04 | 9E-27 | 2E-25 | 0.6343612 | n.a. |
| LSM5 | 1.028 | 9.5848 | 113.4 | 2E-26 | 4E-25 | 0.8810573 | n.a. |
| ESCO2 | 1.123 | 9.2429 | 911.58 | 3E-26 | 5E-25 | 0.2863436 | n.a. |
| CENPH | 1.389 | 9.2558 | 379.79 | 5E-26 | 1E-24 | 0.3920705 | n.a. |
| ENSECAG00000036511 | 0.965 | 9.3303 | 123.35 | 6E-26 | 1E-24 | 0.5991189 | n.a. |
| FAM89B | 0.964 | 9.3314 | 145.03 | 9E-26 | 2E-24 | 0.5154185 | n.a. |
| ERCC1 | 0.979 | 9.3771 | 109.63 | 1E-25 | 2E-24 | 0.6960352 | n.a. |
| RANBP1 | 1.065 | 9.5221 | 108.07 | 3E-25 | 5E-24 | 0.8546256 | n.a. |
| CENPP | 1.326 | 9.2531 | 413.08 | 3E-25 | 7E-24 | 0.4185022 | n.a. |
| TKT | 0.744 | 9.6069 | 106.7 | 6E-25 | 1E-23 | 0.6960352 | n.a. |
| ETHE1 | 0.824 | 9.3855 | 105.81 | 9E-25 | 2E-23 | 0.5682819 | n.a. |
| ACP1 | 1.068 | 9.4501 | 105.14 | 1E-24 | 2E-23 | 0.8414097 | n.a. |
| COX6B1 | 0.78 | 9.9121 | 105.03 | 1E-24 | 2E-23 | 0.9955947 | n.a. |
| STK39 | 1.243 | 9.3069 | 197.31 | 1E-24 | 3E-23 | 0.6035242 | n.a. |
| TUBA4A | 1.083 | 9.5479 | 104.86 | 1E-24 | 3E-23 | 0.7753304 | n.a. |
| FTL | 0.661 | 11.187 | 104.51 | 2E-24 | 3E-23 | 0.9559471 | n.a. |
| PFKP | 0.881 | 9.3233 | 108.36 | 3E-24 | 6E-23 | 0.4757709 | n.a. |
| SRSF3 | 1.019 | 9.545 | 103.17 | 3E-24 | 6E-23 | 0.8810573 | n.a. |
| RNASEH2B | 0.923 | 9.3711 | 102 | 6E-24 | 1E-22 | 0.6079295 | n.a. |
| IDH2 | 0.879 | 9.6001 | 101.52 | 8E-24 | 1E-22 | 0.8722467 | n.a. |
| PLK4 | 1.206 | 9.2404 | 1004.6 | 1E-23 | 2E-22 | 0.2555066 | n.a. |
| CALM2 | 0.728 | 10.275 | 100.98 | 1E-23 | 2E-22 | 0.9295154 | n.a. |
| HELLS | 1.384 | 9.2492 | 552.51 | 2E-23 | 3E-22 | 0.339207 | n.a. |
| DAZAP1 | 1.075 | 9.3388 | 121.24 | 2E-23 | 4E-22 | 0.6696035 | n.a. |
| CENPK | 1.41 | 9.253 | 360.67 | 3E-23 | 6E-22 | 0.4185022 | n.a. |
| CAPZA1 | 0.883 | 9.7535 | 98.65 | 3E-23 | 6E-22 | 0.907489 | n.a. |
| NSD2 | 1.14 | 9.2457 | 701.64 | 3E-23 | 6E-22 | 0.30837 | n.a. |
| CRYL1 | 0.879 | 9.2872 | 164.85 | 5E-23 | 8E-22 | 0.3876652 | n.a. |
| COX5B | 0.779 | 9.881 | 96.899 | 8E-23 | 1E-21 | 0.9735683 | n.a. |
| ENSECAG00000007197 | 1.104 | 9.2815 | 216.52 | 9E-23 | 2E-21 | 0.4493392 | n.a. |
| ITGB1 | 0.737 | 9.5309 | 96.621 | 9E-23 | 2E-21 | 0.5506608 | surface |
| DCTPP1 | 0.935 | 9.3868 | 96.272 | 1E-22 | 2E-21 | 0.6828194 | n.a. |
| MTHFD2 | 1.015 | 9.2787 | 173.84 | 1E-22 | 2E-21 | 0.4405286 | n.a. |
| ENSECAG00000018918 | 1.166 | 9.237 | 2831.4 | 2E-22 | 3E-21 | 0.2863436 | n.a. |
| ACOT7 | 1.28 | 9.2472 | 517.66 | 4E-22 | 7E-21 | 0.4229075 | n.a. |
| ALYREF | 0.946 | 9.2553 | 377.15 | 5E-22 | 8E-21 | 0.2643172 | n.a. |

| | | | | | | | |
|---|---|---|---|---|---|---|---|
| RFC5 | 1.17 | 9.2754 | 222.93 | 5E-22 | 8E-21 | 0.4229075 | n.a. |
| ATP5MC3 | 0.729 | 9.9993 | 92.653 | 7E-22 | 1E-20 | 0.969163 | n.a. |
| ENSECAG00000015558 | 0.866 | 9.662 | 91.939 | 1E-21 | 2E-20 | 0.907489 | n.a. |
| ENSECAG00000009520 | 1.091 | 9.3369 | 91.621 | 1E-21 | 2E-20 | 0.6123348 | n.a. |
| TALDO1 | 0.749 | 9.9099 | 90.707 | 2E-21 | 3E-20 | 0.938326 | n.a. |
| ENSECAG00000020136 | 0.824 | 9.7014 | 90.317 | 2E-21 | 4E-20 | 0.8061674 | n.a. |
| LSM2 | 0.954 | 9.4051 | 89.691 | 3E-21 | 5E-20 | 0.7444934 | n.a. |
| SMC6 | 0.973 | 9.3447 | 89.191 | 4E-21 | 6E-20 | 0.6035242 | n.a. |
| ITGB3BP | 1.099 | 9.2893 | 162.6 | 5E-21 | 8E-20 | 0.4625551 | n.a. |
| RAD51AP1 | 1.089 | 9.2691 | 222.34 | 5E-21 | 9E-20 | 0.3215859 | n.a. |
| WDR76 | 0.952 | 9.2583 | 369.82 | 6E-21 | 9E-20 | 0.2599119 | n.a. |
| RAC1 | 0.726 | 9.9054 | 87.724 | 8E-21 | 1E-19 | 0.9471366 | n.a. |
| UBE2E3 | 0.616 | 9.3915 | 112.82 | 9E-21 | 2E-19 | 0.3480176 | n.a. |
| COX7A2 | 0.821 | 9.6535 | 87.315 | 1E-20 | 2E-19 | 0.907489 | n.a. |
| HNRNPA3 | 0.906 | 9.6289 | 87.314 | 1E-20 | 2E-19 | 0.8678414 | n.a. |
| BANF1 | 0.774 | 9.8742 | 86.598 | 1E-20 | 2E-19 | 0.9559471 | n.a. |
| H1FX | 0.92 | 9.3523 | 88.546 | 1E-20 | 2E-19 | 0.4801762 | n.a. |
| IQGAP2 | 0.852 | 9.3027 | 132.7 | 2E-20 | 2E-19 | 0.3832599 | n.a. |
| SEC11C | 0.877 | 9.493 | 86.252 | 2E-20 | 3E-19 | 0.7929515 | n.a. |
| WDR1 | 0.756 | 9.7464 | 85.524 | 2E-20 | 4E-19 | 0.9427313 | n.a. |
| GYG1 | 0.825 | 9.5017 | 85.364 | 3E-20 | 4E-19 | 0.7136564 | n.a. |
| CARHSP1 | 0.899 | 9.4061 | 84.868 | 3E-20 | 5E-19 | 0.6299559 | n.a. |
| HP1BP3 | 0.956 | 9.4292 | 84.606 | 4E-20 | 6E-19 | 0.6563877 | n.a. |
| COX17 | 0.612 | 10.369 | 84.358 | 4E-20 | 7E-19 | 0.9867841 | n.a. |
| CKAP2L | 1.183 | 9.2529 | 410.18 | 9E-20 | 1E-18 | 0.2951542 | n.a. |
| COX6A1 | 0.693 | 10.007 | 82.544 | 1E-19 | 2E-18 | 0.9779736 | n.a. |
| TXN | 0.712 | 9.9313 | 82.181 | 1E-19 | 2E-18 | 0.9427313 | surface |
| MCM2 | 1.29 | 9.2744 | 146.49 | 1E-19 | 2E-18 | 0.4757709 | n.a. |
| SNRPD1 | 0.924 | 9.5235 | 81.631 | 2E-19 | 3E-18 | 0.8898678 | n.a. |
| NASP | 1.116 | 9.2743 | 185.1 | 2E-19 | 3E-18 | 0.4229075 | n.a. |
| KNSTRN | 1.196 | 9.2554 | 253.48 | 2E-19 | 3E-18 | 0.3171806 | n.a. |
| ORC6 | 1.232 | 9.2453 | 539.9 | 3E-19 | 4E-18 | 0.3303965 | n.a. |
| CBFB | 0.929 | 9.3467 | 86.6 | 3E-19 | 5E-18 | 0.5594714 | n.a. |
| eca-mir-1892 | 0.783 | 9.4799 | 79.738 | 4E-19 | 7E-18 | 0.7356828 | n.a. |
| ATP5PD | 0.812 | 9.5841 | 79.569 | 5E-19 | 7E-18 | 0.9295154 | n.a. |
| PSMB6 | 0.841 | 9.4843 | 79.313 | 6E-19 | 8E-18 | 0.8590308 | n.a. |
| XPO1 | 1.02 | 9.29 | 172.2 | 6E-19 | 9E-18 | 0.339207 | n.a. |
| PIGY | 0.813 | 9.3782 | 78.68 | 8E-19 | 1E-17 | 0.5859031 | n.a. |
| PPP4C | 0.865 | 9.5172 | 78.297 | 9E-19 | 1E-17 | 0.8590308 | n.a. |
| SNRPA | 0.976 | 9.305 | 116.02 | 1E-18 | 2E-17 | 0.5022026 | n.a. |
| ENSECAG00000029897 | 0.925 | 9.3348 | 102.01 | 2E-18 | 2E-17 | 0.5550661 | n.a. |
| DUSP22 | 0.787 | 9.2879 | 145.5 | 2E-18 | 2E-17 | 0.2731278 | n.a. |
| NCAPD3 | 1.136 | 9.2575 | 320.82 | 2E-18 | 3E-17 | 0.2907489 | n.a. |
| PPP1CA | 0.746 | 9.7882 | 76.3 | 3E-18 | 4E-17 | 0.9647577 | n.a. |
| BAX | 0.828 | 9.3106 | 80.904 | 4E-18 | 5E-17 | 0.5198238 | n.a. |
| LSM4 | 0.937 | 9.4444 | 75.523 | 4E-18 | 5E-17 | 0.845815 | n.a. |
| NCAPH2 | 1.11 | 9.2984 | 120.46 | 5E-18 | 8E-17 | 0.5198238 | n.a. |
| CENPX | 0.89 | 9.4182 | 74.189 | 7E-18 | 1E-16 | 0.7004405 | n.a. |

| | | | | | | | |
|---|---|---|---|---|---|---|---|
| PGP | 0.759 | 9.2896 | 165.44 | 8E-18 | 1E-16 | 0.2995595 | n.a. |
| ERP29 | 0.625 | 9.3518 | 74.129 | 8E-18 | 1E-16 | 0.4757709 | n.a. |
| PSMA7 | 0.736 | 9.8217 | 73.587 | 1E-17 | 1E-16 | 0.9471366 | n.a. |
| SPC24 | 1.386 | 9.2428 | 528.27 | 1E-17 | 2E-16 | 0.3832599 | n.a. |
| ENSECAG00000019073 | 0.689 | 9.919 | 73.331 | 1E-17 | 2E-16 | 0.9735683 | n.a. |
| PSMA2 | 0.787 | 9.6232 | 72.343 | 2E-17 | 3E-16 | 0.9162996 | n.a. |
| CDCA4 | 0.93 | 9.2785 | 129.14 | 2E-17 | 3E-16 | 0.3744493 | n.a. |
| MIS18BP1 | 1.07 | 9.2452 | 446.18 | 3E-17 | 4E-16 | 0.2819383 | n.a. |
| NSMCE4A | 0.885 | 9.3632 | 71.014 | 4E-17 | 5E-16 | 0.6035242 | n.a. |
| ENSECAG00000015993 | 0.776 | 9.3388 | 71.969 | 5E-17 | 7E-16 | 0.5506608 | n.a. |
| BRCA2 | 1.229 | 9.2516 | 312.55 | 7E-17 | 9E-16 | 0.3215859 | n.a. |
| ENSECAG00000033515 | 0.729 | 9.8609 | 69.605 | 8E-17 | 1E-15 | 0.9295154 | n.a. |
| BRI3BP | 1.034 | 9.2708 | 121.86 | 8E-17 | 1E-15 | 0.4493392 | n.a. |
| LBR | 0.848 | 9.6093 | 68.868 | 1E-16 | 1E-15 | 0.8502203 | n.a. |
| RHNO1 | 1.358 | 9.2779 | 103.64 | 1E-16 | 2E-15 | 0.5374449 | n.a. |
| RBM17 | 0.822 | 9.4985 | 68.359 | 1E-16 | 2E-15 | 0.8281938 | n.a. |
| RPA1 | 1.053 | 9.2965 | 122.1 | 2E-16 | 2E-15 | 0.4625551 | n.a. |
| CIB1 | 0.799 | 9.4458 | 67.772 | 2E-16 | 3E-15 | 0.8105727 | n.a. |
| SUPT4H1 | 0.794 | 9.5879 | 67.011 | 3E-16 | 4E-15 | 0.8590308 | n.a. |
| HSBP1 | 0.679 | 9.5012 | 66.939 | 3E-16 | 4E-15 | 0.7048458 | n.a. |
| NDUFAB1 | 0.832 | 9.4852 | 66.868 | 3E-16 | 4E-15 | 0.8414097 | n.a. |
| ENSECAG00000024654 | 0.633 | 9.3419 | 80.52 | 3E-16 | 4E-15 | 0.3259912 | n.a. |
| WRAP73 | 0.977 | 9.2804 | 124.26 | 4E-16 | 5E-15 | 0.3876652 | n.a. |
| PSMB1 | 0.701 | 9.7091 | 66.244 | 4E-16 | 5E-15 | 0.9515419 | n.a. |
| ENSECAG00000038550 | 0.738 | 9.537 | 66.09 | 4E-16 | 6E-15 | 0.7973568 | n.a. |
| GNAI2 | 0.673 | 9.598 | 65.713 | 5E-16 | 7E-15 | 0.7929515 | n.a. |
| ENSECAG00000003916 | 1.045 | 9.2637 | 186.22 | 8E-16 | 1E-14 | 0.3524229 | n.a. |
| ENSECAG00000036034 | 0.675 | 9.8269 | 64.988 | 8E-16 | 1E-14 | 0.907489 | n.a. |
| ACTR2 | 0.759 | 9.4911 | 64.357 | 1E-15 | 1E-14 | 0.7312775 | n.a. |
| ENSECAG00000010480 | 0.85 | 9.3685 | 81.405 | 1E-15 | 1E-14 | 0.4845815 | n.a. |
| ATP5F1D | 0.643 | 9.6297 | 64.067 | 1E-15 | 2E-14 | 0.8325991 | n.a. |
| AP3S1 | 0.772 | 9.534 | 63.932 | 1E-15 | 2E-14 | 0.7797357 | n.a. |
| COX5A | 0.707 | 9.6605 | 63.714 | 1E-15 | 2E-14 | 0.9118943 | n.a. |
| CTBP1 | 0.806 | 9.3234 | 71.161 | 2E-15 | 2E-14 | 0.4845815 | n.a. |
| RAP1A | 0.752 | 9.5234 | 63.504 | 2E-15 | 2E-14 | 0.8414097 | n.a. |
| TERF1 | 0.857 | 9.2933 | 119 | 2E-15 | 3E-14 | 0.3612335 | n.a. |
| ATP1B3 | 0.699 | 9.5391 | 62.862 | 2E-15 | 3E-14 | 0.7400881 | surface |
| SRSF7 | 0.829 | 9.5036 | 62.821 | 2E-15 | 3E-14 | 0.7621145 | n.a. |
| DPY30 | 0.823 | 9.5067 | 62.806 | 2E-15 | 3E-14 | 0.8193833 | n.a. |
| TRA2B | 0.804 | 9.5305 | 62.564 | 3E-15 | 3E-14 | 0.8017621 | n.a. |
| ANP32E | 0.975 | 9.2631 | 160.97 | 3E-15 | 4E-14 | 0.3436123 | n.a. |
| KIF20B | 1.061 | 9.2938 | 101.11 | 3E-15 | 4E-14 | 0.4493392 | n.a. |
| NIPSNAP2 | 0.85 | 9.3016 | 96.331 | 5E-15 | 7E-14 | 0.4229075 | n.a. |
| ENSECAG00000019430 | 0.695 | 9.3323 | 84.226 | 7E-15 | 8E-14 | 0.4008811 | n.a. |
| IL2RB | 0.91 | 9.3166 | 121.97 | 1E-14 | 1E-13 | 0.3568282 | surface |
| SNF8 | 0.65 | 9.3578 | 66.957 | 1E-14 | 1E-13 | 0.5594714 | n.a. |
| ENSECAG00000016499 | 0.707 | 9.4863 | 59.181 | 1E-14 | 2E-13 | 0.7268722 | n.a. |
| PSMB2 | 0.7 | 9.5898 | 59.153 | 1E-14 | 2E-13 | 0.9207048 | n.a. |

| | | | | | | | |
|---|---|---|---|---|---|---|---|
| NMRAL1 | 0.763 | 9.3318 | 60.491 | 2E-14 | 2E-13 | 0.5947137 | n.a. |
| ENSECAG00000035110 | 0.685 | 9.6734 | 58.69 | 2E-14 | 2E-13 | 0.8810573 | n.a. |
| BLMH | 0.673 | 9.3179 | 58.502 | 2E-14 | 2E-13 | 0.5638767 | n.a. |
| NXT2 | 0.892 | 9.2747 | 132.24 | 2E-14 | 3E-13 | 0.3612335 | n.a. |
| NABP2 | 0.907 | 9.293 | 106.4 | 3E-14 | 3E-13 | 0.4713656 | n.a. |
| PCNP | 0.744 | 9.6512 | 57.909 | 3E-14 | 3E-13 | 0.9030837 | n.a. |
| SHMT1 | 0.742 | 9.3192 | 57.585 | 3E-14 | 4E-13 | 0.4757709 | n.a. |
| ARPC5L | 0.783 | 9.4373 | 57.45 | 4E-14 | 4E-13 | 0.7973568 | n.a. |
| PRDX3 | 0.849 | 9.3339 | 57.219 | 4E-14 | 5E-13 | 0.6035242 | n.a. |
| NFYB | 0.766 | 9.358 | 56.826 | 5E-14 | 6E-13 | 0.4581498 | n.a. |
| ENSECAG00000019585 | 0.607 | 9.2793 | 114.1 | 7E-14 | 8E-13 | 0.2819383 | n.a. |
| ARL6IP6 | 0.772 | 9.3181 | 70.353 | 7E-14 | 8E-13 | 0.4581498 | n.a. |
| ITM2C | 0.596 | 9.5994 | 55.826 | 8E-14 | 9E-13 | 0.5462555 | surface |
| TMEM141 | 0.811 | 9.2799 | 117.56 | 8E-14 | 1E-12 | 0.3524229 | n.a. |
| ATP5IF1 | 0.69 | 9.6248 | 55.383 | 1E-13 | 1E-12 | 0.9339207 | n.a. |
| NME1 | 0.873 | 9.4477 | 54.879 | 1E-13 | 1E-12 | 0.7929515 | n.a. |
| RBBP4 | 0.808 | 9.4357 | 54.731 | 1E-13 | 2E-12 | 0.7004405 | n.a. |
| TIPIN | 0.95 | 9.2518 | 242.7 | 1E-13 | 2E-12 | 0.2863436 | n.a. |
| DTYMK | 1.146 | 9.2491 | 315.75 | 2E-13 | 2E-12 | 0.3436123 | n.a. |
| YWHAH | 0.706 | 9.5952 | 54.216 | 2E-13 | 2E-12 | 0.814978 | n.a. |
| ACTR3 | 0.598 | 9.9778 | 54.123 | 2E-13 | 2E-12 | 0.9603524 | n.a. |
| RCC2 | 0.692 | 9.2681 | 163.12 | 2E-13 | 2E-12 | 0.2643172 | n.a. |
| STAG1 | 0.809 | 9.2936 | 109.63 | 2E-13 | 2E-12 | 0.3480176 | n.a. |
| HTATSF1 | 0.799 | 9.3418 | 62.874 | 2E-13 | 2E-12 | 0.4801762 | n.a. |
| HDGF | 0.776 | 9.2879 | 98.265 | 2E-13 | 2E-12 | 0.3480176 | n.a. |
| HNRNPU | 0.744 | 9.4008 | 53.763 | 2E-13 | 3E-12 | 0.5242291 | n.a. |
| RFC4 | 1.042 | 9.2587 | 182.59 | 3E-13 | 3E-12 | 0.3524229 | n.a. |
| CBX5 | 0.822 | 9.3148 | 53.152 | 3E-13 | 3E-12 | 0.5814978 | n.a. |
| POLD2 | 0.899 | 9.2564 | 224.73 | 3E-13 | 4E-12 | 0.277533 | n.a. |
| RPS27L | 0.601 | 9.8828 | 52.908 | 4E-13 | 4E-12 | 0.969163 | n.a. |
| CIAO2B | 0.689 | 9.4282 | 52.889 | 4E-13 | 4E-12 | 0.6387665 | n.a. |
| ENSECAG00000036180 | 0.673 | 9.4114 | 52.73 | 4E-13 | 4E-12 | 0.6211454 | n.a. |
| CHCHD5 | 0.719 | 9.4351 | 52.665 | 4E-13 | 4E-12 | 0.7444934 | n.a. |
| HNRNPDL | 0.604 | 9.9694 | 52.124 | 5E-13 | 6E-12 | 0.9735683 | n.a. |
| PSMB9 | 0.624 | 9.6927 | 51.627 | 7E-13 | 7E-12 | 0.938326 | n.a. |
| SNRPB | 0.722 | 9.5647 | 51.555 | 7E-13 | 8E-12 | 0.9207048 | n.a. |
| SRSF9 | 0.618 | 9.4229 | 51.458 | 7E-13 | 8E-12 | 0.6211454 | n.a. |
| HSD17B10 | 0.773 | 9.3945 | 51.422 | 8E-13 | 8E-12 | 0.7004405 | n.a. |
| MYO1F | 0.585 | 9.3593 | 51.392 | 8E-13 | 8E-12 | 0.4933921 | n.a. |
| NTAN1 | 0.68 | 9.412 | 50.635 | 1E-12 | 1E-11 | 0.6740088 | n.a. |
| ENSECAG00000015596 | 0.612 | 9.3128 | 60.202 | 1E-12 | 1E-11 | 0.3964758 | n.a. |
| ENSECAG00000035182 | 1.195 | 9.2638 | 118.85 | 1E-12 | 1E-11 | 0.4537445 | n.a. |
| LDHA | 0.58 | 9.6788 | 50.167 | 1E-12 | 2E-11 | 0.8502203 | n.a. |
| DBF4 | 0.789 | 9.3047 | 72.54 | 2E-12 | 2E-11 | 0.4317181 | n.a. |
| ENSECAG00000026987 | 0.591 | 9.8596 | 49.696 | 2E-12 | 2E-11 | 0.9603524 | n.a. |
| BUD23 | 0.753 | 9.4581 | 49.328 | 2E-12 | 2E-11 | 0.753304 | n.a. |
| RTRAF | 0.671 | 9.5693 | 49.079 | 3E-12 | 3E-11 | 0.8810573 | n.a. |
| TRAPPC1 | 0.687 | 9.5261 | 49.008 | 3E-12 | 3E-11 | 0.8193833 | n.a. |

| | | | | | | | |
|---|---|---|---|---|---|---|---|
| ARFRP1 | 0.742 | 9.3836 | 48.789 | 3E-12 | 3E-11 | 0.6828194 | n.a. |
| SERPINB9 | 0.689 | 9.3883 | 67.273 | 3E-12 | 3E-11 | 0.3876652 | n.a. |
| ATP5F1B | 0.608 | 9.6895 | 48.667 | 3E-12 | 3E-11 | 0.8986784 | n.a. |
| BUB3 | 0.696 | 9.3285 | 48.26 | 4E-12 | 4E-11 | 0.4889868 | n.a. |
| NONO | 0.679 | 9.4716 | 47.732 | 5E-12 | 5E-11 | 0.753304 | n.a. |
| TRAPPC2B | 0.776 | 9.3102 | 65.853 | 6E-12 | 6E-11 | 0.5110132 | n.a. |
| ZDHHC24 | 0.63 | 9.314 | 52.614 | 6E-12 | 6E-11 | 0.4493392 | n.a. |
| SPCS3 | 0.585 | 9.6267 | 47.339 | 6E-12 | 6E-11 | 0.7753304 | n.a. |
| RANBP3 | 0.638 | 9.6437 | 46.8 | 8E-12 | 8E-11 | 0.8986784 | n.a. |
| ATP5MF | 0.623 | 9.5918 | 46.755 | 8E-12 | 8E-11 | 0.9427313 | n.a. |
| SH2D1A | 0.95 | 9.4515 | 56.534 | 9E-12 | 9E-11 | 0.6431718 | n.a. |
| PSMA5 | 0.677 | 9.4172 | 46.372 | 1E-11 | 1E-10 | 0.753304 | n.a. |
| CTDSP1 | 0.679 | 9.3715 | 46.069 | 1E-11 | 1E-10 | 0.4933921 | n.a. |
| RAB5IF | 0.651 | 9.5719 | 45.972 | 1E-11 | 1E-10 | 0.8325991 | n.a. |
| ENSECAG00000032735 | 0.691 | 9.3125 | 75.94 | 1E-11 | 1E-10 | 0.3127753 | n.a. |
| ELOC | 0.655 | 9.4415 | 45.797 | 1E-11 | 1E-10 | 0.6784141 | n.a. |
| CENPC | 0.752 | 9.3187 | 68.068 | 1E-11 | 1E-10 | 0.4317181 | n.a. |
| ANAPC15 | 0.803 | 9.2777 | 103.8 | 2E-11 | 2E-10 | 0.3480176 | n.a. |
| CYCS | 0.716 | 9.3968 | 45.28 | 2E-11 | 2E-10 | 0.6872247 | n.a. |
| POMP | 0.656 | 9.5439 | 45.051 | 2E-11 | 2E-10 | 0.8678414 | n.a. |
| RTN3 | 0.684 | 9.3937 | 44.645 | 2E-11 | 2E-10 | 0.6784141 | n.a. |
| SRI | 0.648 | 9.4117 | 44.482 | 3E-11 | 3E-10 | 0.5991189 | n.a. |
| HNRNPM | 0.678 | 9.6078 | 44.297 | 3E-11 | 3E-10 | 0.8414097 | n.a. |
| FAM136A | 0.703 | 9.381 | 43.789 | 4E-11 | 4E-10 | 0.5374449 | n.a. |
| TUBB | 0.813 | 9.2558 | 242.92 | 4E-11 | 4E-10 | 0.2511013 | n.a. |
| ENSECAG00000001661 | 0.607 | 9.3692 | 43.246 | 5E-11 | 5E-10 | 0.4581498 | n.a. |
| IDI1 | 0.702 | 9.4385 | 42.358 | 8E-11 | 7E-10 | 0.6651982 | n.a. |
| ENSECAG00000022750 | 0.702 | 9.3515 | 48.938 | 8E-11 | 7E-10 | 0.5638767 | n.a. |
| HTATIP2 | 0.774 | 9.3 | 59.989 | 9E-11 | 8E-10 | 0.4229075 | n.a. |
| TRIM59 | 0.724 | 9.3315 | 52.395 | 1E-10 | 9E-10 | 0.3788546 | n.a. |
| RASSF7 | 0.701 | 9.3343 | 57.991 | 1E-10 | 1E-09 | 0.4757709 | n.a. |
| MRPL18 | 0.657 | 9.4099 | 41.514 | 1E-10 | 1E-09 | 0.7665198 | n.a. |
| KIF2A | 0.673 | 9.3544 | 42.121 | 1E-10 | 1E-09 | 0.5638767 | n.a. |
| UBXN4 | 0.599 | 9.4336 | 40.96 | 2E-10 | 1E-09 | 0.6784141 | n.a. |
| NDUFB4 | 0.59 | 9.5448 | 40.491 | 2E-10 | 2E-09 | 0.8502203 | n.a. |
| PTPA | 0.654 | 9.292 | 69.841 | 2E-10 | 2E-09 | 0.30837 | n.a. |
| SRSF1 | 0.671 | 9.4694 | 40.393 | 2E-10 | 2E-09 | 0.6696035 | n.a. |
| HNRNPH1 | 0.646 | 9.5404 | 40.372 | 2E-10 | 2E-09 | 0.7577093 | n.a. |
| TRAPPC6A | 0.651 | 9.3262 | 41.627 | 3E-10 | 2E-09 | 0.5418502 | n.a. |
| RNASEH2C | 0.852 | 9.3304 | 48.648 | 3E-10 | 3E-09 | 0.5770925 | n.a. |
| NELFCD | 0.693 | 9.2933 | 59.246 | 3E-10 | 3E-09 | 0.3788546 | n.a. |
| MRPS18C | 0.778 | 9.2984 | 63.105 | 3E-10 | 3E-09 | 0.4669604 | n.a. |
| COMMD7 | 0.583 | 9.6123 | 38.473 | 6E-10 | 5E-09 | 0.8502203 | n.a. |
| ENSECAG00000010814 | 0.584 | 9.2955 | 61.135 | 6E-10 | 5E-09 | 0.3436123 | n.a. |
| GNG2 | 0.678 | 9.4802 | 38.334 | 6E-10 | 5E-09 | 0.753304 | n.a. |
| PDS5B | 0.731 | 9.287 | 92.68 | 6E-10 | 5E-09 | 0.2863436 | n.a. |
| HNRNPD | 0.625 | 9.4221 | 38.112 | 7E-10 | 6E-09 | 0.6960352 | n.a. |
| PSMD8 | 0.594 | 9.543 | 37.964 | 7E-10 | 6E-09 | 0.8678414 | n.a. |

| | | | | | | | |
|---|---|---|---|---|---|---|---|
| TBC1D10B | 0.592 | 9.2823 | 73.044 | 8E-10 | 7E-09 | 0.3348018 | n.a. |
| SSRP1 | 0.663 | 9.3608 | 37.362 | 1E-09 | 8E-09 | 0.6035242 | n.a. |
| CCDC82 | 0.623 | 9.3286 | 56.38 | 1E-09 | 9E-09 | 0.3480176 | n.a. |
| NDUFA1 | 0.584 | 9.4956 | 37.203 | 1E-09 | 9E-09 | 0.7709251 | n.a. |
| TMEM126A | 0.645 | 9.3424 | 36.766 | 1E-09 | 1E-08 | 0.5506608 | n.a. |
| PSMB3 | 0.614 | 9.5646 | 36.532 | 2E-09 | 1E-08 | 0.876652 | n.a. |
| RCC1 | 0.688 | 9.2505 | 178.81 | 2E-09 | 1E-08 | 0.2511013 | n.a. |
| DCAKD | 0.698 | 9.2565 | 152.07 | 2E-09 | 1E-08 | 0.2643172 | n.a. |
| DELE1 | 0.803 | 9.3169 | 59.785 | 2E-09 | 1E-08 | 0.5066079 | n.a. |
| ENSECAG00000035662 | 0.613 | 9.4047 | 35.913 | 2E-09 | 2E-08 | 0.6343612 | n.a. |
| SSBP1 | 0.653 | 9.3194 | 47.843 | 3E-09 | 2E-08 | 0.4537445 | n.a. |
| HIRIP3 | 0.783 | 9.2693 | 91.527 | 3E-09 | 2E-08 | 0.30837 | n.a. |
| HIST1H1E | 0.71 | 9.278 | 90.018 | 3E-09 | 3E-08 | 0.30837 | n.a. |
| MPHOSPH6 | 0.679 | 9.423 | 34.66 | 4E-09 | 3E-08 | 0.7885463 | n.a. |
| MRPL13 | 0.726 | 9.326 | 42.625 | 4E-09 | 3E-08 | 0.5726872 | n.a. |
| ASF1A | 0.723 | 9.3664 | 34.403 | 5E-09 | 4E-08 | 0.6475771 | n.a. |
| SRSF10 | 0.634 | 9.3878 | 34.279 | 5E-09 | 4E-08 | 0.5550661 | n.a. |
| MAZ | 0.638 | 9.275 | 80.726 | 5E-09 | 4E-08 | 0.2555066 | n.a. |
| SNAPC2 | 0.702 | 9.2728 | 82.879 | 6E-09 | 4E-08 | 0.3259912 | n.a. |
| PPA2 | 0.609 | 9.2617 | 116.01 | 6E-09 | 5E-08 | 0.2599119 | n.a. |
| ETFB | 0.608 | 9.4454 | 33.92 | 6E-09 | 5E-08 | 0.7621145 | n.a. |
| PAICS | 0.719 | 9.2802 | 67.335 | 6E-09 | 5E-08 | 0.3832599 | n.a. |
| SFPQ | 0.603 | 9.3179 | 55.06 | 6E-09 | 5E-08 | 0.3568282 | n.a. |
| MPC1 | 0.599 | 9.3332 | 33.592 | 7E-09 | 5E-08 | 0.5154185 | n.a. |
| ENSECAG00000011754 | 0.608 | 9.3294 | 37.567 | 7E-09 | 6E-08 | 0.5991189 | n.a. |
| NAA10 | 0.627 | 9.5568 | 33.456 | 7E-09 | 6E-08 | 0.9207048 | n.a. |
| PTGES3 | 0.592 | 9.6195 | 33.3 | 8E-09 | 6E-08 | 0.8017621 | n.a. |
| HLTF | 0.678 | 9.2914 | 67.535 | 9E-09 | 7E-08 | 0.3215859 | n.a. |
| NUDT21 | 0.586 | 9.4899 | 32.564 | 1E-08 | 9E-08 | 0.8105727 | n.a. |
| ENSECAG00000018072 | 0.662 | 9.2931 | 68.016 | 1E-08 | 1E-07 | 0.3832599 | n.a. |
| FAM111A | 0.714 | 9.2868 | 78.48 | 2E-08 | 1E-07 | 0.3259912 | n.a. |
| PSMD14 | 0.647 | 9.3754 | 32.02 | 2E-08 | 1E-07 | 0.6740088 | n.a. |
| VRK1 | 0.709 | 9.2911 | 43.278 | 2E-08 | 1E-07 | 0.4449339 | n.a. |
| ENSECAG00000036779 | 0.584 | 9.5478 | 31.8 | 2E-08 | 1E-07 | 0.7488987 | n.a. |
| COQ7 | 0.595 | 9.2706 | 92.877 | 2E-08 | 2E-07 | 0.2643172 | n.a. |
| NSMCE2 | 0.624 | 9.2863 | 55.437 | 2E-08 | 2E-07 | 0.3612335 | n.a. |
| TXNL4A | 0.62 | 9.3794 | 31.148 | 2E-08 | 2E-07 | 0.6740088 | n.a. |
| MPHOSPH9 | 0.737 | 9.2536 | 125.42 | 3E-08 | 2E-07 | 0.2511013 | n.a. |
| PTS | 0.702 | 9.2895 | 56.931 | 3E-08 | 3E-07 | 0.3612335 | n.a. |
| TEX49 | 0.788 | 9.2532 | 167.55 | 4E-08 | 3E-07 | 0.2819383 | n.a. |
| ABRAXAS1 | 0.598 | 9.2896 | 55.515 | 4E-08 | 3E-07 | 0.2643172 | n.a. |
| MALSU1 | 0.612 | 9.4042 | 30.049 | 4E-08 | 3E-07 | 0.6828194 | n.a. |
| TPM3 | 0.627 | 9.3323 | 43.776 | 5E-08 | 4E-07 | 0.4625551 | n.a. |
| TFDP1 | 0.655 | 9.2786 | 48.597 | 5E-08 | 4E-07 | 0.4008811 | n.a. |
| PSMC3 | 0.619 | 9.4071 | 29.541 | 6E-08 | 4E-07 | 0.6784141 | n.a. |
| PRPF38A | 0.655 | 9.304 | 54.767 | 8E-08 | 5E-07 | 0.3656388 | n.a. |
| ERI1 | 0.704 | 9.2807 | 68.453 | 8E-08 | 6E-07 | 0.3348018 | n.a. |
| CDK4 | 0.629 | 9.4089 | 28.702 | 8E-08 | 6E-07 | 0.6696035 | n.a. |

| | | | | | | | |
|---|---|---|---|---|---|---|---|
| E2F4 | 0.627 | 9.3142 | 48.374 | 9E-08 | 6E-07 | 0.4317181 | n.a. |
| RRP7 | 0.583 | 9.3191 | 31.596 | 1E-07 | 7E-07 | 0.4493392 | n.a. |
| PCK2 | 0.734 | 9.2641 | 45.546 | 1E-07 | 7E-07 | 0.3612335 | n.a. |
| HAUS2 | 0.707 | 9.2577 | 96.808 | 1E-07 | 7E-07 | 0.2731278 | n.a. |
| POLE4 | 0.614 | 9.3299 | 36.784 | 1E-07 | 7E-07 | 0.4933921 | n.a. |
| ENSECAG00000015913 | 0.842 | 9.2635 | 107.95 | 1E-07 | 7E-07 | 0.3568282 | n.a. |
| TRIM37 | 0.754 | 9.2555 | 105.69 | 1E-07 | 9E-07 | 0.2643172 | n.a. |
| C10H6orf203 | 0.607 | 9.2937 | 47.344 | 3E-07 | 2E-06 | 0.3436123 | n.a. |
| RSRC1 | 0.631 | 9.3219 | 39.942 | 3E-07 | 2E-06 | 0.3920705 | n.a. |
| KHSRP | 0.622 | 9.2759 | 85.698 | 3E-07 | 2E-06 | 0.2819383 | n.a. |
| RAD50 | 0.589 | 9.314 | 36.375 | 4E-07 | 3E-06 | 0.3876652 | n.a. |
| SUZ12 | 0.669 | 9.2656 | 82.178 | 5E-07 | 3E-06 | 0.2731278 | n.a. |
| RBM14 | 0.625 | 9.3051 | 40.325 | 5E-07 | 3E-06 | 0.3700441 | n.a. |
| RBM38 | 0.642 | 9.2701 | 93.478 | 5E-07 | 3E-06 | 0.2643172 | n.a. |
| AIFM1 | 0.641 | 9.2611 | 87.743 | 1E-06 | 7E-06 | 0.2819383 | n.a. |
| ANKRD39 | 0.614 | 9.3428 | 32.738 | 1E-06 | 7E-06 | 0.5242291 | n.a. |
| CISD3 | 0.589 | 9.317 | 30.756 | 1E-06 | 8E-06 | 0.4273128 | n.a. |
| ENSECAG00000028298 | 0.621 | 9.2564 | 115.76 | 2E-06 | 1E-05 | 0.2643172 | n.a. |
| PRXL2A | 0.712 | 9.2641 | 65.057 | 2E-06 | 1E-05 | 0.2951542 | n.a. |
| RBBP7 | 0.608 | 9.3321 | 24.422 | 2E-06 | 1E-05 | 0.5462555 | n.a. |
| AKIP1 | 0.596 | 9.2726 | 47.571 | 3E-06 | 1E-05 | 0.3127753 | n.a. |
| RF00288 | 0.615 | 9.276 | 51.936 | 3E-06 | 2E-05 | 0.3436123 | n.a. |
| PTPMT1 | 0.606 | 9.4113 | 21.084 | 4E-06 | 2E-05 | 0.69163 | n.a. |
| NT5C3B | 0.62 | 9.3153 | 19.508 | 1E-05 | 5E-05 | 0.4977974 | n.a. |
| NDC80 | 0.793 | 9.2498 | 33.72 | 5E-05 | 2E-04 | 0.2731278 | n.a. |
| ENSECAG00000019698 | 0.59 | 9.4686 | 15.567 | 8E-05 | 4E-04 | 0.753304 | n.a. |
| MSH6 | 0.646 | 9.261 | 55.35 | 2E-04 | 6E-04 | 0.2643172 | n.a. |
| NQO1 | 0.651 | 9.2863 | 28.66 | 3E-04 | 0.001 | 0.4273128 | n.a. |
| SLA2 | 0.632 | 9.2804 | 31.806 | 3E-04 | 0.001 | 0.3039648 | n.a. |
| ARL4C | 0.585 | 9.3066 | 19.482 | 9E-04 | 0.003 | 0.3656388 | n.a. |

| genes | logFC | logCPM | F | PValue | FDR | percent.exp | Surfacome.Label |
|---|---|---|---|---|---|---|---|
| GZMM | 2.61 | 9.4961 | 11282 | 0 | 0 | 0.6028862 | n.a. |
| CCL5 | 2.221 | 11.908 | 4079.1 | 0 | 0 | 0.5611972 | n.a. |
| LGALS1 | 1.783 | 11.109 | 3523.6 | 0 | 0 | 0.8701229 | n.a. |
| ENSECAG00000031322 | 1.68 | 10.499 | 2168.1 | 0 | 0 | 0.6397648 | n.a. |
| CXCR3 | 1.478 | 9.3103 | 4533.5 | 0 | 0 | 0.3083912 | surface |
| CTSW | 1.471 | 9.899 | 2507.3 | 0 | 0 | 0.3655799 | n.a. |
| ZNF683 | 1.381 | 9.478 | 3050.2 | 0 | 0 | 0.2693747 | n.a. |
| IDO1 | 1.363 | 9.5783 | 1579.4 | 0 | 0 | 0.6424372 | n.a. |
| ENSECAG00000029287 | 1.265 | 10.043 | 1502.6 | 0 | 0 | 0.4532336 | n.a. |
| ENSECAG00000028035 | 1.36 | 9.3107 | 3853.6 | 1E-264 | 7E-262 | 0.251737 | n.a. |
| INPP4B | 1.171 | 9.4661 | 1189.6 | 2E-235 | 1E-232 | 0.5077499 | n.a. |
| GLIPR1 | 0.954 | 9.887 | 803.74 | 8E-175 | 3E-172 | 0.7161945 | surface |
| COTL1 | 0.761 | 10.529 | 793.55 | 1E-172 | 5E-170 | 0.9380011 | n.a. |
| ENSECAG00000007681 | 0.945 | 9.6603 | 791.58 | 3E-172 | 1E-169 | 0.6750401 | n.a. |
| ITGB1 | 0.89 | 9.5309 | 779.33 | 1E-169 | 4E-167 | 0.4019241 | surface |
| GIMAP7 | 0.638 | 10.513 | 777.21 | 4E-169 | 1E-166 | 0.9764832 | n.a. |
| TSPO | 1.17 | 9.7323 | 1163.4 | 9E-168 | 3E-165 | 0.3308391 | n.a. |
| PLAC8B | 0.79 | 11.113 | 768.72 | 2E-167 | 7E-165 | 0.9203634 | n.a. |
| CD2 | 0.835 | 9.8338 | 765.19 | 1E-166 | 4E-164 | 0.8027793 | surface |
| ENSECAG00000007668 | 1.108 | 9.3879 | 1003.1 | 1E-153 | 3E-151 | 0.364511 | n.a. |
| CYBA | 0.824 | 10.509 | 680.87 | 1E-148 | 3E-146 | 0.7204703 | n.a. |
| IL7R | 0.915 | 9.3731 | 1318.7 | 2E-114 | 4E-112 | 0.6723677 | surface |
| CD40LG | 0.88 | 9.3506 | 942.87 | 3E-112 | 5E-110 | 0.2971673 | surface |
| GLIPR2 | 0.744 | 9.47 | 736.04 | 6E-105 | 1E-102 | 0.3270978 | n.a. |
| ENSECAG00000032959 | 0.829 | 9.6944 | 497.65 | 5E-101 | 9E-99 | 0.3180118 | n.a. |
| IFI27 | 0.729 | 9.7314 | 447.59 | 1E-98 | 2E-96 | 0.4152859 | n.a. |
| CTSC | 0.759 | 9.6133 | 443.31 | 8E-98 | 1E-95 | 0.5783004 | n.a. |
| ID2 | 0.812 | 9.6451 | 466.17 | 1E-95 | 2E-93 | 0.2677712 | n.a. |
| CD37 | 0.735 | 9.891 | 384.11 | 5E-85 | 7E-83 | 0.676109 | surface |
| ENSECAG00000040180 | 0.643 | 9.7881 | 377.34 | 1E-83 | 2E-81 | 0.6723677 | n.a. |
| GBP5 | 0.685 | 9.7163 | 375.58 | 3E-83 | 4E-81 | 0.6488509 | n.a. |
| MARCKSL1 | 0.773 | 9.5078 | 352.63 | 3E-78 | 4E-76 | 0.4227686 | n.a. |
| ADGRE5 | 0.782 | 9.5791 | 342.82 | 4E-76 | 5E-74 | 0.3468733 | n.a. |
| CISH | 0.724 | 9.3863 | 471.41 | 6E-74 | 8E-72 | 0.2779262 | n.a. |
| IGFLR1 | 0.597 | 9.4383 | 382.35 | 1E-72 | 1E-70 | 0.3928381 | surface |
| CD4 | 0.72 | 9.36 | 508.78 | 1E-72 | 2E-70 | 0.3142704 | surface |
| LGALS3 | 0.691 | 9.5239 | 502.16 | 2E-72 | 3E-70 | 0.2699091 | n.a. |
| ENSECAG00000039088 | 0.649 | 9.7274 | 310.45 | 4E-69 | 4E-67 | 0.5013362 | n.a. |
| ALDOA | 0.599 | 9.99 | 299.13 | 1E-66 | 1E-64 | 0.6509888 | n.a. |
| TALDO1 | 0.581 | 9.9099 | 273.17 | 4E-61 | 4E-59 | 0.6119722 | n.a. |
| KLF6 | 0.64 | 9.5982 | 251.47 | 2E-56 | 2E-54 | 0.4024586 | n.a. |
| TESC | 0.582 | 9.3671 | 345.33 | 3E-56 | 3E-54 | 0.2816676 | n.a. |
| PIM1 | 0.631 | 9.3916 | 314.47 | 8E-51 | 7E-49 | 0.2656334 | n.a. |
| TEX30 | 0.607 | 9.4443 | 279.92 | 4E-49 | 3E-47 | 0.2779262 | n.a. |
| AKIRIN2 | 0.616 | 9.4269 | 239.83 | 5E-48 | 4E-46 | 0.2886157 | n.a. |
| STAMBPL1 | 0.605 | 9.3697 | 292.96 | 7E-47 | 5E-45 | 0.2677712 | n.a. |
| SCML4 | 0.598 | 9.3954 | 269.71 | 2E-42 | 1E-40 | 0.2683057 | n.a. |

| genes | logFC | logCPM | F | PValue | FDR | percent.exp | Surfacome.Label |
|---|---|---|---|---|---|---|---|
| ENSECAG00000000775 | 2.862 | 9.4621 | 7517.9 | 0 | 0 | 0.4877518 | n.a. |
| CD8A | 2.314 | 9.4281 | 4866.9 | 0 | 0 | 0.3832335 | surface |
| SELL | 1.627 | 9.6904 | 1720.3 | 0 | 0 | 0.586282 | surface |
| CCR7 | 1.373 | 9.4631 | 1668.4 | 2E-275 | 2E-272 | 0.4915623 | surface |
| ID3 | 1.309 | 9.6697 | 1273.6 | 6E-274 | 4E-271 | 0.7044094 | n.a. |
| ENSECAG00000030502 | 1.176 | 9.3968 | 2175.8 | 5E-212 | 2E-209 | 0.279804 | n.a. |
| TXNIP | 1.008 | 10.204 | 804.93 | 5E-175 | 1E-172 | 0.8530212 | n.a. |
| PLAC8B | 0.811 | 11.113 | 631.7 | 4E-138 | 9E-136 | 0.8769733 | n.a. |
| OXNAD1 | 1.032 | 9.3622 | 897.04 | 2E-114 | 5E-112 | 0.3004899 | n.a. |
| CD27 | 0.817 | 9.4777 | 469.99 | 3E-102 | 5E-100 | 0.4812194 | surface |
| DRA | 0.598 | 11.742 | 374.22 | 6E-83 | 7E-81 | 0.8633642 | n.a. |
| LEF1 | 0.781 | 9.4498 | 523.95 | 2E-80 | 2E-78 | 0.442025 | n.a. |
| ENSECAG00000037934 | 0.88 | 9.4027 | 418.73 | 1E-76 | 1E-74 | 0.2591181 | n.a. |
| FKBP5 | 0.818 | 9.3784 | 473.31 | 3E-74 | 2E-72 | 0.328797 | n.a. |
| HCST | 0.717 | 9.5985 | 314.57 | 5E-70 | 4E-68 | 0.4273272 | n.a. |
| WARS | 0.65 | 9.8298 | 298.16 | 2E-66 | 1E-64 | 0.6744692 | n.a. |
| MHCB3 | 0.608 | 10.265 | 298.15 | 2E-66 | 1E-64 | 0.7512248 | n.a. |
| RGS10 | 0.682 | 9.5903 | 258.31 | 1E-55 | 7E-54 | 0.3881328 | n.a. |
| CD69 | 0.696 | 9.4382 | 296.78 | 3E-55 | 2E-53 | 0.3505716 | surface |
| TCF7 | 0.642 | 9.3798 | 347.34 | 9E-54 | 6E-52 | 0.3081111 | n.a. |
| ENSECAG00000037985 | 0.75 | 9.4315 | 299.67 | 1E-52 | 6E-51 | 0.3015787 | n.a. |
| ENSECAG00000037904 | 0.696 | 9.482 | 231.55 | 4E-52 | 3E-50 | 0.3217202 | n.a. |
| RF00213 | 0.727 | 9.3791 | 316.47 | 6E-52 | 4E-50 | 0.2536745 | n.a. |
| TMC6 | 0.595 | 9.4027 | 192.58 | 6E-39 | 3E-37 | 0.2629287 | n.a. |
| ACOT13 | 0.587 | 9.4038 | 182.25 | 1E-34 | 5E-33 | 0.26184 | n.a. |

| genes | logFC | logCPM | F | PValue | FDR | percent.exp | Surfacome.Label |
|---|---|---|---|---|---|---|---|
| ENSECAG00000030502 | 2.376 | 9.3968 | 6899.2 | 0 | 0 | 0.5390399 | n.a. |
| CCR7 | 1.731 | 9.4631 | 2564.1 | 0 | 0 | 0.6078658 | surface |
| SELL | 1.508 | 9.6904 | 1405.9 | 1E-301 | 7E-299 | 0.615963 | surface |
| ENSECAG00000033316 | 0.96 | 10.74 | 890.22 | 4E-193 | 8E-191 | 0.9976865 | n.a. |
| ENSECAG00000028304 | 1.219 | 9.3449 | 1436.2 | 9E-178 | 2E-175 | 0.2834008 | n.a. |
| OXNAD1 | 1.303 | 9.3622 | 1339.8 | 8E-171 | 1E-168 | 0.3759398 | n.a. |
| TXNIP | 0.984 | 10.204 | 740.66 | 2E-161 | 3E-159 | 0.86524 | n.a. |
| LEF1 | 1.053 | 9.4498 | 907.77 | 1E-146 | 1E-144 | 0.5135917 | n.a. |
| RGS10 | 1.077 | 9.5903 | 669.87 | 3E-146 | 4E-144 | 0.5054945 | n.a. |
| ENSECAG00000034569 | 0.789 | 9.7713 | 446.77 | 2E-98 | 1E-96 | 0.6096009 | n.a. |
| ENSECAG00000032138 | 1.018 | 9.3299 | 754.45 | 1E-92 | 9E-91 | 0.264893 | n.a. |
| ENSECAG00000030387 | 0.974 | 9.433 | 779.09 | 1E-91 | 1E-89 | 0.2753036 | n.a. |
| PLAC8B | 0.657 | 11.113 | 387.71 | 8E-86 | 6E-84 | 0.8889532 | n.a. |
| FKBP5 | 0.871 | 9.3784 | 525.88 | 3E-83 | 2E-81 | 0.3337189 | n.a. |
| TRAT1 | 0.699 | 9.7859 | 339.17 | 2E-75 | 1E-73 | 0.626952 | surface |
| CD27 | 0.653 | 9.4777 | 293.73 | 1E-65 | 9E-64 | 0.4650087 | surface |
| ADGB | 0.755 | 9.3514 | 476.05 | 1E-61 | 6E-60 | 0.2672065 | n.a. |
| CD69 | 0.71 | 9.4382 | 297.21 | 4E-57 | 2E-55 | 0.3678427 | surface |
| RF00213 | 0.685 | 9.3791 | 273.72 | 7E-46 | 3E-44 | 0.2527473 | n.a. |
| SATB1 | 0.626 | 9.3659 | 274.63 | 1E-40 | 4E-39 | 0.2504338 | n.a. |
| PLA2G16 | 0.606 | 9.6144 | 221.3 | 8E-36 | 3E-34 | 0.3233083 | n.a. |
